# Generating clonal seeds using *non-reduction in female4* (*nrf4*), a novel meiotic mutant of maize

**DOI:** 10.1101/2025.10.24.684334

**Authors:** Nina Chumak, G. Arco Brunner, Mark E. Williams, Wenjing She, Philippa J. Barrell, Valeria Gagliardini, Frédérique Pasquer, Tim W. Fox, Marc C. Albertsen, Ueli Grossniklaus

**Author notes:** Corresponding author. (U.G.). Seestrasse 286, 8810 Horgen, Switzerland (G. Arco Brunner). Springer Nature, Unit 10-11, The Center, 989 Changle Road, Shanghai 200031, China (Wenjing She). New Zealand Institute for Plant & Food Research, Private Bag 4704, Christchurch 8140, New Zealand (Philippa J. Barrell). Food Brewer, Bachtobelstrasse 5, 8810 Horgen, Switzerland (Frédérique Pasquer). Albertsen Crop Genetics for Humanity, Johnston, IA 50131, USA.

## Abstract

The discovery of increased vigor and yield in hybrid corn compared to inbred lines dates to the early 20^th^ century. If hybrid vigor could be maintained across generations, it would revolutionize agriculture. A potential method to achieve this goal is asexual reproduction through seed (apomixis). We searched for mutants mimicking components of apomixis and here, we describe a novel meiotic mutant, *non-reduction in female4* (*nrf4*) of *Zea mays* (maize), that produces unreduced female gametes – an essential component of apomixis. By combining the *nrf4* mutant with the previously described *matrilineal*/*not like dad* (*mtl*/*nld*) mutant, which leads to embryo development without paternal contribution, we successfully produced clonal seeds. This is an important step towards the engineering of clonally propagated hybrids in maize, the most widely grown crop world-wide. Our findings provide new avenues for the engineering of synthetic apomixis in crops.

**One-Sentence Summary:** This manuscript provides a proof-of-principle that synthetic apomixis can be engineered in maize.

## Main Text

Apomixis, defined as the asexual reproduction through seed (*1*), results in offspring that are genetically identical to the mother plant (*2*, *3*). Plant breeding would greatly benefit from harnessing apomixis in crops, as it would allow the propagation of highly vigorous F_1_ hybrid genotypes over generations without loss of hybrid vigor (*4*) (Fig. S1). There are three principal differences between sexual and apomictic reproduction (*5*): First, the absence of reduction and recombination (apomeiosis); second, the development of an embryo without paternal contribution (parthenogenesis); and third, functional endosperm formation. The study of natural apomicts revealed that apomixis evolved as a deviation from the sexual pathway and that the individual components of apomixis are likely of simple genetic inheritance (*1*, *6*). Thus, engineering apomixis in a sexual species through combination of its components seems possible (*7–9*). Mutations affecting the maize *Elongate1* and *Argonaute104* genes lead to phenotypes mimicking apomeiosis and produced unreduced female gametes (*10*, *11*). However, another essential aspect of apomeiosis, namely the loss of recombination, is either unaffected (*12*) or its status remains unclear(*11*). In *Arabidopsis* and rice, it was shown that *MiMe,* a combination of the loss-of-function mutants *rec8*, *spo11* and *osd1*, displays all features of apomeiosis, namely the absence of reduction and recombination (*13*, *14*). In *Arabidopsis* and rice, the combination of *MiMe* with a paternal chromosome elimination system (*CenH3*-tail-swap or *mtl*/*nld*) (*15–18*) or a *BABY-BOOM-LIKE (BBML)* gene that is ectopically expressed in the egg cell (*19*, *20*) results in the production of clonal offspring, albeit at low frequency or with reduced fertility (*21*, *22*). Although *MiMe* can be used to engineer an apomeiotic phenotype in crops, the application of the triple mutant in a breeding scheme might be challenging.

This study aimed to identify mutations in maize that mimic apomeiosis. Therefore, we performed a forward genetic screen that exploited the inviability of seeds derived from interploidy crosses (ploidy barrier) (Fig. 1A-C). In flowering plants, gametes are formed by the haploid gametophytes, which develop post-meiotically through mitotic divisions. The female gametophyte (embryo sac) produces two gametes, the haploid egg and homo-diploid central cell, while the male gametophyte (pollen) carries two haploid sperm. During double fertilization, one sperm fertilizes the egg cell to generate a diploid embryo, while the other fuses with the central cell to form a triploid endosperm (*23*). Thus, the endosperm, which makes up about 80% of the seed, carries one paternal and two maternal genomes. If the ploidies of embryo sac and pollen differ, the parental genomes of the endosperm are imbalanced and its development is compromised, resulting in shrunken, inviable kernels (Fig. 1B) (*24*). We exploited this ploidy barrier and pollinated diploid plants from about 3’200 families segregating for active *Mutator* (*Mu*)-transposons (*25*) with pollen from tetraploid plants carrying the *R1-Navajo* (*26*) anthocyanin marker (4n *R1-nj*). Wild-type diploid plants are expected to produce only shrunken kernels (Fig. 1B; Fig. S2), whereas a mutant with an unreduced embryo sac should develop plump kernels containing normally formed endosperm. We recovered several families with the desired phenotype. If the plump kernels resulted from fertilization of an unreduced embryo sac with diploid pollen, they should contain a tetraploid embryo and a hexaploid endosperm. To assess this, we performed ploidy analysis of kernels by flow cytometry (Fig. S3) and recovered four heritable mutants with good penetrance that we named *<u>n</u>on-<u>r</u>eduction in female* (*nrf*). Here, we describe the *nrf4* mutant at the genetic and molecular level and show that it can be used for clonal seed production in maize.

**Fig. 1.**
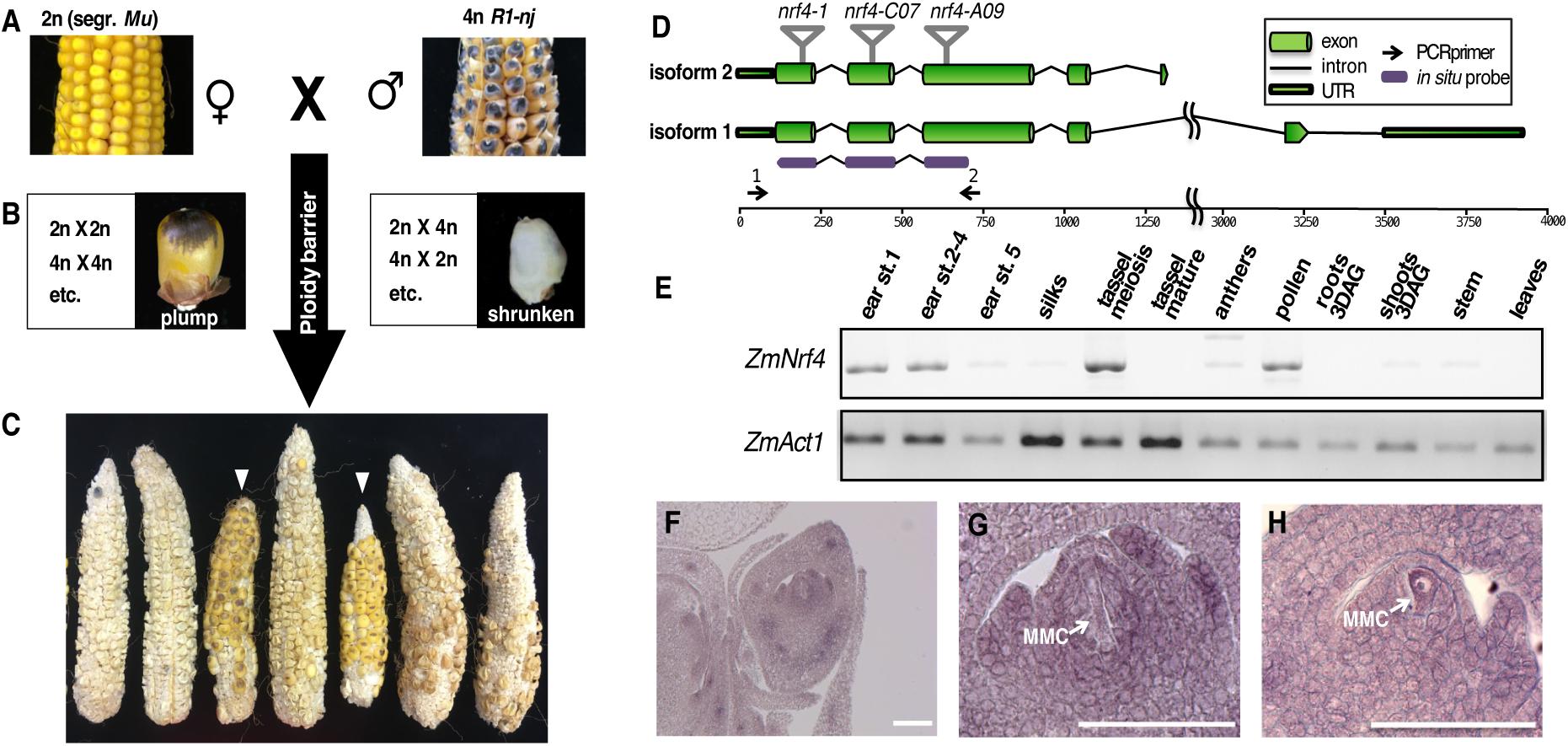
*Nrf4* identification and expression. (**A-C**) Schematic representation of the mutant screen to identify *non-reduction in female* (*nrf*) mutants. (**A**) Diploid families segregating active *Mu* were pollinated by tetraploid males. (**B**) The ploidy barrier in maize leads to endosperm abortion in interploidy crosses. In contrast, individuals with a mutation leading to unreduced female gametophyte formation form normal seed in crosses with tetraploid males. (**C**) A family segregating for the *nrf4* mutation. Asterisks mark ears from nrf4 homozygous individuals (**D**) Structure of the *Nrf4* gene. The positions of 3 transposon insertions are marked on the top, the location of the primers for RT-PCR and the position of the probe for *in situ* hybridization are shown below. (**E**) RT-PCR results of *Nrf4* expression in different plant tissues. *ZmActin1 (ZmAct1)* expression was used as a loading control. (**F-H**), RNA *in situ* hybridization of *Nrf4* mRNA with antisense probe on maize ovules. MMC - megaspore mother cell. Scale bars = 100 mm.

To test if the *nrf4* mutant has a heritable phenotype, a population segregating for *nrf4* was crossed to the tetraploid 4n *R1-nj* tester line. Out of 109 segregating *nrf4* plants, 86 displayed <20% of plump kernels and 23 showed >80% of plump kernels (Table 1), suggesting a recessive inheritance of the mutation. When we pollinated wild-type diploid and wild-type tetraploid plants with the 4n *R1-nj* tester, we observed ears with shrunken kernels and ears with plump kernels, respectively, as expected (Table 1, Fig. S2). Homozygous mutant *nrf4* plants produced a normal amount of pollen. Interestingly, when *nrf4* pollen fertilized wild-type diploid females, more than 95% of plump, diploid, viable kernels resulted, suggesting that the *nrf4* mutation does not affect male meiosis (Fig. S2). This is unique for meiotic mutants in maize, which usually affect both sexes (*27*).

**Table 1.** Genetic segregation of the *nrf4* mutant phenotype in three different mutant lines.

| Cross | Phenotype | | Expected ratio | $\chi^2$<br>P-value |
| --- | --- | --- | --- | --- |
|  | Shrunken kernels<sup>a</sup> | Plump kernels<sup>a</sup> |  |  |
| *nrf4-1* F<sub>2</sub> x 4n *R1-nj* | 86 | 23 | 3:1 | 0.3472 |
| 2n WT x 4n *R1-nj* | 55 | 0 | 1:0 | NA |
| 4n WT x 4n *R1-nj* | 0 | 34 | 0:1 | NA |
| *nrf4-C07* F<sub>2</sub> x 4n *R1-nj* | 17 | 5 | 3:1 | 0.8055 |
| *nrf4-A09* F<sub>2</sub> x 4n *R1-nj* | 40 | 12 | 3:1 | 0.7488 |
| (*nrf4-1* x *nrf4-C07*) x 4n *R1-nj* | 87 | 17 | 3:1 | 0.0415 |
| (*nrf4-1* x *nrf4-A09*) x 4n *R1-nj* | 45 | 16 | 3:1 | 0.8245 |

<sup>a</sup>The two columns show the number of individuals with mostly shrunken or mostly plump kernels.

To clone the *Nrf4* gene, we employed Selective Amplification of Insertion-Flanking Fragments (SAIFF) technology (*28*) combined with next-generation sequencing. We identified a *Mu* transposon insertion in the long arm of chromosome 7 in the *GRMZM2G148133* (*Zm00001eb329530*) gene, associated with the mutant phenotype. The gene was annotated to have 2 isoforms (Fig. 1D). In the original mutant allele affecting *Nrf4*, *nrf4-1*, the *Mu* transposon is inserted at the end of the first exon, which is common to both isoforms. The two isoforms encoded by *Nrf4* are 235 and 248 amino acids long, respectively, and show no homology to protein domains of known function. To confirm that we identified the correct gene affected in the *nrf4-1* mutant, we obtained two independent mutant alleles affecting the same gene from the Trait Utility System for Corn (TUSC) (*29*): PV0363-C07 and PV03129-A09, which we named *nrf4-C07* and *nrf4-A09,* respectively. Plants homozygous for either, the *nrf4-C07* or *nrf4-A09* allele, show the *non-reduction in female* mutant phenotype; they produce plump kernels when crossed to 4n *R1-nj* and are allelic to *nrf4-1* as established by non-complementation (Table 1).

Phylogenetic analysis revealed that NRF4 is a grass-specific protein with very little homology to other monocots or dicots. In grasses, *Nrf4* is present either as a unique gene or as a member of a small two-or three-member gene family (Fig. S4). Interestingly, the maize genome encodes a close homolog of *Nrf4*, *Nrf4-like,* but the full penetrance of the *non-reduction in female* phenotype of the *nrf4* mutant suggests that *Nrf4-like* is unlikely to have a major function in reduction.

To identify the spatiotemporal expression pattern of *Nrf4* in different plant organs, we first performed *in silico* analysis of RNA-Seq and microarray data available in public databases. In general, *Nrf4,* as well as *Nrf4-like,* shows low expression levels (Fig. S5) and is likely to be expressed in the developing ear, embryo, endosperm, and tassel. To confirm the results of this *in silico* analysis, we performed RT-PCR (Fig. 1E). *Nrf4* is expressed in young reproductive organs (developing ears and tassels) and pollen, but only very weakly if at all in vegetative tissues. Using RNA *in situ* hybridization, we detected abundant *Nrf4* transcripts in a portion of the nucellus around the megaspore mother cell and ovule (Fig. 1F-H). In developing and functional megaspore mother cells, the cells that will undergo meiosis, the intensity of the *Nrf4* signal varied from very strong to intermediate (Fig. 1G, H). Taken together, these results indicate that *Nrf4* is expressed in the reproductive cell lineages before and during meiosis.

To gain insights into the *nrf4* mutant phenotype during reproduction, we performed confocal laser scanning microscopy on ovules during female sporogenesis and gametogenesis. While the differentiation of the megaspore mother cell was not affected in *nrf4-1,* meiosis was severely disturbed and not a single megaspore tetrad was observed (Fig. 2A), indicating that the *nrf4-1* mutant completes none or only one of the two meiotic divisions. Furthermore, in contrast to the wild type, the chromosomes in metaphase were not fully synapsed but just connected in the centromeric region in some of the *nrf4-1* ovules (Fig. 2B-D), indicating at least a partial absence of recombination, which requires synapsis. Interestingly, gametogenesis was not affected by this abnormal meiosis and led to the development of a morphologically normal 8-nuclear embryo sac. Apparently, the *nrf4-1* mutant undergoes mitosis instead of meiosis. Specifically, the megaspore mother cell seems to divide to directly form a 2-nuclear embryo sac, a form apomeiosis known as mitotic diplospory in natural apomicts (*6*). That, in *nrf4-1* ovules, we only observed megaspore mother cells and 2-nuclear embryo sacs, but hardly any dyads and no tetrads or 1-nuclear functional megaspores, corroborates this hypothesis. This apomeiotic phenotype is expected to lead to the formation of unreduced and, at least in some cases, unrecombined female gametes with a complete set of maternal chromosomes.

**Fig. 2.**
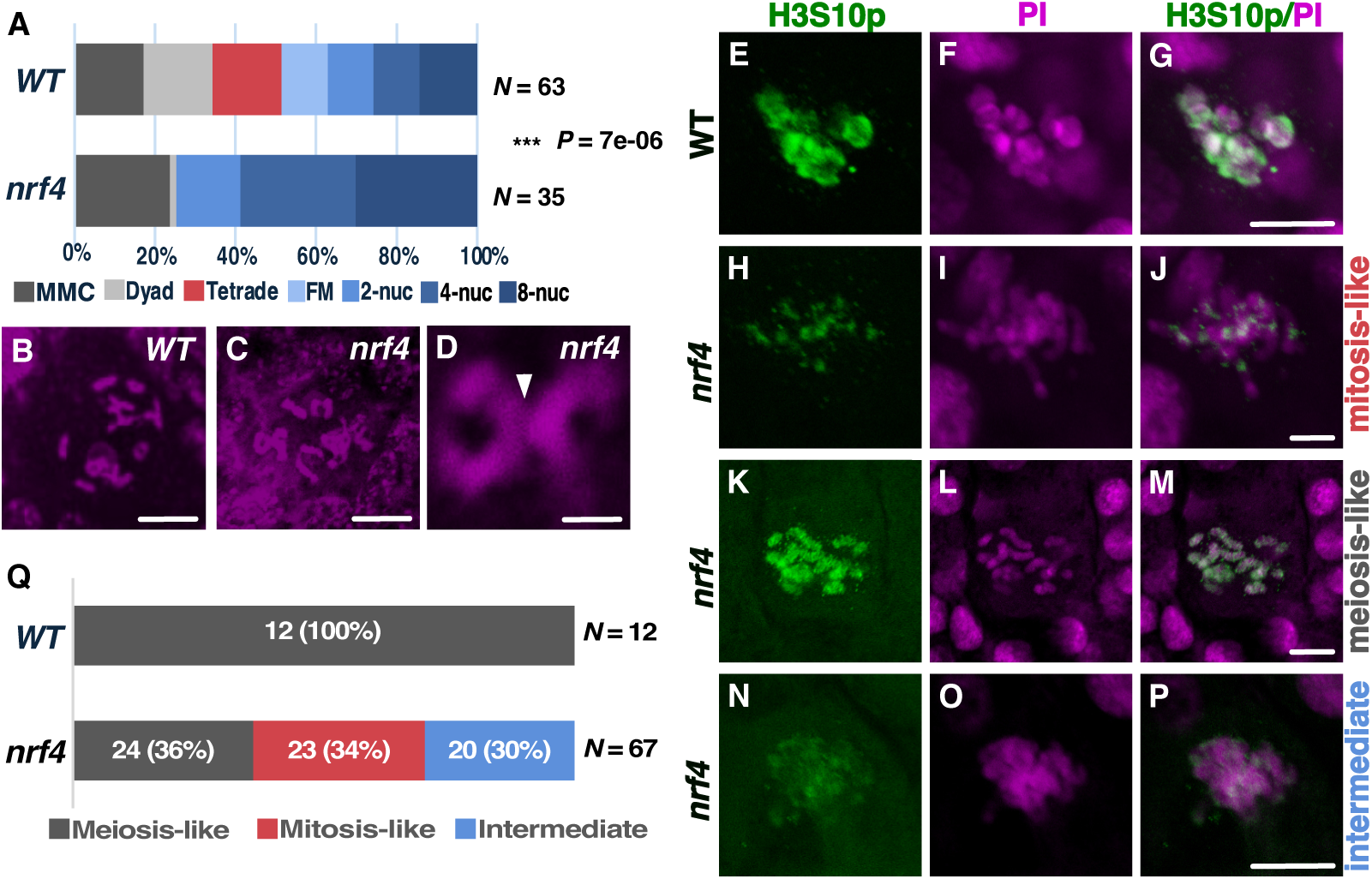
The *nrf4-1* mutation leads to the production of unreduced embryo sacs and abnormal chromosome pairing in meiosis I. (**A**) Percentage of different stages of megasporogenesis and female gametophyte development of the *nrf4-1* mutant in comparison to the wild type. The number of analyzed ovules and the P-value of a chi-square test are indicated on the right side. (**B-D**) Chromosome behavior in wild-type and *nrf4-1* ovules during the first division of the megaspore mother cell (MMC). (**B**) Metaphase of meiosis I in the wild type. (**C**) Metaphase of the first division in the *nrf4-1* mutant. (**D**) In *nrf4-1* mutant megaspore mother cells, metaphase chromosomes are only connected at the centromere (arrow). (**E-P**) Immunostaining of wild-type and *nrf4* mutant megaspore mother cells at metaphase/anaphase of meiosis decorated with H3S10p antibodies (green); DNA is counterstained with propidium iodine (magenta). Genotypes are indicated on the left side of the panel. Three patterns of H3S10p distribution during meiosis were observed in *nrf4* mutant megaspore mother cells, indicated on the right side of the panels. (H-J) – mitosis-like distribution, (K-M) – meiosis-like distribution, (N-P) – intermediate with a weak patchy signal. (Q) Stacked bar plot of the percentage of ovules displaying the three different H3S10p distribution patterns in *nrf4* mutants and the wild type. The number of analyzed ovules is indicated on the right side. MMC – megaspore mother cell; FM – functional megaspore; 2-nuc – two-nuclear embryo sac; 4-nuc – four-nuclear embryo sac; 8-nuc – 8 nuclear embryo sac. Scale bars: B&C – 25 μm; D – 5 μm; G, J, M, P – 10 μm.

To further investigate the meiotic phenotype of *nrf4-1*, we immunostained *nrf4-1* ovules with an antibody against histone H3 that is phosphorylated at serine 10 (H3S10p). H3S10p is a marker of chromosome synapsis and is localized at the centromeric regions of chromosomes only in the metaphase of mitosis and meiosis II, while it decorates both chromosome arms and centromeric regions in meiosis I ((*30*) and Fig. 2E-M). We observed that the anti-H3S10p antibody localized only to the centromeric regions of the chromosomes (Fig. 2H-J&Q) in 23/67 (35%) of *nrf4-1* megaspore mother cells, as expected for mitotic divisions. In the remaining ovules, the anti-H3S10p antibody was either distributed along the arms of the chromosomes as in the wild type (24/67=35%) (Fig. 2K-M&Q) or showed an intermediate pattern with only a partial reduction on the chromosome arms (20/67=30%) (Fig. 2 N-P&Q). Taken together, these results suggest that of the >95% of *nrf4-1* female gametes that are viable and unreduced, at least 35% are derived from a mitotic division and are expected to carry unrecombined chromosomes that are genetically identical to those of the mother plant. But it is unclear whether and to what extent recombination is suppressed in megaspore mother cells that show an intermediate or even wild-type pattern of the anti-H3S10p antibody.

Our genetic, cytological, and molecular characterizations suggested that *nrf4* is a good candidate to engineer the apomeiotic component of apomixis in maize. To demonstrate that clonal seeds can be produced, we introgressed the *nrf4-1* mutation into four genetically diverse inbred lines and obtained F_1_ hybrids in different combinations. We pollinated hybrid plants homozygous for the *nrf4-1* mutation with tetraploid males homozygous for the *R1-nj* marker and the *mtl*/*nld* mutation. The latter causes paternal genome elimination and thus the production of maternal haploid progeny in maize (*16–18*). The pollen donor had to be tetraploid to avoid kernel abortion due to an imbalanced endosperm (Fig. 1B). Among the progeny of this cross, we screened for individuals with (i) no paternal contribution by the absence of *R1-nj* pigmentation in the embryo; (ii) a diploid genome, revealed through ploidy analysis by flow cytometry, and (iii) homozygosity for the *nrf4-1* mutation confirmed by genotyping (Fig. 3A).

**Fig. 3.**
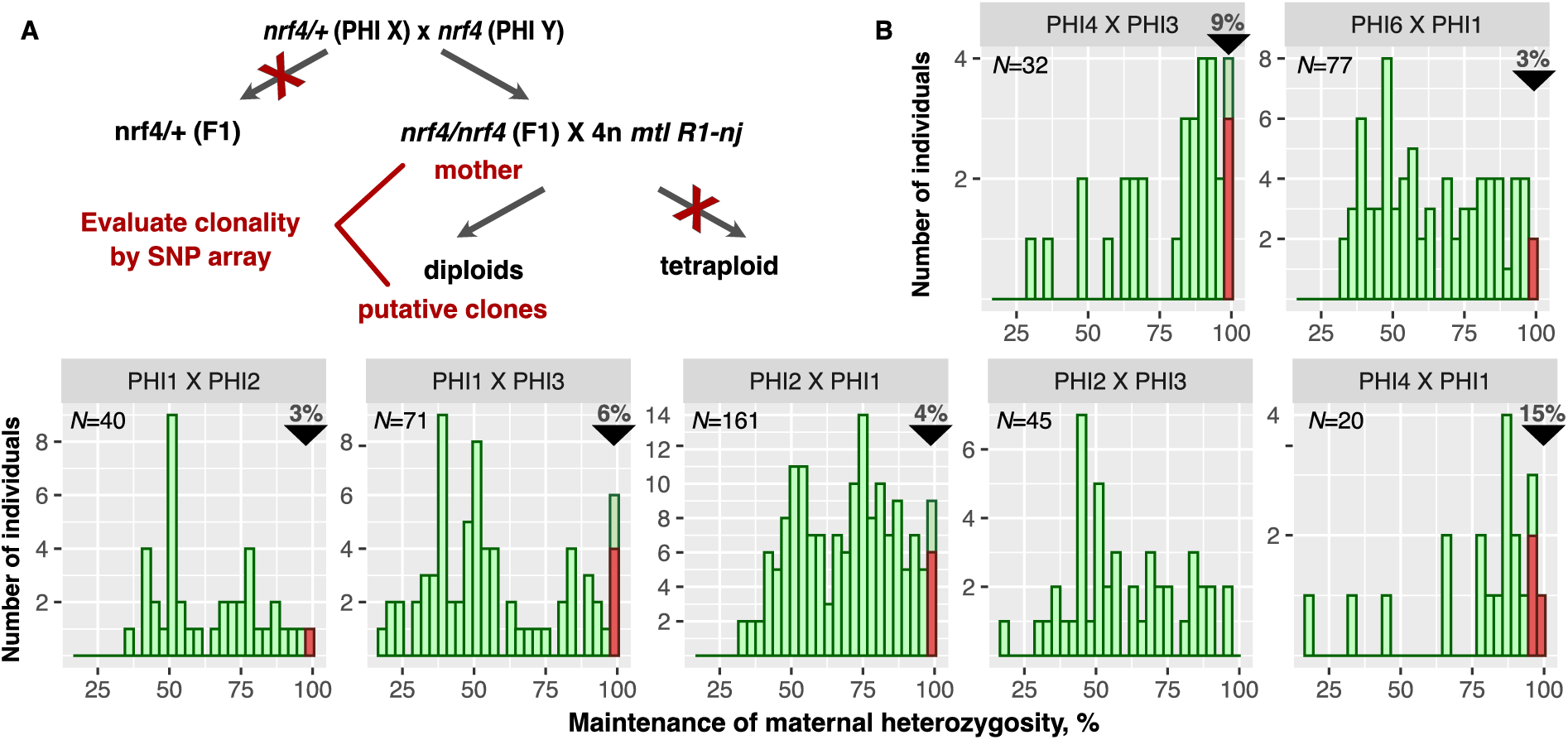
Production of clonal seed by means of *nrf4* and the *mtl*/*nld* mutation. (**A**) Crossing scheme to generate and identify clonal seeds. (**B**) Histograms showing the number of individuals that maintain a certain percentage (%) of maternal heterozygosity in 7 different hybrid combinations. The clonal individuals (see Supplementary Table 10) are high-lighted in red and their percentage is indicated in the top right corner of the plot for each hybrid. The total number of individuals analyzed (N) for each hybrid is shown in the top left corner of each plot.

Using this approach, we obtained between 20 and 161 putative clonal seeds from seven different F_1_ hybrid combinations (Fig. 3B). To test whether maternal heterozygosity was maintained (absence of recombination), we analyzed the putative clonal seeds together with the corresponding mother plants using a single nucleotide polymorphism (SNP) array. We determined the heterozygosity status of 192 to 384 SNP markers distributed across the genome of each plant (Table S1-S8). The number of informative markers (heterozygous in the mother plants and assessed in the progeny) varied from 122 to 306 SNPs per individual. The number of individuals that maintained a certain percentage of maternal heterozygosity is shown in Fig. 3B and Table S9 for each hybrid combination. We identified some individuals that completely maintained maternal heterozygosity in all hybrid combinations, except PHI2 x PHI3. Up to 15% of the diploid progeny of a *nrf4-1* homozygous hybrid crossed to tetraploid *mtl*/*nld* pollen <u>donor</u> were clonal seeds with a genotype identical to that of the mother plant (Table S10). The remaining individuals lost maternal heterozygosity to some extent. Two hybrids, PHI4xPHI1 and PHI4xPHI3 (Fig. 3B) stand out by comprising a vast majority of individuals maintaining a very high level of maternal heterozygosity, indicating a suppression of recombination also in those individuals that were not clones. A possible explanation for the differences in maintaining maternal heterozygosity may be the presence of enhancers of the *nrf4* phenotype in the PHI4 inbred line.

In summary, we were able to obtain clonal progeny independently of genetic background of the maternal hybrid. The *nrf4* mutation led to the production of unreduced gametes without affecting fertility and seed set in all tested genetic backgrounds. Recombination is suppressed in the *nrf4* mutant to different extents, depending on the genetic background. It is possible that *Nrf4-like* plays a partially redundant role with *Nrf4* in recombination, while it does not appear to have a major function in reduction. The respective activities or expression patterns of *Nrf4* and *Nrf4-like* may differ between inbred lines, leading to potential differences in suppressing recombination among the hybrids tested. Paternal genome elimination using the *mtl*/*nld* mutation is not very efficient; given that 35% of the megaspore mother cells undergo a mitotic instead of meiotic division, higher frequencies of clonal seeds can be expected in future experiments that may combine *nrf4* with *BBML* expression in the egg cell.

Using a forward genetic approach, we discovered and characterized the *nrf4* mutation which forms viable, unreduced female gametes and displays no or a reduced level of recombination. The *nrf4-1* mutant is defective in the pairing of homologous chromosome arms during meiosis I similar to the *asynaptic1* (*as1*) and *desynaptic1 (dy1)* mutants of maize (*31–33*). But unlike these highly sterile mutants, *nrf4-1* homozygous plants are fertile and produce a nearly full seed set if crossed with an appropriate pollen donor warranting the formation of balanced endosperm. While we have exploited *nrf4* to demonstrate that clonal seed production is possible in maize, it has to become more efficient for applications in plant breeding and seed production. In rice, experiments using *MiMe* in combination either *mtl*/*nld* or *BBML* expression in the egg cell produced ≤ 6% and ≤ 30% clonal seeds, respectively (*22*,*34*). Generating additional events of the latter in hybrid rice lines led to an increase in clonal seed production to ∼ 95% but was still associated with a significant decrease in seed set (*35*). In fact, the *OSD1* component of the *MiMe* mutant combination is also required for mitotic divisions, which can result in reduced seed set or lethality (*13*,*14,36–38*). Furthermore, depending on copy number, *MiMe* requires the combination of three or more meiotic mutants. In contrast, *nrf4* on its own displays apomeiosis and has a simple genetic inheritance, a highly penetrant non-reduction phenotype, and produces viable unreduced gametes, of which a significant percentage maintains maternal heterozygosity. These properties thus make *nrf4* a good candidate to engineer and further improve synthetic apomixis in maize.

## Materials and Methods

### Plant material and growth conditions

Seeds of the *bz1-Mum9* line (*39*) used to mobilize *Mu* elements by crossing it to the W22 inbred line were kindly provided by R. A. Martienssen (Cold Spring Harbor Laboratory); the W22 inbred line was received from E. Vollbrecht (Plant Gene Expression Center). W22 was crossed to *bz1-Mum9* and the F_1_ progeny were self-fertilized to generate about 3200 families segregating for new *Mu* insertions. 20 kernels from each family were planted and all fertile individuals were crossed to a tetraploid pollen donor carrying the *R1-nj* allele either by hand-pollination or open pollination in isolation fields. The 4n *R1-nj* line was created by crossing pollen of the diploid W23 *R1-nj* line (M142V from the Maize Genetics Cooperation Stock Center) with tetraploid females (N107B from Maize Genetics Cooperation Stock Center), selecting the few plump kernels that resulted from fertilization with spontaneously unreduced pollen, and breeding these tetraploids for several generations through sib- and self-pollinations to increase seed set, which was poor in the original tetraploids that were recovered. Two additional mutant *nrf4* alleles, namely *Mu* insertions PV0363-C07 and PV03129-A09, were obtained from Pioneer Hi-Bred’s Trait Utility System for Corn (TUSC). Inbred lines PHI1 to PHI8, into which *nrf4-1* was introgressed by > 6 backcrosses, were provided by DuPont-Pioneer. To generate the tetraploid *mtl*/*nld* mutant, we first treated the haploid-inducer line IHVR232 (DuPont-Pioneer) with colchicine using a standard procedure (*40*). The plants were male sterile and were thus pollinated with pollen from the tetraploid W23 *R1-nj* plants described above. The F_2_ generation of this cross was genotyped for homozygosity of the *mlt*/*nld* locus using markers and were confirmed as tetraploids by flow cytometry. Fast-flowering mini-maize (*41*) seeds were obtained from J. A. Birchler (University of Missouri). For immunostaining experiments, *nrf4-1* was introgressed into the mini-maize background by six backcrosses to have materials available year-round. The plants to generate the materials for the genetic screen were grown at Uplands Farm, New York (Cold Spring Habor Laboratory). The screens were performed at Uplands Farm and in fields of FAL Reckenholz near Zurich, Switzerland (Agroscope), and plants for genetic analyses were grown at FAL Reckenholz, in Juana Diaz, Puerto Rico (Illinois Crop Improvement Association), in Nayarit, Mexico (PV Winter Seed Services), or the former DuPont Stine-Haskell Research Center. Material for microscopy was grown in the ZOM greenhouse of the University of Zurich, Switzerland, under a 16 h light / 8 h dark cycle at 25°C during the day and 22°C during the night.

### Flow cytometry

To assess the ploidy level of dry kernels, the latter were ground for 5 s in a coffee bean grinder, Eldom Type EL 10. Around 50 mg of the powdered material was transferred to 20 mm Petri dishes. 600 μl of Otto I buffer, containing 0.1M citric acid and 0.1% Tween 20, was added and pipetted up and down several times. The nuclei suspension was filtered through CellTrics® 30 µm (Partec). 800 μl of CyStain UV Precise P Staining Buffer (Partec) containing DAPI was added to 200 μl of the nuclei suspension to stain the extracted nuclei. After 5 min incubation at room temperature, the nuclei suspension was analysed by CytoFlexS flow cytometer, using the NUV-diod as a light source and a 450/45 filter for detection.

### Cloning the *Nrf4* gene using SAIFF by sequencing

Fifteen plants with the *nrf4-1* phenotype producing a high percentage of plump seeds when crossed to tetraploid males were evaluated by high-throughput sequencing of their *Mu*-flanking regions. Briefly, DNA was isolated from leaf tissue and sheared on a Covaris E210 ultrasound sonicator (Covaris). An indexed adapter was then ligated to DNA fragments. The libraries underwent two rounds of PCR amplification using the 2x Phusion DNA Polymerase Mastermix (New England Biolabs), for 18 and 20 cycles of amplification, respectively. The first round of PCR was performed with a forward primer that matches the common region in all *Mu* elements (PHN130859-Muint26 AGA AGC CAA CGC CAW CGC CTC YAT TTC) and an adapter-specific reverse primer (PE1.0 AAT GAT ACG GCG ACC ACC GAG ATC TAC ACT CTT TCC CTA CAC GAC GCT CTT CCG ATC T). The second round of PCR was completed with the same primers, except that the flowcell site was attached to the forward primer (PHN130859-Muint26-PE2.0 CAA GCA GAA GAC GGC ATA CGA GAT AGA AGC CAA CGC CAW CGC CTC YAT TTC) to enable Illumina sequencing. Products of all samples were pooled and sequenced on an Illumina HiSeq2000 sequencer with a run configuration of single read 108 bases. *nrf4-1* samples had approximately 1.5-10 million reads for each sample. One *Mu*-flanking sequence was found to be present in all 15 plants, and thus most likely represented the insertion causing the *nrf4-1* non-reduction phenotype.

### Genotyping

DNA extractions were conducted using the CTAB method (*42*) with minor modifications. Briefly, a piece of a young leaf (0.5 cm^2^) was placed in a 1.2 ml tube and supplemented with glass beads (Roth). Tubes were cooled in liquid nitrogen and ground using a Retch ball mill for 1 min at 30 Hz in 96 tube racks. DNA was extracted with 2x CTAB buffer for 15 min at 65°C. After centrifugation, the supernatant was transferred into a new tube and extracted with an equal volume of chloroform. After phase separation by centrifugation, the upper phase was transferred to a new tube and DNA was precipitated by adding 0.7 volumes of isopropyl alcohol and incubated for 10 min at room temperature. The DNA was pelleted by centrifugation, rinsed in 75% ethanol, and diluted in 100ul of 1x TE. PCR was performed using the KAPA3G Plant PCR kit (Kapabiosystems), according to the manufacturer’s instructions. The PCR primers and conditions for wild-type and mutant alleles are summarized in Table S11.

### RT-PCR

The following tissues and organs were collected from mini-maize plants: ears stage 1 (approximately meiotic stage); ears stage 2-4 (approximately 2-8 nuclear embryo sac); ears stage 5 (mature embryo sac); silks; developing tassel (meiotic stage); tassel shedding pollen; mature anthers; pollen; seedling roots (4 days after germination); seedling shoots (4 days after germination); stalk of an adult plant, leaves. All samples were frozen in liquid nitrogen and cryogenically pulverized using glass beads and a Silamat S6 grinder (Ivoclar Vivadent AG). RNA was extracted using the TRIzol ^®^ Reagent (Ambion) and treated with TURBO DNA-*free*^TM^ DNase (Invitrogen) according to the manufacturer’s instructions. One mg of total RNA was subsequently reverse-transcribed to cDNA using SuperScript^TM^II Reverse Transcriptase (Invitrogen) with oligo-dT as a primer for cDNA synthesis. PCR was performed using KAPA3G Plant PCR kit (Kapabiosystems) according to the manufacturer’s recommendations. Primers and PCR conditions are shown in Table S11. The PCR products were separated by agarose gel electrophoresis in 1x TBE buffer, stained by ethidium bromide and documented using the GelDoc-It*^e^* Imaging System (UVP LLC **Analytik Jena AG)**.

### RNA *in situ* hybridization

Ribo-probe preparation: Total RNA extraction and cDNA synthesis was performed similarly to RT-PCR (see above). An *Nrf4* fragment was PCR-amplified with the following primers: pco657953f 3’-AAA CTT TTC GAA AAT CCC CCA CAG AAT CCC-5’ and pco657953r 3’-AAA ACC AC TGC ATG ATC ACC TCC GC-5’.

Subsequently, it was cloned into the pDRIVE vector between the Sp6 and T7 RNA polymerase promoters using the QIAGEN PCR cloning kit (Qiagen). Sense and antisense digoxygenin-UTP–labeled (DIG) riboprobes were generated by *in vitro* transcription using T7 or Sp6 RNA polymerase, respectively, and the DIG RNA labeling mix (Roche Diagnostics).

*In situ* hybridization: Ears of the W22 inbred line at megaspore mother cell stage were dissected into 4-8 longitudinal sections and fixed in 3:1 ethanol:acetic acid for 12-14 h at 4°C. Subsequently, the tissue was dehydrated in an ethanol series (70%, 90%, 100% 1 h each). Ethanol was substituted by 100% xylol in three consequent 1h incubations at RT, and the tissue was embedded in Paraplast extra at 56°C. Dehydration steps, xylol treatment, and Paraplast embedding were performed in a Leica ASP200 Vacuum tissue processor (Leica Microsystems). Nine μm thick sections of fixed and embedded *maize* florets were prepared using a microtome and transferred to ProbeOn Plus microscopy slides (Fischer). The tissue was dewaxed in Histo-Clear (National Diagnostics) by incubation of the slides for 2x 10 min at room temperature, followed by rehydration and Proteinase K treatment for 30 min at 37°C. Slides were post-fixed in 4% paraformaldehyde for 10 min, dehydrated in an ethanol series, and subjected to hybridization with the probes for 16 h at 55°C in a humid chamber. After hybridization, subsequent washes were performed, slides were blocked with Roche blocking buffer (Roche) and incubated with anti-DIG antibodies conjugated to alkaline phosphatase. We used NBT/BCIP substrate with 1 mM Levamisole for signal detection (Sigma-Aldrich). Pictures were captured on a Leica DMR microscope (Leica Microsystems), cropped, and processed in ImageJ.

### Phylogenetic analysis

The NRF4 protein sequence was used as a query to identify similar proteins in the non-redundant NCBI database of protein sequences using the BlastP algorithm. Protein sequence alignments and the phylogenetic tree were made using the CLC Main workbench suit.

### SNP array and analysis of the results

Two 6-mm leaf tissue samples were collected per maize seedling, and the leaf samples were placed in a polypropylene 96-well deep-well storage plate (VWR) containing several 4.6 mm steel beads. The plates were covered with a blotter pad secured in place using rubber bands, and the tissue was lyophilized and then ground using a GenoGrinder (Spex). DNA was isolated using a HotSHOT DNA Extraction Kit (Bento Bioworks) following the manufacturer’s instructions. SNP genotyping was performed using TaqMan SNP genotyping assays (ThermoFisher). Each assay contains sequence-specific forward and reverse primers to amplify the polymorphic sequence of interest. Two TaqMan minor groove binder probes with non-fluorescent quenchers were used: a first fluorophore-labelled probe to detect allele 1 sequence (for example, VIC) and a second fluorophore-labelled probe to detect allele 2 sequence (for example, FAM). PCR reactions containing 2.5 µl of TaqMan Universal PCR Master Mix (2x), 0.5 µl of each a forward and reverse primer (50 nM final concentration), 0.5 µl of assay probes, 0.5 µl of DNA and 0.5 µl of sterile, deionized water. The 5 µl reaction volumes were dispensed onto the Array Tape platform (LGC Limited) following the manufacturer’s instructions. PCR amplification was performed using a Nexar assay processing system, and assay results were scored resulting in bi-allelic polymorphism scores per SNP assay. The raw results of the SNP array can be found in Tables S1-S8. We built an SQL database for storing and analysing results of the SNP arrays. First, we identified the number of informative markers (M_I_) for each individual progeny plant as the number of markers that were called as heterozygous in the corresponding mother plant, which were also scored in the progeny plant, differed among them. Then, we identified the number of heterozygous markers (M_H_) in the progeny plant based on those that were heterozygous in the mother plant and remained heterozygous in the progeny plant. From these two values, we calculated the percentage of maintained maternal heterozygosity (MMH) as MMH=M_H_/M_I_*100% (Table S9). We considered 29 individuals with over 97% of maternal heterozygosity as clonal, because the SNP array has an error rate of approximately 2-3%. All 29 individuals were manually curated. If they had either two or more homozygous markers in the row at the end of any chromosome or three or more markers in a row in the middle of any chromosome arm, they were not qualified as clonal individuals (Table S10). Maternal heterozygosity maintenance histograms displayed in Fig.3 were plotted using the gplot2 package of *R*.

### Cytology of developing female florets

Fixation, Feulgen staining with Shiff’s solution, dissection, mounting, and microscopical analysis were performed as described previously (*42*).

### Immunolabelling of H3S10p

Immunostaining of H3S10p was performed using a protocol slightly differing from the previously described (*42,43*). Ears were fixed with freshly made BVO buffer containg 1% formaldehyde and 10% DMSO in PBS-Tween (0.1%) on ice. The samples were then dissected to release the ovule primordia and embedded in 5% acrylamide pads on microscope slides. The following steps included tissue clearing and fixation, cell wall digestion and permeabilization before incubation with the primary antibody against H3S10p with 1:100 dilution for 13-24 h, and then secondary antibody for around 48 h at 4°C. The samples were counterstained with propidium iodide and then mounted in Prolong Gold supplemented with 0.04mg/ml propidium iodide. Fluorescent signals in ovule primordia were recorded by Leica SP5 confocal laser-scanning microscopy.

## Acknowledgments

We thank the Maize Genetics Cooperation Stock Center (University of Illinois), the Trait Utility System for Corn (DuPont-Pioneer), R. A. Martienssen (Cold Spring Harbor Laboratory), and E. Vollbrecht (Iowa State University) for seeds, C. Westermann (University of Zurich) for editing of the manuscript, and C. Eichenberger, D. Guthörl, A. Bolaños, P. Kopf and S. Sowinski (DuPont-Pioneer) for general lab support. We are indebted to our greenhouse and field managers K. Huwiler (University of Zurich), T. Mulligan (Cold Spring Habor Laboratory), M. Menzi, J. Hiltbrunner, U. Buchmann, T. Huber (Agroscope), M. Widmann (DuPont-Pioneer) and C. Robledo (PV Winter Services) for taking care of the plants, and to numerous summer students and civil servants for help with field work.

## Author contributions

Conceptualization: U.G.; Formal analysis: N.C., G.A.B., M.W., V,G., U.G.; Funding acquisition: U.G., M.C.A.; Investigation: N.C., A.B., M.E.W., W.S., P.J.B, V.G., F.P., T.W.F., U.G.; Methodology: N.C., G.A.B., M.E.W., P.J.B, V.G., T.W.F., U.G.; Project administration: N.C., M.C.A., U.G.; Resources: U.G., M.C.A.; Supervision: U.G., M.C.A.; Visualization: N.C., G.A.B., W.S.; Writing – original draft: N.C., G.A.B., U.G.; Writing – review & editing: M.E.W., W.S., V.G., M.C.A.

## Data and materials availability

All data are available in the main text or the supplementary materials.

## Supplementary Materials

### The PDF file includes

Materials and Methods Figures S1 to S5 References (*39–44*)

### Other Supplementary Material for this manuscript includes the following

Tables S1 to S11

## Supplementary Figures

**Fig. S1.**
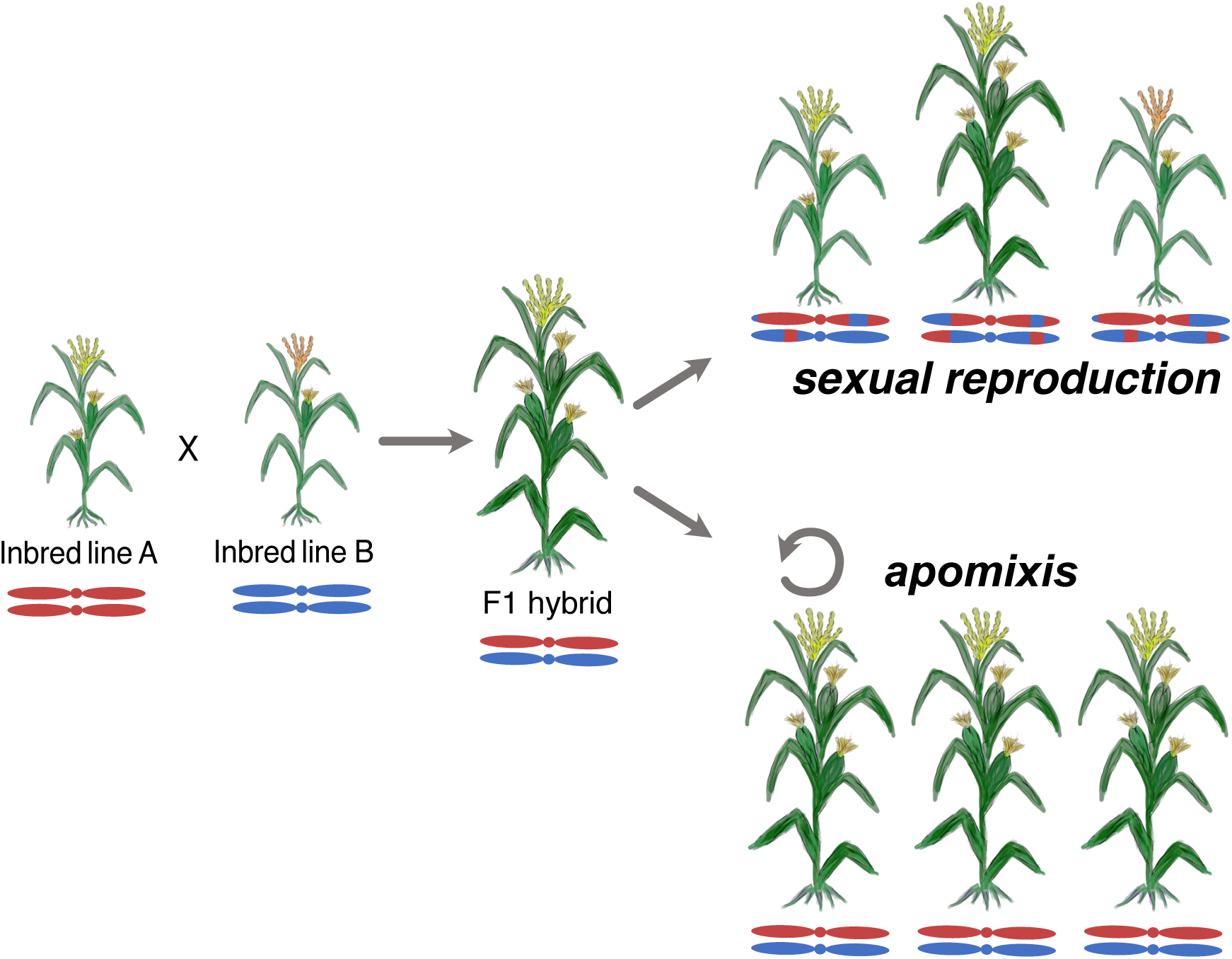
Schematic model of plant phenotypes and genotypes from different crosses. Left: The cross of two different homozygous inbred lines results in a F_1_ hybrid that displays drastic increase in yield compared to the two homozygous parental lines due to hybrid vigor. Right, top: Self-or cross-pollination of hybrid plants leads to a segregating population that loses heterozygosity in up to 50% of the loci and is accompanied by trait segregation and reduced yield. Right bottom: Apomixis produces clonal offspring. Apomictic hybrids maintain the heterozygous genetic composition at 100% of the loci, thus maintaining hybrid vigor.

**Fig. S2.**
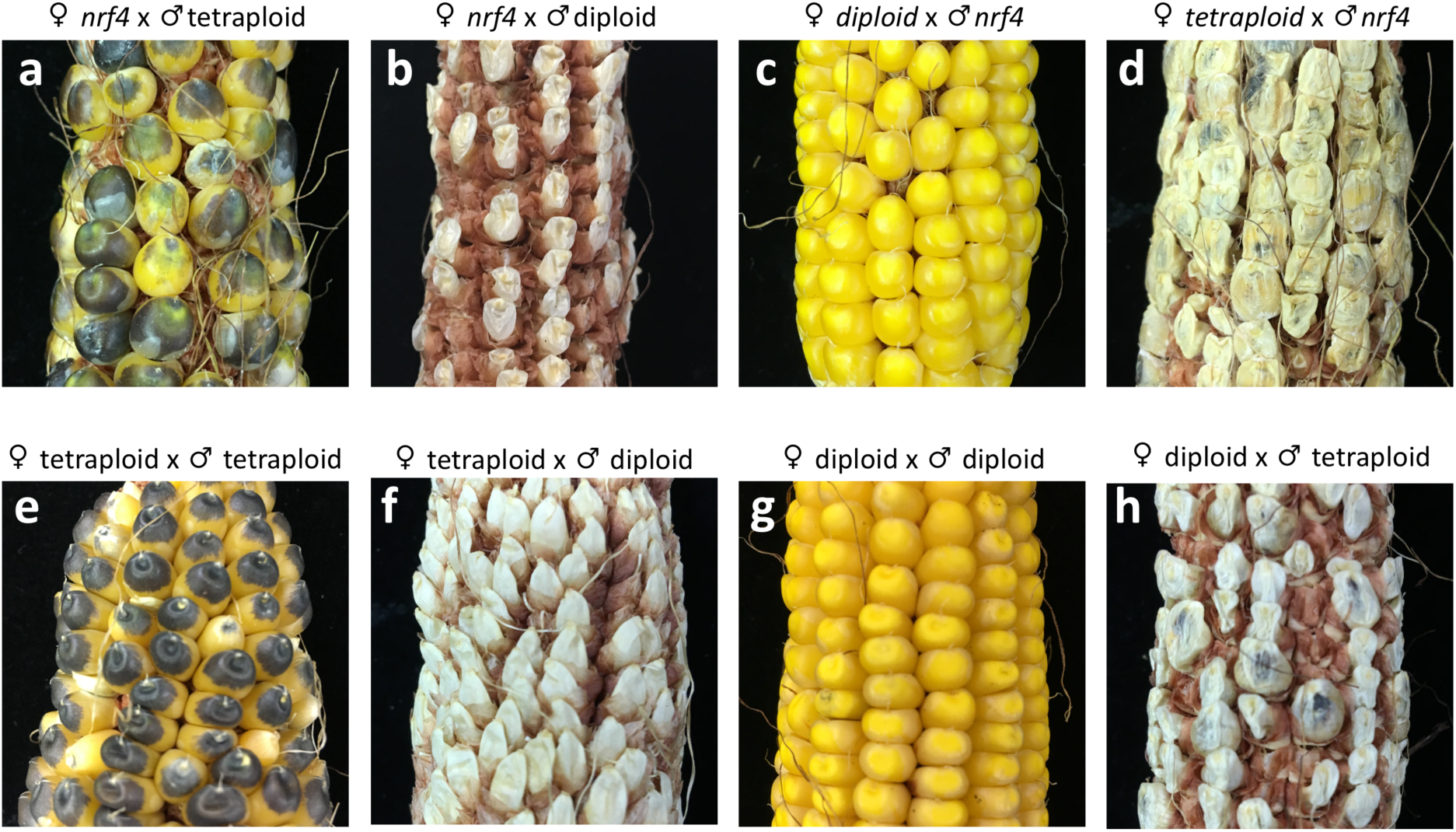
*nrf4* mutant phenotype when crossed to diploid or tetraploid tester lines in a reciprocal manner. a, Ear of a *nrf4* mutant plant fertilized by tetraploid (4n) *R1-nj* pollen. All kernels are plump and have the *R1-nj* pattern of pigmentation similar to the situation when wild-type 4n is crossed to pollen from a 4n pollen donor (**e**), indicating non-reduction of the female gametophyte in the *nrf4* mutant. **b**, Ear from a *nrf4* mutant plant crossed with pollen from a wild-type diploid (2n) plant. All kernels aborted because of the different ploidies of female (2n) and male (1n) gametophytes, similar to the cross between 4n and 2n parents (**f**). **c**, Ear from wild-type 2n plant crossed with *nrf4* pollen carries normal kernels, similar to a cross between two 2n plants (**g**), but not to a cross between a 2n female and a 4n male (**h**), suggesting that male gametophyte is reduced in the *nrf4* mutant. **d**, Ear from 4n plant pollinated with *nrf4* pollen. All kernels abort development, because the ploidies of female (2n) and male (1n) gametophytes do not match.

**Fig. S3.**
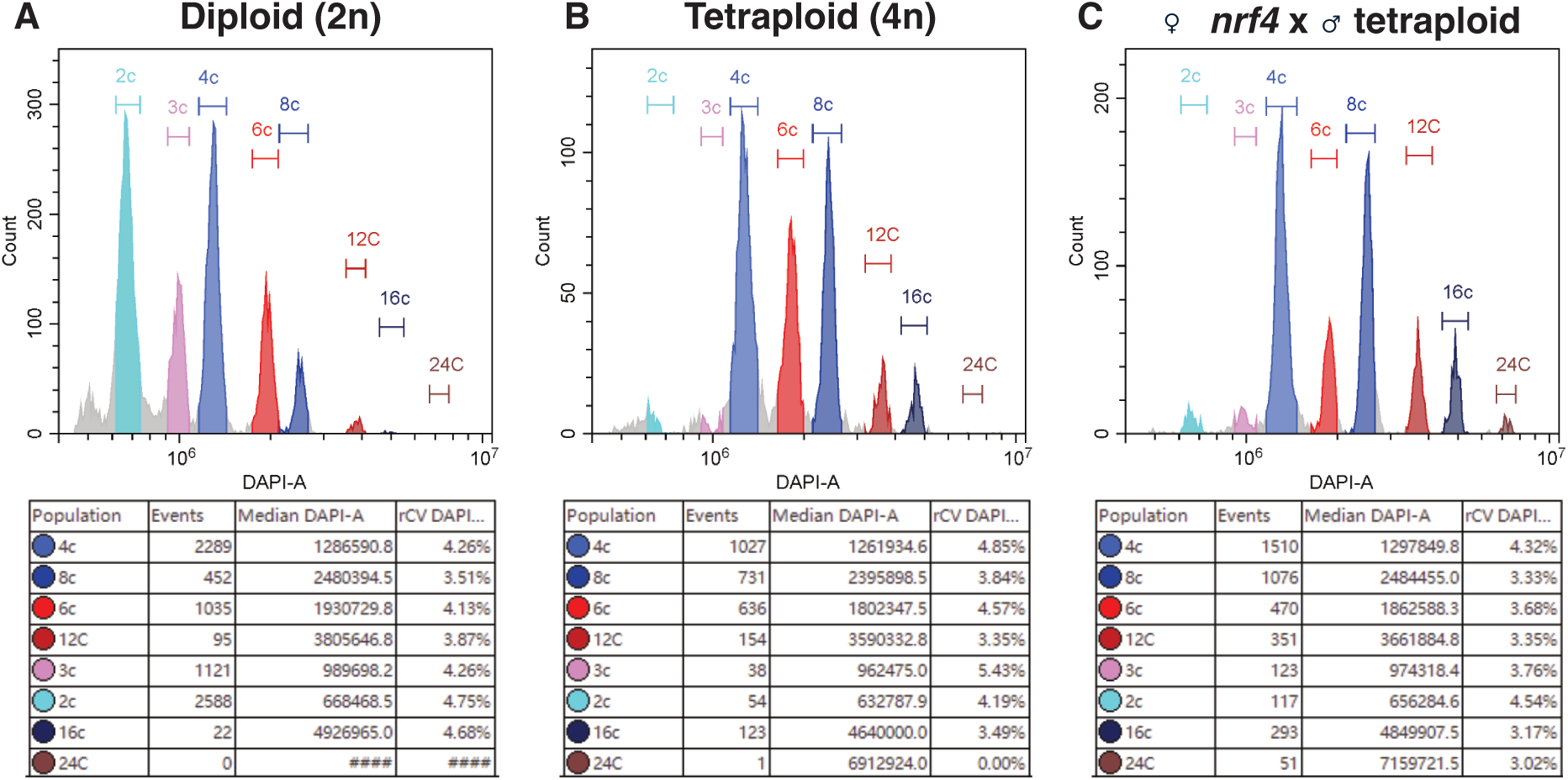
Ploidy histograms of plump kernels. **(A**) Diploid (2n) W22; (**B**) Tetraploid (4n) W23 *R1-nj*; (**C**) ♀*nrf4* x ♂4n W23 *R1-nj*. Each histogram represents the number of nuclei (Y-axis) with a specific genome size, reflected by relative fluorescence of DAPI stained nuclei (X-axis). Peaks of the blue and red palette represent embryo and endosperm nuclei, respectively, in different phases of the cell cycle (G1, G2, endoreduplication). The 2n kernel contained a 2n embryo (major peaks at 2c and 4c, corresponding to the G1 and G2 phases of cell cycle) and triploid (3n) endosperm (3c and 6c peaks). In contrast, both the 4n kernel and *nrf4* x 4n *R1-nj* kernel contained a 4n embryo (major peaks are 4c and 8c, whereas the 2c peak is not present) and a hexaploidy (6n) endosperm (6c and 12c peaks). Statistical data for the histograms is displayed below each histogram.

**Fig. S4.**
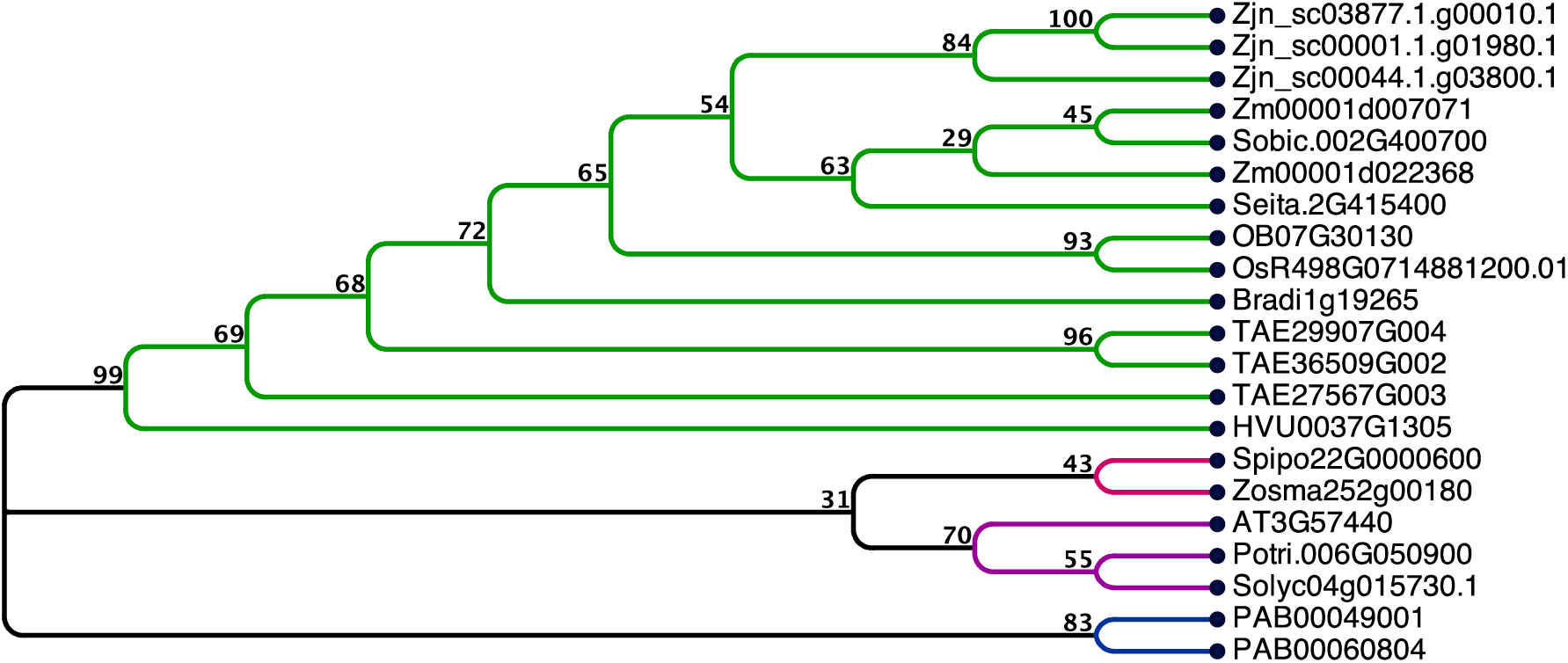
Phylogenetic tree of NRF4 homologs in different species. Green branch – proteins from grass (Poaceae) species, red branch – non-grass monocots, magenta – dicots, blue – gymnosperms. Zjn - *Zoysia japonica*; Zm - *Zea mays*; Sobic - *Sorghum bicolor*; Seita - *Setaria italica*; OB - *Oryza brachyantha*; Os - *Oryza sativa*; Brsdi - *Brachypodium distachyon*; TAE - *Triticum aestivum*; HVU - *Hordeum vulgare*; Spipo - *Spirodela polyrhiza*; Zosma - *Zostera marina*; AT - *Arabidopsis thaliana*; Potri - *Populus trichocarpa*, Solic - *Solanum lycopersicum*; PAB - *Picea abies.* The *Nrf4* gene is marked by a star.

**Fig. S5.**
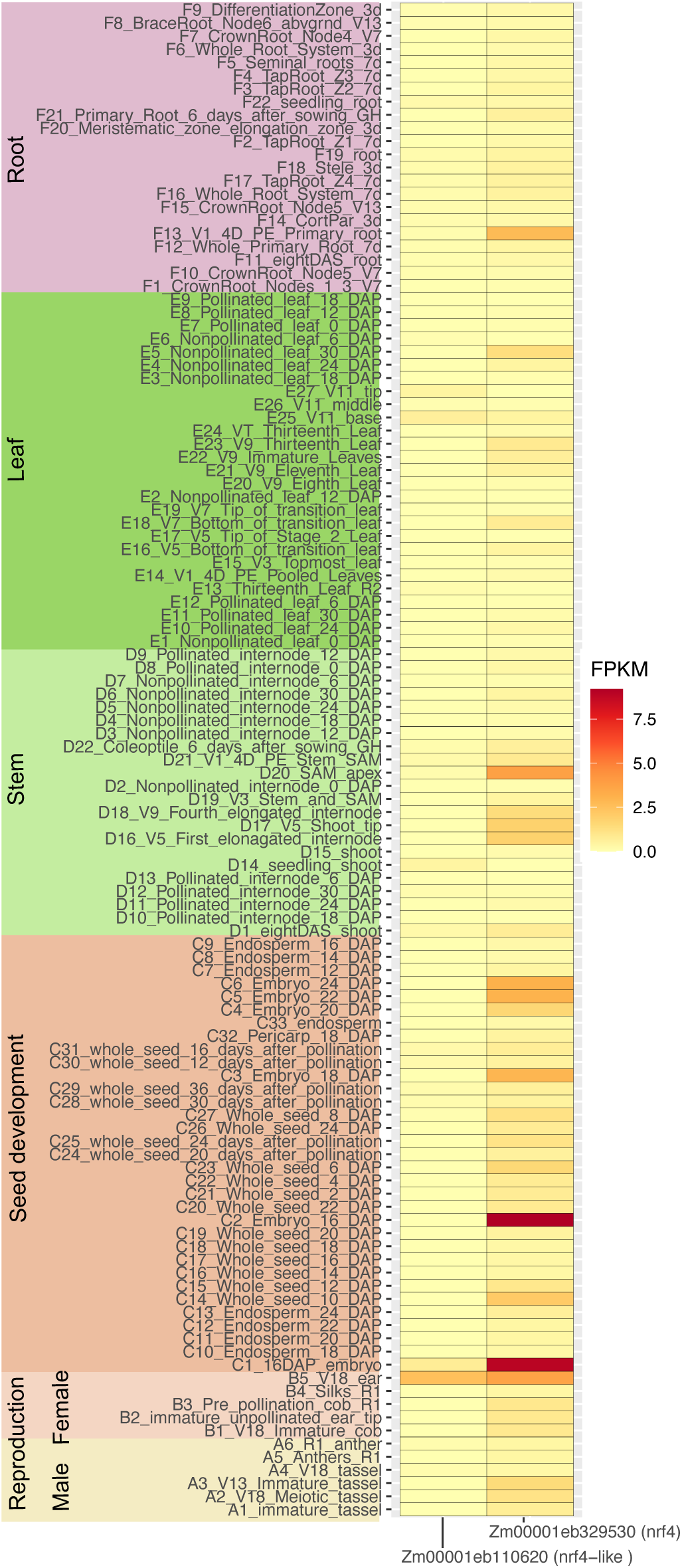
Expression levels of *Nrf4 (Zm00001eb329530)* and the dulicate *Nrf4-like* gene (Zm00001eb110620) in different plant tissues based on *in silico* analysis of diverse RNA-Seq experiments. Expression values (FPKM) were obtained from the MaizeMine v1.6 web-tool and visualized by ggplot2 package for R.

