## Supplementary tables 1-11 for "Generating clonal seeds using *non-reduction in female4* (*nrf4*), a novel meiotic mutant of maize"

### Table S1. Results of the SNP array analasis

| | | | | | Chr | 1 | 1 | 1 | 1 | 1 | 1 | 1 | 1 | 1 | 1 | 1 | 1 | 1 | 1 | 1 | 1 | 1 | 1 | 1 | 1 | 1 | 1 | 1 | 1 | 1 | 1 | 1 | 1 | 1 | 1 | 1 | 1 | 1 | 1 | 1 | 1 | 1 | 1 | 1 | 1 | 1 | 1 | 1 | 1 | 1 | 1 | 1 | 1 |
|---|---|---|---|---|---|---|---|---|---|---|---|---|---|---|---|---|---|---|---|---|---|---|---|---|---|---|---|---|---|---|---|---|---|---|---|---|---|---|---|---|---|---|---|---|---|---|---|---|---|---|---|---|---|
| | | | | | Distance, cM | 0 | 2.6 | 4.5 | 4.8 | 4.8 | 13 | 23 | 32 | 42 | 52 | 59 | 69 | 80 | 90 | 99 | 110 | 120 | 128 | 137 | 137 | 138 | 138 | 138 | 147 | 157 | 165 | 173 | 182 | 193 | 203 | 207 | 211 | 220 | 230 | 240 | 249 | 260 | 268 | 277 | 287 | 296 | 300 | 306 | 314 | 316 | 317 | 319 | 319 |
| plant_id | mom_gen | dad_gen | mother_progeny | family_ | | | | | | | | | | | | | | | | | | | | | | | | | | | | | | | | | | | | | | | | | | | | | | | | | |
| 2013_P1-1 | PHI1 | PHI2 | mother | 2013_P1-1 | | H | T | G | T | T | H | G | H | H | G | T | H | A | G | H | H | H | T | C | C | T | T | G | H | T | H | H | H | G | G | A | T | T | T | T | EQV | C | H | G | G | H | H | A | T | G | H | A | C |
| 2013_P1-1-1 | PHI1 | PHI2 | progeny | 2013_P1-1 | | H | T | G | T | T | H | G | H | H | G | T | H | A | G | H | H | H | T | C | C | T | T | G | H | T | H | H | H | G | G | A | T | T | T | T | T | C | H | G | G | H | H | A | T | G | H | A | C |
| 2013_P1-3 | PHI1 | PHI2 | mother | 2013_P1-1 | | H | T | G | T | T | H | G | H | H | G | T | H | A | G | H | H | H | T | C | C | T | T | G | H | T | H | H | H | G | G | A | T | EQV | T | T | H | C | LS | G | G | H | H | A | T | G | H | A | C |
| 2013_P1-3-1 | PHI1 | PHI2 | progeny | 2013_P1-3 | | H | T | G | H | T | H | G | H | H | G | T | H | A | G | H | A | EQV | T | C | C | T | T | G | H | T | H | H | H | H | G | H | EQV | EQV | T | T | H | C | LS | G | G | C | A | A | T | G | H | A | H |
| 2013_P1-3-2 | PHI1 | PHI2 | progeny | 2013_P1-3 | | H | T | G | H | T | H | G | H | H | G | T | H | A | G | H | A | H | T | C | C | T | T | G | H | EQV | H | H | G | G | G | H | T | EQV | T | T | H | H | LS | G | G | H | H | A | T | G | T | A | H |
| 2013_P1-3-3 | PHI1 | PHI2 | progeny | 2013_P1-3 | | H | T | G | H | T | H | G | H | H | G | T | H | A | G | H | H | H | T | C | C | T | T | G | H | EQV | H | H | H | G | G | H | T | EQV | T | T | H | H | EQV | G | G | H | H | A | T | G | EQV | A | H |
| 2013_P1-3-4 | PHI1 | PHI2 | progeny | 2013_P1-3 | | H | T | G | H | T | H | G | H | H | G | T | H | A | G | H | H | H | T | C | C | T | T | G | H | EQV | H | H | H | G | G | H | T | T | T | T | EQV | H | LS | G | G | EQV | H | A | T | G | H | A | H |
| 2013_P1-3-5 | PHI1 | PHI2 | progeny | 2013_P1-3 | | H | T | G | H | T | H | G | EQV | H | G | T | H | A | H | EQV | H | EQV | T | C | C | T | T | G | H | EQV | H | G | H | G | G | H | EQV | EQV | T | T | EQV | C | EQV | G | G | H | H | A | T | G | EQV | A | H |
| 2013_P1-6 | PHI1 | PHI2 | mother | 2013_P1-6 | | H | T | G | T | T | H | G | H | H | G | T | H | A | A | H | H | H | T | C | C | T | T | G | H | T | H | H | H | G | G | A | T | T | T | T | EQV | C | H | G | G | H | H | A | T | G | H | A | C |
| 2013_P1-6-1 | PHI1 | PHI2 | progeny | 2013_P1-6 | | H | T | G | T | T | H | G | H | H | G | T | H | A | A | H | H | H | T | C | C | T | T | G | H | T | H | H | H | G | G | A | T | T | T | T | H | C | H | G | G | C | A | A | T | G | H | A | C |
| 2013_P1-7 | PHI1 | PHI2 | mother | 2013_P1-7 | | H | T | G | T | T | H | G | H | H | G | T | H | A | G | H | H | H | T | C | C | T | T | G | H | T | H | H | H | G | G | A | T | T | T | T | EQV | C | H | G | G | H | H | A | T | G | H | A | C |
| 2013_P1-7-1 | PHI1 | PHI2 | progeny | 2013_P1-7 | | H | T | G | T | T | H | G | H | H | G | T | H | A | G | H | H | H | T | C | C | T | T | G | H | T | H | H | H | G | G | A | T | T | T | T | H | C | LS | G | G | C | A | A | T | G | C | A | C |
| 2013_P1-8 | PHI1 | PHI2 | mother | 2013_P1-8 | | H | T | G | T | T | H | G | H | H | G | T | H | A | G | H | H | H | T | C | C | T | T | G | H | T | H | H | H | G | G | A | T | T | T | T | H | C | H | EQV | G | H | H | A | T | G | H | A | C |
| 2013_P1-8-1 | PHI1 | PHI2 | progeny | 2013_P1-8 | | H | T | G | H | T | H | G | H | H | G | T | H | A | G | H | A | H | T | C | C | T | T | G | H | EQV | H | H | H | H | G | H | T | T | T | T | H | H | H | G | G | H | H | A | T | G | H | A | H |
| 2013_P1-8-2 | PHI1 | PHI2 | progeny | 2013_P1-8 | | G | T | G | T | T | C | G | G | H | G | T | H | A | G | H | H | H | T | C | C | T | T | G | H | T | H | H | H | G | G | A | T | T | T | T | H | C | H | G | G | C | A | A | T | G | H | A | C |
| 2013_P1-8-3 | PHI1 | PHI2 | progeny | 2013_P1-8 | | H | T | G | H | T | H | G | H | H | G | T | C | A | G | H | H | C | T | C | C | T | T | G | A | EQV | G | H | H | H | G | H | T | T | T | T | T | H | H | G | G | C | A | A | T | G | H | A | H |
| 2013_P1-8-4 | PHI1 | PHI2 | progeny | 2013_P1-8 | | H | T | G | T | T | H | G | H | H | G | T | H | A | G | H | H | H | T | C | C | T | T | G | H | EQV | H | H | H | G | G | EQV | T | T | T | T | T | C | LS | G | G | H | H | A | T | G | H | A | C |
| 2013_P1-8-5 | PHI1 | PHI2 | progeny | 2013_P1-8 | | H | T | G | H | T | H | G | H | H | G | T | H | A | G | H | H | H | T | C | C | T | T | G | H | EQV | H | H | EQV | H | G | H | T | T | T | T | H | C | H | G | G | H | H | A | T | G | H | A | H |
| 2013_P1-9 | PHI1 | PHI2 | mother | 2013_P1-9 | | H | T | G | T | T | H | G | H | H | G | T | H | A | A | H | H | H | T | C | C | T | T | G | H | EQV | H | H | H | G | G | A | T | T | T | EQV | T | C | H | G | G | H | H | A | T | G | H | A | C |
| 2013_P1-9-1 | PHI1 | PHI2 | progeny | 2013_P1-9 | | A | T | G | T | T | G | G | A | H | G | T | H | A | A | C | C | C | T | C | C | T | T | G | A | T | G | G | A | G | G | A | T | T | T | T | T | C | G | G | G | T | G | A | T | G | T | A | C |
| 2013_P1-9-2 | PHI1 | PHI2 | progeny | 2013_P1-9 | | H | T | G | T | T | H | G | H | H | G | T | H | A | A | H | H | H | T | C | C | T | T | G | A | T | G | G | A | G | G | A | T | T | T | T | H | C | LS | G | G | H | H | A | T | G | C | A | C |
| 2013_P1-9-3 | PHI1 | PHI2 | progeny | 2013_P1-9 | | H | T | G | T | T | H | G | H | H | G | T | H | A | A | C | C | C | T | C | C | T | T | G | H | T | C | A | G | G | G | A | T | T | T | T | H | C | H | G | G | T | G | A | T | G | T | A | C |
| 2013_P1-10 | PHI1 | PHI2 | mother | 2013_P1-10 | | H | T | G | T | T | H | G | H | H | G | T | H | A | G | H | H | H | T | C | C | T | T | G | H | T | H | EQV | H | G | G | A | T | T | T | T | H | C | H | G | G | H | H | A | T | G | H | A | C |
| 2013_P1-10-1 | PHI1 | PHI2 | progeny | 2013_P1-10 | | H | T | G | T | T | H | G | H | G | G | T | C | A | G | H | H | H | T | C | C | T | T | G | T | T | H | H | H | G | G | A | T | T | T | T | H | C | H | G | G | H | H | A | T | G | H | A | C |
| 2013_P1-16 | PHI1 | PHI2 | mother | 2013_P1-16 | | H | T | G | T | T | H | G | H | H | G | T | H | A | G | H | H | H | T | C | C | T | T | G | H | T | H | H | H | G | G | A | T | T | T | T | EQV | C | H | EQV | G | H | H | A | T | G | H | A | C |
| 2013_P1-16-1 | PHI1 | PHI2 | progeny | 2013_P1-16 | | H | T | G | T | T | H | G | H | H | G | T | A | A | G | H | H | H | T | C | C | T | T | G | T | T | C | A | G | G | G | A | T | T | T | T | T | C | LS | G | G | C | A | A | T | G | C | A | C |
| 2013_P1-16-2 | PHI1 | PHI2 | progeny | 2013_P1-16 | | G | T | G | T | T | C | G | G | G | G | T | C | A | G | H | H | H | T | C | C | T | T | G | T | T | C | A | G | G | G | A | T | T | T | T | T | C | H | G | G | H | H | A | T | G | H | A | C |
| 2013_P1-17 | PHI1 | PHI2 | mother | 2013_P1-17 | | H | T | G | T | T | H | G | H | H | G | T | H | A | G | H | H | H | T | C | C | T | T | G | H | T | H | H | H | G | G | A | T | T | T | T | T | C | LS | G | G | H | H | A | T | G | H | A | C |
| 2013_P1-17-1 | PHI1 | PHI2 | progeny | 2013_P1-17 | | H | T | G | T | T | H | G | H | H | G | T | C | A | A | C | C | C | T | C | C | T | T | G | A | T | G | H | H | G | G | A | T | T | T | T | T | C | LS | G | G | C | A | A | T | G | C | A | C |
| 2013_P1-17-2 | PHI1 | PHI2 | progeny | 2013_P1-17 | | H | T | G | T | T | H | G | H | H | G | T | H | A | G | H | A | G | T | C | C | T | T | G | T | T | C | H | H | G | G | A | T | T | T | T | H | C | LS | G | G | H | H | A | T | G | H | A | C |
| 2013_P1-19 | PHI1 | PHI2 | mother | 2013_P1-19 | | H | T | G | T | T | H | G | H | H | G | T | H | A | G | H | H | H | T | C | C | T | T | G | H | T | H | H | H | G | G | A | T | T | T | T | H | C | LS | G | G | H | H | A | T | G | H | A | C |
| 2013_P1-19-1 | PHI1 | PHI2 | progeny | 2013_P1-19 | | G | T | G | T | T | C | G | H | H | G | T | H | A | A | C | C | C | T | C | C | T | T | G | H | T | C | A | G | G | G | A | T | T | T | T | T | C | LS | G | G | H | H | A | T | G | H | A | C |
| 2013_P1-19-2 | PHI1 | PHI2 | progeny | 2013_P1-19 | | H | T | G | T | T | H | G | H | H | G | T | H | A | G | C | C | C | T | C | C | T | T | G | H | T | H | H | H | G | G | A | T | T | T | T | H | C | H | G | G | H | H | A | T | G | H | A | C |
| 2013_P1-19-3 | PHI1 | PHI2 | progeny | 2013_P1-19 | | H | T | G | EQV | T | H | G | H | H | G | T | H | A | G | H | H | H | T | C | C | T | T | G | H | T | H | H | H | G | G | A | T | T | T | T | EQV | C | H | G | G | H | H | A | T | G | H | A | C |
| 2013_P1-19-4 | PHI1 | PHI2 | progeny | 2013_P1-19 | | H | EQV | G | T | T | H | G | H | H | G | T | H | A | G | H | H | H | T | C | C | T | T | G | H | T | H | H | H | G | G | A | T | T | T | T | H | C | LS | G | G | C | A | A | T | G | H | A | C |
| 2013_P1-19-5 | PHI1 | PHI2 | progeny | 2013_P1-19 | | H | T | EQV | T | T | H | G | H | A | G | T | A | A | G | T | A | G | T | C | C | T | T | G | T | T | C | A | G | G | G | A | T | T | T | T | T | C | H | G | G | C | A | A | T | G | C | A | C |
| 2013_P1-20 | PHI1 | PHI2 | mother | 2013_P1-20 | | H | T | G | T | T | H | G | H | H | G | T | H | A | G | H | H | H | T | C | C | T | T | G | H | EQV | H | H | H | G | G | A | T | T | T | T | EQV | C | H | G | G | H | H | A | T | G | H | A | C |
| 2013_P1-20-1 | PHI1 | PHI2 | progeny | 2013_P1-20 | | H | T | G | T | T | H | G | H | H | G | T | H | A | G | H | H | H | T | C | C | T | T | G | H | T | H | H | H | G | G | A | T | T | T | T | H | C | LS | G | G | H | H | A | T | G | H | A | C |
| 2013_P1-20-2 | PHI1 | PHI2 | progeny | 2013_P1-20 | | A | T | G | T | T | G | G | A | A | G | T | H | A | G | H | H | H | T | C | C | T | T | G | H | T | H | H | H | G | G | A | T | T | T | T | T | C | LS | G | G | C | A | A | T | G | C | A | C |
| 2013_P1-21 | PHI1 | PHI2 | mother | 2013_P1-21 | | H | T | G | T | T | H | G | H | H | G | T | H | A | G | H | H | H | T | C | C | T | T | G | H | T | H | H | H | G | G | A | T | T | T | T | H | C | LS | G | G | H | H | A | T | G | H | A | C |
| 2013_P1-21-1 | PHI1 | PHI2 | progeny | 2013_P1-21 | | H | T | G | T | T | H | G | G | G | G | T | C | A | G | C | C | C | T | C | C | T | T | G | A | T | H | H | H | G | G | A | T | T | T | T | G | C | LS | G | G | H | H | A | T | G | H | A | C |
| 2013_P1-21-2 | PHI1 | PHI2 | progeny | 2013_P1-21 | | H | T | G | T | T | H | G | G | G | G | T | C | A | A | C | H | C | T | C | C | T | T | G | A | T | H | H | H | G | G | A | T | T | T | T | T | C | LS | G | G | C | A | A | T | G | H | A | C |
| 2013_P1-21-3 | PHI1 | PHI2 | progeny | 2013_P1-21 | | H | T | G | T | T | H | G | H | H | G | T | A | A | G | T | A | G | T | C | C | T | T | G | T | T | C | A | G | G | G | A | T | T | T | T | G | C | G | G | G | H | H | A | T | G | H | A | C |
| 2013_P1-21-4 | PHI1 | PHI2 | progeny | 2013_P1-21 | | A | T | G | T | T | G | G | A | H | G | T | H | A | A | C | C | C | T | C | C | T | T | G | A | T | G | G | A | G | G | A | T | T | T | T | H | C | LS | G | G | H | H | A | T | G | H | A | C |
| 2013_P1-22 | PHI1 | PHI2 | mother | 2013_P1-22 | | H | T | G | T | T | H | G | H | H | G | T | H | A | G | H | H | H | T | C | C | T | T | G | H | T | H | H | H | G | G | A | T | T | T | T | EQV | C | LS | G | G | H | H | A | T | G | H | A | C |
| 2013_P1-22-1 | PHI1 | PHI2 | progeny | 2013_P1-22 | | H | T | G | H | T | H | G | H | H | G | T | H | A | EQV | H | H | H | T | C | C | T | T | G | H | EQV | H | H | H | G | G | H | EQV | T | T | T | H | H | EQV | G | G | H | H | A | T | H | H | A | H |
| 2013_P1-22-2 | PHI1 | PHI2 | progeny | 2013_P1-22 | | H | T | G | T | T | H | G | H | H | G | T | H | A | G | H | H | G | T | C | C | T | T | G | T | T | C | A | G | G | G | A | T | T | T | T | H | C | LS | G | G | H | H | A | T | G | H | A | C |
| 2013_P1-25 | PHI1 | PHI2 | mother | 2013_P1-25 | | H | T | G | T | T | H | G | LS | H | G | T | H | A | A | H | H | H | T | C | C | T | T | G | H | T | H | H | H | G | G | A | T | T | T | T | H | C | LS | G | G | H | H | A | T | G | H | A | C |
| 2013_P1-25-1 | PHI1 | PHI2 | progeny | 2013_P1-25 | | H | T | G | T | T | H | G | H | H | G | T | H | A | A | H | H | H | T | C | C | T | T | G | H | T | H | H | H | G | G | A | T | T | T | T | EQV | C | LS | G | G | H | H | A | T | G | H | A | C |
| 2013_P1-25-2 | PHI1 | PHI2 | progeny | 2013_P1-25 | | G | T | G | T | T | C | G | G | H | G | T | H | A | A | H | H | H | T | C | C | T | T | G | H | T | H | H | H | G | G | A | T | T | T | T | H | C | H | G | G | C | A | A | T | G | C | A | C |
| 2013_P1-25-3 | PHI1 | PHI2 | progeny | 2013_P1-25 | | H | T | G | T | T | H | G | H | H | G | T | A | A | A | T | A | G | T | C | C | T | T | G | T | T | C | A | G | G | G | A | T | T | T | T | T | C | H | G | G | H | H | A | T | G | H | A | C |
| 2013_P1-25-4 | PHI1 | PHI2 | progeny | 2013_P1-25 | | H | T | G | T | T | H | G | H | H | G | T | H | A | A | H | H | H | T | C | C | T | T | G | H | T | H | H | H | G | G | A | T | T | T | T | EQV | C | LS | G | G | H | H | A | T | G | H | A | C |
| 2013_P1-25-5 | PHI1 | PHI2 | progeny | 2013_P1-25 | | H | T | G | T | T | H | G | G | G | G | T | C | A | A | H | H | G | T | C | C | T | T | G | H | T | H | H | H | G | G | A | T | T | T | T | T | C | H | G | G | H | H | A | T | G | H | A | C |
| 2013_P1-25-6 | PHI1 | PHI2 | progeny | 2013_P1-25 | | G | T | G | T | T | C | G | G | G | G | T | H | A | A | H | H | H | T | C | C | T | T | G | H | T | H | H | H | G | G | A | T | T | T | T | H | C | H | G | G | H | H | A | T | G | H | A | C |
| 2013_P1-26 | PHI1 | PHI2 | mother | 2013_P1-26 | | H | T | G | T | T | H | G | H | H | G | T | H | A | G | H | H | H | T | C | C | T | T | G | H | T | H | H | H | G | G | A | T | T | T | T | H | C | H | G | G | H | H | A | T | G | H | A | C |
| 2013_P1-26-1 | PHI1 | PHI2 | progeny | 2013_P1-26 | | H | T | G | T | T | H | G | H | H | G | T | H | A | G | H | C | C | T | C | C | T | T | G | A | T | G | G | A | G | G | A | T | T | T | T | T | C | EQV | G | G | C | A | A | T | G | C | A | C |
| 2013_P1-27 | PHI1 | PHI2 | mother | 2013_P1-27 | | H | T | G | T | T | H | G | H | H | G | T | H | A | A | H | H | H | T | C | C | T | T | G | H | T | H | H | H | G | G | A | T | T | T | T | H | C | H | G | G | H | H | A | T | G | H | A | C |
| 2013_P1-27-1 | PHI1 | PHI2 | progeny | 2013_P1-27 | | A | T | G | T | T | G | G | H | H | G | T | H | A | A | H | C | C | T | C | C | T | T | G | A | T | G | G | A | G | G | A | T | T | T | T | H | C | LS | G | G | H | G | A | T | G | T | A | C |
| 2013_P1-28 | PHI1 | PHI2 | mother | 2013_P1-28 | | H | T | G | T | T | H | G | H | H | G | T | H | A | A | H | H | H | T | C | C | T | T | G | H | T | H | H | H | G | G | A | T | T | T | T | H | C | H | G | G | H | H | A | T | G | H | A | C |
| 2013_P1-28-1 | PHI1 | PHI2 | progeny | 2013_P1-28 | | H | T | G | T | T | H | G | H | H | G | T | H | A | A | H | H | C | T | C | C | T | T | G | A | T | G | G | A | G | G | A | T | T | T | T | H | C | H | G | G | H | H | A | T | G | H | A | C |
| 2013_P1-31 | PHI1 | PHI2 | mother | 2013_P1-31 | | H | T | G | T | T | H | G | H | H | G | T | H | A | A | H | H | H | T | C | C | T | T | G | H | T | EQV | H | H | G | G | A | T | T | T | T | H | C | LS | EQV | G | H | H | A | T | G | H | A | C |
| 2013_P1-31-1 | PHI1 | PHI2 | progeny | 2013_P1-31 | | H | T | G | T | T | H | G | H | H | G | T | H | A | A | T | A | G | T | C | C | T | T | G | H | T | H | H | H | G | G | A | T | T | T | T | H | C | H | G | G | H | H | A | T | G | H | A | C |
| 2013_P1-31-2 | PHI1 | PHI2 | progeny | 2013_P1-31 | | H | T | G | T | T | H | G | H | H | G | T | H | A | A | H | H | H | T | C | C | T | T | G | A | T | G | G | A | G | G | A | T | T | T | T | T | C | G | G | G | H | H | A | T | G | H | A | C |
| 2013_O4 | PHI2 | PHI2 | ancestor | 2013_O4 | | G | T | G | T | T | C | G | G | G | G | T | C | A | A | C | C | C | T | C | C | T | T | G | A | T | G | G | A | G | G | A | T | T | T | T | G | C | LS | G | G | T | G | A | T | G | T | A | C |
| 2013_O1 | PHI1 | PHI1 | ancestor | 2013_O1 | | A | T | G | T | T | G | G | A | A | G | T | A | A | G | T | A | G | T | C | C | T | T | G | T | T | C | A | G | G | G | A | T | T | T | T | T | C | LS | G | G | C | A | A | T | G | C | A | C |
| 2013_O1_O4 | PHI1 | PHI2 | ancestor | 2013_O1_O4 | | H | T | G | T | T | H | G | H | H | G | T | H | A | G | H | H | H | T | C | C | T | T | G | H | T | H | H | H | G | G | A | T | T | T | T | H | C | H | G | G | H | H | A | T | G | H | A | C |

| | Chr | 2 | 2 | 2 | 2 | 2 | 2 | 2 | 2 | 2 | 2 | 2 | 2 | 2 | 2 | 2 | 2 | 2 | 2 | 2 | 2 | 2 | 2 | 2 | 2 | 2 | 2 | 2 | 2 | 2 |
|---|---|---|---|---|---|---|---|---|---|---|---|---|---|---|---|---|---|---|---|---|---|---|---|---|---|---|---|---|---|---|
| | Distance, cM | 1.7 | 2.2 | 6.1 | 7.6 | 7.7 | 17 | 27 | 35 | 44 | 55 | 64 | 73 | 81 | 86 | 94 | 104 | 114 | 123 | 129 | 129 | 129 | 129 | 129 | 137 | 146 | 156 | 164 | 173 | 183 |
| plant_id | | | | | | | | | | | | | | | | | | | | | | | | | | | | | | |
| 2013_P1-1 | | H | G | H | H | G | G | A | A | H | C | H | G | H | T | C | H | H | H | C | H | G | C | T | H | H | T | T | G | H |
| 2013_P1-1-1 | | H | G | H | H | G | G | A | A | H | C | H | G | H | T | C | H | H | H | C | H | G | C | T | H | H | T | T | G | H |
| 2013_P1-3 | | H | G | H | H | G | G | A | A | H | C | H | H | H | T | C | H | H | H | C | H | G | C | T | H | H | T | T | EQV | H |
| 2013_P1-3-1 | | H | G | H | H | G | H | A | A | H | H | H | EQV | H | H | C | H | H | H | C | G | G | C | T | G | A | T | T | G | C |
| 2013_P1-3-2 | | H | G | H | H | G | H | A | A | H | C | H | H | T | H | C | EQV | H | H | C | EQV | G | H | T | G | A | T | T | G | H |
| 2013_P1-3-3 | | H | G | H | EQV | G | G | A | A | A | C | EQV | G | H | H | C | EQV | H | EQV | C | H | G | EQV | T | H | H | T | T | G | H |
| 2013_P1-3-4 | | H | G | H | H | G | H | A | A | H | EQV | C | H | H | H | C | H | H | H | C | H | G | C | T | H | H | T | T | G | EQV |
| 2013_P1-3-5 | | G | G | A | C | G | G | A | A | H | C | EQV | G | T | H | C | H | H | H | C | G | G | EQV | T | G | A | T | T | EQV | EQV |
| 2013_P1-6 | | H | G | EQV | H | G | G | A | A | H | C | H | G | H | T | C | H | H | H | C | H | G | C | T | H | H | T | T | G | H |
| 2013_P1-6-1 | | H | G | H | H | G | G | A | A | G | C | T | G | T | T | C | H | H | H | C | H | G | C | T | H | H | T | T | G | H |
| 2013_P1-7 | | H | G | H | H | G | G | A | A | H | C | H | EQV | H | T | C | H | H | H | C | H | G | C | T | H | H | T | T | G | H |
| 2013_P1-7-1 | | H | G | H | H | G | G | A | A | G | C | T | H | T | H | C | H | H | T | C | G | G | C | T | G | A | T | T | G | H |
| 2013_P1-8 | | H | G | H | H | G | G | A | A | H | C | H | G | H | T | C | H | H | H | C | H | G | C | T | H | H | T | T | G | H |
| 2013_P1-8-1 | | H | G | H | H | G | G | A | A | H | H | EQV | EQV | H | H | C | H | H | H | C | H | G | H | T | H | H | T | T | G | H |
| 2013_P1-8-2 | | H | G | H | H | G | G | A | A | G | C | H | G | H | T | C | H | H | H | C | G | G | EQV | T | G | A | T | T | G | C |
| 2013_P1-8-3 | | H | G | H | H | G | H | A | A | H | H | H | G | H | H | C | G | G | C | C | H | G | C | T | H | EQV | T | T | G | EQV |
| 2013_P1-8-4 | | H | G | H | H | G | G | A | A | H | C | H | G | H | T | C | H | H | H | C | H | G | C | T | H | EQV | T | T | G | H |
| 2013_P1-8-5 | | H | G | H | H | G | H | A | A | H | H | H | G | H | H | C | H | H | H | C | H | G | H | T | H | H | T | T | G | EQV |
| 2013_P1-9 | | H | G | H | H | G | G | A | A | H | C | H | H | H | T | C | H | H | H | C | H | G | C | T | H | EQV | T | T | G | H |
| 2013_P1-9-1 | | A | G | C | A | G | G | A | A | EQV | C | H | H | H | T | C | H | H | H | C | H | G | C | T | H | H | T | T | G | H |
| 2013_P1-9-2 | | H | G | H | H | G | G | A | A | G | C | T | C | T | T | C | H | A | T | C | G | G | C | T | G | A | T | T | G | H |
| 2013_P1-9-3 | | H | G | H | H | G | G | A | A | H | C | H | H | H | T | C | H | H | H | C | H | G | C | T | H | H | T | T | G | H |
| 2013_P1-10 | | H | G | H | H | G | G | A | A | H | C | H | H | H | T | C | H | H | H | C | H | G | C | T | H | H | T | T | G | H |
| 2013_P1-10-1 | | H | G | H | H | G | G | A | A | H | C | H | H | H | T | C | H | H | H | C | H | G | C | T | H | H | T | T | G | H |
| 2013_P1-16 | | H | G | H | H | G | G | A | A | H | C | H | EQV | H | T | C | H | H | H | C | H | G | C | T | H | H | T | T | G | H |
| 2013_P1-16-1 | | H | G | H | H | G | G | A | A | H | C | H | EQV | H | T | C | G | G | C | C | C | G | C | T | A | C | T | T | G | A |
| 2013_P1-16-2 | | H | G | H | H | G | G | A | A | H | C | H | EQV | T | T | C | T | A | T | C | G | G | C | T | G | A | T | T | G | H |
| 2013_P1-17 | | H | G | H | H | G | G | A | A | H | C | H | H | H | T | C | H | H | H | C | H | G | C | T | H | H | T | T | G | H |
| 2013_P1-17-1 | | H | G | H | H | G | G | A | A | H | C | H | H | H | T | C | H | A | EQV | C | G | G | C | T | G | A | T | T | G | C |
| 2013_P1-17-2 | | A | G | C | A | G | G | A | A | G | C | H | H | H | T | C | EQV | H | H | C | C | G | C | T | A | C | T | T | G | H |
| 2013_P1-19 | | EQV | G | H | H | G | G | A | A | H | C | H | EQV | H | T | C | H | H | H | C | H | G | C | T | H | H | T | T | G | H |
| 2013_P1-19-1 | | H | G | H | H | G | G | A | A | H | C | T | H | H | T | C | H | H | H | C | C | G | C | T | A | C | T | T | G | H |
| 2013_P1-19-2 | | G | G | A | C | G | G | A | A | H | C | H | H | H | T | C | H | H | H | C | H | G | C | T | H | H | T | T | G | H |
| 2013_P1-19-3 | | A | G | C | A | G | G | A | A | H | C | EQV | H | H | T | C | T | H | H | C | H | G | C | T | H | H | T | T | G | H |
| 2013_P1-19-4 | | H | G | H | H | G | G | A | A | G | C | T | C | T | T | C | H | H | H | C | H | G | C | T | H | H | T | T | G | EQV |
| 2013_P1-19-5 | | G | G | A | C | G | G | A | A | H | C | H | H | H | T | C | H | A | T | C | G | G | C | T | G | H | T | T | G | H |
| 2013_P1-20 | | H | G | H | H | G | G | A | A | H | C | EQV | G | H | T | C | H | H | H | C | H | G | C | T | H | H | T | T | G | H |
| 2013_P1-20-1 | | H | G | H | H | G | G | A | A | H | C | H | G | H | T | C | H | H | T | C | G | G | C | T | G | A | T | T | G | H |
| 2013_P1-20-2 | | H | G | H | H | G | G | A | A | H | C | H | G | H | T | C | H | G | C | C | C | G | C | T | A | C | T | T | G | A |
| 2013_P1-21 | | H | G | H | H | G | G | A | A | H | C | H | H | H | T | C | H | H | H | C | H | G | C | T | H | H | T | T | G | H |
| 2013_P1-21-1 | | A | G | C | A | G | G | A | A | G | C | H | H | H | T | C | H | H | H | C | C | G | C | T | A | C | T | T | G | A |
| 2013_P1-21-2 | | H | G | H | H | G | G | A | A | G | C | T | C | T | T | C | H | G | C | C | C | G | C | T | A | H | T | T | G | H |
| 2013_P1-21-3 | | H | G | H | H | G | G | A | A | H | C | H | H | H | T | C | G | G | C | C | C | G | C | T | H | H | T | T | G | H |
| 2013_P1-21-4 | | A | G | C | A | G | G | A | A | G | C | H | H | H | T | C | G | G | C | C | C | G | C | T | A | C | T | T | G | H |
| 2013_P1-22 | | H | G | H | H | G | G | A | A | H | C | H | EQV | H | T | C | H | H | H | C | H | G | C | T | H | H | T | T | G | H |
| 2013_P1-22-1 | | H | G | EQV | H | G | G | A | A | H | C | H | H | H | H | C | H | H | H | C | H | G | H | T | H | H | T | T | G | C |
| 2013_P1-22-2 | | H | G | H | H | G | G | A | A | H | C | H | EQV | H | T | C | T | A | T | C | G | G | C | T | G | A | T | T | G | C |
| 2013_P1-25 | | H | G | H | H | G | G | A | A | H | C | H | G | H | T | C | H | H | H | C | H | G | C | T | H | H | T | T | G | H |
| 2013_P1-25-1 | | H | G | H | H | G | G | A | A | H | C | H | EQV | H | T | C | H | H | H | C | H | G | C | T | H | H | T | T | G | H |
| 2013_P1-25-2 | | H | G | H | H | G | G | A | A | A | C | H | G | H | T | C | H | G | C | C | C | G | C | T | A | C | T | T | G | H |
| 2013_P1-25-3 | | H | G | H | H | G | G | A | A | H | C | T | G | T | T | C | T | H | H | C | H | G | C | T | H | H | T | T | G | H |
| 2013_P1-25-4 | | G | G | A | C | G | G | A | A | A | C | C | G | H | T | C | H | H | H | C | H | G | C | T | H | H | T | T | G | C |
| 2013_P1-25-5 | | H | G | H | H | G | G | A | A | H | C | H | G | H | T | C | H | H | H | C | G | G | C | T | H | H | T | T | G | H |
| 2013_P1-25-6 | | A | G | C | A | G | G | A | A | H | C | H | G | H | T | C | H | H | T | C | G | G | C | T | G | A | T | T | G | C |
| 2013_P1-26 | | H | G | H | H | G | G | A | A | H | C | H | H | H | T | C | H | H | H | C | H | G | C | T | H | H | T | T | G | H |
| 2013_P1-26-1 | | H | G | H | H | G | G | A | A | H | C | H | H | H | T | C | T | A | T | C | G | G | C | T | G | A | T | T | G | C |
| 2013_P1-27 | | H | G | H | H | G | G | A | A | H | C | H | EQV | H | T | C | H | H | H | C | H | G | C | T | H | H | T | T | G | H |
| 2013_P1-27-1 | | G | G | A | C | G | G | A | A | A | C | C | G | H | T | C | H | H | T | C | G | G | C | T | G | A | T | T | G | H |
| 2013_P1-28 | | H | G | H | H | G | G | A | A | H | C | H | H | H | T | C | H | H | H | C | H | G | C | T | H | H | T | T | G | H |
| 2013_P1-28-1 | | H | G | H | H | G | G | A | A | H | C | C | EQV | C | T | C | G | H | H | C | C | G | C | T | A | C | T | T | G | H |
| 2013_P1-31 | | H | G | H | H | G | G | A | A | H | C | H | G | H | T | C | H | H | H | C | H | G | C | T | H | H | T | T | G | H |
| 2013_P1-31-1 | | H | G | H | H | G | G | A | A | H | C | H | G | H | T | C | H | G | C | C | C | G | C | T | A | C | T | T | G | A |
| 2013_P1-31-2 | | H | G | H | H | G | G | A | A | H | C | H | G | H | T | C | H | H | C | C | C | G | C | T | A | C | T | T | G | A |
| 2013_O4 | | G | G | A | C | G | G | A | A | A | C | C | G | C | T | C | G | G | C | C | C | G | C | T | A | C | T | T | G | A |
| 2013_O1 | | A | G | C | A | G | G | A | A | G | C | T | C | T | T | C | T | A | T | C | G | G | C | T | G | A | T | T | G | C |
| 2013_O1_O4 | | H | G | H | H | G | G | A | A | H | C | H | H | H | T | C | H | H | H | C | H | G | C | T | H | H | T | T | G | H |

plant_id – unique identifier of the plant
mom_gen – genetic backgroiund of mother in the hybrid
dad_gen – genetic background of father in the hybrid
family_ – identify mother and all the progeny plants from the same mother
mother_progeny – identify each plant as mother of the family or progeny
Chr – number of the chromosome, where marker is located
Distance – position of the marker on the corresponding chromosome in cM
H – heterozygous marker
A, T, G and C – marker homozygous for the corresponding nucleotide
EQV, LS, -9 , etc. – missing values

### Table S1. Results of the SNP array analasis (continue)

| | | | | | Chr | 2 | 2 | 2 | 2 | 2 | 2 | 2 | 2 | 2 | 2 | 2 | 2 | 2 | 3 | 3 | 3 | 3 | 3 | 3 | 3 | 3 | 3 | 3 | 3 | 3 | 3 | 3 | 3 | 3 | 3 | 3 | 3 | 3 | 3 | 3 | 3 | 3 | 3 | 3 | 3 | 3 | 3 | 3 | 3 | 3 | 3 | 3 | 3 | 3 | 3 | 3 | 3 | 4 | 4 | 4 | 4 | 4 | 4 | 4 | 4 | 4 | 4 | 4 | 4 | 4 | 4 | 4 | 4 | 4 | 4 | 4 | 4 | 4 | 4 | 4 | 4 | 4 |
|---|---|---|---|---|---|---|---|---|---|---|---|---|---|---|---|---|---|---|---|---|---|---|---|---|---|---|---|---|---|---|---|---|---|---|---|---|---|---|---|---|---|---|---|---|---|---|---|---|---|---|---|---|---|---|---|---|---|---|---|---|---|---|---|---|---|---|---|---|---|---|---|---|---|---|---|---|---|---|---|---|---|---|
| | | | | | Distance, cM | 196 | 201 | 208 | 216 | 225 | 230 | 235 | 243 | 253 | 253 | 253 | 253 | 254 | 4.4 | 5.9 | 5.9 | 6.7 | 12 | 20 | 28 | 38 | 49 | 54 | 60 | 66 | 77 | 79 | 79 | 80 | 80 | 80 | 87 | 96 | 106 | 115 | 126 | 135 | 145 | 153 | 163 | 173 | 180 | 189 | 199 | 208 | 219 | 228 | 245 | 245 | 245 | 246 | 246 | 4.3 | 5.3 | 7.1 | 7.4 | 11 | 19 | 29 | 37 | 48 | 57 | 67 | 75 | 85 | 94 | 100 | 108 | 109 | 110 | 111 | 111 | 117 | 129 | 134 | 143 | 153 |
| plant_id | mom_gen | dad_gen | mother_progeny | family_ | | | | | | | | | | | | | | | | | | | | | | | | | | | | | | | | | | | | | | | | | | | | | | | | | | | | | | | | | | | | | | | | | | | | | | | | | | | | | | |
| 2013_P1-1 | PHI1 | PHI2 | mother | 2013_P1-1 | | G | H | G | T | A | A | C | C | A | G | G | A | EQV | H | C | C | H | T | T | G | H | H | H | A | A | T | A | C | H | G | H | C | A | H | H | H | EQV | G | C | H | H | H | H | G | H | H | C | A | A | G | G | G | G | H | C | G | H | A | T | C | T | H | T | H | T | G | H | H | G | A | T | H | C | H | T | H | H |
| 2013_P1-1-1 | PHI1 | PHI2 | progeny | 2013_P1-1 | | G | H | G | T | A | A | C | C | A | G | G | A | EQV | A | C | C | A | T | T | G | H | H | H | A | A | T | A | C | H | G | H | C | A | H | H | H | EQV | EQV | C | H | H | H | H | G | H | H | C | A | A | G | G | G | G | H | C | G | H | A | T | C | T | H | T | H | T | G | H | H | G | A | T | H | C | H | T | H | H |
| 2013_P1-3 | PHI1 | PHI2 | mother | 2013_P1-1 | | G | H | G | T | A | A | C | C | A | G | G | A | EQV | H | C | C | H | T | T | G | H | G | H | A | A | T | A | C | H | G | H | C | A | G | T | A | C | G | C | H | H | H | H | G | H | H | C | A | A | G | G | G | G | H | C | G | H | A | T | C | T | H | T | H | T | G | H | H | G | A | T | H | C | H | T | H | H |
| 2013_P1-3-1 | PHI1 | PHI2 | progeny | 2013_P1-3 | | H | H | G | T | A | A | C | C | A | G | G | A | EQV | A | H | C | H | T | T | G | H | G | G | A | A | T | H | C | C | G | H | H | A | H | T | H | C | G | C | A | EQV | H | EQV | H | C | C | H | H | H | H | G | G | G | H | C | H | H | A | H | C | T | H | T | H | H | G | H | C | G | A | T | T | C | H | H | A | C |
| 2013_P1-3-2 | PHI1 | PHI2 | progeny | 2013_P1-3 | | H | H | H | T | A | A | C | C | A | G | G | A | G | H | H | C | EQV | T | T | G | G | G | H | A | A | T | H | C | H | G | C | H | A | H | T | H | C | G | C | A | EQV | H | H | G | H | H | H | A | A | H | G | G | G | H | C | H | H | A | H | C | T | H | T | H | H | G | EQV | H | G | A | H | H | C | A | H | H | H |
| 2013_P1-3-3 | PHI1 | PHI2 | progeny | 2013_P1-3 | | H | EQV | G | T | A | A | C | C | A | G | G | A | EQV | EQV | H | C | EQV | T | T | G | H | G | G | A | A | T | H | C | C | EQV | H | H | A | H | T | EQV | C | G | C | A | EQV | H | H | G | H | H | H | A | A | G | G | G | G | A | C | EQV | H | A | H | C | T | H | T | H | H | G | H | H | G | A | T | H | C | EQV | H | H | H |
| 2013_P1-3-4 | PHI1 | PHI2 | progeny | 2013_P1-3 | | H | H | G | T | A | A | C | C | A | G | G | A | EQV | H | H | C | EQV | T | T | G | H | G | EQV | A | A | T | H | C | H | G | H | H | A | H | T | A | C | G | C | A | T | H | H | EQV | H | H | H | H | H | H | G | G | G | A | C | H | H | A | H | C | T | H | T | H | H | G | H | H | G | A | H | H | C | H | H | H | H |
| 2013_P1-3-5 | PHI1 | PHI2 | progeny | 2013_P1-3 | | H | EQV | H | T | A | A | C | EQV | A | G | G | A | EQV | EQV | H | C | H | T | T | G | G | G | H | A | A | T | H | C | H | EQV | C | H | EQV | H | T | A | C | G | C | H | H | H | H | H | EQV | H | H | H | H | H | G | G | G | EQV | C | G | H | A | H | EQV | T | H | T | T | H | G | A | EQV | G | A | T | T | C | EQV | H | H | H |
| 2013_P1-6 | PHI1 | PHI2 | mother | 2013_P1-6 | | G | H | G | T | A | A | C | C | A | G | G | A | EQV | H | C | C | H | T | T | H | G | G | H | A | A | T | A | C | H | EQV | H | C | A | H | H | H | H | G | C | H | H | H | H | G | H | H | C | A | A | G | G | G | G | H | C | H | C | A | T | C | T | H | T | H | T | G | H | H | G | A | T | H | C | H | T | H | H |
| 2013_P1-6-1 | PHI1 | PHI2 | progeny | 2013_P1-6 | | G | H | G | T | A | A | C | C | A | G | G | A | EQV | A | C | C | A | T | T | A | G | G | H | A | A | T | A | C | H | G | H | C | A | H | H | H | H | G | C | H | H | H | H | G | H | H | C | A | A | G | G | G | G | T | C | A | C | A | T | C | T | H | T | H | T | G | H | H | G | A | T | H | LS | H | T | H | H |
| 2013_P1-7 | PHI1 | PHI2 | mother | 2013_P1-7 | | G | H | G | T | A | A | C | C | A | G | G | A | EQV | H | C | C | H | T | T | H | G | G | H | A | A | T | A | C | H | EQV | H | C | A | H | H | H | H | G | C | H | H | H | H | G | H | H | C | A | A | G | G | G | G | H | C | G | H | A | T | C | T | H | T | H | T | G | H | H | G | A | T | H | C | H | T | H | H |
| 2013_P1-7-1 | PHI1 | PHI2 | progeny | 2013_P1-7 | | G | A | G | T | A | A | C | C | A | G | G | A | EQV | A | C | C | A | T | T | H | G | G | H | A | A | T | A | C | G Text | EQV | C | C | A | G | T | A | C | G | C | G | C | G | H | G | H | H | C | A | A | G | G | G | G | H | C | G | H | A | T | C | T | H | T | H | T | G | G | T | G | A | T | G | C | A | T | G | A |
| 2013_P1-8 | PHI1 | PHI2 | mother | 2013_P1-8 | | G | H | G | T | A | A | C | C | A | G | G | A | EQV | H | C | C | H | T | T | H | G | G | H | A | A | T | A | C | H | G | H | C | A | H | T | A | C | G | C | H | H | H | H | G | H | T | C | A | A | G | G | G | G | H | C | H | C | A | EQV | C | T | H | T | H | T | G | H | H | G | A | T | H | C | H | T | H | H |
| 2013_P1-8-1 | PHI1 | PHI2 | progeny | 2013_P1-8 | | H | H | G | T | A | A | C | C | A | G | G | A | EQV | A | H | C | H | T | T | H | G | G | H | A | A | T | H | C | C | EQV | H | H | A | A | T | H | C | G | C | H | H | H | H | H | H | H | H | A | A | G | G | G | G | H | C | H | H | A | H | C | T | H | T | H | H | G | H | H | G | A | H | H | C | EQV | H | H | H |
| 2013_P1-8-2 | PHI1 | PHI2 | progeny | 2013_P1-8 | | G | A | G | T | A | A | C | C | A | G | G | A | T | H | C | C | H | T | T | H | G | G | H | A | A | T | A | C | H | EQV | H | C | A | H | T | A | C | G | C | H | H | H | H | G | H | T | C | A | A | G | G | G | G | H | C | H | C | A | T | C | T | A | T | C | T | G | G | H | G | A | T | H | C | H | T | H | H |
| 2013_P1-8-3 | PHI1 | PHI2 | progeny | 2013_P1-8 | | H | G | H | T | A | A | C | C | A | G | G | A | EQV | H | H | C | H | T | T | H | G | G | EQV | A | A | T | H | C | H | EQV | H | H | A | H | T | A | C | G | C | A | T | H | H | G | H | H | H | H | H | H | G | G | G | H | C | H | H | A | H | C | T | A | T | H | H | G | H | C | G | A | H | T | C | H | H | A | C |
| 2013_P1-8-4 | PHI1 | PHI2 | progeny | 2013_P1-8 | | G | H | G | T | A | A | C | C | A | G | G | A | EQV | H | C | C | H | T | T | H | G | G | H | A | A | T | A | C | H | G | H | C | A | H | T | A | C | G | C | H | H | H | H | G | H | T | C | A | A | G | G | G | G | H | C | H | C | A | T | C | T | H | T | H | T | G | H | H | G | A | T | H | C | H | T | H | H |
| 2013_P1-8-5 | PHI1 | PHI2 | progeny | 2013_P1-8 | | H | H | H | T | A | A | C | C | A | G | G | A | EQV | H | H | C | EQV | T | T | EQV | G | G | H | A | A | T | H | C | H | EQV | H | H | EQV | H | T | A | C | G | C | H | EQV | H | T | G | H | T | H | A | A | H | G | G | G | H | C | H | H | A | H | C | T | H | T | H | H | G | H | H | G | A | T | H | C | EQV | H | H | H |
| 2013_P1-9 | PHI1 | PHI2 | mother | 2013_P1-9 | | G | H | G | T | A | A | C | C | A | G | G | A | EQV | H | C | C | H | T | T | G | H | H | H | A | A | T | A | C | H | G | H | C | A | H | H | H | H | G | C | H | H | H | H | G | H | H | C | A | A | G | G | G | G | H | C | H | C | A | T | C | T | H | T | H | T | G | H | H | G | A | T | H | C | H | T | H | H |
| 2013_P1-9-1 | PHI1 | PHI2 | progeny | 2013_P1-9 | | G | H | G | T | A | A | C | C | A | G | G | A | EQV | H | C | C | H | T | T | G | H | H | H | A | A | T | A | C | C | G | G | C | EQV | A | C | G | A | G | C | H | H | H | H | G | EQV | H | C | A | A | G | G | G | G | EQV | C | A | C | A | T | C | T | H | T | H | T | G | H | H | G | A | T | EQV | C | H | T | H | H |
| 2013_P1-9-2 | PHI1 | PHI2 | progeny | 2013_P1-9 | | G | H | G | T | A | A | C | C | A | G | G | A | EQV | H | C | C | H | T | T | G | H | H | H | A | A | T | A | C | G | G | C | C | A | G | H | H | H | G | C | H | C | G | T | G | T | T | C | A | A | G | G | G | G | H | C | H | C | A | T | C | T | H | T | H | T | G | G | T | G | A | T | G | C | A | T | G | A |
| 2013_P1-9-3 | PHI1 | PHI2 | progeny | 2013_P1-9 | | G | H | G | T | A | A | C | C | A | G | G | A | EQV | H | C | C | H | T | T | G | H | H | H | A | A | T | A | C | H | G | H | C | A | H | H | H | EQV | G | C | H | H | H | H | G | H | H | C | A | A | G | G | G | G | H | C | H | C | A | T | EQV | T | H | T | H | T | G | H | C | G | A | T | T | C | G | T | A | C |
| 2013_P1-10 | PHI1 | PHI2 | mother | 2013_P1-10 | | G | H | G | T | A | A | C | C | A | G | G | A | EQV | H | C | C | H | T | T | H | G | G | H | A | A | T | A | C | H | G | H | C | A | H | H | H | H | G | C | H | H | H | H | G | H | H | C | A | A | G | G | G | G | H | C | H | H | A | T | C | T | H | T | H | T | G | H | H | G | A | T | H | C | H | T | H | H |
| 2013_P1-10-1 | PHI1 | PHI2 | progeny | 2013_P1-10 | | G | H | G | T | A | A | C | C | A | G | G | A | EQV | H | C | C | H | T | T | H | G | G | H | A | A | T | A | C | H | G | H | C | A | H | H | H | C | G | C | G | C | H | H | G | H | H | C | A | A | G | G | G | G | H | C | H | H | A | T | C | T | H | T | H | T | G | H | H | G | A | T | H | C | G | T | A | C |
| 2013_P1-16 | PHI1 | PHI2 | mother | 2013_P1-16 | | G | H | G | T | A | A | C | C | A | G | G | A | EQV | H | C | C | H | T | T | H | G | G | H | A | A | T | A | C | H | EQV | H | C | A | H | H | H | H | G | C | H | H | H | H | G | H | H | C | A | A | G | G | G | G | H | C | H | C | A | T | C | T | H | T | H | T | G | H | H | G | A | T | H | C | H | T | H | H |
| 2013_P1-16-1 | PHI1 | PHI2 | progeny | 2013_P1-16 | | G | H | G | T | A | A | C | C | A | G | G | A | G | H | C | C | H | T | T | H | G | G | H | A | A | T | A | C | G | EQV | C | C | A | G | T | A | C | G | C | G | C | G | T | G | T | H | C | A | A | G | G | G | G | A | C | G | C | A | T | C | T | H | T | H | T | G | H | T | G | A | T | G | C | A | T | H | H |
| 2013_P1-16-2 | PHI1 | PHI2 | progeny | 2013_P1-16 | | G | H | G | T | A | A | C | C | EQV | G | G | A | EQV | H | C | C | EQV | T | T | H | G | G | H | A | A | T | A | C | H | G | H | C | A | H | H | H | EQV | G | C | H | H | H | H | G | H | H | C | A | A | G | G | G | G | H | C | H | C | A | T | C | T | A | T | C | T | G | G | T | G | A | T | G | C | H | T | H | H |
| 2013_P1-17 | PHI1 | PHI2 | mother | 2013_P1-17 | | G | H | G | T | A | A | C | C | A | G | G | A | EQV | H | C | C | H | T | T | G | H | H | H | A | A | T | A | C | G | G | C | C | A | G | H | H | H | G | C | H | H | H | H | G | H | H | C | A | A | G | G | G | G | H | C | H | C | A | T | C | T | H | T | H | T | G | H | H | G | A | T | H | C | H | T | H | H |
| 2013_P1-17-1 | PHI1 | PHI2 | progeny | 2013_P1-17 | | H | H | G | T | A | A | C | C | A | G | G | A | EQV | H | C | C | H | T | T | G | C | C | G | A | A | T | A | C | G | EQV | C | C | EQV | G | H | H | H | G | C | H | H | H | H | G | H | H | C | A | A | G | G | G | G | H | C | H | C | A | T | C | T | H | T | T | T | G | EQV | C | G | A | H | H | C | H | H | H | H |
| 2013_P1-17-2 | PHI1 | PHI2 | progeny | 2013_P1-17 | | G | H | G | T | A | A | C | C | A | G | G | A | EQV | G | C | C | T | T | T | G | G | G | A | A | A | T | A | C | G | EQV | C | C | A | G | H | H | H | G | C | G | C | G | T | G | T | H | C | A | A | G | G | G | G | H | C | H | C | A | T | C | T | H | T | C | T | G | G | T | G | A | T | G | C | A | T | G | A |
| 2013_P1-19 | PHI1 | PHI2 | mother | 2013_P1-19 | | G | H | G | T | A | A | C | C | A | G | G | A | EQV | H | C | C | H | T | T | H | H | H | H | A | A | T | A | C | G | EQV | C | C | A | H | H | H | H | G | C | H | H | H | H | G | H | H | C | A | A | G | G | G | G | H | C | H | C | A | T | C | T | H | T | H | T | G | H | H | G | A | T | H | C | H | T | H | H |
| 2013_P1-19-1 | PHI1 | PHI2 | progeny | 2013_P1-19 | | G | H | G | T | A | A | C | C | A | G | G | A | G | H | C | C | H | T | T | G | G | G | A | A | A | T | A | C | G | G | C | C | A | A | C | G | A | G | C | H | H | H | H | G | H | C | C | A | A | G | G | G | G | H | C | H | C | A | T | C | T | A | T | C | T | G | H | C | G | A | T | T | C | G | T | A | C |
| 2013_P1-19-2 | PHI1 | PHI2 | progeny | 2013_P1-19 | | G | H | G | T | A | A | C | C | A | G | G | A | EQV | H | C | C | H | T | T | G | G | G | A | A | A | T | A | C | G | G | C | C | A | H | H | H | H | G | C | H | H | H | H | G | H | H | C | A | A | G | G | G | G | H | C | H | C | A | T | C | T | T | T | H | T | G | H | H | G | A | T | H | C | H | T | H | H |
| 2013_P1-19-3 | PHI1 | PHI2 | progeny | 2013_P1-19 | | G | H | G | T | A | A | C | C | A | G | G | A | EQV | H | C | C | H | T | T | H | H | G | A | A | A | T | A | C | G | G | C | C | A | G | H | H | H | G | C | H | H | G | T | G | H | H | C | A | A | G | G | G | G | H | C | H | C | A | T | C | T | H | T | H | T | G | H | H | G | A | T | H | C | H | T | H | H |
| 2013_P1-19-4 | PHI1 | PHI2 | progeny | 2013_P1-19 | | G | H | G | T | A | A | C | C | A | G | G | A | T | H | C | C | H | T | T | H | H | H | H | A | A | T | A | C | G | EQV | C | C | A | H | H | H | H | G | C | H | H | T | C | G | H | H | C | A | A | G | G | G | G | H | C | EQV | C | A | T | C | T | H | T | H | T | G | H | H | G | A | T | EQV | C | H | T | H | H |
| 2013_P1-19-5 | PHI1 | PHI2 | progeny | 2013_P1-19 | | G | A | G | T | A | A | C | C | A | G | G | A | T | A | C | C | A | T | T | A | C | H | H | A | A | T | A | C | G | EQV | C | C | A | G | T | A | C | G | C | H | H | H | H | G | H | H | C | A | A | G | G | G | G | H | C | H | C | A | T | C | T | H | T | H | T | G | A | C | G | A | T | T | C | H | T | H | A |
| 2013_P1-20 | PHI1 | PHI2 | mother | 2013_P1-20 | | G | H | G | T | A | A | C | C | A | G | G | A | EQV | H | C | C | H | T | T | H | G | G | H | A | A | T | A | C | H | EQV | H | C | A | H | H | H | H | G | C | H | H | H | H | G | H | H | C | A | A | G | G | G | G | H | C | H | C | A | T | C | T | H | T | H | T | G | H | H | G | A | T | H | C | H | T | H | H |
| 2013_P1-20-1 | PHI1 | PHI2 | progeny | 2013_P1-20 | | G | H | G | T | A | A | C | C | A | G | G | A | T | A | C | C | A | T | T | A | G | G | G | A | A | T | A | C | C | G | G | C | A | H | H | H | H | G | C | H | T | T | C | G | C | C | C | A | A | G | G | G | G | H | C | H | C | A | T | C | T | H | T | T | T | G | A | C | G | A | T | T | C | G | T | A | C |
| 2013_P1-20-2 | PHI1 | PHI2 | progeny | 2013_P1-20 | | G | G | G | T | A | A | C | C | A | G | G | A | G | H | C | C | H | T | T | H | G | G | G | A | A | T | A | C | C | G | G | C | A | A | C | G | A | G | C | H | H | H | H | G | H | H | C | A | A | G | G | G | G | H | C | H | C | A | T | C | T | T | T | T | T | G | A | C | G | A | T | T | C | G | T | A | C |
| 2013_P1-21 | PHI1 | PHI2 | mother | 2013_P1-21 | | G | H | G | T | A | A | C | C | A | G | G | A | EQV | H | C | C | H | T | T | G | H | H | H | A | A | T | A | C | G | EQV | C | C | A | G | T | A | C | G | C | H | H | H | H | G | H | H | C | A | A | G | G | G | G | H | C | H | C | A | T | C | T | H | T | H | T | G | H | H | G | A | T | H | C | H | T | H | H |
| 2013_P1-21-1 | PHI1 | PHI2 | progeny | 2013_P1-21 | | G | G | G | T | A | A | C | C | A | G | G | A | EQV | H | C | C | H | T | T | G | H | H | H | A | A | T | A | C | G | EQV | C | C | A | G | T | A | C | G | C | H | H | H | H | G | H | H | C | A | A | G | G | G | G | EQV | C | H | C | A | T | C | T | H | T | C | T | G | G | T | G | A | T | G | C | A | T | G | A |
| 2013_P1-21-2 | PHI1 | PHI2 | progeny | 2013_P1-21 | | G | H | G | T | A | A | C | C | A | G | G | A | EQV | H | C | C | H | T | T | G | H | H | G | A | A | T | A | C | G | EQV | C | C | A | G | T | A | C | G | C | H | H | H | H | G | C | C | C | A | A | G | G | G | G | H | C | H | C | A | T | C | T | T | T | T | T | G | A | C | G | A | T | H | C | H | T | H | H |
| 2013_P1-21-3 | PHI1 | PHI2 | progeny | 2013_P1-21 | | G | H | G | T | A | A | C | C | A | G | G | A | T | H | C | C | H | T | T | G | C | C | G | A | A | T | A | C | G | EQV | C | C | A | G | T | A | C | G | C | A | T | T | H | G | H | H | C | A | A | G | G | G | G | H | C | H | C | A | T | C | T | H | T | H | T | G | G | T | G | A | T | G | C | A | T | H | H |
| 2013_P1-21-4 | PHI1 | PHI2 | progeny | 2013_P1-21 | | G | H | G | T | A | A | C | C | A | G | G | A | G | H | C | C | H | T | T | G | G | G | A | A | A | T | A | C | G | G | C | C | A | G | T | A | C | G | C | G | H | H | T | G | T | T | C | A | A | G | G | G | G | H | C | H | C | A | T | C | T | H | T | T | T | G | A | C | G | A | T | T | C | G | T | H | H |
| 2013_P1-22 | PHI1 | PHI2 | mother | 2013_P1-22 | | G | H | G | T | A | A | C | C | A | G | G | A | EQV | H | C | C | H | T | T | G | H | H | H | A | A | T | A | C | G | EQV | C | C | A | G | T | A | H | G | C | H | H | H | H | G | H | T | C | A | A | G | G | G | G | H | C | H | C | A | T | C | T | H | T | H | T | G | H | H | G | A | T | H | C | H | T | H | H |
| 2013_P1-22-1 | PHI1 | PHI2 | progeny | 2013_P1-22 | | H | H | H | EQV | A | A | C | C | A | G | G | A | EQV | H | H | C | H | T | T | G | H | H | H | A | A | T | H | C | H | EQV | C | H | A | H | T | H | C | G | C | H | EQV | H | EQV | G | EQV | T | H | A | A | G | G | G | G | H | C | H | H | A | H | C | T | H | T | H | H | G | EQV | H | G | A | H | H | C | EQV | H | H | H |
| 2013_P1-22-2 | PHI1 | PHI2 | progeny | 2013_P1-22 | | G | A | G | T | A | A | C | C | A | G | G | A | T | G | C | C | T | T | T | G | G | G | H | A | A | T | A | C | G | EQV | C | C | A | G | T | A | A | G | C | A | T | H | H | G | H | T | C | A | A | G | G | G | G | H | C | H | C | A | T | C | T | H | T | H | T | G | G | T | G | A | T | G | C | A | T | G | A |
| 2013_P1-25 | PHI1 | PHI2 | mother | 2013_P1-25 | | G | H | G | T | A | A | C | C | A | G | G | A | EQV | H | C | C | H | T | T | G | H | H | H | A | A | T | A | C | H | EQV | H | C | A | H | H | H | EQV | G | C | H | H | H | H | G | H | H | C | A | A | G | G | G | G | H | C | H | C | A | T | C | LS | H | T | H | T | G | H | H | G | A | T | H | C | H | T | H | H |
| 2013_P1-25-1 | PHI1 | PHI2 | progeny | 2013_P1-25 | | G | H | G | T | A | A | C | C | A | G | G | A | EQV | H | C | C | H | T | T | G | H | H | H | A | A | T | A | C | H | G | H | C | A | H | H | H | H | G | C | H | C | G | T | G | T | T | C | A | A | G | G | G | G | H | C | H | C | A | T | C | T | H | T | H | T | G | H | H | G | A | T | H | C | H | T | H | H |
| 2013_P1-25-2 | PHI1 | PHI2 | progeny | 2013_P1-25 | | G | H | G | T | A | A | C | C | A | G | G | A | EQV | H | C | C | H | T | T | G | G | G | A | A | A | T | A | C | G | EQV | C | C | A | H | H | H | EQV | G | C | H | T | T | C | EQV | H | H | C | A | A | G | G | G | G | H | C | EQV | C | A | T | C | T | H | T | T | T | G | A | C | G | A | T | H | EQV | G | T | A | C |
| 2013_P1-25-3 | PHI1 | PHI2 | progeny | 2013_P1-25 | | G | H | G | T | A | A | C | C | A | G | G | A | EQV | H | C | C | H | T | T | G | H | H | H | A | A | T | A | C | H | G | H | C | A | H | H | H | H | G | C | G | H | H | H | G | H | H | C | A | A | G | G | G | G | A | C | G | C | A | T | C | T | T | T | T | T | G | H | H | G | A | T | H | C | H | T | H | H |
| 2013_P1-25-4 | PHI1 | PHI2 | progeny | 2013_P1-25 | | G | A | G | T | A | A | C | C | A | G | G | A | T | H | C | C | H | T | T | G | H | H | H | A | A | T | A | C | H | EQV | H | C | A | H | H | H | H | G | C | H | H | H | H | G | H | H | C | A | A | G | G | G | G | A | C | G | C | A | T | C | T | H | T | T | T | G | A | H | G | A | T | H | C | H | T | H | H |
| 2013_P1-25-5 | PHI1 | PHI2 | progeny | 2013_P1-25 | | G | H | G | T | A | A | C | C | A | G | G | A | EQV | A | C | C | A | T | T | G | H | C | G | A | A | T | A | C | C | EQV | G | C | A | H | H | H | H | G | C | H | T | H | H | G | H | H | C | A | A | G | G | G | G | H | C | H | C | A | T | C | T | A | T | C | T | G | G | T | G | A | T | G | C | H | T | H | H |
| 2013_P1-25-6 | PHI1 | PHI2 | progeny | 2013_P1-25 | | G | A | G | T | A | A | C | C | A | G | G | A | EQV | A | C | C | A | T | T | G | C | C | G | A | A | T | A | C | H | G | H | C | A | H | H | H | H | G | C | H | H | H | H | G | T | T | C | A | A | G | G | G | G | H | C | H | C | A | T | C | T | H | T | H | T | G | H | H | G | A | T | H | C | H | T | H | H |
| 2013_P1-26 | PHI1 | PHI2 | mother | 2013_P1-26 | | G | H | G | T | A | A | C | C | A | G | G | A | EQV | H | C | C | H | T | T | G | H | H | H | A | A | T | A | C | G | G | C | C | A | G | T | A | C | G | C | H | H | H | H | G | H | H | C | A | A | G | G | G | G | H | C | H | C | A | T | C | T | H | T | H | T | G | H | H | G | A | T | H | C | H | T | H | H |
| 2013_P1-26-1 | PHI1 | PHI2 | progeny | 2013_P1-26 | | G | A | G | T | A | A | C | C | A | G | G | A | EQV | H | C | C | H | T | T | G | C | C | G | A | A | T | A | C | G | G | C | C | A | G | T | A | C | G | C | H | H | T | C | G | C | C | C | A | A | G | G | G | G | H | C | H | C | A | T | C | T | H | T | H | T | G | A | C | G | A | T | T | C | G | T | A | C |
| 2013_P1-27 | PHI1 | PHI2 | mother | 2013_P1-27 | | G | H | G | T | A | A | C | C | A | G | G | A | EQV | H | C | C | H | T | T | H | G | H | H | A | A | T | A | C | G | EQV | C | C | A | G | T | A | C | G | C | H | H | H | H | G | H | H | C | A | A | G | G | G | G | H | C | H | H | A | T | C | T | H | T | H | T | G | H | H | G | A | T | H | C | H | T | H | H |
| 2013_P1-27-1 | PHI1 | PHI2 | progeny | 2013_P1-27 | | G | H | G | T | A | A | C | C | A | G | G | A | EQV | A | C | C | A | T | T | A | G | C | G | A | A | T | A | C | G | EQV | C | C | A | G | T | A | C | G | C | G | H | H | H | G | H | H | C | A | A | G | G | G | G | H | C | H | H | A | T | C | T | H | T | T | T | G | A | C | G | A | T | T | C | H | T | H | H |
| 2013_P1-28 | PHI1 | PHI2 | mother | 2013_P1-28 | | G | H | G | T | A | A | C | C | A | G | G | A | EQV | H | C | C | H | T | T | G | H | H | H | A | A | T | A | C | G | EQV | C | C | A | G | T | A | C | G | C | H | H | H | H | G | H | T | C | A | A | G | G | G | G | H | C | G | H | A | T | C | T | H | T | H | T | G | H | H | G | A | T | H | C | H | T | H | H |
| 2013_P1-28-1 | PHI1 | PHI2 | progeny | 2013_P1-28 | | G | G | G | T | A | A | C | C | A | G | G | A | G | H | C | C | H | T | T | G | C | C | G | A | A | T | A | C | G | EQV | C | C | A | G | T | A | C | G | C | H | H | H | H | G | H | T | C | A | A | G | G | G | G | H | C | G | H | A | T | C | T | H | T | H | T | G | A | C | G | A | T | T | C | G | T | A | C |
| 2013_P1-31 | PHI1 | PHI2 | mother | 2013_P1-31 | | G | H | G | T | A | A | C | C | A | G | G | A | T | H | C | C | H | T | T | G | H | H | H | A | A | T | A | C | G | EQV | C | C | A | G | T | H | H | G | C | H | H | H | H | G | EQV | H | C | A | A | G | G | G | G | H | C | H | C | A | T | C | T | H | T | H | T | G | H | H | G | A | T | H | C | H | T | H | H |
| 2013_P1-31-1 | PHI1 | PHI2 | progeny | 2013_P1-31 | | G | G | G | T | A | A | C | C | A | G | G | A | EQV | H | C | C | H | T | T | G | H | H | H | A | A | T | A | C | G | G | C | C | A | G | T | H | H | G | C | A | T | H | H | G | H | H | C | A | A | G | G | G | G | H | C | H | C | A | T | C | T | H | T | H | T | G | H | H | G | A | T | H | C | H | T | H | H |
| 2013_P1-31-2 | PHI1 | PHI2 | progeny | 2013_P1-31 | | G | G | G | T | A | A | C | C | A | G | G | A | EQV | A | C | C | H | T | T | G | H | H | H | A | A | T | A | C | G | EQV | C | C | A | G | T | A | C | G | C | G | C | H | H | G | T | T | C | A | A | G | G | G | G | H | C | H | C | A | T | C | T | H | T | C | T | G | G | T | G | A | T | G | C | A | T | G | H |
| 2013_O4 | PHI2 | PHI2 | ancestor | 2013_O4 | | G | G | G | T | A | A | C | C | A | G | G | A | G | G | C | C | T | T | T | G | G | G | A | A | A | T | A | C | G | EQV | C | C | A | G | T | A | C | G | C | G | C | G | T | G | T | T | C | A | A | G | G | G | G | A | C | G | C | A | T | C | T | T | T | T | T | G | A | C | G | A | T | T | C | G | T | A | C |
| 2013_O1 | PHI1 | PHI1 | ancestor | 2013_O1 | | G | A | G | T | A | A | C | C | A | G | G | A | T | A | C | C | A | T | T | A | G | G | G | A | A | T | A | C | C | G | G | C | A | A | C | G | A | G | C | A | T | T | C | G | C | C | C | A | A | G | G | G | G | T | C | A | C | A | T | C | T | A | T | C | T | G | G | T | G | A | T | G | C | A | T | G | A |
| 2013_O1_O4 | PHI1 | PHI2 | ancestor | 2013_O1_O4 | | G | H | G | T | A | A | C | C | A | G | G | A | EQV | H | C | C | H | T | T | H | G | G | H | A | A | T | A | C | H | G | H | C | A | H | H | H | H | G | C | H | H | H | H | G | H | H | C | A | A | G | G | G | G | H | C | H | C | A | T | C | T | H | T | H | T | G | H | H | G | A | T | H | C | H | T | H | H |

plant_id – unique identifier of the plant
mom_gen – genetic backgroiund of mother in the hy
dad_gen – genetic background of father in the hybı
family_ – identify mother and all the progeny plar
mother_progeny – identify each plant as mother of the fam
Chr – number of the chromosome, where mar
Distance – position of the marker on the correspon
H – heterozygous marker
A, T, G and C – marker homozygous for the correspond
EQV, LS, -9 , etc. – missing values

### Table S1. Results of the SNP array analasis (continue)

| | | | | | Chr | 4 | 4 | 4 | 4 | 4 | 4 | 4 | 4 | 4 | 4 | 4 | 4 | 4 | 4 | 4 | 5 | 5 | 5 | 5 | 5 | 5 | 5 | 5 | 5 | 5 | 5 | 5 | 5 | 5 | 5 | 5 | 5 | 5 | 5 | 5 | 5 | 5 | 5 | 5 | 5 | 5 | 5 | 5 | 5 | 5 | 5 | 5 | 5 | 5 | 5 | 5 | 5 | 5 | 5 | 6 | 6 | 6 | 6 | 6 | 6 | 6 | 6 | 6 | 6 | 6 | 6 | 6 | 6 | 6 | 6 | 6 | 6 | 6 | 6 | 6 | 6 | 6 |
|---|---|---|---|---|---|---|---|---|---|---|---|---|---|---|---|---|---|---|---|---|---|---|---|---|---|---|---|---|---|---|---|---|---|---|---|---|---|---|---|---|---|---|---|---|---|---|---|---|---|---|---|---|---|---|---|---|---|---|---|---|---|---|---|---|---|---|---|---|---|---|---|---|---|---|---|---|---|---|---|---|---|---|
| | | | | | Distance, cM | 160 | 170 | 179 | 188 | 193 | 198 | 207 | 216 | 226 | 230 | 240 | 240 | 242 | 246 | 246 | 2.8 | 4.9 | 4.9 | 5.3 | 5.5 | 13 | 24 | 30 | 40 | 50 | 59 | 67 | 77 | 85 | 94 | 97 | 101 | 102 | 102 | 103 | 103 | 111 | 120 | 127 | 132 | 140 | 148 | 157 | 167 | 176 | 185 | 193 | 203 | 209 | 218 | 218 | 219 | 224 | 227 | 0.3 | 1.4 | 4 | 5.4 | 5.4 | 8.9 | 18 | 22 | 23 | 23 | 23 | 24 | 24 | 32 | 43 | 50 | 57 | 66 | 76 | 85 | 90 | 96 | 105 |
| plant_id | mom_gen | dad_gen | mother_progeny | family_ | | | | | | | | | | | | | | | | | | | | | | | | | | | | | | | | | | | | | | | | | | | | | | | | | | | | | | | | | | | | | | | | | | | | | | | | | | | | | | |
| 2013_P1-1 | PHI1 | PHI2 | mother | 2013_P1-1 | | C | G | A | T | C | C | H | H | H | H | C | T | H | T | H | H | H | A | H | G | H | H | T | C | H | T | H | C | G | C | H | EQV | A | A | H | H | A | EQV | H | C | H | C | H | H | H | H | H | G | H | G | H | T | G | T | A | G | H | G | C | C | C | H | C | H | G | H | G | G | C | T | G | H | A | G | H | T | H |
| 2013_P1-1-1 | PHI1 | PHI2 | progeny | 2013_P1-1 | | C | G | A | T | C | C | H | H | H | H | C | T | C | T | G | H | H | A | H | G | H | H | T | C | H | T | H | C | G | C | H | EQV | A | A | H | H | A | H | H | C | H | C | H | H | H | H | H | G | H | G | C | T | G | T | A | G | H | G | C | C | C | H | C | H | G | H | G | G | C | T | G | H | A | G | H | T | H |
| 2013_P1-3 | PHI1 | PHI2 | mother | 2013_P1-1 | | C | G | A | T | C | H | A | A | H | H | H | H | H | H | G | H | H | A | H | G | G | C | H | C | H | T | H | C | G | C | H | EQV | A | A | H | EQV | A | H | H | C | H | C | H | C | H | C | H | G | H | G | G | T | H | H | A | G | H | G | C | C | C | H | C | H | G | H | G | G | C | T | G | H | A | G | H | T | H |
| 2013_P1-3-1 | PHI1 | PHI2 | progeny | 2013_P1-3 | | C | EQV | H | T | C | C | A | A | H | EQV | C | T | H | H | G | H | H | A | H | G | H | C | A | C | T | T | H | C | G | C | H | EQV | A | H | T | G | H | C | H | C | H | C | H | C | H | C | H | H | H | G | G | T | H | H | A | G | H | G | C | C | H | H | C | H | H | H | EQV | G | C | T | G | H | H | G | H | T | EQV |
| 2013_P1-3-2 | PHI1 | PHI2 | progeny | 2013_P1-3 | | C | EQV | H | T | C | H | H | A | H | H | H | T | C | T | G | C | A | A | G | G | H | C | H | C | H | T | EQV | C | G | C | H | C | A | H | T | G | H | C | H | C | H | C | H | C | C | C | H | H | H | G | G | T | H | H | A | G | H | G | C | C | H | G | C | H | H | T | EQV | G | C | T | G | H | H | H | H | T | H |
| 2013_P1-3-3 | PHI1 | PHI2 | progeny | 2013_P1-3 | | C | EQV | H | T | C | H | A | A | H | H | H | EQV | H | H | G | H | H | A | H | G | H | C | H | C | H | T | C | C | G | C | EQV | EQV | A | H | H | EQV | H | H | H | C | EQV | C | H | C | H | C | H | H | H | G | G | EQV | H | H | A | G | H | G | C | C | H | H | C | H | H | H | EQV | G | C | T | G | H | H | H | H | T | EQV |
| 2013_P1-3-4 | PHI1 | PHI2 | progeny | 2013_P1-3 | | C | H | H | T | C | C | H | A | H | H | H | EQV | H | H | G | C | A | A | G | G | H | C | H | C | H | T | C | C | G | C | EQV | EQV | A | H | H | H | H | H | H | C | EQV | C | EQV | C | H | C | H | H | H | G | G | H | H | H | A | G | H | G | C | C | C | H | C | H | H | T | EQV | G | C | T | G | G | H | H | H | T | G |
| 2013_P1-3-5 | PHI1 | PHI2 | progeny | 2013_P1-3 | | C | EQV | H | T | C | C | H | A | H | H | H | EQV | H | H | G | C | A | A | G | G | H | C | H | C | H | T | EQV | H | G | C | H | EQV | A | H | EQV | G | H | C | H | C | H | C | H | C | C | C | G | H | H | G | G | T | H | H | A | G | G | G | C | C | H | H | C | H | H | H | EQV | G | C | T | G | H | H | H | H | T | EQV |
| 2013_P1-6 | PHI1 | PHI2 | mother | 2013_P1-6 | | C | G | A | T | C | H | A | A | H | H | H | H | H | H | G | H | H | A | H | G | H | H | T | C | C | T | H | C | G | C | H | EQV | A | A | H | H | A | H | H | C | H | C | H | H | H | H | H | G | H | G | G | T | H | H | A | G | H | G | C | C | C | H | C | H | G | H | G | G | C | T | G | H | A | G | H | T | H |
| 2013_P1-6-1 | PHI1 | PHI2 | progeny | 2013_P1-6 | | C | G | A | T | C | H | A | A | H | H | H | H | H | H | G | H | H | A | H | G | A | T | T | C | C | T | T | C | G | C | H | EQV | A | A | H | H | A | H | H | C | H | C | H | H | H | H | H | G | H | G | G | T | H | H | A | G | H | G | C | C | C | H | C | H | G | H | G | G | C | T | G | H | A | G | H | T | H |
| 2013_P1-7 | PHI1 | PHI2 | mother | 2013_P1-7 | | C | G | A | T | C | C | H | H | H | A | H | H | H | T | H | H | H | A | H | G | H | H | T | C | C | T | H | C | G | C | H | EQV | A | A | H | H | A | H | H | C | H | C | H | H | H | H | H | G | H | G | G | T | H | H | A | G | H | G | C | C | C | H | C | H | G | H | G | G | C | T | G | H | A | G | H | T | H |
| 2013_P1-7-1 | PHI1 | PHI2 | progeny | 2013_P1-7 | | C | G | A | T | C | C | G | G | H | A | H | H | H | T | H | H | H | A | H | G | H | H | T | C | C | T | H | C | G | C | H | C | A | A | H | EQV | A | H | H | C | H | C | H | H | H | C | A | G | C | G | G | T | G | T | A | G | H | G | C | C | C | G | C | T | G | T | G | G | C | T | G | H | A | G | H | T | T |
| 2013_P1-8 | PHI1 | PHI2 | mother | 2013_P1-8 | | C | G | A | T | C | H | A | A | H | H | C | T | H | T | H | H | H | A | H | G | G | C | H | C | H | T | H | C | G | C | H | EQV | A | A | H | EQV | A | H | H | C | H | C | H | H | H | C | H | G | H | G | H | T | G | T | A | G | H | G | C | C | C | H | C | H | G | H | G | G | C | T | G | H | A | G | H | T | H |
| 2013_P1-8-1 | PHI1 | PHI2 | progeny | 2013_P1-8 | | C | EQV | H | T | C | EQV | H | A | H | H | C | T | H | T | H | H | H | A | H | G | H | C | A | C | H | T | H | H | G | C | H | C | A | H | T | G | H | C | H | C | EQV | C | H | H | H | C | H | H | H | G | H | T | G | T | A | G | H | G | C | C | H | G | C | H | H | H | EQV | G | C | T | G | H | H | H | H | T | H |
| 2013_P1-8-2 | PHI1 | PHI2 | progeny | 2013_P1-8 | | C | G | A | T | C | T | A | A | H | H | C | T | H | T | H | H | H | A | H | G | G | C | A | C | T | T | T | C | G | C | H | C | A | A | H | H | A | H | H | C | H | C | H | H | H | C | G | G | G | G | C | T | G | T | A | G | G | G | C | C | C | A | C | C | G | C | G | G | C | T | G | C | A | G | H | T | H |
| 2013_P1-8-3 | PHI1 | PHI2 | progeny | 2013_P1-8 | | C | EQV | H | T | C | C | H | A | H | EQV | H | T | C | T | G | H | H | A | H | G | H | C | H | C | H | T | H | H | G | C | H | C | A | H | EQV | G | H | C | H | C | H | C | G | C | H | C | H | H | H | G | H | T | G | T | A | G | H | G | C | C | H | H | C | H | H | H | EQV | G | C | T | G | H | H | G | H | T | H |
| 2013_P1-8-4 | PHI1 | PHI2 | progeny | 2013_P1-8 | | C | G | A | T | C | H | A | A | H | H | C | T | H | T | H | H | H | A | H | G | G | C | H | C | H | T | H | C | G | C | H | EQV | A | A | H | H | A | H | H | C | H | C | H | H | H | C | H | G | H | G | H | T | G | T | A | G | H | G | C | C | C | H | C | H | G | H | G | G | C | T | G | H | A | G | H | T | H |
| 2013_P1-8-5 | PHI1 | PHI2 | progeny | 2013_P1-8 | | C | H | H | T | C | EQV | A | A | C | A | C | T | C | T | G | C | A | A | G | G | H | C | EQV | C | H | T | H | C | G | C | H | EQV | A | H | H | H | H | H | C | C | A | C | G | C | H | C | H | H | H | G | G | T | G | T | A | G | EQV | G | C | C | H | H | C | H | H | H | EQV | G | C | T | G | EQV | H | H | H | T | H |
| 2013_P1-9 | PHI1 | PHI2 | mother | 2013_P1-9 | | C | G | A | T | C | H | A | A | H | H | C | T | H | T | H | H | H | A | H | G | H | H | EQV | C | C | T | H | C | G | C | H | EQV | A | H | EQV | H | A | H | H | C | H | C | H | H | H | H | H | G | H | G | G | T | H | H | A | G | H | G | C | C | C | H | C | H | G | H | G | G | C | T | G | H | A | G | H | T | H |
| 2013_P1-9-1 | PHI1 | PHI2 | progeny | 2013_P1-9 | | C | G | A | T | C | T | A | A | H | H | C | T | H | T | H | H | G | A | A | G | A | T | T | C | C | T | H | C | G | C | H | EQV | A | EQV | H | H | A | H | H | C | H | C | G | C | T | C | A | G | C | G | G | T | H | H | A | G | H | G | C | C | C | H | C | T | G | T | G | G | C | T | G | H | A | G | H | T | H |
| 2013_P1-9-2 | PHI1 | PHI2 | progeny | 2013_P1-9 | | C | G | EQV | T | C | H | A | A | H | H | C | T | H | T | H | C | A | A | G | G | G | C | T | C | C | T | H | C | G | C | H | C | A | A | T | G | A | C | A | C | H | C | A | T | C | A | H | G | H | G | G | T | H | H | A | G | H | G | C | C | C | G | C | T | G | T | G | G | C | T | G | G | A | G | H | T | H |
| 2013_P1-9-3 | PHI1 | PHI2 | progeny | 2013_P1-9 | | C | G | A | T | C | H | A | A | H | A | C | T | C | T | G | H | H | A | H | G | H | H | T | C | C | T | H | C | G | C | H | C | A | A | H | H | A | H | H | C | H | C | H | H | H | H | H | G | H | G | G | T | H | H | A | G | H | G | C | C | C | A | C | C | G | C | G | G | C | T | G | H | A | G | G | T | G |
| 2013_P1-10 | PHI1 | PHI2 | mother | 2013_P1-10 | | C | G | A | T | C | H | A | A | H | H | C | T | H | T | H | H | H | A | H | G | H | H | H | C | H | T | H | C | G | C | H | EQV | A | A | H | H | A | H | H | C | H | C | H | C | H | C | H | G | H | G | H | T | G | T | A | G | H | G | C | C | C | H | C | H | G | H | G | G | C | T | G | H | A | G | H | T | H |
| 2013_P1-10-1 | PHI1 | PHI2 | progeny | 2013_P1-10 | | C | G | A | T | C | C | A | A | H | H | C | T | H | T | H | H | H | A | H | G | H | H | H | C | H | T | T | C | G | C | H | EQV | A | A | H | EQV | A | H | H | C | H | C | H | C | H | C | A | G | C | G | G | T | G | T | A | G | H | G | C | C | C | H | C | T | G | T | G | G | C | T | G | G | A | G | H | T | T |
| 2013_P1-16 | PHI1 | PHI2 | mother | 2013_P1-16 | | C | G | A | T | C | H | H | H | H | A | H | H | H | H | G | H | H | A | H | G | G | C | T | C | C | T | H | C | G | C | H | EQV | A | A | H | H | A | H | H | C | H | C | H | C | H | C | H | G | H | G | H | T | G | T | A | G | H | G | C | C | C | H | C | H | G | H | G | G | C | T | G | H | A | G | H | T | H |
| 2013_P1-16-1 | PHI1 | PHI2 | progeny | 2013_P1-16 | | C | G | A | T | C | H | H | H | H | A | H | H | H | H | G | A | G | A | A | G | G | C | EQV | C | C | T | H | C | G | C | A | C | A | A | T | G | A | C | A | C | H | C | A | C | C | C | G | G | G | G | H | T | G | T | A | G | T | G | C | C | C | G | C | T | G | T | G | G | C | T | G | G | A | G | G | T | H |
| 2013_P1-16-2 | PHI1 | PHI2 | progeny | 2013_P1-16 | | C | G | A | T | C | H | H | H | H | A | H | H | H | H | G | C | A | A | G | G | G | C | T | C | C | T | T | C | G | C | A | C | A | EQV | EQV | G | A | C | A | C | H | C | A | C | H | C | H | G | H | G | H | T | G | T | A | G | H | G | C | C | C | A | C | C | G | C | G | G | C | T | G | C | A | G | A | T | H |
| 2013_P1-17 | PHI1 | PHI2 | mother | 2013_P1-17 | | C | G | A | T | C | H | A | A | H | H | C | H | H | H | G | H | H | A | H | G | G | C | H | C | C | T | H | C | G | C | EQV | EQV | A | A | H | H | A | H | H | C | H | C | H | C | H | C | H | G | H | G | G | T | H | H | A | G | H | G | C | C | C | H | C | H | G | H | G | G | C | T | G | H | A | G | H | T | H |
| 2013_P1-17-1 | PHI1 | PHI2 | progeny | 2013_P1-17 | | C | EQV | H | T | C | H | H | A | H | EQV | H | EQV | H | H | G | C | A | A | G | G | G | C | T | C | C | T | H | C | G | C | A | C | A | EQV | T | G | A | C | A | C | C | C | H | C | H | C | H | G | G | G | G | T | A | A | A | G | H | G | C | C | C | G | C | T | G | T | G | G | C | T | G | G | A | G | H | T | H |
| 2013_P1-17-2 | PHI1 | PHI2 | progeny | 2013_P1-17 | | C | G | A | T | C | H | A | A | H | H | C | H | H | H | G | C | A | A | G | G | G | C | H | C | C | T | H | C | G | C | H | EQV | A | A | H | H | A | H | H | C | H | C | A | C | C | C | H | G | H | G | G | T | H | A | A | G | H | G | C | C | C | A | C | C | G | C | G | G | C | T | G | C | A | G | A | T | H |
| 2013_P1-19 | PHI1 | PHI2 | mother | 2013_P1-19 | | C | G | A | T | C | H | A | A | H | H | C | T | H | T | H | H | H | A | H | G | G | H | T | C | C | T | H | C | G | C | H | EQV | A | A | H | H | A | H | H | C | H | C | H | C | H | C | H | G | H | G | G | T | H | H | A | G | H | G | C | C | C | H | C | H | G | H | G | G | C | T | G | H | A | G | H | T | H |
| 2013_P1-19-1 | PHI1 | PHI2 | progeny | 2013_P1-19 | | C | G | A | T | C | C | A | A | H | H | C | T | H | T | H | H | H | A | H | G | G | H | T | C | C | T | H | C | G | C | A | C | A | A | T | G | A | H | H | C | H | C | G | C | T | C | A | G | G | G | G | T | A | A | A | G | G | G | C | C | C | A | C | C | G | C | G | G | C | T | G | C | A | G | A | T | H |
| 2013_P1-19-2 | PHI1 | PHI2 | progeny | 2013_P1-19 | | C | G | A | T | C | H | A | A | H | H | C | T | H | T | H | C | A | A | G | G | G | C | T | C | C | T | C | C | G | C | H | EQV | A | A | H | EQV | A | H | H | C | H | C | H | C | H | C | H | G | H | G | G | T | H | H | A | G | H | G | C | C | C | H | C | H | G | H | G | G | C | T | G | H | A | G | H | T | H |
| 2013_P1-19-3 | PHI1 | PHI2 | progeny | 2013_P1-19 | | C | G | A | T | C | H | A | A | H | H | C | T | H | T | H | H | H | A | H | G | G | H | T | C | C | T | H | C | G | C | H | EQV | A | A | H | H | A | H | H | C | H | C | H | C | C | C | H | G | H | G | G | T | H | H | A | G | H | G | C | C | C | H | C | H | G | H | G | G | C | T | G | H | A | G | G | T | H |
| 2013_P1-19-4 | PHI1 | PHI2 | progeny | 2013_P1-19 | | C | G | A | T | C | H | A | A | H | H | C | T | H | T | H | H | H | A | H | G | G | H | T | C | C | T | H | C | G | C | H | EQV | A | A | H | H | A | H | H | C | H | C | H | C | H | C | A | G | C | G | G | T | H | H | A | G | G | G | C | C | C | A | C | C | G | C | G | G | C | T | G | H | A | G | A | T | H |
| 2013_P1-19-5 | PHI1 | PHI2 | progeny | 2013_P1-19 | | C | G | EQV | T | C | H | A | A | H | H | C | T | H | T | H | H | H | A | H | G | G | H | T | C | C | T | H | C | G | C | A | C | A | A | T | G | A | C | A | C | C | C | A | C | C | C | G | G | H | G | G | T | H | H | A | G | H | G | C | C | C | G | C | T | G | T | G | G | C | T | G | H | A | G | A | T | T |
| 2013_P1-20 | PHI1 | PHI2 | mother | 2013_P1-20 | | C | G | A | T | C | C | H | H | H | A | H | H | H | H | G | H | H | A | H | G | H | H | T | C | H | T | H | C | G | C | H | EQV | A | A | H | H | A | H | H | C | H | C | H | C | H | C | H | G | H | G | H | T | G | T | A | G | H | G | C | EQV | C | H | C | H | G | H | G | G | C | T | G | H | A | G | H | T | H |
| 2013_P1-20-1 | PHI1 | PHI2 | progeny | 2013_P1-20 | | C | G | A | T | C | C | G | G | T | A | T | A | G | C | G | H | H | A | H | G | H | H | T | C | H | T | H | C | G | C | H | EQV | A | A | H | H | A | H | H | C | H | C | G | C | T | C | A | G | H | G | H | T | G | T | A | G | H | G | C | C | C | H | C | C | G | C | G | G | C | T | G | C | A | G | A | T | H |
| 2013_P1-20-2 | PHI1 | PHI2 | progeny | 2013_P1-20 | | C | G | A | T | C | C | H | H | T | A | T | A | G | C | G | EQV | A | A | G | G | G | C | T | C | H | T | H | C | G | C | A | C | A | A | T | G | A | C | A | C | H | C | H | C | H | C | H | G | H | G | H | T | G | T | A | G | G | G | C | C | C | A | C | C | G | C | G | G | C | T | G | C | A | G | A | T | T |
| 2013_P1-21 | PHI1 | PHI2 | mother | 2013_P1-21 | | C | G | A | T | C | H | A | A | H | H | C | T | H | T | H | H | H | A | H | G | G | C | EQV | C | H | T | H | C | G | C | H | EQV | A | A | H | H | A | H | H | C | H | C | H | C | H | C | H | G | H | G | H | T | G | T | A | G | H | G | C | C | C | H | C | H | G | H | G | G | C | T | G | H | A | G | H | T | H |
| 2013_P1-21-1 | PHI1 | PHI2 | progeny | 2013_P1-21 | | C | G | A | T | C | T | A | A | T | C | C | T | G | T | A | H | H | A | H | G | G | C | H | C | H | T | H | C | G | C | H | EQV | A | A | T | G | A | C | A | C | C | C | A | C | C | C | A | G | C | G | G | T | G | T | A | G | G | G | C | C | C | A | C | C | G | C | G | G | C | T | G | H | A | G | H | T | G |
| 2013_P1-21-2 | PHI1 | PHI2 | progeny | 2013_P1-21 | | C | G | A | T | C | H | A | A | H | H | C | T | H | T | H | H | H | A | H | G | G | C | H | C | H | T | T | C | G | C | A | C | A | A | T | G | A | C | A | C | C | C | H | C | H | C | H | G | G | G | C | T | G | T | A | G | H | G | C | C | C | A | C | C | G | C | G | G | C | T | G | C | A | G | H | T | H |
| 2013_P1-21-3 | PHI1 | PHI2 | progeny | 2013_P1-21 | | C | G | A | T | C | C | A | A | C | A | C | T | C | T | G | H | H | A | H | G | G | C | H | C | T | T | T | C | G | C | A | C | A | A | T | G | A | C | H | C | H | C | H | C | H | C | H | G | H | G | H | T | G | T | A | G | H | G | C | C | C | A | C | C | G | C | G | G | C | T | G | H | A | G | H | T | H |
| 2013_P1-21-4 | PHI1 | PHI2 | progeny | 2013_P1-21 | | C | G | A | T | C | H | A | A | C | A | C | T | C | T | G | C | A | A | G | G | G | C | T | C | C | T | C | C | G | C | G | G | A | A | G | C | A | T | H | C | H | C | H | C | H | C | H | G | H | G | H | T | G | T | A | G | H | G | C | C | C | A | C | C | G | C | G | G | C | T | G | H | A | G | H | T | H |
| 2013_P1-22 | PHI1 | PHI2 | mother | 2013_P1-22 | | C | G | A | T | C | C | H | EQV | H | A | H | H | H | H | G | H | H | A | H | G | G | C | H | C | H | T | H | C | G | C | EQV | C | A | A | H | H | A | H | H | C | H | C | H | H | H | H | H | G | H | G | G | T | H | H | A | G | H | G | C | C | C | H | C | H | G | H | G | G | C | T | G | H | A | G | H | T | H |
| 2013_P1-22-1 | PHI1 | PHI2 | progeny | 2013_P1-22 | | C | EQV | H | T | C | C | G | H | T | H | T | H | H | H | G | H | H | A | H | G | H | C | EQV | C | H | T | H | C | G | C | EQV | EQV | A | H | H | H | H | H | H | C | EQV | C | G | C | H | C | H | H | H | G | G | T | H | EQV | A | G | H | G | C | C | H | H | C | H | H | H | EQV | G | C | T | G | G | H | G | G | T | G |
| 2013_P1-22-2 | PHI1 | PHI2 | progeny | 2013_P1-22 | | C | G | A | T | C | C | G | G | T | A | T | A | G | C | G | A | G | A | A | G | G | C | A | C | T | T | T | C | G | C | A | C | A | A | T | G | A | C | A | C | C | C | A | H | H | H | H | G | G | G | G | T | A | A | A | G | G | G | C | C | C | A | C | C | G | C | G | G | C | T | G | C | A | G | H | T | H |
| 2013_P1-25 | PHI1 | PHI2 | mother | 2013_P1-25 | | C | G | A | T | C | C | H | H | H | A | H | H | H | H | G | H | H | A | H | G | G | C | EQV | C | H | T | H | C | G | C | H | EQV | A | A | H | H | A | H | H | C | H | C | H | H | H | H | H | G | H | G | G | T | H | H | A | G | H | G | C | C | C | H | C | H | G | H | G | G | C | T | G | H | A | G | H | T | H |
| 2013_P1-25-1 | PHI1 | PHI2 | progeny | 2013_P1-25 | | C | G | A | T | C | C | H | H | H | A | H | H | H | H | G | H | H | A | H | G | G | C | H | C | H | T | H | C | G | C | H | EQV | A | A | H | EQV | A | H | H | C | H | C | H | H | H | H | H | G | H | G | G | T | H | H | A | G | H | G | C | C | C | H | C | H | G | H | G | G | C | T | G | H | A | G | H | T | H |
| 2013_P1-25-2 | PHI1 | PHI2 | progeny | 2013_P1-25 | | C | G | A | T | C | LS | H | H | C | A | C | T | C | T | G | H | H | A | H | G | G | C | H | C | H | T | T | C | G | C | H | C | A | A | H | H | A | H | H | C | H | C | H | H | C | A | EQV | G | H | G | G | T | H | H | A | G | G | G | C | C | C | G | C | T | G | T | G | G | C | T | G | G | A | G | H | T | H |
| 2013_P1-25-3 | PHI1 | PHI2 | progeny | 2013_P1-25 | | C | G | A | T | C | C | H | H | T | A | T | A | G | C | G | H | H | A | H | G | G | C | H | C | H | T | T | C | G | C | H | EQV | A | A | H | H | A | H | H | C | H | C | A | H | H | H | H | G | H | G | G | T | A | A | A | G | H | G | C | C | C | H | C | H | G | H | G | G | C | T | G | H | A | G | H | T | T |
| 2013_P1-25-4 | PHI1 | PHI2 | progeny | 2013_P1-25 | | C | G | A | T | C | C | H | H | H | A | H | H | H | H | G | A | G | A | A | G | G | C | A | C | T | T | T | C | G | C | H | EQV | A | A | H | H | A | H | H | C | H | C | H | H | H | H | H | G | H | G | G | T | H | H | A | G | H | G | C | C | C | H | C | H | G | H | G | G | C | T | G | H | A | G | H | T | H |
| 2013_P1-25-5 | PHI1 | PHI2 | progeny | 2013_P1-25 | | C | G | A | T | C | C | H | H | C | A | C | T | C | T | G | C | H | A | H | G | G | C | T | C | C | T | H | C | G | C | EQV | G | A | A | G | C | A | T | C | C | H | C | H | H | H | H | H | G | H | G | G | T | H | H | A | G | H | G | C | C | C | H | C | H | G | H | G | G | C | T | G | H | A | G | H | T | H |
| 2013_P1-25-6 | PHI1 | PHI2 | progeny | 2013_P1-25 | | C | G | A | T | C | C | H | H | H | A | H | H | H | H | G | H | H | A | H | G | G | C | H | C | T | T | H | C | G | C | H | EQV | A | A | T | G | A | C | A | C | C | C | A | T | C | A | H | G | H | G | G | T | H | H | A | G | G | G | C | C | C | H | C | H | G | H | G | G | C | T | G | H | A | G | H | T | H |
| 2013_P1-26 | PHI1 | PHI2 | mother | 2013_P1-26 | | C | G | A | T | C | H | A | H | H | A | H | H | H | H | G | EQV | H | A | H | G | G | C | H | C | H | T | H | C | G | C | H | EQV | A | A | H | H | A | H | H | C | H | C | H | C | H | C | H | G | H | G | H | T | G | T | A | G | H | G | C | C | C | H | C | H | G | H | G | G | C | T | G | H | A | G | H | T | H |
| 2013_P1-26-1 | PHI1 | PHI2 | progeny | 2013_P1-26 | | C | G | A | T | C | C | A | H | H | A | H | H | H | H | G | H | H | A | H | G | G | C | H | C | H | T | T | C | G | C | A | C | A | A | T | G | A | C | A | C | C | C | H | C | T | C | A | G | C | G | G | T | G | T | A | G | H | G | C | C | C | G | C | T | G | T | G | G | C | T | G | H | A | G | H | T | H |
| 2013_P1-27 | PHI1 | PHI2 | mother | 2013_P1-27 | | C | G | A | T | C | H | A | A | H | H | C | T | H | T | H | H | H | A | H | G | G | C | H | C | H | T | H | C | G | C | H | C | A | A | H | H | A | H | H | C | H | C | H | H | H | H | H | G | H | G | G | T | H | H | A | G | H | G | C | C | C | H | C | H | G | H | G | G | C | T | G | H | A | G | H | T | H |
| 2013_P1-27-1 | PHI1 | PHI2 | progeny | 2013_P1-27 | | C | G | A | T | C | H | A | A | C | A | C | T | C | T | G | H | H | A | H | G | G | C | H | C | H | T | T | C | G | C | A | C | A | A | T | G | A | H | H | C | H | C | H | H | H | H | H | G | H | G | G | T | A | A | A | G | H | G | C | C | C | H | C | H | G | H | G | G | C | T | G | H | A | G | H | T | H |
| 2013_P1-28 | PHI1 | PHI2 | mother | 2013_P1-28 | | C | G | A | T | C | C | H | H | H | A | H | H | H | H | G | H | H | A | H | G | H | H | T | C | C | T | H | C | G | C | H | EQV | A | A | H | H | A | H | H | C | H | C | H | H | H | H | H | G | H | G | G | T | H | H | A | G | H | G | C | C | C | H | C | H | G | H | G | G | C | T | G | H | A | G | H | T | H |
| 2013_P1-28-1 | PHI1 | PHI2 | progeny | 2013_P1-28 | | C | G | EQV | T | C | C | G | G | T | A | H | H | H | H | G | H | H | A | H | G | H | H | T | C | C | T | C | C | G | C | G | G | A | A | G | C | A | T | C | C | A | C | H | H | H | H | H | G | H | G | G | T | H | H | A | G | H | G | C | C | C | A | C | C | G | C | G | G | C | T | G | H | A | G | H | T | T |
| 2013_P1-31 | PHI1 | PHI2 | mother | 2013_P1-31 | | C | G | A | T | C | H | A | A | H | A | H | H | H | H | G | H | H | A | H | G | H | H | T | C | C | T | H | C | G | C | H | EQV | A | A | H | EQV | A | H | H | C | H | C | H | C | H | H | H | G | H | G | G | T | G | T | A | G | H | G | C | C | C | H | C | H | G | H | G | G | C | T | G | H | A | G | H | T | H |
| 2013_P1-31-1 | PHI1 | PHI2 | progeny | 2013_P1-31 | | C | G | A | T | C | H | A | A | C | A | C | T | C | T | G | H | H | A | H | G | H | H | T | C | C | T | H | C | G | C | H | EQV | A | A | H | H | A | T | C | C | A | C | G | C | T | C | A | G | H | G | G | T | G | T | A | G | H | G | C | C | C | H | C | H | G | H | G | G | C | T | G | H | A | G | H | T | H |
| 2013_P1-31-2 | PHI1 | PHI2 | progeny | 2013_P1-31 | | C | G | A | T | C | H | EQV | A | H | A | H | H | G | C | G | C | A | A | G | G | H | H | T | C | C | T | H | C | G | C | A | C | A | A | T | G | A | C | A | C | C | C | H | C | H | H | A | G | C | G | G | T | G | T | A | G | T | G | C | C | C | G | C | T | G | T | G | G | C | T | G | G | A | G | H | T | H |
| 2013_O4 | PHI2 | PHI2 | ancestor | 2013_O4 | | C | G | A | T | C | C | A | A | C | A | C | T | C | T | G | C | A | A | G | G | G | C | EQV | C | C | T | C | C | G | C | G | G | A | A | G | C | A | T | C | C | A | C | G | C | T | C | A | G | C | G | G | T | G | T | A | G | T | G | C | C | C | G | C | T | G | T | G | G | C | T | G | G | A | G | G | T | G |
| 2013_O1 | PHI1 | PHI1 | ancestor | 2013_O1 | | C | G | A | T | C | C | G | G | T | A | T | A | G | C | G | A | G | A | A | G | A | T | T | C | C | T | T | C | G | C | A | C | A | A | T | G | A | C | A | C | C | C | A | T | C | A | H | G | G | G | G | T | A | A | A | G | G | G | C | C | C | A | C | C | G | C | G | G | C | T | G | C | A | G | A | T | T |
| 2013_O1_O4 | PHI1 | PHI2 | ancestor | 2013_O1_O4 | | C | G | A | T | C | C | H | H | H | A | H | H | H | H | G | H | H | A | H | G | H | H | T | C | C | T | H | C | G | C | H | EQV | A | A | H | H | A | H | H | C | H | C | H | EQV | H | H | H | G | H | G | G | T | H | H | A | G | H | G | C | C | C | H | C | H | G | H | G | G | C | T | G | H | A | G | H | T | H |

plant_id – unique identifier of the plant
mom_gen – genetic backgroiund of mother in the hy
dad_gen – genetic background of father in the hybı
family_ – identify mother and all the progeny plar
mother_progeny – identify each plant as mother of the fam
Chr – number of the chromosome, where mar
Distance – position of the marker on the correspon
H – heterozygous marker
A, T, G and C – marker homozygous for the correspond
EQV, LS, -9 , etc. – missing values

### Table S1. Results of the SNP array analasis (continue)

| | | | | | Chr | 6 | 6 | 6 | 6 | 6 | 6 | 6 | 6 | 6 | 6 | 6 | 6 | 7 | 7 | 7 | 7 | 7 | 7 | 7 | 7 | 7 | 7 | 7 | 7 | 7 | 7 | 7 | 7 | 7 | 7 | 7 | 7 | 7 | 7 | 7 | 7 | 7 | 7 | 7 | 7 | 7 | 7 | 7 | 7 | 7 | 7 | 7 | 7 | 7 | 8 | 8 | 8 | 8 | 8 | 8 | 8 | 8 | 8 | 8 | 8 | 8 | 8 | 8 | 8 | 8 | 8 | 8 | 8 | 8 | 8 | 8 | 8 | 8 | 8 | 8 | 8 | 8 |
|---|---|---|---|---|---|---|---|---|---|---|---|---|---|---|---|---|---|---|---|---|---|---|---|---|---|---|---|---|---|---|---|---|---|---|---|---|---|---|---|---|---|---|---|---|---|---|---|---|---|---|---|---|---|---|---|---|---|---|---|---|---|---|---|---|---|---|---|---|---|---|---|---|---|---|---|---|---|---|---|---|---|---|
| | | | | | Distance, cM | 114 | 124 | 134 | 143 | 150 | 160 | 170 | 180 | 180 | 180 | 181 | 183 | 1 | 1.3 | 1.7 | 4.2 | 5.6 | 8 | 17 | 26 | 34 | 43 | 52 | 62 | 70 | 75 | 75 | 76 | 76 | 77 | 86 | 94 | 103 | 112 | 121 | 130 | 139 | 146 | 152 | 162 | 173 | 182 | 192 | 200 | 205 | 206 | 206 | 207 | 211 | 1.8 | 2.1 | 3 | 8 | 11 | 19 | 30 | 40 | 50 | 56 | 64 | 74 | 83 | 84 | 84 | 84 | 85 | 93 | 98 | 109 | 118 | 123 | 127 | 132 | 137 | 145 | 153 | 164 |
| plant_id | mom_gen | dad_gen | mother_progeny | family_ | | | | | | | | | | | | | | | | | | | | | | | | | | | | | | | | | | | | | | | | | | | | | | | | | | | | | | | | | | | | | | | | | | | | | | | | | | | | | | |
| 2013_P1-1 | PHI1 | PHI2 | mother | 2013_P1-1 | | H | T | G | H | G | C | C | A | C | A | H | C | G | H | H | C | G | C | G | A | EQV | H | C | C | H | A | H | H | H | C | A | C | G | G | H | C | G | G | C | H | H | C | A | H | G | C | C | C | G | H | G | T | T | A | C | H | T | T | G | T | C | C | H | G | A | H | C | H | C | C | G | H | G | H | T | T | A |
| 2013_P1-1-1 | PHI1 | PHI2 | progeny | 2013_P1-1 | | H | T | G | H | G | C | C | A | C | A | H | C | G | H | H | C | G | C | G | A | EQV | H | C | C | H | A | H | H | H | C | A | C | G | G | H | C | G | G | C | H | H | C | A | H | G | C | C | C | G | H | G | T | T | A | C | H | T | T | G | T | C | C | H | G | A | H | C | H | C | C | G | H | G | H | T | T | A |
| 2013_P1-3 | PHI1 | PHI2 | mother | 2013_P1-3 | | H | T | G | H | G | C | C | A | C | A | H | C | G | H | H | C | G | C | G | A | EQV | H | C | C | H | A | H | H | H | C | A | C | G | G | H | C | G | G | C | H | H | C | A | H | G | C | C | C | G | H | G | T | T | A | C | H | T | T | G | T | C | C | H | G | A | H | C | H | C | C | G | H | G | H | T | T | A |
| 2013_P1-3-1 | PHI1 | PHI2 | progeny | 2013_P1-3 | | H | T | G | H | G | C | H | A | C | A | H | C | G | H | H | C | G | C | H | A | EQV | H | C | LS | G | A | C | C | G | H | A | C | G | G | A | C | G | G | C | H | H | H | H | G | H | H | C | H | H | A | G | T | H | H | C | T | T | T | G | T | H | H | H | H | A | C | C | H | C | EQV | H | C | G | H | T | H | H |
| 2013_P1-3-2 | PHI1 | PHI2 | progeny | 2013_P1-3 | | H | T | G | H | G | C | H | A | C | A | H | C | G | C | H | C | G | C | H | A | EQV | A | C | LS | G | H | C | C | H | H | A | C | G | G | H | C | G | G | C | G | G | H | H | EQV | H | H | C | H | H | A | G | T | H | H | C | T | T | T | G | T | EQV | H | H | H | A | C | C | H | C | C | G | H | G | A | T | H | H |
| 2013_P1-3-3 | PHI1 | PHI2 | progeny | 2013_P1-3 | | H | T | G | EQV | G | C | H | A | C | A | C | C | G | H | EQV | C | G | C | H | A | EQV | H | C | C | H | H | H | EQV | H | C | A | C | G | G | H | C | G | G | C | G | G | H | H | H | EQV | H | C | H | H | A | G | T | H | H | C | T | T | T | G | T | EQV | H | H | H | A | H | C | H | C | H | G | H | G | H | T | H | H |
| 2013_P1-3-4 | PHI1 | PHI2 | progeny | 2013_P1-3 | | C | T | G | H | G | C | H | A | C | A | H | C | G | H | H | C | G | C | H | A | EQV | H | C | C | H | A | H | C | H | H | A | C | G | G | H | C | G | G | C | G | G | H | H | H | H | H | C | C | G | A | G | T | H | H | C | T | T | T | G | T | EQV | H | H | EQV | A | H | C | H | C | C | H | H | G | H | T | H | H |
| 2013_P1-3-5 | PHI1 | PHI2 | progeny | 2013_P1-3 | | EQV | T | G | EQV | G | C | H | A | C | A | C | C | G | H | H | LS | G | C | H | A | A | H | C | C | H | H | H | EQV | T | C | A | C | G | G | H | C | G | G | C | H | H | H | H | G | EQV | H | C | H | H | A | G | T | H | H | C | T | T | T | G | T | C | H | G | G | A | C | C | H | C | EQV | G | H | G | EQV | T | H | EQV |
| 2013_P1-6 | PHI1 | PHI2 | mother | 2013_P1-6 | | H | T | G | H | G | C | C | A | C | A | H | C | G | H | H | C | G | C | G | A | EQV | H | C | C | H | A | H | H | H | C | A | C | G | G | H | C | G | G | C | H | H | C | A | H | G | C | C | C | G | H | G | T | T | A | C | H | T | T | G | T | C | C | H | G | A | H | C | H | C | C | G | H | G | H | T | T | A |
| 2013_P1-6-1 | PHI1 | PHI2 | progeny | 2013_P1-6 | | H | T | G | H | G | C | C | A | C | A | C | C | G | T | C | C | G | C | G | A | EQV | H | C | C | H | A | H | H | H | C | A | C | G | G | H | C | G | G | C | H | H | C | A | A | G | C | C | C | G | H | G | T | T | A | EQV | H | T | T | G | T | C | C | H | G | A | H | C | H | C | C | G | H | G | H | T | T | A |
| 2013_P1-7 | PHI1 | PHI2 | mother | 2013_P1-7 | | H | T | G | H | G | C | C | A | C | A | H | C | G | H | H | C | G | C | G | A | EQV | H | C | C | H | A | H | H | H | C | A | C | G | G | H | C | G | G | C | H | H | C | A | H | G | C | C | C | G | H | G | T | T | A | C | H | T | T | G | T | C | C | H | G | A | H | C | H | C | C | G | H | G | H | T | T | A |
| 2013_P1-7-1 | PHI1 | PHI2 | progeny | 2013_P1-7 | | T | T | G | H | G | C | C | A | C | A | H | C | G | H | C | C | G | C | H | A | EQV | H | C | C | G | H | C | C | G | C | A | C | G | C | H | C | G | G | C | A | T | C | A | G | G | C | C | C | G | H | G | T | T | A | C | H | T | T | G | T | C | C | H | G | A | H | C | H | C | C | G | H | G | H | T | T | A |
| 2013_P1-8 | PHI1 | PHI2 | mother | 2013_P1-8 | | H | T | G | H | G | C | C | A | C | A | H | C | G | H | H | C | G | C | G | A | EQV | H | C | C | H | A | H | H | H | C | A | C | G | G | H | C | G | G | C | H | H | C | A | H | G | C | C | C | G | H | G | T | T | A | C | H | T | T | G | T | C | C | H | G | A | H | C | H | C | C | G | H | G | H | T | T | A |
| 2013_P1-8-1 | PHI1 | PHI2 | progeny | 2013_P1-8 | | H | T | G | EQV | G | C | H | A | C | A | H | C | G | H | H | C | G | C | H | A | EQV | H | C | C | H | A | H | H | H | H | A | C | G | EQV | H | C | G | G | C | H | H | H | H | H | H | H | C | H | H | H | G | T | H | H | C | H | T | T | G | T | EQV | C | H | G | A | H | C | H | C | C | G | EQV | G | H | T | H | H |
| 2013_P1-8-2 | PHI1 | PHI2 | progeny | 2013_P1-8 | | H | T | G | H | G | C | C | A | C | A | T | C | G | H | H | C | G | C | G | A | EQV | H | C | C | H | A | H | H | H | C | A | C | G | G | H | C | G | G | C | H | H | C | A | H | G | C | C | C | G | G | G | T | T | A | C | G | T | T | G | T | C | C | G | G | A | C | C | A | C | C | G | C | G | G | T | T | A |
| 2013_P1-8-3 | PHI1 | PHI2 | progeny | 2013_P1-8 | | H | T | G | H | G | C | H | A | C | A | H | C | G | H | H | C | G | C | H | A | EQV | H | C | C | H | H | H | H | H | H | A | C | G | G | H | C | G | G | C | H | H | H | H | H | H | H | C | H | H | H | G | T | H | H | C | EQV | T | T | G | T | C | C | G | G | A | C | C | H | C | C | G | C | G | H | T | H | H |
| 2013_P1-8-4 | PHI1 | PHI2 | progeny | 2013_P1-8 | | H | T | G | H | G | C | C | A | C | A | H | C | G | H | H | C | G | C | G | A | EQV | H | C | C | H | A | H | H | H | C | A | C | G | G | H | C | G | G | C | H | H | C | A | H | G | C | C | C | G | H | G | T | T | A | C | H | T | T | G | T | C | C | H | G | A | H | C | H | C | C | G | H | G | H | T | T | A |
| 2013_P1-8-5 | PHI1 | PHI2 | progeny | 2013_P1-8 | | H | T | G | H | G | C | H | A | C | A | H | C | G | H | H | C | G | C | H | A | EQV | H | C | C | H | H | H | H | H | H | A | C | G | G | H | C | G | G | C | H | H | H | H | G | H | H | C | H | H | H | G | T | H | H | H | H | T | T | G | T | C | C | H | G | A | H | C | H | C | EQV | H | H | G | A | T | H | H |
| 2013_P1-9 | PHI1 | PHI2 | mother | 2013_P1-9 | | H | T | G | H | G | C | C | A | C | A | H | C | G | H | H | C | G | C | G | A | EQV | H | C | C | H | A | H | H | H | C | A | C | G | G | H | C | G | G | C | H | EQV | C | A | H | G | C | C | C | G | H | G | T | T | A | EQV | H | T | T | G | T | C | C | H | G | A | H | C | H | C | C | G | H | G | H | T | T | A |
| 2013_P1-9-1 | PHI1 | PHI2 | progeny | 2013_P1-9 | | H | T | G | H | G | C | C | A | C | A | H | C | G | H | H | C | G | C | G | A | EQV | H | C | C | H | A | C | C | G | C | A | C | G | G | H | C | G | G | C | A | T | C | A | G | G | C | C | C | G | G | G | T | T | A | H | G | T | T | G | T | C | C | A | G | A | A | C | C | C | C | G | T | G | A | T | T | A |
| 2013_P1-9-2 | PHI1 | PHI2 | progeny | 2013_P1-9 | | H | T | G | H | G | C | C | A | C | A | H | C | G | T | C | C | G | C | G | A | EQV | H | C | C | H | A | C | C | G | C | A | C | G | C | H | C | G | G | C | A | H | C | A | H | G | C | C | C | G | A | G | T | T | A | C | T | T | T | G | T | C | C | G | G | A | C | C | A | C | C | G | C | G | H | T | T | A |
| 2013_P1-9-3 | PHI1 | PHI2 | progeny | 2013_P1-9 | | C | T | G | H | G | C | C | A | C | A | H | C | G | H | H | C | G | C | G | A | EQV | H | C | C | C | A | G | T | T | C | A | C | G | G | H | C | G | G | C | H | H | C | A | H | G | C | C | C | G | H | G | T | T | A | C | H | T | T | G | T | C | C | A | G | A | A | C | H | C | C | G | H | G | H | T | T | A |
| 2013_P1-10 | PHI1 | PHI2 | mother | 2013_P1-10 | | H | T | G | H | G | C | C | A | C | A | H | C | G | H | EQV | C | G | C | G | A | EQV | H | C | C | H | A | H | H | H | C | A | C | G | G | H | C | G | G | C | H | H | C | A | H | G | C | H | C | G | H | G | T | T | A | C | H | T | T | G | T | C | C | H | G | A | H | C | H | C | C | G | H | G | H | T | T | A |
| 2013_P1-10-1 | PHI1 | PHI2 | progeny | 2013_P1-10 | | T | T | G | H | G | C | C | A | C | A | T | C | G | H | H | C | G | C | G | A | EQV | H | C | C | H | A | H | H | H | C | A | C | G | G | H | C | G | G | C | H | H | C | A | H | G | C | H | C | G | A | G | T | T | A | C | T | T | T | G | T | C | C | H | G | A | H | C | H | C | C | G | H | G | H | T | T | A |
| 2013_P1-16 | PHI1 | PHI2 | mother | 2013_P1-16 | | H | T | G | H | G | C | C | A | C | A | H | C | G | H | H | C | G | C | G | A | EQV | H | C | C | H | A | H | H | H | C | A | C | G | G | H | C | G | G | C | H | H | C | A | H | G | C | C | C | G | H | G | T | T | A | C | H | T | T | G | T | C | C | H | G | A | H | C | H | C | C | G | H | G | H | T | T | A |
| 2013_P1-16-1 | PHI1 | PHI2 | progeny | 2013_P1-16 | | H | T | G | H | G | C | C | A | C | A | T | C | G | C | A | C | G | C | G | A | G | A | C | LS | H | A | H | T | T | C | A | C | G | G | G | C | G | G | C | H | T | C | A | G | G | C | C | C | G | A | G | T | T | A | C | T | T | T | G | T | C | C | G | G | A | C | C | A | C | C | G | C | G | H | T | T | EQV |
| 2013_P1-16-2 | PHI1 | PHI2 | progeny | 2013_P1-16 | | H | T | G | H | G | C | C | A | C | A | T | C | G | H | H | C | G | C | G | A | EQV | G | C | C | C | A | G | T | T | C | A | C | G | G | G | C | G | G | C | G | G | C | A | H | G | C | C | C | G | H | G | T | T | A | C | H | T | T | G | T | C | C | G | G | A | C | C | A | C | C | G | C | G | G | T | T | A |
| 2013_P1-17 | PHI1 | PHI2 | mother | 2013_P1-17 | | H | T | G | H | G | C | C | A | C | A | H | C | G | H | H | C | G | C | G | A | EQV | H | C | C | H | A | H | H | H | C | A | C | G | G | H | C | G | G | C | H | H | C | A | H | G | C | C | C | G | H | G | T | T | A | C | H | T | T | G | T | C | C | H | G | A | H | C | H | C | C | G | H | G | H | T | T | A |
| 2013_P1-17-1 | PHI1 | PHI2 | progeny | 2013_P1-17 | | H | T | G | H | G | C | C | A | C | A | C | C | G | H | H | C | G | C | H | A | EQV | H | C | C | H | A | H | C | G | H | A | C | G | C | A | C | G | G | C | H | H | C | A | H | G | C | C | C | G | A | G | T | T | A | EQV | T | T | T | G | T | C | C | G | G | A | C | C | A | C | C | G | C | G | G | T | T | A |
| 2013_P1-17-2 | PHI1 | PHI2 | progeny | 2013_P1-17 | | H | T | G | H | G | C | C | A | C | A | T | C | G | H | H | C | G | C | G | A | EQV | H | C | C | H | A | H | H | H | C | A | C | G | G | H | C | G | G | C | H | H | C | A | H | G | C | C | C | G | G | G | T | T | A | C | G | T | T | G | T | C | C | G | G | A | C | C | A | C | C | G | H | G | H | T | T | A |
| 2013_P1-19 | PHI1 | PHI2 | mother | 2013_P1-19 | | H | T | G | H | G | C | C | A | C | A | H | C | G | H | H | C | G | C | G | A | EQV | H | C | C | H | A | H | H | H | C | A | C | EQV | G | H | C | G | G | C | EQV | H | C | A | H | G | C | C | C | G | H | G | T | T | A | C | H | T | T | G | T | C | C | H | G | A | H | C | H | C | C | G | H | G | H | T | T | A |
| 2013_P1-19-1 | PHI1 | PHI2 | progeny | 2013_P1-19 | | H | T | G | H | G | C | C | A | C | A | C | C | G | H | H | C | G | C | G | A | EQV | H | C | C | C | A | G | T | T | C | A | C | G | G | G | C | G | G | C | H | H | C | A | A | G | C | C | C | G | H | G | T | T | A | C | H | T | T | G | T | C | C | A | G | A | A | C | C | C | C | G | T | G | A | T | T | A |
| 2013_P1-19-2 | PHI1 | PHI2 | progeny | 2013_P1-19 | | H | T | G | H | G | C | C | A | C | A | H | C | G | T | C | C | G | C | G | A | A | H | C | C | H | A | H | H | H | C | A | C | G | G | H | C | G | G | C | H | H | C | A | H | G | C | C | C | G | H | G | T | T | A | C | H | T | T | G | T | C | C | A | G | A | A | C | C | C | C | G | H | G | H | T | T | A |
| 2013_P1-19-3 | PHI1 | PHI2 | progeny | 2013_P1-19 | | H | T | G | H | G | C | C | A | C | A | H | C | G | H | H | C | G | C | G | A | EQV | H | C | C | H | A | H | H | H | C | A | C | G | G | H | C | G | G | C | A | T | C | A | H | G | C | C | C | G | H | G | T | T | A | C | H | T | T | G | T | C | C | H | G | A | H | C | H | C | C | EQV | C | G | G | T | T | A |
| 2013_P1-19-4 | PHI1 | PHI2 | progeny | 2013_P1-19 | | H | T | G | C | G | C | C | A | C | A | T | C | G | H | H | C | G | C | G | A | EQV | A | C | LS | H | A | H | H | H | C | A | C | G | G | H | C | G | G | C | H | H | C | A | H | G | C | C | C | G | H | G | T | T | A | C | H | T | T | G | T | C | C | A | G | A | A | C | C | C | C | G | H | G | H | T | T | A |
| 2013_P1-19-5 | PHI1 | PHI2 | progeny | 2013_P1-19 | | T | T | G | H | G | C | C | A | C | A | C | C | G | H | H | C | G | C | G | A | EQV | H | C | C | H | A | H | C | G | C | A | C | G | C | H | C | G | G | C | H | H | C | A | H | G | C | C | C | G | G | G | T | T | A | C | G | T | T | G | T | C | C | A | G | A | A | C | H | C | C | G | H | G | H | T | T | A |
| 2013_P1-20 | PHI1 | PHI2 | mother | 2013_P1-20 | | H | T | G | H | G | C | C | A | C | A | H | C | G | H | H | C | G | C | G | A | EQV | H | C | C | H | A | H | H | H | C | A | C | G | G | H | C | G | G | C | H | H | C | A | H | G | C | C | C | G | H | G | T | T | A | C | H | T | T | G | T | C | C | H | G | A | H | C | H | C | C | G | H | G | H | T | T | A |
| 2013_P1-20-1 | PHI1 | PHI2 | progeny | 2013_P1-20 | | H | T | G | H | G | C | C | A | C | A | T | C | G | H | H | C | G | C | G | A | EQV | H | C | C | H | A | G | T | T | C | A | C | G | G | G | C | G | G | C | G | G | C | A | H | G | C | C | C | G | G | G | T | T | A | C | H | T | T | G | T | C | C | A | G | A | A | C | C | C | C | G | H | G | H | T | T | A |
| 2013_P1-20-2 | PHI1 | PHI2 | progeny | 2013_P1-20 | | H | T | G | H | G | C | C | A | C | A | C | C | G | T | C | C | G | C | G | A | EQV | G | C | C | G | A | C | C | G | C | A | C | G | C | A | C | G | G | C | H | H | C | A | G | G | C | C | C | G | H | G | T | T | A | C | H | T | T | G | T | C | C | G | G | A | C | C | H | C | C | G | H | G | G | T | T | A |
| 2013_P1-21 | PHI1 | PHI2 | mother | 2013_P1-21 | | H | T | G | H | G | C | C | A | C | A | H | C | G | H | H | C | G | C | G | A | EQV | H | C | C | H | A | H | H | H | C | A | C | G | G | H | C | G | G | C | H | H | C | A | H | G | C | C | C | G | H | G | T | T | A | C | H | T | T | G | T | C | C | H | G | A | H | C | H | C | C | G | H | G | H | T | T | A |
| 2013_P1-21-1 | PHI1 | PHI2 | progeny | 2013_P1-21 | | C | T | G | C | G | C | C | A | C | A | H | C | G | H | H | C | G | C | G | A | EQV | H | C | C | C | A | G | T | T | C | A | C | G | G | A | C | G | G | C | A | T | C | A | H | G | C | C | C | G | H | G | T | T | A | EQV | H | T | T | G | T | C | C | H | G | A | H | C | H | C | C | G | H | G | H | T | T | A |
| 2013_P1-21-2 | PHI1 | PHI2 | progeny | 2013_P1-21 | | H | T | G | H | G | C | C | A | C | A | H | C | G | H | H | C | G | C | G | A | G | A | C | LS | G | A | C | C | G | C | A | C | G | G | H | C | G | G | C | H | H | C | A | A | G | C | C | C | G | H | G | T | T | A | C | H | T | T | G | T | C | C | G | G | A | C | C | A | C | C | G | H | G | H | T | T | A |
| 2013_P1-21-3 | PHI1 | PHI2 | progeny | 2013_P1-21 | | H | T | G | H | G | C | C | A | C | A | C | C | G | H | H | C | G | C | G | A | EQV | G | C | C | C | A | G | T | T | C | A | C | G | G | EQV | C | G | G | C | G | G | C | A | H | G | C | C | C | G | H | G | T | T | A | EQV | H | T | T | G | T | C | C | G | G | A | C | C | A | C | C | G | C | G | G | T | T | A |
| 2013_P1-21-4 | PHI1 | PHI2 | progeny | 2013_P1-21 | | H | T | G | H | G | C | C | A | C | A | H | C | G | H | H | C | G | C | G | A | EQV | A | C | LS | G | A | C | C | G | C | A | C | G | C | H | C | G | G | C | H | H | C | A | H | G | C | C | C | G | H | G | T | T | A | C | H | T | T | G | T | C | C | G | G | A | C | C | A | C | C | G | C | G | H | T | T | A |
| 2013_P1-22 | PHI1 | PHI2 | mother | 2013_P1-22 | | H | T | G | H | G | C | C | A | C | A | H | C | G | H | H | C | G | C | G | A | EQV | H | C | C | H | A | H | H | H | C | A | C | G | G | H | C | G | G | C | H | H | C | A | H | G | C | C | C | G | H | G | T | T | A | C | H | T | T | G | T | C | C | H | G | A | H | C | H | C | C | G | H | G | H | T | T | A |
| 2013_P1-22-1 | PHI1 | PHI2 | progeny | 2013_P1-22 | | C | T | G | H | G | C | H | A | C | A | H | C | G | H | H | EQV | G | C | H | A | A | H | C | C | H | H | H | H | H | C | A | C | G | G | H | C | G | G | C | G | G | H | H | H | H | H | C | H | H | H | G | T | H | H | C | H | T | T | G | T | EQV | H | H | H | A | H | C | C | C | C | G | H | G | A | T | H | H |
| 2013_P1-22-2 | PHI1 | PHI2 | progeny | 2013_P1-22 | | H | T | G | T | G | C | C | A | C | A | C | C | G | H | H | C | G | C | G | A | EQV | H | C | C | C | A | G | T | T | C | A | C | G | G | G | C | G | G | C | G | G | C | A | A | G | C | C | C | G | H | G | T | T | A | C | H | T | T | G | T | C | C | G | G | A | C | C | A | C | C | G | C | G | H | T | T | A |
| 2013_P1-25 | PHI1 | PHI2 | mother | 2013_P1-25 | | H | T | G | H | G | C | C | A | C | A | LS | C | G | H | H | C | G | C | G | A | EQV | H | C | C | H | A | H | H | H | C | A | C | G | G | H | C | G | G | C | H | H | C | A | H | G | C | C | C | G | H | G | T | T | A | C | H | T | T | G | T | C | C | H | G | A | H | C | H | C | C | G | H | G | H | T | T | A |
| 2013_P1-25-1 | PHI1 | PHI2 | progeny | 2013_P1-25 | | H | T | G | H | G | C | C | A | C | A | H | C | G | H | H | C | G | C | G | A | EQV | H | C | C | H | A | H | H | H | C | A | C | G | G | H | C | G | G | C | H | H | C | A | G | G | C | C | C | G | H | G | T | T | A | C | H | T | T | G | T | C | C | H | G | A | H | C | H | C | C | G | H | G | H | T | T | A |
| 2013_P1-25-2 | PHI1 | PHI2 | progeny | 2013_P1-25 | | H | T | G | T | G | C | C | A | C | A | C | C | G | H | H | C | G | C | G | A | EQV | H | C | C | H | A | H | H | H | C | A | C | G | G | A | C | G | G | C | H | H | C | A | H | G | C | C | C | G | G | G | T | T | A | C | G | T | T | G | T | C | C | G | G | A | C | C | A | C | C | G | H | G | A | T | T | A |
| 2013_P1-25-3 | PHI1 | PHI2 | progeny | 2013_P1-25 | | T | T | G | H | G | C | C | A | C | A | H | C | G | C | A | C | G | C | G | A | G | A | C | LS | G | A | C | C | G | C | A | C | G | G | G | C | G | G | C | H | H | C | A | H | G | C | C | C | G | H | G | T | T | A | C | H | T | T | G | T | C | C | H | G | A | H | C | H | C | C | G | H | G | A | T | T | A |
| 2013_P1-25-4 | PHI1 | PHI2 | progeny | 2013_P1-25 | | H | T | G | C | G | C | C | A | C | A | T | C | G | H | H | C | G | C | G | A | EQV | H | C | C | H | A | H | H | H | C | A | C | G | G | H | C | G | G | C | G | G | C | A | A | G | C | C | C | G | G | G | T | T | A | C | G | T | T | G | T | C | C | H | G | A | H | C | H | C | C | G | H | G | H | T | T | A |
| 2013_P1-25-5 | PHI1 | PHI2 | progeny | 2013_P1-25 | | H | T | G | C | G | C | C | A | C | A | H | C | G | H | H | C | G | C | G | A | EQV | H | C | C | H | A | H | H | H | C | A | C | G | G | H | C | G | G | C | H | H | C | A | H | G | C | C | C | G | H | G | T | T | A | EQV | H | T | T | G | T | C | C | H | G | A | H | C | H | C | C | G | H | G | H | T | T | A |
| 2013_P1-25-6 | PHI1 | PHI2 | progeny | 2013_P1-25 | | H | T | G | H | G | C | C | A | C | A | H | C | G | C | A | C | G | C | G | A | A | G | C | C | G | A | C | C | G | C | A | C | G | C | A | C | G | G | C | A | T | C | A | A | G | C | C | C | G | H | G | T | T | A | C | H | T | T | G | T | C | C | H | G | A | H | C | H | C | C | G | H | G | H | T | T | A |
| 2013_P1-26 | PHI1 | PHI2 | mother | 2013_P1-26 | | H | T | G | H | G | C | C | A | C | A | H | C | G | H | H | C | G | C | G | A | EQV | H | C | C | H | A | H | H | H | C | A | C | G | G | H | C | G | G | C | H | H | C | A | H | G | C | C | C | G | H | G | T | T | A | C | H | T | T | G | T | C | C | H | G | A | H | C | H | C | C | G | H | G | H | T | T | A |
| 2013_P1-26-1 | PHI1 | PHI2 | progeny | 2013_P1-26 | | H | T | G | H | G | C | C | A | C | A | H | C | G | H | H | C | G | C | G | A | G | EQV | C | LS | G | A | C | C | G | C | A | C | G | C | A | C | G | G | C | H | H | C | A | H | G | C | C | C | G | H | G | T | T | A | C | H | T | T | G | T | C | C | A | G | A | A | C | C | C | C | G | T | G | H | T | T | A |
| 2013_P1-27 | PHI1 | PHI2 | mother | 2013_P1-27 | | H | T | G | H | G | C | C | A | C | A | H | C | G | H | H | C | G | C | G | A | EQV | H | C | C | H | A | H | H | H | C | A | C | G | G | H | C | G | G | C | H | H | C | A | H | G | C | C | C | G | H | G | T | T | A | C | H | T | T | G | T | C | C | H | G | A | H | C | H | C | C | G | H | G | H | T | T | A |
| 2013_P1-27-1 | PHI1 | PHI2 | progeny | 2013_P1-27 | | H | T | G | H | G | C | C | A | C | A | T | C | G | H | H | C | G | C | G | A | G | A | C | LS | G | A | C | C | G | C | A | C | G | G | H | C | G | G | C | H | H | C | A | G | G | C | C | C | G | H | G | T | T | A | C | H | T | T | G | T | C | C | H | G | A | H | C | H | C | C | G | H | G | H | T | T | A |
| 2013_P1-28 | PHI1 | PHI2 | mother | 2013_P1-28 | | H | T | G | H | G | C | C | A | C | A | H | C | G | H | H | C | G | C | G | A | EQV | H | C | C | H | A | H | H | H | C | A | C | G | G | H | C | G | G | C | H | H | C | A | H | G | C | H | C | G | H | G | T | T | A | C | H | T | T | G | T | C | C | H | G | A | H | C | H | C | C | G | H | G | H | T | T | A |
| 2013_P1-28-1 | PHI1 | PHI2 | progeny | 2013_P1-28 | | T | T | G | T | G | C | C | A | C | A | C | C | G | H | H | C | G | C | G | A | G | A | C | LS | G | A | C | C | G | C | A | C | G | G | H | C | G | G | C | H | H | C | A | H | G | C | H | C | G | H | G | T | T | A | C | H | T | T | G | T | C | C | A | G | A | A | C | C | C | C | G | H | G | G | T | T | A |
| 2013_P1-31 | PHI1 | PHI2 | mother | 2013_P1-31 | | H | T | G | H | G | C | C | A | C | A | H | C | G | H | H | C | G | C | G | A | EQV | H | C | C | H | A | H | H | H | C | A | C | G | G | H | C | G | G | C | H | H | C | A | H | G | C | C | C | G | H | G | T | T | A | C | H | T | T | G | T | C | C | H | G | A | H | C | H | C | C | G | H | G | H | T | T | A |
| 2013_P1-31-1 | PHI1 | PHI2 | progeny | 2013_P1-31 | | H | T | G | H | G | C | C | A | C | A | C | C | G | H | H | C | G | C | G | A | EQV | H | C | C | C | A | G | T | T | C | A | C | G | G | H | C | G | G | C | H | H | C | A | A | G | C | C | C | G | H | G | T | T | A | C | H | T | T | G | T | C | C | G | G | A | C | C | A | C | C | G | H | G | A | T | T | A |
| 2013_P1-31-2 | PHI1 | PHI2 | progeny | 2013_P1-31 | | H | T | G | H | G | C | C | A | C | A | H | C | G | H | H | C | G | C | G | A | EQV | H | C | C | H | A | G | T | T | C | A | C | G | G | H | C | G | G | C | H | H | C | A | G | G | C | C | C | G | H | G | T | T | A | C | H | T | T | G | T | C | C | A | G | A | A | C | C | C | C | G | H | G | H | T | T | A |
| 2013_O4 | PHI2 | PHI2 | ancestor | 2013_O4 | | C | T | G | C | G | C | C | A | C | A | T | C | G | C | A | C | G | C | G | A | EQV | A | C | EQV | G | A | C | C | G | C | A | C | G | C | A | C | G | G | C | A | G | T | A | A | G | C | C | C | G | A | G | T | T | A | C | T | T | T | G | T | C | C | G | G | A | C | C | A | C | C | G | C | G | G | T | T | A |
| 2013_O1 | PHI1 | PHI1 | ancestor | 2013_O1 | | T | T | G | T | G | C | C | A | C | A | C | C | G | T | C | C | G | C | G | A | EQV | G | C | C | C | A | G | T | T | C | A | C | G | G | G | C | G | G | C | G | G | T | A | A | G | C | A | C | G | G | G | T | T | A | C | G | T | T | G | T | C | C | A | G | A | A | C | C | C | C | G | T | G | A | T | T | A |
| 2013_O1_O4 | PHI1 | PHI2 | ancestor | 2013_O1_O4 | | H | T | G | H | G | C | C | A | C | A | H | C | G | H | H | C | G | C | G | A | EQV | H | C | C | H | A | H | H | H | C | A | C | G | G | H | C | G | G | C | H | G | T | A | A | G | C | H | C | G | H | G | T | T | A | C | H | T | T | G | T | C | C | H | G | A | H | C | H | C | C | G | H | G | H | T | T | A |

plant_id – unique identifier of the plant
mom_gen – genetic backgroiund of mother in the hy
dad_gen – genetic background of father in the hybı
family_ – identify mother and all the progeny plar
mother_progeny – identify each plant as mother of the fam
Chr – number of the chromosome, where mar
Distance – position of the marker on the correspon
H – heterozygous marker
A, T, G and C – marker homozygous for the correspond
EQV, LS, -9 , etc. – missing values

**Table S1. Results of the SNP array analasis (continue)**

| | | | | | Chr | 8 | 8 | 8 | 8 | 8 | 8 | 8 | 8 | 8 | 9 | 9 | 9 | 9 | 9 | 9 | 9 | 9 | 9 | 9 | 9 | 9 | 9 | 9 | 9 | 9 | 9 | 9 | 9 | 9 | 9 | 9 | 9 | 9 | 9 | 9 | 9 | 9 | 9 | 9 | 9 | 9 | 9 | 9 | 10 | 10 | 10 | 10 | 10 | 10 | 10 | 10 | 10 | 10 | 10 | 10 | 10 | 10 | 10 | 10 | 10 | 10 | 10 | 10 | 10 | 10 | 10 | 10 | 10 | 10 | 10 | 10 | 10 | 10 | 10 | 10 |
|---|---|---|---|---|---|---|---|---|---|---|---|---|---|---|---|---|---|---|---|---|---|---|---|---|---|---|---|---|---|---|---|---|---|---|---|---|---|---|---|---|---|---|---|---|---|---|---|---|---|---|---|---|---|---|---|---|---|---|---|---|---|---|---|---|---|---|---|---|---|---|---|---|---|---|---|---|---|---|---|---|
| | | | | | Distance, cM | 168 | 174 | 182 | 192 | 199 | 200 | 205 | 207 | 210 | 0.3 | 9 | 12 | 12 | 12 | 18 | 26 | 35 | 45 | 52 | 62 | 73 | 76 | 76 | 76 | 76 | 76 | 85 | 93 | 101 | 112 | 119 | 129 | 138 | 143 | 152 | 163 | 172 | 172 | 180 | 181 | 181 | 181 | 181 | 4.1 | 6.3 | 7.8 | 9.7 | 11 | 18 | 29 | 40 | 48 | 55 | 59 | 61 | 61 | 61 | 61 | 61 | 71 | 82 | 87 | 93 | 101 | 109 | 118 | 128 | 139 | 148 | 154 | 160 | 161 | 162 | 162 | 162 |
| plant_id | mom_gen | dad_gen | mother_progeny | family_ | | | | | | | | | | | | | | | | | | | | | | | | | | | | | | | | | | | | | | | | | | | | | | | | | | | | | | | | | | | | | | | | | | | | | | | | | | |
| 2013_P1-1 | PHI1 | PHI2 | mother | 2013_P1-1 | | H | H | H | H | G | C | C | G | H | EQV | H | H | H | H | G | H | H | A | H | A | C | C | H | T | C | A | C | G | H | H | T | H | T | T | H | H | H | C | A | C | C | G | T | G | A | G | T | G | A | T | C | H | T | H | H | G | H | A | H | H | H | C | H | T | H | C | H | T | T | C | A | C | G | C | H |
| 2013_P1-1-1 | PHI1 | PHI2 | progeny | 2013_P1-1 | | H | H | H | H | G | C | C | G | H | EQV | H | H | H | H | G | H | H | A | H | A | C | C | H | T | C | EQV | C | G | H | H | T | H | T | T | H | H | H | C | A | C | C | G | T | G | A | G | T | G | A | T | C | EQV | T | H | H | G | H | A | H | H | H | C | H | T | H | C | H | T | T | C | A | C | G | C | H |
| 2013_P1-3 | PHI1 | PHI2 | mother | 2013_P1-1 | | H | H | H | H | G | C | C | G | T | EQV | H | H | H | H | G | H | H | A | H | A | H | C | C | H | C | H | H | G | A | H | H | H | T | T | H | H | H | C | A | C | C | G | T | G | A | G | T | G | H | T | C | EQV | T | H | H | G | H | A | H | A | C | H | H | T | H | C | H | T | T | C | A | C | G | C | H |
| 2013_P1-3-1 | PHI1 | PHI2 | progeny | 2013_P1-3 | | H | H | H | H | G | C | C | G | T | EQV | H | EQV | EQV | H | G | H | H | A | C | H | T | C | C | H | C | H | G | G | H | T | A | G | T | T | H | H | H | C | H | H | EQV | G | T | G | A | G | T | G | H | T | C | EQV | T | H | A | G | H | A | H | A | H | H | H | T | H | C | H | T | T | C | A | H | G | C | H |
| 2013_P1-3-2 | PHI1 | PHI2 | progeny | 2013_P1-3 | | H | H | H | H | G | C | C | G | T | EQV | H | H | G | A | G | H | T | A | H | H | H | C | C | T | C | EQV | H | G | H | EQV | H | H | T | T | EQV | H | H | C | H | H | H | G | T | G | A | G | T | G | H | T | C | EQV | T | C | H | G | T | A | EQV | A | H | A | H | T | H | C | H | T | T | C | A | H | G | C | EQV |
| 2013_P1-3-3 | PHI1 | PHI2 | progeny | 2013_P1-3 | | H | H | H | H | G | C | C | G | T | EQV | C | EQV | EQV | H | G | H | H | A | H | H | H | C | C | T | C | EQV | H | G | H | EQV | H | H | T | T | EQV | EQV | H | C | H | H | H | G | T | G | A | G | T | G | H | T | C | EQV | T | C | H | G | T | A | C | A | H | A | H | T | EQV | C | G | T | T | C | A | H | G | C | EQV |
| 2013_P1-3-4 | PHI1 | PHI2 | progeny | 2013_P1-3 | | H | H | H | H | G | C | C | G | T | C | H | EQV | EQV | H | G | T | H | A | EQV | H | H | C | C | H | C | H | EQV | G | T | EQV | H | H | T | T | EQV | H | H | C | H | H | H | G | T | G | A | G | T | G | H | T | C | H | T | H | H | G | H | A | EQV | A | H | H | H | T | H | C | EQV | T | T | C | A | H | G | C | H |
| 2013_P1-3-5 | PHI1 | PHI2 | progeny | 2013_P1-3 | | H | H | H | H | G | C | C | G | T | EQV | EQV | EQV | EQV | H | G | T | H | A | H | H | H | C | C | T | C | EQV | EQV | G | T | EQV | EQV | H | T | T | EQV | H | H | C | H | H | H | G | T | G | A | G | T | G | H | T | C | EQV | T | C | H | G | T | A | C | A | H | A | C | T | H | C | H | T | T | C | A | H | G | C | H |
| 2013_P1-6 | PHI1 | PHI2 | mother | 2013_P1-6 | | H | H | H | H | G | C | C | G | H | EQV | H | H | H | H | G | H | H | A | H | A | H | C | C | H | C | H | C | G | H | H | T | T | H | T | H | H | H | C | A | C | C | G | T | G | A | G | T | G | H | T | C | H | T | H | H | G | H | A | H | A | C | H | H | T | H | C | H | T | T | C | A | C | G | C | H |
| 2013_P1-6-1 | PHI1 | PHI2 | progeny | 2013_P1-6 | | H | H | H | H | G | C | C | G | H | EQV | H | H | H | H | G | H | H | A | H | A | C | C | C | T | C | EQV | C | G | H | H | T | T | H | T | H | G | A | C | A | C | C | G | T | G | A | G | T | G | A | T | C | H | T | H | H | G | H | A | H | A | C | H | H | T | H | C | H | T | T | C | A | C | G | C | H |
| 2013_P1-7 | PHI1 | PHI2 | mother | 2013_P1-7 | | H | H | H | H | G | C | C | G | H | EQV | H | H | H | H | G | H | H | A | H | A | H | C | C | H | C | H | H | G | A | EQV | H | H | H | T | H | H | H | C | A | C | C | G | T | G | A | G | T | G | H | T | C | EQV | T | H | H | G | H | A | H | A | C | H | H | T | H | C | H | T | T | C | A | C | G | C | EQV |
| 2013_P1-7-1 | PHI1 | PHI2 | progeny | 2013_P1-7 | | H | H | H | H | G | C | C | G | C | EQV | H | EQV | H | H | G | H | H | A | H | A | H | C | C | H | C | H | EQV | G | A | H | H | H | T | T | EQV | H | H | C | A | C | C | G | T | G | A | G | T | G | H | T | C | G | T | H | H | G | H | A | H | A | C | H | H | T | EQV | C | H | T | T | C | A | C | G | C | H |
| 2013_P1-8 | PHI1 | PHI2 | mother | 2013_P1-8 | | H | H | H | H | G | C | C | G | T | C | C | C | H | H | G | H | H | A | H | A | H | C | C | H | C | H | H | G | A | H | T | T | H | T | H | H | H | C | A | C | C | G | T | G | A | G | T | G | A | T | C | EQV | T | H | H | G | H | A | H | A | C | H | H | T | H | C | H | T | T | C | A | C | G | C | EQV |
| 2013_P1-8-1 | PHI1 | PHI2 | progeny | 2013_P1-8 | | T | T | G | H | G | C | C | G | T | C | C | C | G | A | G | H | T | A | H | H | H | C | C | T | C | EQV | H | G | H | EQV | H | H | T | T | G | H | G | C | H | H | H | G | T | G | A | G | T | G | A | T | C | H | T | H | H | G | H | A | EQV | A | H | H | H | T | H | C | H | T | T | C | A | H | G | C | A |
| 2013_P1-8-2 | PHI1 | PHI2 | progeny | 2013_P1-8 | | T | T | G | G | G | C | C | G | T | C | C | C | A | G | G | T | C | A | H | A | C | C | C | T | C | EQV | C | G | A | EQV | T | T | H | T | H | H | H | C | A | C | C | G | T | G | A | G | T | G | A | T | C | A | T | T | A | G | A | A | A | A | C | C | A | T | H | C | H | T | T | C | A | C | G | C | H |
| 2013_P1-8-3 | PHI1 | PHI2 | progeny | 2013_P1-8 | | H | H | H | H | G | C | C | G | T | C | C | C | EQV | H | G | H | H | A | H | H | H | C | H | T | C | EQV | H | G | EQV | EQV | H | H | H | T | EQV | EQV | H | C | H | H | H | G | T | G | A | G | T | G | A | T | C | H | T | C | H | G | T | A | C | A | H | H | H | T | H | C | H | T | T | C | A | H | G | C | H |
| 2013_P1-8-4 | PHI1 | PHI2 | progeny | 2013_P1-8 | | H | H | H | H | G | C | C | G | T | C | C | C | H | H | G | H | H | A | H | A | H | C | C | H | C | H | H | G | EQV | H | T | T | H | T | H | H | H | C | A | C | C | G | T | G | A | G | T | G | A | T | C | H | T | H | H | G | H | A | H | A | C | H | H | T | H | C | H | T | T | C | A | C | G | C | H |
| 2013_P1-8-5 | PHI1 | PHI2 | progeny | 2013_P1-8 | | H | H | H | H | G | C | C | G | T | C | C | C | H | H | EQV | T | H | A | H | H | H | C | C | H | C | H | EQV | G | EQV | H | H | H | EQV | T | EQV | H | EQV | C | H | H | H | G | T | G | A | G | T | G | A | T | C | H | T | H | H | G | H | A | EQV | A | H | H | H | T | H | C | H | T | T | C | A | H | G | C | H |
| 2013_P1-9 | PHI1 | PHI2 | mother | 2013_P1-9 | | H | H | H | H | G | C | C | G | T | C | C | C | H | H | G | H | H | A | EQV | A | C | C | H | T | C | A | C | G | EQV | EQV | T | H | EQV | T | H | H | A | C | A | C | C | G | T | G | A | G | T | G | H | T | C | H | T | H | H | G | H | A | H | A | C | H | H | T | H | C | H | T | T | C | A | C | G | C | H |
| 2013_P1-9-1 | PHI1 | PHI2 | progeny | 2013_P1-9 | | H | H | H | H | G | C | C | G | T | C | C | C | H | H | G | H | H | A | H | A | C | C | T | T | C | EQV | C | G | T | T | T | H | EQV | T | H | H | EQV | C | A | C | C | G | T | G | A | G | T | G | A | T | C | A | T | T | A | G | A | A | A | A | C | H | H | T | T | C | G | T | T | C | A | C | G | C | G |
| 2013_P1-9-2 | PHI1 | PHI2 | progeny | 2013_P1-9 | | T | T | H | H | G | C | C | G | T | C | C | C | H | H | G | H | H | A | C | A | C | C | T | T | C | EQV | C | G | H | H | T | H | T | T | H | H | G | C | A | C | C | G | T | G | A | G | T | G | H | T | C | H | T | C | G | G | T | A | C | A | C | A | C | T | H | C | H | T | T | C | A | C | G | C | H |
| 2013_P1-9-3 | PHI1 | PHI2 | progeny | 2013_P1-9 | | H | H | H | H | G | C | C | G | T | C | C | C | A | G | G | H | H | A | H | A | C | C | C | T | C | EQV | C | G | H | H | T | H | T | T | H | H | H | C | A | C | C | G | T | G | A | G | T | G | H | T | C | H | T | T | A | G | A | A | A | A | C | C | A | T | T | C | H | T | T | C | A | C | G | C | H |
| 2013_P1-10 | PHI1 | PHI2 | mother | 2013_P1-10 | | H | H | H | H | G | C | C | G | T | C | C | C | H | H | G | H | H | A | H | A | C | C | H | T | C | A | C | G | H | H | T | T | H | T | H | H | H | C | A | C | C | G | T | G | A | G | T | G | A | T | C | EQV | T | H | H | G | H | A | H | H | H | C | H | T | H | C | H | T | T | C | A | C | G | C | H |
| 2013_P1-10-1 | PHI1 | PHI2 | progeny | 2013_P1-10 | | A | A | C | C | G | C | C | G | T | C | C | C | G | A | G | C | T | A | T | A | C | C | H | T | C | EQV | C | G | H | H | T | T | H | T | H | H | H | C | A | C | C | G | T | G | A | G | T | G | A | T | C | G | T | C | G | G | T | A | C | G | G | C | C | T | C | C | H | T | T | C | A | C | G | C | H |
| 2013_P1-16 | PHI1 | PHI2 | mother | 2013_P1-16 | | H | H | H | H | G | C | C | G | H | EQV | H | H | H | H | G | H | H | A | H | A | H | C | C | H | C | H | C | G | EQV | H | T | T | H | T | H | H | H | C | A | C | C | G | T | G | A | G | T | G | H | T | C | EQV | T | H | H | G | H | A | H | A | C | H | H | T | H | C | H | T | T | C | A | C | G | C | H |
| 2013_P1-16-1 | PHI1 | PHI2 | progeny | 2013_P1-16 | | A | A | C | C | G | C | C | G | H | EQV | H | H | H | H | G | H | H | A | T | A | C | C | C | T | C | EQV | C | G | A | C | T | T | H | T | H | H | H | C | A | C | C | G | T | G | A | G | T | G | H | T | C | H | T | H | H | G | H | A | H | A | C | A | C | T | C | C | H | T | T | C | A | C | G | C | H |
| 2013_P1-16-2 | PHI1 | PHI2 | progeny | 2013_P1-16 | | T | T | G | G | G | C | C | G | T | EQV | H | H | H | H | G | C | T | A | T | A | C | C | C | T | C | EQV | C | G | EQV | H | T | T | EQV | T | H | H | A | C | A | C | C | G | T | G | A | G | T | G | A | T | C | EQV | T | C | G | G | T | A | H | A | C | EQV | H | T | H | C | H | T | T | C | A | C | G | C | H |
| 2013_P1-17 | PHI1 | PHI2 | mother | 2013_P1-17 | | H | H | H | H | G | C | C | G | T | C | C | C | H | H | G | H | H | A | H | A | H | C | C | H | C | H | H | G | A | C | H | H | EQV | T | H | H | H | C | A | C | C | G | T | G | A | G | T | G | A | T | C | EQV | T | H | H | G | H | A | H | A | C | H | H | T | H | C | H | T | T | C | A | C | G | C | H |
| 2013_P1-17-1 | PHI1 | PHI2 | progeny | 2013_P1-17 | | H | H | H | H | G | C | C | G | T | C | C | C | H | H | G | C | T | A | T | A | C | C | C | T | C | EQV | C | G | A | H | H | H | EQV | T | H | G | A | C | A | C | C | G | T | G | A | G | T | G | A | T | C | EQV | T | H | A | G | H | A | EQV | A | H | H | H | T | H | C | H | T | T | C | A | C | G | C | H |
| 2013_P1-17-2 | PHI1 | PHI2 | progeny | 2013_P1-17 | | H | A | C | C | G | C | C | G | T | C | C | C | H | H | G | H | H | A | H | A | C | C | C | T | C | EQV | C | G | A | H | A | G | T | T | C | G | EQV | C | A | C | C | G | T | G | A | G | T | G | A | T | C | G | T | C | G | G | T | A | C | A | C | A | C | T | C | C | C | T | T | C | A | C | G | C | H |
| 2013_P1-19 | PHI1 | PHI2 | mother | 2013_P1-19 | | H | H | H | H | G | C | C | G | T | C | C | C | H | H | G | H | H | A | H | A | C | C | H | T | C | A | C | G | EQV | H | T | T | H | T | H | H | H | C | A | C | C | G | T | G | A | G | T | G | H | T | C | H | T | H | H | G | H | A | H | A | C | H | H | T | H | C | H | T | T | C | A | C | G | C | H |
| 2013_P1-19-1 | PHI1 | PHI2 | progeny | 2013_P1-19 | | H | H | H | H | G | C | C | G | T | C | C | C | H | H | G | H | H | A | T | A | C | C | C | T | C | EQV | C | G | A | EQV | T | T | T | T | G | C | G | C | A | C | C | G | T | G | A | G | T | G | H | T | C | G | T | C | G | G | T | A | C | A | C | A | C | T | C | C | C | T | T | C | A | C | G | C | H |
| 2013_P1-19-2 | PHI1 | PHI2 | progeny | 2013_P1-19 | | H | H | H | H | G | C | C | G | T | C | C | C | H | H | G | H | H | A | H | A | C | C | H | T | C | EQV | C | G | H | H | T | T | H | T | H | H | H | C | A | C | C | G | T | G | A | G | T | G | H | T | C | G | T | H | H | G | H | A | H | A | C | H | H | T | H | C | H | T | T | C | A | C | G | C | H |
| 2013_P1-19-3 | PHI1 | PHI2 | progeny | 2013_P1-19 | | H | H | H | H | G | C | C | G | T | C | C | C | H | H | G | H | H | A | H | A | C | C | H | T | C | EQV | C | G | H | H | T | T | H | T | H | H | H | C | A | C | C | G | T | G | A | G | T | G | T | T | C | EQV | T | H | H | G | H | A | H | A | C | H | H | T | H | C | H | T | T | C | A | C | G | C | EQV |
| 2013_P1-19-4 | PHI1 | PHI2 | progeny | 2013_P1-19 | | H | H | H | H | G | C | C | G | T | C | C | C | A | G | G | H | H | A | H | A | C | C | C | T | C | EQV | C | G | A | EQV | T | T | H | T | H | H | H | C | A | C | C | G | T | G | A | G | T | G | H | T | C | H | T | H | H | G | H | A | H | A | C | H | H | T | C | C | C | T | T | C | A | C | G | C | A |
| 2013_P1-19-5 | PHI1 | PHI2 | progeny | 2013_P1-19 | | H | H | H | H | G | C | C | G | T | C | C | C | H | H | G | H | H | A | H | A | C | C | T | T | C | EQV | C | G | T | T | T | T | H | T | H | H | H | C | A | C | C | G | T | G | A | G | T | G | H | T | C | H | T | T | A | G | A | A | A | A | C | H | H | T | H | C | H | T | T | C | A | C | G | C | H |
| 2013_P1-20 | PHI1 | PHI2 | mother | 2013_P1-20 | | H | H | H | H | G | C | C | G | H | C | C | C | H | H | G | H | H | A | H | A | C | C | H | T | C | EQV | C | G | A | H | H | T | H | T | H | H | H | C | A | C | C | G | T | G | A | G | T | G | A | T | C | H | T | H | H | G | H | A | H | A | C | H | H | T | H | C | H | T | T | C | A | C | G | C | H |
| 2013_P1-20-1 | PHI1 | PHI2 | progeny | 2013_P1-20 | | T | T | G | G | G | C | C | G | T | C | C | C | A | G | G | T | H | A | H | A | C | C | C | T | C | EQV | C | G | EQV | EQV | T | T | T | T | G | C | H | C | A | C | C | G | T | G | A | G | T | G | A | T | C | A | T | T | A | G | A | A | A | A | C | C | A | T | T | C | G | T | T | C | A | C | G | C | H |
| 2013_P1-20-2 | PHI1 | PHI2 | progeny | 2013_P1-20 | | H | H | H | H | G | C | C | G | H | C | C | C | H | H | G | H | H | A | H | A | C | C | T | T | C | EQV | C | G | EQV | H | H | T | EQV | T | H | H | H | C | A | C | C | G | T | G | A | G | T | G | A | T | C | EQV | T | C | G | G | T | A | C | A | C | A | C | T | C | C | C | T | T | C | A | C | G | C | H |
| 2013_P1-21 | PHI1 | PHI2 | mother | 2013_P1-21 | | H | H | H | H | G | C | C | G | H | EQV | H | H | H | H | G | H | H | A | H | A | C | C | H | T | C | A | C | G | EQV | H | T | T | EQV | T | H | H | H | C | A | C | C | G | T | G | A | G | T | G | A | T | C | EQV | T | H | H | G | H | A | H | H | H | C | H | T | H | C | H | T | T | C | A | C | G | C | H |
| 2013_P1-21-1 | PHI1 | PHI2 | progeny | 2013_P1-21 | | H | H | H | H | G | C | C | G | H | EQV | T | T | G | A | G | C | T | A | H | A | C | C | C | T | C | EQV | C | G | A | EQV | T | T | EQV | T | H | H | H | C | A | C | C | G | T | G | A | G | T | G | A | T | C | H | T | T | A | G | A | A | A | A | H | C | H | T | H | C | H | T | T | C | A | C | G | C | H |
| 2013_P1-21-2 | PHI1 | PHI2 | progeny | 2013_P1-21 | | H | H | H | H | G | C | C | G | H | EQV | H | H | H | H | G | H | H | A | C | A | C | C | T | T | C | EQV | C | G | EQV | H | T | T | T | T | G | C | G | C | A | C | C | G | T | G | A | G | T | G | A | T | C | A | T | T | A | G | A | A | A | A | C | C | A | T | H | C | H | T | T | C | A | C | G | C | H |
| 2013_P1-21-3 | PHI1 | PHI2 | progeny | 2013_P1-21 | | H | H | H | H | G | C | C | G | H | EQV | H | H | H | H | G | T | C | A | C | A | C | C | T | T | C | EQV | C | G | T | T | T | T | H | T | H | H | H | C | A | C | C | G | T | G | A | G | T | G | A | T | C | G | T | C | G | G | T | A | C | H | H | C | H | T | H | C | H | T | T | C | A | C | G | C | A |
| 2013_P1-21-4 | PHI1 | PHI2 | progeny | 2013_P1-21 | | T | T | G | G | G | C | C | G | H | EQV | H | H | H | H | G | C | T | A | T | A | C | C | C | T | C | EQV | C | G | H | H | T | T | H | T | H | H | H | C | A | C | C | G | T | G | A | G | T | G | A | T | C | EQV | T | T | A | G | A | A | A | A | C | C | H | T | H | C | H | T | T | C | A | C | G | C | H |
| 2013_P1-22 | PHI1 | PHI2 | mother | 2013_P1-22 | | H | H | H | H | G | C | C | G | H | EQV | H | H | H | H | G | H | H | A | H | A | C | C | H | T | C | A | C | G | H | H | T | H | T | T | H | H | H | C | A | C | C | G | T | G | A | G | T | G | A | T | C | G | T | H | H | G | H | A | H | A | H | C | H | T | H | C | H | T | T | C | A | C | G | C | H |
| 2013_P1-22-1 | PHI1 | PHI2 | progeny | 2013_P1-22 | | H | H | H | C | G | C | C | G | T | EQV | EQV | EQV | EQV | H | G | H | H | A | H | H | H | C | C | T | C | EQV | H | G | H | H | H | H | T | T | G | H | H | C | EQV | EQV | H | G | T | G | A | G | T | G | A | T | C | G | T | C | H | G | T | A | C | A | H | H | H | T | EQV | C | H | T | T | C | A | H | G | C | H |
| 2013_P1-22-2 | PHI1 | PHI2 | progeny | 2013_P1-22 | | H | H | H | H | G | C | C | G | H | EQV | H | H | H | H | G | H | T | A | T | A | C | C | C | T | C | EQV | C | G | A | EQV | T | H | T | T | H | H | H | C | A | C | C | G | T | G | A | G | T | G | A | T | C | H | T | T | A | G | A | A | A | A | C | C | A | T | T | C | G | T | T | C | A | C | G | C | G |
| 2013_P1-25 | PHI1 | PHI2 | mother | 2013_P1-25 | | H | H | H | H | G | C | C | G | T | EQV | EQV | H | H | H | G | H | H | A | H | A | C | C | H | T | C | A | H | G | A | H | H | H | T | T | H | H | H | C | A | C | C | G | T | G | A | G | T | G | H | T | C | H | T | H | H | G | H | A | H | A | C | H | H | T | H | C | H | T | T | C | A | C | G | C | H |
| 2013_P1-25-1 | PHI1 | PHI2 | progeny | 2013_P1-25 | | H | H | H | H | G | C | C | G | T | EQV | H | H | H | H | G | H | H | A | H | A | C | C | H | T | C | EQV | H | G | A | H | H | H | T | T | H | H | A | C | A | C | C | G | T | G | A | G | T | G | H | T | C | H | T | H | H | G | H | A | H | A | C | H | H | T | H | C | H | T | T | C | A | C | G | C | H |
| 2013_P1-25-2 | PHI1 | PHI2 | progeny | 2013_P1-25 | | H | H | H | H | G | C | C | G | T | EQV | C | C | A | G | G | T | H | A | H | A | C | C | T | T | C | EQV | G | G | A | H | H | H | T | T | H | H | H | C | A | C | C | G | T | G | A | G | T | G | H | T | C | A | T | T | A | G | A | A | A | A | C | H | H | T | H | C | H | T | T | C | A | C | G | C | A |
| 2013_P1-25-3 | PHI1 | PHI2 | progeny | 2013_P1-25 | | A | A | C | H | G | C | C | G | T | EQV | H | H | H | H | G | H | H | A | H | A | C | C | H | T | C | EQV | H | G | A | H | H | H | T | T | H | H | H | C | A | C | C | G | T | G | A | G | T | G | H | T | C | EQV | T | C | G | G | T | A | C | A | C | A | C | T | H | C | H | T | T | C | A | C | G | C | H |
| 2013_P1-25-4 | PHI1 | PHI2 | progeny | 2013_P1-25 | | H | H | H | H | G | C | C | G | T | C | C | C | A | G | G | T | H | A | H | A | C | C | H | T | C | EQV | H | G | A | H | H | H | T | T | H | H | H | C | A | C | C | G | T | G | A | G | T | G | H | T | C | EQV | T | H | H | G | H | A | H | A | C | H | H | T | H | C | H | T | T | C | A | C | G | C | H |
| 2013_P1-25-5 | PHI1 | PHI2 | progeny | 2013_P1-25 | | H | H | H | H | G | C | C | G | T | EQV | H | H | H | H | G | H | H | A | T | A | C | C | C | T | C | EQV | H | G | A | H | H | H | EQV | T | H | H | A | C | A | C | C | G | T | G | A | G | T | G | H | T | C | H | T | H | H | G | H | A | H | A | C | H | H | T | C | C | C | T | T | C | A | C | G | C | A |
| 2013_P1-25-6 | PHI1 | PHI2 | progeny | 2013_P1-25 | | H | H | G | G | G | C | C | G | T | EQV | H | H | H | H | G | H | H | A | C | A | C | C | H | T | C | EQV | H | G | A | EQV | H | H | T | T | H | H | H | C | A | C | C | G | T | G | A | G | T | G | H | T | C | H | T | H | H | G | H | A | H | A | C | H | H | T | H | C | H | T | T | C | A | C | G | C | H |
| 2013_P1-26 | PHI1 | PHI2 | mother | 2013_P1-26 | | H | H | H | H | G | C | C | G | H | C | H | H | H | H | G | H | H | A | H | A | H | C | C | H | C | H | H | G | A | H | H | H | T | T | H | H | H | C | A | C | C | G | T | G | A | G | T | G | A | T | C | H | T | H | H | G | H | A | H | H | H | C | H | T | H | C | H | T | T | C | A | C | G | C | H |
| 2013_P1-26-1 | PHI1 | PHI2 | progeny | 2013_P1-26 | | H | H | H | H | G | C | C | G | H | C | H | H | H | H | G | H | H | A | H | A | C | C | C | T | C | EQV | C | G | A | H | H | H | T | T | H | H | H | C | A | C | C | G | T | G | A | G | T | G | A | T | C | G | T | C | G | G | T | A | C | G | H | C | H | T | H | C | H | T | T | C | A | C | G | C | G |
| 2013_P1-27 | PHI1 | PHI2 | mother | 2013_P1-27 | | H | H | H | H | G | C | C | G | T | EQV | H | H | H | H | G | H | H | A | H | A | C | C | H | T | C | A | C | G | H | H | T | T | H | T | H | H | H | C | A | C | C | G | T | G | A | G | T | G | A | T | C | H | T | H | H | G | H | A | H | H | H | C | H | T | H | C | H | T | T | C | A | C | G | C | H |
| 2013_P1-27-1 | PHI1 | PHI2 | progeny | 2013_P1-27 | | H | H | G | G | G | C | C | G | T | EQV | H | H | H | H | G | H | C | A | C | A | C | C | T | T | C | EQV | C | G | T | H | T | T | H | T | C | G | A | C | A | C | C | G | T | G | A | G | T | G | A | T | C | A | T | T | A | G | A | A | A | A | H | C | H | T | H | C | H | T | T | C | A | C | G | C | A |
| 2013_P1-28 | PHI1 | PHI2 | mother | 2013_P1-28 | | H | H | H | H | G | C | C | G | T | C | H | H | H | H | G | H | H | A | H | A | H | C | C | H | C | H | H | G | H | H | H | H | T | T | H | H | H | C | A | C | C | G | T | G | A | G | T | G | H | T | C | H | T | H | H | G | H | A | H | A | C | H | H | T | H | C | H | T | T | C | A | C | G | C | H |
| 2013_P1-28-1 | PHI1 | PHI2 | progeny | 2013_P1-28 | | T | T | G | G | G | C | C | G | T | C | C | C | A | G | G | T | C | A | H | A | C | C | C | T | C | EQV | C | G | A | H | H | H | T | T | C | G | A | C | A | C | C | G | T | G | A | G | T | G | H | T | C | EQV | T | T | A | G | A | A | A | A | C | C | A | T | T | C | G | T | T | C | A | C | G | C | H |
| 2013_P1-31 | PHI1 | PHI2 | mother | 2013_P1-31 | | H | H | H | H | G | C | C | G | T | C | C | C | H | H | G | H | H | A | H | A | C | C | EQV | T | C | A | C | G | A | H | H | H | T | T | H | H | H | C | A | C | C | G | T | G | A | G | T | G | H | T | C | EQV | T | H | H | G | H | A | H | H | H | C | H | T | H | C | H | T | T | C | A | C | G | C | H |
| 2013_P1-31-1 | PHI1 | PHI2 | progeny | 2013_P1-31 | | H | H | H | H | G | C | C | G | T | C | C | C | H | H | G | H | H | A | H | A | C | C | H | T | C | EQV | C | G | A | H | H | H | T | T | H | H | H | C | A | C | C | G | T | G | A | G | T | G | T | T | C | H | T | H | H | G | H | A | H | H | H | C | H | T | H | C | C | T | T | C | A | C | G | C | H |
| 2013_P1-31-2 | PHI1 | PHI2 | progeny | 2013_P1-31 | | H | H | H | H | G | C | C | G | T | C | C | C | A | G | G | T | C | A | C | A | C | C | T | T | C | EQV | C | G | EQV | H | H | H | T | T | H | H | H | C | A | C | C | G | T | G | A | G | T | G | H | T | C | A | T | T | A | G | A | A | A | H | H | C | H | T | H | C | H | T | T | C | A | C | G | C | H |
| 2013_O4 | PHI2 | PHI2 | ancestor | 2013_O4 | | T | T | G | G | G | C | C | G | T | C | C | C | G | A | G | C | T | A | T | A | C | C | C | T | C | EQV | C | G | A | C | T | T | T | T | G | C | G | C | A | C | C | G | T | G | A | G | T | G | A | T | C | G | T | T | A | G | A | A | A | A | C | C | A | T | T | C | G | T | T | C | A | C | G | C | G |
| 2013_O1 | PHI1 | PHI1 | ancestor | 2013_O1 | | A | A | C | C | G | C | C | G | T | C | C | C | A | G | G | T | C | A | C | A | T | C | C | C | C | G | G | G | A | T | A | G | EQV | T | C | G | A | C | A | C | C | G | T | G | A | G | T | G | T | T | C | G | T | C | G | G | T | A | C | A | EQV | A | C | T | C | C | C | T | T | C | A | C | G | C | A |
| 2013_O1_O4 | PHI1 | PHI2 | ancestor | 2013_O1_O4 | | H | H | H | H | G | C | C | G | T | C | C | C | H | H | G | H | H | A | H | A | H | C | C | H | C | H | H | G | A | H | H | H | T | T | H | H | H | C | A | C | C | G | T | G | A | G | T | G | H | T | C | G | T | H | H | G | H | A | H | A | C | H | H | T | H | C | H | T | T | C | A | C | G | C | H |

plant_id – unique identifier of the plant
mom_gen – genetic backgroiund of mother in the hy
dad_gen – genetic background of father in the hybi
family_ – identify mother and all the progeny plar
mother_progeny – identify each plant as mother of the fam
Chr – number of the chromosome, where mar
Distance – position of the marker on the correspon
H – heterozygous marker
A, T, G and C – marker homozygous for the correspond
EQV, LS, -9 , etc. – missing values

### Table S2. Results of the SNP array analasis

| | | | | | | Chr | 1 | 1 | 1 | 1 | 1 | 1 | 1 | 1 | 1 | 1 | 1 | 1 | 1 | 1 | 1 | 1 | 1 | 1 | 1 | 1 | 1 | 1 | 1 | [illegible] |
|---|---|---|---|---|---|---|---|---|---|---|---|---|---|---|---|---|---|---|---|---|---|---|---|---|---|---|---|---|---|---|
| | | | | | | Position, cM | 9.4 | 16 | 23 | 27 | 52 | 52 | 57 | 59 | 59 | 80 | 81 | 81 | 85 | 89 | 126 | 131 | 132 | 135 | 136 | 137 | 137 | 137 | 137 | [illegible] |
| plant_id | mom_gen | dad_gen | mother_progeny | family_ | plant_id | | | | | | | | | | | | | | | | | | | | | | | | | |
| 2013_P1-1 | PHI1 | PHI2 | mother | 2013_P1-1 | 2013-P1-1 | | H | H | H | LS | T | H | H | H | G | A | G | T | T | H | H | C | A | A | C | A | G | A | T | [illegible] |
| 2013_P1-1-1 | PHI1 | PHI2 | progeny | 2013_P1-1 | 2013-P1-1-1 | | H | H | H | H | T | H | H | H | G | A | G | T | T | H | H | C | A | A | C | A | G | A | T | [illegible] |
| 2013_P1-3 | PHI1 | PHI2 | mother | 2013_P1-1 | 2013-P1-3 | | EQV | H | H | LS | T | H | H | H | G | LS | G | T | T | H | H | C | A | A | C | EQV | G | A | T | [illegible] |
| 2013_P1-3-1 | PHI1 | PHI2 | progeny | 2013_P1-3 | 2013-P1-3-1 | | H | H | H | H | T | H | H | H | G | A | G | EQV | H | H | H | C | A | A | C | H | G | A | T | [illegible] |
| 2013_P1-3-2 | PHI1 | PHI2 | progeny | 2013_P1-3 | 2013-P1-3-2 | | H | H | H | H | T | H | H | H | G | A | G | EQV | H | H | H | C | A | A | C | H | G | A | T | [illegible] |
| 2013_P1-3-3 | PHI1 | PHI2 | progeny | 2013_P1-3 | 2013-P1-3-3 | | H | EQV | H | EQV | T | A | H | T | G | A | G | EQV | H | T | EQV | C | A | A | C | EQV | EQV | A | H | [illegible] |
| 2013_P1-3-4 | PHI1 | PHI2 | progeny | 2013_P1-3 | 2013-P1-3-4 | | H | EQV | EQV | EQV | T | H | H | H | G | A | G | EQV | H | T | H | C | A | A | C | H | G | A | T | [illegible] |
| 2013_P1-3-5 | PHI1 | PHI2 | progeny | 2013_P1-3 | 2013-P1-3-5 | | H | EQV | EQV | LS | T | H | A | T | G | A | G | EQV | T | LS | T | C | A | A | C | EQV | EQV | A | T | [illegible] |
| 2013_P1-6 | PHI1 | PHI2 | mother | 2013_P1-6 | 2013-P1-6 | | EQV | H | G | LS | EQV | H | H | A | G | A | G | T | T | LS | C | C | EQV | A | C | EQV | G | A | EQV | [illegible] |
| 2013_P1-6-1 | PHI1 | PHI2 | progeny | 2013_P1-6 | 2013-P1-6-1 | | H | H | H | T | T | H | H | H | G | A | G | T | T | T | H | C | A | A | C | A | G | A | T | [illegible] |
| 2013_P1-7 | PHI1 | PHI2 | mother | 2013_P1-7 | 2013-P1-7 | | H | H | H | T | T | H | H | H | G | A | G | T | T | T | H | C | A | A | C | A | G | A | T | [illegible] |
| 2013_P1-7-1 | PHI1 | PHI2 | progeny | 2013_P1-7 | 2013-P1-7-1 | | H | H | H | C | T | H | H | H | G | A | G | T | T | C | H | C | A | A | C | A | G | A | T | [illegible] |
| 2013_P1-8 | PHI1 | PHI2 | mother | 2013_P1-8 | 2013-P1-8 | | G | H | H | LS | T | H | H | H | G | A | G | T | T | EQV | H | C | A | A | C | A | G | A | EQV | [illegible] |
| 2013_P1-8-1 | PHI1 | PHI2 | progeny | 2013_P1-8 | 2013-P1-8-1 | | H | H | H | EQV | T | H | H | EQV | G | A | G | EQV | H | H | H | C | A | A | C | H | EQV | A | T | [illegible] |
| 2013_P1-8-2 | PHI1 | PHI2 | progeny | 2013_P1-8 | 2013-P1-8-2 | | T | C | A | C | T | H | H | H | G | A | G | T | T | C | EQV | C | A | A | C | A | G | A | T | [illegible] |
| 2013_P1-8-3 | PHI1 | PHI2 | progeny | 2013_P1-8 | 2013-P1-8-3 | | H | T | H | H | T | H | H | EQV | G | A | G | T | T | EQV | T | C | A | A | C | H | G | A | T | [illegible] |
| 2013_P1-8-4 | PHI1 | PHI2 | progeny | 2013_P1-8 | 2013-P1-8-4 | | H | H | H | H | T | H | H | H | G | A | G | T | T | EQV | H | C | A | A | C | A | G | A | T | [illegible] |
| 2013_P1-8-5 | PHI1 | PHI2 | progeny | 2013_P1-8 | 2013-P1-8-5 | | H | H | H | H | T | H | H | H | G | A | G | T | H | H | H | C | A | A | C | EQV | G | A | T | [illegible] |
| 2013_P1-9 | PHI1 | PHI2 | mother | 2013_P1-9 | 2013-P1-9 | | H | H | EQV | LS | T | EQV | H | H | G | A | G | EQV | T | LS | T | C | EQV | A | C | EQV | G | A | T | [illegible] |
| 2013_P1-9-1 | PHI1 | PHI2 | progeny | 2013_P1-9 | 2013-P1-9-1 | | G | EQV | G | LS | T | H | H | T | G | A | G | T | T | T | T | C | A | A | C | A | G | A | T | [illegible] |
| 2013_P1-9-2 | PHI1 | PHI2 | progeny | 2013_P1-9 | 2013-P1-9-2 | | H | H | H | LS | T | H | H | A | G | A | G | T | T | LS | T | C | A | A | C | A | G | A | T | [illegible] |
| 2013_P1-9-3 | PHI1 | PHI2 | progeny | 2013_P1-9 | 2013-P1-9-3 | | H | H | H | H | T | H | H | H | G | A | G | T | T | T | T | C | A | A | C | A | G | A | T | [illegible] |
| 2013_P1-10 | PHI1 | PHI2 | mother | 2013_P1-10 | 2013-P1-10 | | H | H | EQV | EQV | T | H | H | H | G | A | G | T | T | H | EQV | C | A | A | C | A | G | A | T | [illegible] |
| 2013_P1-10-1 | PHI1 | PHI2 | progeny | 2013_P1-10 | 2013-P1-10-1 | | H | H | H | H | T | A | A | T | G | A | G | T | T | H | H | C | A | A | C | A | G | A | T | [illegible] |
| 2013_P1-16 | PHI1 | PHI2 | mother | 2013_P1-16 | 2013-P1-16 | | H | H | H | EQV | T | H | H | H | G | A | G | T | T | H | H | C | A | A | C | A | G | A | T | [illegible] |
| 2013_P1-16-1 | PHI1 | PHI2 | progeny | 2013_P1-16 | 2013-P1-16-1 | | H | H | H | LS | T | G | H | A | G | A | G | T | T | H | C | C | A | A | C | A | G | A | T | [illegible] |
| 2013_P1-16-2 | PHI1 | PHI2 | progeny | 2013_P1-16 | 2013-P1-16-2 | | T | C | A | EQV | T | A | A | T | G | A | G | T | T | H | C | C | A | A | C | A | G | A | T | [illegible] |
| 2013_P1-17 | PHI1 | PHI2 | mother | 2013_P1-17 | 2013-P1-17 | | H | H | H | LS | T | H | H | H | G | A | G | T | T | T | H | C | EQV | A | C | EQV | G | A | LS | [illegible] |
| 2013_P1-17-1 | PHI1 | PHI2 | progeny | 2013_P1-17 | 2013-P1-17-1 | | H | H | H | H | T | A | A | T | G | A | G | T | T | T | T | C | A | A | C | A | G | A | T | [illegible] |
| 2013_P1-17-2 | PHI1 | PHI2 | progeny | 2013_P1-17 | 2013-P1-17-2 | | G | H | G | LS | T | H | H | H | G | A | G | T | T | LS | C | C | A | A | C | A | G | A | T | [illegible] |
| 2013_P1-19 | PHI1 | PHI2 | mother | 2013_P1-19 | 2013-P1-19 | | H | H | H | EQV | T | H | H | H | G | A | G | T | T | H | H | C | A | A | C | A | G | A | T | [illegible] |
| 2013_P1-19-1 | PHI1 | PHI2 | progeny | 2013_P1-19 | 2013-P1-19-1 | | T | C | H | H | T | H | H | H | G | A | G | T | T | T | T | C | A | A | C | A | G | A | T | [illegible] |
| 2013_P1-19-2 | PHI1 | PHI2 | progeny | 2013_P1-19 | 2013-P1-19-2 | | H | H | H | H | T | H | H | H | G | A | G | T | T | H | H | C | A | A | C | A | G | A | T | [illegible] |
| 2013_P1-19-3 | PHI1 | PHI2 | progeny | 2013_P1-19 | 2013-P1-19-3 | | H | H | H | H | T | H | H | H | G | A | G | T | T | H | H | C | A | A | C | A | G | A | T | [illegible] |
| 2013_P1-19-4 | PHI1 | PHI2 | progeny | 2013_P1-19 | 2013-P1-19-4 | | H | H | H | H | T | H | H | H | G | A | G | T | T | H | H | C | A | A | C | A | G | A | T | [illegible] |
| 2013_P1-19-5 | PHI1 | PHI2 | progeny | 2013_P1-19 | 2013-P1-19-5 | | H | H | H | H | T | G | H | A | G | A | G | T | T | H | C | C | A | A | C | A | G | A | T | [illegible] |
| 2013_P1-20 | PHI1 | PHI2 | mother | 2013_P1-20 | 2013-P1-20 | | H | H | H | LS | T | H | H | H | G | A | G | T | T | EQV | H | C | A | A | C | A | G | A | T | [illegible] |
| 2013_P1-20-1 | PHI1 | PHI2 | progeny | 2013_P1-20 | 2013-P1-20-1 | | H | H | H | H | T | H | H | H | G | A | G | T | T | H | H | C | A | A | C | A | G | A | T | [illegible] |
| 2013_P1-20-2 | PHI1 | PHI2 | progeny | 2013_P1-20 | 2013-P1-20-2 | | G | T | G | T | T | H | H | H | G | A | G | T | T | H | H | C | A | A | C | A | G | A | T | [illegible] |
| 2013_P1-21 | PHI1 | PHI2 | mother | 2013_P1-21 | 2013-P1-21 | | H | H | H | EQV | T | H | H | H | G | A | G | T | T | H | EQV | C | A | A | C | A | G | A | T | [illegible] |
| 2013_P1-21-1 | PHI1 | PHI2 | progeny | 2013_P1-21 | 2013-P1-21-1 | | H | H | H | EQV | T | A | A | T | G | A | G | T | T | EQV | T | C | A | A | C | A | G | A | T | [illegible] |
| 2013_P1-21-2 | PHI1 | PHI2 | progeny | 2013_P1-21 | 2013-P1-21-2 | | H | H | A | C | T | A | A | T | G | A | G | T | T | T | T | C | A | A | C | A | G | A | T | [illegible] |
| 2013_P1-21-3 | PHI1 | PHI2 | progeny | 2013_P1-21 | 2013-P1-21-3 | | H | H | H | H | T | EQV | H | H | G | A | G | T | T | C | C | C | A | A | C | A | G | A | T | [illegible] |
| 2013_P1-21-4 | PHI1 | PHI2 | progeny | 2013_P1-21 | 2013-P1-21-4 | | G | T | G | T | T | H | H | H | G | A | G | T | T | H | T | C | A | A | C | A | G | A | T | [illegible] |
| 2013_P1-22 | PHI1 | PHI2 | mother | 2013_P1-22 | 2013-P1-22 | | H | H | EQV | H | T | H | H | H | G | A | G | T | T | T | H | C | A | A | C | EQV | G | A | T | [illegible] |
| 2013_P1-22-1 | PHI1 | PHI2 | progeny | 2013_P1-22 | 2013-P1-22-1 | | H | T | H | H | T | EQV | H | H | G | A | G | EQV | T | T | H | C | A | A | C | H | G | A | T | [illegible] |
| 2013_P1-22-2 | PHI1 | PHI2 | progeny | 2013_P1-22 | 2013-P1-22-2 | | H | H | H | H | T | H | H | H | G | A | G | T | T | H | C | C | A | A | C | A | G | A | T | [illegible] |
| 2013_P1-25 | PHI1 | PHI2 | mother | 2013_P1-25 | 2013-P1-25 | | H | H | H | H | T | H | H | H | G | A | G | T | T | T | H | C | A | A | C | A | G | A | T | [illegible] |
| 2013_P1-25-1 | PHI1 | PHI2 | progeny | 2013_P1-25 | 2013-P1-25-1 | | H | H | H | H | T | H | H | H | G | A | G | T | T | T | H | C | A | A | C | A | G | A | T | [illegible] |
| 2013_P1-25-2 | PHI1 | PHI2 | progeny | 2013_P1-25 | 2013-P1-25-2 | | T | C | A | C | T | H | H | H | G | A | G | T | T | T | H | C | A | A | C | A | G | A | T | [illegible] |
| 2013_P1-25-3 | PHI1 | PHI2 | progeny | 2013_P1-25 | 2013-P1-25-3 | | H | H | H | H | T | G | H | A | G | A | G | T | T | T | C | C | A | A | C | A | G | A | T | [illegible] |
| 2013_P1-25-4 | PHI1 | PHI2 | progeny | 2013_P1-25 | 2013-P1-25-4 | | H | H | H | H | T | H | H | H | G | A | G | T | T | T | H | C | A | A | C | A | G | A | T | [illegible] |
| 2013_P1-25-5 | PHI1 | PHI2 | progeny | 2013_P1-25 | 2013-P1-25-5 | | H | H | H | C | T | A | A | T | G | A | G | T | T | T | C | C | A | A | C | A | G | A | T | [illegible] |
| 2013_P1-25-6 | PHI1 | PHI2 | progeny | 2013_P1-25 | 2013-P1-25-6 | | T | C | A | C | T | A | H | H | G | A | G | T | T | T | H | C | A | A | C | A | G | A | T | [illegible] |
| 2013_P1-26 | PHI1 | PHI2 | mother | 2013_P1-26 | 2013-P1-26 | | H | H | H | LS | T | H | H | H | G | A | G | T | T | LS | H | C | A | A | C | A | G | A | T | [illegible] |
| 2013_P1-26-1 | PHI1 | PHI2 | progeny | 2013_P1-26 | 2013-P1-26-1 | | H | H | H | H | T | H | H | H | G | A | G | T | T | H | T | C | A | A | C | A | G | A | T | [illegible] |
| 2013_P1-27 | PHI1 | PHI2 | mother | 2013_P1-27 | 2013-P1-27 | | EQV | H | H | H | T | H | H | H | G | A | G | T | T | T | H | C | A | A | C | A | G | A | T | [illegible] |
| 2013_P1-27-1 | PHI1 | PHI2 | progeny | 2013_P1-27 | 2013-P1-27-1 | | G | T | G | H | T | H | H | H | G | A | G | T | T | T | T | C | A | A | C | A | G | A | T | [illegible] |
| 2013_P1-28 | PHI1 | PHI2 | mother | 2013_P1-28 | 2013-P1-28 | | H | H | EQV | EQV | T | H | H | H | G | A | G | T | T | T | H | C | A | A | C | A | G | A | T | [illegible] |
| 2013_P1-28-1 | PHI1 | PHI2 | progeny | 2013_P1-28 | 2013-P1-28-1 | | H | H | H | H | T | H | H | H | G | A | G | T | T | T | T | C | A | A | C | A | G | A | T | [illegible] |
| 2013_P1-31 | PHI1 | PHI2 | mother | 2013_P1-31 | 2013-P1-31 | | H | H | H | C | T | H | H | H | G | A | G | T | T | T | H | C | A | A | C | A | G | A | T | [illegible] |
| 2013_P1-31-1 | PHI1 | PHI2 | progeny | 2013_P1-31 | 2013-P1-31-1 | | H | H | H | H | T | H | H | H | G | A | G | T | T | T | C | C | A | A | C | A | G | A | T | [illegible] |
| 2013_P1-31-2 | PHI1 | PHI2 | progeny | 2013_P1-31 | 2013-P1-31-2 | | H | H | H | H | T | A | H | H | G | A | G | T | T | T | T | C | A | A | C | A | G | A | T | [illegible] |
| 2013_O4 | PHI2 | PHI2 | ancestor | 2013_O4 | 2013-O4 | | T | C | A | C | T | A | A | T | G | A | G | T | T | T | T | C | A | A | C | A | G | A | T | [illegible] |
| 2013_O1 | PHI1 | PHI1 | ancestor | 2013_O1 | 2013-O1 | | G | T | G | T | T | G | H | A | G | A | G | T | T | C | C | C | A | A | C | A | G | A | T | [illegible] |
| 2013_O1_O4 | PHI1 | PHI2 | ancestor | 2013_O1_O4 | 2013-O1_O4 | | H | H | H | LS | T | H | H | H | G | A | G | T | T | LS | H | C | EQV | A | C | A | G | A | LS | [illegible] |

plant_id – unique identifier of the plant
mom_gen – genetic backgroiund of mother in the hybred
dad_gen – genetic background of father in the hybred
family_ – identify mother and all the progeny plants from the same mother
mother_progeny – identify each plant as mother of the family or progeny
Chr – number of the chromosome, where marker is located
Distance – position of the marker on the corresponding chromosome in cM
H – heterozygous marker
A, T, G and C – marker homozygous for the corresponding nucleotide
EQV, LS, -9 , etc. – missing values

### Table S2. Results of the SNP array analasis (continue)

| | | | | | | 3 | 3 | 3 | 3 | |
|---|---|---|---|---|---|---|---|---|---|---|
| | | | | | | 203 | 224 | 227 | 228 | [illegible] |
| plant_id | mom_gen | dad_gen | mother_progeny | family_ | plant_id | | | | | |
| 2013_P1-1 | PHI1 | PHI2 | mother | 2013_P1-1 | 2013-P1-1 | H | H | C | A | [illegible] |
| 2013_P1-1-1 | PHI1 | PHI2 | progeny | 2013_P1-1 | 2013-P1-1-1 | H | H | C | A | [illegible] |
| 2013_P1-3 | PHI1 | PHI2 | mother | 2013_P1-1 | 2013-P1-3 | A | EQV | C | A | [illegible] |
| 2013_P1-3-1 | PHI1 | PHI2 | progeny | 2013_P1-3 | 2013-P1-3-1 | H | H | C | H | [illegible] |
| 2013_P1-3-2 | PHI1 | PHI2 | progeny | 2013_P1-3 | 2013-P1-3-2 | H | EQV | C | EQV | [illegible] |
| 2013_P1-3-3 | PHI1 | PHI2 | progeny | 2013_P1-3 | 2013-P1-3-3 | G | EQV | C | H | [illegible] |
| 2013_P1-3-4 | PHI1 | PHI2 | progeny | 2013_P1-3 | 2013-P1-3-4 | H | EQV | C | EQV | [illegible] |
| 2013_P1-3-5 | PHI1 | PHI2 | progeny | 2013_P1-3 | 2013-P1-3-5 | G | LS | C | LS | [illegible] |
| 2013_P1-6 | PHI1 | PHI2 | mother | 2013_P1-6 | 2013-P1-6 | EQV | EQV | C | A | [illegible] |
| 2013_P1-6-1 | PHI1 | PHI2 | progeny | 2013_P1-6 | 2013-P1-6-1 | H | H | C | A | [illegible] |
| 2013_P1-7 | PHI1 | PHI2 | mother | 2013_P1-7 | 2013-P1-7 | EQV | H | C | A | [illegible] |
| 2013_P1-7-1 | PHI1 | PHI2 | progeny | 2013_P1-7 | 2013-P1-7-1 | H | H | C | A | [illegible] |
| 2013_P1-8 | PHI1 | PHI2 | mother | 2013_P1-8 | 2013-P1-8 | H | T | C | A | [illegible] |
| 2013_P1-8-1 | PHI1 | PHI2 | progeny | 2013_P1-8 | 2013-P1-8-1 | H | T | C | EQV | [illegible] |
| 2013_P1-8-2 | PHI1 | PHI2 | progeny | 2013_P1-8 | 2013-P1-8-2 | H | T | C | A | [illegible] |
| 2013_P1-8-3 | PHI1 | PHI2 | progeny | 2013_P1-8 | 2013-P1-8-3 | G | T | C | EQV | [illegible] |
| 2013_P1-8-4 | PHI1 | PHI2 | progeny | 2013_P1-8 | 2013-P1-8-4 | H | T | C | A | [illegible] |
| 2013_P1-8-5 | PHI1 | PHI2 | progeny | 2013_P1-8 | 2013-P1-8-5 | H | T | C | EQV | [illegible] |
| 2013_P1-9 | PHI1 | PHI2 | mother | 2013_P1-9 | 2013-P1-9 | H | T | C | A | [illegible] |
| 2013_P1-9-1 | PHI1 | PHI2 | progeny | 2013_P1-9 | 2013-P1-9-1 | H | H | C | A | [illegible] |
| 2013_P1-9-2 | PHI1 | PHI2 | progeny | 2013_P1-9 | 2013-P1-9-2 | G | EQV | C | A | [illegible] |
| 2013_P1-9-3 | PHI1 | PHI2 | progeny | 2013_P1-9 | 2013-P1-9-3 | H | H | C | A | [illegible] |
| 2013_P1-10 | PHI1 | PHI2 | mother | 2013_P1-10 | 2013-P1-10 | H | EQV | C | A | [illegible] |
| 2013_P1-10-1 | PHI1 | PHI2 | progeny | 2013_P1-10 | 2013-P1-10-1 | H | H | C | A | [illegible] |
| 2013_P1-16 | PHI1 | PHI2 | mother | 2013_P1-16 | 2013-P1-16 | H | T | C | A | [illegible] |
| 2013_P1-16-1 | PHI1 | PHI2 | progeny | 2013_P1-16 | 2013-P1-16-1 | G | T | C | A | [illegible] |
| 2013_P1-16-2 | PHI1 | PHI2 | progeny | 2013_P1-16 | 2013-P1-16-2 | H | T | C | A | [illegible] |
| 2013_P1-17 | PHI1 | PHI2 | mother | 2013_P1-17 | 2013-P1-17 | EQV | H | C | A | [illegible] |
| 2013_P1-17-1 | PHI1 | PHI2 | progeny | 2013_P1-17 | 2013-P1-17-1 | H | EQV | C | A | [illegible] |
| 2013_P1-17-2 | PHI1 | PHI2 | progeny | 2013_P1-17 | 2013-P1-17-2 | G | T | C | A | [illegible] |
| 2013_P1-19 | PHI1 | PHI2 | mother | 2013_P1-19 | 2013-P1-19 | H | EQV | C | A | [illegible] |
| 2013_P1-19-1 | PHI1 | PHI2 | progeny | 2013_P1-19 | 2013-P1-19-1 | H | T | C | A | [illegible] |
| 2013_P1-19-2 | PHI1 | PHI2 | progeny | 2013_P1-19 | 2013-P1-19-2 | H | T | C | A | [illegible] |
| 2013_P1-19-3 | PHI1 | PHI2 | progeny | 2013_P1-19 | 2013-P1-19-3 | H | T | C | A | [illegible] |
| 2013_P1-19-4 | PHI1 | PHI2 | progeny | 2013_P1-19 | 2013-P1-19-4 | H | T | C | A | [illegible] |
| 2013_P1-19-5 | PHI1 | PHI2 | progeny | 2013_P1-19 | 2013-P1-19-5 | H | T | C | A | [illegible] |
| 2013_P1-20 | PHI1 | PHI2 | mother | 2013_P1-20 | 2013-P1-20 | H | H | C | A | [illegible] |
| 2013_P1-20-1 | PHI1 | PHI2 | progeny | 2013_P1-20 | 2013-P1-20-1 | A | C | C | A | [illegible] |
| 2013_P1-20-2 | PHI1 | PHI2 | progeny | 2013_P1-20 | 2013-P1-20-2 | H | H | C | A | [illegible] |
| 2013_P1-21 | PHI1 | PHI2 | mother | 2013_P1-21 | 2013-P1-21 | H | EQV | C | A | [illegible] |
| 2013_P1-21-1 | PHI1 | PHI2 | progeny | 2013_P1-21 | 2013-P1-21-1 | H | H | C | A | [illegible] |
| 2013_P1-21-2 | PHI1 | PHI2 | progeny | 2013_P1-21 | 2013-P1-21-2 | A | C | C | A | [illegible] |
| 2013_P1-21-3 | PHI1 | PHI2 | progeny | 2013_P1-21 | 2013-P1-21-3 | H | H | C | A | [illegible] |
| 2013_P1-21-4 | PHI1 | PHI2 | progeny | 2013_P1-21 | 2013-P1-21-4 | G | T | C | A | [illegible] |
| 2013_P1-22 | PHI1 | PHI2 | mother | 2013_P1-22 | 2013-P1-22 | H | T | C | A | [illegible] |
| 2013_P1-22-1 | PHI1 | PHI2 | progeny | 2013_P1-22 | 2013-P1-22-1 | H | T | C | EQV | [illegible] |
| 2013_P1-22-2 | PHI1 | PHI2 | progeny | 2013_P1-22 | 2013-P1-22-2 | H | T | C | A | [illegible] |
| 2013_P1-25 | PHI1 | PHI2 | mother | 2013_P1-25 | 2013-P1-25 | H | H | C | A | [illegible] |
| 2013_P1-25-1 | PHI1 | PHI2 | progeny | 2013_P1-25 | 2013-P1-25-1 | G | T | C | A | [illegible] |
| 2013_P1-25-2 | PHI1 | PHI2 | progeny | 2013_P1-25 | 2013-P1-25-2 | EQV | H | C | A | [illegible] |
| 2013_P1-25-3 | PHI1 | PHI2 | progeny | 2013_P1-25 | 2013-P1-25-3 | H | H | C | A | [illegible] |
| 2013_P1-25-4 | PHI1 | PHI2 | progeny | 2013_P1-25 | 2013-P1-25-4 | H | H | C | A | [illegible] |
| 2013_P1-25-5 | PHI1 | PHI2 | progeny | 2013_P1-25 | 2013-P1-25-5 | H | H | C | A | [illegible] |
| 2013_P1-25-6 | PHI1 | PHI2 | progeny | 2013_P1-25 | 2013-P1-25-6 | G | T | C | A | [illegible] |
| 2013_P1-26 | PHI1 | PHI2 | mother | 2013_P1-26 | 2013-P1-26 | H | EQV | C | A | [illegible] |
| 2013_P1-26-1 | PHI1 | PHI2 | progeny | 2013_P1-26 | 2013-P1-26-1 | A | C | C | A | [illegible] |
| 2013_P1-27 | PHI1 | PHI2 | mother | 2013_P1-27 | 2013-P1-27 | H | H | C | A | [illegible] |
| 2013_P1-27-1 | PHI1 | PHI2 | progeny | 2013_P1-27 | 2013-P1-27-1 | H | C | C | A | [illegible] |
| 2013_P1-28 | PHI1 | PHI2 | mother | 2013_P1-28 | 2013-P1-28 | H | T | C | A | [illegible] |
| 2013_P1-28-1 | PHI1 | PHI2 | progeny | 2013_P1-28 | 2013-P1-28-1 | H | T | C | A | [illegible] |
| 2013_P1-31 | PHI1 | PHI2 | mother | 2013_P1-31 | 2013-P1-31 | H | T | C | A | [illegible] |
| 2013_P1-31-1 | PHI1 | PHI2 | progeny | 2013_P1-31 | 2013-P1-31-1 | H | H | C | A | [illegible] |
| 2013_P1-31-2 | PHI1 | PHI2 | progeny | 2013_P1-31 | 2013-P1-31-2 | G | T | C | A | [illegible] |
| 2013_O4 | PHI2 | PHI2 | ancestor | 2013_O4 | 2013-O4 | G | T | C | A | [illegible] |
| 2013_O1 | PHI1 | PHI1 | ancestor | 2013_O1 | 2013-O1 | A | C | C | A | [illegible] |
| 2013_O1_O4 | PHI1 | PHI2 | ancestor | 2013_O1_O4 | 2013-O1_O4 | H | EQV | C | A | [illegible] |

plant_id – unique identifier of the plant
mom_gen – genetic backgroiund of mother in the hyb
dad_gen – genetic background of father in the hybre
family_ – identify mother and all the progeny plants
mother_progeny – identify each plant as mother of the family
Chr – number of the chromosome, where mark
Distance – position of the marker on the correspond
H – heterozygous marker
A, T, G and C – marker homozygous for the correspondir
EQV, LS, -9 , etc. – missing values

### Table S2. Results of the SNP array analasis (continue)

| | | | | | | 7 | 7 | 7 | 7 | 7 | 7 | 7 | 7 | 7 | 7 | 7 | 7 | 7 | 7 | 7 | 7 | 7 | 7 | 7 | 7 | 7 | 7 | 8 | 8 | 8 | 8 | 8 | 8 | 8 | 8 | 8 | 8 | 8 | 8 | 8 | 8 | 8 | 8 | 8 | 8 | 8 | 8 | 8 | 8 | 8 | 8 | 8 | 8 | 8 | 8 | 8 | 8 | 8 |
|---|---|---|---|---|---|---|---|---|---|---|---|---|---|---|---|---|---|---|---|---|---|---|---|---|---|---|---|---|---|---|---|---|---|---|---|---|---|---|---|---|---|---|---|---|---|---|---|---|---|---|---|---|---|---|---|---|---|---|
| | | | | | | 87 | 92 | 95 | 98 | 100 | 111 | 112 | 112 | 129 | 135 | 141 | 145 | 151 | 170 | 175 | 182 | 191 | 196 | 201 | 201 | 205 | 205 | 15 | 20 | 20 | 43 | 46 | 48 | 51 | 51 | 54 | 67 | 73 | 74 | 77 | 84 | 85 | 85 | 85 | 92 | 95 | 108 | 117 | 123 | 133 | 145 | 150 | 155 | 164 | 164 | 195 | 197 | 199 |
| plant_id | mom_gen | dad_gen | mother_progeny | family_ | plant_id | | | | | | | | | | | | | | | | | | | | | | | | | | | | | | | | | | | | | | | | | | | | | | | | | | | | | |
| 2013_P1-1 | PHI1 | PHI2 | mother | 2013_P1-1 | 2013-P1-1 | EQV | C | G | H | A | T | A | C | C | EQV | LS | H | EQV | H | LS | C | C | H | EQV | H | T | C | EQV | T | LS | T | H | H | LS | A | T | H | T | A | H | H | H | G | H | A | H | G | G | T | T | H | EQV | LS | C | G | A | H | G |
| 2013_P1-1-1 | PHI1 | PHI2 | progeny | 2013_P1-1 | 2013-P1-1-1 | A | C | G | H | A | T | A | C | C | H | G | H | H | H | T | C | C | H | C | H | T | C | H | T | G | T | H | H | H | A | T | H | T | A | H | H | H | G | H | A | H | G | G | T | T | H | A | EQV | C | G | A | H | G |
| 2013_P1-3 | PHI1 | PHI2 | mother | 2013_P1-1 | 2013-P1-3 | EQV | C | G | EQV | A | T | A | C | C | EQV | LS | H | T | H | LS | C | C | H | C | H | T | C | T | LS | LS | T | H | H | A | LS | T | H | T | A | H | H | EQV | G | H | A | H | G | G | T | T | H | A | LS | C | G | A | C | G |
| 2013_P1-3-1 | PHI1 | PHI2 | progeny | 2013_P1-3 | 2013-P1-3-1 | EQV | C | G | T | A | H | A | C | C | EQV | LS | H | T | G | C | C | H | T | H | A | H | H | T | T | G | T | C | C | A | A | T | H | T | A | H | H | C | G | H | H | G | G | G | EQV | T | H | EQV | LS | H | G | A | C | H |
| 2013_P1-3-2 | PHI1 | PHI2 | progeny | 2013_P1-3 | 2013-P1-3-2 | EQV | C | G | T | A | H | A | C | C | H | LS | C | EQV | H | C | C | H | H | H | H | H | H | T | T | G | T | C | C | A | A | T | H | T | A | H | H | C | G | H | H | G | G | G | T | T | A | LS | LS | H | G | A | C | H |
| 2013_P1-3-3 | PHI1 | PHI2 | progeny | 2013_P1-3 | 2013-P1-3-3 | EQV | C | G | T | A | EQV | A | C | C | H | LS | EQV | EQV | H | C | C | H | H | EQV | H | H | H | T | T | LS | T | H | C | EQV | A | T | H | EQV | A | H | H | H | G | H | A | H | G | LS | EQV | T | H | EQV | LS | H | G | A | C | H |
| 2013_P1-3-4 | PHI1 | PHI2 | progeny | 2013_P1-3 | 2013-P1-3-4 | LS | C | G | H | A | H | A | C | C | H | LS | C | LS | EQV | C | C | H | H | H | H | H | H | LS | T | EQV | T | C | EQV | EQV | A | T | H | T | A | H | H | H | G | H | H | H | G | G | H | T | H | LS | LS | H | G | A | H | H |
| 2013_P1-3-5 | PHI1 | PHI2 | progeny | 2013_P1-3 | 2013-P1-3-5 | LS | C | G | H | A | H | A | C | C | H | LS | H | T | G | LS | C | H | H | EQV | EQV | H | EQV | LS | LS | LS | T | C | EQV | A | LS | T | A | EQV | A | T | A | C | G | G | A | G | G | G | T | T | H | LS | LS | H | G | A | LS | EQV |
| 2013_P1-6 | PHI1 | PHI2 | mother | 2013_P1-6 | 2013-P1-6 | EQV | C | G | EQV | A | T | A | C | EQV | EQV | LS | LS | EQV | LS | LS | C | C | H | C | EQV | T | C | EQV | LS | EQV | T | H | T | EQV | LS | T | EQV | T | A | H | H | LS | G | H | A | G | G | G | EQV | T | H | EQV | LS | C | G | A | EQV | G |
| 2013_P1-6-1 | PHI1 | PHI2 | progeny | 2013_P1-6 | 2013-P1-6-1 | A | C | G | EQV | A | T | A | C | H | H | G | T | H | H | EQV | C | C | H | C | G | T | C | EQV | T | G | T | H | H | H | A | T | H | T | A | H | H | H | G | H | A | H | G | G | T | T | H | A | EQV | C | G | A | H | G |
| 2013_P1-7 | PHI1 | PHI2 | mother | 2013_P1-7 | 2013-P1-7 | EQV | C | G | EQV | A | T | A | C | C | H | LS | H | T | T | EQV | C | C | H | C | H | T | C | EQV | T | LS | T | H | T | EQV | LS | T | H | T | A | H | H | EQV | G | H | A | H | G | G | T | T | H | A | LS | C | G | A | H | G |
| 2013_P1-7-1 | PHI1 | PHI2 | progeny | 2013_P1-7 | 2013-P1-7-1 | A | C | G | T | A | T | A | C | C | T | EQV | H | C | G | LS | C | C | T | C | A | T | C | T | T | G | T | H | H | EQV | A | T | H | T | A | H | H | H | G | H | A | H | G | G | T | T | H | A | EQV | C | G | A | A | G |
| 2013_P1-8 | PHI1 | PHI2 | mother | 2013_P1-8 | 2013-P1-8 | EQV | C | G | H | A | T | A | C | C | H | LS | H | EQV | EQV | EQV | C | C | H | C | H | T | C | EQV | T | LS | T | H | H | A | LS | T | H | T | A | H | H | EQV | G | H | A | H | G | G | T | T | H | EQV | LS | C | G | A | EQV | G |
| 2013_P1-8-1 | PHI1 | PHI2 | progeny | 2013_P1-8 | 2013-P1-8-1 | EQV | C | G | EQV | A | H | A | C | C | H | LS | EQV | H | H | EQV | C | H | H | H | H | H | H | T | T | G | T | H | H | EQV | A | T | H | T | A | H | H | H | G | H | A | H | G | G | T | T | H | EQV | LS | H | G | A | C | H |
| 2013_P1-8-2 | PHI1 | PHI2 | progeny | 2013_P1-8 | 2013-P1-8-2 | A | C | G | EQV | A | T | A | C | C | H | LS | H | EQV | G | T | C | C | H | C | H | T | C | G | T | G | T | H | EQV | LS | A | T | G | T | A | T | A | C | G | G | A | G | G | G | T | T | G | A | LS | C | G | A | H | G |
| 2013_P1-8-3 | PHI1 | PHI2 | progeny | 2013_P1-8 | 2013-P1-8-3 | EQV | C | G | H | A | H | A | C | C | H | LS | EQV | EQV | EQV | H | C | H | H | H | H | H | H | T | T | LS | T | H | H | A | A | T | H | H | A | T | A | C | G | G | A | G | G | G | T | T | H | EQV | LS | H | G | A | C | H |
| 2013_P1-8-4 | PHI1 | PHI2 | progeny | 2013_P1-8 | 2013-P1-8-4 | A | C | G | H | A | T | A | C | C | H | LS | H | H | H | T | C | C | H | C | H | T | C | EQV | T | LS | T | H | H | EQV | A | T | H | T | A | H | H | H | G | H | A | H | G | G | T | T | H | EQV | LS | C | G | A | H | G |
| 2013_P1-8-5 | PHI1 | PHI2 | progeny | 2013_P1-8 | 2013-P1-8-5 | LS | C | G | EQV | A | H | A | C | C | H | LS | H | T | G | EQV | C | H | T | H | A | H | H | EQV | T | G | T | H | EQV | EQV | A | T | EQV | H | A | H | H | C | G | H | A | H | G | G | EQV | T | A | EQV | EQV | H | G | A | H | H |
| 2013_P1-9 | PHI1 | PHI2 | mother | 2013_P1-9 | 2013-P1-9 | LS | C | G | EQV | A | T | A | C | C | H | LS | H | LS | H | LS | C | C | H | C | H | T | C | LS | T | LS | T | H | T | EQV | LS | T | A | T | A | H | H | C | G | H | A | H | G | LS | T | T | H | EQV | LS | C | G | A | A | G |
| 2013_P1-9-1 | PHI1 | PHI2 | progeny | 2013_P1-9 | 2013-P1-9-1 | EQV | C | G | EQV | A | T | A | C | C | H | LS | T | LS | G | EQV | C | C | T | C | A | T | C | G | LS | G | T | G | H | H | A | T | H | T | A | C | C | A | G | EQV | A | A | G | G | T | T | A | A | T | C | G | A | C | G |
| 2013_P1-9-2 | PHI1 | PHI2 | progeny | 2013_P1-9 | 2013-P1-9-2 | EQV | C | G | T | A | T | A | C | C | T | G | T | EQV | LS | LS | C | C | H | C | H | T | C | EQV | LS | LS | T | C | C | A | LS | T | LS | T | A | T | A | C | G | G | EQV | G | G | G | T | T | A | A | G | C | G | A | LS | G |
| 2013_P1-9-3 | PHI1 | PHI2 | progeny | 2013_P1-9 | 2013-P1-9-3 | A | C | G | H | A | T | A | C | C | H | G | H | H | H | T | C | C | H | C | H | T | C | EQV | T | G | T | H | H | G | A | T | A | T | A | C | C | A | G | A | A | H | G | G | T | T | H | A | EQV | C | G | A | H | G |
| 2013_P1-10 | PHI1 | PHI2 | mother | 2013_P1-10 | 2013-P1-10 | EQV | C | G | H | A | T | A | C | C | H | G | EQV | LS | EQV | LS | C | C | H | C | H | T | C | EQV | T | G | T | H | T | LS | A | T | EQV | T | A | H | H | A | G | H | A | H | G | G | T | T | H | A | T | C | G | A | H | G |
| 2013_P1-10-1 | PHI1 | PHI2 | progeny | 2013_P1-10 | 2013-P1-10-1 | A | C | G | H | A | T | A | C | C | H | G | H | H | H | T | C | C | H | C | H | T | C | T | T | G | T | C | C | A | A | T | H | T | A | H | H | H | G | H | A | H | G | G | T | T | H | A | T | C | G | A | A | G |
| 2013_P1-16 | PHI1 | PHI2 | mother | 2013_P1-16 | 2013-P1-16 | EQV | C | G | EQV | A | EQV | A | C | H | H | LS | T | C | H | EQV | C | C | H | C | H | T | C | T | T | G | T | H | EQV | EQV | A | T | H | T | A | H | H | H | G | H | A | H | G | G | T | T | H | EQV | LS | C | G | A | H | G |
| 2013_P1-16-1 | PHI1 | PHI2 | progeny | 2013_P1-16 | 2013-P1-16-1 | A | C | G | C | A | T | A | C | T | H | G | T | H | G | T | C | C | T | C | A | T | C | T | T | G | T | C | C | A | A | T | G | T | A | T | A | C | G | G | A | G | G | G | T | T | A | A | T | C | G | A | A | G |
| 2013_P1-16-2 | PHI1 | PHI2 | progeny | 2013_P1-16 | 2013-P1-16-2 | A | C | G | C | A | T | A | C | T | C | G | T | C | T | T | C | C | H | C | H | T | C | EQV | T | G | T | H | H | A | A | T | H | T | A | T | A | C | G | G | A | G | G | G | T | T | G | A | G | C | G | A | C | G |
| 2013_P1-17 | PHI1 | PHI2 | mother | 2013_P1-17 | 2013-P1-17 | LS | C | G | H | A | T | A | C | EQV | EQV | LS | H | LS | G | LS | C | C | H | C | H | T | C | LS | LS | LS | T | H | T | EQV | LS | T | A | T | A | H | H | A | G | H | A | H | G | G | T | T | H | LS | LS | C | G | A | C | G |
| 2013_P1-17-1 | PHI1 | PHI2 | progeny | 2013_P1-17 | 2013-P1-17-1 | A | C | G | T | A | T | A | C | C | T | G | H | H | H | T | C | C | H | C | H | T | C | T | T | G | T | C | C | A | A | T | G | T | A | H | H | C | G | G | A | G | G | G | T | T | G | A | H | C | G | A | H | G |
| 2013_P1-17-2 | PHI1 | PHI2 | progeny | 2013_P1-17 | 2013-P1-17-2 | A | C | G | C | A | T | A | C | EQV | EQV | G | H | C | EQV | LS | C | C | H | C | H | T | C | LS | LS | EQV | T | G | T | G | LS | T | G | T | A | T | A | C | G | G | A | G | G | G | T | T | H | A | LS | C | G | A | A | G |
| 2013_P1-19 | PHI1 | PHI2 | mother | 2013_P1-19 | 2013-P1-19 | A | C | G | H | A | T | A | C | C | H | G | H | LS | G | LS | C | C | H | C | H | T | C | EQV | T | EQV | T | H | H | EQV | A | T | H | T | A | H | H | H | G | H | A | H | G | G | T | T | H | A | EQV | C | G | A | H | G |
| 2013_P1-19-1 | PHI1 | PHI2 | progeny | 2013_P1-19 | 2013-P1-19-1 | A | C | G | C | A | T | A | C | C | C | G | C | C | H | T | C | C | C | C | G | T | C | H | T | G | T | H | H | H | A | T | A | T | A | C | C | A | G | A | A | A | G | G | T | T | A | A | T | C | G | A | H | G |
| 2013_P1-19-2 | PHI1 | PHI2 | progeny | 2013_P1-19 | 2013-P1-19-2 | A | C | G | H | A | T | A | C | C | H | G | H | H | H | T | C | C | H | C | H | T | C | H | T | G | T | G | T | G | A | T | A | T | A | C | C | A | G | A | A | A | G | G | T | T | H | A | H | C | G | A | H | G |
| 2013_P1-19-3 | PHI1 | PHI2 | progeny | 2013_P1-19 | 2013-P1-19-3 | A | C | G | H | A | T | A | C | C | T | G | T | T | G | T | C | C | H | C | H | T | C | H | T | G | T | H | H | H | A | T | H | T | A | H | H | H | G | H | A | H | G | G | T | T | G | A | G | C | G | A | H | G |
| 2013_P1-19-4 | PHI1 | PHI2 | progeny | 2013_P1-19 | 2013-P1-19-4 | A | C | G | H | A | T | A | C | C | H | G | H | H | H | T | C | C | H | C | H | T | C | H | T | G | T | H | H | H | A | T | A | T | A | C | C | A | G | A | A | A | G | G | T | T | H | A | H | C | G | A | H | G |
| 2013_P1-19-5 | PHI1 | PHI2 | progeny | 2013_P1-19 | 2013-P1-19-5 | A | C | G | T | A | T | A | C | C | H | G | H | H | H | T | C | C | H | C | H | T | C | G | T | G | T | H | H | H | A | T | A | T | A | C | C | A | G | A | A | H | G | G | T | T | H | A | H | C | G | A | H | G |
| 2013_P1-20 | PHI1 | PHI2 | mother | 2013_P1-20 | 2013-P1-20 | A | C | G | H | A | T | A | C | C | T | G | H | T | H | EQV | C | C | H | C | H | T | C | EQV | T | G | T | H | EQV | LS | LS | T | H | T | A | H | H | A | G | H | A | H | G | G | T | T | H | A | EQV | C | G | A | H | G |
| 2013_P1-20-1 | PHI1 | PHI2 | progeny | 2013_P1-20 | 2013-P1-20-1 | A | C | G | C | A | T | A | C | C | C | G | C | C | T | T | C | C | H | C | H | T | C | G | T | G | T | H | H | H | A | T | H | T | A | H | H | A | G | A | A | A | G | G | T | T | H | A | G | C | G | A | C | G |
| 2013_P1-20-2 | PHI1 | PHI2 | progeny | 2013_P1-20 | 2013-P1-20-2 | A | C | G | T | A | T | A | C | C | H | G | H | H | H | T | C | C | T | C | A | T | C | H | T | G | T | H | H | H | A | T | H | T | A | T | A | C | G | G | A | H | G | G | T | T | G | A | G | C | G | A | H | G |
| 2013_P1-21 | PHI1 | PHI2 | mother | 2013_P1-21 | 2013-P1-21 | LS | C | G | H | A | T | A | C | C | T | LS | H | T | G | EQV | C | C | H | C | H | T | C | EQV | LS | G | T | H | H | LS | A | T | H | T | A | H | H | H | G | H | A | H | G | G | T | T | H | A | LS | C | G | A | A | G |
| 2013_P1-21-1 | PHI1 | PHI2 | progeny | 2013_P1-21 | 2013-P1-21-1 | A | C | G | H | A | T | A | C | C | T | G | T | T | G | T | C | C | H | C | H | T | C | EQV | T | G | T | H | H | H | A | T | H | T | A | H | H | H | G | H | A | H | G | G | T | T | H | A | H | C | G | A | H | G |
| 2013_P1-21-2 | PHI1 | PHI2 | progeny | 2013_P1-21 | 2013-P1-21-2 | A | C | G | LS | A | T | A | C | C | H | LS | H | H | H | T | C | C | C | C | G | T | C | H | T | G | T | H | C | A | A | T | G | T | A | T | A | C | G | G | A | G | G | G | T | T | H | EQV | EQV | C | G | A | EQV | G |
| 2013_P1-21-3 | PHI1 | PHI2 | progeny | 2013_P1-21 | 2013-P1-21-3 | A | C | G | C | A | T | A | C | C | C | G | C | C | T | T | C | C | H | C | H | T | C | H | T | G | T | C | C | A | A | T | G | T | A | T | A | C | G | G | A | G | G | G | T | T | H | A | H | C | G | A | H | G |
| 2013_P1-21-4 | PHI1 | PHI2 | progeny | 2013_P1-21 | 2013-P1-21-4 | A | C | G | T | A | T | A | C | C | H | G | H | H | H | T | C | C | H | C | H | T | C | H | T | G | T | H | H | H | A | T | G | T | A | T | A | C | G | G | A | G | G | G | T | T | G | A | G | C | G | A | C | G |
| 2013_P1-22 | PHI1 | PHI2 | mother | 2013_P1-22 | 2013-P1-22 | A | C | G | H | A | T | A | C | C | EQV | G | H | H | EQV | EQV | C | C | H | C | H | T | C | T | T | G | T | H | H | EQV | A | T | H | T | A | H | H | H | G | H | A | H | G | G | T | T | H | A | H | C | G | A | H | G |
| 2013_P1-22-1 | PHI1 | PHI2 | progeny | 2013_P1-22 | 2013-P1-22-1 | H | C | G | H | A | H | A | C | C | H | G | C | C | H | T | C | H | H | H | H | H | H | H | T | G | T | H | H | H | A | T | A | H | A | H | H | H | G | EQV | H | H | G | G | T | T | A | EQV | EQV | H | G | A | EQV | H |
| 2013_P1-22-2 | PHI1 | PHI2 | progeny | 2013_P1-22 | 2013-P1-22-2 | A | C | G | C | A | T | A | C | C | C | G | C | C | T | T | C | C | C | C | G | T | C | H | T | G | T | H | H | H | A | T | G | T | A | T | A | C | G | G | A | G | G | G | T | T | H | A | H | C | G | A | H | G |
| 2013_P1-25 | PHI1 | PHI2 | mother | 2013_P1-25 | 2013-P1-25 | A | C | G | H | A | T | A | C | C | H | G | H | H | H | EQV | C | C | H | C | H | T | C | EQV | T | G | T | H | H | H | A | T | H | T | A | H | H | H | G | H | A | H | G | G | T | T | H | A | EQV | C | G | A | H | G |
| 2013_P1-25-1 | PHI1 | PHI2 | progeny | 2013_P1-25 | 2013-P1-25-1 | A | C | G | H | A | T | A | C | C | H | G | H | H | H | T | C | C | T | C | A | T | C | H | T | G | T | H | H | H | A | T | H | T | A | H | H | H | G | H | A | H | G | G | T | T | H | A | H | C | G | A | H | G |
| 2013_P1-25-2 | PHI1 | PHI2 | progeny | 2013_P1-25 | 2013-P1-25-2 | EQV | C | G | EQV | A | T | A | C | C | H | LS | H | C | EQV | T | C | C | H | C | H | T | C | LS | LS | G | T | H | T | EQV | A | T | H | T | A | T | A | C | G | G | A | G | G | G | T | T | A | A | EQV | C | G | A | H | G |
| 2013_P1-25-3 | PHI1 | PHI2 | progeny | 2013_P1-25 | 2013-P1-25-3 | A | C | G | H | A | T | A | C | C | C | G | C | C | H | T | C | C | H | C | H | T | C | H | T | G | T | H | H | EQV | A | T | H | T | A | H | H | H | G | H | A | H | G | G | T | T | A | A | T | C | G | A | H | G |
| 2013_P1-25-4 | PHI1 | PHI2 | progeny | 2013_P1-25 | 2013-P1-25-4 | A | C | G | H | A | T | A | C | C | H | G | H | C | T | T | C | C | C | C | G | T | C | G | T | G | T | H | H | H | A | T | H | T | A | H | H | H | G | H | A | H | G | G | T | T | H | A | H | EQV | G | A | A | G |
| 2013_P1-25-5 | PHI1 | PHI2 | progeny | 2013_P1-25 | 2013-P1-25-5 | A | C | G | H | A | T | A | C | C | H | G | H | H | T | LS | C | C | H | C | H | T | C | T | LS | G | T | H | H | A | A | T | H | T | A | H | H | H | G | H | A | H | G | G | T | T | H | A | T | C | G | A | H | G |
| 2013_P1-25-6 | PHI1 | PHI2 | progeny | 2013_P1-25 | 2013-P1-25-6 | A | C | G | LS | A | T | A | C | C | T | G | T | T | G | EQV | C | C | T | C | G | T | C | EQV | T | G | T | H | H | EQV | A | T | H | T | A | H | H | H | G | H | A | H | G | G | T | T | H | A | H | C | G | A | C | G |
| 2013_P1-26 | PHI1 | PHI2 | mother | 2013_P1-26 | 2013-P1-26 | EQV | C | G | H | A | T | A | C | C | H | EQV | H | T | H | LS | C | C | H | C | EQV | T | C | T | T | LS | T | H | H | LS | A | T | EQV | T | EQV | H | H | C | G | H | A | H | G | G | T | T | H | A | H | C | G | A | A | G |
| 2013_P1-26-1 | PHI1 | PHI2 | progeny | 2013_P1-26 | 2013-P1-26-1 | A | C | G | T | A | T | A | C | C | T | G | T | T | H | T | C | C | H | C | H | T | C | H | T | G | T | G | T | G | A | T | A | T | A | C | C | A | G | A | A | A | G | G | T | T | H | A | H | C | G | A | H | G |
| 2013_P1-27 | PHI1 | PHI2 | mother | 2013_P1-27 | 2013-P1-27 | A | C | G | H | A | T | A | C | H | H | G | T | EQV | H | EQV | C | C | H | C | H | T | C | EQV | T | G | T | H | H | EQV | A | T | H | T | A | H | H | A | G | EQV | A | H | G | G | T | T | H | A | EQV | C | G | A | H | G |
| 2013_P1-27-1 | PHI1 | PHI2 | progeny | 2013_P1-27 | 2013-P1-27-1 | A | C | G | T | A | T | A | C | H | H | G | T | H | H | T | C | C | T | C | A | T | C | H | T | G | T | H | H | H | A | T | H | T | A | H | H | A | G | EQV | A | H | G | G | T | T | H | EQV | EQV | C | G | A | C | G |
| 2013_P1-28 | PHI1 | PHI2 | mother | 2013_P1-28 | 2013-P1-28 | A | C | G | H | A | T | A | C | C | EQV | G | H | H | H | EQV | C | C | H | C | H | T | C | EQV | T | G | T | H | H | EQV | A | T | H | T | A | H | H | H | G | H | A | H | G | G | T | T | H | A | EQV | C | G | A | EQV | G |
| 2013_P1-28-1 | PHI1 | PHI2 | progeny | 2013_P1-28 | 2013-P1-28-1 | A | C | G | H | A | T | A | C | C | H | G | H | H | H | T | C | C | H | C | H | T | C | H | T | G | T | H | H | H | A | T | A | T | A | C | C | A | G | A | A | A | G | G | T | T | G | A | G | C | G | A | C | G |
| 2013_P1-31 | PHI1 | PHI2 | mother | 2013_P1-31 | 2013-P1-31 | EQV | C | G | H | A | T | A | C | C | H | LS | H | EQV | H | EQV | C | C | H | C | H | T | C | EQV | T | G | T | H | H | EQV | A | T | H | T | A | H | H | H | G | H | A | H | G | G | T | T | H | A | T | C | G | A | H | G |
| 2013_P1-31-1 | PHI1 | PHI2 | progeny | 2013_P1-31 | 2013-P1-31-1 | A | C | G | EQV | A | T | A | C | C | H | G | H | H | H | T | C | C | C | C | G | T | C | EQV | T | G | T | H | H | H | A | T | G | T | A | T | A | C | G | G | A | G | G | G | T | T | A | A | T | EQV | G | A | H | G |
| 2013_P1-31-2 | PHI1 | PHI2 | progeny | 2013_P1-31 | 2013-P1-31-2 | A | C | G | C | A | T | A | C | C | H | G | H | T | H | T | C | C | T | C | A | T | C | H | T | G | T | H | H | H | A | T | A | T | A | C | C | A | G | A | A | A | G | G | T | T | H | A | EQV | EQV | G | A | H | G |
| 2013_O4 | PHI2 | PHI2 | ancestor | 2013_O4 | 2013-O4 | A | C | G | T | A | T | A | C | C | T | G | T | T | G | C | LS | C | C | C | G | T | C | T | T | G | T | C | C | A | A | T | G | T | A | T | A | C | G | G | A | G | G | G | T | T | G | A | G | C | G | A | C | G |
| 2013_O1 | PHI1 | PHI1 | ancestor | 2013_O1 | 2013-O1 | A | C | G | C | A | T | A | C | T | C | G | T | C | T | T | A | C | C | C | G | T | C | G | T | G | T | G | T | G | A | T | A | T | A | C | C | A | G | A | A | A | G | G | T | T | A | A | T | C | G | A | A | G |
| 2013_O1_O4 | PHI1 | PHI2 | ancestor | 2013_O1_O4 | 2013-O1_O4 | A | C | G | H | A | T | A | C | H | H | G | T | H | H | LS | LS | C | C | C | G | T | C | EQV | T | LS | T | H | H | A | A | T | H | T | A | H | H | A | G | H | A | H | G | G | T | T | H | A | EQV | C | G | A | LS | G |

| | 9 | 9 | 9 | 9 | 9 | 9 | 9 | 9 | 9 | 9 | 9 | 9 | 9 | 9 | 9 | 9 | 9 | 9 | 9 | 9 | 10 | 10 | 10 | 10 | 10 | 10 | 10 | 10 | 10 | 10 | 10 | 10 | 10 | 10 | 10 | 10 | 10 | 10 | 10 | 10 | 10 | 10 | 10 | 10 | 10 | 10 | 10 | 10 | 10 | 10 | 10 | |
|---|---|---|---|---|---|---|---|---|---|---|---|---|---|---|---|---|---|---|---|---|---|---|---|---|---|---|---|---|---|---|---|---|---|---|---|---|---|---|---|---|---|---|---|---|---|---|---|---|---|---|---|---|
| | 18 | 41 | 42 | 62 | 62 | 76 | 76 | 76 | 76 | 76 | 90 | 92 | 94 | 99 | 100 | 143 | 144 | 179 | 179 | 180 | 8 | 8 | 9.4 | 9.5 | 10 | 25 | 28 | 30 | 36 | 40 | 40 | 45 | 46 | 48 | 56 | 61 | 61 | 61 | 69 | 71 | 72 | 79 | 81 | 103 | 106 | 121 | 142 | 143 | 150 | 161 | 161 | |
| plant_id | | | | | | | | | | | | | | | | | | | | | | | | | | | | | | | | | | | | | | | | | | | | | | | | | | | | |
| 2013_P1-1 | EQV | G | C | H | A | H | T | T | H | C | H | H | C | EQV | H | LS | LS | C | G | T | EQV | EQV | C | A | G | T | H | EQV | H | T | EQV | T | T | LS | LS | EQV | H | T | G | EQV | H | A | T | T | H | H | LS | LS | G | T | G | G |
| 2013_P1-1-1 | H | G | C | H | A | H | T | T | H | H | H | H | C | H | H | A | T | C | G | T | T | A | C | A | G | T | H | H | H | T | H | T | T | H | H | H | H | T | G | H | H | A | T | T | H | H | A | G | H | T | G | G |
| 2013_P1-3 | EQV | LS | C | H | A | EQV | T | T | T | C | G | G | C | H | H | A | T | C | G | T | EQV | EQV | C | A | EQV | T | H | LS | T | H | A | T | T | EQV | EQV | H | H | T | G | C | H | A | H | T | H | H | LS | LS | H | T | G | G |
| 2013_P1-3-1 | H | H | C | C | H | H | EQV | H | T | C | G | G | C | T | G | A | T | H | G | H | EQV | T | C | A | G | T | EQV | A | H | T | A | H | T | C | C | H | H | T | G | C | C | A | H | H | EQV | H | A | G | H | T | H | G |
| 2013_P1-3-2 | C | EQV | C | H | H | G | EQV | H | T | C | G | G | C | H | H | A | T | H | G | H | EQV | T | C | A | G | T | EQV | A | H | T | A | H | T | H | H | C | C | T | G | C | H | A | H | H | EQV | H | A | LS | H | T | H | G |
| 2013_P1-3-3 | H | EQV | C | H | H | G | EQV | H | T | C | LS | G | C | H | H | A | T | H | G | H | EQV | EQV | C | A | LS | T | H | EQV | H | H | A | EQV | T | LS | H | C | C | T | G | C | H | EQV | H | H | H | T | LS | EQV | H | T | H | G |
| 2013_P1-3-4 | H | EQV | C | H | H | H | EQV | H | T | C | G | G | C | H | H | LS | T | H | G | H | EQV | EQV | C | A | G | T | H | T | H | H | A | H | T | EQV | H | C | EQV | T | G | C | H | A | H | H | H | T | LS | LS | H | T | H | G |
| 2013_P1-3-5 | H | LS | C | H | H | G | EQV | H | T | C | LS | G | C | H | EQV | LS | EQV | H | G | H | EQV | EQV | LS | A | EQV | T | C | LS | H | H | A | H | T | EQV | EQV | C | C | T | G | C | H | A | H | H | EQV | T | LS | LS | H | T | H | G |
| 2013_P1-6 | LS | LS | LS | H | A | T | T | T | EQV | C | EQV | EQV | C | EQV | H | LS | EQV | EQV | G | T | EQV | EQV | C | A | H | T | H | EQV | T | H | A | T | T | LS | C | EQV | EQV | LS | G | C | H | A | H | T | EQV | H | LS | LS | H | T | G | G |
| 2013_P1-6-1 | H | G | C | A | A | G | T | T | T | C | G | H | C | H | H | A | T | C | G | T | T | A | C | A | G | T | C | T | T | H | A | T | T | C | C | H | H | T | G | C | H | A | H | T | H | H | A | G | H | T | G | G |
| 2013_P1-7 | H | G | C | H | A | EQV | T | T | T | C | G | G | C | H | H | A | T | C | G | T | EQV | LS | C | A | EQV | T | H | EQV | T | H | A | T | T | C | C | H | H | T | G | C | H | A | H | T | EQV | H | LS | G | H | T | G | G |
| 2013_P1-7-1 | H | G | C | H | A | EQV | T | T | T | C | G | G | C | EQV | H | A | T | C | G | T | H | EQV | C | A | H | T | H | A | H | H | A | H | T | C | C | H | H | T | G | C | H | A | H | T | H | H | A | LS | H | T | G | G |
| 2013_P1-8 | H | G | C | H | A | H | T | T | T | C | G | G | C | H | H | A | T | C | G | T | EQV | EQV | C | A | G | T | H | LS | H | T | EQV | T | T | C | C | H | H | T | G | C | H | A | H | T | H | H | EQV | LS | H | T | G | G |
| 2013_P1-8-1 | C | H | C | H | H | G | EQV | H | T | C | G | G | C | H | H | A | LS | H | G | H | EQV | H | C | A | G | T | H | LS | H | T | H | H | T | C | C | H | H | T | G | C | EQV | A | H | H | H | H | A | G | H | T | H | G |
| 2013_P1-8-2 | G | G | C | H | A | G | T | T | T | C | G | G | C | C | A | A | T | C | G | T | T | A | C | A | G | T | H | T | T | T | A | T | T | C | C | A | A | T | G | C | C | A | T | T | H | H | LS | G | H | T | G | G |
| 2013_P1-8-3 | H | EQV | C | H | H | G | EQV | H | T | C | G | G | C | H | H | A | T | H | G | H | H | EQV | C | A | G | T | H | LS | H | T | H | H | T | C | C | C | C | T | G | C | H | A | H | H | H | H | A | G | H | T | H | G |
| 2013_P1-8-4 | H | G | C | H | A | H | T | T | T | C | G | G | C | H | H | A | T | C | G | T | T | A | C | A | G | T | H | T | H | T | H | T | T | C | C | H | H | T | G | C | H | A | H | T | H | H | A | EQV | H | T | G | G |
| 2013_P1-8-5 | H | H | C | H | H | H | EQV | H | T | C | G | G | C | EQV | EQV | A | T | H | G | H | H | T | C | A | G | T | H | EQV | H | T | H | H | T | C | C | H | EQV | T | G | C | EQV | A | H | H | H | H | A | G | H | T | H | G |
| 2013_P1-9 | EQV | G | C | H | A | EQV | T | T | EQV | EQV | LS | H | C | H | H | A | EQV | C | G | T | EQV | EQV | C | A | H | T | H | EQV | T | H | A | T | T | C | LS | H | EQV | T | G | C | H | A | H | T | H | H | LS | LS | H | T | G | G |
| 2013_P1-9-1 | H | G | C | H | A | T | T | T | C | T | A | C | C | T | G | A | T | C | G | T | H | T | C | A | H | T | C | EQV | T | H | A | T | T | C | EQV | A | A | T | G | C | H | A | H | T | A | T | A | G | T | T | G | G |
| 2013_P1-9-2 | H | G | C | C | A | T | T | T | C | T | H | H | C | H | H | A | T | C | G | T | EQV | EQV | C | A | H | T | H | T | T | H | A | T | T | C | C | C | C | T | G | C | T | LS | C | T | H | EQV | A | G | H | T | G | G |
| 2013_P1-9-3 | H | G | C | H | A | H | T | T | T | C | H | H | C | H | H | A | T | C | G | T | H | EQV | C | A | H | T | H | H | T | H | A | T | T | C | C | A | A | T | G | C | H | A | T | T | A | T | A | G | H | T | G | G |
| 2013_P1-10 | H | G | C | H | A | G | T | T | EQV | H | H | EQV | C | H | H | A | T | C | G | T | T | EQV | C | A | G | T | H | EQV | H | T | G | T | T | H | H | EQV | H | T | G | H | H | A | T | T | H | H | EQV | EQV | H | T | G | G |
| 2013_P1-10-1 | C | G | C | H | A | H | T | T | H | H | H | H | C | H | H | A | T | C | G | T | T | A | C | A | G | T | H | H | H | T | H | T | T | A | EQV | C | C | T | G | T | T | A | T | T | C | C | A | G | H | T | G | G |
| 2013_P1-16 | H | G | C | H | A | H | T | T | T | C | H | H | C | H | H | A | T | C | G | T | EQV | EQV | C | A | H | T | H | EQV | T | H | H | T | T | H | H | H | H | T | G | C | H | A | H | T | H | H | A | EQV | H | T | G | G |
| 2013_P1-16-1 | C | G | C | A | A | G | T | T | T | C | G | G | C | C | A | A | T | C | G | T | H | EQV | C | A | H | T | H | EQV | T | H | T | T | H | H | H | H | T | G | C | H | A | C | T | C | H | H | A | G | H | T | G | G |
| 2013_P1-16-2 | C | G | C | A | A | G | T | T | T | C | H | H | C | H | H | A | T | C | G | T | T | EQV | C | A | G | T | C | T | T | T | G | T | T | EQV | EQV | C | C | T | G | C | H | A | H | T | H | H | A | G | H | T | G | G |
| 2013_P1-17 | EQV | G | C | H | A | H | T | T | T | C | LS | G | C | H | H | LS | T | C | G | T | EQV | EQV | C | A | G | T | H | LS | T | H | EQV | T | T | C | EQV | EQV | H | T | G | C | H | A | H | T | H | H | LS | LS | H | T | G | G |
| 2013_P1-17-1 | EQV | G | C | A | A | G | T | T | T | C | G | G | C | H | H | A | T | C | G | T | H | H | C | A | G | T | H | EQV | H | H | A | H | T | C | C | H | H | T | G | C | H | A | H | H | H | H | A | G | H | T | G | G |
| 2013_P1-17-2 | H | G | C | H | A | G | T | T | EQV | C | G | G | C | H | H | LS | T | C | G | T | T | LS | C | A | G | T | H | T | T | C | A | T | T | C | LS | C | C | T | G | C | T | A | C | T | C | C | A | G | H | T | G | G |
| 2013_P1-19 | EQV | G | C | H | A | H | T | T | H | H | H | EQV | C | H | H | A | T | C | G | T | T | EQV | C | A | H | T | H | T | T | H | A | T | T | EQV | EQV | H | H | T | G | C | H | A | H | T | H | H | A | G | H | T | G | G |
| 2013_P1-19-1 | H | G | C | A | A | G | T | T | T | C | G | G | C | C | A | A | LS | C | G | T | T | H | C | A | H | T | H | A | T | C | A | T | T | C | C | C | C | T | G | C | T | A | C | T | C | C | A | G | G | T | G | G |
| 2013_P1-19-2 | H | G | C | H | A | H | T | T | H | H | H | H | C | H | H | A | EQV | C | G | T | H | H | C | A | H | T | H | EQV | T | H | A | T | T | C | C | H | H | T | G | C | H | A | H | T | H | H | A | G | H | T | G | G |
| 2013_P1-19-3 | H | G | C | H | A | H | T | T | H | H | H | H | C | H | H | A | EQV | C | G | T | C | T | C | A | A | T | T | A | T | C | A | T | T | C | C | H | H | T | G | C | H | A | H | T | H | H | A | G | H | T | G | G |
| 2013_P1-19-4 | G | G | C | H | A | G | T | T | T | C | G | G | C | C | A | A | T | C | G | T | H | H | C | A | H | T | H | H | T | H | A | T | T | C | C | H | H | T | G | C | H | A | H | T | H | C | A | G | G | T | G | G |
| 2013_P1-19-5 | H | G | C | C | A | T | T | T | C | T | A | C | C | T | G | A | T | C | G | T | H | H | C | A | H | T | H | H | T | H | A | T | T | C | C | A | A | T | G | C | C | A | H | T | H | H | A | G | H | T | G | G |
| 2013_P1-20 | H | G | C | H | A | H | T | T | EQV | H | H | H | C | H | H | A | T | C | G | T | T | A | C | A | G | T | H | EQV | H | T | H | T | T | C | C | H | H | T | G | C | H | A | EQV | T | H | H | A | G | H | T | G | G |
| 2013_P1-20-1 | G | G | C | H | A | H | T | T | T | C | G | G | C | C | A | A | LS | C | G | T | T | A | C | A | G | T | C | T | T | T | A | T | T | C | C | A | A | T | G | C | C | A | T | T | A | T | A | G | H | T | G | G |
| 2013_P1-20-2 | H | G | C | H | A | T | T | T | C | T | H | H | C | H | H | A | T | C | G | T | T | A | C | A | G | T | T | A | C | T | G | T | T | C | C | C | C | T | G | C | T | A | C | T | C | C | A | G | H | T | G | G |
| 2013_P1-21 | EQV | G | C | H | A | EQV | T | T | H | H | G | H | C | H | H | A | T | C | G | T | T | A | C | A | G | T | H | EQV | H | T | EQV | T | T | EQV | LS | H | H | T | G | EQV | H | A | T | T | H | H | A | LS | H | T | G | G |
| 2013_P1-21-1 | C | G | C | H | A | G | T | T | T | C | H | H | C | C | A | A | T | C | G | T | T | A | C | A | G | T | H | H | H | T | H | T | T | H | C | A | A | T | G | C | C | A | T | T | H | H | A | G | H | T | G | G |
| 2013_P1-21-2 | H | G | C | C | A | T | T | T | C | T | H | H | C | H | H | A | LS | C | G | T | T | A | C | A | G | T | H | A | T | T | A | T | T | C | C | A | A | T | G | C | C | A | T | T | H | H | A | G | H | T | G | G |
| 2013_P1-21-3 | G | G | C | C | A | T | T | T | C | T | A | C | C | T | G | A | T | C | G | T | T | A | C | A | G | T | T | A | C | T | G | T | T | A | G | C | C | T | G | H | H | A | T | T | H | H | A | G | H | T | G | G |
| 2013_P1-21-4 | C | G | C | A | A | G | T | T | T | C | H | H | C | H | H | A | T | C | G | T | T | A | C | A | G | T | H | H | H | T | H | T | T | H | C | A | A | T | G | C | C | A | T | T | H | H | A | G | H | T | G | G |
| 2013_P1-22 | EQV | G | C | H | A | H | T | T | H | H | H | H | C | H | H | A | T | C | G | T | T | A | C | A | G | T | H | EQV | H | T | H | T | T | H | C | H | H | T | G | C | H | A | T | T | H | H | A | G | H | T | G | G |
| 2013_P1-22-1 | H | H | C | H | H | G | EQV | H | T | C | G | G | C | H | H | A | LS | EQV | G | EQV | H | H | C | A | G | T | H | A | H | T | H | H | T | EQV | H | C | C | T | G | C | H | A | H | H | H | H | A | G | H | T | H | G |
| 2013_P1-22-2 | H | G | C | A | A | G | T | T | T | C | G | G | C | C | A | A | T | C | G | T | T | A | C | A | G | T | H | EQV | H | T | H | T | T | H | C | A | A | T | G | C | C | A | T | T | A | T | A | G | T | T | G | G |
| 2013_P1-25 | H | G | C | H | A | H | T | T | H | H | G | G | C | H | H | A | T | C | G | T | H | H | C | A | H | T | H | H | T | H | A | T | T | C | C | H | H | T | G | C | H | A | H | T | H | H | A | EQV | H | T | G | G |
| 2013_P1-25-1 | H | G | C | H | A | H | T | T | H | H | G | G | C | H | H | A | T | C | G | T | H | H | C | A | H | T | H | EQV | T | H | A | T | T | C | C | H | H | T | G | C | H | A | H | T | H | H | A | G | H | T | G | G |
| 2013_P1-25-2 | G | G | C | H | A | T | T | T | C | T | G | G | C | T | G | A | T | C | G | T | H | A | C | A | H | T | H | T | T | T | A | T | T | C | C | A | A | T | G | C | C | A | T | T | H | H | A | G | H | T | G | G |
| 2013_P1-25-3 | H | G | C | H | A | H | T | T | H | H | G | G | C | H | H | A | T | C | G | T | H | H | C | A | H | T | T | A | T | C | A | T | T | C | C | C | C | T | G | C | T | A | C | T | H | H | A | G | T | T | G | G |
| 2013_P1-25-4 | G | G | C | H | A | H | T | T | H | H | G | G | C | H | H | A | T | C | G | T | C | EQV | C | A | H | T | H | A | T | H | A | T | T | C | C | H | H | T | G | C | H | A | H | T | H | H | A | G | H | T | G | G |
| 2013_P1-25-5 | H | G | C | A | A | G | T | T | T | C | G | G | C | H | H | LS | T | C | G | T | H | H | C | A | H | T | H | T | T | H | A | T | T | C | C | H | H | T | G | C | H | A | H | T | C | C | LS | G | G | T | G | G |
| 2013_P1-25-6 | H | G | C | C | A | H | T | T | H | H | G | G | C | H | H | A | H | C | G | T | H | H | C | A | H | T | H | H | T | H | A | T | T | C | C | H | H | T | G | C | H | A | H | T | H | H | A | G | H | T | G | G |
| 2013_P1-26 | H | G | C | H | A | H | T | T | T | C | G | G | C | H | H | A | EQV | C | G | T | EQV | EQV | C | A | G | T | H | EQV | H | T | H | T | T | EQV | EQV | H | EQV | T | G | EQV | H | A | T | T | H | H | LS | LS | H | T | G | G |
| 2013_P1-26-1 | H | G | C | A | A | G | T | T | T | C | G | G | C | H | H | A | T | C | G | T | T | A | C | A | G | T | H | H | H | T | G | T | T | A | G | C | C | T | G | T | T | A | T | T | H | H | A | G | H | T | G | G |
| 2013_P1-27 | C | G | C | H | A | H | T | T | H | H | H | H | C | H | H | A | T | C | G | T | T | A | C | A | G | T | H | EQV | H | T | H | T | T | H | H | H | H | T | G | EQV | H | A | T | T | H | H | A | EQV | H | T | G | G |
| 2013_P1-27-1 | H | G | C | C | A | T | T | T | C | T | A | C | C | T | G | A | T | C | G | T | T | A | C | A | G | T | H | H | T | T | A | T | T | C | C | A | A | T | G | C | C | A | T | T | H | H | A | G | G | T | G | G |
| 2013_P1-28 | H | G | C | H | A | H | T | T | T | C | G | H | C | H | H | A | EQV | C | G | T | EQV | H | C | A | H | T | H | T | T | H | A | T | T | C | C | H | H | T | G | C | H | A | H | T | H | H | EQV | G | H | T | G | G |
| 2013_P1-28-1 | G | G | C | H | A | G | T | T | T | C | G | G | C | C | A | A | T | C | G | T | C | T | C | A | A | T | H | H | T | H | A | T | T | C | C | A | A | T | G | C | C | A | T | T | A | T | A | G | H | T | G | G |
| 2013_P1-31 | H | G | C | H | A | EQV | T | T | H | EQV | G | G | C | H | H | LS | T | C | G | T | EQV | EQV | C | A | H | T | H | LS | T | H | A | T | T | C | H | H | H | T | G | H | H | A | T | T | H | H | A | EQV | H | T | G | G |
| 2013_P1-31-1 | H | G | C | H | A | H | T | T | H | H | G | G | C | H | H | A | T | C | G | T | C | T | C | A | A | T | T | A | T | H | A | T | T | C | H | H | H | T | G | H | H | A | T | T | H | H | A | G | H | T | G | G |
| 2013_P1-31-2 | G | G | C | C | A | T | T | T | C | T | G | G | C | H | H | A | T | C | G | T | H | H | C | A | H | T | H | EQV | T | H | A | T | T | C | C | A | A | T | G | H | H | A | T | T | H | H | A | G | H | T | G | G |
| 2013_O4 | C | G | C | A | A | G | T | T | T | C | G | G | C | C | A | A | LS | C | G | T | T | A | C | A | G | T | C | EQV | T | T | A | T | T | A | C | A | A | T | G | C | C | A | T | T | A | T | A | G | T | T | G | G |
| 2013_O1 | G | G | C | C | A | T | T | T | C | G | G | G | C | T | G | A | T | C | G | T | C | T | C | A | A | T | T | A | T | C | A | T | T | C | C | C | C | T | G | C | T | A | C | T | C | C | A | G | G | T | G | G |
| 2013_O1_O4 | H | G | C | H | A | H | T | T | T | C | G | G | C | H | H | A | T | C | G | T | C | LS | C | A | H | T | H | T | T | H | A | T | T | H | LS | H | H | T | G | C | H | A | H | T | EQV | H | A | LS | EQV | T | G | G |

plant_id – unique identifier of the plant
mom_gen – genetic backgroiund of mother in the hyb
dad_gen – genetic background of father in the hybre
family_ – identify mother and all the progeny plant
mother_progeny – identify each plant as mother of the famil
Chr – number of the chromosome, where mark
Distance – position of the marker on the correspond
H – heterozygous marker
A, T, G and C – marker homozygous for the correspondi
EQV, LS, -9 , etc. – missing values

### Table S3. Results of the SNP array analasis

| | | | | | Chr | 1 | (remaining marker columns) |
|---|---|---|---|---|---|---|---|
| | | | | | Position, cM | 0 | [illegible] |
| plant_id | mom_gen | dad_gen | mother_progeny | family_ | | | |
| 2015_P3-A11-1 | PHI1 | PHI3 | progeny | 2015_P3-A11 | | A | [illegible] |
| 2015_P3-A11-2 | PHI1 | PHI3 | progeny | 2015_P3-A11 | | A | [illegible] |
| 2015_P3-A11-3 | PHI1 | PHI3 | progeny | 2015_P3-A11 | | A | [illegible] |
| 2015_P3-A10-1 | PHI1 | PHI3 | progeny | 2015_P3-A10 | | A | [illegible] |
| 2015_P3-A8-1 | PHI1 | PHI3 | progeny | 2015_P3-A8 | | G | [illegible] |
| 2015_P3-A8-2 | PHI1 | PHI3 | progeny | 2015_P3-A8 | | H | [illegible] |
| 2015_P3-A8-3 | PHI1 | PHI3 | progeny | 2015_P3-A8 | | H | [illegible] |
| 2015_P3-A7-1 | PHI1 | PHI3 | progeny | 2015_P3-A7 | | H | [illegible] |
| 2015_P3-A6-1 | PHI1 | PHI3 | progeny | 2015_P3-A6 | | A | [illegible] |
| 2015_P3-A2-1 | PHI1 | PHI3 | progeny | 2015_P3-A2 | | A | [illegible] |
| 2015_P3-A2-2 | PHI1 | PHI3 | progeny | 2015_P3-A2 | | A | [illegible] |
| 2015_P3-A2-3 | PHI1 | PHI3 | progeny | 2015_P3-A2 | | A | [illegible] |
| 2015_P3-A2-4 | PHI1 | PHI3 | progeny | 2015_P3-A2 | | A | [illegible] |
| 2015_P3-A1-1 | PHI1 | PHI3 | progeny | 2015_P3-A1 | | A | [illegible] |
| 2015_P3-A3-1 | PHI1 | PHI3 | progeny | 2015_P3-A3 | | A | [illegible] |
| 2015_P3-B11-1 | PHI1 | PHI3 | progeny | 2015_P3-B11 | | A | [illegible] |
| 2015_P3-C1-1 | PHI1 | PHI3 | progeny | 2015_P3-C1 | | A | [illegible] |
| 2015_P3-C2-1 | PHI1 | PHI3 | progeny | 2015_P3-C2 | | G | [illegible] |
| 2015_P3-C8-1 | PHI1 | PHI3 | progeny | 2015_P3-C8 | | H | [illegible] |
| 2015_P3-C8-2 | PHI1 | PHI3 | progeny | 2015_P3-C8 | | A | [illegible] |
| 2015_P3-C8-3 | PHI1 | PHI3 | progeny | 2015_P3-C8 | | A | [illegible] |
| 2015_P3-C7-1 | PHI1 | PHI3 | progeny | 2015_P3-C7 | | A | [illegible] |
| 2015_P3-C10-2 | PHI1 | PHI3 | progeny | 2015_P3-C10 | | A | [illegible] |
| 2015_P3-C11-3 | PHI1 | PHI3 | progeny | 2015_P3-C11 | | H | [illegible] |
| 2015_P3-D3-1 | PHI1 | PHI3 | progeny | 2015_P3-D3 | | H | [illegible] |
| 2015_P3-D3-2 | PHI1 | PHI3 | progeny | 2015_P3-D3 | | H | [illegible] |
| 2015_P3-D4-1 | PHI1 | PHI3 | progeny | 2015_P3-D4 | | A | [illegible] |
| 2015_P3-D4-2 | PHI1 | PHI3 | progeny | 2015_P3-D4 | | A | [illegible] |
| 2015_P3-D4-3 | PHI1 | PHI3 | progeny | 2015_P3-D4 | | A | [illegible] |
| 2015_P3-D6-1 | PHI1 | PHI3 | progeny | 2015_P3-D6 | | A | [illegible] |
| 2015_P3-B5-1 | PHI1 | PHI3 | progeny | 2015_P3-B5 | | A | [illegible] |
| 2015_P3-B4-1 | PHI1 | PHI3 | progeny | 2015_P3-B4 | | A | [illegible] |
| 2015_P3-D5-1 | PHI1 | PHI3 | progeny | 2015_P3-D5 | | A | [illegible] |
| 2015_P3-D10-1 | PHI1 | PHI3 | progeny | 2015_P3-D10 | | A | [illegible] |
| 2015_P3-E4-1 | PHI1 | PHI3 | progeny | 2015_P3-E4 | | A | [illegible] |
| 2015_P3-E4-2 | PHI1 | PHI3 | progeny | 2015_P3-E4 | | A | [illegible] |
| 2015_P3-E5-1 | PHI1 | PHI3 | progeny | 2015_P3-E5 | | A | [illegible] |
| 2015_P3-E5-2 | PHI1 | PHI3 | progeny | 2015_P3-E5 | | A | [illegible] |
| 2015_P3-E5-3 | PHI1 | PHI3 | progeny | 2015_P3-E5 | | A | [illegible] |
| 2015_P3-E7-1 | PHI1 | PHI3 | progeny | 2015_P3-E7 | | A | [illegible] |
| 2015_P3-E9-1 | PHI1 | PHI3 | progeny | 2015_P3-E9 | | A | [illegible] |
| 2015_P3-E9-2 | PHI1 | PHI3 | progeny | 2015_P3-E9 | | A | [illegible] |
| 2015_P3-E9-3 | PHI1 | PHI3 | progeny | 2015_P3-E9 | | A | [illegible] |
| 2015_P3-E9-4 | PHI1 | PHI3 | progeny | 2015_P3-E9 | | A | [illegible] |
| 2015_P3-E8-1 | PHI1 | PHI3 | progeny | 2015_P3-E8 | | A | [illegible] |
| 2015_P3-E8-2 | PHI1 | PHI3 | progeny | 2015_P3-E8 | | A | [illegible] |
| 2015_P3-E8-3 | PHI1 | PHI3 | progeny | 2015_P3-E8 | | A | [illegible] |
| 2015_P3-E10-1 | PHI1 | PHI3 | progeny | 2015_P3-E10 | | G | [illegible] |
| 2015_P3-E11-1 | PHI1 | PHI3 | progeny | 2015_P3-E11 | | A | [illegible] |
| 2015_P3-E12-1 | PHI1 | PHI3 | progeny | 2015_P3-E12 | | A | [illegible] |
| 2015_P3-F1-1 | PHI1 | PHI3 | progeny | 2015_P3-F1 | | A | [illegible] |
| 2015_P3-F2-1 | PHI1 | PHI3 | progeny | 2015_P3-F2 | | A | [illegible] |
| 2015_P3-F4-1 | PHI1 | PHI3 | progeny | 2015_P3-F4 | | A | [illegible] |
| 2015_P3-F4-2 | PHI1 | PHI3 | progeny | 2015_P3-F4 | | A | [illegible] |
| 2015_P3-F4-3 | PHI1 | PHI3 | progeny | 2015_P3-F4 | | A | [illegible] |
| 2015_P3-F4-4 | PHI1 | PHI3 | progeny | 2015_P3-F4 | | A | [illegible] |
| 2015_P3-F3-1 | PHI1 | PHI3 | progeny | 2015_P3-F3 | | A | [illegible] |
| 2015_P3-F3-2 | PHI1 | PHI3 | progeny | 2015_P3-F3 | | A | [illegible] |
| 2015_P3-B3-1 | PHI1 | PHI3 | progeny | 2015_P3-B3 | | A | [illegible] |
| 2015_P3-B7-1 | PHI1 | PHI3 | progeny | 2015_P3-B7 | | A | [illegible] |
| 2015_P3-B2-1 | PHI1 | PHI3 | progeny | 2015_P3-B2 | | A | [illegible] |
| 2015_P3-B2-2 | PHI1 | PHI3 | progeny | 2015_P3-B2 | | A | [illegible] |
| 2015_P3-D7-1 | PHI1 | PHI3 | progeny | 2015_P3-D7 | | A | [illegible] |
| 2015_P3-D7-2 | PHI1 | PHI3 | progeny | 2015_P3-D7 | | A | [illegible] |
| 2015_P4-C8-1 | PHI2 | PHI3 | progeny | 2015_P4-C8 | | H | [illegible] |
| 2015_P4-C8-2 | PHI2 | PHI3 | progeny | 2015_P4-C8 | | G | [illegible] |
| 2015_P4-B11-1 | PHI2 | PHI3 | progeny | 2015_P4-B11 | | H | [illegible] |
| 2015_P4-B11-3 | PHI2 | PHI3 | progeny | 2015_P4-B11 | | H | [illegible] |
| 2015_P4-C11-1 | PHI2 | PHI3 | progeny | 2015_P4-C11 | | H | [illegible] |
| 2015_P4-C11-2 | PHI2 | PHI3 | progeny | 2015_P4-C11 | | H | [illegible] |
| 2015_P4-E6-1 | PHI2 | PHI3 | progeny | 2015_P4-E6 | | H | [illegible] |
| 2015_P4-E6-2 | PHI2 | PHI3 | progeny | 2015_P4-E6 | | G | [illegible] |
| 2015_P4-D12-1 | PHI2 | PHI3 | progeny | 2015_P4-D12 | | G | [illegible] |
| 2015_P4-D12-2 | PHI2 | PHI3 | progeny | 2015_P4-D12 | | G | [illegible] |
| 2015_P4-E1-1 | PHI2 | PHI3 | progeny | 2015_P4-E1 | | G | [illegible] |
| 2015_P4-E1-2 | PHI2 | PHI3 | progeny | 2015_P4-E1 | | G | [illegible] |
| 2015_P4-C10-1 | PHI2 | PHI3 | progeny | 2015_P4-C10 | | H | [illegible] |
| 2015_P4-B9-1 | PHI2 | PHI3 | progeny | 2015_P4-B9 | | H | [illegible] |
| 2015_P4-B9-2 | PHI2 | PHI3 | progeny | 2015_P4-B9 | | H | [illegible] |
| 2015_P4-B7-1 | PHI2 | PHI3 | progeny | 2015_P4-B7 | | H | [illegible] |
| 2015_P4-B7-2 | PHI2 | PHI3 | progeny | 2015_P4-B7 | | H | [illegible] |
| 2015_P4-C2-1 | PHI2 | PHI3 | progeny | 2015_P4-C2 | | H | [illegible] |
| 2015_P4-E5-1 | PHI2 | PHI3 | progeny | 2015_P4-E5 | | G | [illegible] |
| 2015_P4-E5-2 | PHI2 | PHI3 | progeny | 2015_P4-E5 | | H | [illegible] |
| 2015_P4-E5-3 | PHI2 | PHI3 | progeny | 2015_P4-E5 | | H | [illegible] |
| 2015_P4-C9-1 | PHI2 | PHI3 | progeny | 2015_P4-C9 | | H | [illegible] |
| 2015_P4-C9-2 | PHI2 | PHI3 | progeny | 2015_P4-C9 | | H | [illegible] |
| 2015_P4-D4-1 | PHI2 | PHI3 | progeny | 2015_P4-D4 | | H | [illegible] |
| 2015_P4-C1-1 | PHI2 | PHI3 | progeny | 2015_P4-C1 | | G | [illegible] |
| 2015_P4-C1-2 | PHI2 | PHI3 | progeny | 2015_P4-C1 | | A | [illegible] |
| 2015_P4-D7-1 | PHI2 | PHI3 | progeny | 2015_P4-D7 | | G | [illegible] |
| 2015_P4-D3-1 | PHI2 | PHI3 | progeny | 2015_P4-D3 | | H | [illegible] |
| 2015_P4-F2-1 | PHI8 | PHI3 | progeny | 2015_P4-F2 | | H | [illegible] |
| 2015_P4-F2-2 | PHI8 | PHI3 | progeny | 2015_P4-F2 | | H | [illegible] |
| 2015_P4-F4-1 | PHI8 | PHI3 | progeny | 2015_P4-F4 | | H | [illegible] |
| 2015_P4-F4-2 | PHI8 | PHI3 | progeny | 2015_P4-F4 | | G | [illegible] |
| 2015_P4-F4-3 | PHI8 | PHI3 | progeny | 2015_P4-F4 | | H | [illegible] |
| 2015_P4-F4-4 | PHI8 | PHI3 | progeny | 2015_P4-F4 | | H | [illegible] |
| 2015_P4-E9-1 | PHI8 | PHI3 | progeny | 2015_P4-E9 | | H | [illegible] |
| 2015_P4-E9-2 | PHI8 | PHI3 | progeny | 2015_P4-E9 | | H | [illegible] |
| 2015_P4-F1-1 | PHI8 | PHI3 | progeny | 2015_P4-F1 | | A | [illegible] |
| 2015_P4-E11-1 | PHI8 | PHI3 | progeny | 2015_P4-E11 | | H | [illegible] |
| 2015_P4-E11-2 | PHI8 | PHI3 | progeny | 2015_P4-E11 | | H | [illegible] |
| 2015_P4-E10-1 | PHI8 | PHI3 | progeny | 2015_P4-E10 | | H | [illegible] |
| 2015_P4-E10-2 | PHI8 | PHI3 | progeny | 2015_P4-E10 | | H | [illegible] |
| 2015_P4-E10-3 | PHI8 | PHI3 | progeny | 2015_P4-E10 | | H | [illegible] |
| 2015_P4-E10-4 | PHI8 | PHI3 | progeny | 2015_P4-E10 | | H | [illegible] |
| 2015_P4-E8-1 | PHI8 | PHI3 | progeny | 2015_P4-E8 | | H | [illegible] |
| 2015_P4-E12-1 | PHI8 | PHI3 | progeny | 2015_P4-E12 | | H | [illegible] |
| 2015_P4-E12-2 | PHI8 | PHI3 | progeny | 2015_P4-E12 | | H | [illegible] |
| 2015_P4-F5-1 | PHI2 | PHI8 | progeny | 2015_P4-F5 | | G | [illegible] |
| 2015_P4-F5-2 | PHI2 | PHI8 | progeny | 2015_P4-F5 | | G | [illegible] |
| 2015_P4-F5-3 | PHI2 | PHI8 | progeny | 2015_P4-F5 | | G | [illegible] |
| 2015_P4-F5-4 | PHI2 | PHI8 | progeny | 2015_P4-F5 | | G | [illegible] |
| 2015_P4-F5-5 | PHI2 | PHI8 | progeny | 2015_P4-F5 | | G | [illegible] |
| 2015_P4-F5-6 | PHI2 | PHI8 | progeny | 2015_P4-F5 | | G | [illegible] |
| 2015_P4-F5-7 | PHI2 | PHI8 | progeny | 2015_P4-F5 | | G | [illegible] |
| 2015_P4-F5 | PHI2 | PHI8 | mother | 2015_P4-F5 | | G | [illegible] |
| 2015_P3-G1-1 | PHI2 | PHI1 | progeny | 2015_P3-G1 | | H | [illegible] |
| 2015_P3-G1-2 | PHI2 | PHI1 | progeny | 2015_P3-G1 | | A | [illegible] |
| 2015_P3-G1-3 | PHI2 | PHI1 | progeny | 2015_P3-G1 | | G | [illegible] |
| 2015_P3-G1-4 | PHI2 | PHI1 | progeny | 2015_P3-G1 | | H | [illegible] |
| 2015_P4-A1-1 | PHI2 | PHI1 | progeny | 2015_P4-A1 | | H | [illegible] |
| 2015_P4-A1-2 | PHI2 | PHI1 | progeny | 2015_P4-A1 | | G | [illegible] |
| 2015_P3-F11-1 | PHI2 | PHI1 | progeny | 2015_P3-F11 | | G | [illegible] |
| 2015_P3-H5-1 | PHI2 | PHI1 | progeny | 2015_P3-H5 | | G | [illegible] |
| 2015_P3-H5-2 | PHI2 | PHI1 | progeny | 2015_P3-H5 | | G | [illegible] |
| 2015_P3-H5-3 | PHI2 | PHI1 | progeny | 2015_P3-H5 | | G | [illegible] |
| 2015_P3-H5-4 | PHI2 | PHI1 | progeny | 2015_P3-H5 | | G | [illegible] |
| 2015_P3-H3-1 | PHI2 | PHI1 | progeny | 2015_P3-H3 | | G | [illegible] |
| 2015_P3-H3-2 | PHI2 | PHI1 | progeny | 2015_P3-H3 | | G | [illegible] |
| 2015_P3-H6-1 | PHI2 | PHI1 | progeny | 2015_P3-H6 | | G | [illegible] |
| 2015_P3-H7-1 | PHI2 | PHI1 | progeny | 2015_P3-H7 | | H | [illegible] |
| 2015_P4-B6-1 | PHI2 | PHI1 | progeny | 2015_P4-B6 | | G | [illegible] |
| 2015_P4-B6-2 | PHI2 | PHI1 | progeny | 2015_P4-B6 | | G | [illegible] |
| 2015_P3-G6-1 | PHI2 | PHI1 | progeny | 2015_P3-G6 | | H | [illegible] |
| 2015_P3-G6-2 | PHI2 | PHI1 | progeny | 2015_P3-G6 | | H | [illegible] |
| 2015_P3-G3-1 | PHI2 | PHI1 | progeny | 2015_P3-G3 | | A | [illegible] |
| 2015_P4-A2-1 | PHI2 | PHI1 | progeny | 2015_P4-A2 | | G | [illegible] |
| 2015_P4-A9-1 | PHI2 | PHI1 | progeny | 2015_P4-A9 | | H | [illegible] |
| 2015_P4-A9-2 | PHI2 | PHI1 | progeny | 2015_P4-A9 | | H | [illegible] |
| 2015_P4-A9-3 | PHI2 | PHI1 | progeny | 2015_P4-A9 | | H | [illegible] |
| 2015_P4-A9-4 | PHI2 | PHI1 | progeny | 2015_P4-A9 | | H | [illegible] |
| 2015_P4-B4-1 | PHI2 | PHI1 | progeny | 2015_P4-B4 | | H | [illegible] |
| 2015_P4-B4-2 | PHI2 | PHI1 | progeny | 2015_P4-B4 | | H | [illegible] |
| 2015_P3-F5-1 | PHI2 | PHI1 | progeny | 2015_P3-F5 | | A | [illegible] |
| 2015_P3-F6-1 | PHI2 | PHI1 | progeny | 2015_P3-F6 | | H | [illegible] |
| 2015_P3-F6-2 | PHI2 | PHI1 | progeny | 2015_P3-F6 | | A | [illegible] |
| 2015_P3-F6-3 | PHI2 | PHI1 | progeny | 2015_P3-F6 | | A | [illegible] |
| 2015_P4-A3-1 | PHI2 | PHI1 | progeny | 2015_P4-A3 | | H | [illegible] |
| 2015_P3-H9-1 | PHI2 | PHI1 | progeny | 2015_P3-H9 | | G | [illegible] |
| 2015_P3-H9-2 | PHI2 | PHI1 | progeny | 2015_P3-H9 | | G | [illegible] |
| 2015_P3-H9-3 | PHI2 | PHI1 | progeny | 2015_P3-H9 | | G | [illegible] |
| 2015_P3-H9-4 | PHI2 | PHI1 | progeny | 2015_P3-H9 | | G | [illegible] |
| 2015_P3-H9-5 | PHI2 | PHI1 | progeny | 2015_P3-H9 | | G | [illegible] |
| 2015_P3-H9-6 | PHI2 | PHI1 | progeny | 2015_P3-H9 | | G | [illegible] |
| 2015_P4-A4-1 | PHI2 | PHI1 | progeny | 2015_P4-A4 | | H | [illegible] |
| 2015_P4-A4-2 | PHI2 | PHI1 | progeny | 2015_P4-A4 | | H | [illegible] |
| 2015_P4-A4-3 | PHI2 | PHI1 | progeny | 2015_P4-A4 | | H | [illegible] |
| 2015_P4-A4-4 | PHI2 | PHI1 | progeny | 2015_P4-A4 | | A | [illegible] |
| 2015_P3-G10-1 | PHI2 | PHI1 | progeny | 2015_P3-G10 | | G | [illegible] |
| 2015_P3-G10-2 | PHI2 | PHI1 | progeny | 2015_P3-G10 | | H | [illegible] |
| 2015_P3-G10-3 | PHI2 | PHI1 | progeny | 2015_P3-G10 | | G | [illegible] |
| 2015_P3-G10-4 | PHI2 | PHI1 | progeny | 2015_P3-G10 | | H | [illegible] |
| 2015_P3-G10-5 | PHI2 | PHI1 | progeny | 2015_P3-G10 | | H | [illegible] |
| 2015_P3-G10-6 | PHI2 | PHI1 | progeny | 2015_P3-G10 | | H | [illegible] |
| 2015_P3-G10-7 | PHI2 | PHI1 | progeny | 2015_P3-G10 | | H | [illegible] |
| 2015_P3-G10-8 | PHI2 | PHI1 | progeny | 2015_P3-G10 | | H | [illegible] |
| 2015_P3-G10-9 | PHI2 | PHI1 | progeny | 2015_P3-G10 | | H | [illegible] |
| 2015_P3-G10-10 | PHI2 | PHI1 | progeny | 2015_P3-G10 | | H | [illegible] |
| 2015_P3-G9-1 | PHI2 | PHI1 | progeny | 2015_P3-G9 | | G | [illegible] |
| 2015_P3-G9-2 | PHI2 | PHI1 | progeny | 2015_P3-G9 | | G | [illegible] |
| 2015_P3-G9-3 | PHI2 | PHI1 | progeny | 2015_P3-G9 | | G | [illegible] |
| 2015_P3-G9-4 | PHI2 | PHI1 | progeny | 2015_P3-G9 | | G | [illegible] |
| 2015_P3-G9-5 | PHI2 | PHI1 | progeny | 2015_P3-G9 | | G | [illegible] |
| 2015_P3-G9-6 | PHI2 | PHI1 | progeny | 2015_P3-G9 | | G | [illegible] |
| 2015_P3-G9-7 | PHI2 | PHI1 | progeny | 2015_P3-G9 | | G | [illegible] |
| 2015_P3-G9-8 | PHI2 | PHI1 | progeny | 2015_P3-G9 | | G | [illegible] |
| 2015_P3-G9-9 | PHI2 | PHI1 | progeny | 2015_P3-G9 | | G | [illegible] |

**Table S3. Results of the SNP array analasis (continue)**

| | | | | | | | |
|---|---|---|---|---|---|---|---|
| | | | | | Chr | 1 | [illegible] |
| | | | | | Position, cM | 0 | [illegible] |
| plant_id | mom_gen | dad_gen | mother_progeny | family_ | | | |
| 2015_P3-G9-10 | PHI2 | PHI1 | progeny | 2015_P3-G9 | | G | [illegible] |
| 2015_P3-G9-11 | PHI2 | PHI1 | progeny | 2015_P3-G9 | | G | [illegible] |
| 2015_P3-G9-12 | PHI2 | PHI1 | progeny | 2015_P3-G9 | | G | [illegible] |
| 2015_P3-H8-1 | PHI2 | PHI1 | progeny | 2015_P3-H8 | | H | [illegible] |
| 2015_P4-A5-1 | PHI2 | PHI1 | progeny | 2015_P4-A5 | | H | [illegible] |
| 2015_P4-A5-2 | PHI2 | PHI1 | progeny | 2015_P4-A5 | | H | [illegible] |
| 2015_P4-A5-3 | PHI2 | PHI1 | progeny | 2015_P4-A5 | | H | [illegible] |
| 2015_P3-G2-1 | PHI2 | PHI1 | progeny | 2015_P3-G2 | | G | [illegible] |
| 2015_P3-F9-1 | PHI2 | PHI1 | progeny | 2015_P3-F9 | | H | [illegible] |
| 2015_P3-F9-2 | PHI2 | PHI1 | progeny | 2015_P3-F9 | | H | [illegible] |
| 2015_P3-F9-3 | PHI2 | PHI1 | progeny | 2015_P3-F9 | | H | [illegible] |
| 2015_P3-F9-4 | PHI2 | PHI1 | progeny | 2015_P3-F9 | | A | [illegible] |
| 2015_P3-F9-5 | PHI2 | PHI1 | progeny | 2015_P3-F9 | | H | [illegible] |
| 2015_P4-A7-1 | PHI2 | PHI1 | progeny | 2015_P4-A7 | | A | [illegible] |
| 2015_P3-F8-1 | PHI2 | PHI1 | progeny | 2015_P3-F8 | | G | [illegible] |
| 2015_P3-H10-1 | PHI2 | PHI1 | progeny | 2015_P3-H10 | | A | [illegible] |
| 2015_P3-H10-2 | PHI2 | PHI1 | progeny | 2015_P3-H10 | | H | [illegible] |
| 2015_P3-H10-3 | PHI2 | PHI1 | progeny | 2015_P3-H10 | | H | [illegible] |
| 2015_P3-H10-4 | PHI2 | PHI1 | progeny | 2015_P3-H10 | | H | [illegible] |
| 2015_P3-H10-5 | PHI2 | PHI1 | progeny | 2015_P3-H10 | | H | [illegible] |
| 2015_P3-H10-6 | PHI2 | PHI1 | progeny | 2015_P3-H10 | | A | [illegible] |
| 2015_P3-H10-7 | PHI2 | PHI1 | progeny | 2015_P3-H10 | | H | [illegible] |
| 2015_P3-H11-1 | PHI2 | PHI1 | progeny | 2015_P3-H11 | | H | [illegible] |
| 2015_P4-A6-1 | PHI2 | PHI1 | progeny | 2015_P4-A6 | | H | [illegible] |
| 2015_P4-A6-2 | PHI2 | PHI1 | progeny | 2015_P4-A6 | | H | [illegible] |
| 2015_P4-A6-3 | PHI2 | PHI1 | progeny | 2015_P4-A6 | | H | [illegible] |
| 2015_P4-A6-4 | PHI2 | PHI1 | progeny | 2015_P4-A6 | | H | [illegible] |
| 2015_P3-G4-1 | PHI2 | PHI1 | progeny | 2015_P3-G4 | | G | [illegible] |
| 2015_P3-G4-2 | PHI2 | PHI1 | progeny | 2015_P3-G4 | | H | [illegible] |
| 2015_P3-G4-3 | PHI2 | PHI1 | progeny | 2015_P3-G4 | | H | [illegible] |
| 2015_P3-G4-4 | PHI2 | PHI1 | progeny | 2015_P3-G4 | | H | [illegible] |
| 2015_P3-G4-5 | PHI2 | PHI1 | progeny | 2015_P3-G4 | | H | [illegible] |
| 2015_P3-G12-1 | PHI2 | PHI1 | progeny | 2015_P3-G12 | | H | [illegible] |
| 2015_P3-G12-2 | PHI2 | PHI1 | progeny | 2015_P3-G12 | | G | [illegible] |
| 2015_P3-G12-3 | PHI2 | PHI1 | progeny | 2015_P3-G12 | | H | [illegible] |
| 2015_P3-G12-4 | PHI2 | PHI1 | progeny | 2015_P3-G12 | | H | [illegible] |
| 2015_P3-G8-1 | PHI2 | PHI1 | progeny | 2015_P3-G8 | | H | [illegible] |
| 2015_P3-G8-2 | PHI2 | PHI1 | progeny | 2015_P3-G8 | | H | [illegible] |
| 2015_P3-F7-1 | PHI2 | PHI1 | progeny | 2015_P3-F7 | | G | [illegible] |
| 2015_P3-F7-2 | PHI2 | PHI1 | progeny | 2015_P3-F7 | | G | [illegible] |
| 2015_P3-F7-3 | PHI2 | PHI1 | progeny | 2015_P3-F7 | | G | [illegible] |
| 2015_P3-F7-4 | PHI2 | PHI1 | progeny | 2015_P3-F7 | | G | [illegible] |
| 2015_P3-F7-5 | PHI2 | PHI1 | progeny | 2015_P3-F7 | | G | [illegible] |
| 2015_P3-F12-1 | PHI2 | PHI1 | progeny | 2015_P3-F12 | | A | [illegible] |
| 2015_P3-F12-2 | PHI2 | PHI1 | progeny | 2015_P3-F12 | | H | [illegible] |
| 2015_P3-F12-3 | PHI2 | PHI1 | progeny | 2015_P3-F12 | | G | [illegible] |
| 2015_P3-F12-4 | PHI2 | PHI1 | progeny | 2015_P3-F12 | | H | [illegible] |
| 2015_P4-B3-1 | PHI2 | PHI1 | progeny | 2015_P4-B3 | | A | [illegible] |
| 2015_P4-B3-2 | PHI2 | PHI1 | progeny | 2015_P4-B3 | | H | [illegible] |
| 2015_P4-B3-3 | PHI2 | PHI1 | progeny | 2015_P4-B3 | | H | [illegible] |
| 2015_P3-A1 | PHI1 | PHI3 | mother | 2015_P3-A1 | | A | [illegible] |
| 2015_P3-A2 | PHI1 | PHI3 | mother | 2015_P3-A2 | | A | [illegible] |
| 2015_P3-A3 | PHI1 | PHI3 | mother | 2015_P3-A3 | | A | [illegible] |
| 2015_P3-A6 | PHI1 | PHI3 | mother | 2015_P3-A6 | | A | [illegible] |
| 2015_P3-A7 | PHI1 | PHI3 | mother | 2015_P3-A7 | | H | [illegible] |
| 2015_P3-A8 | PHI1 | PHI3 | mother | 2015_P3-A8 | | H | [illegible] |
| 2015_P3-H9 | PHI2 | PHI1 | mother | 2015_P3-H9 | | A | [illegible] |
| 2015_P3-A10 | PHI1 | PHI3 | mother | 2015_P3-A10 | | A | [illegible] |
| 2015_P3-A11 | PHI1 | PHI3 | mother | 2015_P3-A11 | | A | [illegible] |
| 2015_P3-B2 | PHI1 | PHI3 | mother | 2015_P3-B2 | | A | [illegible] |
| 2015_P3-B3 | PHI1 | PHI3 | mother | 2015_P3-B3 | | A | [illegible] |
| 2015_P3-B4 | PHI1 | PHI3 | mother | 2015_P3-B4 | | A | [illegible] |
| 2015_P3-B5 | PHI1 | PHI3 | mother | 2015_P3-B5 | | A | [illegible] |
| 2015_P3-B7 | PHI1 | PHI3 | mother | 2015_P3-B7 | | A | [illegible] |
| 2015_P3-B11 | PHI1 | PHI3 | mother | 2015_P3-B11 | | A | [illegible] |
| 2015_P3-C1 | PHI1 | PHI3 | mother | 2015_P3-C1 | | A | [illegible] |
| 2015_P3-C2 | PHI1 | PHI3 | mother | 2015_P3-C2 | | H | [illegible] |
| 2015_P3-C7 | PHI1 | PHI3 | mother | 2015_P3-C7 | | A | [illegible] |
| 2015_P3-C8 | PHI1 | PHI3 | mother | 2015_P3-C8 | | A | [illegible] |
| 2015_P3-C10 | PHI1 | PHI3 | mother | 2015_P3-C10 | | A | [illegible] |
| 2015_P3-C11 | PHI1 | PHI3 | mother | 2015_P3-C11 | | H | [illegible] |
| 2015_P3-D3 | PHI1 | PHI3 | mother | 2015_P3-D3 | | H | [illegible] |
| 2015_P3-D4 | PHI1 | PHI3 | mother | 2015_P3-D4 | | A | [illegible] |
| 2015_P3-D5 | PHI1 | PHI3 | mother | 2015_P3-D5 | | A | [illegible] |
| 2015_P3-D6 | PHI1 | PHI3 | mother | 2015_P3-D6 | | H | [illegible] |
| 2015_P3-D7 | PHI1 | PHI3 | mother | 2015_P3-D7 | | A | [illegible] |
| 2015_P3-D8 | PHI1 | PHI3 | mother | 2015_P3-D8 | | A | [illegible] |
| 2015_P3-D10 | PHI1 | PHI3 | mother | 2015_P3-D10 | | A | [illegible] |
| 2015_P3-E4 | PHI1 | PHI3 | mother | 2015_P3-E4 | | A | [illegible] |
| 2015_P3-E5 | PHI1 | PHI3 | mother | 2015_P3-E5 | | A | [illegible] |
| 2015_P3-E7 | PHI1 | PHI3 | mother | 2015_P3-E7 | | A | [illegible] |
| 2015_P3-E8 | PHI1 | PHI3 | mother | 2015_P3-E8 | | A | [illegible] |
| 2015_P3-E9 | PHI1 | PHI3 | mother | 2015_P3-E9 | | A | [illegible] |
| 2015_P3-E10 | PHI1 | PHI3 | mother | 2015_P3-E10 | | H | [illegible] |
| 2015_P3-E11 | PHI1 | PHI3 | mother | 2015_P3-E11 | | A | [illegible] |
| 2015_P3-E12 | PHI1 | PHI3 | mother | 2015_P3-E12 | | A | [illegible] |
| 2015_P3-F1 | PHI1 | PHI3 | mother | 2015_P3-F1 | | A | [illegible] |
| 2015_P3-F2 | PHI1 | PHI3 | mother | 2015_P3-F2 | | A | [illegible] |
| 2015_P3-F3 | PHI1 | PHI3 | mother | 2015_P3-F3 | | A | [illegible] |
| 2015_P3-F4 | PHI1 | PHI3 | mother | 2015_P3-F4 | | A | [illegible] |
| 2015_P3-F5 | PHI2 | PHI1 | mother | 2015_P3-F5 | | H | [illegible] |
| 2015_P3-F6 | PHI2 | PHI1 | mother | 2015_P3-F6 | | H | [illegible] |
| 2015_P3-F7 | PHI2 | PHI1 | mother | 2015_P3-F7 | | G | [illegible] |
| 2015_P3-F8 | PHI2 | PHI1 | mother | 2015_P3-F8 | | G | [illegible] |
| 2015_P3-F9 | PHI2 | PHI1 | mother | 2015_P3-F9 | | H | [illegible] |
| 2015_P3-F11 | PHI2 | PHI1 | mother | 2015_P3-F11 | | G | [illegible] |
| 2015_P3-F12 | PHI2 | PHI1 | mother | 2015_P3-F12 | | H | [illegible] |
| 2015_P3-G1 | PHI2 | PHI1 | mother | 2015_P3-G1 | | H | [illegible] |
| 2015_P3-G2 | PHI2 | PHI1 | mother | 2015_P3-G2 | | H | [illegible] |
| 2015_P3-G3 | PHI2 | PHI1 | mother | 2015_P3-G3 | | H | [illegible] |
| 2015_P3-G4 | PHI2 | PHI1 | mother | 2015_P3-G4 | | -9 | [illegible] |
| 2015_P3-G6 | PHI2 | PHI1 | mother | 2015_P3-G6 | | H | [illegible] |
| 2015_P3-G8 | PHI2 | PHI1 | mother | 2015_P3-G8 | | H | [illegible] |
| 2015_P3-G9 | PHI2 | PHI1 | mother | 2015_P3-G9 | | G | [illegible] |
| 2015_P3-G10 | PHI2 | PHI1 | mother | 2015_P3-G10 | | H | [illegible] |
| 2015_P3-G12 | PHI2 | PHI1 | mother | 2015_P3-G12 | | H | [illegible] |
| 2015_P3-H3 | PHI2 | PHI1 | mother | 2015_P3-H3 | | G | [illegible] |
| 2015_P3-H5 | PHI2 | PHI1 | mother | 2015_P3-H5 | | G | [illegible] |
| 2015_P3-H6 | PHI2 | PHI1 | mother | 2015_P3-H6 | | G | [illegible] |
| 2015_P3-H7 | PHI2 | PHI1 | mother | 2015_P3-H7 | | A | [illegible] |
| 2015_P3-H8 | PHI2 | PHI1 | mother | 2015_P3-H8 | | H | [illegible] |
| 2015_P3-H10 | PHI2 | PHI1 | mother | 2015_P3-H10 | | H | [illegible] |
| 2015_P3-H11 | PHI2 | PHI1 | mother | 2015_P3-H11 | | H | [illegible] |
| 2015_P4-A1 | PHI2 | PHI1 | mother | 2015_P4-A1 | | H | [illegible] |
| 2015_P4-A2 | PHI2 | PHI1 | mother | 2015_P4-A2 | | G | [illegible] |
| 2015_P4-A3 | PHI2 | PHI1 | mother | 2015_P4-A3 | | H | [illegible] |
| 2015_P4-A4 | PHI2 | PHI1 | mother | 2015_P4-A4 | | H | [illegible] |
| 2015_P4-A5 | PHI2 | PHI1 | mother | 2015_P4-A5 | | H | [illegible] |
| 2015_P4-A6 | PHI2 | PHI1 | mother | 2015_P4-A6 | | H | [illegible] |
| 2015_P4-A7 | PHI2 | PHI1 | mother | 2015_P4-A7 | | H | [illegible] |
| 2015_P4-A9 | PHI2 | PHI1 | mother | 2015_P4-A9 | | H | [illegible] |
| 2015_P4-B3 | PHI2 | PHI1 | mother | 2015_P4-B3 | | H | [illegible] |
| 2015_P4-B4 | PHI2 | PHI1 | mother | 2015_P4-B4 | | H | [illegible] |
| 2015_P4-B6 | PHI2 | PHI1 | mother | 2015_P4-B6 | | G | [illegible] |
| 2015_P4-B7 | PHI2 | PHI3 | mother | 2015_P4-B7 | | H | [illegible] |
| 2015_P4-B9 | PHI2 | PHI3 | mother | 2015_P4-B9 | | H | [illegible] |
| 2015_P4-B11 | PHI2 | PHI3 | mother | 2015_P4-B11 | | H | [illegible] |
| 2015_P4-C1 | PHI2 | PHI3 | mother | 2015_P4-C1 | | H | [illegible] |
| 2015_P4-C2 | PHI2 | PHI3 | mother | 2015_P4-C2 | | H | [illegible] |
| 2015_P4-C8 | PHI2 | PHI3 | mother | 2015_P4-C8 | | H | [illegible] |
| 2015_P4-C9 | PHI2 | PHI3 | mother | 2015_P4-C9 | | H | [illegible] |
| 2015_P4-C10 | PHI2 | PHI3 | mother | 2015_P4-C10 | | H | [illegible] |
| 2015_P4-C11 | PHI2 | PHI3 | mother | 2015_P4-C11 | | H | [illegible] |
| 2015_P4-D3 | PHI2 | PHI3 | mother | 2015_P4-D3 | | H | [illegible] |
| 2015_P4-D4 | PHI2 | PHI3 | mother | 2015_P4-D4 | | H | [illegible] |
| 2015_P4-D7 | PHI2 | PHI3 | mother | 2015_P4-D7 | | H | [illegible] |
| 2015_P4-D12 | PHI2 | PHI3 | mother | 2015_P4-D12 | | H | [illegible] |
| 2015_P4-E1 | PHI2 | PHI3 | mother | 2015_P4-E1 | | H | [illegible] |
| 2015_P4-E5 | PHI2 | PHI3 | mother | 2015_P4-E5 | | H | [illegible] |
| 2015_P4-E6 | PHI2 | PHI3 | mother | 2015_P4-E6 | | H | [illegible] |
| 2015_P4-E8 | PHI8 | PHI3 | mother | 2015_P4-E8 | | H | [illegible] |
| 2015_P4-E9 | PHI8 | PHI3 | mother | 2015_P4-E9 | | H | [illegible] |
| 2015_P4-E10 | PHI8 | PHI3 | mother | 2015_P4-E10 | | H | [illegible] |
| 2015_P4-E11 | PHI8 | PHI3 | mother | 2015_P4-E11 | | H | [illegible] |
| 2015_P4-E12 | PHI8 | PHI3 | mother | 2015_P4-E12 | | H | [illegible] |
| 2015_P4-F1 | PHI8 | PHI3 | mother | 2015_P4-F1 | | H | [illegible] |
| 2015_P4-F2 | PHI8 | PHI3 | mother | 2015_P4-F2 | | H | [illegible] |
| 2015_P4-F4 | PHI8 | PHI3 | mother | 2015_P4-F4 | | H | [illegible] |
| 2013_O4 | PHI2 | PHI2 | ancestor | 2013_O4 | | G | [illegible] |
| 2013_O1 | PHI1 | PHI1 | ancestor | 2013_O1 | | A | [illegible] |
| 2015_O2 | PHI3 | PHI3 | ancestor | 2015_O2 | | A | [illegible] |
| 2015_O3 | PHI8 | PHI8 | ancestor | 2015_O3 | | G | [illegible] |

plant_id – unique identifier of the plant
mom_gen – genetic backgroiund of mother in the hybred
dad_gen – genetic background of father in the hybred
family_ – identify mother and all the progeny plants from the same mother
mother_progeny – identify each plant as mother of the family or progeny
Chr – number of the chromosome, where marker is located
Distance – position of the marker on the corresponding chromosome in cM
H – heterozygous marker
A, T, G and C – marker homozygous for the corresponding nucleotide
EQV, LS, -9 , etc. – missing values

### Table S3. Results of the SNP array analasis (continue)

| plant_id | mom_gen | dad_gen | mother_progeny | family_ | Chr / Position, cM | SNP genotype calls |
|---|---|---|---|---|---|---|
| 2015_P3-A11-1 | PHI1 | PHI3 | progeny | 2015_P3-A11 | | [illegible] |
| 2015_P3-A11-2 | PHI1 | PHI3 | progeny | 2015_P3-A11 | | [illegible] |
| 2015_P3-A11-3 | PHI1 | PHI3 | progeny | 2015_P3-A11 | | [illegible] |
| 2015_P3-A10-1 | PHI1 | PHI3 | progeny | 2015_P3-A10 | | [illegible] |
| 2015_P3-A8-1 | PHI1 | PHI3 | progeny | 2015_P3-A8 | | [illegible] |
| 2015_P3-A8-2 | PHI1 | PHI3 | progeny | 2015_P3-A8 | | [illegible] |
| 2015_P3-A8-3 | PHI1 | PHI3 | progeny | 2015_P3-A8 | | [illegible] |
| 2015_P3-A7-1 | PHI1 | PHI3 | progeny | 2015_P3-A7 | | [illegible] |
| 2015_P3-A6-1 | PHI1 | PHI3 | progeny | 2015_P3-A6 | | [illegible] |
| 2015_P3-A2-1 | PHI1 | PHI3 | progeny | 2015_P3-A2 | | [illegible] |
| 2015_P3-A2-2 | PHI1 | PHI3 | progeny | 2015_P3-A2 | | [illegible] |
| 2015_P3-A2-3 | PHI1 | PHI3 | progeny | 2015_P3-A2 | | [illegible] |
| 2015_P3-A2-4 | PHI1 | PHI3 | progeny | 2015_P3-A2 | | [illegible] |
| 2015_P3-A1-1 | PHI1 | PHI3 | progeny | 2015_P3-A1 | | [illegible] |
| 2015_P3-A3-1 | PHI1 | PHI3 | progeny | 2015_P3-A3 | | [illegible] |
| 2015_P3-B11-1 | PHI1 | PHI3 | progeny | 2015_P3-B11 | | [illegible] |
| 2015_P3-C1-1 | PHI1 | PHI3 | progeny | 2015_P3-C1 | | [illegible] |
| 2015_P3-C2-1 | PHI1 | PHI3 | progeny | 2015_P3-C2 | | [illegible] |
| 2015_P3-C8-1 | PHI1 | PHI3 | progeny | 2015_P3-C8 | | [illegible] |
| 2015_P3-C8-2 | PHI1 | PHI3 | progeny | 2015_P3-C8 | | [illegible] |
| 2015_P3-C8-3 | PHI1 | PHI3 | progeny | 2015_P3-C8 | | [illegible] |
| 2015_P3-C7-1 | PHI1 | PHI3 | progeny | 2015_P3-C7 | | [illegible] |
| 2015_P3-C10-2 | PHI1 | PHI3 | progeny | 2015_P3-C10 | | [illegible] |
| 2015_P3-C11-3 | PHI1 | PHI3 | progeny | 2015_P3-C11 | | [illegible] |
| 2015_P3-D3-1 | PHI1 | PHI3 | progeny | 2015_P3-D3 | | [illegible] |
| 2015_P3-D3-2 | PHI1 | PHI3 | progeny | 2015_P3-D3 | | [illegible] |
| 2015_P3-D4-1 | PHI1 | PHI3 | progeny | 2015_P3-D4 | | [illegible] |
| 2015_P3-D4-2 | PHI1 | PHI3 | progeny | 2015_P3-D4 | | [illegible] |
| 2015_P3-D4-3 | PHI1 | PHI3 | progeny | 2015_P3-D4 | | [illegible] |
| 2015_P3-D6-1 | PHI1 | PHI3 | progeny | 2015_P3-D6 | | [illegible] |
| 2015_P3-B5-1 | PHI1 | PHI3 | progeny | 2015_P3-B5 | | [illegible] |
| 2015_P3-B4-1 | PHI1 | PHI3 | progeny | 2015_P3-B4 | | [illegible] |
| 2015_P3-D5-1 | PHI1 | PHI3 | progeny | 2015_P3-D5 | | [illegible] |
| 2015_P3-D10-1 | PHI1 | PHI3 | progeny | 2015_P3-D10 | | [illegible] |
| 2015_P3-E4-1 | PHI1 | PHI3 | progeny | 2015_P3-E4 | | [illegible] |
| 2015_P3-E4-2 | PHI1 | PHI3 | progeny | 2015_P3-E4 | | [illegible] |
| 2015_P3-E5-1 | PHI1 | PHI3 | progeny | 2015_P3-E5 | | [illegible] |
| 2015_P3-E5-2 | PHI1 | PHI3 | progeny | 2015_P3-E5 | | [illegible] |
| 2015_P3-E5-3 | PHI1 | PHI3 | progeny | 2015_P3-E5 | | [illegible] |
| 2015_P3-E7-1 | PHI1 | PHI3 | progeny | 2015_P3-E7 | | [illegible] |
| 2015_P3-E9-1 | PHI1 | PHI3 | progeny | 2015_P3-E9 | | [illegible] |
| 2015_P3-E9-2 | PHI1 | PHI3 | progeny | 2015_P3-E9 | | [illegible] |
| 2015_P3-E9-3 | PHI1 | PHI3 | progeny | 2015_P3-E9 | | [illegible] |
| 2015_P3-E9-4 | PHI1 | PHI3 | progeny | 2015_P3-E9 | | [illegible] |
| 2015_P3-E8-1 | PHI1 | PHI3 | progeny | 2015_P3-E8 | | [illegible] |
| 2015_P3-E8-2 | PHI1 | PHI3 | progeny | 2015_P3-E8 | | [illegible] |
| 2015_P3-E8-3 | PHI1 | PHI3 | progeny | 2015_P3-E8 | | [illegible] |
| 2015_P3-E10-1 | PHI1 | PHI3 | progeny | 2015_P3-E10 | | [illegible] |
| 2015_P3-E11-1 | PHI1 | PHI3 | progeny | 2015_P3-E11 | | [illegible] |
| 2015_P3-E12-1 | PHI1 | PHI3 | progeny | 2015_P3-E12 | | [illegible] |
| 2015_P3-F1-1 | PHI1 | PHI3 | progeny | 2015_P3-F1 | | [illegible] |
| 2015_P3-F2-1 | PHI1 | PHI3 | progeny | 2015_P3-F2 | | [illegible] |
| 2015_P3-F4-1 | PHI1 | PHI3 | progeny | 2015_P3-F4 | | [illegible] |
| 2015_P3-F4-2 | PHI1 | PHI3 | progeny | 2015_P3-F4 | | [illegible] |
| 2015_P3-F4-3 | PHI1 | PHI3 | progeny | 2015_P3-F4 | | [illegible] |
| 2015_P3-F4-4 | PHI1 | PHI3 | progeny | 2015_P3-F4 | | [illegible] |
| 2015_P3-F3-1 | PHI1 | PHI3 | progeny | 2015_P3-F3 | | [illegible] |
| 2015_P3-F3-2 | PHI1 | PHI3 | progeny | 2015_P3-F3 | | [illegible] |
| 2015_P3-B3-1 | PHI1 | PHI3 | progeny | 2015_P3-B3 | | [illegible] |
| 2015_P3-B7-1 | PHI1 | PHI3 | progeny | 2015_P3-B7 | | [illegible] |
| 2015_P3-B2-1 | PHI1 | PHI3 | progeny | 2015_P3-B2 | | [illegible] |
| 2015_P3-B2-2 | PHI1 | PHI3 | progeny | 2015_P3-B2 | | [illegible] |
| 2015_P3-D7-1 | PHI1 | PHI3 | progeny | 2015_P3-D7 | | [illegible] |
| 2015_P3-D7-2 | PHI1 | PHI3 | progeny | 2015_P3-D7 | | [illegible] |
| 2015_P4-C8-1 | PHI2 | PHI3 | progeny | 2015_P4-C8 | | [illegible] |
| 2015_P4-C8-2 | PHI2 | PHI3 | progeny | 2015_P4-C8 | | [illegible] |
| 2015_P4-B11-1 | PHI2 | PHI3 | progeny | 2015_P4-B11 | | [illegible] |
| 2015_P4-B11-3 | PHI2 | PHI3 | progeny | 2015_P4-B11 | | [illegible] |
| 2015_P4-C11-1 | PHI2 | PHI3 | progeny | 2015_P4-C11 | | [illegible] |
| 2015_P4-C11-2 | PHI2 | PHI3 | progeny | 2015_P4-C11 | | [illegible] |
| 2015_P4-E6-1 | PHI2 | PHI3 | progeny | 2015_P4-E6 | | [illegible] |
| 2015_P4-E6-2 | PHI2 | PHI3 | progeny | 2015_P4-E6 | | [illegible] |
| 2015_P4-D12-1 | PHI2 | PHI3 | progeny | 2015_P4-D12 | | [illegible] |
| 2015_P4-D12-2 | PHI2 | PHI3 | progeny | 2015_P4-D12 | | [illegible] |
| 2015_P4-E1-1 | PHI2 | PHI3 | progeny | 2015_P4-E1 | | [illegible] |
| 2015_P4-E1-2 | PHI2 | PHI3 | progeny | 2015_P4-E1 | | [illegible] |
| 2015_P4-C10-1 | PHI2 | PHI3 | progeny | 2015_P4-C10 | | [illegible] |
| 2015_P4-B9-1 | PHI2 | PHI3 | progeny | 2015_P4-B9 | | [illegible] |
| 2015_P4-B9-2 | PHI2 | PHI3 | progeny | 2015_P4-B9 | | [illegible] |
| 2015_P4-B7-1 | PHI2 | PHI3 | progeny | 2015_P4-B7 | | [illegible] |
| 2015_P4-B7-2 | PHI2 | PHI3 | progeny | 2015_P4-B7 | | [illegible] |
| 2015_P4-C2-1 | PHI2 | PHI3 | progeny | 2015_P4-C2 | | [illegible] |
| 2015_P4-E5-1 | PHI2 | PHI3 | progeny | 2015_P4-E5 | | [illegible] |
| 2015_P4-E5-2 | PHI2 | PHI3 | progeny | 2015_P4-E5 | | [illegible] |
| 2015_P4-E5-3 | PHI2 | PHI3 | progeny | 2015_P4-E5 | | [illegible] |
| 2015_P4-C9-1 | PHI2 | PHI3 | progeny | 2015_P4-C9 | | [illegible] |
| 2015_P4-C9-2 | PHI2 | PHI3 | progeny | 2015_P4-C9 | | [illegible] |
| 2015_P4-D4-1 | PHI2 | PHI3 | progeny | 2015_P4-D4 | | [illegible] |
| 2015_P4-C1-1 | PHI2 | PHI3 | progeny | 2015_P4-C1 | | [illegible] |
| 2015_P4-C1-2 | PHI2 | PHI3 | progeny | 2015_P4-C1 | | [illegible] |
| 2015_P4-D7-1 | PHI2 | PHI3 | progeny | 2015_P4-D7 | | [illegible] |
| 2015_P4-D3-1 | PHI2 | PHI3 | progeny | 2015_P4-D3 | | [illegible] |
| 2015_P4-F2-1 | PHI8 | PHI3 | progeny | 2015_P4-F2 | | [illegible] |
| 2015_P4-F2-2 | PHI8 | PHI3 | progeny | 2015_P4-F2 | | [illegible] |
| 2015_P4-F4-1 | PHI8 | PHI3 | progeny | 2015_P4-F4 | | [illegible] |
| 2015_P4-F4-2 | PHI8 | PHI3 | progeny | 2015_P4-F4 | | [illegible] |
| 2015_P4-F4-3 | PHI8 | PHI3 | progeny | 2015_P4-F4 | | [illegible] |
| 2015_P4-F4-4 | PHI8 | PHI3 | progeny | 2015_P4-F4 | | [illegible] |
| 2015_P4-E9-1 | PHI8 | PHI3 | progeny | 2015_P4-E9 | | [illegible] |
| 2015_P4-E9-2 | PHI8 | PHI3 | progeny | 2015_P4-E9 | | [illegible] |
| 2015_P4-F1-1 | PHI8 | PHI3 | progeny | 2015_P4-F1 | | [illegible] |
| 2015_P4-E11-1 | PHI8 | PHI3 | progeny | 2015_P4-E11 | | [illegible] |
| 2015_P4-E11-2 | PHI8 | PHI3 | progeny | 2015_P4-E11 | | [illegible] |
| 2015_P4-E10-1 | PHI8 | PHI3 | progeny | 2015_P4-E10 | | [illegible] |
| 2015_P4-E10-2 | PHI8 | PHI3 | progeny | 2015_P4-E10 | | [illegible] |
| 2015_P4-E10-3 | PHI8 | PHI3 | progeny | 2015_P4-E10 | | [illegible] |
| 2015_P4-E10-4 | PHI8 | PHI3 | progeny | 2015_P4-E10 | | [illegible] |
| 2015_P4-E8-1 | PHI8 | PHI3 | progeny | 2015_P4-E8 | | [illegible] |
| 2015_P4-E12-1 | PHI8 | PHI3 | progeny | 2015_P4-E12 | | [illegible] |
| 2015_P4-E12-2 | PHI8 | PHI3 | progeny | 2015_P4-E12 | | [illegible] |
| 2015_P4-F5-1 | PHI2 | PHI8 | progeny | 2015_P4-F5 | | [illegible] |
| 2015_P4-F5-2 | PHI2 | PHI8 | progeny | 2015_P4-F5 | | [illegible] |
| 2015_P4-F5-3 | PHI2 | PHI8 | progeny | 2015_P4-F5 | | [illegible] |
| 2015_P4-F5-4 | PHI2 | PHI8 | progeny | 2015_P4-F5 | | [illegible] |
| 2015_P4-F5-5 | PHI2 | PHI8 | progeny | 2015_P4-F5 | | [illegible] |
| 2015_P4-F5-6 | PHI2 | PHI8 | progeny | 2015_P4-F5 | | [illegible] |
| 2015_P4-F5-7 | PHI2 | PHI8 | progeny | 2015_P4-F5 | | [illegible] |
| 2015_P4-F5 | PHI2 | PHI8 | mother | 2015_P4-F5 | | [illegible] |
| 2015_P3-G1-1 | PHI2 | PHI1 | progeny | 2015_P3-G1 | | [illegible] |
| 2015_P3-G1-2 | PHI2 | PHI1 | progeny | 2015_P3-G1 | | [illegible] |
| 2015_P3-G1-3 | PHI2 | PHI1 | progeny | 2015_P3-G1 | | [illegible] |
| 2015_P3-G1-4 | PHI2 | PHI1 | progeny | 2015_P3-G1 | | [illegible] |
| 2015_P4-A1-1 | PHI2 | PHI1 | progeny | 2015_P4-A1 | | [illegible] |
| 2015_P4-A1-2 | PHI2 | PHI1 | progeny | 2015_P4-A1 | | [illegible] |
| 2015_P3-F11-1 | PHI2 | PHI1 | progeny | 2015_P3-F11 | | [illegible] |
| 2015_P3-H5-1 | PHI2 | PHI1 | progeny | 2015_P3-H5 | | [illegible] |
| 2015_P3-H5-2 | PHI2 | PHI1 | progeny | 2015_P3-H5 | | [illegible] |
| 2015_P3-H5-3 | PHI2 | PHI1 | progeny | 2015_P3-H5 | | [illegible] |
| 2015_P3-H5-4 | PHI2 | PHI1 | progeny | 2015_P3-H5 | | [illegible] |
| 2015_P3-H3-1 | PHI2 | PHI1 | progeny | 2015_P3-H3 | | [illegible] |
| 2015_P3-H3-2 | PHI2 | PHI1 | progeny | 2015_P3-H3 | | [illegible] |
| 2015_P3-H6-1 | PHI2 | PHI1 | progeny | 2015_P3-H6 | | [illegible] |
| 2015_P3-H7-1 | PHI2 | PHI1 | progeny | 2015_P3-H7 | | [illegible] |
| 2015_P4-B6-1 | PHI2 | PHI1 | progeny | 2015_P4-B6 | | [illegible] |
| 2015_P4-B6-2 | PHI2 | PHI1 | progeny | 2015_P4-B6 | | [illegible] |
| 2015_P3-G6-1 | PHI2 | PHI1 | progeny | 2015_P3-G6 | | [illegible] |
| 2015_P3-G6-2 | PHI2 | PHI1 | progeny | 2015_P3-G6 | | [illegible] |
| 2015_P3-G3-1 | PHI2 | PHI1 | progeny | 2015_P3-G3 | | [illegible] |
| 2015_P4-A2-1 | PHI2 | PHI1 | progeny | 2015_P4-A2 | | [illegible] |
| 2015_P4-A9-1 | PHI2 | PHI1 | progeny | 2015_P4-A9 | | [illegible] |
| 2015_P4-A9-2 | PHI2 | PHI1 | progeny | 2015_P4-A9 | | [illegible] |
| 2015_P4-A9-3 | PHI2 | PHI1 | progeny | 2015_P4-A9 | | [illegible] |
| 2015_P4-A9-4 | PHI2 | PHI1 | progeny | 2015_P4-A9 | | [illegible] |
| 2015_P4-B4-1 | PHI2 | PHI1 | progeny | 2015_P4-B4 | | [illegible] |
| 2015_P4-B4-2 | PHI2 | PHI1 | progeny | 2015_P4-B4 | | [illegible] |
| 2015_P3-F5-1 | PHI2 | PHI1 | progeny | 2015_P3-F5 | | [illegible] |
| 2015_P3-F6-1 | PHI2 | PHI1 | progeny | 2015_P3-F6 | | [illegible] |
| 2015_P3-F6-2 | PHI2 | PHI1 | progeny | 2015_P3-F6 | | [illegible] |
| 2015_P3-F6-3 | PHI2 | PHI1 | progeny | 2015_P3-F6 | | [illegible] |
| 2015_P4-A3-1 | PHI2 | PHI1 | progeny | 2015_P4-A3 | | [illegible] |
| 2015_P3-H9-1 | PHI2 | PHI1 | progeny | 2015_P3-H9 | | [illegible] |
| 2015_P3-H9-2 | PHI2 | PHI1 | progeny | 2015_P3-H9 | | [illegible] |
| 2015_P3-H9-3 | PHI2 | PHI1 | progeny | 2015_P3-H9 | | [illegible] |
| 2015_P3-H9-4 | PHI2 | PHI1 | progeny | 2015_P3-H9 | | [illegible] |
| 2015_P3-H9-5 | PHI2 | PHI1 | progeny | 2015_P3-H9 | | [illegible] |
| 2015_P3-H9-6 | PHI2 | PHI1 | progeny | 2015_P3-H9 | | [illegible] |
| 2015_P4-A4-1 | PHI2 | PHI1 | progeny | 2015_P4-A4 | | [illegible] |
| 2015_P4-A4-2 | PHI2 | PHI1 | progeny | 2015_P4-A4 | | [illegible] |
| 2015_P4-A4-3 | PHI2 | PHI1 | progeny | 2015_P4-A4 | | [illegible] |
| 2015_P4-A4-4 | PHI2 | PHI1 | progeny | 2015_P4-A4 | | [illegible] |
| 2015_P3-G10-1 | PHI2 | PHI1 | progeny | 2015_P3-G10 | | [illegible] |
| 2015_P3-G10-2 | PHI2 | PHI1 | progeny | 2015_P3-G10 | | [illegible] |
| 2015_P3-G10-3 | PHI2 | PHI1 | progeny | 2015_P3-G10 | | [illegible] |
| 2015_P3-G10-4 | PHI2 | PHI1 | progeny | 2015_P3-G10 | | [illegible] |
| 2015_P3-G10-5 | PHI2 | PHI1 | progeny | 2015_P3-G10 | | [illegible] |
| 2015_P3-G10-6 | PHI2 | PHI1 | progeny | 2015_P3-G10 | | [illegible] |
| 2015_P3-G10-7 | PHI2 | PHI1 | progeny | 2015_P3-G10 | | [illegible] |
| 2015_P3-G10-8 | PHI2 | PHI1 | progeny | 2015_P3-G10 | | [illegible] |
| 2015_P3-G10-9 | PHI2 | PHI1 | progeny | 2015_P3-G10 | | [illegible] |
| 2015_P3-G10-10 | PHI2 | PHI1 | progeny | 2015_P3-G10 | | [illegible] |
| 2015_P3-G9-1 | PHI2 | PHI1 | progeny | 2015_P3-G9 | | [illegible] |
| 2015_P3-G9-2 | PHI2 | PHI1 | progeny | 2015_P3-G9 | | [illegible] |
| 2015_P3-G9-3 | PHI2 | PHI1 | progeny | 2015_P3-G9 | | [illegible] |
| 2015_P3-G9-4 | PHI2 | PHI1 | progeny | 2015_P3-G9 | | [illegible] |
| 2015_P3-G9-5 | PHI2 | PHI1 | progeny | 2015_P3-G9 | | [illegible] |
| 2015_P3-G9-6 | PHI2 | PHI1 | progeny | 2015_P3-G9 | | [illegible] |
| 2015_P3-G9-7 | PHI2 | PHI1 | progeny | 2015_P3-G9 | | [illegible] |
| 2015_P3-G9-8 | PHI2 | PHI1 | progeny | 2015_P3-G9 | | [illegible] |
| 2015_P3-G9-9 | PHI2 | PHI1 | progeny | 2015_P3-G9 | | [illegible] |

**Table S3. Results of the SNP array analasis (continue)**

[illegible]

plant_id – unique identifier of the plant
mom_gen – genetic backgroiund of mother in the hybre
dad_gen – genetic background of father in the hybred
family_ – identify mother and all the progeny plants f
mother_progeny – identify each plant as mother of the family c
Chr – number of the chromosome, where marker
Distance – position of the marker on the correspondin
H – heterozygous marker
A, T, G and C – marker homozygous for the corresponding
EQV, LS, -9 , etc. – missing values

### Table S4. Results of the SNP array analasis

| | | | | | Chr | 1 | 1 | 1 | 1 | 1 | 1 | 1 | 1 | 1 | 1 | 1 | 1 | 1 | 1 | 2 | 2 | 2 | 2 | 2 | 2 | 2 | 3 | 3 | 3 | 3 | 3 | 3 | 3 | 3 | 3 | 3 | 3 | 3 | 3 | 3 | 4 | 4 | 4 | 4 | 4 | 4 | 4 | 4 | 5 | 5 | 5 | 5 | 5 | 5 | 5 | 5 | 5 | 5 | 5 | 5 | 5 | 6 | 6 | 6 | 6 | 6 | 7 | 7 | 7 | 7 | 7 | 7 | 7 | 7 | 7 | 7 | 7 | 7 | 7 | 7 | 7 | 7 | 8 | 8 | 8 | 8 | 8 | 8 | 8 | 8 | 8 | 9 | 9 | 9 | 9 | 9 | 10 | 10 | 10 | 10 | 10 |
|---|---|---|---|---|---|---|---|---|---|---|---|---|---|---|---|---|---|---|---|---|---|---|---|---|---|---|---|---|---|---|---|---|---|---|---|---|---|---|---|---|---|---|---|---|---|---|---|---|---|---|---|---|---|---|---|---|---|---|---|---|---|---|---|---|---|---|---|---|---|---|---|---|---|---|---|---|---|---|---|---|---|---|---|---|---|---|---|---|---|---|---|---|---|---|---|---|---|---|---|---|---|
| | | | | | Position, cM | 48 | 59 | 80 | 85 | 97 | 113 | 116 | 137 | 137 | 143 | 162 | 197 | 218 | 285 | 25 | 33 | 142 | 142 | 162 | 193 | 202 | 20 | 39 | 75 | 75 | 75 | 102 | 104 | 127 | 140 | 199 | 213 | 228 | 237 | 246 | 30 | 73 | 83 | 91 | 128 | 146 | 166 | 197 | 20 | 65 | 71 | 77 | 102 | 107 | 108 | 132 | 154 | 164 | 191 | 208 | 218 | 27 | 34 | 86 | 146 | 150 | 5.6 | 37 | 52 | 57 | 74 | 78 | 79 | 80 | 86 | 93 | 109 | 112 | 166 | 173 | 182 | 192 | 50 | 131 | 135 | 140 | 149 | 153 | 164 | 186 | 191 | 60 | 76 | 78 | 144 | 164 | 77 | 89 | 101 | 148 | 154 |
| plant_id | mom_gen | dad_gen | mother_progeny | family_ | | | | | | | | | | | | | | | | | | | | | | | | | | | | | | | | | | | | | | | | | | | | | | | | | | | | | | | | | | | | | | | | | | | | | | | | | | | | | | | | | | | | | | | | | | | | | | | | | |
| 2013_O1 | PHI1 | PHI1 | ancestor | 2013_O1 | | A | T | T | G | C | G | G | C | A | C | G | A | A | H | G | C | G | G | C | A | T | T | T | C | C | C | T | A | G | A | G | C | C | A | G | C | G | T | C | G | T | G | G | C | A | G | C | G | T | T | C | A | A | T | G | G | G | G | A | A | G | G | C | A | T | T | C | A | A | A | G | G | A | T | T | A | A | T | C | A | C | T | T | A | C | T | A | C | T | G | A | T | A | T | T | G |
| 2015_O2 | PHI3 | PHI3 | ancestor | 2015_O2 | | A | C | C | A | C | T | G | C | A | C | G | G | A | A | A | T | G | A | C | A | T | T | A | C | C | T | C | T | A | T | G | A | T | A | T | G | G | C | C | G | C | T | A | C | G | G | C | C | C | T | T | G | A | T | G | G | T | -9 | G | G | T | C | A | G | C | C | G | G | G | A | A | C | C | T | T | A | G | T | C | A | T | C | C | G | T | G | G | T | G | G | A | C | A | T | -9 | G |
| 2015_O3 | PHI8 | PHI8 | ancestor | 2015_O3 | | T | T | T | G | -9 | G | A | T | C | T | A | A | C | G | A | T | T | G | G | G | C | A | T | T | G | C | T | A | A | T | T | C | C | G | T | G | C | T | C | A | T | T | A | T | A | A | T | C | T | C | C | A | G | G | T | G | G | -9 | A | A | G | C | C | A | T | T | C | A | G | C | G | G | A | G | C | C | A | G | A | G | C | T | T | A | C | T | A | C | T | C | G | T | G | C | -9 | G |
| 2013_O4 | PHI2 | PHI2 | ancestor | 2013_O4 | | T | T | T | H | C | G | G | H | A | C | H | H | A | G | G | T | T | A | C | A | H | T | T | H | H | C | C | T | G | H | G | C | C | A | G | G | C | T | T | H | T | -9 | A | H | G | A | H | G | T | T | C | G | H | T | G | G | H | G | H | -9 | T | G | C | A | C | T | G | -9 | G | A | G | G | H | G | T | -9 | H | T | A | A | T | -9 | -9 | H | C | H | H | C | H | H | H | T | G | H | T | A |
| 2015_P3-A1 | PHI1 | PHI3 | mother | 2015_P3-A1 | | A | C | H | H | C | H | G | C | A | C | G | H | A | A | H | H | G | H | C | A | T | T | A | H | H | H | H | H | H | T | G | H | H | A | T | H | G | H | C | H | H | T | A | C | H | G | C | H | C | H | T | H | A | T | G | G | H | G | G | H | H | H | H | H | H | H | H | H | H | A | H | H | H | H | T | C | H | T | C | A | H | H | T | A | H | H | H | H | G | G | A | H | A | T | T | G |
| 2015_P3-A1-1 | PHI1 | PHI3 | progeny | 2015_P3-A1 | | A | H | H | H | C | G | -9 | C | H | C | H | H | A | H | G | H | G | H | C | A | H | T | H | T | G | C | T | A | G | H | G | C | H | H | H | G | G | H | H | G | C | H | A | H | H | G | H | H | H | C | T | A | A | T | G | G | H | G | H | H | H | -9 | H | H | H | H | H | H | A | A | G | G | H | G | T | C | H | T | C | A | H | -9 | H | -9 | -9 | H | H | H | H | H | A | H | A | T | T | H |
| 2015_P3-A10 | PHI1 | PHI3 | mother | 2015_P3-A10 | | A | H | H | H | C | H | G | C | A | C | G | H | A | A | H | H | G | H | C | A | T | T | H | C | C | H | C | T | H | H | G | H | H | A | H | G | G | H | C | H | H | H | A | C | H | G | C | H | H | T | H | H | A | T | G | G | H | G | H | G | H | H | H | H | H | H | H | H | H | A | H | H | H | T | T | C | H | T | C | A | H | C | T | A | H | H | H | H | H | G | A | T | H | H | T | G |
| 2015_P3-A10-1 | PHI1 | PHI3 | progeny | 2015_P3-A10 | | A | H | H | H | C | H | G | C | A | C | G | H | A | A | H | H | G | H | C | A | T | T | H | C | C | H | C | T | H | H | G | C | C | A | G | G | G | T | C | A | T | G | A | C | H | G | C | H | H | T | H | H | A | T | G | G | H | G | H | G | H | H | H | H | H | H | H | H | H | A | H | H | H | T | T | C | H | T | C | A | C | C | T | A | C | T | H | H | H | G | A | T | H | H | T | G |
| 2015_P3-A11 | PHI1 | PHI3 | mother | 2015_P3-A11 | | A | C | H | H | C | H | G | C | A | C | G | H | A | A | H | H | G | H | C | A | T | T | H | H | H | H | H | H | H | H | G | H | H | A | H | G | H | H | H | H | H | T | A | C | H | G | C | H | H | T | H | H | A | T | G | G | H | G | H | H | H | H | H | H | H | H | H | H | H | A | H | H | H | H | T | C | A | T | C | A | H | H | H | H | H | H | H | H | G | G | A | H | A | T | T | G |
| 2015_P3-A11-1 | PHI1 | PHI3 | progeny | 2015_P3-A11 | | -9 | -9 | -9 | G | -9 | G | -9 | C | A | C | -9 | A | A | A | -9 | H | G | G | -9 | -9 | T | -9 | T | T | -9 | -9 | -9 | A | -9 | A | G | A | T | -9 | T | -9 | C | T | -9 | -9 | T | -9 | -9 | C | G | -9 | C | C | -9 | -9 | T | -9 | A | T | G | -9 | T | -9 | -9 | G | -9 | H | -9 | A | -9 | -9 | -9 | A | A | A | G | G | -9 | H | -9 | C | A | T | C | A | H | C | T | -9 | -9 | -9 | A | C | -9 | G | -9 | H | -9 | -9 | -9 | G |
| 2015_P3-A11-2 | PHI1 | PHI3 | progeny | 2015_P3-A11 | | A | C | T | G | C | G | G | C | A | C | G | A | A | A | A | T | G | A | C | A | T | T | A | C | C | T | C | T | A | -9 | G | A | T | A | T | G | G | C | C | G | C | T | A | C | H | G | C | G | T | T | C | A | A | T | G | G | G | G | A | A | G | G | A | G | C | -9 | G | G | G | A | A | C | C | H | T | C | A | T | C | A | C | T | T | A | C | T | H | C | G | G | A | T | A | T | T | G |
| 2015_P3-A11-3 | PHI1 | PHI3 | progeny | 2015_P3-A11 | | A | C | C | A | C | T | G | C | A | C | G | G | A | A | H | T | G | A | C | A | T | T | A | C | C | T | C | T | A | T | G | A | T | A | T | G | G | C | C | G | C | T | A | C | A | G | C | G | T | T | C | A | A | T | G | G | T | -9 | G | H | H | H | H | H | H | C | G | G | G | A | A | C | C | H | T | C | A | T | C | A | T | C | -9 | G | T | G | G | T | G | G | A | H | A | T | T | G |
| 2015_P3-A2 | PHI1 | PHI3 | mother | 2015_P3-A2 | | A | H | H | H | C | H | G | C | A | C | G | H | A | A | H | H | G | H | C | A | T | T | H | C | C | H | C | T | H | H | G | H | H | A | H | G | G | H | C | H | H | H | H | C | H | G | C | H | H | T | H | H | A | T | G | G | H | G | G | H | H | H | H | H | H | H | H | H | H | A | H | H | H | T | T | C | A | T | C | A | H | C | H | H | H | H | A | H | H | G | A | H | A | T | T | G |
| 2015_P3-A2-1 | PHI1 | PHI3 | progeny | 2015_P3-A2 | | [illegible] |
| 2015_P3-A2-2 | PHI1 | PHI3 | progeny | 2015_P3-A2 | | [illegible] |
| 2015_P3-A2-3 | PHI1 | PHI3 | progeny | 2015_P3-A2 | | [illegible] |
| 2015_P3-A2-4 | PHI1 | PHI3 | progeny | 2015_P3-A2 | | [illegible] |
| 2015_P3-A3 | PHI1 | PHI3 | mother | 2015_P3-A3 | | [illegible] |
| 2015_P3-A3-1 | PHI1 | PHI3 | progeny | 2015_P3-A3 | | [illegible] |
| 2015_P3-A6 | PHI1 | PHI3 | mother | 2015_P3-A6 | | [illegible] |
| 2015_P3-A6-1 | PHI1 | PHI3 | progeny | 2015_P3-A6 | | [illegible] |
| 2015_P3-A7 | PHI1 | PHI3 | mother | 2015_P3-A7 | | [illegible] |
| 2015_P3-A7-1 | PHI1 | PHI3 | progeny | 2015_P3-A7 | | [illegible] |
| 2015_P3-A8 | PHI1 | PHI3 | mother | 2015_P3-A8 | | [illegible] |
| 2015_P3-A8-1 | PHI1 | PHI3 | progeny | 2015_P3-A8 | | [illegible] |
| 2015_P3-A8-2 | PHI1 | PHI3 | progeny | 2015_P3-A8 | | [illegible] |
| 2015_P3-A8-3 | PHI1 | PHI3 | progeny | 2015_P3-A8 | | [illegible] |
| 2015_P3-B11 | PHI1 | PHI3 | mother | 2015_P3-B11 | | [illegible] |
| 2015_P3-B11-1 | PHI1 | PHI3 | progeny | 2015_P3-B11 | | [illegible] |
| 2015_P3-B2 | PHI1 | PHI3 | mother | 2015_P3-B2 | | [illegible] |
| 2015_P3-B2-1 | PHI1 | PHI3 | progeny | 2015_P3-B2 | | [illegible] |
| 2015_P3-B2-2 | PHI1 | PHI3 | progeny | 2015_P3-B2 | | [illegible] |
| 2015_P3-B3 | PHI1 | PHI3 | mother | 2015_P3-B3 | | [illegible] |
| 2015_P3-B3-1 | PHI1 | PHI3 | progeny | 2015_P3-B3 | | [illegible] |
| 2015_P3-B4 | PHI1 | PHI3 | mother | 2015_P3-B4 | | [illegible] |
| 2015_P3-B4-1 | PHI1 | PHI3 | progeny | 2015_P3-B4 | | [illegible] |
| 2015_P3-B5 | PHI1 | PHI3 | mother | 2015_P3-B5 | | [illegible] |
| 2015_P3-B5-1 | PHI1 | PHI3 | progeny | 2015_P3-B5 | | [illegible] |
| 2015_P3-B7 | PHI1 | PHI3 | mother | 2015_P3-B7 | | [illegible] |
| 2015_P3-B7-1 | PHI1 | PHI3 | progeny | 2015_P3-B7 | | [illegible] |
| 2015_P3-C1 | PHI1 | PHI3 | mother | 2015_P3-C1 | | [illegible] |
| 2015_P3-C1-1 | PHI1 | PHI3 | progeny | 2015_P3-C1 | | [illegible] |
| 2015_P3-C10 | PHI1 | PHI3 | mother | 2015_P3-C10 | | [illegible] |
| 2015_P3-C10-2 | PHI1 | PHI3 | progeny | 2015_P3-C10 | | [illegible] |
| 2015_P3-C11 | PHI1 | PHI3 | mother | 2015_P3-C11 | | [illegible] |
| 2015_P3-C11-3 | PHI1 | PHI3 | progeny | 2015_P3-C11 | | [illegible] |
| 2015_P3-C2 | PHI1 | PHI3 | mother | 2015_P3-C2 | | [illegible] |
| 2015_P3-C2-1 | PHI1 | PHI3 | progeny | 2015_P3-C2 | | [illegible] |
| 2015_P3-C7 | PHI1 | PHI3 | mother | 2015_P3-C7 | | [illegible] |
| 2015_P3-C7-1 | PHI1 | PHI3 | progeny | 2015_P3-C7 | | [illegible] |
| 2015_P3-C8 | PHI1 | PHI3 | mother | 2015_P3-C8 | | [illegible] |
| 2015_P3-C8-1 | PHI1 | PHI3 | progeny | 2015_P3-C8 | | [illegible] |
| 2015_P3-C8-2 | PHI1 | PHI3 | progeny | 2015_P3-C8 | | [illegible] |
| 2015_P3-C8-3 | PHI1 | PHI3 | progeny | 2015_P3-C8 | | [illegible] |
| 2015_P3-D10 | PHI1 | PHI3 | mother | 2015_P3-D10 | | [illegible] |
| 2015_P3-D10-1 | PHI1 | PHI3 | progeny | 2015_P3-D10 | | [illegible] |
| 2015_P3-D3 | PHI1 | PHI3 | mother | 2015_P3-D3 | | [illegible] |
| 2015_P3-D3-1 | PHI1 | PHI3 | progeny | 2015_P3-D3 | | [illegible] |
| 2015_P3-D3-2 | PHI1 | PHI3 | progeny | 2015_P3-D3 | | [illegible] |
| 2015_P3-D4 | PHI1 | PHI3 | mother | 2015_P3-D4 | | [illegible] |
| 2015_P3-D4-1 | PHI1 | PHI3 | progeny | 2015_P3-D4 | | [illegible] |
| 2015_P3-D4-2 | PHI1 | PHI3 | progeny | 2015_P3-D4 | | [illegible] |
| 2015_P3-D4-3 | PHI1 | PHI3 | progeny | 2015_P3-D4 | | [illegible] |
| 2015_P3-D5 | PHI1 | PHI3 | mother | 2015_P3-D5 | | [illegible] |
| 2015_P3-D5-1 | PHI1 | PHI3 | progeny | 2015_P3-D5 | | [illegible] |
| 2015_P3-D6 | PHI1 | PHI3 | mother | 2015_P3-D6 | | [illegible] |
| 2015_P3-D6-1 | PHI1 | PHI3 | progeny | 2015_P3-D6 | | [illegible] |
| 2015_P3-D7 | PHI1 | PHI3 | mother | 2015_P3-D7 | | [illegible] |
| 2015_P3-D7-1 | PHI1 | PHI3 | progeny | 2015_P3-D7 | | [illegible] |
| 2015_P3-D7-2 | PHI1 | PHI3 | progeny | 2015_P3-D7 | | [illegible] |
| 2015_P3-D8 | PHI1 | PHI3 | mother | 2015_P3-D8 | | [illegible] |
| 2015_P3-E10 | PHI1 | PHI3 | mother | 2015_P3-E10 | | [illegible] |
| 2015_P3-E10-1 | PHI1 | PHI3 | progeny | 2015_P3-E10 | | [illegible] |
| 2015_P3-E11 | PHI1 | PHI3 | mother | 2015_P3-E11 | | [illegible] |
| 2015_P3-E11-1 | PHI1 | PHI3 | progeny | 2015_P3-E11 | | [illegible] |
| 2015_P3-E12 | PHI1 | PHI3 | mother | 2015_P3-E12 | | [illegible] |
| 2015_P3-E12-1 | PHI1 | PHI3 | progeny | 2015_P3-E12 | | [illegible] |
| 2015_P3-E4 | PHI1 | PHI3 | mother | 2015_P3-E4 | | [illegible] |
| 2015_P3-E4-1 | PHI1 | PHI3 | progeny | 2015_P3-E4 | | [illegible] |
| 2015_P3-E4-2 | PHI1 | PHI3 | progeny | 2015_P3-E4 | | [illegible] |
| 2015_P3-E5 | PHI1 | PHI3 | mother | 2015_P3-E5 | | [illegible] |
| 2015_P3-E5-1 | PHI1 | PHI3 | progeny | 2015_P3-E5 | | [illegible] |
| 2015_P3-E5-2 | PHI1 | PHI3 | progeny | 2015_P3-E5 | | [illegible] |
| 2015_P3-E5-3 | PHI1 | PHI3 | progeny | 2015_P3-E5 | | [illegible] |
| 2015_P3-E7 | PHI1 | PHI3 | mother | 2015_P3-E7 | | [illegible] |
| 2015_P3-E7-1 | PHI1 | PHI3 | progeny | 2015_P3-E7 | | [illegible] |
| 2015_P3-E8 | PHI1 | PHI3 | mother | 2015_P3-E8 | | [illegible] |
| 2015_P3-E8-1 | PHI1 | PHI3 | progeny | 2015_P3-E8 | | [illegible] |
| 2015_P3-E8-2 | PHI1 | PHI3 | progeny | 2015_P3-E8 | | [illegible] |
| 2015_P3-E8-3 | PHI1 | PHI3 | progeny | 2015_P3-E8 | | [illegible] |
| 2015_P3-E9 | PHI1 | PHI3 | mother | 2015_P3-E9 | | [illegible] |
| 2015_P3-E9-1 | PHI1 | PHI3 | progeny | 2015_P3-E9 | | [illegible] |
| 2015_P3-E9-2 | PHI1 | PHI3 | progeny | 2015_P3-E9 | | [illegible] |
| 2015_P3-E9-3 | PHI1 | PHI3 | progeny | 2015_P3-E9 | | [illegible] |
| 2015_P3-E9-4 | PHI1 | PHI3 | progeny | 2015_P3-E9 | | [illegible] |
| 2015_P3-F1 | PHI1 | PHI3 | mother | 2015_P3-F1 | | [illegible] |
| 2015_P3-F1-1 | PHI1 | PHI3 | progeny | 2015_P3-F1 | | [illegible] |
| 2015_P3-F11 | PHI2 | PHI1 | mother | 2015_P3-F11 | | [illegible] |
| 2015_P3-F11-1 | PHI2 | PHI1 | progeny | 2015_P3-F11 | | [illegible] |
| 2015_P3-F12 | PHI2 | PHI1 | mother | 2015_P3-F12 | | [illegible] |
| 2015_P3-F12-1 | PHI2 | PHI1 | progeny | 2015_P3-F12 | | [illegible] |
| 2015_P3-F12-2 | PHI2 | PHI1 | progeny | 2015_P3-F12 | | [illegible] |
| 2015_P3-F12-3 | PHI2 | PHI1 | progeny | 2015_P3-F12 | | [illegible] |
| 2015_P3-F12-4 | PHI2 | PHI1 | progeny | 2015_P3-F12 | | [illegible] |
| 2015_P3-F2 | PHI1 | PHI3 | mother | 2015_P3-F2 | | [illegible] |
| 2015_P3-F2-1 | PHI1 | PHI3 | progeny | 2015_P3-F2 | | [illegible] |
| 2015_P3-F3 | PHI1 | PHI3 | mother | 2015_P3-F3 | | [illegible] |
| 2015_P3-F3-1 | PHI1 | PHI3 | progeny | 2015_P3-F3 | | [illegible] |
| 2015_P3-F3-2 | PHI1 | PHI3 | progeny | 2015_P3-F3 | | [illegible] |
| 2015_P3-F4 | PHI1 | PHI3 | mother | 2015_P3-F4 | | [illegible] |
| 2015_P3-F4-1 | PHI1 | PHI3 | progeny | 2015_P3-F4 | | [illegible] |
| 2015_P3-F4-2 | PHI1 | PHI3 | progeny | 2015_P3-F4 | | [illegible] |
| 2015_P3-F4-3 | PHI1 | PHI3 | progeny | 2015_P3-F4 | | [illegible] |
| 2015_P3-F4-4 | PHI1 | PHI3 | progeny | 2015_P3-F4 | | [illegible] |
| 2015_P3-F5 | PHI2 | PHI1 | mother | 2015_P3-F5 | | [illegible] |
| 2015_P3-F5-1 | PHI2 | PHI1 | progeny | 2015_P3-F5 | | [illegible] |
| 2015_P3-F6 | PHI2 | PHI1 | mother | 2015_P3-F6 | | [illegible] |
| 2015_P3-F6-1 | PHI2 | PHI1 | progeny | 2015_P3-F6 | | [illegible] |
| 2015_P3-F6-2 | PHI2 | PHI1 | progeny | 2015_P3-F6 | | [illegible] |

### Table S4. Results of the SNP array analasis (continue)

| plant_id | mom_gen | dad_gen | mother_progeny | family_ |
|---|---|---|---|---|
| 2015_P3-F6-3 | PHI2 | PHI1 | progeny | 2015_P3-F6 |
| 2015_P3-F7 | PHI2 | PHI1 | mother | 2015_P3-F7 |
| 2015_P3-F7-1 | PHI2 | PHI1 | progeny | 2015_P3-F7 |
| 2015_P3-F7-2 | PHI2 | PHI1 | progeny | 2015_P3-F7 |
| 2015_P3-F7-3 | PHI2 | PHI1 | progeny | 2015_P3-F7 |
| 2015_P3-F7-4 | PHI2 | PHI1 | progeny | 2015_P3-F7 |
| 2015_P3-F7-5 | PHI2 | PHI1 | progeny | 2015_P3-F7 |
| 2015_P3-F8 | PHI2 | PHI1 | mother | 2015_P3-F8 |
| 2015_P3-F8-1 | PHI2 | PHI1 | progeny | 2015_P3-F8 |
| 2015_P3-F9 | PHI2 | PHI1 | mother | 2015_P3-F9 |
| 2015_P3-F9-1 | PHI2 | PHI1 | progeny | 2015_P3-F9 |
| 2015_P3-F9-2 | PHI2 | PHI1 | progeny | 2015_P3-F9 |
| 2015_P3-F9-3 | PHI2 | PHI1 | progeny | 2015_P3-F9 |
| 2015_P3-F9-4 | PHI2 | PHI1 | progeny | 2015_P3-F9 |
| 2015_P3-F9-5 | PHI2 | PHI1 | progeny | 2015_P3-F9 |
| 2015_P3-G1 | PHI2 | PHI1 | mother | 2015_P3-G1 |
| 2015_P3-G1-1 | PHI2 | PHI1 | progeny | 2015_P3-G1 |
| 2015_P3-G1-2 | PHI2 | PHI1 | progeny | 2015_P3-G1 |
| 2015_P3-G1-3 | PHI2 | PHI1 | progeny | 2015_P3-G1 |
| 2015_P3-G1-4 | PHI2 | PHI1 | progeny | 2015_P3-G1 |
| 2015_P3-G10 | PHI2 | PHI1 | mother | 2015_P3-G10 |
| 2015_P3-G10-1 | PHI2 | PHI1 | progeny | 2015_P3-G10 |
| 2015_P3-G10-10 | PHI2 | PHI1 | progeny | 2015_P3-G10 |
| 2015_P3-G10-2 | PHI2 | PHI1 | progeny | 2015_P3-G10 |
| 2015_P3-G10-3 | PHI2 | PHI1 | progeny | 2015_P3-G10 |
| 2015_P3-G10-4 | PHI2 | PHI1 | progeny | 2015_P3-G10 |
| 2015_P3-G10-5 | PHI2 | PHI1 | progeny | 2015_P3-G10 |
| 2015_P3-G10-6 | PHI2 | PHI1 | progeny | 2015_P3-G10 |
| 2015_P3-G10-7 | PHI2 | PHI1 | progeny | 2015_P3-G10 |
| 2015_P3-G10-8 | PHI2 | PHI1 | progeny | 2015_P3-G10 |
| 2015_P3-G10-9 | PHI2 | PHI1 | progeny | 2015_P3-G10 |
| 2015_P3-G12 | PHI2 | PHI1 | mother | 2015_P3-G12 |
| 2015_P3-G12-1 | PHI2 | PHI1 | progeny | 2015_P3-G12 |
| 2015_P3-G12-2 | PHI2 | PHI1 | progeny | 2015_P3-G12 |
| 2015_P3-G12-3 | PHI2 | PHI1 | progeny | 2015_P3-G12 |
| 2015_P3-G12-4 | PHI2 | PHI1 | progeny | 2015_P3-G12 |
| 2015_P3-G2 | PHI2 | PHI1 | mother | 2015_P3-G2 |
| 2015_P3-G2-1 | PHI2 | PHI1 | progeny | 2015_P3-G2 |
| 2015_P3-G3 | PHI2 | PHI1 | mother | 2015_P3-G3 |
| 2015_P3-G3-1 | PHI2 | PHI1 | progeny | 2015_P3-G3 |
| 2015_P3-G4 | PHI2 | PHI1 | mother | 2015_P3-G4 |
| 2015_P3-G4-1 | PHI2 | PHI1 | progeny | 2015_P3-G4 |
| 2015_P3-G4-2 | PHI2 | PHI1 | progeny | 2015_P3-G4 |
| 2015_P3-G4-3 | PHI2 | PHI1 | progeny | 2015_P3-G4 |
| 2015_P3-G4-4 | PHI2 | PHI1 | progeny | 2015_P3-G4 |
| 2015_P3-G4-5 | PHI2 | PHI1 | progeny | 2015_P3-G4 |
| 2015_P3-G6 | PHI2 | PHI1 | mother | 2015_P3-G6 |
| 2015_P3-G6-1 | PHI2 | PHI1 | progeny | 2015_P3-G6 |
| 2015_P3-G6-2 | PHI2 | PHI1 | progeny | 2015_P3-G6 |
| 2015_P3-G8 | PHI2 | PHI1 | mother | 2015_P3-G8 |
| 2015_P3-G8-1 | PHI2 | PHI1 | progeny | 2015_P3-G8 |
| 2015_P3-G8-2 | PHI2 | PHI1 | progeny | 2015_P3-G8 |
| 2015_P3-G9 | PHI2 | PHI1 | mother | 2015_P3-G9 |
| 2015_P3-G9-1 | PHI2 | PHI1 | progeny | 2015_P3-G9 |
| 2015_P3-G9-10 | PHI2 | PHI1 | progeny | 2015_P3-G9 |
| 2015_P3-G9-11 | PHI2 | PHI1 | progeny | 2015_P3-G9 |
| 2015_P3-G9-12 | PHI2 | PHI1 | progeny | 2015_P3-G9 |
| 2015_P3-G9-2 | PHI2 | PHI1 | progeny | 2015_P3-G9 |
| 2015_P3-G9-3 | PHI2 | PHI1 | progeny | 2015_P3-G9 |
| 2015_P3-G9-4 | PHI2 | PHI1 | progeny | 2015_P3-G9 |
| 2015_P3-G9-5 | PHI2 | PHI1 | progeny | 2015_P3-G9 |
| 2015_P3-G9-6 | PHI2 | PHI1 | progeny | 2015_P3-G9 |
| 2015_P3-G9-7 | PHI2 | PHI1 | progeny | 2015_P3-G9 |
| 2015_P3-G9-8 | PHI2 | PHI1 | progeny | 2015_P3-G9 |
| 2015_P3-G9-9 | PHI2 | PHI1 | progeny | 2015_P3-G9 |
| 2015_P3-H10 | PHI2 | PHI1 | mother | 2015_P3-H10 |
| 2015_P3-H10-1 | PHI2 | PHI1 | progeny | 2015_P3-H10 |
| 2015_P3-H10-2 | PHI2 | PHI1 | progeny | 2015_P3-H10 |
| 2015_P3-H10-3 | PHI2 | PHI1 | progeny | 2015_P3-H10 |
| 2015_P3-H10-4 | PHI2 | PHI1 | progeny | 2015_P3-H10 |
| 2015_P3-H10-5 | PHI2 | PHI1 | progeny | 2015_P3-H10 |
| 2015_P3-H10-6 | PHI2 | PHI1 | progeny | 2015_P3-H10 |
| 2015_P3-H10-7 | PHI2 | PHI1 | progeny | 2015_P3-H10 |
| 2015_P3-H11 | PHI2 | PHI1 | mother | 2015_P3-H11 |
| 2015_P3-H11-1 | PHI2 | PHI1 | progeny | 2015_P3-H11 |
| 2015_P3-H3 | PHI2 | PHI1 | mother | 2015_P3-H3 |
| 2015_P3-H3-1 | PHI2 | PHI1 | progeny | 2015_P3-H3 |
| 2015_P3-H3-2 | PHI2 | PHI1 | progeny | 2015_P3-H3 |
| 2015_P3-H5 | PHI2 | PHI1 | mother | 2015_P3-H5 |
| 2015_P3-H5-1 | PHI2 | PHI1 | progeny | 2015_P3-H5 |
| 2015_P3-H5-2 | PHI2 | PHI1 | progeny | 2015_P3-H5 |
| 2015_P3-H5-3 | PHI2 | PHI1 | progeny | 2015_P3-H5 |
| 2015_P3-H5-4 | PHI2 | PHI1 | progeny | 2015_P3-H5 |
| 2015_P3-H6 | PHI2 | PHI1 | mother | 2015_P3-H6 |
| 2015_P3-H6-1 | PHI2 | PHI1 | progeny | 2015_P3-H6 |
| 2015_P3-H7 | PHI2 | PHI1 | mother | 2015_P3-H7 |
| 2015_P3-H7-1 | PHI2 | PHI1 | progeny | 2015_P3-H7 |
| 2015_P3-H8 | PHI2 | PHI1 | mother | 2015_P3-H8 |
| 2015_P3-H8-1 | PHI2 | PHI1 | progeny | 2015_P3-H8 |
| 2015_P3-H9 | PHI2 | PHI1 | mother | 2015_P3-H9 |
| 2015_P3-H9-1 | PHI2 | PHI1 | progeny | 2015_P3-H9 |
| 2015_P3-H9-2 | PHI2 | PHI1 | progeny | 2015_P3-H9 |
| 2015_P3-H9-3 | PHI2 | PHI1 | progeny | 2015_P3-H9 |
| 2015_P3-H9-4 | PHI2 | PHI1 | progeny | 2015_P3-H9 |
| 2015_P3-H9-5 | PHI2 | PHI1 | progeny | 2015_P3-H9 |
| 2015_P3-H9-6 | PHI2 | PHI1 | progeny | 2015_P3-H9 |
| 2015_P4-A1 | PHI2 | PHI1 | mother | 2015_P4-A1 |
| 2015_P4-A1-1 | PHI2 | PHI1 | progeny | 2015_P4-A1 |
| 2015_P4-A1-2 | PHI2 | PHI1 | progeny | 2015_P4-A1 |
| 2015_P4-A2 | PHI2 | PHI1 | mother | 2015_P4-A2 |
| 2015_P4-A2-1 | PHI2 | PHI1 | progeny | 2015_P4-A2 |
| 2015_P4-A3 | PHI2 | PHI1 | mother | 2015_P4-A3 |
| 2015_P4-A3-1 | PHI2 | PHI1 | progeny | 2015_P4-A3 |
| 2015_P4-A4 | PHI2 | PHI1 | mother | 2015_P4-A4 |
| 2015_P4-A4-1 | PHI2 | PHI1 | progeny | 2015_P4-A4 |
| 2015_P4-A4-2 | PHI2 | PHI1 | progeny | 2015_P4-A4 |
| 2015_P4-A4-3 | PHI2 | PHI1 | progeny | 2015_P4-A4 |
| 2015_P4-A4-4 | PHI2 | PHI1 | progeny | 2015_P4-A4 |
| 2015_P4-A5 | PHI2 | PHI1 | mother | 2015_P4-A5 |
| 2015_P4-A5-1 | PHI2 | PHI1 | progeny | 2015_P4-A5 |
| 2015_P4-A5-2 | PHI2 | PHI1 | progeny | 2015_P4-A5 |
| 2015_P4-A5-3 | PHI2 | PHI1 | progeny | 2015_P4-A5 |
| 2015_P4-A6 | PHI2 | PHI1 | mother | 2015_P4-A6 |
| 2015_P4-A6-1 | PHI2 | PHI1 | progeny | 2015_P4-A6 |
| 2015_P4-A6-2 | PHI2 | PHI1 | progeny | 2015_P4-A6 |
| 2015_P4-A6-3 | PHI2 | PHI1 | progeny | 2015_P4-A6 |
| 2015_P4-A6-4 | PHI2 | PHI1 | progeny | 2015_P4-A6 |
| 2015_P4-A7 | PHI2 | PHI1 | mother | 2015_P4-A7 |
| 2015_P4-A7-1 | PHI2 | PHI1 | progeny | 2015_P4-A7 |

| Chr | 1 | 1 | 1 | 1 | 1 | 1 | 1 | 1 | 1 | 1 | 1 | 1 | 1 | 1 | 2 | 2 | 2 | 2 | 2 | 2 | 2 | 3 | 3 | 3 | 3 | 3 | 3 | 3 | 3 | 3 | 3 | 3 | 3 | 3 | 3 | 4 | 4 | 4 | 4 | 4 | 4 |
|---|---|---|---|---|---|---|---|---|---|---|---|---|---|---|---|---|---|---|---|---|---|---|---|---|---|---|---|---|---|---|---|---|---|---|---|---|---|---|---|---|---|
| Position, cM | 48 | 59 | 80 | 85 | 97 | 113 | 116 | 137 | 137 | 143 | 162 | 197 | 218 | 285 | 25 | 33 | 142 | 142 | 162 | 193 | 202 | 20 | 39 | 75 | 75 | 75 | 102 | 104 | 127 | 140 | 199 | 213 | 228 | 237 | 246 | 30 | 73 | 83 | 91 | 128 | 146 |
| 2015_P3-F6-3 | H | T | T | G | C | G | G | C | A | C | G | H | A | H | G | H | H | H | C | A | T | T | T | C | C | C | H | H | G | H | G | C | C | A | G | C | H | T | T | A | T |
| 2015_P3-F7 | H | H | T | G | C | G | G | C | A | C | G | H | A | H | G | H | H | H | C | A | T | T | T | C | C | C | H | H | G | H | G | C | C | A | H | G | H | T | H | H | T |
| 2015_P3-F7-1 | H | H | T | G | C | G | G | C | A | C | G | A | A | A | G | H | H | H | C | A | T | T | T | C | C | C | H | H | G | H | G | C | C | A | H | G | H | T | H | H | T |
| 2015_P3-F7-2 | H | H | T | G | C | G | G | C | A | C | G | A | A | A | G | T | T | A | C | A | T | T | T | C | C | C | T | A | G | H | G | C | C | A | H | G | H | T | H | G | T |
| 2015_P3-F7-3 | H | C | T | G | C | G | G | C | A | C | G | G | A | H | G | H | T | A | C | A | T | T | T | C | C | C | C | T | G | H | G | C | C | A | G | G | H | T | C | G | T |
| 2015_P3-F7-4 | A | C | T | G | C | G | G | C | A | C | G | A | A | H | G | H | G | G | C | A | T | T | T | C | C | C | C | T | G | T | G | C | C | A | G | G | G | T | C | H | T |
| 2015_P3-F7-5 | H | H | T | G | C | G | G | C | A | C | G | H | A | G | G | H | H | H | C | A | T | T | T | C | C | C | H | H | G | H | G | C | C | A | H | G | H | T | H | H | T |

| Chr | 4 | 4 | 5 | 5 | 5 | 5 | 5 | 5 | 5 | 5 | 5 | 5 | 5 | 5 | 5 | 6 | 6 | 6 | 6 | 6 | 7 | 7 | 7 | 7 | 7 | 7 | 7 | 7 | 7 | 7 | 7 | 7 | 7 | 7 | 7 | 7 | 8 | 8 | 8 | 8 | 8 | 8 | 8 | 8 | 8 | 9 | 9 | 9 | 9 | 9 | 10 | 10 | 10 | 10 | 10 |
|---|---|---|---|---|---|---|---|---|---|---|---|---|---|---|---|---|---|---|---|---|---|---|---|---|---|---|---|---|---|---|---|---|---|---|---|---|---|---|---|---|---|---|---|---|---|---|---|---|---|---|---|---|---|---|---|
| Position, cM | 166 | 197 | 20 | 65 | 71 | 77 | 102 | 107 | 108 | 132 | 154 | 164 | 191 | 208 | 218 | 27 | 34 | 86 | 146 | 150 | 5.6 | 37 | 52 | 57 | 74 | 78 | 79 | 80 | 86 | 93 | 109 | 112 | 166 | 173 | 182 | 192 | 50 | 131 | 135 | 140 | 149 | 153 | 164 | 186 | 191 | 60 | 76 | 78 | 144 | 164 | 77 | 89 | 101 | 148 | 154 |
| 2015_P3-F6-3 | -9 | A | H | H | H | C | G | T | T | C | H | H | T | G | G | -9 | G | A | A | G | G | C | A | H | T | H | H | H | A | G | G | H | G | T | C | A | T | A | A | T | C | T | A | C | G | A | C | G | H | H | T | H | T | T | A |
| 2015_P3-F7 | -9 | A | H | H | H | C | G | H | H | C | H | H | T | G | G | G | G | A | A | G | G | C | A | H | T | H | H | H | A | G | G | H | G | T | C | A | T | H | A | H | H | T | A | -9 | H | A | C | H | H | H | T | H | T | T | H |
| 2015_P3-F7-1 | -9 | A | H | H | H | C | G | H | H | C | H | H | T | G | G | G | G | A | A | G | G | C | A | H | T | H | H | H | A | G | G | H | G | T | C | A | T | H | A | H | H | T | A | C | H | A | C | H | H | H | T | H | T | T | H |
| 2015_P3-F7-2 | -9 | A | C | A | G | C | G | C | C | C | H | H | T | G | G | G | G | A | A | G | G | C | A | H | T | C | A | A | A | G | G | H | G | T | C | A | T | H | A | H | H | T | A | C | H | A | C | T | G | A | T | A | T | T | H |
| 2015_P3-F7-3 | -9 | A | C | A | G | C | G | C | C | C | G | G | T | G | G | G | G | A | A | G | G | C | A | T | T | C | A | A | A | G | G | A | G | T | C | A | T | H | A | H | H | T | A | C | G | A | C | T | H | A | T | H | T | T | H |
| 2015_P3-F7-4 | -9 | A | H | H | -9 | C | G | -9 | H | C | H | H | T | G | G | G | G | A | A | G | G | C | A | T | T | C | A | A | A | G | G | A | G | T | C | A | T | A | A | T | C | T | A | C | G | A | C | G | C | G | T | H | T | T | H |
| 2015_P3-F7-5 | -9 | A | H | H | H | C | G | H | H | C | H | H | T | G | G | G | G | A | A | G | G | C | A | H | T | H | H | H | A | G | G | H | G | T | C | A | T | H | A | H | H | T | -9 | -9 | -9 | A | -9 | H | H | H | T | H | T | T | H |

### Table S4. Results of the SNP array analasis (continue)

| | | | | | Chr | 1 | 1 | 1 | 1 | 1 | 1 | 1 | 1 | 1 | 1 | 1 | 1 | 1 | 1 | 2 | 2 | 2 | 2 | 2 | 2 | 2 | 3 | 3 | 3 | 3 | 3 | 3 | 3 | 3 | 3 | 3 | 3 | 3 | 3 | 3 | 4 | 4 | 4 | 4 | 4 | 4 | 4 | 4 | 5 | 5 | 5 | 5 | 5 | 5 | 5 | 5 | 5 | 5 | 5 | 5 | 5 | 6 | 6 | 6 | 6 | 6 | 7 | 7 | 7 | 7 | 7 | 7 | 7 | 7 | 7 | 7 | 7 | 7 | 7 | 7 | 7 | 7 | 8 | 8 | 8 | 8 | 8 | 8 | 8 | 8 | 8 | 9 | 9 | 9 | 9 | 9 | 10 | 10 | 10 | 10 | 10 |
|---|---|---|---|---|---|---|---|---|---|---|---|---|---|---|---|---|---|---|---|---|---|---|---|---|---|---|---|---|---|---|---|---|---|---|---|---|---|---|---|---|---|---|---|---|---|---|---|---|---|---|---|---|---|---|---|---|---|---|---|---|---|---|---|---|---|---|---|---|---|---|---|---|---|---|---|---|---|---|---|---|---|---|---|---|---|---|---|---|---|---|---|---|---|---|---|---|---|---|---|---|---|
| | | | | | Position, cM | 48 | 59 | 80 | 85 | 97 | 113 | 116 | 137 | 137 | 143 | 162 | 197 | 218 | 285 | 25 | 33 | 142 | 142 | 162 | 193 | 202 | 20 | 39 | 75 | 75 | 75 | 102 | 104 | 127 | 140 | 199 | 213 | 228 | 237 | 246 | 30 | 73 | 83 | 91 | 128 | 146 | 166 | 197 | 20 | 65 | 71 | 77 | 102 | 107 | 108 | 132 | 154 | 164 | 191 | 208 | 218 | 27 | 34 | 86 | 146 | 150 | 5.6 | 37 | 52 | 57 | 74 | 78 | 79 | 80 | 86 | 93 | 109 | 112 | 166 | 173 | 182 | 192 | 50 | 131 | 135 | 140 | 149 | 153 | 164 | 186 | 191 | 60 | 76 | 78 | 144 | 164 | 77 | 89 | 101 | 148 | 154 |
| plant_id | mom_gen | dad_gen | mother_progen | family_ | | | | | | | | | | | | | | | | | | | | | | | | | | | | | | | | | | | | | | | | | | | | | | | | | | | | | | | | | | | | | | | | | | | | | | | | | | | | | | | | | | | | | | | | | | | | | | | | | | |
| 2015_P4-A9 | PHI2 | PHI1 | mother | 2015_P4-A9 | | H | H | T | G | C | G | G | C | A | C | G | H | A | H | G | H | H | H | C | A | T | T | H | H | H | C | H | H | G | T | G | C | C | A | H | G | C | T | T | A | T | -9 | A | H | H | H | C | G | T | T | C | H | H | T | G | G | G | G | H | A | G | G | C | A | H | T | H | H | H | A | G | G | H | G | T | C | A | T | A | A | H | H | T | A | C | H | A | C | H | H | H | T | H | T | T | H |
| 2015_P4-A9-1 | PHI2 | PHI1 | progeny | 2015_P4-A9 | | H | H | T | G | C | G | G | C | A | C | G | H | A | A | G | H | H | H | C | A | -9 | T | H | H | H | C | H | H | G | T | G | C | C | A | T | G | C | T | T | A | T | -9 | A | T | G | A | C | G | T | T | C | G | G | T | G | G | G | G | H | A | G | G | C | A | H | T | H | H | H | A | G | G | H | G | T | C | A | T | A | A | H | H | T | A | C | G | A | C | G | H | G | T | H | T | T | H |
| 2015_P4-A9-2 | PHI2 | PHI1 | progeny | 2015_P4-A9 | | H | H | T | G | C | G | G | C | A | C | G | H | A | H | G | C | H | H | C | A | T | T | H | H | H | C | H | H | G | T | G | C | C | A | H | G | C | T | T | A | T | -9 | A | H | H | H | C | G | T | T | C | G | H | T | G | G | G | G | G | A | G | G | C | A | H | T | H | H | H | A | G | G | H | G | T | C | A | T | A | A | T | C | T | A | C | T | A | C | T | G | H | T | H | T | T | H |
| 2015_P4-A9-3 | PHI2 | PHI1 | progeny | 2015_P4-A9 | | H | H | T | G | C | G | G | C | A | C | G | H | A | H | G | H | H | H | C | A | T | T | H | H | H | C | -9 | H | G | T | G | C | C | A | -9 | G | C | T | T | A | T | -9 | A | H | H | H | C | G | T | T | C | H | H | T | G | G | G | G | H | A | G | G | C | A | H | T | H | H | H | A | G | G | H | G | T | C | A | T | A | A | H | H | T | A | C | H | A | C | H | H | H | T | H | T | T | H |
| 2015_P4-A9-4 | PHI2 | PHI1 | progeny | 2015_P4-A9 | | H | H | T | G | C | G | G | C | A | C | G | H | A | -9 | G | -9 | H | H | C | A | T | T | A | H | -9 | C | H | H | G | T | G | C | C | A | H | G | C | T | T | A | T | -9 | A | H | H | H | C | G | T | T | C | G | H | T | G | G | G | G | H | A | G | G | C | A | H | T | H | H | -9 | A | G | G | H | G | T | C | A | T | A | A | T | C | T | A | C | H | A | C | H | H | H | T | H | T | T | H |
| 2015_P4-B11 | PHI2 | PHI3 | mother | 2015_P4-B11 | | H | H | H | H | C | H | G | C | A | C | G | G | A | H | H | T | H | A | C | A | T | T | H | C | C | H | C | T | H | T | G | H | H | A | H | G | H | H | H | H | H | T | A | H | G | H | H | H | H | T | H | G | H | T | G | G | H | G | H | H | T | H | H | H | C | H | G | G | G | A | H | H | C | H | T | C | A | T | H | A | T | C | H | H | H | G | H | H | G | H | H | H | H | T | T | H |
| 2015_P4-B11-1 | PHI2 | PHI3 | progeny | 2015_P4-B11 | | H | H | -9 | A | C | H | G | C | A | C | G | -9 | -9 | H | G | T | T | -9 | -9 | A | T | T | T | C | C | -9 | -9 | T | G | T | -9 | H | H | A | T | G | -9 | T | T | H | H | -9 | A | C | G | -9 | C | C | C | T | T | G | H | T | G | G | H | G | H | H | T | H | H | H | C | T | G | -9 | H | H | G | H | C | T | H | C | A | T | H | A | T | H | H | H | H | -9 | G | T | G | C | G | -9 | -9 | -9 | T | H |
| 2015_P4-B11-3 | PHI2 | PHI3 | progeny | 2015_P4-B11 | | H | H | H | H | C | H | G | C | A | C | G | G | A | H | G | T | H | A | C | A | T | T | T | C | C | C | C | T | H | T | G | A | H | A | G | G | H | H | H | A | T | -9 | A | C | G | G | C | H | H | T | H | G | H | T | G | G | T | G | H | H | T | H | H | H | C | H | G | G | G | A | H | H | C | H | T | C | A | T | H | A | T | C | H | H | H | G | G | H | G | H | H | H | H | T | T | H |
| 2015_P4-B3 | PHI2 | PHI1 | mother | 2015_P4-B3 | | H | T | T | G | C | G | G | C | A | C | G | H | A | H | G | H | H | H | C | A | T | T | -9 | H | H | C | H | H | G | T | G | C | C | A | H | G | C | T | T | -9 | T | -9 | A | H | H | H | C | G | T | T | C | H | H | T | G | G | G | G | A | A | G | G | C | A | H | T | H | H | H | A | G | G | H | G | T | C | A | T | A | A | H | H | T | A | C | H | A | C | H | H | H | T | H | T | T | H |
| 2015_P4-B3-1 | PHI2 | PHI1 | progeny | 2015_P4-B3 | | A | T | T | G | C | G | G | C | A | C | G | A | A | G | G | H | H | H | C | A | T | T | H | C | C | C | -9 | T | G | T | G | C | C | A | H | G | C | T | T | A | T | -9 | A | H | A | G | C | G | T | T | C | H | A | T | G | G | G | G | A | A | G | G | C | A | H | T | H | H | H | A | G | G | A | G | T | C | A | T | A | A | H | H | T | A | C | H | A | C | H | H | H | T | H | T | T | H |
| 2015_P4-B3-2 | PHI2 | PHI1 | progeny | 2015_P4-B3 | | A | T | T | G | C | G | G | C | A | C | G | H | A | H | G | H | H | H | C | A | T | T | A | T | G | C | T | A | G | T | G | C | C | A | H | G | C | T | T | A | T | -9 | A | H | H | H | C | G | T | T | C | H | H | T | G | G | G | G | A | A | G | G | C | A | H | T | H | H | H | A | G | G | H | G | T | C | A | T | A | A | C | T | T | A | C | H | A | C | G | H | H | T | H | T | T | H |
| 2015_P4-B3-3 | PHI2 | PHI1 | progeny | 2015_P4-B3 | | T | T | T | G | C | G | G | C | A | C | G | A | A | A | G | H | H | H | C | A | T | T | H | H | H | C | T | A | G | T | G | C | C | -9 | H | G | C | T | T | A | T | -9 | A | H | H | H | C | G | T | T | C | A | A | T | G | G | G | G | A | A | -9 | G | C | A | H | T | H | H | G | A | G | G | H | G | T | C | A | T | A | A | H | H | T | -9 | -9 | H | A | C | G | C | G | T | H | -9 | T | A |
| 2015_P4-B4 | PHI2 | PHI1 | mother | 2015_P4-B4 | | H | T | T | G | C | G | G | C | A | C | G | H | A | H | G | H | H | H | C | A | T | T | T | C | C | C | T | H | G | H | G | C | C | A | G | H | C | T | H | H | T | G | -9 | H | H | H | C | G | T | T | C | H | H | T | G | G | G | G | A | A | G | G | C | A | H | T | H | H | H | A | G | G | H | H | T | C | A | T | H | A | H | H | T | A | C | H | A | C | G | H | H | T | H | T | T | H |
| 2015_P4-B4-1 | PHI2 | PHI1 | progeny | 2015_P4-B4 | | H | T | T | G | C | G | G | C | A | C | G | A | A | H | G | H | G | G | C | A | T | T | T | C | C | C | T | A | G | A | G | C | C | A | G | H | C | T | T | A | T | -9 | -9 | H | G | A | C | G | T | T | C | H | H | T | G | G | G | G | A | A | G | G | C | A | C | T | G | -9 | -9 | A | G | G | C | H | T | C | A | T | C | A | C | T | T | A | C | H | A | C | G | H | H | T | H | T | T | A |
| 2015_P4-B4-2 | PHI2 | PHI1 | progeny | 2015_P4-B4 | | A | T | T | G | C | G | G | C | A | C | G | H | A | H | G | H | T | A | C | A | T | T | T | C | C | C | C | T | G | H | G | C | C | A | G | C | C | T | C | G | T | G | A | H | H | H | C | G | T | T | C | H | H | T | G | G | G | G | A | A | G | G | C | A | T | T | C | A | A | A | G | G | A | H | T | C | A | T | H | A | H | H | T | A | C | G | A | C | G | G | A | T | H | T | T | G |
| 2015_P4-B6 | PHI2 | PHI1 | mother | 2015_P4-B6 | | H | H | T | G | C | G | G | C | A | C | G | H | A | H | G | H | H | H | C | A | T | T | H | H | H | C | H | H | -9 | T | G | C | C | A | H | G | C | T | T | A | T | -9 | H | H | H | H | C | G | T | T | C | H | H | T | G | G | G | G | H | A | G | G | C | A | H | T | H | H | H | A | G | G | H | G | T | C | A | T | A | A | H | H | T | A | C | H | A | C | H | H | H | T | H | T | T | H |
| 2015_P4-B6-1 | PHI2 | PHI1 | progeny | 2015_P4-B6 | | H | H | T | G | C | G | G | C | A | C | G | H | A | H | G | C | G | G | C | A | T | T | H | T | G | C | T | A | G | T | G | C | C | A | G | G | C | T | T | A | T | -9 | H | H | H | H | C | G | T | T | C | H | G | T | G | G | G | G | H | A | G | G | C | A | H | T | H | H | H | A | G | G | H | G | T | C | A | T | A | A | T | C | T | A | C | G | A | C | T | H | H | T | H | T | T | H |
| 2015_P4-B6-2 | PHI2 | PHI1 | progeny | 2015_P4-B6 | | H | H | T | G | C | G | G | C | A | C | G | H | A | H | G | H | H | H | C | A | T | T | H | H | H | C | H | H | G | T | G | C | C | A | H | G | C | T | T | A | T | -9 | H | H | H | H | C | G | T | T | C | H | H | T | G | G | G | G | H | A | G | G | C | A | H | T | H | H | H | A | G | G | H | G | T | C | A | T | A | A | H | H | T | A | C | H | A | C | H | H | G | T | H | T | T | A |
| 2015_P4-B7 | PHI2 | PHI3 | mother | 2015_P4-B7 | | H | H | H | H | C | H | G | C | A | C | G | G | A | H | H | T | H | A | C | A | T | T | H | C | C | H | C | T | H | T | G | H | H | A | H | G | H | H | H | H | H | T | A | H | G | H | C | H | H | T | H | G | H | T | G | G | H | G | H | H | T | H | H | H | C | H | G | G | G | A | H | H | C | H | T | C | H | T | H | A | T | C | T | A | H | G | H | H | G | H | H | T | G | T | T | H |
| 2015_P4-B7-1 | PHI2 | PHI3 | progeny | 2015_P4-B7 | | H | H | H | H | C | H | G | C | A | C | G | G | A | H | G | T | G | A | C | A | T | T | H | C | C | C | C | T | G | T | G | H | H | A | H | G | C | T | T | H | H | T | A | H | G | H | C | C | C | T | T | G | H | T | G | G | G | G | A | H | T | H | H | H | C | C | G | G | G | A | H | H | C | H | T | C | H | T | C | A | T | C | T | A | H | G | G | T | G | H | H | T | G | T | T | H |
| 2015_P4-B7-2 | PHI2 | PHI3 | progeny | 2015_P4-B7 | | H | H | H | H | C | H | G | C | A | C | G | G | A | H | H | T | H | A | C | A | T | T | T | C | C | C | C | T | -9 | T | G | C | C | A | G | G | H | H | H | H | H | T | A | H | G | H | C | H | H | T | H | G | H | T | G | G | H | G | H | H | T | H | H | H | C | H | G | G | G | A | H | H | C | H | T | C | H | T | H | A | T | -9 | T | A | H | G | H | H | G | H | H | T | G | T | T | H |
| 2015_P4-B9 | PHI2 | PHI3 | mother | 2015_P4-B9 | | H | H | H | H | C | H | G | C | A | C | G | G | A | H | H | T | H | A | C | A | T | T | H | C | C | H | C | T | H | T | G | H | H | A | H | G | H | H | H | H | H | T | A | H | G | H | C | H | H | T | H | G | H | T | G | G | H | G | H | H | T | H | H | H | C | H | G | G | G | A | H | H | C | H | T | C | A | T | H | A | T | C | H | H | H | G | H | H | G | H | H | H | H | T | T | H |
| 2015_P4-B9-1 | PHI2 | PHI3 | progeny | 2015_P4-B9 | | H | H | H | H | C | H | G | C | A | C | G | G | A | H | H | T | H | A | C | A | T | T | H | C | C | H | C | T | H | T | G | H | H | A | H | G | H | H | H | H | H | T | A | H | G | H | C | H | H | T | H | G | H | T | G | G | T | -9 | H | H | T | H | H | H | C | H | G | G | G | A | H | H | C | H | T | C | A | T | H | A | T | C | H | H | H | G | H | H | G | H | H | H | H | T | T | H |
| 2015_P4-B9-2 | PHI2 | PHI3 | progeny | 2015_P4-B9 | | H | H | H | H | C | H | G | C | A | C | G | G | A | G | H | T | H | A | C | A | T | T | H | C | C | H | C | T | H | T | G | H | H | A | H | G | H | H | H | H | H | T | A | H | G | H | C | H | H | T | H | G | H | T | G | G | H | G | H | H | T | H | H | H | C | H | G | G | G | A | H | H | C | H | T | C | A | T | H | A | T | C | H | H | H | G | H | H | G | H | H | H | H | T | T | H |
| 2015_P4-C1 | PHI2 | PHI3 | mother | 2015_P4-C1 | | H | H | H | H | C | H | G | C | A | C | G | G | A | H | H | T | H | A | C | A | T | T | H | C | C | H | C | T | H | T | G | H | H | A | H | G | H | H | H | H | H | T | A | H | G | H | C | -9 | H | T | H | G | H | T | G | G | H | G | H | H | T | H | H | H | C | H | G | G | G | A | H | H | C | H | T | C | H | T | H | A | T | C | H | H | H | G | A | C | G | H | H | T | G | H | T | H |
| 2015_P4-C1-1 | PHI2 | PHI3 | progeny | 2015_P4-C1 | | H | H | H | H | C | H | G | C | A | C | G | G | A | H | A | T | H | A | C | A | T | T | H | C | C | H | C | T | H | T | G | H | H | A | H | G | H | H | H | H | H | T | A | T | G | A | C | H | H | T | H | G | H | T | G | G | H | G | H | H | T | H | H | H | C | T | G | G | G | A | G | G | C | H | T | C | H | T | H | A | T | C | H | H | T | G | A | C | G | H | H | T | G | C | T | H |
| 2015_P4-C1-2 | PHI2 | PHI3 | progeny | 2015_P4-C1 | | H | H | H | H | C | H | G | C | A | C | G | -9 | A | H | H | T | H | A | C | -9 | T | T | H | C | C | C | C | T | H | -9 | G | H | H | A | H | G | H | H | H | H | H | T | A | H | G | H | C | C | C | T | T | G | A | T | G | G | H | G | H | H | T | H | H | H | C | H | G | G | G | A | H | H | C | H | T | C | H | T | H | A | T | C | H | H | -9 | G | A | C | G | H | H | T | G | C | T | A |
| 2015_P4-C10 | PHI2 | PHI3 | mother | 2015_P4-C10 | | H | H | H | H | C | H | G | C | A | C | G | G | A | H | H | T | H | A | C | A | T | -9 | H | C | C | H | C | T | -9 | T | G | H | -9 | A | H | G | H | H | H | H | H | T | A | H | G | H | C | H | H | T | H | G | H | T | G | -9 | H | G | H | H | T | H | H | H | C | -9 | G | G | G | A | H | H | C | H | T | C | A | T | H | A | T | C | H | [illegible] | -9 | G | A | C | G | H | H | T | G | H | T | H |
| 2015_P4-C10-1 | PHI2 | PHI3 | progeny | 2015_P4-C10 | | H | H | H | H | C | H | G | C | A | C | G | G | A | H | G | T | H | A | C | A | T | T | T | C | C | H | C | T | H | T | G | H | H | A | T | G | C | T | H | H | H | T | A | H | G | H | C | H | H | T | H | G | H | T | G | G | H | G | H | G | T | G | H | H | C | H | G | G | G | A | H | H | C | H | T | C | A | T | H | A | T | C | H | A | C | G | A | C | G | H | H | T | G | H | T | H |
| 2015_P4-C11 | PHI2 | PHI3 | mother | 2015_P4-C11 | | H | H | -9 | H | C | H | G | C | A | C | G | G | A | H | H | T | H | A | C | A | T | T | H | C | C | H | C | T | H | T | G | H | H | A | H | G | H | H | H | H | H | T | A | H | G | H | C | H | H | T | H | G | H | T | G | G | H | G | H | H | T | H | H | H | C | H | G | G | G | A | H | H | C | H | T | C | A | T | H | A | T | C | T | A | H | G | H | H | G | H | H | T | G | H | T | H |
| 2015_P4-C11-1 | PHI2 | PHI3 | progeny | 2015_P4-C11 | | H | H | H | H | C | H | G | C | A | C | G | G | A | H | G | T | T | A | C | A | T | T | H | C | C | H | -9 | T | H | T | G | H | H | A | H | G | G | C | C | G | C | T | A | H | G | H | C | H | H | T | H | G | H | T | G | G | T | -9 | G | G | T | H | H | G | C | C | G | G | G | A | A | C | C | T | T | C | A | T | H | A | T | C | T | A | H | G | H | H | G | H | H | T | G | H | T | H |
| 2015_P4-C11-2 | PHI2 | PHI3 | progeny | 2015_P4-C11 | | H | H | H | H | C | G | G | C | A | C | G | G | A | G | A | T | G | A | C | A | T | T | H | C | C | H | C | T | -9 | -9 | G | H | H | A | H | G | H | H | H | H | H | T | A | H | G | H | C | H | H | T | H | G | H | T | G | G | T | -9 | G | G | T | H | H | H | C | H | G | G | G | A | H | H | C | H | T | C | A | T | H | A | T | C | T | A | H | G | G | T | G | G | A | T | -9 | H | T | H |
| 2015_P4-C2 | PHI2 | PHI3 | mother | 2015_P4-C2 | | H | H | H | H | C | H | G | C | A | C | G | G | A | H | H | T | H | A | C | A | T | T | H | C | C | H | C | T | H | T | G | H | H | A | H | G | H | H | H | H | H | T | A | H | G | H | C | H | H | T | H | G | H | T | G | G | H | G | H | H | T | H | H | H | C | H | G | G | G | A | H | H | C | H | T | C | H | T | H | A | T | C | H | H | H | G | A | C | G | H | H | T | G | H | T | H |
| 2015_P4-C2-1 | PHI2 | PHI3 | progeny | 2015_P4-C2 | | A | C | H | H | C | H | G | C | A | C | G | G | A | H | H | T | G | A | C | A | T | T | T | C | C | C | C | T | G | -9 | G | C | C | A | H | G | C | T | T | A | T | -9 | A | C | G | H | C | G | T | T | C | G | G | T | G | G | H | G | H | H | T | G | H | H | C | T | G | G | G | A | G | G | C | H | T | C | H | T | A | A | T | C | H | H | -9 | G | A | C | G | C | H | T | G | H | T | H |
| 2015_P4-C8 | PHI2 | PHI3 | mother | 2015_P4-C8 | | H | H | H | H | C | H | G | C | A | C | G | G | A | H | H | T | H | A | C | A | T | T | H | C | C | H | C | T | H | T | G | H | H | A | H | G | H | H | H | H | H | T | A | H | G | H | H | H | H | T | H | G | H | T | G | G | H | G | H | H | T | H | H | H | C | H | G | G | G | A | H | H | C | H | T | C | H | T | H | A | -9 | C | T | A | H | G | A | C | G | H | H | H | G | H | T | H |
| 2015_P4-C8-1 | PHI2 | PHI3 | progeny | 2015_P4-C8 | | H | H | H | H | C | H | G | C | A | C | G | G | A | H | H | T | H | A | C | A | T | T | H | C | C | H | C | T | H | T | G | H | H | A | H | G | H | H | H | H | H | T | A | H | G | A | H | G | T | T | C | G | H | T | G | G | G | G | H | H | T | H | H | H | C | H | G | G | G | A | H | H | C | H | T | C | H | T | H | A | T | C | T | A | T | G | A | C | G | C | H | H | G | C | T | H |
| 2015_P4-C8-2 | PHI2 | PHI3 | progeny | 2015_P4-C8 | | H | H | H | A | C | T | G | C | A | C | G | G | A | A | A | T | G | A | C | A | T | T | H | C | C | C | C | T | G | T | G | H | H | A | H | G | H | C | C | G | C | T | A | H | G | H | H | C | C | T | T | G | H | T | G | G | G | G | H | G | -9 | H | H | H | C | T | G | G | G | A | G | H | C | T | T | C | G | T | C | A | T | C | T | A | H | G | A | C | G | H | H | C | G | C | T | H |
| 2015_P4-C9 | PHI2 | PHI3 | mother | 2015_P4-C9 | | H | H | H | H | C | H | G | C | A | C | G | G | A | H | H | T | H | A | C | A | T | T | H | C | C | H | C | T | H | T | G | H | H | A | H | G | H | H | H | H | H | T | A | H | G | H | C | H | H | T | H | G | H | T | G | G | H | G | H | H | T | H | H | H | C | H | G | G | G | A | H | H | C | H | T | C | A | T | H | A | T | C | T | A | H | G | H | H | G | H | H | T | G | H | T | H |
| 2015_P4-C9-1 | PHI2 | PHI3 | progeny | 2015_P4-C9 | | H | H | C | A | C | T | G | C | A | C | G | G | A | H | A | T | T | A | C | A | T | T | H | C | C | T | C | T | H | T | G | H | H | A | H | G | C | T | T | A | T | T | A | H | G | A | H | G | T | T | C | -9 | G | T | G | G | G | G | A | H | T | H | A | G | C | C | G | G | G | A | A | C | C | H | T | C | A | T | C | A | T | C | T | -9 | H | G | G | T | G | G | H | T | G | C | T | H |
| 2015_P4-C9-2 | PHI2 | PHI3 | progeny | 2015_P4-C9 | | H | H | H | H | C | H | G | C | A | C | G | G | A | H | H | T | H | A | C | A | T | T | A | C | C | T | C | T | H | T | G | C | C | A | G | G | H | H | H | A | T | -9 | A | H | G | H | C | C | C | T | T | G | A | T | G | G | H | G | H | H | T | H | H | H | C | H | G | G | G | A | H | H | C | H | T | C | A | T | A | A | T | C | T | -9 | C | G | H | T | G | G | H | T | G | T | T | A |
| 2015_P4-D12 | PHI2 | PHI3 | mother | 2015_P4-D12 | | H | H | H | H | C | H | G | C | A | C | G | G | A | H | H | T | H | A | C | A | T | T | H | C | C | H | C | T | A | -9 | G | H | H | A | H | G | H | H | H | H | H | T | A | H | G | H | H | H | H | T | H | G | H | T | G | G | H | G | H | H | T | H | H | H | C | H | G | G | G | A | H | H | C | H | T | C | A | T | H | A | T | C | H | H | H | G | A | C | G | H | H | H | H | T | T | H |
| 2015_P4-D12-1 | PHI2 | PHI3 | progeny | 2015_P4-D12 | | T | T | T | G | C | G | G | C | A | C | G | G | A | H | H | T | G | A | C | A | T | T | H | C | C | T | C | T | A | T | G | H | H | A | H | G | G | C | C | G | C | T | A | H | G | G | C | C | C | T | T | G | A | T | G | G | T | -9 | G | H | T | C | A | G | C | C | G | G | G | A | H | H | C | H | T | C | A | T | H | A | T | C | C | G | H | G | A | C | G | C | H | T | G | T | T | A |
| 2015_P4-D12-2 | PHI2 | PHI3 | progeny | 2015_P4-D12 | | H | H | H | H | C | H | G | C | A | C | G | G | A | H | H | T | H | A | C | A | T | T | H | C | C | H | C | T | H | T | G | C | C | A | H | G | C | H | H | H | H | T | A | H | G | H | C | H | H | T | H | G | G | T | G | G | H | G | H | H | T | H | H | H | C | H | G | G | G | A | H | H | C | T | T | C | A | T | C | A | -9 | C | C | -9 | T | G | A | C | G | G | A | H | H | T | T | H |
| 2015_P4-D3 | PHI2 | PHI3 | mother | 2015_P4-D3 | | H | H | H | H | C | H | G | C | A | C | G | G | A | H | H | T | H | A | C | A | T | -9 | H | C | C | H | C | T | H | T | G | H | H | A | H | G | H | H | H | H | H | T | A | H | G | H | H | H | H | T | H | G | H | T | G | G | H | G | H | H | T | H | H | H | C | H | G | G | G | A | H | H | C | H | T | C | A | T | H | A | T | C | T | A | H | G | H | H | G | H | H | H | G | H | T | H |
| 2015_P4-D3-1 | PHI2 | PHI3 | progeny | 2015_P4-D3 | | H | H | C | A | C | T | G | C | A | C | G | G | A | A | G | T | G | A | C | A | T | T | T | C | C | C | C | T | G | T | G | H | H | A | G | G | C | T | T | A | T | -9 | A | H | G | H | C | C | C | T | T | G | A | T | G | G | G | G | H | H | T | C | A | G | C | C | G | G | G | A | A | C | C | T | T | C | A | T | A | A | T | -9 | -9 | A | C | G | A | C | G | H | H | T | G | C | T | H |
| 2015_P4-D4 | PHI2 | PHI3 | mother | 2015_P4-D4 | | H | H | H | H | C | H | G | C | A | C | G | G | A | H | H | T | H | A | C | A | T | T | H | C | C | H | C | T | H | T | G | H | H | A | H | G | H | H | H | H | H | T | A | H | G | H | C | H | H | T | H | G | H | T | G | G | H | G | H | H | T | H | H | H | C | H | G | G | G | A | H | H | C | H | T | C | H | T | H | A | T | C | T | H | H | G | H | H | G | H | H | H | H | T | T | H |
| 2015_P4-D4-1 | PHI2 | PHI3 | progeny | 2015_P4-D4 | | A | H | H | H | C | H | G | C | A | C | G | G | A | H | H | T | H | A | C | A | T | T | H | C | C | H | C | T | H | T | G | H | H | A | H | G | H | H | H | H | H | T | A | H | G | H | C | H | H | T | H | G | H | T | G | G | H | G | H | G | T | H | H | H | C | H | G | G | G | A | H | H | C | H | T | C | H | T | H | A | T | C | T | A | C | G | H | H | G | H | G | T | G | T | T | A |
| 2015_P4-D7 | PHI2 | PHI3 | mother | 2015_P4-D7 | | H | H | H | H | C | H | G | C | A | C | G | G | A | H | H | T | H | A | C | A | T | T | H | C | C | H | C | T | H | T | G | H | H | A | H | G | H | H | H | H | H | T | A | H | G | H | C | H | H | T | H | G | H | T | G | G | H | G | H | H | T | H | H | H | C | H | G | G | G | A | H | H | C | H | T | C | A | T | H | A | T | C | T | A | H | G | H | H | G | H | H | H | G | H | T | H |
| 2015_P4-D7-1 | PHI2 | PHI3 | progeny | 2015_P4-D7 | | T | T | H | H | C | H | G | C | A | C | G | G | A | H | H | T | H | A | C | A | T | T | H | C | C | H | C | T | H | T | G | H | H | A | H | G | H | H | H | H | H | T | A | H | G | H | H | H | T | T | C | G | G | T | G | G | G | G | A | H | T | G | C | A | C | T | G | G | G | A | G | G | C | G | T | C | A | T | H | A | T | -9 | H | H | H | G | A | C | G | C | G | H | A | T | T | H |
| 2015_P4-E1 | PHI2 | PHI3 | mother | 2015_P4-E1 | | H | H | H | H | C | H | G | C | A | C | G | G | A | H | H | T | H | A | C | A | T | T | H | C | C | H | -9 | T | H | T | G | H | H | A | H | G | H | H | H | H | H | T | A | H | G | H | C | H | H | T | H | G | H | T | G | G | H | G | H | H | T | H | H | H | C | H | -9 | G | G | A | H | H | C | H | T | C | A | T | H | A | T | C | H | H | H | G | H | H | G | H | H | H | H | T | H |
| 2015_P4-E1-1 | PHI2 | PHI3 | progeny | 2015_P4-E1 | | T | H | H | A | C | H | G | C | A | C | G | G | A | A | H | T | T | A | C | A | T | T | A | C | C | C | C | T | H | T | G | H | H | A | H | G | H | H | H | H | H | T | A | C | G | H | C | H | H | T | H | G | H | T | G | G | H | G | G | G | T | C | A | G | C | C | G | G | G | A | A | C | C | H | T | C | A | T | H | A | -9 | C | T | A | T | G | A | T | G | H | H | T | G | H | T | H |
| 2015_P4-E1-2 | PHI2 | PHI3 | progeny | 2015_P4-E1 | | H | H | C | A | C | T | G | C | A | C | G | G | A | G | H | T | G | A | C | A | T | [illegible] | [illegible] | [illegible] | [illegible] | [illegible] | [illegible] | [illegible] | [illegible] | [illegible] | [illegible] | [illegible] | [illegible] | [illegible] | [illegible] | [illegible] | [illegible] | [illegible] | [illegible] | [illegible] | [illegible] | [illegible] | [illegible] | [illegible] | [illegible] | [illegible] | [illegible] | [illegible] | [illegible] | [illegible] | [illegible] | [illegible] | [illegible] | [illegible] | [illegible] | [illegible] | [illegible] | [illegible] | [illegible] | [illegible] | [illegible] | [illegible] | [illegible] | [illegible] | [illegible] | [illegible] | [illegible] | [illegible] | [illegible] | [illegible] | [illegible] | [illegible] | [illegible] | [illegible] | [illegible] | [illegible] | [illegible] | [illegible] | [illegible] | [illegible] | [illegible] | [illegible] | [illegible] | [illegible] | [illegible] | [illegible] | [illegible] | [illegible] | [illegible] | [illegible] | [illegible] | [illegible] | [illegible] | [illegible] | [illegible] | [illegible] | [illegible] | [illegible] | [illegible] |
| 2015_P4-E10 | PHI8 | PHI3 | mother | 2015_P4-E10 | | H | H | H | H | H | H | H | H | H | H | H | H | H | H | H | H | H | H | H | H | H | H | H | H | H | H | H | H | A | H | G | H | H | H | T | G | H | H | C | H | H | T | A | H | H | H | H | C | H | H | H | H | H | H | H | G | H | G | H | H | H | C | H | H | H | H | H | H | G | H | H | H | H | H | H | C | A | H | A | H | H | H | H | H | H | H | H | H | H | H | H | H | T | G |
| 2015_P4-E10-1 | PHI8 | PHI3 | progeny | 2015_P4-E10 | | H | H | H | H | H | H | H | H | H | H | H | H | H | H | H | H | H | H | H | H | H | H | H | H | H | H | H | H | A | H | G | H | H | H | T | G | H | H | C | H | H | T | A | H | H | H | H | C | H | H | H | H | H | H | H | G | H | G | H | H | H | C | H | H | H | H | H | H | G | H | H | H | H | H | H | C | A | H | A | H | H | H | H | H | H | H | H | H | H | H | H | H | T | G |
| 2015_P4-E10-2 | PHI8 | PHI3 | progeny | 2015_P4-E10 | | H | H | C | A | C | T | G | C | A | C | G | G | A | A | H | T | G | G | C | A | T | [illegible] | [illegible] | [illegible] | [illegible] | [illegible] | [illegible] | [illegible] | [illegible] | [illegible] | [illegible] | [illegible] | [illegible] | [illegible] | [illegible] | [illegible] | [illegible] | [illegible] | [illegible] | [illegible] | [illegible] | [illegible] | [illegible] | [illegible] | [illegible] | [illegible] | [illegible] | [illegible] | [illegible] | [illegible] | [illegible] | [illegible] | [illegible] | [illegible] | [illegible] | [illegible] | [illegible] | [illegible] | [illegible] | [illegible] | [illegible] | [illegible] | [illegible] | [illegible] | [illegible] | [illegible] | [illegible] | [illegible] | [illegible] | [illegible] | [illegible] | [illegible] | [illegible] | [illegible] | [illegible] | [illegible] | [illegible] | [illegible] | [illegible] | [illegible] | [illegible] | [illegible] | [illegible] | [illegible] | [illegible] | [illegible] | [illegible] | [illegible] | [illegible] | [illegible] | [illegible] | [illegible] | [illegible] | [illegible] | [illegible] | [illegible] | [illegible] | [illegible] | [illegible] |
| 2015_P4-E10-3 | PHI8 | PHI3 | progeny | 2015_P4-E10 | | H | H | H | H | H | H | H | H | H | H | H | H | H | H | H | H | H | H | H | H | H | H | H | H | H | H | H | H | A | H | G | H | H | H | T | G | H | H | C | H | H | T | A | H | H | H | H | C | H | H | H | H | H | H | H | G | H | G | H | H | H | C | H | H | H | H | H | H | G | H | H | H | H | H | H | C | A | H | A | H | H | H | H | H | H | H | H | H | H | H | H | H | T | G |
| 2015_P4-E10-4 | PHI8 | PHI3 | progeny | 2015_P4-E10 | | H | H | H | H | H | H | H | H | H | H | H | H | H | H | H | H | H | H | H | H | H | H | H | H | H | H | H | H | A | H | G | H | H | H | T | G | H | H | C | H | H | T | A | H | H | H | -9 | C | C | T | T | G | A | T | G | G | H | G | H | H | H | C | H | H | H | H | H | H | G | H | H | H | H | H | H | C | A | H | A | H | H | H | H | -9 | H | H | H | H | H | H | H | H | H | T | G |
| 2015_P4-E11 | PHI8 | PHI3 | mother | 2015_P4-E11 | | H | H | H | H | H | H | H | H | H | H | H | H | H | H | H | H | H | H | H | H | H | H | H | H | H | H | H | H | H | H | G | H | H | H | T | G | H | H | C | H | H | T | A | H | H | H | H | C | H | H | H | H | H | H | H | G | H | G | H | H | H | C | H | H | H | H | H | H | G | H | H | H | H | H | H | C | H | H | A | H | H | H | H | H | H | H | H | H | H | H | H | H | T | G |
| 2015_P4-E11-1 | PHI8 | PHI3 | progeny | 2015_P4-E11 | | H | H | H | H | H | T | G | C | A | C | G | G | A | A | A | T | G | A | C | A | T | T | A | C | C | T | H | H | G | A | G | C | C | G | T | G | C | T | C | H | H | T | A | H | G | H | H | C | T | C | C | A | G | H | H | G | T | G | H | G | T | C | H | H | H | C | G | G | G | A | A | C | C | T | T | C | G | H | A | A | T | C | C | G | T | G | H | C | T | G | A | H | H | H | T | G |
| 2015_P4-E11-2 | PHI8 | PHI3 | progeny | 2015_P4-E11 | | T | H | T | G | -9 | G | A | T | C | T | A | A | C | H | H | H | H | H | H | H | H | H | H | H | H | H | C | T | A | T | G | A | T | A | T | G | H | H | C | H | H | T | A | C | H | H | H | C | H | -9 | H | H | H | H | H | G | G | -9 | H | H | H | C | A | G | C | C | G | G | G | A | A | C | C | T | T | C | G | -9 | A | H | H | H | H | H | H | H | H | H | H | H | H | C | A | T | T | G |
| 2015_P4-E12 | PHI8 | PHI3 | mother | 2015_P4-E12 | | H | H | H | H | H | H | H | -9 | H | H | H | H | H | H | H | H | H | H | H | H | H | H | H | H | H | H | H | T | A | H | G | H | H | H | T | G | H | H | C | H | H | T | A | H | H | H | H | C | H | H | H | H | H | H | H | G | H | -9 | H | H | H | C | H | H | H | H | H | H | G | H | H | H | H | H | H | C | A | H | A | H | H | H | H | H | H | H | H | H | H | H | H | H | T | G |
| 2015_P4-E12-1 | PHI8 | PHI3 | progeny | 2015_P4-E12 | | A | H | H | A | C | T | G | C | A | C | G | G | A | A | H | H | H | H | H | H | H | H | H | H | H | H | H | T | A | H | G | H | H | H | T | G | H | H | C | H | H | T | A | H | H | H | H | C | H | H | H | H | H | H | H | G | T | -9 | G | G | T | C | H | H | H | H | H | H | G | H | H | H | H | H | H | C | A | H | A | H | H | H | H | H | H | H | H | H | H | H | [illegible] | T | G | C | T | G |
| 2015_P4-E12-2 | PHI8 | PHI3 | progeny | 2015_P4-E12 | | H | H | H | H | H | H | H | H | H | H | H | H | H | H | H | H | G | A | C | A | T | H | H | H | H | H | H | T | A | H | G | H | H | H | T | G | H | H | C | H | H | T | A | H | H | H | H | C | H | H | H | H | H | H | H | G | T | -9 | G | G | T | C | H | H | H | H | H | H | G | H | H | H | H | H | H | C | A | G | A | G | H | H | H | -9 | -9 | H | H | H | T | C | G | H | H | H | T | G |
| 2015_P4-E5 | PHI2 | PHI3 | mother | 2015_P4-E5 | | H | H | H | H | C | H | G | C | A | C | G | G | A | H | H | T | H | A | C | A | T | T | H | C | C | H | C | T | H | T | G | H | H | A | H | G | H | H | H | H | H | T | A | H | G | H | C | H | H | T | H | G | H | T | G | G | H | G | H | H | T | H | H | H | C | H | G | G | G | A | H | H | C | H | T | C | A | T | H | A | T | C | H | H | H | G | H | H | G | H | H | H | H | G | T | H |
| 2015_P4-E5-1 | PHI2 | PHI3 | progeny | 2015_P4-E5 | | H | H | T | G | C | G | G | C | A | C | G | G | A | A | H | T | G | A | C | A | T | T | H | C | C | T | C | T | A | T | G | A | H | A | H | G | H | T | T | A | T | -9 | A | C | G | H | C | G | T | T | C | G | G | T | G | G | G | G | A | H | T | G | H | A | C | T | G | G | G | A | G | G | C | H | T | C | A | T | H | A | T | C | H | H | H | G | H | C | G | H | H | H | H | G | T | H |
| 2015_P4-E5-2 | PHI2 | PHI3 | progeny | 2015_P4-E5 | | H | H | H | H | C | H | G | C | A | C | G | G | A | H | A | T | H | A | C | A | T | T | H | C | C | H | C | T | G | T | G | C | C | A | G | G | H | H | H | H | H | T | A | H | G | H | C | H | H | T | H | G | H | T | G | G | H | G | H | H | T | H | H | H | C | H | G | G | G | A | H | H | C | H | T | C | A | T | H | A | T | C | H | H | H | G | H | H | G | H | H | H | H | G | T | H |
| 2015_P4-E5-3 | PHI2 | PHI3 | progeny | 2015_P4-E5 | | T | H | H | G | C | G | G | C | A | C | G | G | A | G | H | T | H | A | C | A | T | T | T | C | C | C | C | T | H | -9 | G | H | H | A | H | G | H | C | C | G | C | T | A | H | G | H | C | C | C | T | T | G | A | T | G | G | G | G | A | G | T | G | C | A | C | T | G | G | G | A | G | G | C | G | T | C | A | T | C | A | T | C | C | G | H | G | G | T | G | C | G | T | G | T | T | A |
| 2015_P4-E6 | PHI2 | PHI3 | mother | 2015_P4-E6 | | H | H | H | H | C | H | G | C | A | C | G | G | A | H | H | T | H | A | C | A | T | T | H | C | C | H | C | T | H | T | G | H | H | A | H | G | H | H | H | H | H | T | A | H | G | H | C | H | H | T | H | G | H | T | G | G | H | G | H | H | T | H | H | H | C | H | G | G | G | A | H | H | C | H | T | C | A | T | H | A | T | C | T | A | H | G | A | H | G | H | H | T | G | H | T | H |
| 2015_P4-E6-1 | PHI2 | PHI3 | progeny | 2015_P4-E6 | | A | C | C | A | -9 | T | G | C | A | C | G | G | A | A | A | T | G | A | C | A | T | T | T | C | C | C | C | T | G | T | G | H | H | A | T | G | C | T | T | A | T | -9 | A | C | G | G | C | C | C | T | T | G | A | T | G | G | T | -9 | G | G | T | H | A | G | C | C | G | G | G | A | A | C | C | T | T | C | A | T | A | A | T | C | T | A | C | G | A | T | G | H | H | T | G | C | T | G |
| 2015_P4-E6-2 | PHI2 | PHI3 | progeny | 2015_P4-E6 | | T | T | T | H | C | T | G | C | A | C | G | G | A | A | H | T | T | A | C | A | T | T | A | C | C | C | C | T | H | T | G | H | H | A | H | G | H | H | H | H | H | T | A | C | G | H | C | H | H | T | H | G | H | T | G | G | H | G | G | G | T | C | A | G | C | C | G | G | G | A | A | C | C | H | T | C | A | T | H | A | -9 | C | T | A | T | G | A | T | G | H | H | T | G | H | T | H |
| 2015_P4-E8 | PHI8 | PHI3 | mother | 2015_P4-E8 | | H | H | H | H | H | H | H | H | H | H | H | H | H | H | A | H | H | H | H | H | H | H | H | H | H | H | H | T | H | H | G | H | H | H | T | G | H | H | C | H | H | T | A | H | H | H | H | C | H | H | H | H | H | H | H | G | H | -9 | H | H | H | C | H | H | H | H | H | H | G | H | H | H | H | H | H | C | A | H | H | H | H | H | T | A | H | H | H | H | G | H | H | T | G | C | T | G |
| 2015_P4-E8-1 | PHI8 | PHI3 | progeny | 2015_P4-E8 | | H | H | H | H | C | T | G | C | A | C | G | G | A | A | A | H | H | H | H | H | H | H | H | H | H | H | H | T | H | H | G | H | H | H | T | G | H | H | C | H | H | T | A | H | G | G | H | C | C | T | T | G | H | H | G | G | T | -9 | H | H | H | C | H | H | H | H | H | H | G | H | H | H | H | H | H | C | A | H | H | H | H | H | T | A | H | H | H | H | G | H | H | T | G | C | T | G |
| 2015_P4-E9 | PHI8 | PHI3 | mother | 2015_P4-E9 | | H | H | H | H | H | H | H | H | H | H | H | H | H | H | A | H | H | H | H | H | H | H | H | H | H | H | H | T | H | T | H | H | H | H | T | G | H | H | C | H | H | T | A | H | H | H | H | C | H | H | H | H | H | H | H | G | H | -9 | H | H | H | C | H | H | H | H | H | H | G | H | H | H | H | H | H | C | A | H | H | H | H | H | T | A | H | H | A | C | G | H | H | H | H | H | T | G |
| 2015_P4-E9-1 | PHI8 | PHI3 | progeny | 2015_P4-E9 | | H | -9 | H | A | -9 | -9 | -9 | -9 | H | -9 | H | H | -9 | H | A | H | H | A | G | -9 | H | -9 | T | T | G | C | T | T | -9 | T | T | C | C | G | T | G | H | H | C | H | H | T | A | -9 | H | H | -9 | C | -9 | H | H | H | H | H | -9 | G | H | -9 | H | H | H | C | -9 | -9 | H | H | H | H | G | H | -9 | H | H | H | -9 | C | A | H | H | H | H | H | T | A | H | H | A | C | G | H | H | H | H | T | -9 | G |
| 2015_P4-E9-2 | PHI8 | PHI3 | progeny | 2015_P4-E9 | | H | H | H | H | H | H | H | H | H | H | H | H | H | H | A | C | H | H | H | H | H | H | H | H | H | H | H | T | H | T | H | H | H | H | T | G | G | C | C | G | C | T | A | H | H | H | H | C | H | H | H | H | H | H | H | G | H | -9 | H | H | H | C | H | H | H | -9 | H | H | G | H | H | H | H | H | H | C | A | H | H | H | H | H | T | A | H | H | A | C | G | H | H | H | H | H | T | G |
| 2015_P4-F1 | PHI8 | PHI3 | mother | 2015_P4-F1 | | H | H | H | H | H | H | H | H | H | H | H | H | H | H | H | H | H | H | H | H | H | H | H | H | H | H | H | H | A | H | G | H | H | H | T | G | H | H | C | H | H | T | A | H | H | H | H | C | H | H | H | H | H | H | H | G | H | G | H | H | H | C | H | H | H | H | H | H | G | H | H | H | H | H | H | C | A | -9 | A | H | H | H | H | H | H | H | H | H | H | H | H | H | H | T | G |
| 2015_P4-F1-1 | PHI8 | PHI3 | progeny | 2015_P4-F1 | | -9 | -9 | C | G | -9 | -9 | -9 | T | -9 | -9 | -9 | A | -9 | G | -9 | -9 | -9 | -9 | -9 | -9 | -9 | -9 | -9 | -9 | -9 | C | -9 | -9 | -9 | -9 | -9 | -9 | C | -9 | H | -9 | -9 | -9 | -9 | -9 | -9 | -9 | A | T | G | A | H | C | T | T | -9 | -9 | -9 | H | -9 | -9 | -9 | -9 | -9 | -9 | -9 | C | C | -9 | -9 | T | -9 | H | -9 | -9 | -9 | -9 | -9 | H | -9 | C | A | -9 | -9 | -9 | T | -9 | -9 | -9 | -9 | -9 | H | -9 | -9 | -9 | -9 | -9 | A | -9 | -9 | G |
| 2015_P4-F2 | PHI8 | PHI3 | mother | 2015_P4-F2 | | H | H | H | H | H | H | H | H | H | H | H | H | H | H | H | H | H | H | H | H | H | H | H | H | H | H | H | H | A | H | H | H | H | H | T | G | H | H | C | H | H | T | A | H | H | H | H | C | H | H | H | H | H | H | H | G | H | -9 | H | H | H | C | H | H | H | H | H | H | G | H | H | H | H | H | H | C | A | H | A | H | H | H | H | H | H | H | H | H | H | H | H | H | H | T | G |
| 2015_P4-F2-1 | PHI8 | PHI3 | progeny | 2015_P4-F2 | | -9 | H | -9 | H | -9 | H | H | H | H | T | H | H | H | H | A | C | H | A | -9 | G | -9 | H | H | -9 | C | T | C | T | A | H | -9 | H | H | G | T | -9 | C | T | C | A | H | T | A | H | H | H | -9 | C | -9 | H | H | H | H | H | -9 | G | G | -9 | -9 | -9 | H | C | A | G | C | -9 | G | G | G | A | A | -9 | -9 | H | -9 | C | A | G | A | A | H | -9 | -9 | -9 | T | H | G | -9 | G | H | G | T | G | -9 | T | G |
| 2015_P4-F2-2 | PHI8 | PHI3 | progeny | 2015_P4-F2 | | H | C | C | A | C | T | G | C | A | C | G | G | A | H | A | C | T | -9 | G | G | -9 | T | A | C | C | T | C | T | A | T | H | H | H | H | T | G | G | C | C | G | C | T | A | H | H | A | T | -9 | T | C | C | A | G | H | -9 | G | -9 | -9 | A | A | G | C | C | A | T | T | C | A | G | C | G | G | A | G | C | C | A | G | A | G | C | T | T | A | C | T | A | C | T | C | -9 | C | H | T | T | G |
| 2015_P4-F4 | PHI8 | PHI3 | mother | 2015_P4-F4 | | H | H | H | H | H | H | H | H | H | H | H | H | H | H | H | H | H | H | H | H | H | H | H | H | H | H | H | T | A | H | G | H | H | H | T | G | H | H | C | H | H | T | A | H | H | H | H | C | H | H | H | H | H | H | H | G | H | G | H | H | H | C | H | H | H | H | H | H | G | H | H | H | H | H | H | C | A | H | A | H | H | H | H | H | H | H | H | H | H | H | H | H | H | T | G |
| 2015_P4-F4-1 | PHI8 | PHI3 | progeny | 2015_P4-F4 | | H | H | H | H | H | H | H | H | H | H | H | H | H | H | G | C | T | G | G | G | C | H | H | H | H | H | H | T | A | H | G | H | H | H | T | G | H | H | C | H | H | T | A | T | A | A | T | C | T | C | C | A | G | G | H | G | H | G | H | H | H | C | A | G | C | C | G | G | G | A | A | C | C | T | T | C | A | H | A | H | H | H | H | H | H | H | H | G | T | G | G | H | H | H | T | G |
| 2015_P4-F4-2 | PHI8 | PHI3 | progeny | 2015_P4-F4 | | H | H | H | H | H | H | H | C | A | C | G | G | A | A | H | H | H | H | H | H | H | H | H | H | H | H | H | T | A | H | G | H | H | H | T | G | H | H | C | G | C | T | A | H | G | G | C | C | C | T | T | G | A | T | H | G | H | G | H | H | H | C | H | H | H | H | H | H | G | H | H | H | H | H | H | C | A | G | A | G | C | T | T | A | C | T | H | H | H | H | H | H | H | H | T | G |
| 2015_P4-F4-3 | PHI8 | PHI3 | progeny | 2015_P4-F4 | | H | H | H | H | -9 | H | H | H | H | H | H | H | H | H | H | H | H | H | H | H | H | H | H | H | H | H | H | T | A | H | G | H | H | H | T | G | H | H | C | H | H | T | A | H | H | H | H | C | H | H | H | H | H | H | H | G | H | G | H | H | H | C | H | H | H | H | H | H | G | H | H | H | H | H | H | C | A | H | A | H | H | H | H | -9 | H | H | H | H | H | H | H | H | H | T | G |
| 2015_P4-F4-4 | PHI8 | PHI3 | progeny | 2015_P4-F4 | | H | H | H | H | H | H | H | H | H | H | H | H | H | H | H | H | H | H | H | H | H | H | H | H | H | H | H | T | A | H | G | H | H | H | T | G | G | C | C | G | C | T | A | H | H | H | H | C | H | H | H | H | H | H | H | G | H | G | H | H | H | C | H | H | H | H | H | H | G | H | H | H | H | H | H | C | A | H | A | H | H | H | H | -9 | H | H | H | H | H | H | C | A | T | T | G |
| 2015_P4-F5 | PHI2 | PHI8 | mother | 2015_P4-F5 | | T | T | T | G | H | G | H | H | H | H | H | H | G | G | H | T | H | H | H | H | H | T | -9 | H | C | -9 | H | G | H | G | C | C | H | H | G | [illegible] | C | T | H | A | T | T | A | T | H | A | H | H | T | H | C | H | G | H | H | G | G | G | A | A | G | H | C | A | H | T | H | H | G | H | G | G | H | G | H | C | A | H | A | -9 | H | H | T | A | C | H | A | C | H | C | G | T | G | H | T | H |
| 2015_P4-F5-1 | PHI2 | PHI8 | progeny | 2015_P4-F5 | | T | T | T | G | -9 | G | A | T | C | T | A | H | H | G | G | H | T | A | C | A | T | H | T | T | G | C | T | A | G | A | G | C | C | H | T | G | C | T | T | A | T | -9 | A | T | G | A | C | G | T | T | C | H | G | H | H | G | G | -9 | A | A | G | C | C | A | T | T | C | A | G | C | G | G | A | G | H | C | A | T | A | A | T | C | T | A | C | H | A | C | H | C | G | T | G | C | T | G |
| 2015_P4-F5-2 | PHI2 | PHI8 | progeny | 2015_P4-F5 | | T | T | T | G | C | G | G | C | A | C | G | G | A | G | G | H | T | G | G | G | C | -9 | T | T | G | C | T | A | -9 | H | G | C | C | G | T | G | C | T | T | A | T | -9 | A | T | A | A | T | C | T | C | C | A | G | G | -9 | G | G | G | A | A | G | H | C | A | T | T | C | A | G | C | G | G | H | G | C | C | A | -9 | A | -9 | H | H | T | -9 | C | H | A | C | T | C | G | T | G | C | T | G |
| 2015_P4-F5-3 | PHI2 | PHI8 | progeny | 2015_P4-F5 | | T | T | T | G | H | G | H | H | H | H | H | H | G | G | C | T | H | H | H | H | H | T | H | H | C | H | H | G | H | G | C | C | H | H | G | [illegible] | C | T | H | A | T | T | A | T | H | A | H | H | T | H | C | G | G | T | G | G | G | G | A | A | G | H | C | A | C | T | G | G | G | A | G | G | C | G | C | C | A | H | A | G | C | H | T | A | C | H | A | C | H | C | G | T | G | C | H | H |
| 2015_P4-F5-4 | PHI2 | PHI8 | progeny | 2015_P4-F5 | | T | T | T | G | -9 | G | H | C | A | C | G | H | -9 | G | G | H | T | G | G | G | H | T | T | T | G | C | T | A | G | T | G | C | C | G | T | G | C | T | H | A | T | T | -9 | T | H | A | H | C | T | C | C | H | G | H | G | G | -9 | G | A | A | G | H | C | A | H | T | G | G | G | A | G | G | C | G | T | C | A | G | A | -9 | C | T | T | -9 | C | H | A | C | G | C | G | T | G | C | T | H |
| 2015_P4-F5-5 | PHI2 | PHI8 | progeny | 2015_P4-F5 | | T | T | T | G | H | G | H | H | H | H | H | H | G | G | T | T | A | C | A | T | H | T | C | C | C | C | T | G | T | G | C | C | H | H | G | [illegible] | C | T | H | A | T | T | A | T | H | A | H | C | T | H | C | H | G | H | H | G | G | G | A | A | G | H | C | A | T | T | C | A | G | C | G | G | A | G | C | C | A | T | A | A | H | H | T | A | C | G | A | C | G | C | G | T | G | H | T | A |
| 2015_P4-F5-6 | PHI2 | PHI8 | progeny | 2015_P4-F5 | | T | T | T | G | C | G | H | C | A | C | G | G | A | G | G | C | T | A | C | A | T | H | T | C | C | C | C | T | G | T | G | C | C | H | H | G | C | T | C | A | T | T | A | T | G | A | C | G | T | T | C | H | G | T | G | G | G | G | A | A | G | G | C | A | C | T | G | G | G | A | G | G | C | G | H | C | A | H | A | A | T | C | T | A | C | H | A | C | T | C | G | T | G | C | T | A |
| 2015_P4-F5-7 | PHI2 | PHI8 | progeny | 2015_P4-F5 | | T | T | T | G | -9 | G | A | T | C | T | A | H | H | G | G | H | T | G | H | A | T | H | T | H | H | C | H | H | G | H | G | C | C | H | H | G | C | T | T | A | T | T | A | T | A | A | T | C | T | C | C | A | G | G | H | G | G | -9 | A | A | G | H | C | A | C | T | G | G | G | A | G | G | C | G | T | C | A | H | A | H | H | H | T | A | C | H | A | C | T | C | G | T | G | H | T | H |

plant_id – unique identifier of the plant
mom_gen – genetic backgroiund of mother in the hybred
dad_gen – genetic background of father in the hybred
family_ – identify mother and all the progeny plants from the same mother
mother_progeny – identify each plant as mother of the family or progeny
Chr – number of the chromosome, where marker is located
Distance – position of the marker on the corresponding chromosome in cM
H – heterozygous marker
A, T, G and C – marker homozygous for the corresponding nucleotide
EQV, LS, -9 , etc. – missing values

### Table S5. Results of the SNP array analasis

SNP columns (Chr / Position, cM):

| Chr | 1 | 1 | 1 | 1 | 1 | 1 | 1 | 1 | 1 | 1 | 1 | 1 | 1 | 1 | 1 | 1 | 1 | 1 | 1 | 1 | 1 | 1 | 1 | 1 | 1 | 1 | 1 | 1 | 1 | 1 | 1 | 1 | 1 | 1 | 1 | 1 | 1 | 1 | 1 | 1 | 1 | 1 | 1 | 2 | 2 | 2 | 2 | 2 | 2 | 2 | 2 | 2 | 2 | 2 | 2 | 2 | 2 | 2 | 2 | 2 | 2 | 2 | 2 | 2 | 2 | 2 | 2 | 2 | 2 | 2 | 2 | 2 | 2 | 2 | 2 | 2 | 2 | 2 | 2 | 2 | 3 | 3 | 3 | 3 | 3 | 3 | 3 | 3 | 3 | 3 | 3 | 3 | 3 | 3 | 3 | 3 | 3 | 3 | 3 | 3 | 3 | 3 |
|---|---|---|---|---|---|---|---|---|---|---|---|---|---|---|---|---|---|---|---|---|---|---|---|---|---|---|---|---|---|---|---|---|---|---|---|---|---|---|---|---|---|---|---|---|---|---|---|---|---|---|---|---|---|---|---|---|---|---|---|---|---|---|---|---|---|---|---|---|---|---|---|---|---|---|---|---|---|---|---|---|---|---|---|---|---|---|---|---|---|---|---|---|---|---|---|---|---|---|---|---|---|---|
| Position, cM | 2.6 | 9.4 | 16 | 23 | 30 | 30 | 32 | 36 | 42 | 48 | 52 | 52 | 57 | 57 | 62 | 62 | 63 | 77 | 83 | 93 | 99 | 111 | 122 | 136 | 137 | 138 | 147 | 161 | 181 | 200 | 207 | 216 | 226 | 230 | 242 | 251 | 264 | 277 | 281 | 295 | 298 | 307 | 317 | 7.6 | 10 | 14 | 27 | 33 | 44 | 52 | 65 | 81 | 92 | 104 | 114 | 120 | 126 | 127 | 129 | 129 | 134 | 145 | 148 | 159 | 162 | 164 | 164 | 170 | 170 | 171 | 190 | 201 | 201 | 201 | 208 | 216 | 227 | 231 | 240 | 243 | 249 | 4.4 | 10 | 26 | 28 | 42 | 43 | 54 | 54 | 55 | 56 | 62 | 75 | 75 | 77 | 77 | 78 | 82 | 84 | 84 | 84 | 84 | 91 |

| plant_id | mom_gen | dad_gen | mother_progeny | family_ | SNP genotypes |
|---|---|---|---|---|---|
| 2016A_P1-A3 | PHI1 | PHI3 | mother | 2016A_P1-A3 | [illegible] |
| 2016A_P1-A3_1 | PHI1 | PHI3 | progeny | 2016A_P1-A3 | [illegible] |
| 2016A_P1-D5 | PHI1 | PHI3 | mother | 2016A_P1-D5 | [illegible] |
| 2016A_P1-D5_1 | PHI1 | PHI3 | progeny | 2016A_P1-D5 | [illegible] |
| 2016A_P1-D8 | PHI1 | PHI3 | mother | 2016A_P1-D8 | [illegible] |
| 2016A_P1-D8_3 | PHI1 | PHI3 | progeny | 2016A_P1-D8 | [illegible] |
| 2016A_P1-E3 | PHI1 | PHI3 | mother | 2016A_P1-E3 | [illegible] |
| 2016A_P1-E3_3 | PHI1 | PHI3 | progeny | 2016A_P1-E3 | [illegible] |
| 2016A_P1-E3_5 | PHI1 | PHI3 | progeny | 2016A_P1-E3 | [illegible] |
| 2016A_P1-E9 | PHI1 | PHI3 | mother | 2016A_P1-E9 | [illegible] |
| 2016A_P1-E9_1 | PHI1 | PHI3 | progeny | 2016A_P1-E9 | [illegible] |
| 2016A_P1-F7 | PHI1 | PHI3 | mother | 2016A_P1-F7 | [illegible] |
| 2016A_P1-F7_4 | PHI1 | PHI3 | progeny | 2016A_P1-F7 | [illegible] |
| 2016A_P1-F8 | PHI2 | PHI1 | mother | 2016A_P1-F8 | [illegible] |
| 2016A_P1-F8_4 | PHI2 | PHI1 | progeny | 2016A_P1-F8 | [illegible] |
| 2016A_P1-G12 | PHI2 | PHI1 | mother | 2016A_P1-G12 | [illegible] |
| 2016A_P1-G12_1 | PHI2 | PHI1 | progeny | 2016A_P1-G12 | [illegible] |
| 2016A_P1-G12_2 | PHI2 | PHI1 | progeny | 2016A_P1-G12 | [illegible] |
| 2016A_P1-G12_6 | PHI2 | PHI1 | progeny | 2016A_P1-G12 | [illegible] |
| 2016A_P1-H11 | PHI2 | PHI1 | mother | 2016A_P1-H11 | [illegible] |
| 2016A_P1-H11_3 | PHI2 | PHI1 | progeny | 2016A_P1-H11 | [illegible] |
| 2016A_P1-H11_4 | PHI2 | PHI1 | progeny | 2016A_P1-H11 | [illegible] |
| 2016A_P1-H8 | PHI2 | PHI1 | mother | 2016A_P1-H8 | [illegible] |
| 2016A_P1-H8_1 | PHI2 | PHI1 | progeny | 2016A_P1-H8 | [illegible] |
| 2016A_P1-H8_16 | PHI2 | PHI1 | progeny | 2016A_P1-H8 | [illegible] |
| 2016A_P1-H8_4 | PHI2 | PHI1 | progeny | 2016A_P1-H8 | [illegible] |
| 2016A_P1-H8_5 | PHI2 | PHI1 | progeny | 2016A_P1-H8 | [illegible] |
| 2016A_P1-H8_7 | PHI2 | PHI1 | progeny | 2016A_P1-H8 | [illegible] |
| 2016A_P2-A11 | PHI2 | PHI1 | mother | 2016A_P2-A11 | [illegible] |
| 2016A_P2-A11_4 | PHI2 | PHI1 | progeny | 2016A_P2-A11 | [illegible] |
| 2016A_P2-A11_5 | PHI2 | PHI1 | progeny | 2016A_P2-A11 | [illegible] |
| 2016A_P2-A12 | PHI2 | PHI1 | mother | 2016A_P2-A12 | [illegible] |
| 2016A_P2-A12_12 | PHI2 | PHI1 | progeny | 2016A_P2-A12 | [illegible] |
| 2016A_P2-A12_4 | PHI2 | PHI1 | progeny | 2016A_P2-A12 | [illegible] |
| 2016A_P2-A12_7 | PHI2 | PHI1 | progeny | 2016A_P2-A12 | [illegible] |
| 2016A_P2-A12_8 | PHI2 | PHI1 | progeny | 2016A_P2-A12 | [illegible] |
| 2016A_P2-A12_9 | PHI2 | PHI1 | progeny | 2016A_P2-A12 | [illegible] |
| 2016A_P2-A7 | PHI2 | PHI1 | mother | 2016A_P2-A7 | [illegible] |
| 2016A_P2-A7_9 | PHI2 | PHI1 | progeny | 2016A_P2-A7 | [illegible] |
| 2016A_P2-B10 | PHI2 | PHI1 | mother | 2016A_P2-B10 | [illegible] |
| 2016A_P2-B10_6 | PHI2 | PHI1 | progeny | 2016A_P2-B10 | [illegible] |
| 2016A_P2-B4 | PHI2 | PHI1 | mother | 2016A_P2-B4 | [illegible] |
| 2016A_P2-B4_1 | PHI2 | PHI1 | progeny | 2016A_P2-B4 | [illegible] |
| 2016A_P2-B4_5 | PHI2 | PHI1 | progeny | 2016A_P2-B4 | [illegible] |
| 2016A_P2-B4_6 | PHI2 | PHI1 | progeny | 2016A_P2-B4 | [illegible] |
| 2016A_P2-B7 | PHI2 | PHI1 | mother | 2016A_P2-B7 | [illegible] |
| 2016A_P2-B7_1 | PHI2 | PHI1 | progeny | 2016A_P2-B7 | [illegible] |
| 2016A_P2-C11 | PHI2 | PHI1 | mother | 2016A_P2-C11 | [illegible] |
| 2016A_P2-C11_1 | PHI2 | PHI1 | progeny | 2016A_P2-C11 | [illegible] |
| 2016A_P2-C11_4 | PHI2 | PHI1 | progeny | 2016A_P2-C11 | [illegible] |
| 2016A_P2-D1 | PHI2 | PHI1 | mother | 2016A_P2-D1 | [illegible] |
| 2016A_P2-D1_10 | PHI2 | PHI1 | progeny | 2016A_P2-D1 | [illegible] |
| 2016A_P2-D1_12 | PHI2 | PHI1 | progeny | 2016A_P2-D1 | [illegible] |
| 2016A_P2-D1_3 | PHI2 | PHI1 | progeny | 2016A_P2-D1 | [illegible] |
| 2016A_P2-D1_4 | PHI2 | PHI1 | progeny | 2016A_P2-D1 | [illegible] |
| 2016A_P2-D1_7 | PHI2 | PHI1 | progeny | 2016A_P2-D1 | [illegible] |
| 2016A_P2-D1_8 | PHI2 | PHI1 | progeny | 2016A_P2-D1 | [illegible] |
| 2016A_P2-D10 | PHI2 | PHI1 | mother | 2016A_P2-D10 | [illegible] |
| 2016A_P2-D10_3 | PHI2 | PHI1 | progeny | 2016A_P2-D10 | [illegible] |
| 2016A_P2-D5 | PHI2 | PHI1 | mother | 2016A_P2-D5 | [illegible] |
| 2016A_P2-D5_4 | PHI2 | PHI1 | progeny | 2016A_P2-D5 | [illegible] |
| 2016A_P2-D7 | PHI2 | PHI1 | mother | 2016A_P2-D7 | [illegible] |
| 2016A_P2-D7_3 | PHI2 | PHI1 | progeny | 2016A_P2-D7 | [illegible] |
| 2016A_P2-F2 | PHI2 | PHI1 | mother | 2016A_P2-F2 | [illegible] |
| 2016A_P2-F2_2 | PHI2 | PHI1 | progeny | 2016A_P2-F2 | [illegible] |
| 2016A_P2-F7 | PHI2 | PHI1 | mother | 2016A_P2-F7 | [illegible] |
| 2016A_P2-F7_1 | PHI2 | PHI1 | progeny | 2016A_P2-F7 | [illegible] |
| 2016A_P2-G1 | PHI2 | PHI1 | mother | 2016A_P2-G1 | [illegible] |
| 2016A_P2-G1_4 | PHI2 | PHI1 | progeny | 2016A_P2-G1 | [illegible] |
| 2016A_P2-G1_6 | PHI2 | PHI1 | progeny | 2016A_P2-G1 | [illegible] |
| 2016A_P2-G2 | PHI2 | PHI1 | mother | 2016A_P2-G2 | [illegible] |
| 2016A_P2-G2_5 | PHI2 | PHI1 | progeny | 2016A_P2-G2 | [illegible] |
| 2016A_P2-G2_6 | PHI2 | PHI1 | progeny | 2016A_P2-G2 | [illegible] |
| 2016A_P2-G7 | PHI2 | PHI1 | mother | 2016A_P2-G7 | [illegible] |
| 2016A_P2-G7_13 | PHI2 | PHI1 | progeny | 2016A_P2-G7 | [illegible] |
| 2016A_P2-G7_15 | PHI2 | PHI1 | progeny | 2016A_P2-G7 | [illegible] |
| 2016A_P2-G7_6 | PHI2 | PHI1 | progeny | 2016A_P2-G7 | [illegible] |
| 2016A_P2-G7_8 | PHI2 | PHI1 | progeny | 2016A_P2-G7 | [illegible] |
| 2016A_P2-G9 | PHI2 | PHI1 | mother | 2016A_P2-G9 | [illegible] |
| 2016A_P2-G9_3 | PHI2 | PHI1 | progeny | 2016A_P2-G9 | [illegible] |
| 2016A_P2-G9_5 | PHI2 | PHI1 | progeny | 2016A_P2-G9 | [illegible] |
| 2016A_P2-G9_6 | PHI2 | PHI1 | progeny | 2016A_P2-G9 | [illegible] |
| 2016A_P2-H6 | PHI2 | PHI1 | mother | 2016A_P2-H6 | [illegible] |
| 2016A_P2-H6_1 | PHI2 | PHI1 | progeny | 2016A_P2-H6 | [illegible] |
| 2016A_P2-H6_16 | PHI2 | PHI1 | progeny | 2016A_P2-H6 | [illegible] |
| 2016A_P2-H6_17 | PHI2 | PHI1 | progeny | 2016A_P2-H6 | [illegible] |
| 2016A_P2-H6_9 | PHI2 | PHI1 | progeny | 2016A_P2-H6 | [illegible] |
| 2016A_P3-B5 | PHI10 | PHI1 | mother | 2016A_P3-B5 | [illegible] |
| 2016A_P3-B5_1 | PHI10 | PHI1 | progeny | 2016A_P3-B5 | [illegible] |
| 2016A_P3-B8 | PHI10 | PHI1 | mother | 2016A_P3-B8 | [illegible] |
| 2016A_P3-B8_1 | PHI10 | PHI1 | progeny | 2016A_P3-B8 | [illegible] |
| 2016A_P3-B8_2 | PHI10 | PHI1 | progeny | 2016A_P3-B8 | [illegible] |
| 2016A_P3-B8_4 | PHI10 | PHI1 | progeny | 2016A_P3-B8 | [illegible] |
| 2016A_P3-C12 | PHI10 | PHI1 | mother | 2016A_P3-C12 | [illegible] |
| 2016A_P3-C12_2 | PHI10 | PHI1 | progeny | 2016A_P3-C12 | [illegible] |
| 2016A_P3-D9 | PHI10 | PHI1 | mother | 2016A_P3-D9 | [illegible] |
| 2016A_P3-D9_2 | PHI10 | PHI1 | progeny | 2016A_P3-D9 | [illegible] |
| 2016A_P3-H12 | PHI4 | PHI1 | mother | 2016A_P3-H12 | [illegible] |
| 2016A_P3-H12_1 | PHI4 | PHI1 | progeny | 2016A_P3-H12 | [illegible] |
| 2016A_P3-H8 | PHI4 | PHI1 | mother | 2016A_P3-H8 | [illegible] |
| 2016A_P3-H8_1 | PHI4 | PHI1 | progeny | 2016A_P3-H8 | [illegible] |
| 2016A_P4-A1 | PHI4 | PHI1 | mother | 2016A_P4-A1 | [illegible] |
| 2016A_P4-A1_1 | PHI4 | PHI1 | progeny | 2016A_P4-A1 | [illegible] |
| 2016A_P4-A3 | PHI4 | PHI1 | mother | 2016A_P4-A3 | [illegible] |
| 2016A_P4-A3_1 | PHI4 | PHI1 | progeny | 2016A_P4-A3 | [illegible] |
| 2016A_P4-A3_5 | PHI4 | PHI1 | progeny | 2016A_P4-A3 | [illegible] |
| 2016A_P4-B4 | PHI4 | PHI1 | mother | 2016A_P4-B4 | [illegible] |
| 2016A_P4-B4_3 | PHI4 | PHI1 | progeny | 2016A_P4-B4 | [illegible] |
| 2016A_P4-C11 | PHI4 | PHI1 | mother | 2016A_P4-C11 | [illegible] |
| 2016A_P4-C11_3 | PHI4 | PHI1 | progeny | 2016A_P4-C11 | [illegible] |
| 2016A_P4-C2 | PHI4 | PHI1 | mother | 2016A_P4-C2 | [illegible] |
| 2016A_P4-C2_2 | PHI4 | PHI1 | progeny | 2016A_P4-C2 | [illegible] |
| 2016A_P4-C2_3 | PHI4 | PHI1 | progeny | 2016A_P4-C2 | [illegible] |
| 2016A_P4-C2_5 | PHI4 | PHI1 | progeny | 2016A_P4-C2 | [illegible] |
| 2016A_P4-C2_6 | PHI4 | PHI1 | progeny | 2016A_P4-C2 | [illegible] |
| 2016A_P4-C4 | PHI4 | PHI1 | mother | 2016A_P4-C4 | [illegible] |
| 2016A_P4-C4_9 | PHI4 | PHI1 | progeny | 2016A_P4-C4 | [illegible] |
| 2016A_P4-C9 | PHI4 | PHI1 | mother | 2016A_P4-C9 | [illegible] |
| 2016A_P4-C9_1 | PHI4 | PHI1 | progeny | 2016A_P4-C9 | [illegible] |
| 2016A_P4-C9_3 | PHI4 | PHI1 | progeny | 2016A_P4-C9 | [illegible] |
| 2016A_P4-D12 | PHI4 | PHI1 | mother | 2016A_P4-D12 | [illegible] |
| 2016A_P4-D12_5 | PHI4 | PHI1 | progeny | 2016A_P4-D12 | [illegible] |
| 2016A_P4-D5 | PHI4 | PHI1 | mother | 2016A_P4-D5 | [illegible] |
| 2016A_P4-D5_1 | PHI4 | PHI1 | progeny | 2016A_P4-D5 | [illegible] |
| 2016A_P4-F5 | PHI4 | PHI1 | mother | 2016A_P4-F5 | [illegible] |
| 2016A_P4-F5_1 | PHI4 | PHI1 | progeny | 2016A_P4-F5 | [illegible] |
| 2016A_P4-F5_2 | PHI4 | PHI1 | progeny | 2016A_P4-F5 | [illegible] |
| 2016A_P4-F5_5 | PHI4 | PHI1 | progeny | 2016A_P4-F5 | [illegible] |
| 2016A_P4-F5_6 | PHI4 | PHI1 | progeny | 2016A_P4-F5 | [illegible] |
| 2016A_P4-F7 | PHI7 | PHI1 | mother | 2016A_P4-F7 | [illegible] |
| 2016A_P4-F7_1 | PHI7 | PHI1 | progeny | 2016A_P4-F7 | [illegible] |
| 2016A_P4-G11 | PHI7 | PHI1 | mother | 2016A_P4-G11 | [illegible] |
| 2016A_P4-G11_2 | PHI7 | PHI1 | progeny | 2016A_P4-G11 | [illegible] |
| 2016A_P4-G11_4 | PHI7 | PHI1 | progeny | 2016A_P4-G11 | [illegible] |

### Table S5. Results of the SNP array analasis (continue)

| plant_id | mom_gen | dad_gen | mother_progeny | family_ |
|---|---|---|---|---|
| 2016A_P4-H1 | PHI7 | PHI1 | mother | 2016A_P4-H1 |
| 2016A_P4-H1_2 | PHI7 | PHI1 | progeny | 2016A_P4-H1 |
| 2016A_P4-H1_3 | PHI7 | PHI1 | progeny | 2016A_P4-H1 |
| 2016A_P4-H1_4 | PHI7 | PHI1 | progeny | 2016A_P4-H1 |
| 2016A_P4-H10 | PHI7 | PHI1 | mother | 2016A_P4-H10 |
| 2016A_P4-H10_4 | PHI7 | PHI1 | progeny | 2016A_P4-H10 |
| 2016A_P4-H5 | PHI7 | PHI1 | mother | 2016A_P4-H5 |
| 2016A_P4-H5_1 | PHI7 | PHI1 | progeny | 2016A_P4-H5 |
| 2016A_P4-H5_2 | PHI7 | PHI1 | progeny | 2016A_P4-H5 |
| 2016A_P5-A1 | PHI7 | PHI1 | mother | 2016A_P5-A1 |
| 2016A_P5-A1_3 | PHI7 | PHI1 | progeny | 2016A_P5-A1 |
| 2016A_P5-A1_4 | PHI7 | PHI1 | progeny | 2016A_P5-A1 |
| 2016A_P5-B11 | PHI7 | PHI1 | mother | 2016A_P5-B11 |
| 2016A_P5-B11_1 | PHI7 | PHI1 | progeny | 2016A_P5-B11 |
| 2016A_P5-B11_7 | PHI7 | PHI1 | progeny | 2016A_P5-B11 |
| 2016A_P5-C8 | PHI7 | PHI1 | mother | 2016A_P5-C8 |
| 2016A_P5-C8_1 | PHI7 | PHI1 | progeny | 2016A_P5-C8 |
| 2016A_P5-C8_2 | PHI7 | PHI1 | progeny | 2016A_P5-C8 |
| 2016A_P5-F1 | PHI6 | PHI1 | mother | 2016A_P5-F1 |
| 2016A_P5-F1_1 | PHI6 | PHI1 | progeny | 2016A_P5-F1 |
| 2016A_P5-F10 | PHI6 | PHI1 | mother | 2016A_P5-F10 |
| 2016A_P5-F10_2 | PHI6 | PHI1 | progeny | 2016A_P5-F10 |
| 2016A_P5-F10_4 | PHI6 | PHI1 | progeny | 2016A_P5-F10 |
| 2016A_P5-F10_7 | PHI6 | PHI1 | progeny | 2016A_P5-F10 |
| 2016A_P5-F2 | PHI6 | PHI1 | mother | 2016A_P5-F2 |
| 2016A_P5-F2_3 | PHI6 | PHI1 | progeny | 2016A_P5-F2 |
| 2016A_P5-G10 | PHI6 | PHI1 | mother | 2016A_P5-G10 |
| 2016A_P5-G10_10 | PHI6 | PHI1 | progeny | 2016A_P5-G10 |
| 2016A_P5-G10_11 | PHI6 | PHI1 | progeny | 2016A_P5-G10 |
| 2016A_P5-G10_6 | PHI6 | PHI1 | progeny | 2016A_P5-G10 |
| 2016A_P5-G10_7 | PHI6 | PHI1 | progeny | 2016A_P5-G10 |
| 2016A_P5-G10_8 | PHI6 | PHI1 | progeny | 2016A_P5-G10 |
| 2016A_P5-G10_9 | PHI6 | PHI1 | progeny | 2016A_P5-G10 |
| 2016A_P5-G12 | PHI6 | PHI1 | mother | 2016A_P5-G12 |
| 2016A_P5-G12_1 | PHI6 | PHI1 | progeny | 2016A_P5-G12 |
| 2016A_P5-G8 | PHI6 | PHI1 | mother | 2016A_P5-G8 |
| 2016A_P5-G8_3 | PHI6 | PHI1 | progeny | 2016A_P5-G8 |
| 2016A_P5-G8_4 | PHI6 | PHI1 | progeny | 2016A_P5-G8 |
| 2016A_P5-G8_5 | PHI6 | PHI1 | progeny | 2016A_P5-G8 |
| 2016A_P5-G9 | PHI6 | PHI1 | mother | 2016A_P5-G9 |
| 2016A_P5-G9_1 | PHI6 | PHI1 | progeny | 2016A_P5-G9 |
| 2016A_P5-G9_5 | PHI6 | PHI1 | progeny | 2016A_P5-G9 |
| 2016A_P5-G9_6 | PHI6 | PHI1 | progeny | 2016A_P5-G9 |
| 2016A_P5-H10 | PHI6 | PHI1 | mother | 2016A_P5-H10 |
| 2016A_P5-H10_2 | PHI6 | PHI1 | progeny | 2016A_P5-H10 |
| 2016A_P5-H10_3 | PHI6 | PHI1 | progeny | 2016A_P5-H10 |
| 2016A_P5-H10_5 | PHI6 | PHI1 | progeny | 2016A_P5-H10 |
| 2016A_P5-H11 | PHI6 | PHI1 | mother | 2016A_P5-H11 |
| 2016A_P5-H11_2 | PHI6 | PHI1 | progeny | 2016A_P5-H11 |
| 2016A_P5-H11_3 | PHI6 | PHI1 | progeny | 2016A_P5-H11 |
| 2016A_P5-H11_5 | PHI6 | PHI1 | progeny | 2016A_P5-H11 |
| 2016A_P5-H11_6 | PHI6 | PHI1 | progeny | 2016A_P5-H11 |
| 2016A_P5-H12_11 | PHI6 | PHI1 | progeny | 2016A_P5-H12 |
| 2016A_P5-H12_14 | PHI6 | PHI1 | progeny | 2016A_P5-H12 |
| 2016A_P5-H12_16 | PHI6 | PHI1 | progeny | 2016A_P5-H12 |
| 2016A_P5-H12_17 | PHI6 | PHI1 | progeny | 2016A_P5-H12 |
| 2016A_P5-H12_3 | PHI6 | PHI1 | progeny | 2016A_P5-H12 |
| 2016A_P5-H12_9 | PHI6 | PHI1 | progeny | 2016A_P5-H12 |
| 2016A_P5-H3 | PHI6 | PHI1 | mother | 2016A_P5-H3 |
| 2016A_P5-H3_1 | PHI6 | PHI1 | progeny | 2016A_P5-H3 |
| 2016A_P5-H3_11 | PHI6 | PHI1 | progeny | 2016A_P5-H3 |
| 2016A_P5-H3_2 | PHI6 | PHI1 | progeny | 2016A_P5-H3 |
| 2016A_P5-H3_3 | PHI6 | PHI1 | progeny | 2016A_P5-H3 |
| 2016A_P5-H3_4 | PHI6 | PHI1 | progeny | 2016A_P5-H3 |
| 2016A_P5-H3_5 | PHI6 | PHI1 | progeny | 2016A_P5-H3 |
| 2016A_P5-H3_6 | PHI6 | PHI1 | progeny | 2016A_P5-H3 |
| 2016A_P5-H3_9 | PHI6 | PHI1 | progeny | 2016A_P5-H3 |
| 2016A_P5-H5 | PHI6 | PHI1 | mother | 2016A_P5-H5 |
| 2016A_P5-H5_1 | PHI6 | PHI1 | progeny | 2016A_P5-H5 |
| 2016A_P5-H5_10 | PHI6 | PHI1 | progeny | 2016A_P5-H5 |
| 2016A_P5-H5_12 | PHI6 | PHI1 | progeny | 2016A_P5-H5 |
| 2016A_P5-H5_3 | PHI6 | PHI1 | progeny | 2016A_P5-H5 |
| 2016A_P5-H5_4 | PHI6 | PHI1 | progeny | 2016A_P5-H5 |
| 2016A_P5-H5_5 | PHI6 | PHI1 | progeny | 2016A_P5-H5 |
| 2016A_P5-H5_6 | PHI6 | PHI1 | progeny | 2016A_P5-H5 |
| 2016A_P5-H5_7 | PHI6 | PHI1 | progeny | 2016A_P5-H5 |
| 2016A_P5-H5_8 | PHI6 | PHI1 | progeny | 2016A_P5-H5 |
| 2016A_P5-H6 | PHI6 | PHI1 | mother | 2016A_P5-H6 |
| 2016A_P5-H6_1 | PHI6 | PHI1 | progeny | 2016A_P5-H6 |
| 2016A_P5-H8 | PHI6 | PHI1 | mother | 2016A_P5-H8 |
| 2016A_P5-H8_1 | PHI6 | PHI1 | progeny | 2016A_P5-H8 |
| 2016A_P5-H8_10 | PHI6 | PHI1 | progeny | 2016A_P5-H8 |
| 2016A_P5-H8_16 | PHI6 | PHI1 | progeny | 2016A_P5-H8 |
| 2016A_P5-H8_4 | PHI6 | PHI1 | progeny | 2016A_P5-H8 |
| 2016A_P5H12 | PHI6 | PHI1 | mother | 2016A_P5H12 |
| 2016A_P6-A8 | PHI6 | PHI1 | mother | 2016A_P6-A8 |
| 2016A_P6-A8_1 | PHI6 | PHI1 | progeny | 2016A_P6-A8 |
| 2016A_P6-A8_11 | PHI6 | PHI1 | progeny | 2016A_P6-A8 |
| 2016A_P6-A8_12 | PHI6 | PHI1 | progeny | 2016A_P6-A8 |
| 2016A_P6-A8_16 | PHI6 | PHI1 | progeny | 2016A_P6-A8 |
| 2016A_P6-A8_8 | PHI6 | PHI1 | progeny | 2016A_P6-A8 |
| 2016A_P6-A9 | PHI6 | PHI1 | mother | 2016A_P6-A9 |
| 2016A_P6-A9_1 | PHI6 | PHI1 | progeny | 2016A_P6-A9 |
| 2016A_P6-A9_2 | PHI6 | PHI1 | progeny | 2016A_P6-A9 |
| 2016A_P6-A9_4 | PHI6 | PHI1 | progeny | 2016A_P6-A9 |
| 2016A_P6-A9_6 | PHI6 | PHI1 | progeny | 2016A_P6-A9 |
| 2016A_P6-A9_8 | PHI6 | PHI1 | progeny | 2016A_P6-A9 |
| 2016A_P6-B10 | PHI6 | PHI1 | mother | 2016A_P6-B10 |
| 2016A_P6-B10_11 | PHI6 | PHI1 | progeny | 2016A_P6-B10 |
| 2016A_P6-B10_12 | PHI6 | PHI1 | progeny | 2016A_P6-B10 |
| 2016A_P6-B10_13 | PHI6 | PHI1 | progeny | 2016A_P6-B10 |
| 2016A_P6-B10_15 | PHI6 | PHI1 | progeny | 2016A_P6-B10 |
| 2016A_P6-B10_4 | PHI6 | PHI1 | progeny | 2016A_P6-B10 |
| 2016A_P6-B10_8 | PHI6 | PHI1 | progeny | 2016A_P6-B10 |
| 2016A_P6-B12 | PHI6 | PHI1 | mother | 2016A_P6-B12 |
| 2016A_P6-B12_4 | PHI6 | PHI1 | progeny | 2016A_P6-B12 |
| 2016A_P6-B12_7 | PHI6 | PHI1 | progeny | 2016A_P6-B12 |
| 2016A_P6-B7 | PHI6 | PHI1 | mother | 2016A_P6-B7 |
| 2016A_P6-B7_1 | PHI6 | PHI1 | progeny | 2016A_P6-B7 |
| 2016A_P6-B7_2 | PHI6 | PHI1 | progeny | 2016A_P6-B7 |
| 2016A_P6-B8 | PHI6 | PHI1 | mother | 2016A_P6-B8 |
| 2016A_P6-B8_2 | PHI6 | PHI1 | progeny | 2016A_P6-B8 |
| 2016A_P6-B8_3 | PHI6 | PHI1 | progeny | 2016A_P6-B8 |
| 2016A_P6-C10 | PHI6 | PHI1 | mother | 2016A_P6-C10 |
| 2016A_P6-C10_1 | PHI6 | PHI1 | progeny | 2016A_P6-C10 |
| 2016A_P6-C10_2 | PHI6 | PHI1 | progeny | 2016A_P6-C10 |
| 2016A_P6-C10_3 | PHI6 | PHI1 | progeny | 2016A_P6-C10 |
| 2016A_P6-C12 | PHI6 | PHI1 | mother | 2016A_P6-C12 |
| 2016A_P6-C12_1 | PHI6 | PHI1 | progeny | 2016A_P6-C12 |
| 2016A_P6-C12_2 | PHI6 | PHI1 | progeny | 2016A_P6-C12 |
| 2016A_P6-C12_3 | PHI6 | PHI1 | progeny | 2016A_P6-C12 |
| 2016A_P6-C12_4 | PHI6 | PHI1 | progeny | 2016A_P6-C12 |
| 2016A_P6-D11 | PHI6 | PHI1 | mother | 2016A_P6-D11 |
| 2016A_P6-D11_1 | PHI6 | PHI1 | progeny | 2016A_P6-D11 |
| 2016A_P6-E5 | PHI8 | PHI1 | mother | 2016A_P6-E5 |
| 2016A_P6-E5_4 | PHI8 | PHI1 | progeny | 2016A_P6-E5 |
| 2016A_P6-F7 | PHI8 | PHI1 | mother | 2016A_P6-F7 |
| 2016A_P6-F7_12 | PHI8 | PHI1 | progeny | 2016A_P6-F7 |
| 2016A_P6-G9 | PHI8 | PHI1 | mother | 2016A_P6-G9 |
| 2016A_P6-G9_4 | PHI8 | PHI1 | progeny | 2016A_P6-G9 |
| 2016A_P6-H1 | PHI8 | PHI1 | mother | 2016A_P6-H1 |
| 2016A_P6-H1_3 | PHI8 | PHI1 | progeny | 2016A_P6-H1 |
| 2016A_P6-H3_3 | PHI8 | PHI1 | progeny | 2016A_P6-H3 |
| 2016A_P7-A8 | PHI8 | PHI1 | mother | 2016A_P7-A8 |

| Chr | 1 | 1 | 1 | 1 | 1 | 1 | 1 | 1 | 1 | 1 | 1 | 1 | 1 | 1 | 1 | 1 | 1 | 1 | 1 | 1 | 1 | 1 | 1 | 1 | 1 | 1 | 1 | 1 | 1 | 1 | 1 | 1 | 1 | 1 | 1 | 1 | 1 | 1 | 1 | 1 | 1 | 1 | 1 | 2 | 2 | 2 | 2 | 2 | 2 | 2 | 2 | 2 | 2 | 2 | 2 | 2 | 2 | 2 | 2 | 2 | 2 | 2 | 2 | 2 | 2 | 2 | 2 | 2 | 2 | 2 | 2 | 2 | 2 | 2 | 2 | 2 | 2 | 2 | 2 | 2 | 3 | 3 | 3 | 3 | 3 | 3 | 3 | 3 | 3 | 3 | 3 | 3 | 3 | 3 | 3 | 3 | 3 | 3 | 3 | 3 | 3 | 3 |
|---|---|---|---|---|---|---|---|---|---|---|---|---|---|---|---|---|---|---|---|---|---|---|---|---|---|---|---|---|---|---|---|---|---|---|---|---|---|---|---|---|---|---|---|---|---|---|---|---|---|---|---|---|---|---|---|---|---|---|---|---|---|---|---|---|---|---|---|---|---|---|---|---|---|---|---|---|---|---|---|---|---|---|---|---|---|---|---|---|---|---|---|---|---|---|---|---|---|---|---|---|---|---|---|
| Position, cM | 2.6 | 9.4 | 16 | 23 | 30 | 30 | 32 | 36 | 42 | 48 | 52 | 52 | 57 | 57 | 62 | 62 | 63 | 77 | 83 | 93 | 99 | 111 | 122 | 136 | 137 | 138 | 147 | 161 | 181 | 200 | 207 | 216 | 226 | 230 | 242 | 251 | 264 | 277 | 281 | 295 | 298 | 307 | 317 | 7.6 | 10 | 14 | 27 | 33 | 44 | 52 | 65 | 81 | 92 | 104 | 114 | 120 | 126 | 127 | 129 | 129 | 134 | 145 | 148 | 159 | 162 | 164 | 164 | 170 | 170 | 171 | 190 | 201 | 201 | 201 | 208 | 216 | 227 | 231 | 240 | 243 | 249 | 4.4 | 10 | 26 | 28 | 42 | 43 | 54 | 54 | 55 | 56 | 62 | 75 | 75 | 77 | 77 | 78 | 82 | 84 | 84 | 84 | 84 | 91 |
| 2016A_P4-H1 | H | T | C | G | H | H | H | H | H | H | H | H | H | H | H | H | H | H | H | H | H | H | H | H | H | H | H | C | H | H | H | H | H | H | H | H | H | H | H | H | H | H | H | A | H | H | H | H | H | C | H | H | H | H | H | H | H | G | H | H | H | H | H | H | H | H | H | H | H | H | H | H | H | H | H | T | H | H | H | H | H | H | H | H | H | A | H | A | A | T | H | A | G | T | H | C | H | T | G | C | C | C | H |
| 2016A_P4-H1_2 | H | T | C | G | H | H | H | H | H | H | H | H | H | H | H | H | H | H | H | H | H | A | T | C | C | T | T | C | H | H | H | H | H | H | H | H | H | H | H | C | T | T | T | A | H | H | G | C | G | C | A | T | G | G | A | A | A | G | G | T | C | C | C | G | G | G | H | H | H | H | H | H | H | H | H | T | H | H | C | C | T | H | H | G | A | A | A | A | A | T | A | A | G | T | G | C | T | T | G | C | C | C | T |
| 2016A_P4-H1_3 | H | T | C | G | H | H | H | H | H | H | H | H | H | H | H | H | H | H | H | H | H | H | H | H | H | H | H | C | H | H | H | H | H | H | H | H | H | H | H | H | H | H | H | A | G | H | H | H | H | C | H | H | H | H | H | H | H | G | H | H | H | H | H | H | H | H | H | H | H | H | H | H | H | H | H | T | H | H | H | G | H | H | H | H | H | A | H | A | A | T | H | A | G | T | H | C | H | T | G | C | C | C | H |
| 2016A_P4-H1_4 | H | T | C | G | H | H | H | H | H | H | H | H | H | H | H | H | H | H | H | H | H | H | H | H | H | H | H | C | H | H | H | H | H | H | H | H | H | H | H | H | H | H | H | A | H | H | H | H | H | C | H | H | H | H | H | H | A | G | G | H | C | C | C | G | G | G | A | A | G | C | G | T | T | A | T | T | G | G | C | C | T | H | H | H | H | A | H | A | A | T | H | A | G | T | H | C | H | T | G | C | C | C | H |
| 2016A_P4-H10 | H | H | H | G | H | H | H | H | H | H | H | H | H | H | H | H | H | H | H | H | H | H | H | H | H | H | H | H | H | H | H | H | H | H | H | H | H | H | H | H | H | H | H | A | H | H | H | H | H | C | H | H | H | H | H | A | A | H | H | T | H | H | H | H | H | H | H | H | H | H | H | H | H | H | H | T | H | H | H | H | H | H | H | H | H | H | H | H | H | H | H | H | H | H | H | H | H | T | G | C | C | C | H |
| 2016A_P4-H10_4 | H | H | H | G | H | H | H | H | H | H | H | H | H | H | H | H | H | H | H | H | H | H | H | H | H | H | H | H | H | H | H | H | H | H | H | H | G | G | C | T | G | C | C | A | H | H | H | H | H | C | H | H | H | H | H | A | A | H | H | T | H | H | H | H | H | H | H | H | H | H | G | T | T | A | T | T | G | G | C | C | T | H | H | H | H | H | H | H | H | H | H | H | H | H | H | H | H | T | G | C | C | C | H |
| 2016A_P4-H5 | T | H | H | G | H | H | H | H | H | H | H | H | H | H | H | H | C | H | H | H | H | H | H | H | H | H | H | C | H | H | H | H | H | H | H | H | H | H | H | H | H | H | H | A | H | H | H | H | H | C | H | H | H | H | H | H | H | G | H | H | H | H | H | H | H | H | H | H | H | H | H | H | H | H | H | H | H | H | H | C | H | H | H | H | H | A | H | H | H | H | H | H | H | H | H | H | H | H | H | H | H | H | H |
| 2016A_P4-H5_1 | T | H | H | G | H | H | H | H | H | H | H | H | H | H | H | H | C | H | H | H | H | H | H | H | H | H | H | C | H | H | H | H | H | H | H | H | H | H | H | H | H | H | H | A | H | H | H | H | H | C | H | H | H | H | H | H | H | G | H | H | H | H | H | H | H | H | H | H | H | H | H | H | H | H | H | H | H | T | T | C | G | H | H | H | H | A | H | H | H | H | H | H | H | H | H | H | H | H | H | H | H | H | H |

### Table S5. Results of the SNP array analasis (continue)

| | | | | | Chr | 1 | 1 | 1 | 1 | 1 | 1 | 1 | 1 | 1 | 1 | 1 | 1 | 1 | 1 | 1 | 1 | 1 | 1 | 1 | 1 | 1 | 1 | 1 | 1 | 1 | 1 | 1 | 1 | 1 | 1 | 1 | 1 | 1 | 1 | 1 | 1 | 1 | 1 | 1 | 2 | 2 | 2 | 2 | 2 | 2 | 2 | 2 | 2 | 2 | 2 | 2 | 2 | 2 | 2 | 2 | 2 | 2 | 2 | 2 | 2 | 2 | 2 | 2 | 2 | 2 | 2 | 2 | 2 | 2 | 2 | 2 | 2 | 2 | 2 | 2 | 2 | 2 | 3 | 3 | 3 | 3 | 3 | 3 | 3 | 3 | 3 | 3 | 3 | 3 | 3 | 3 | 3 | 3 | 3 | 3 | 3 | 3 | 3 | 3 |
|---|---|---|---|---|---|---|---|---|---|---|---|---|---|---|---|---|---|---|---|---|---|---|---|---|---|---|---|---|---|---|---|---|---|---|---|---|---|---|---|---|---|---|---|---|---|---|---|---|---|---|---|---|---|---|---|---|---|---|---|---|---|---|---|---|---|---|---|---|---|---|---|---|---|---|---|---|---|---|---|---|---|---|---|---|---|---|---|---|---|---|---|---|---|---|---|---|---|---|---|---|---|---|---|---|---|---|---|---|
| | | | | | Position, cM | 2.6 | 9.4 | 16 | 23 | 30 | 30 | 32 | 36 | 42 | 48 | 52 | 52 | 57 | 57 | 62 | 62 | 63 | 77 | 83 | 93 | 99 | 111 | 122 | 136 | 137 | 138 | 147 | 161 | 181 | 200 | 207 | 216 | 226 | 230 | 242 | 251 | 264 | 277 | 281 | 295 | 298 | 307 | 317 | 7.6 | 10 | 14 | 27 | 33 | 44 | 52 | 65 | 81 | 92 | 104 | 114 | 120 | 126 | 127 | 129 | 129 | 134 | 145 | 148 | 159 | 162 | 164 | 164 | 170 | 170 | 171 | 190 | 201 | 201 | 201 | 208 | 216 | 227 | 231 | 240 | 243 | 249 | 4.4 | 10 | 26 | 28 | 42 | 43 | 54 | 54 | 55 | 56 | 62 | 75 | 75 | 77 | 77 | 78 | 82 | 84 | 84 | 84 | 84 | 91 |
| plant_id | mom_gen | dad_gen | mother_progeny | family_ | | | | | | | | | | | | | | | | | | | | | | | | | | | | | | | | | | | | | | | | | | | | | | | | | | | | | | | | | | | | | | | | | | | | | | | | | | | | | | | | | | | | | | | | | | | | | | | | | | | | | | | | | |
| 2016A_P7-A8_1 | PHI8 | PHI1 | progeny | 2016A_P7-A8 | | H | H | H | G | H | H | H | H | H | H | H | H | H | H | H | T | C | H | H | H | T | H | H | H | H | H | H | H | T | H | H | H | H | H | H | C | H | H | H | H | H | C | H | H | H | T | H | C | G | H | H | H | H | H | H | H | H | G | H | H | H | H | H | H | H | H | H | H | H | H | H | H | H | H | C | H | H | G | C | C | H | H | H | H | H | A | G | A | A | T | G | G | G | T | G | G | T | T | G | C | C | C | C |
| 2016A_P7-C11 | PHI5 | PHI1 | mother | 2016A_P7-C11 | | H | T | C | G | G | G | G | G | G | G | H | H | H | H | H | H | C | H | H | H | H | H | H | H | H | H | H | H | H | H | H | H | H | T | G | C | H | G | C | H | H | H | C | H | H | H | H | H | G | H | H | H | H | H | H | H | A | G | G | H | H | H | H | G | H | G | H | H | H | H | H | H | H | H | H | H | H | H | H | C | H | H | H | H | H | H | A | G | H | H | H | G | G | T | H | C | T | T | G | C | C | C | H |
| 2016A_P7-C11_1 | PHI5 | PHI1 | progeny | 2016A_P7-C11 | | C | T | C | G | G | G | G | G | G | G | A | C | G | A | A | T | C | C | C | T | T | G | C | T | T | G | A | A | C | T | G | A | C | T | G | C | H | G | C | T | G | C | C | H | H | H | H | H | G | H | H | H | H | H | H | H | A | G | G | H | H | H | H | G | H | G | H | H | H | H | H | H | H | H | H | H | H | H | H | C | H | A | A | H | A | G | A | G | G | C | A | G | G | T | G | C | T | T | G | C | H | C | T |
| 2016A_P7-D10 | PHI5 | PHI1 | mother | 2016A_P7-D10 | | H | H | H | G | H | H | H | H | H | H | H | H | H | H | H | T | C | H | H | H | H | H | H | H | H | H | H | H | H | H | H | H | H | H | G | H | H | G | C | H | H | H | C | H | H | H | H | H | G | H | H | H | H | H | H | H | A | G | G | H | H | H | H | H | H | H | H | H | H | H | H | H | H | H | H | H | H | H | H | H | H | H | H | H | H | H | H | G | H | H | H | G | H | H | H | H | T | H | H | H | H | C | H |
| 2016A_P7-D10_1 | PHI5 | PHI1 | progeny | 2016A_P7-D10 | | T | G | T | G | C | A | A | A | A | A | H | A | A | C | G | C | G | T | A | T | T | G | C | T | T | G | A | A | C | G | A | T | T | T | G | H | G | G | C | T | G | C | C | A | A | T | G | C | G | C | A | T | G | A | G | G | A | G | G | H | T | G | H | G | G | G | A | A | G | C | G | H | H | H | T | T | T | T | T | G | G | G | G | H | G | A | G | A | A | T | G | A | G | T | T | C | T | T | G | C | C | C | C |
| 2016A_P7-D2 | PHI5 | PHI1 | mother | 2016A_P7-D2 | | C | T | C | G | H | G | H | H | A | H | H | H | H | H | H | H | G | H | H | H | H | H | H | H | H | H | H | H | H | H | H | H | H | T | G | C | G | G | C | H | H | T | C | C | H | H | H | H | G | C | H | H | H | H | H | H | A | G | G | H | H | G | A | G | C | G | H | A | H | H | H | T | H | H | H | H | H | H | T | H | T | H | A | H | H | A | H | A | A | T | H | A | G | T | H | C | T | T | G | C | C | C | H |
| 2016A_P7-D2_1 | PHI5 | PHI1 | progeny | 2016A_P7-D2 | | T | T | C | G | C | A | A | A | A | A | G | A | A | C | G | C | G | T | A | A | H | A | T | C | C | T | T | C | T | G | G | A | C | T | G | T | C | G | C | C | T | C | C | C | G | C | A | T | G | T | G | T | G | G | A | A | A | G | G | T | C | G | A | G | G | G | A | A | G | C | G | T | T | A | T | T | G | G | C | C | T | A | H | H | A | A | G | A | A | T | G | A | G | T | T | C | T | T | G | C | C | C | C |
| 2016A_P7-D8 | PHI5 | PHI1 | mother | 2016A_P7-D8 | | H | T | C | G | H | H | H | H | H | H | H | H | H | H | H | H | H | H | H | H | H | H | H | H | H | H | H | H | H | H | H | H | H | H | G | H | H | G | C | H | H | H | C | H | H | H | H | H | G | H | H | H | H | H | H | H | A | G | G | H | H | H | H | G | H | G | H | H | H | H | H | H | H | H | H | H | H | H | H | H | H | H | H | H | H | H | A | H | A | A | T | H | H | G | T | H | C | T | T | G | C | C | C | H |
| 2016A_P7-D8_2 | PHI5 | PHI1 | progeny | 2016A_P7-D8 | | H | T | C | G | H | H | H | H | H | H | H | H | H | H | H | H | H | H | H | H | G | A | T | C | C | T | T | C | T | G | A | T | T | T | G | C | G | G | C | H | H | H | C | H | H | H | H | H | G | H | H | H | H | G | A | A | A | G | G | T | C | C | C | G | G | G | A | A | G | C | G | T | T | A | T | T | G | G | C | C | T | H | G | H | G | A | G | A | A | T | G | G | G | T | T | C | T | T | G | C | C | C | C |
| 2016A_P7-G3 | PHI5 | PHI1 | mother | 2016A_P7-G3 | | H | T | C | G | H | H | H | H | H | H | H | H | H | H | H | H | H | H | H | H | T | H | H | H | H | H | H | H | H | H | H | H | H | H | G | H | H | G | C | H | H | H | C | H | H | H | H | H | G | H | H | H | H | H | H | H | A | G | G | H | H | G | A | G | H | G | H | H | H | H | H | H | H | H | H | H | H | H | H | H | H | H | H | H | H | H | A | H | A | A | T | H | H | G | T | H | C | T | T | G | C | C | C | H |
| 2016A_P7-G3_1 | PHI5 | PHI1 | progeny | 2016A_P7-G3 | | H | T | C | G | H | H | H | H | H | H | H | H | H | H | H | H | H | H | H | H | T | H | H | H | H | H | H | H | H | H | H | H | H | H | G | H | H | G | C | H | H | H | C | A | A | H | H | H | G | H | H | H | H | H | H | H | A | G | G | H | T | G | H | G | C | G | T | G | A | T | T | C | G | G | C | C | H | T | T | H | G | H | H | H | H | A | H | A | A | T | H | H | G | T | H | C | T | T | G | C | C | C | H |
| 2016A_P7-G3_3 | PHI5 | PHI1 | progeny | 2016A_P7-G3 | | C | T | C | G | G | G | G | G | G | H | H | H | H | H | H | H | H | H | H | H | H | H | H | T | T | G | A | A | C | T | G | H | H | H | G | H | H | G | C | C | T | T | C | H | H | H | H | H | G | H | H | H | A | A | G | G | A | G | G | C | T | G | A | G | H | G | H | H | H | H | H | H | H | H | H | H | H | H | H | H | H | H | H | H | H | H | A | A | A | A | T | A | A | G | T | G | C | T | T | G | C | C | C | H |
| 2016A_P7-G3_4 | PHI5 | PHI1 | progeny | 2016A_P7-G3 | | C | T | C | G | G | G | G | G | G | G | A | C | G | A | A | T | C | H | H | H | H | H | C | T | T | G | A | A | C | T | G | H | H | H | G | T | C | G | C | C | T | H | C | A | A | T | G | C | G | H | H | H | H | A | G | G | A | G | G | C | T | G | A | G | C | G | T | G | A | T | T | C | G | G | C | C | T | T | H | G | H | H | H | H | G | A | H | H | A | T | G | G | G | T | T | C | T | T | G | C | C | C | C |
| 2016A_P7-G7 | PHI5 | PHI1 | mother | 2016A_P7-G7 | | H | H | H | G | H | H | H | H | H | H | H | H | H | H | H | H | H | H | H | H | H | H | H | H | H | H | H | H | H | H | H | H | H | H | G | H | H | G | C | C | H | H | C | H | H | H | H | H | G | H | H | H | H | H | H | H | A | G | G | H | H | H | H | H | H | H | H | H | H | H | H | H | H | H | H | H | H | H | T | G | H | H | H | H | H | A | H | A | A | T | H | A | G | T | H | C | T | H | H | H | H | C | H |
| 2016A_P7-G7_1 | PHI5 | PHI1 | progeny | 2016A_P7-G7 | | T | H | H | G | H | H | H | H | H | H | H | H | H | H | H | H | H | H | H | H | H | H | H | H | H | H | H | H | H | G | H | H | H | H | G | H | H | G | C | H | H | H | C | H | H | H | H | H | G | H | H | H | H | H | H | H | A | G | G | H | H | H | H | H | H | H | H | H | H | H | H | H | H | H | H | H | T | T | T | G | G | H | H | H | H | A | H | A | A | T | H | A | G | T | H | H | H | H | H | H | H | H | H |
| 2016A_P7-H9 | PHI5 | PHI1 | mother | 2016A_P7-H9 | | H | H | C | G | H | H | H | H | H | H | H | H | H | H | H | T | C | H | H | H | H | H | H | H | H | H | H | H | H | H | H | H | H | H | G | H | H | G | C | H | H | H | C | H | H | H | H | H | G | H | H | H | H | H | H | H | A | G | G | H | H | H | H | H | H | H | H | H | H | H | H | C | H | H | H | H | H | H | H | C | H | H | H | H | H | H | A | A | A | T | A | H | G | T | H | C | T | T | G | C | C | C | H |
| 2016A_P7-H9_1 | PHI5 | PHI1 | progeny | 2016A_P7-H9 | | H | T | C | G | G | G | G | G | G | G | H | H | H | H | H | H | H | H | H | H | H | H | H | H | H | H | H | H | H | H | H | H | H | T | G | H | H | G | C | H | H | H | C | H | H | H | H | H | G | H | H | H | H | H | H | H | A | G | G | H | H | H | H | G | H | G | H | H | H | H | H | H | H | H | H | H | H | H | H | C | H | A | A | G | A | G | A | A | A | T | A | A | G | T | G | C | T | T | G | C | C | C | H |
| 2016A_P7-H9_2 | PHI5 | PHI1 | progeny | 2016A_P7-H9 | | H | T | C | G | G | G | G | G | G | G | H | H | H | H | H | H | H | H | H | H | H | H | H | H | H | H | H | H | H | H | H | H | H | T | G | H | H | G | C | H | H | H | C | H | H | H | H | H | G | H | H | H | H | H | H | H | A | G | G | H | H | H | H | G | H | G | H | H | H | H | H | H | H | H | H | H | H | H | H | C | H | H | H | H | H | H | H | A | A | T | H | A | G | T | H | C | T | T | G | C | C | C | H |
| 2016A_P8-A1 | PHI5 | PHI1 | mother | 2016A_P8-A1 | | H | H | H | G | H | H | H | H | H | G | H | H | G | A | H | T | C | H | H | H | H | H | H | H | H | H | H | H | H | H | H | H | H | T | G | H | H | G | C | H | H | H | C | H | H | H | H | H | G | H | H | H | H | H | H | H | A | G | G | H | H | H | H | G | H | G | H | H | H | H | H | H | H | H | H | H | H | H | H | C | H | H | H | H | H | H | A | H | H | H | H | H | G | T | H | C | T | T | G | C | C | C | H |
| 2016A_P8-A1_1 | PHI5 | PHI1 | progeny | 2016A_P8-A1 | | C | T | H | G | G | G | G | H | H | G | H | H | G | A | H | T | C | H | H | H | H | H | H | H | H | H | H | H | H | H | H | H | H | T | G | H | H | G | C | H | H | H | C | H | H | H | H | H | G | H | H | H | H | H | H | H | A | G | G | H | H | H | H | G | H | G | H | H | H | H | H | H | H | H | H | H | H | H | H | C | H | H | H | H | H | H | H | H | H | H | H | H | G | T | H | C | T | T | G | C | C | C | H |
| 2016A_P8-A5 | PHI5 | PHI1 | mother | 2016A_P8-A5 | | H | T | C | G | H | H | H | H | H | H | H | H | H | H | H | H | H | H | H | H | H | H | H | H | H | H | H | H | H | H | H | H | H | H | G | H | H | G | C | H | H | H | C | H | H | H | H | H | G | H | H | H | H | H | H | H | A | G | G | H | H | H | H | H | H | H | H | H | H | H | H | H | H | H | H | H | H | H | H | H | H | H | H | H | H | H | H | H | H | H | H | H | H | H | H | H | T | H | H | H | H | H | H |
| 2016A_P8-A5_1 | PHI5 | PHI1 | progeny | 2016A_P8-A5 | | H | T | H | G | H | H | H | H | H | G | H | H | H | H | H | H | H | H | H | H | H | H | H | H | C | H | H | H | H | H | H | H | H | H | H | H | H | G | C | H | H | H | H | H | H | H | H | H | H | H | H | H | H | H | H | H | H | G | H | H | H | H | H | H | H | H | H | H | H | H | H | H | H | H | H | H | H | H | H | H | H | H | H | H | H | H | H | H | H | H | H | H | H | H | H | H | H | H | H | H | H | H | H |
| 2016A_P8-A6 | PHI5 | PHI1 | mother | 2016A_P8-A6 | | H | H | H | G | H | H | H | H | H | H | H | H | H | H | H | H | H | H | H | H | H | H | H | H | H | H | H | H | H | H | H | H | H | H | G | H | H | G | C | H | H | H | C | H | H | H | H | H | G | H | H | H | H | H | H | H | A | G | G | H | H | H | H | H | H | H | H | H | H | H | H | H | H | H | H | H | H | H | H | H | H | H | H | H | H | H | H | H | H | H | H | H | G | H | H | H | H | T | H | H | H | H | H |
| 2016A_P8-A6_1 | PHI5 | PHI1 | progeny | 2016A_P8-A6 | | H | H | H | G | C | A | A | A | A | A | G | A | A | C | G | C | G | T | A | A | G | A | T | C | C | T | T | C | T | G | A | T | H | H | G | H | H | G | C | T | G | C | C | H | H | H | H | H | G | H | H | C | A | A | G | G | A | G | G | H | T | G | A | H | H | H | H | H | H | H | H | H | H | H | H | H | H | H | H | H | H | H | H | H | H | H | H | G | G | C | A | G | A | G | G | G | T | C | A | T | T | T | T |
| 2016A_P8-D5 | PHI9 | PHI1 | mother | 2016A_P8-D5 | | T | T | H | G | G | G | G | G | G | G | H | H | G | A | H | T | C | C | A | H | H | H | H | C | H | T | H | C | H | H | H | H | H | H | G | H | H | G | H | T | H | C | H | C | G | H | H | H | H | H | H | H | H | H | H | A | H | G | H | H | H | H | H | H | H | H | H | H | H | H | H | G | H | H | H | H | H | H | H | H | H | T | H | A | H | H | H | H | A | H | T | H | A | G | T | H | C | H | T | G | C | C | C | H |
| 2016A_P8-D5_3 | PHI9 | PHI1 | progeny | 2016A_P8-D5 | | T | T | H | G | G | G | G | G | G | G | H | H | G | A | H | T | C | C | A | H | H | H | H | C | H | T | H | C | H | H | H | H | H | H | G | C | G | G | C | T | G | C | C | C | H | C | A | T | A | T | G | C | A | A | G | A | G | G | C | C | T | G | A | A | C | T | T | G | A | T | G | C | G | G | C | C | T | T | T | G | T | H | A | H | H | H | H | A | H | T | H | A | G | T | H | C | H | T | G | C | C | C | H |
| 2016A_P8-D5_5 | PHI9 | PHI1 | progeny | 2016A_P8-D5 | | T | T | H | G | G | G | G | G | G | G | H | H | G | A | H | T | C | C | A | H | G | A | T | C | C | T | T | C | T | H | H | H | H | H | G | T | C | G | T | T | T | C | T | C | G | H | H | H | H | H | H | H | H | H | A | A | A | G | G | T | H | C | C | G | G | G | A | A | G | C | G | T | T | A | T | T | G | H | C | C | T | H | A | H | H | G | A | A | A | T | A | A | G | T | G | C | T | T | G | C | C | C | T |
| 2016A_P8-E1 | PHI9 | PHI1 | mother | 2016A_P8-E1 | | T | T | H | G | G | G | G | G | G | G | H | H | G | A | H | T | C | H | A | H | H | H | H | C | H | T | H | C | H | H | H | H | H | H | G | H | H | G | H | T | H | C | H | H | H | H | H | H | H | H | H | T | H | H | H | A | H | G | H | H | H | G | A | H | H | H | H | H | H | H | G | H | H | H | H | H | H | G | H | H | T | H | A | H | H | A | H | H | G | H | H | H | H | H | H | H | T | H | H | H | H | H | H |
| 2016A_P8-E1_2 | PHI9 | PHI1 | progeny | 2016A_P8-E1 | | T | T | H | G | G | G | G | G | G | G | H | H | G | A | H | T | C | H | A | H | H | H | H | C | H | T | H | C | H | H | H | H | H | H | G | H | H | G | H | T | H | C | H | C | G | H | H | H | H | H | H | T | H | H | H | A | H | G | H | H | H | G | A | H | H | H | H | H | H | H | G | H | H | H | H | H | H | G | H | H | T | H | A | H | H | A | H | A | H | T | H | A | H | H | H | H | H | H | G | C | C | C | H |
| 2016A_P9-A10 | PHI9 | PHI1 | mother | 2016A_P9-A10 | | T | T | H | G | G | G | G | G | G | G | H | H | G | A | H | T | C | C | A | H | H | H | H | C | H | T | H | C | H | H | H | H | H | H | G | H | H | G | H | T | H | C | H | C | G | H | H | H | H | H | H | T | H | H | H | A | H | G | H | H | H | G | A | H | H | H | H | H | H | H | G | H | H | H | H | H | H | G | H | H | T | H | A | H | H | H | H | H | G | H | H | H | H | H | H | H | H | H | H | H | H | H | H |
| 2016A_P9-A10_1 | PHI9 | PHI1 | progeny | 2016A_P9-A10 | | T | T | T | G | G | G | G | G | G | G | A | C | G | A | A | T | C | C | A | T | T | G | C | C | T | T | A | C | C | T | G | A | C | A | G | T | C | G | T | T | T | C | T | C | G | T | G | C | G | C | A | T | A | A | G | A | G | G | C | C | T | G | A | A | C | T | T | G | A | T | G | C | G | G | C | T | G | G | C | C | T | A | A | A | G | A | G | A | G | T | G | A | G | T | T | C | G | T | G | C | C | C | C |
| 2016A_P9-A10_3 | PHI9 | PHI1 | progeny | 2016A_P9-A10 | | T | T | T | G | G | G | G | G | G | G | A | C | G | A | A | T | C | C | A | T | T | G | C | C | T | T | A | C | C | T | G | A | C | A | G | C | G | G | C | T | G | C | C | C | G | C | A | T | A | T | G | T | A | A | G | A | G | G | C | C | T | G | A | A | C | T | T | G | A | T | G | C | G | G | C | C | T | G | T | G | T | G | A | G | A | G | A | G | G | C | A | G | A | G | G | G | T | C | A | T | T | T | T |
| 2016A_P9-A7 | PHI9 | PHI1 | mother | 2016A_P9-A7 | | T | T | H | G | G | G | G | G | G | G | H | H | G | A | H | T | C | C | A | H | H | H | H | C | H | T | H | C | H | H | H | H | H | H | G | H | H | G | H | T | H | C | H | H | H | H | H | H | H | H | H | T | H | H | H | A | H | G | H | H | H | G | A | H | H | H | H | H | H | H | G | H | H | H | H | H | H | G | H | H | T | H | A | H | H | A | H | A | H | T | H | A | G | T | H | H | H | H | H | H | H | H | H |
| 2016A_P9-A7_1 | PHI9 | PHI1 | progeny | 2016A_P9-A7 | | T | T | H | G | G | G | G | G | G | G | H | H | G | A | H | T | C | C | A | H | H | H | H | C | H | T | H | C | H | H | H | H | H | H | G | C | H | G | H | T | H | C | H | H | H | H | H | H | H | H | A | T | G | G | A | A | H | G | H | H | H | G | A | H | H | H | H | H | H | H | G | T | H | H | H | H | H | G | T | H | T | H | A | H | H | A | H | A | G | T | H | A | G | T | H | H | H | H | H | H | H | H | H |
| 2016A_P9-A7_2 | PHI9 | PHI1 | progeny | 2016A_P9-A7 | | T | T | C | G | G | G | G | G | G | G | G | A | G | A | G | T | C | C | A | H | G | H | H | C | H | T | H | C | H | H | H | H | H | H | G | H | H | G | H | T | H | C | H | H | H | H | H | H | H | H | H | T | H | H | H | A | H | G | H | H | H | G | A | H | H | H | H | H | H | H | G | H | H | H | H | H | H | G | H | C | T | H | A | H | H | A | G | A | H | T | H | A | G | T | H | H | H | H | H | H | H | H | H |
| 2016A_P9-B1 | PHI9 | PHI1 | mother | 2016A_P9-B1 | | T | T | H | G | G | G | G | G | G | G | H | H | H | A | G | T | C | H | A | H | H | H | H | C | H | T | H | C | H | H | H | H | H | H | G | H | H | G | H | T | H | C | H | C | G | H | H | H | H | H | H | T | H | H | H | A | H | G | H | H | H | H | H | G | H | G | H | H | H | H | H | H | H | H | H | H | H | G | H | H | T | H | A | H | H | H | H | H | G | H | H | H | H | H | H | H | T | H | H | H | H | H | H |
| 2016A_P9-B1_2 | PHI9 | PHI1 | progeny | 2016A_P9-B1 | | T | T | T | G | G | G | G | G | G | G | A | C | A | A | G | T | C | T | A | T | T | G | C | C | T | T | A | C | C | G | A | T | T | T | G | C | G | G | C | T | T | C | T | C | G | T | G | C | G | T | G | T | A | A | G | A | G | G | C | C | T | G | A | G | C | G | T | G | A | T | T | C | G | G | C | C | T | G | T | G | T | A | A | G | A | G | A | G | G | C | A | G | A | G | G | G | T | C | A | T | T | T | T |
| 2016A_P9-B7 | PHI9 | PHI1 | mother | 2016A_P9-B7 | | T | H | T | G | H | H | H | H | H | G | H | H | G | A | H | T | C | C | A | H | H | H | H | C | H | T | H | C | H | H | H | H | H | H | G | H | H | G | H | T | H | C | H | H | H | H | H | H | H | H | H | H | H | H | H | A | H | G | H | H | H | H | A | H | H | H | H | H | H | H | G | H | H | H | H | H | H | H | H | H | T | H | A | H | H | H | H | H | G | H | H | H | H | H | H | H | H | H | H | H | H | H | H |
| 2016A_P9-B7_10 | PHI9 | PHI1 | progeny | 2016A_P9-B7 | | T | T | T | G | G | G | G | G | G | G | A | C | G | A | A | T | C | C | A | A | G | A | T | C | C | T | A | C | T | G | A | T | T | T | G | C | G | G | C | T | G | C | C | C | G | C | A | T | A | T | G | C | A | A | G | A | G | G | C | C | T | G | A | A | C | T | T | G | A | T | G | C | G | G | C | C | T | T | T | G | T | G | A | A | A | G | A | G | G | C | A | G | A | G | G | G | T | C | A | T | T | T | T |
| 2016A_P9-B7_5 | PHI9 | PHI1 | progeny | 2016A_P9-B7 | | T | G | T | G | C | A | A | G | G | G | A | C | G | A | A | T | C | C | A | T | T | G | C | C | T | T | A | C | C | T | G | A | C | A | G | T | C | G | T | T | T | C | C | A | A | T | G | C | G | C | A | T | A | G | A | A | A | G | G | T | C | C | A | G | G | G | A | A | G | C | G | T | T | A | T | T | G | G | C | C | T | G | A | A | G | A | G | A | G | T | G | A | A | G | G | G | T | C | A | T | T | T | T |
| 2016A_P9-B7_9 | PHI9 | PHI1 | progeny | 2016A_P9-B7 | | T | G | T | G | G | G | G | G | G | G | A | C | G | A | A | T | C | C | A | T | T | G | T | C | C | T | T | C | T | G | A | T | T | T | G | C | G | G | C | T | G | C | C | C | G | C | A | T | A | T | G | C | A | A | G | A | A | G | G | T | C | C | A | G | G | G | A | A | G | C | G | T | T | A | T | T | G | G | T | G | T | G | A | A | G | A | G | A | G | T | G | A | G | T | T | C | G | T | G | C | C | C | C |

plant_id – unique identifier of the plant
mom_gen – genetic backgroiund of mother in the hybred
dad_gen – genetic background of father in the hybred
family_ – identify mother and all the progeny plants from the same mother
mother_progeny – identify each plant as mother of the family or progeny
Chr – number of the chromosome, where marker is located
Distance – position of the marker on the corresponding chromosome in cM
H – heterozygous marker
A, T, G and C – marker homozygous for the corresponding nucleotide
EQV, LS, -9 , etc. – missing values

**Table S5. Results of the SNP array analasis (continue)**

| | | | | | | | | | |
|---|---|---|---|---|---|---|---|---|---|
| | | | | | Chr | 3 | 3 | 3 | [illegible] |
| | | | | | Position, cM | 94 | 95 | 95 | [illegible] |
| plant_id | mom_gen | dad_gen | mother_progeny | family_ | | | | | |
| 2016A_P1-A3 | PHI1 | PHI3 | mother | 2016A_P1-A3 | | H | H | H | [illegible] |
| 2016A_P1-A3_1 | PHI1 | PHI3 | progeny | 2016A_P1-A3 | | T | C | A | [illegible] |
| 2016A_P1-D5 | PHI1 | PHI3 | mother | 2016A_P1-D5 | | H | H | H | [illegible] |
| 2016A_P1-D5_1 | PHI1 | PHI3 | progeny | 2016A_P1-D5 | | H | H | H | [illegible] |
| 2016A_P1-D8 | PHI1 | PHI3 | mother | 2016A_P1-D8 | | H | H | H | [illegible] |
| 2016A_P1-D8_3 | PHI1 | PHI3 | progeny | 2016A_P1-D8 | | H | H | H | [illegible] |
| 2016A_P1-E3 | PHI1 | PHI3 | mother | 2016A_P1-E3 | | H | H | H | [illegible] |
| 2016A_P1-E3_3 | PHI1 | PHI3 | progeny | 2016A_P1-E3 | | T | C | A | [illegible] |
| 2016A_P1-E3_5 | PHI1 | PHI3 | progeny | 2016A_P1-E3 | | T | C | A | [illegible] |
| 2016A_P1-E9 | PHI1 | PHI3 | mother | 2016A_P1-E9 | | H | H | H | [illegible] |
| 2016A_P1-E9_1 | PHI1 | PHI3 | progeny | 2016A_P1-E9 | | C | T | G | [illegible] |
| 2016A_P1-F7 | PHI1 | PHI3 | mother | 2016A_P1-F7 | | H | H | H | [illegible] |
| 2016A_P1-F7_4 | PHI1 | PHI3 | progeny | 2016A_P1-F7 | | C | T | G | [illegible] |
| 2016A_P1-F8 | PHI2 | PHI1 | mother | 2016A_P1-F8 | | H | T | H | [illegible] |
| 2016A_P1-F8_4 | PHI2 | PHI1 | progeny | 2016A_P1-F8 | | C | T | G | [illegible] |
| 2016A_P1-G12 | PHI2 | PHI1 | mother | 2016A_P1-G12 | | H | T | H | [illegible] |
| 2016A_P1-G12_1 | PHI2 | PHI1 | progeny | 2016A_P1-G12 | | T | T | A | [illegible] |
| 2016A_P1-G12_2 | PHI2 | PHI1 | progeny | 2016A_P1-G12 | | H | T | H | [illegible] |
| 2016A_P1-G12_6 | PHI2 | PHI1 | progeny | 2016A_P1-G12 | | H | T | H | [illegible] |
| 2016A_P1-H11 | PHI2 | PHI1 | mother | 2016A_P1-H11 | | H | T | H | [illegible] |
| 2016A_P1-H11_3 | PHI2 | PHI1 | progeny | 2016A_P1-H11 | | H | T | H | [illegible] |
| 2016A_P1-H11_4 | PHI2 | PHI1 | progeny | 2016A_P1-H11 | | T | T | A | [illegible] |
| 2016A_P1-H8 | PHI2 | PHI1 | mother | 2016A_P1-H8 | | H | T | H | [illegible] |
| 2016A_P1-H8_1 | PHI2 | PHI1 | progeny | 2016A_P1-H8 | | H | T | H | [illegible] |
| 2016A_P1-H8_16 | PHI2 | PHI1 | progeny | 2016A_P1-H8 | | H | T | H | [illegible] |
| 2016A_P1-H8_4 | PHI2 | PHI1 | progeny | 2016A_P1-H8 | | H | T | H | [illegible] |
| 2016A_P1-H8_5 | PHI2 | PHI1 | progeny | 2016A_P1-H8 | | H | T | H | [illegible] |
| 2016A_P1-H8_7 | PHI2 | PHI1 | progeny | 2016A_P1-H8 | | H | T | H | [illegible] |
| 2016A_P2-A11 | PHI2 | PHI1 | mother | 2016A_P2-A11 | | H | T | H | [illegible] |
| 2016A_P2-A11_4 | PHI2 | PHI1 | progeny | 2016A_P2-A11 | | H | T | H | [illegible] |
| 2016A_P2-A11_5 | PHI2 | PHI1 | progeny | 2016A_P2-A11 | | C | T | G | [illegible] |
| 2016A_P2-A12 | PHI2 | PHI1 | mother | 2016A_P2-A12 | | H | T | H | [illegible] |
| 2016A_P2-A12_12 | PHI2 | PHI1 | progeny | 2016A_P2-A12 | | H | H | H | [illegible] |
| 2016A_P2-A12_4 | PHI2 | PHI1 | progeny | 2016A_P2-A12 | | T | T | A | [illegible] |
| 2016A_P2-A12_7 | PHI2 | PHI1 | progeny | 2016A_P2-A12 | | C | T | G | [illegible] |
| 2016A_P2-A12_8 | PHI2 | PHI1 | progeny | 2016A_P2-A12 | | H | T | H | [illegible] |
| 2016A_P2-A12_9 | PHI2 | PHI1 | progeny | 2016A_P2-A12 | | H | T | A | [illegible] |
| 2016A_P2-A7 | PHI2 | PHI1 | mother | 2016A_P2-A7 | | H | T | H | [illegible] |
| 2016A_P2-A7_9 | PHI2 | PHI1 | progeny | 2016A_P2-A7 | | T | T | A | [illegible] |
| 2016A_P2-B10 | PHI2 | PHI1 | mother | 2016A_P2-B10 | | T | T | A | [illegible] |
| 2016A_P2-B10_6 | PHI2 | PHI1 | progeny | 2016A_P2-B10 | | T | T | A | [illegible] |
| 2016A_P2-B4 | PHI2 | PHI1 | mother | 2016A_P2-B4 | | H | T | H | [illegible] |
| 2016A_P2-B4_1 | PHI2 | PHI1 | progeny | 2016A_P2-B4 | | C | T | G | [illegible] |
| 2016A_P2-B4_5 | PHI2 | PHI1 | progeny | 2016A_P2-B4 | | T | T | A | [illegible] |
| 2016A_P2-B4_6 | PHI2 | PHI1 | progeny | 2016A_P2-B4 | | T | T | A | [illegible] |
| 2016A_P2-B7 | PHI2 | PHI1 | mother | 2016A_P2-B7 | | H | T | H | [illegible] |
| 2016A_P2-B7_1 | PHI2 | PHI1 | progeny | 2016A_P2-B7 | | H | T | H | [illegible] |
| 2016A_P2-C11 | PHI2 | PHI1 | mother | 2016A_P2-C11 | | H | T | H | [illegible] |
| 2016A_P2-C11_1 | PHI2 | PHI1 | progeny | 2016A_P2-C11 | | H | T | A | [illegible] |
| 2016A_P2-C11_4 | PHI2 | PHI1 | progeny | 2016A_P2-C11 | | T | T | A | [illegible] |
| 2016A_P2-D1 | PHI2 | PHI1 | mother | 2016A_P2-D1 | | H | T | H | [illegible] |
| 2016A_P2-D1_10 | PHI2 | PHI1 | progeny | 2016A_P2-D1 | | T | T | A | [illegible] |
| 2016A_P2-D1_12 | PHI2 | PHI1 | progeny | 2016A_P2-D1 | | T | T | A | [illegible] |
| 2016A_P2-D1_3 | PHI2 | PHI1 | progeny | 2016A_P2-D1 | | H | T | H | [illegible] |
| 2016A_P2-D1_4 | PHI2 | PHI1 | progeny | 2016A_P2-D1 | | C | T | G | [illegible] |
| 2016A_P2-D1_7 | PHI2 | PHI1 | progeny | 2016A_P2-D1 | | H | T | H | [illegible] |
| 2016A_P2-D1_8 | PHI2 | PHI1 | progeny | 2016A_P2-D1 | | H | T | H | [illegible] |
| 2016A_P2-D10 | PHI2 | PHI1 | mother | 2016A_P2-D10 | | T | T | A | [illegible] |
| 2016A_P2-D10_3 | PHI2 | PHI1 | progeny | 2016A_P2-D10 | | T | T | A | [illegible] |
| 2016A_P2-D5 | PHI2 | PHI1 | mother | 2016A_P2-D5 | | H | T | H | [illegible] |
| 2016A_P2-D5_4 | PHI2 | PHI1 | progeny | 2016A_P2-D5 | | H | T | H | [illegible] |
| 2016A_P2-D7 | PHI2 | PHI1 | mother | 2016A_P2-D7 | | H | T | H | [illegible] |
| 2016A_P2-D7_3 | PHI2 | PHI1 | progeny | 2016A_P2-D7 | | H | T | H | [illegible] |
| 2016A_P2-F2 | PHI2 | PHI1 | mother | 2016A_P2-F2 | | H | T | H | [illegible] |
| 2016A_P2-F2_2 | PHI2 | PHI1 | progeny | 2016A_P2-F2 | | C | T | H | [illegible] |
| 2016A_P2-F7 | PHI2 | PHI1 | mother | 2016A_P2-F7 | | H | T | H | [illegible] |
| 2016A_P2-F7_1 | PHI2 | PHI1 | progeny | 2016A_P2-F7 | | C | T | G | [illegible] |
| 2016A_P2-G1 | PHI2 | PHI1 | mother | 2016A_P2-G1 | | H | T | H | [illegible] |
| 2016A_P2-G1_4 | PHI2 | PHI1 | progeny | 2016A_P2-G1 | | T | T | A | [illegible] |
| 2016A_P2-G1_6 | PHI2 | PHI1 | progeny | 2016A_P2-G1 | | T | H | H | [illegible] |
| 2016A_P2-G2 | PHI2 | PHI1 | mother | 2016A_P2-G2 | | H | T | H | [illegible] |
| 2016A_P2-G2_5 | PHI2 | PHI1 | progeny | 2016A_P2-G2 | | C | T | G | [illegible] |
| 2016A_P2-G2_6 | PHI2 | PHI1 | progeny | 2016A_P2-G2 | | C | T | G | [illegible] |
| 2016A_P2-G7 | PHI2 | PHI1 | mother | 2016A_P2-G7 | | H | T | H | [illegible] |
| 2016A_P2-G7_13 | PHI2 | PHI1 | progeny | 2016A_P2-G7 | | H | T | H | [illegible] |
| 2016A_P2-G7_15 | PHI2 | PHI1 | progeny | 2016A_P2-G7 | | H | T | H | [illegible] |
| 2016A_P2-G7_6 | PHI2 | PHI1 | progeny | 2016A_P2-G7 | | H | T | H | [illegible] |
| 2016A_P2-G7_8 | PHI2 | PHI1 | progeny | 2016A_P2-G7 | | H | T | H | [illegible] |
| 2016A_P2-G9 | PHI2 | PHI1 | mother | 2016A_P2-G9 | | T | T | A | [illegible] |
| 2016A_P2-G9_3 | PHI2 | PHI1 | progeny | 2016A_P2-G9 | | T | T | A | [illegible] |
| 2016A_P2-G9_5 | PHI2 | PHI1 | progeny | 2016A_P2-G9 | | T | T | A | [illegible] |
| 2016A_P2-G9_6 | PHI2 | PHI1 | progeny | 2016A_P2-G9 | | T | T | A | [illegible] |
| 2016A_P2-H6 | PHI2 | PHI1 | mother | 2016A_P2-H6 | | H | T | H | [illegible] |
| 2016A_P2-H6_1 | PHI2 | PHI1 | progeny | 2016A_P2-H6 | | C | T | G | [illegible] |
| 2016A_P2-H6_16 | PHI2 | PHI1 | progeny | 2016A_P2-H6 | | H | T | H | [illegible] |
| 2016A_P2-H6_17 | PHI2 | PHI1 | progeny | 2016A_P2-H6 | | H | T | H | [illegible] |
| 2016A_P2-H6_9 | PHI2 | PHI1 | progeny | 2016A_P2-H6 | | H | T | H | [illegible] |
| 2016A_P3-B5 | PHI10 | PHI1 | mother | 2016A_P3-B5 | | H | H | H | [illegible] |
| 2016A_P3-B5_1 | PHI10 | PHI1 | progeny | 2016A_P3-B5 | | C | T | G | [illegible] |
| 2016A_P3-B8 | PHI10 | PHI1 | mother | 2016A_P3-B8 | | H | H | H | [illegible] |
| 2016A_P3-B8_1 | PHI10 | PHI1 | progeny | 2016A_P3-B8 | | C | T | G | [illegible] |
| 2016A_P3-B8_2 | PHI10 | PHI1 | progeny | 2016A_P3-B8 | | T | C | A | [illegible] |
| 2016A_P3-B8_4 | PHI10 | PHI1 | progeny | 2016A_P3-B8 | | C | T | G | [illegible] |
| 2016A_P3-C12 | PHI10 | PHI1 | mother | 2016A_P3-C12 | | H | H | H | [illegible] |
| 2016A_P3-C12_2 | PHI10 | PHI1 | progeny | 2016A_P3-C12 | | C | T | G | [illegible] |
| 2016A_P3-D9 | PHI10 | PHI1 | mother | 2016A_P3-D9 | | H | H | H | [illegible] |
| 2016A_P3-D9_2 | PHI10 | PHI1 | progeny | 2016A_P3-D9 | | C | T | G | [illegible] |
| 2016A_P3-H12 | PHI4 | PHI1 | mother | 2016A_P3-H12 | | H | T | H | [illegible] |
| 2016A_P3-H12_1 | PHI4 | PHI1 | progeny | 2016A_P3-H12 | | H | T | H | [illegible] |
| 2016A_P3-H8 | PHI4 | PHI1 | mother | 2016A_P3-H8 | | H | T | H | [illegible] |
| 2016A_P3-H8_1 | PHI4 | PHI1 | progeny | 2016A_P3-H8 | | H | T | H | [illegible] |
| 2016A_P4-A1 | PHI4 | PHI1 | mother | 2016A_P4-A1 | | H | H | G | [illegible] |
| 2016A_P4-A1_1 | PHI4 | PHI1 | progeny | 2016A_P4-A1 | | H | H | G | [illegible] |
| 2016A_P4-A3 | PHI4 | PHI1 | mother | 2016A_P4-A3 | | H | H | G | [illegible] |
| 2016A_P4-A3_1 | PHI4 | PHI1 | progeny | 2016A_P4-A3 | | H | H | G | [illegible] |
| 2016A_P4-A3_5 | PHI4 | PHI1 | progeny | 2016A_P4-A3 | | T | H | G | [illegible] |
| 2016A_P4-B4 | PHI4 | PHI1 | mother | 2016A_P4-B4 | | H | H | G | [illegible] |
| 2016A_P4-B4_3 | PHI4 | PHI1 | progeny | 2016A_P4-B4 | | H | H | G | [illegible] |
| 2016A_P4-C11 | PHI4 | PHI1 | mother | 2016A_P4-C11 | | H | H | H | [illegible] |
| 2016A_P4-C11_3 | PHI4 | PHI1 | progeny | 2016A_P4-C11 | | H | H | G | [illegible] |
| 2016A_P4-C2 | PHI4 | PHI1 | mother | 2016A_P4-C2 | | H | T | H | [illegible] |
| 2016A_P4-C2_2 | PHI4 | PHI1 | progeny | 2016A_P4-C2 | | C | T | G | [illegible] |
| 2016A_P4-C2_3 | PHI4 | PHI1 | progeny | 2016A_P4-C2 | | H | T | A | [illegible] |
| 2016A_P4-C2_5 | PHI4 | PHI1 | progeny | 2016A_P4-C2 | | T | T | A | [illegible] |
| 2016A_P4-C2_6 | PHI4 | PHI1 | progeny | 2016A_P4-C2 | | T | T | A | [illegible] |
| 2016A_P4-C4 | PHI4 | PHI1 | mother | 2016A_P4-C4 | | H | H | G | [illegible] |
| 2016A_P4-C4_9 | PHI4 | PHI1 | progeny | 2016A_P4-C4 | | H | H | G | [illegible] |
| 2016A_P4-C9 | PHI4 | PHI1 | mother | 2016A_P4-C9 | | H | H | G | [illegible] |
| 2016A_P4-C9_1 | PHI4 | PHI1 | progeny | 2016A_P4-C9 | | H | H | G | [illegible] |
| 2016A_P4-C9_3 | PHI4 | PHI1 | progeny | 2016A_P4-C9 | | H | H | G | [illegible] |
| 2016A_P4-D12 | PHI4 | PHI1 | mother | 2016A_P4-D12 | | H | H | G | [illegible] |
| 2016A_P4-D12_5 | PHI4 | PHI1 | progeny | 2016A_P4-D12 | | H | H | G | [illegible] |
| 2016A_P4-D5 | PHI4 | PHI1 | mother | 2016A_P4-D5 | | H | H | G | [illegible] |
| 2016A_P4-D5_1 | PHI4 | PHI1 | progeny | 2016A_P4-D5 | | H | H | G | [illegible] |
| 2016A_P4-F5 | PHI4 | PHI1 | mother | 2016A_P4-F5 | | H | H | G | [illegible] |
| 2016A_P4-F5_1 | PHI4 | PHI1 | progeny | 2016A_P4-F5 | | C | T | G | [illegible] |
| 2016A_P4-F5_2 | PHI4 | PHI1 | progeny | 2016A_P4-F5 | | H | H | G | [illegible] |
| 2016A_P4-F5_5 | PHI4 | PHI1 | progeny | 2016A_P4-F5 | | H | C | G | [illegible] |
| 2016A_P4-F5_6 | PHI4 | PHI1 | progeny | 2016A_P4-F5 | | H | H | G | [illegible] |
| 2016A_P4-F7 | PHI7 | PHI1 | mother | 2016A_P4-F7 | | H | H | H | [illegible] |
| 2016A_P4-F7_1 | PHI7 | PHI1 | progeny | 2016A_P4-F7 | | T | C | A | [illegible] |
| 2016A_P4-G11 | PHI7 | PHI1 | mother | 2016A_P4-G11 | | H | H | H | [illegible] |
| 2016A_P4-G11_2 | PHI7 | PHI1 | progeny | 2016A_P4-G11 | | H | H | H | [illegible] |
| 2016A_P4-G11_4 | PHI7 | PHI1 | progeny | 2016A_P4-G11 | | H | H | H | [illegible] |

### Table S5. Results of the SNP array analasis (continue)

| Chr | 3 | 3 | 3 | 3 | 3 | 3 | 3 | 3 | 3 | 3 | 3 | 3 | 3 | 3 | 3 | 3 | 3 | 4 | 4 | 4 | 4 | 4 | 4 | 4 | 4 | 4 | 4 | 4 | 4 | 4 | 4 | 4 | 4 | 4 | 4 | 4 | 4 | 4 | 4 | 4 | 4 | 4 | 4 | 5 | 5 | 5 | 5 | 5 | 5 | 5 | 5 | 5 | 5 | 5 | 5 | 5 | 5 | 5 | 5 | 5 | 5 | 5 | 5 | 5 | 5 | 5 | 5 | 5 | 5 | 5 | 5 | 5 | 5 | 6 | 6 | 6 | 6 | 6 | 6 | 6 | 6 | 6 | 6 | 6 | 6 | 6 | 6 | 6 | 6 | 6 | 6 | 6 | 6 | 6 | 6 | 6 | 7 | 7 | 7 | 7 | 7 | 7 | 7 |
|---|---|---|---|---|---|---|---|---|---|---|---|---|---|---|---|---|---|---|---|---|---|---|---|---|---|---|---|---|---|---|---|---|---|---|---|---|---|---|---|---|---|---|---|---|---|---|---|---|---|---|---|---|---|---|---|---|---|---|---|---|---|---|---|---|---|---|---|---|---|---|---|---|---|---|---|---|---|---|---|---|---|---|---|---|---|---|---|---|---|---|---|---|---|---|---|---|---|---|---|---|---|---|---|
| Position, cM | 94 | 95 | 95 | 104 | 113 | 123 | 135 | 149 | 160 | 168 | 182 | 195 | 203 | 224 | 227 | 240 | 243 | 7.4 | 18 | 30 | 33 | 42 | 50 | 75 | 85 | 89 | 104 | 108 | 110 | 111 | 110 | 119 | 128 | 140 | 155 | 163 | 163 | 176 | 188 | 197 | 216 | 238 | 246 | 5.3 | 12 | 21 | 32 | 48 | 51 | 65 | 71 | 89 | 93 | 97 | 98 | 101 | 102 | 102 | 102 | 105 | 111 | 124 | 138 | 148 | 159 | 166 | 177 | 185 | 191 | 198 | 208 | 215 | 224 | 0.3 | 14 | 16 | 22 | 23 | 24 | 23 | 24 | 38 | 52 | 54 | 72 | 79 | 93 | 98 | 114 | 129 | 130 | 135 | 143 | 157 | 180 | 181 | 1.3 | 14 | 26 | 33 | 50 | 62 | 68 |

| plant_id | mom_gen | dad_gen | mother_progeny | family_ | 94 | 95 | 95 |
|---|---|---|---|---|---|---|---|
| 2016A_P4-H1 | PHI7 | PHI1 | mother | 2016A_P4-H1 | H | H | H |
| 2016A_P4-H1_2 | PHI7 | PHI1 | progeny | 2016A_P4-H1 | C | T | G |
| 2016A_P4-H1_3 | PHI7 | PHI1 | progeny | 2016A_P4-H1 | H | H | H |
| 2016A_P4-H1_4 | PHI7 | PHI1 | progeny | 2016A_P4-H1 | H | H | H |
| 2016A_P4-H10 | PHI7 | PHI1 | mother | 2016A_P4-H10 | H | H | H |
| 2016A_P4-H10_4 | PHI7 | PHI1 | progeny | 2016A_P4-H10 | H | H | H |
| 2016A_P4-H5 | PHI7 | PHI1 | mother | 2016A_P4-H5 | H | H | H |
| 2016A_P4-H5_1 | PHI7 | PHI1 | progeny | 2016A_P4-H5 | H | H | H |
| 2016A_P4-H5_2 | PHI7 | PHI1 | progeny | 2016A_P4-H5 | H | H | H |
| 2016A_P5-A1 | PHI7 | PHI1 | mother | 2016A_P5-A1 | H | H | H |
| 2016A_P5-A1_3 | PHI7 | PHI1 | progeny | 2016A_P5-A1 | C | T | G |
| 2016A_P5-A1_4 | PHI7 | PHI1 | progeny | 2016A_P5-A1 | T | C | A |
| 2016A_P5-B11 | PHI7 | PHI1 | mother | 2016A_P5-B11 | H | H | H |
| 2016A_P5-B11_1 | PHI7 | PHI1 | progeny | 2016A_P5-B11 | H | H | H |
| 2016A_P5-B11_7 | PHI7 | PHI1 | progeny | 2016A_P5-B11 | C | H | H |
| 2016A_P5-C8 | PHI7 | PHI1 | mother | 2016A_P5-C8 | H | H | H |
| 2016A_P5-C8_1 | PHI7 | PHI1 | progeny | 2016A_P5-C8 | H | H | H |
| 2016A_P5-C8_2 | PHI7 | PHI1 | progeny | 2016A_P5-C8 | C | T | G |
| 2016A_P5-F1 | PHI6 | PHI1 | mother | 2016A_P5-F1 | H | T | H |
| 2016A_P5-F1_1 | PHI6 | PHI1 | progeny | 2016A_P5-F1 | H | T | H |
| 2016A_P5-F10 | PHI6 | PHI1 | mother | 2016A_P5-F10 | H | H | H |
| 2016A_P5-F10_2 | PHI6 | PHI1 | progeny | 2016A_P5-F10 | H | H | H |
| 2016A_P5-F10_4 | PHI6 | PHI1 | progeny | 2016A_P5-F10 | H | H | H |
| 2016A_P5-F10_7 | PHI6 | PHI1 | progeny | 2016A_P5-F10 | H | H | H |
| 2016A_P5-F2 | PHI6 | PHI1 | mother | 2016A_P5-F2 | H | H | H |
| 2016A_P5-F2_3 | PHI6 | PHI1 | progeny | 2016A_P5-F2 | T | C | A |
| 2016A_P5-G10 | PHI6 | PHI1 | mother | 2016A_P5-G10 | H | T | H |
| 2016A_P5-G10_10 | PHI6 | PHI1 | progeny | 2016A_P5-G10 | H | T | H |
| 2016A_P5-G10_11 | PHI6 | PHI1 | progeny | 2016A_P5-G10 | H | T | H |
| 2016A_P5-G10_6 | PHI6 | PHI1 | progeny | 2016A_P5-G10 | H | T | H |
| 2016A_P5-G10_7 | PHI6 | PHI1 | progeny | 2016A_P5-G10 | H | T | H |
| 2016A_P5-G10_8 | PHI6 | PHI1 | progeny | 2016A_P5-G10 | H | T | H |
| 2016A_P5-G10_9 | PHI6 | PHI1 | progeny | 2016A_P5-G10 | H | T | H |
| 2016A_P5-G12 | PHI6 | PHI1 | mother | 2016A_P5-G12 | H | H | H |
| 2016A_P5-G12_1 | PHI6 | PHI1 | progeny | 2016A_P5-G12 | T | C | A |
| 2016A_P5-G8 | PHI6 | PHI1 | mother | 2016A_P5-G8 | H | T | H |
| 2016A_P5-G8_3 | PHI6 | PHI1 | progeny | 2016A_P5-G8 | T | T | A |
| 2016A_P5-G8_4 | PHI6 | PHI1 | progeny | 2016A_P5-G8 | C | T | G |
| 2016A_P5-G8_5 | PHI6 | PHI1 | progeny | 2016A_P5-G8 | T | T | A |
| 2016A_P5-G9 | PHI6 | PHI1 | mother | 2016A_P5-G9 | H | T | H |
| 2016A_P5-G9_1 | PHI6 | PHI1 | progeny | 2016A_P5-G9 | C | T | G |
| 2016A_P5-G9_5 | PHI6 | PHI1 | progeny | 2016A_P5-G9 | T | T | A |
| 2016A_P5-G9_6 | PHI6 | PHI1 | progeny | 2016A_P5-G9 | H | T | H |
| 2016A_P5-H10 | PHI6 | PHI1 | mother | 2016A_P5-H10 | H | H | H |
| 2016A_P5-H10_2 | PHI6 | PHI1 | progeny | 2016A_P5-H10 | H | H | H |
| 2016A_P5-H10_3 | PHI6 | PHI1 | progeny | 2016A_P5-H10 | H | H | H |
| 2016A_P5-H10_5 | PHI6 | PHI1 | progeny | 2016A_P5-H10 | H | H | H |
| 2016A_P5-H11 | PHI6 | PHI1 | mother | 2016A_P5-H11 | H | T | H |
| 2016A_P5-H11_2 | PHI6 | PHI1 | progeny | 2016A_P5-H11 | H | T | H |
| 2016A_P5-H11_3 | PHI6 | PHI1 | progeny | 2016A_P5-H11 | C | T | G |
| 2016A_P5-H11_5 | PHI6 | PHI1 | progeny | 2016A_P5-H11 | H | T | H |
| 2016A_P5-H11_6 | PHI6 | PHI1 | progeny | 2016A_P5-H11 | H | T | H |
| 2016A_P5-H12_11 | PHI6 | PHI1 | progeny | 2016A_P5-H12 | H | T | H |
| 2016A_P5-H12_14 | PHI6 | PHI1 | progeny | 2016A_P5-H12 | H | T | H |
| 2016A_P5-H12_16 | PHI6 | PHI1 | progeny | 2016A_P5-H12 | H | T | H |
| 2016A_P5-H12_17 | PHI6 | PHI1 | progeny | 2016A_P5-H12 | H | H | H |
| 2016A_P5-H12_3 | PHI6 | PHI1 | progeny | 2016A_P5-H12 | T | T | A |
| 2016A_P5-H12_9 | PHI6 | PHI1 | progeny | 2016A_P5-H12 | H | T | H |
| 2016A_P5-H3 | PHI6 | PHI1 | mother | 2016A_P5-H3 | H | T | H |
| 2016A_P5-H3_1 | PHI6 | PHI1 | progeny | 2016A_P5-H3 | H | T | H |
| 2016A_P5-H3_11 | PHI6 | PHI1 | progeny | 2016A_P5-H3 | T | T | H |
| 2016A_P5-H3_2 | PHI6 | PHI1 | progeny | 2016A_P5-H3 | C | T | G |
| 2016A_P5-H3_3 | PHI6 | PHI1 | progeny | 2016A_P5-H3 | T | T | A |
| 2016A_P5-H3_4 | PHI6 | PHI1 | progeny | 2016A_P5-H3 | C | T | G |
| 2016A_P5-H3_5 | PHI6 | PHI1 | progeny | 2016A_P5-H3 | C | T | G |
| 2016A_P5-H3_6 | PHI6 | PHI1 | progeny | 2016A_P5-H3 | C | T | H |
| 2016A_P5-H3_9 | PHI6 | PHI1 | progeny | 2016A_P5-H3 | H | T | H |
| 2016A_P5-H5 | PHI6 | PHI1 | mother | 2016A_P5-H5 | H | H | H |
| 2016A_P5-H5_1 | PHI6 | PHI1 | progeny | 2016A_P5-H5 | H | H | H |
| 2016A_P5-H5_10 | PHI6 | PHI1 | progeny | 2016A_P5-H5 | H | H | H |
| 2016A_P5-H5_12 | PHI6 | PHI1 | progeny | 2016A_P5-H5 | H | H | H |
| 2016A_P5-H5_3 | PHI6 | PHI1 | progeny | 2016A_P5-H5 | C | T | G |
| 2016A_P5-H5_4 | PHI6 | PHI1 | progeny | 2016A_P5-H5 | H | H | H |
| 2016A_P5-H5_5 | PHI6 | PHI1 | progeny | 2016A_P5-H5 | C | T | G |
| 2016A_P5-H5_6 | PHI6 | PHI1 | progeny | 2016A_P5-H5 | C | T | G |
| 2016A_P5-H5_7 | PHI6 | PHI1 | progeny | 2016A_P5-H5 | H | H | H |
| 2016A_P5-H5_8 | PHI6 | PHI1 | progeny | 2016A_P5-H5 | T | C | A |
| 2016A_P5-H6 | PHI6 | PHI1 | mother | 2016A_P5-H6 | H | T | H |
| 2016A_P5-H6_1 | PHI6 | PHI1 | progeny | 2016A_P5-H6 | H | T | H |
| 2016A_P5-H8 | PHI6 | PHI1 | mother | 2016A_P5-H8 | H | H | H |
| 2016A_P5-H8_1 | PHI6 | PHI1 | progeny | 2016A_P5-H8 | T | H | H |
| 2016A_P5-H8_10 | PHI6 | PHI1 | progeny | 2016A_P5-H8 | C | H | H |
| 2016A_P5-H8_16 | PHI6 | PHI1 | progeny | 2016A_P5-H8 | C | T | G |
| 2016A_P5-H8_4 | PHI6 | PHI1 | progeny | 2016A_P5-H8 | H | H | H |
| 2016A_P5H12 | PHI6 | PHI1 | mother | 2016A_P5H12 | H | T | H |
| 2016A_P6-A8 | PHI6 | PHI1 | mother | 2016A_P6-A8 | H | H | H |
| 2016A_P6-A8_1 | PHI6 | PHI1 | progeny | 2016A_P6-A8 | T | C | H |
| 2016A_P6-A8_11 | PHI6 | PHI1 | progeny | 2016A_P6-A8 | H | H | H |
| 2016A_P6-A8_12 | PHI6 | PHI1 | progeny | 2016A_P6-A8 | C | T | G |
| 2016A_P6-A8_16 | PHI6 | PHI1 | progeny | 2016A_P6-A8 | C | T | H |
| 2016A_P6-A8_8 | PHI6 | PHI1 | progeny | 2016A_P6-A8 | H | H | H |
| 2016A_P6-A9 | PHI6 | PHI1 | mother | 2016A_P6-A9 | H | H | H |
| 2016A_P6-A9_1 | PHI6 | PHI1 | progeny | 2016A_P6-A9 | C | T | G |
| 2016A_P6-A9_2 | PHI6 | PHI1 | progeny | 2016A_P6-A9 | H | H | H |
| 2016A_P6-A9_4 | PHI6 | PHI1 | progeny | 2016A_P6-A9 | T | C | A |
| 2016A_P6-A9_6 | PHI6 | PHI1 | progeny | 2016A_P6-A9 | T | C | A |
| 2016A_P6-A9_8 | PHI6 | PHI1 | progeny | 2016A_P6-A9 | C | T | G |
| 2016A_P6-B10 | PHI6 | PHI1 | mother | 2016A_P6-B10 | H | H | H |
| 2016A_P6-B10_11 | PHI6 | PHI1 | progeny | 2016A_P6-B10 | T | C | A |
| 2016A_P6-B10_12 | PHI6 | PHI1 | progeny | 2016A_P6-B10 | T | C | A |
| 2016A_P6-B10_13 | PHI6 | PHI1 | progeny | 2016A_P6-B10 | C | T | G |
| 2016A_P6-B10_15 | PHI6 | PHI1 | progeny | 2016A_P6-B10 | H | H | H |
| 2016A_P6-B10_4 | PHI6 | PHI1 | progeny | 2016A_P6-B10 | H | H | H |
| 2016A_P6-B10_8 | PHI6 | PHI1 | progeny | 2016A_P6-B10 | C | T | G |
| 2016A_P6-B12 | PHI6 | PHI1 | mother | 2016A_P6-B12 | H | H | H |
| 2016A_P6-B12_4 | PHI6 | PHI1 | progeny | 2016A_P6-B12 | C | T | G |
| 2016A_P6-B12_7 | PHI6 | PHI1 | progeny | 2016A_P6-B12 | T | C | A |
| 2016A_P6-B7 | PHI6 | PHI1 | mother | 2016A_P6-B7 | H | H | H |
| 2016A_P6-B7_1 | PHI6 | PHI1 | progeny | 2016A_P6-B7 | H | C | H |
| 2016A_P6-B7_2 | PHI6 | PHI1 | progeny | 2016A_P6-B7 | H | H | H |
| 2016A_P6-B8 | PHI6 | PHI1 | mother | 2016A_P6-B8 | H | T | H |
| 2016A_P6-B8_2 | PHI6 | PHI1 | progeny | 2016A_P6-B8 | H | T | H |
| 2016A_P6-B8_3 | PHI6 | PHI1 | progeny | 2016A_P6-B8 | H | T | H |
| 2016A_P6-C10 | PHI6 | PHI1 | mother | 2016A_P6-C10 | H | T | H |
| 2016A_P6-C10_1 | PHI6 | PHI1 | progeny | 2016A_P6-C10 | H | T | H |
| 2016A_P6-C10_2 | PHI6 | PHI1 | progeny | 2016A_P6-C10 | H | T | H |
| 2016A_P6-C10_3 | PHI6 | PHI1 | progeny | 2016A_P6-C10 | T | T | A |
| 2016A_P6-C12 | PHI6 | PHI1 | mother | 2016A_P6-C12 | H | T | H |
| 2016A_P6-C12_1 | PHI6 | PHI1 | progeny | 2016A_P6-C12 | H | T | H |
| 2016A_P6-C12_2 | PHI6 | PHI1 | progeny | 2016A_P6-C12 | H | T | H |
| 2016A_P6-C12_3 | PHI6 | PHI1 | progeny | 2016A_P6-C12 | C | T | G |
| 2016A_P6-C12_4 | PHI6 | PHI1 | progeny | 2016A_P6-C12 | H | T | H |
| 2016A_P6-D11 | PHI6 | PHI1 | mother | 2016A_P6-D11 | H | H | H |
| 2016A_P6-D11_1 | PHI6 | PHI1 | progeny | 2016A_P6-D11 | T | C | A |
| 2016A_P6-E5 | PHI8 | PHI1 | mother | 2016A_P6-E5 | H | H | H |
| 2016A_P6-E5_4 | PHI8 | PHI1 | progeny | 2016A_P6-E5 | T | H | H |
| 2016A_P6-F7 | PHI8 | PHI1 | mother | 2016A_P6-F7 | H | H | H |
| 2016A_P6-F7_12 | PHI8 | PHI1 | progeny | 2016A_P6-F7 | H | C | H |
| 2016A_P6-G9 | PHI8 | PHI1 | mother | 2016A_P6-G9 | H | H | H |
| 2016A_P6-G9_4 | PHI8 | PHI1 | progeny | 2016A_P6-G9 | H | H | H |
| 2016A_P6-H1 | PHI8 | PHI1 | mother | 2016A_P6-H1 | H | H | H |
| 2016A_P6-H1_3 | PHI8 | PHI1 | progeny | 2016A_P6-H1 | H | H | H |
| 2016A_P6-H3_3 | PHI8 | PHI1 | progeny | 2016A_P6-H3 | H | H | H |
| 2016A_P7-A8 | PHI8 | PHI1 | mother | 2016A_P7-A8 | H | H | H |

### Table S5. Results of the SNP array analasis (continue)

| | | | | | Chr | 3 | 3 | 3 | 3 | 3 | 3 | 3 | 3 | 3 | 3 | 3 | 3 | 3 | 3 | 3 | 3 | 3 | 4 | 4 | 4 | 4 | 4 | 4 | 4 | 4 | 4 | 4 | 4 | 4 | 4 | 4 | 4 | 4 | 4 | 4 | 4 | 4 | 4 | 4 | 4 | 4 | 4 | 4 | 5 | 5 | 5 | 5 | 5 | 5 | 5 | 5 | 5 | 5 | 5 | 5 | 5 | 5 | 5 | 5 | 5 | 5 | 5 | 5 | 5 | 5 | 5 | 5 | 5 | 5 | 5 | 5 | 5 | 5 | 6 | 6 | 6 | 6 | 6 | 6 | 6 | 6 | 6 | 6 | 6 | 6 | 6 | 6 | 6 | 6 | 6 | 6 | 6 | 6 | 6 | 6 | 6 | 7 | 7 | 7 | 7 | 7 | 7 | 7 |
|---|---|---|---|---|---|---|---|---|---|---|---|---|---|---|---|---|---|---|---|---|---|---|---|---|---|---|---|---|---|---|---|---|---|---|---|---|---|---|---|---|---|---|---|---|---|---|---|---|---|---|---|---|---|---|---|---|---|---|---|---|---|---|---|---|---|---|---|---|---|---|---|---|---|---|---|---|---|---|---|---|---|---|---|---|---|---|---|---|---|---|---|---|---|---|---|---|---|---|---|---|---|---|---|---|---|---|---|---|
| | | | | | Position, cM | 94 | 95 | 95 | 104 | 113 | 123 | 135 | 149 | 160 | 168 | 182 | 195 | 203 | 224 | 227 | 240 | 243 | 7.4 | 18 | 30 | 33 | 42 | 50 | 75 | 85 | 89 | 104 | 108 | 110 | 111 | 110 | 119 | 128 | 140 | 155 | 163 | 163 | 176 | 188 | 197 | 216 | 238 | 246 | 5.3 | 12 | 21 | 32 | 48 | 51 | 65 | 71 | 89 | 93 | 97 | 98 | 101 | 102 | 102 | 102 | 105 | 111 | 124 | 138 | 148 | 159 | 166 | 177 | 185 | 191 | 198 | 208 | 215 | 224 | 0.3 | 14 | 16 | 22 | 23 | 24 | 23 | 24 | 38 | 52 | 54 | 72 | 79 | 93 | 98 | 114 | 129 | 130 | 135 | 143 | 157 | 180 | 181 | 1.3 | 14 | 26 | 33 | 50 | 62 | 68 |
| plant_id | mom_gen | dad_gen | mother_progeny | family_ | | | | | | | | | | | | | | | | | | | | | | | | | | | | | | | | | | | | | | | | | | | | | | | | | | | | | | | | | | | | | | | | | | | | | | | | | | | | | | | | | | | | | | | | | | | | | | | | | | | | | | | | |
| 2016A_P7-A8_1 | PHI8 | PHI1 | progeny | 2016A_P7-A8 | | T | C | A | C | G | T | C | C | H | H | H | H | H | H | H | C | H | A | C | G | H | T | A | T | C | T | C | C | C | H | A | H | H | H | H | H | H | H | H | G | A | C | T | H | C | H | H | H | H | A | H | H | H | H | G | G | A | T | T | T | H | H | G | H | H | G | H | H | H | H | T | H | G | H | H | H | H | H | H | H | H | H | A | G | A | C | H | G | C | H | H | H | H | H | H | C | H | C | G | H | G | G | G |
| 2016A_P7-C11 | PHI5 | PHI1 | mother | 2016A_P7-C11 | | H | T | H | H | H | H | C | H | A | H | H | H | H | H | H | A | H | H | H | G | C | H | A | H | H | H | H | H | H | G | A | H | H | H | H | H | H | H | H | A | H | H | H | H | C | H | H | H | H | H | H | H | H | H | H | H | C | H | H | H | H | H | H | A | H | H | H | H | H | H | H | H | H | H | T | H | G | A | T | A | G | C | H | H | G | H | H | G | H | T | T | H | H | H | H | C | H | H | H | G | H | H | H |
| 2016A_P7-C11_1 | PHI5 | PHI1 | progeny | 2016A_P7-C11 | | C | T | G | A | A | T | C | T | A | T | A | T | G | C | C | A | G | H | H | G | C | H | H | H | H | T | C | C | T | T | A | C | A | T | A | C | G | G | G | A | A | C | T | H | C | H | H | H | H | H | H | T | T | A | C | A | C | C | T | C | T | A | T | A | T | A | C | A | T | A | G | G | A | H | T | H | G | A | T | A | G | H | H | H | G | H | A | G | T | T | T | H | H | H | H | C | H | T | H | G | H | G | G |
| 2016A_P7-D10 | PHI5 | PHI1 | mother | 2016A_P7-D10 | | H | T | H | H | H | H | H | H | H | H | H | H | H | H | H | H | C | H | H | G | C | H | H | H | H | H | H | H | H | H | A | H | H | H | H | H | H | H | H | G | A | C | T | H | C | H | H | H | H | H | H | H | H | H | H | H | C | H | H | H | H | A | H | A | C | H | H | C | H | H | H | H | H | H | T | H | G | H | H | H | H | C | H | H | H | C | H | G | C | H | H | H | H | H | H | C | H | C | G | H | G | H | H |
| 2016A_P7-D10_1 | PHI5 | PHI1 | progeny | 2016A_P7-D10 | | T | C | A | C | G | C | C | C | A | C | G | G | C | T | C | A | C | A | G | G | C | C | A | T | C | T | C | C | T | T | A | C | G | C | G | T | T | C | T | G | G | G | T | G | C | A | A | C | A | G | T | C | G | G | G | G | C | T | C | T | C | G | G | A | C | G | T | C | C | G | T | G | A | C | T | A | G | A | T | A | G | C | A | A | G | C | G | G | C | T | T | C | C | A | A | C | C | C | G | G | G | G | G |
| 2016A_P7-D2 | PHI5 | PHI1 | mother | 2016A_P7-D2 | | T | H | H | H | H | T | C | H | H | T | H | T | H | H | C | A | G | A | G | C | H | H | H | H | H | H | H | H | T | H | A | H | H | H | H | H | H | H | H | A | H | H | H | H | C | H | A | H | A | H | H | H | H | G | G | G | H | T | H | T | T | H | H | A | H | H | H | C | H | H | T | H | H | H | H | H | G | A | T | A | G | C | A | H | A | C | H | G | C | C | G | C | H | H | H | C | H | H | H | C | H | H | H |
| 2016A_P7-D2_1 | PHI5 | PHI1 | progeny | 2016A_P7-D2 | | T | C | A | C | G | T | C | C | A | C | G | G | C | C | C | A | G | A | G | G | C | C | A | T | C | T | C | C | T | T | H | C | A | T | A | T | T | C | T | G | G | G | C | A | C | G | G | T | G | A | C | T | H | G | G | G | A | T | T | T | T | A | T | A | T | A | C | C | T | A | G | G | A | C | C | A | H | A | T | A | G | C | A | G | G | C | G | G | C | C | G | C | H | C | C | C | C | C | G | G | G | G | G |
| 2016A_P7-D8 | PHI5 | PHI1 | mother | 2016A_P7-D8 | | H | H | H | H | H | T | H | H | A | H | H | H | H | H | C | A | H | A | G | H | H | H | H | H | H | H | H | H | H | H | A | H | H | H | H | H | T | H | H | H | H | H | H | H | C | H | H | H | H | H | H | H | G | G | G | G | H | T | H | T | H | H | H | A | H | G | H | C | H | H | H | H | H | H | T | H | G | A | T | A | G | C | A | H | H | C | H | G | H | H | H | C | H | H | H | C | H | H | H | H | H | H | H |
| 2016A_P7-D8_2 | PHI5 | PHI1 | progeny | 2016A_P7-D8 | | T | C | A | C | G | T | C | C | A | C | H | G | H | H | C | A | H | A | G | H | G | T | H | C | H | G | T | T | H | T | A | H | H | H | H | H | T | H | H | H | H | H | H | H | C | H | H | H | H | H | H | C | G | G | G | G | C | T | C | T | C | G | H | A | H | G | H | C | T | A | G | G | A | C | T | A | G | A | T | A | G | C | A | G | A | C | A | G | T | T | H | C | H | H | H | C | T | T | T | C | A | G | G |
| 2016A_P7-G3 | PHI5 | PHI1 | mother | 2016A_P7-G3 | | H | H | H | H | H | H | C | H | A | H | H | H | H | H | C | A | H | H | H | G | C | H | H | H | H | H | H | H | H | H | A | H | H | H | H | H | H | H | H | H | H | H | T | H | C | H | H | H | H | H | H | H | G | G | G | G | H | T | H | T | H | H | G | A | C | H | H | C | H | H | H | H | H | H | T | H | G | A | T | A | G | C | A | H | H | C | H | G | H | H | H | C | H | H | H | C | H | H | H | G | H | H | H |
| 2016A_P7-G3_1 | PHI5 | PHI1 | progeny | 2016A_P7-G3 | | T | H | H | H | H | H | C | H | A | H | A | H | H | H | C | A | H | H | H | G | C | H | T | H | H | H | H | H | H | H | A | H | H | H | H | H | H | H | H | H | H | H | T | H | C | H | H | H | H | H | H | H | H | G | G | G | H | T | H | T | H | H | G | A | C | A | H | C | H | H | H | H | H | H | T | H | G | A | T | A | G | C | A | H | H | C | H | G | H | T | H | C | H | H | H | C | H | H | T | G | H | H | H |
| 2016A_P7-G3_3 | PHI5 | PHI1 | progeny | 2016A_P7-G3 | | H | H | H | H | H | H | C | C | A | C | G | H | H | H | C | A | H | H | H | G | C | H | H | H | C | T | C | C | T | T | A | C | A | T | A | C | G | H | H | H | H | G | T | H | C | H | H | H | H | G | T | C | G | G | G | G | C | T | C | T | C | G | G | A | C | G | T | C | H | G | T | C | G | H | T | H | G | A | T | A | G | C | A | A | G | C | H | G | H | H | H | C | H | H | H | C | H | H | H | G | H | H | H |
| 2016A_P7-G3_4 | PHI5 | PHI1 | progeny | 2016A_P7-G3 | | T | C | A | H | G | C | C | H | A | H | H | H | H | T | C | A | H | H | H | G | C | C | A | T | C | T | C | C | T | T | A | C | H | H | H | H | H | H | H | H | H | H | T | H | C | H | H | H | H | H | H | H | G | G | G | G | C | T | C | T | C | H | G | A | C | H | H | C | H | H | H | H | G | C | T | A | G | A | T | A | G | C | A | G | A | C | H | G | H | H | H | C | C | C | A | C | C | C | G | G | G | G | G |
| 2016A_P7-G7 | PHI5 | PHI1 | mother | 2016A_P7-G7 | | H | H | H | H | H | H | H | H | A | H | H | H | H | H | C | A | H | A | G | H | H | H | A | T | H | T | H | H | H | H | A | H | H | H | H | H | H | H | H | H | H | H | T | H | C | H | H | H | H | H | H | H | G | G | G | G | H | T | H | T | H | H | H | A | C | A | H | C | H | H | H | H | H | H | H | H | G | A | T | A | G | C | A | H | H | C | H | G | C | T | H | H | H | A | H | C | H | H | H | H | H | H | H |
| 2016A_P7-G7_1 | PHI5 | PHI1 | progeny | 2016A_P7-G7 | | H | H | H | H | H | H | H | H | A | H | H | H | H | H | C | A | G | A | G | H | H | H | A | T | H | T | H | H | H | H | A | H | H | H | H | H | H | H | H | H | H | H | T | H | C | H | H | H | H | H | H | H | G | G | G | G | H | T | H | T | H | H | H | A | C | G | H | C | H | H | H | H | H | H | H | H | G | A | T | A | G | C | A | H | A | C | H | G | C | H | H | H | H | A | C | C | H | H | H | H | H | H | H |
| 2016A_P7-H9 | PHI5 | PHI1 | mother | 2016A_P7-H9 | | H | H | H | H | H | H | C | H | A | H | H | H | H | H | H | A | H | A | G | H | G | H | T | H | H | H | H | H | H | H | A | H | H | H | H | H | H | H | H | H | H | H | H | H | C | H | H | H | H | H | H | H | G | G | G | G | H | T | H | T | H | H | H | A | H | A | H | C | H | H | H | H | G | H | T | H | G | A | T | A | G | C | A | H | H | C | H | G | C | C | G | C | H | A | H | C | H | H | H | H | H | H | H |
| 2016A_P7-H9_1 | PHI5 | PHI1 | progeny | 2016A_P7-H9 | | H | H | H | H | H | H | C | H | H | H | H | H | H | H | H | A | H | H | H | G | C | H | A | T | H | H | H | H | H | H | A | H | H | H | H | H | H | H | H | H | H | H | H | A | C | G | G | T | G | A | C | T | T | A | C | A | C | C | T | H | H | H | H | A | H | H | H | H | H | H | H | H | H | H | T | H | G | A | T | A | G | C | H | H | G | H | H | H | H | H | H | H | H | H | H | C | H | H | H | G | H | H | H |
| 2016A_P7-H9_2 | PHI5 | PHI1 | progeny | 2016A_P7-H9 | | H | H | H | H | H | H | C | H | H | H | H | H | H | H | H | A | H | H | H | G | C | H | A | T | H | H | H | H | H | H | A | H | H | H | H | H | H | H | H | H | H | C | H | C | H | H | H | H | H | H | H | H | H | H | H | C | H | H | H | H | H | H | A | H | H | H | H | H | H | H | H | H | H | T | H | G | A | T | A | G | C | H | H | G | H | H | H | H | H | H | H | H | H | H | H | H | H | H | G | H | H | H |
| 2016A_P8-A1 | PHI5 | PHI1 | mother | 2016A_P8-A1 | | H | H | H | H | H | H | C | H | H | H | H | H | H | H | H | A | H | H | H | G | C | H | A | T | H | H | H | H | H | H | A | H | H | T | H | C | H | H | H | H | H | H | H | H | C | H | H | H | H | H | H | H | H | H | H | H | C | H | H | H | H | A | H | A | H | H | H | H | H | H | H | H | H | H | H | H | H | H | H | H | H | C | H | H | G | H | H | G | H | H | H | H | H | H | H | C | H | C | H | G | H | H | H |
| 2016A_P8-A1_1 | PHI5 | PHI1 | progeny | 2016A_P8-A1 | | H | H | H | H | H | H | C | H | H | H | H | H | G | C | C | A | G | H | H | G | C | H | A | T | H | H | H | H | H | H | A | H | H | T | H | C | H | H | T | G | G | G | C | H | C | H | H | H | H | A | C | H | H | H | H | H | C | H | H | H | H | A | H | A | H | H | H | A | T | H | H | H | H | H | H | H | H | H | H | H | H | C | H | H | G | H | H | G | H | H | H | H | H | H | H | C | T | T | T | G | A | T | A |
| 2016A_P8-A5 | PHI5 | PHI1 | mother | 2016A_P8-A5 | | H | H | H | H | H | H | H | H | A | H | H | H | H | H | C | A | H | A | G | H | H | H | H | H | H | H | H | H | H | H | A | H | H | H | H | H | T | H | H | H | H | H | T | H | C | H | H | H | H | H | H | H | G | G | G | G | H | T | H | T | H | H | H | A | H | H | H | C | H | H | H | H | H | H | T | H | G | A | T | A | G | C | A | H | H | C | H | G | H | H | H | H | H | H | H | C | H | H | H | H | H | H | H |
| 2016A_P8-A5_1 | PHI5 | PHI1 | progeny | 2016A_P8-A5 | | H | H | H | H | H | H | H | H | A | H | H | H | H | H | C | A | H | H | G | H | H | H | H | H | H | H | H | H | H | H | A | H | H | H | H | H | H | H | H | H | H | H | T | H | C | H | H | H | H | H | H | H | G | H | H | G | H | T | H | H | H | H | H | H | H | H | H | C | H | H | H | H | H | H | T | H | G | A | T | H | H | H | H | H | H | C | H | G | H | H | H | H | H | H | H | H | H | H | H | H | H | H | H |
| 2016A_P8-A6 | PHI5 | PHI1 | mother | 2016A_P8-A6 | | H | H | H | H | H | T | C | H | H | H | H | H | H | H | H | A | H | A | G | H | H | H | H | H | H | T | H | H | H | H | A | H | H | H | H | H | H | H | H | H | H | H | H | H | C | H | H | H | H | H | H | H | G | G | G | G | H | T | H | T | H | H | H | A | H | H | H | C | H | H | H | H | H | H | H | H | G | A | T | A | G | C | A | H | H | C | H | G | C | T | T | H | H | H | H | C | H | H | H | G | H | H | H |
| 2016A_P8-A6_1 | PHI5 | PHI1 | progeny | 2016A_P8-A6 | | C | T | G | A | A | T | C | H | H | H | H | H | H | T | A | A | C | A | G | H | H | H | H | H | A | T | T | T | C | G | A | A | G | C | H | H | H | H | H | H | H | H | H | H | C | H | H | H | H | H | H | T | G | G | G | G | A | T | T | T | T | A | T | A | T | A | H | C | H | H | H | H | H | A | T | C | G | A | T | A | G | C | A | H | H | C | H | G | C | T | T | H | H | H | A | H | H | H | H | G | H | T | A |
| 2016A_P8-D5 | PHI9 | PHI1 | mother | 2016A_P8-D5 | | H | H | G | H | H | H | C | H | A | H | H | H | H | H | H | H | H | H | H | H | H | T | H | C | H | H | H | H | C | H | A | H | H | H | H | H | H | C | H | H | H | H | C | A | H | H | H | H | H | H | H | T | H | H | H | H | H | C | H | H | T | A | H | A | H | H | H | H | H | A | H | H | A | C | T | H | G | H | H | G | T | H | H | H | H | G | H | H | H | H | H | H | T | H | C | H | T | H | H | G | A | T | H |
| 2016A_P8-D5_3 | PHI9 | PHI1 | progeny | 2016A_P8-D5 | | H | H | G | H | H | H | C | H | A | H | H | H | H | H | H | H | H | H | H | H | H | T | H | C | H | H | H | H | C | H | A | H | H | H | H | H | H | C | H | H | H | H | C | A | H | H | H | H | H | H | H | T | H | H | H | H | H | C | H | H | T | A | H | A | H | H | H | H | H | A | H | H | A | C | T | H | G | H | H | G | T | H | H | H | H | G | H | H | H | H | H | H | T | H | C | H | T | H | H | G | A | T | H |
| 2016A_P8-D5_5 | PHI9 | PHI1 | progeny | 2016A_P8-D5 | | C | T | G | A | A | T | C | H | A | H | H | G | H | H | H | H | H | H | H | H | H | T | T | C | A | G | T | T | C | G | A | A | G | C | H | H | H | C | H | H | H | G | C | A | H | H | H | H | H | H | H | T | G | G | G | G | A | C | C | T | T | A | G | A | C | H | H | H | H | A | H | H | A | C | T | H | G | H | H | G | T | H | H | H | H | G | H | H | H | H | H | H | T | C | C | T | T | T | T | G | A | T | A |
| 2016A_P8-E1 | PHI9 | PHI1 | mother | 2016A_P8-E1 | | H | T | H | H | H | H | H | H | H | H | H | H | H | H | H | H | H | H | H | H | H | T | H | C | C | H | C | H | C | T | A | C | H | T | H | H | T | H | H | H | H | H | C | A | H | H | H | H | H | H | H | H | G | H | H | H | H | C | H | H | T | A | G | A | C | H | H | H | H | A | H | H | A | C | C | H | H | H | C | H | H | H | A | H | A | H | H | H | H | H | H | H | H | A | C | H | T | H | H | H | H | T | H |
| 2016A_P8-E1_2 | PHI9 | PHI1 | progeny | 2016A_P8-E1 | | H | H | G | H | H | H | C | H | A | H | H | H | H | H | H | H | H | H | H | H | H | T | H | C | H | H | H | H | C | H | A | H | G | H | H | C | H | H | H | H | H | H | C | A | H | H | H | H | H | H | H | T | H | H | H | H | H | C | H | H | T | A | H | A | H | H | H | H | H | A | H | H | A | C | C | H | H | H | C | H | H | H | H | H | H | G | H | H | H | H | H | H | T | H | C | H | T | H | H | H | H | T | H |
| 2016A_P9-A10 | PHI9 | PHI1 | mother | 2016A_P9-A10 | | H | H | G | H | H | H | H | H | A | H | H | H | H | H | H | H | H | H | H | H | H | T | H | H | C | H | H | H | C | H | A | H | G | H | H | C | H | H | H | H | H | H | C | A | H | H | H | H | H | H | H | T | H | H | H | H | H | C | H | H | T | A | H | A | H | H | H | H | H | A | H | H | A | C | H | H | H | H | H | G | T | H | H | H | H | G | H | H | H | H | H | H | T | H | C | H | H | H | H | H | H | T | H |
| 2016A_P9-A10_1 | PHI9 | PHI1 | progeny | 2016A_P9-A10 | | T | C | G | C | G | C | C | C | A | C | G | G | C | C | C | A | G | A | G | C | G | T | T | T | C | G | T | T | C | G | A | A | G | T | A | C | G | C | G | A | A | C | C | A | T | A | A | C | A | G | C | T | T | A | C | A | C | C | T | C | T | A | T | A | T | A | C | A | T | A | G | G | A | C | C | A | A | G | C | G | T | C | G | G | G | G | A | A | T | C | G | C | T | C | H | T | C | C | T | G | A | T | A |
| 2016A_P9-A10_3 | PHI9 | PHI1 | progeny | 2016A_P9-A10 | | C | T | G | A | A | T | A | T | A | T | A | T | G | T | A | C | C | G | C | G | C | T | A | C | C | T | C | C | C | T | A | C | G | T | A | C | G | C | G | A | A | C | C | A | T | A | A | C | A | G | T | T | G | A | C | A | C | C | T | C | T | A | T | A | T | G | T | C | C | A | T | C | A | C | T | C | G | A | T | G | T | C | G | G | G | G | A | A | T | T | T | T | T | A | C | T | T | T | T | G | A | T | A |
| 2016A_P9-A7 | PHI9 | PHI1 | mother | 2016A_P9-A7 | | H | H | G | H | H | H | H | H | H | H | H | H | H | H | H | H | H | H | H | H | H | T | T | C | H | H | H | H | C | H | A | H | G | H | H | C | H | H | H | H | H | H | C | A | H | H | H | H | H | H | H | T | H | H | H | H | H | C | H | C | T | A | H | A | H | H | H | H | H | A | H | H | A | C | C | H | H | H | C | H | H | H | H | H | H | G | H | H | H | H | H | H | H | A | C | H | T | H | H | H | H | T | H |
| 2016A_P9-A7_1 | PHI9 | PHI1 | progeny | 2016A_P9-A7 | | H | H | G | H | H | H | H | H | H | H | H | H | H | H | H | H | H | H | H | C | H | T | T | C | H | H | H | H | C | G | A | H | G | C | H | C | H | H | H | H | H | H | C | A | H | H | H | H | H | H | H | T | H | H | H | H | H | C | H | C | T | A | H | A | H | A | C | H | H | A | H | H | A | C | C | H | H | H | C | H | H | H | H | H | H | G | H | H | H | H | H | H | H | A | C | H | T | H | H | H | H | T | A |
| 2016A_P9-A7_2 | PHI9 | PHI1 | progeny | 2016A_P9-A7 | | H | H | G | H | H | H | H | H | H | H | H | H | H | H | H | H | H | H | H | H | H | T | H | C | H | H | H | H | C | H | A | H | G | H | H | C | H | H | H | H | H | H | C | A | H | H | H | H | H | H | H | T | H | H | H | H | H | C | H | C | T | A | H | A | H | H | H | H | H | A | H | H | A | C | C | H | H | H | C | H | H | H | H | H | H | G | H | H | H | H | H | H | H | A | C | H | T | H | H | H | H | T | H |
| 2016A_P9-B1 | PHI9 | PHI1 | mother | 2016A_P9-B1 | | H | T | H | H | H | H | H | H | H | H | H | H | H | H | H | H | H | H | H | H | H | T | H | C | C | H | C | H | C | T | A | C | G | H | H | H | T | H | H | H | H | H | C | A | H | H | H | H | H | H | H | T | H | H | H | H | H | C | H | H | T | A | G | A | C | H | H | H | H | A | H | H | A | C | C | H | H | H | C | H | H | H | A | H | A | H | H | H | H | H | H | H | T | H | C | H | T | H | H | H | H | T | H |
| 2016A_P9-B1_2 | PHI9 | PHI1 | progeny | 2016A_P9-B1 | | C | T | G | A | A | T | A | T | T | T | A | T | G | C | C | A | G | G | C | G | C | T | A | C | C | G | C | T | C | T | A | C | G | C | A | C | T | G | G | A | A | C | C | A | C | G | G | T | G | G | T | T | G | G | G | G | A | C | C | T | T | A | G | A | C | G | T | A | T | A | G | G | A | C | C | A | A | G | C | G | T | C | A | G | A | C | A | A | T | T | T | T | T | A | C | C | T | T | T | G | A | T | G |
| 2016A_P9-B7 | PHI9 | PHI1 | mother | 2016A_P9-B7 | | H | H | G | H | H | H | H | H | H | H | H | H | H | H | H | H | H | H | H | H | H | T | H | H | C | H | C | H | C | T | A | C | A | T | H | H | H | C | H | H | H | H | C | A | H | H | H | H | H | H | H | T | G | H | H | H | H | C | H | H | T | A | H | A | H | H | H | H | H | A | H | H | A | C | H | H | H | H | H | G | H | H | H | H | H | G | H | H | H | H | H | H | T | H | C | H | T | H | H | H | A | T | H |
| 2016A_P9-B7_10 | PHI9 | PHI1 | progeny | 2016A_P9-B7 | | C | T | G | A | A | T | A | T | T | T | A | T | C | T | A | C | C | G | C | C | G | T | T | T | C | G | C | T | C | T | A | C | A | T | G | T | T | C | T | G | G | G | C | A | T | A | A | T | G | A | C | T | G | A | C | A | C | C | T | C | T | A | G | A | C | G | T | C | C | A | T | C | A | C | C | A | A | G | C | G | T | C | A | A | A | G | G | G | T | T | T | T | T | A | C | C | T | H | G | C | A | T | G |
| 2016A_P9-B7_5 | PHI9 | PHI1 | progeny | 2016A_P9-B7 | | C | T | G | A | A | T | A | T | T | C | G | G | C | T | A | C | C | G | C | G | C | T | T | T | C | G | C | T | C | T | A | C | A | T | G | C | G | C | G | A | A | C | C | A | C | G | G | T | G | A | C | T | G | A | C | A | C | C | T | T | T | A | G | A | C | G | T | C | C | A | T | C | A | C | C | A | A | G | C | G | T | C | A | A | A | G | G | G | C | C | G | C | T | A | C | C | T | T | T | G | A | T | A |
| 2016A_P9-B7_9 | PHI9 | PHI1 | progeny | 2016A_P9-B7 | | T | C | G | C | G | C | C | C | A | C | G | G | C | T | A | C | C | A | G | C | G | T | T | T | C | G | C | T | C | T | A | C | A | T | G | T | T | C | T | G | G | G | C | A | C | G | G | C | A | G | T | T | G | G | G | G | A | C | C | T | T | A | G | A | C | G | T | C | C | A | G | G | A | C | T | C | G | A | T | G | G | T | A | A | G | G | A | A | T | C | G | C | T | C | C | T | T | T | T | G | A | T | G |

plant_id – unique identifier of the plant
mom_gen – genetic backgroiund of mother in the hybred
dad_gen – genetic background of father in the hybred
family_ – identify mother and all the progeny plants from the same mother
mother_progeny – identify each plant as mother of the family or progeny
Chr – number of the chromosome, where marker is located
Distance – position of the marker on the corresponding chromosome in cM
H – heterozygous marker
A, T, G and C – marker homozygous for the corresponding nucleotide
EQV, LS, -9 , etc. – missing values

**Table S5. Results of the SNP array analasis (continue)**

| | | | | Chr | 7 | 7 | 7 | 7 | 7 | 7 | 7 | 7 | 7 | 7 | 7 | 7 | 7 | 7 | 7 | 7 | 7 | 7 | 7 | 8 | 8 | 8 | 8 | 8 | 8 | 8 | 8 | 8 | 8 | 8 | 8 | 8 | 8 | 8 | 8 | 8 | 8 | 8 | 8 | 8 | 8 | 8 | 8 | 9 | 9 | 9 | 9 | 9 | 9 | 9 | 9 | 9 | 9 | 9 | 9 | 9 | 9 | 9 | 9 | 9 | 9 | 9 | 9 | 10 | 10 | 10 | 10 | 10 | 10 | 10 | 10 | 10 | 10 | 10 | 10 | 10 | 10 | 10 | 10 | 10 | 10 | 10 |
|---|---|---|---|---|---|---|---|---|---|---|---|---|---|---|---|---|---|---|---|---|---|---|---|---|---|---|---|---|---|---|---|---|---|---|---|---|---|---|---|---|---|---|---|---|---|---|---|---|---|---|---|---|---|---|---|---|---|---|---|---|---|---|---|---|---|---|---|---|---|---|---|---|---|---|---|---|---|---|---|---|---|---|---|---|---|---|---|
| | | | | Position, cM | 76 | 78 | 77 | 80 | 83 | 84 | 84 | 98 | 114 | 121 | 129 | 142 | 151 | 170 | 177 | 186 | 196 | 204 | 206 | 3 | 15 | 31 | 46 | 48 | 54 | 68 | 85 | 84 | 85 | 87 | 95 | 105 | 114 | 122 | 130 | 137 | 149 | 158 | 168 | 174 | 182 | 197 | 210 | 12 | 26 | 38 | 46 | 56 | 60 | 72 | 76 | 76 | 77 | 89 | 101 | 109 | 124 | 134 | 141 | 152 | 163 | 174 | 181 | 4.1 | 10 | 18 | 30 | 44 | 58 | 62 | 61 | 61 | 62 | 71 | 81 | 93 | 98 | 114 | 128 | 140 | 154 | 162 |
| plant_id | mom_gen | dad_gen | mother_progeny | family_ | | | | | | | | | | | | | | | | | | | | | | | | | | | | | | | | | | | | | | | | | | | | | | | | | | | | | | | | | | | | | | | | | | | | | | | | | | | | | | | | | | | |
| 2016A_P1-A3 | PHI1 | PHI3 | mother | 2016A_P1-A3 | | H | H | H | H | H | H | H | H | H | H | H | H | H | H | C | C | H | T | C | H | H | H | H | H | H | G | G | G | C | A | G | H | H | H | C | H | H | H | H | H | H | H | H | H | T | G | H | H | H | H | H | H | H | H | H | H | H | H | H | H | G | H | C | H | H | H | H | H | H | H | H | H | H | H | H | H | H | A | H | H | H | H |
| 2016A_P1-A3_1 | PHI1 | PHI3 | progeny | 2016A_P1-A3 | | H | H | H | H | H | H | H | H | H | H | H | H | H | H | C | C | H | T | C | H | H | H | H | H | H | G | G | G | C | A | G | H | H | H | C | H | H | H | H | H | H | H | H | H | T | G | H | H | H | H | H | H | H | H | H | H | H | H | H | H | G | H | C | H | H | H | H | H | H | H | H | H | H | H | H | H | H | A | H | H | H | H |
| 2016A_P1-D5 | PHI1 | PHI3 | mother | 2016A_P1-D5 | | H | H | H | H | H | H | H | H | H | H | H | H | H | H | C | C | H | T | C | H | H | H | H | H | H | G | G | G | C | A | G | H | H | H | C | C | H | H | H | G | H | H | H | H | T | G | H | H | H | H | H | H | H | H | H | H | H | H | H | H | G | H | C | H | H | H | H | H | H | H | H | H | H | H | H | H | T | A | H | H | H | A |
| 2016A_P1-D5_1 | PHI1 | PHI3 | progeny | 2016A_P1-D5 | | H | H | H | H | H | H | H | H | H | H | H | H | H | H | C | C | H | T | C | H | H | H | H | H | H | G | G | G | C | A | G | H | H | H | C | C | H | H | H | G | H | H | H | H | T | G | H | H | H | H | H | H | H | H | H | H | H | H | H | H | G | H | C | H | H | H | H | H | H | H | H | H | H | H | H | H | T | A | H | H | H | A |
| 2016A_P1-D8 | PHI1 | PHI3 | mother | 2016A_P1-D8 | | H | H | H | H | H | H | H | H | H | H | H | H | H | H | C | C | H | T | C | H | H | H | H | H | H | H | H | H | H | H | G | H | H | H | C | C | H | H | H | H | H | H | H | H | T | G | H | H | H | H | H | H | H | H | H | H | H | H | H | H | G | H | H | H | H | H | H | H | H | H | H | H | C | H | H | H | T | A | H | G | H | H |
| 2016A_P1-D8_3 | PHI1 | PHI3 | progeny | 2016A_P1-D8 | | T | G | C | H | A | C | T | H | H | H | H | H | T | T | C | C | H | T | C | T | G | C | G | T | T | A | A | A | A | G | A | T | T | C | C | C | C | T | A | G | C | A | T | A | T | G | C | A | T | T | G | T | A | T | H | H | H | H | H | G | G | A | G | H | H | H | H | H | T | A | G | A | C | A | C | C | T | A | C | G | A | A |
| 2016A_P1-E3 | PHI1 | PHI3 | mother | 2016A_P1-E3 | | H | H | H | H | H | H | H | H | H | H | H | H | H | H | C | C | H | T | C | H | H | H | H | H | H | G | G | G | C | A | G | H | H | H | H | H | H | H | H | H | H | H | H | H | H | H | H | H | H | H | H | H | H | H | H | H | H | H | H | H | G | H | C | H | H | H | H | H | H | H | H | H | C | H | H | H | T | A | H | G | C | A |
| 2016A_P1-E3_3 | PHI1 | PHI3 | progeny | 2016A_P1-E3 | | T | G | C | H | A | C | T | C | H | H | H | H | H | H | C | C | H | T | C | H | H | H | H | H | H | G | G | G | C | A | G | H | H | H | H | H | H | H | H | H | H | H | H | H | H | H | H | A | T | T | G | T | A | H | H | C | C | H | H | H | G | H | C | H | H | C | T | C | C | G | A | G | C | G | T | A | T | A | G | G | C | A |
| 2016A_P1-E3_5 | PHI1 | PHI3 | progeny | 2016A_P1-E3 | | G | A | T | G | G | G | C | T | T | A | H | C | G | G | C | C | T | T | C | H | G | C | G | T | T | G | G | G | C | A | G | T | T | C | G | T | C | H | A | G | C | A | T | H | H | H | H | H | H | H | H | H | H | H | T | H | H | H | H | H | G | H | C | H | H | H | A | H | C | G | A | G | C | G | T | A | T | A | G | G | C | A |
| 2016A_P1-E9 | PHI1 | PHI3 | mother | 2016A_P1-E9 | | H | H | H | H | H | H | H | H | H | H | H | H | H | H | C | C | H | T | C | H | H | H | H | H | H | G | G | G | C | A | G | H | H | H | C | H | H | H | H | G | H | H | H | H | T | G | H | H | H | H | H | H | H | H | H | H | H | H | H | H | G | H | C | H | H | H | H | H | H | H | H | H | C | H | H | H | T | A | H | H | H | A |
| 2016A_P1-E9_1 | PHI1 | PHI3 | progeny | 2016A_P1-E9 | | T | G | C | H | A | C | T | H | H | H | H | H | H | H | C | C | H | T | C | H | G | C | G | T | T | G | G | G | C | A | G | T | H | C | C | H | H | H | H | G | H | H | H | H | T | G | C | A | T | T | G | T | A | H | H | H | H | H | H | G | G | A | C | H | H | H | H | H | T | A | G | A | C | A | C | C | T | A | C | H | H | A |
| 2016A_P1-F7 | PHI1 | PHI3 | mother | 2016A_P1-F7 | | H | H | H | H | H | H | H | H | H | H | H | H | H | H | C | C | H | H | H | H | H | H | H | H | H | H | H | H | H | H | G | H | C | H | C | C | H | H | H | G | H | H | H | H | T | G | H | H | H | H | H | H | H | H | H | H | H | H | H | H | G | H | H | H | H | H | H | H | H | H | H | H | H | H | H | H | H | A | H | H | H | A |
| 2016A_P1-F7_4 | PHI1 | PHI3 | progeny | 2016A_P1-F7 | | T | G | C | H | A | C | T | C | C | G | T | T | T | T | C | C | C | C | H | H | H | H | H | H | H | H | H | H | H | H | G | H | C | H | C | C | H | H | H | G | H | H | H | A | T | G | H | H | T | H | H | H | H | H | H | H | H | H | H | H | G | H | H | H | H | H | H | H | H | H | H | H | H | H | H | H | H | A | H | H | H | A |
| 2016A_P1-F8 | PHI2 | PHI1 | mother | 2016A_P1-F8 | | H | H | C | H | A | C | T | H | H | H | H | T | H | H | C | C | H | T | C | T | T | C | H | H | T | H | H | H | H | H | H | T | T | C | H | H | H | T | H | G | H | H | T | H | H | H | H | H | H | H | A | H | C | G | A | H | H | G | H | H | H | H | C | G | H | H | H | A | H | H | G | A | C | H | H | H | H | A | H | A | C | G |
| 2016A_P1-F8_4 | PHI2 | PHI1 | progeny | 2016A_P1-F8 | | H | G | C | H | A | C | T | C | H | H | H | T | H | H | C | C | T | T | C | T | T | C | H | H | T | A | A | A | A | G | A | T | T | C | G | T | C | T | T | G | H | C | T | H | H | H | H | H | C | G | A | C | C | G | H | H | T | G | G | H | H | H | C | G | H | C | H | A | C | G | G | H | C | G | H | H | H | A | H | A | C | G |
| 2016A_P1-G12 | PHI2 | PHI1 | mother | 2016A_P1-G12 | | H | H | C | H | A | C | T | H | H | H | H | T | H | H | C | C | H | H | H | T | H | C | H | H | T | G | G | G | C | A | G | T | T | C | C | C | H | T | H | G | H | H | T | H | H | H | H | H | H | H | H | H | H | H | A | H | H | G | H | H | H | H | C | G | H | H | H | A | H | H | G | A | C | H | H | H | H | A | H | A | H | H |
| 2016A_P1-G12_1 | PHI2 | PHI1 | progeny | 2016A_P1-G12 | | G | H | C | H | A | C | T | H | T | H | H | T | H | H | C | C | T | T | C | T | T | C | H | H | T | G | G | G | C | A | G | T | H | C | C | C | H | T | H | G | H | H | T | H | H | H | H | H | H | G | A | C | C | H | A | H | T | G | G | G | H | H | C | G | G | C | T | A | C | G | G | A | C | G | H | H | H | A | H | A | A | H |
| 2016A_P1-G12_2 | PHI2 | PHI1 | progeny | 2016A_P1-G12 | | H | H | H | H | A | C | T | H | H | H | H | T | H | H | C | C | C | C | H | T | H | C | H | H | T | G | G | G | C | A | G | T | T | C | C | C | H | T | H | G | H | H | T | H | H | H | H | H | H | H | H | H | H | H | A | C | H | G | H | H | H | H | C | G | H | H | H | A | C | G | G | A | C | H | H | H | H | A | H | A | H | H |
| 2016A_P1-G12_6 | PHI2 | PHI1 | progeny | 2016A_P1-G12 | | G | A | C | G | A | C | T | T | T | A | C | T | G | G | C | C | H | H | H | T | H | C | H | H | T | G | G | G | C | A | G | T | T | C | C | C | H | T | H | G | H | H | T | H | H | H | C | A | T | T | G | T | A | H | A | H | H | G | H | H | C | A | C | G | G | C | T | A | H | H | G | A | C | H | H | H | H | A | H | A | H | H |
| 2016A_P1-H11 | PHI2 | PHI1 | mother | 2016A_P1-H11 | | H | H | C | H | A | C | T | H | H | H | H | T | H | H | H | H | H | H | H | T | H | C | H | H | T | H | H | H | H | H | H | T | T | C | H | H | H | T | H | G | H | H | T | H | H | H | H | H | H | H | H | H | H | H | A | H | H | G | H | H | H | H | C | G | H | H | H | A | H | H | G | A | C | H | H | H | H | A | H | A | H | H |
| 2016A_P1-H11_3 | PHI2 | PHI1 | progeny | 2016A_P1-H11 | | H | H | C | H | A | C | T | H | H | H | H | T | H | H | H | H | H | H | H | T | H | C | H | H | T | H | A | A | A | G | A | T | T | C | G | H | H | T | H | G | H | H | T | G | C | A | T | G | C | H | H | H | H | H | A | H | H | G | G | G | C | A | C | G | H | H | A | A | T | A | G | A | C | A | C | C | T | A | H | A | H | H |
| 2016A_P1-H11_4 | PHI2 | PHI1 | progeny | 2016A_P1-H11 | | H | H | C | H | A | C | T | H | H | H | H | T | H | H | H | H | H | H | H | T | T | C | H | H | T | H | A | A | A | G | A | T | T | C | G | H | T | T | T | G | G | H | T | G | C | H | T | G | C | G | A | C | C | G | A | H | H | G | H | H | H | H | C | G | G | C | A | A | H | H | G | A | C | H | H | H | H | A | H | A | A | A |
| 2016A_P1-H8 | PHI2 | PHI1 | mother | 2016A_P1-H8 | | H | H | C | H | A | C | T | H | H | H | H | T | H | H | C | C | H | T | C | T | H | C | H | H | T | G | G | G | C | A | G | T | T | C | C | C | H | T | H | G | H | H | T | H | H | H | H | H | H | H | H | H | H | H | A | H | H | G | H | H | H | H | C | G | H | H | H | A | H | H | G | A | C | H | H | H | H | A | H | A | H | H |
| 2016A_P1-H8_1 | PHI2 | PHI1 | progeny | 2016A_P1-H8 | | H | H | C | H | A | C | T | H | H | H | H | T | H | H | C | C | H | T | C | T | T | C | C | C | T | G | G | G | C | A | G | T | T | C | C | C | T | T | T | G | G | C | T | H | H | H | H | H | H | H | H | H | H | H | A | H | H | G | H | C | G | C | C | G | H | H | H | A | H | H | G | A | C | H | H | H | H | A | H | A | H | H |
| 2016A_P1-H8_16 | PHI2 | PHI1 | progeny | 2016A_P1-H8 | | H | H | C | H | A | C | T | C | C | G | T | T | T | H | C | C | H | T | C | T | H | C | G | T | T | G | G | G | C | A | G | T | H | C | C | C | C | T | A | G | C | A | T | H | H | H | H | G | C | G | A | C | C | H | A | H | H | G | H | C | G | H | C | G | H | H | A | A | H | G | G | A | C | G | T | A | H | A | H | A | H | A |
| 2016A_P1-H8_4 | PHI2 | PHI1 | progeny | 2016A_P1-H8 | | H | H | C | H | A | C | T | H | H | H | H | T | H | H | C | C | H | T | C | T | H | C | H | H | T | G | G | G | C | A | G | T | T | C | C | C | H | T | H | G | H | H | T | H | H | H | H | H | H | H | H | H | H | H | A | H | H | G | H | H | H | H | C | G | H | H | H | A | H | H | G | A | C | H | H | H | H | A | H | A | H | H |
| 2016A_P1-H8_5 | PHI2 | PHI1 | progeny | 2016A_P1-H8 | | H | H | C | H | A | C | T | H | H | H | H | T | H | H | C | C | H | T | C | T | H | C | H | H | T | G | G | G | C | A | G | T | T | C | C | C | H | T | H | G | H | H | T | H | H | H | H | H | H | H | H | H | H | H | A | C | H | G | H | H | H | H | C | G | H | H | T | A | H | H | G | A | C | H | H | H | H | A | H | A | H | H |
| 2016A_P1-H8_7 | PHI2 | PHI1 | progeny | 2016A_P1-H8 | | H | H | C | H | A | C | T | H | H | H | H | T | H | H | C | C | H | T | C | T | H | C | H | H | T | G | G | G | C | A | G | H | T | C | C | C | H | T | H | G | H | H | T | H | H | H | H | H | H | H | H | H | H | H | A | H | H | G | H | H | H | H | C | G | H | H | H | A | H | H | G | A | C | H | H | H | H | A | H | A | H | H |
| 2016A_P2-A11 | PHI2 | PHI1 | mother | 2016A_P2-A11 | | H | H | C | H | A | C | T | H | H | H | H | T | H | H | C | C | H | T | C | T | H | C | H | H | T | G | G | G | C | A | G | T | T | C | C | C | H | T | H | G | H | H | T | H | H | H | H | H | H | H | H | H | H | H | A | H | H | G | H | H | H | H | C | G | H | H | H | A | H | H | G | A | C | H | H | H | H | A | H | A | H | H |
| 2016A_P2-A11_4 | PHI2 | PHI1 | progeny | 2016A_P2-A11 | | H | H | C | H | A | C | T | H | H | H | H | T | H | H | C | C | T | T | C | T | H | C | H | H | T | G | G | G | C | A | G | T | H | C | C | C | H | T | H | G | H | H | T | A | T | G | H | H | H | H | H | H | H | G | A | G | T | G | H | H | H | H | C | G | H | H | H | A | H | H | G | A | C | H | H | H | H | A | H | A | H | H |
| 2016A_P2-A11_5 | PHI2 | PHI1 | progeny | 2016A_P2-A11 | | H | H | C | H | A | C | T | H | H | H | H | T | H | H | C | C | H | T | C | T | H | C | H | H | T | G | G | G | C | A | G | T | T | C | C | C | H | T | H | G | H | H | T | H | H | H | H | H | H | H | H | H | H | H | A | H | H | G | H | H | H | H | C | G | H | H | H | A | C | G | G | A | C | G | T | H | H | A | H | A | H | H |
| 2016A_P2-A12 | PHI2 | PHI1 | mother | 2016A_P2-A12 | | H | H | C | H | A | C | T | H | H | H | H | T | H | H | C | C | H | T | C | T | T | C | H | H | T | H | H | H | H | H | H | T | T | C | H | H | H | T | H | G | H | H | T | H | H | H | H | H | H | H | A | H | C | G | A | H | H | G | H | H | H | H | C | G | G | H | H | A | H | H | G | A | C | G | H | H | H | A | H | A | C | G |
| 2016A_P2-A12_12 | PHI2 | PHI1 | progeny | 2016A_P2-A12 | | H | H | H | H | H | C | T | H | H | H | H | T | T | G | C | C | T | T | C | T | T | C | H | H | T | H | H | H | H | H | H | H | T | C | H | H | H | H | H | G | H | H | T | H | H | H | T | G | C | H | A | H | C | G | A | G | H | G | H | H | H | H | H | G | G | H | A | H | C | H | G | A | C | G | H | H | H | H | H | H | C | H |
| 2016A_P2-A12_4 | PHI2 | PHI1 | progeny | 2016A_P2-A12 | | G | A | C | G | A | C | T | T | T | A | C | T | H | H | C | C | H | T | C | T | T | C | C | C | T | A | A | A | A | G | H | T | T | C | H | H | H | T | H | G | G | C | T | H | H | H | H | H | H | G | A | C | C | G | A | C | C | G | A | H | H | H | C | G | G | C | T | A | C | G | G | A | C | G | T | H | H | A | H | A | C | G |
| 2016A_P2-A12_7 | PHI2 | PHI1 | progeny | 2016A_P2-A12 | | G | A | C | G | A | C | T | T | H | H | H | T | H | H | C | C | H | T | C | T | T | C | H | H | T | H | A | A | A | G | H | T | T | C | H | H | H | T | H | G | H | H | T | H | H | A | T | G | C | G | A | C | C | G | A | H | H | G | H | H | H | H | C | G | G | H | T | A | C | H | G | A | C | G | H | C | T | A | C | A | C | G |
| 2016A_P2-A12_8 | PHI2 | PHI1 | progeny | 2016A_P2-A12 | | G | A | C | G | A | C | T | H | H | H | H | T | H | T | C | C | C | T | C | T | T | C | C | T | T | A | A | A | A | G | A | T | T | C | G | T | C | T | A | G | C | A | T | H | H | H | H | H | H | H | A | H | C | G | A | H | H | G | H | H | H | H | C | G | G | H | H | A | H | H | G | A | C | G | H | H | H | A | H | A | C | G |
| 2016A_P2-A12_9 | PHI2 | PHI1 | progeny | 2016A_P2-A12 | | T | G | C | H | A | C | T | H | C | A | H | T | H | H | C | C | H | H | C | T | H | C | H | T | T | A | A | A | A | G | A | T | T | C | H | C | T | T | H | G | H | H | T | H | C | H | T | H | C | G | A | C | C | G | A | C | H | G | G | H | H | H | C | G | G | H | A | A | H | H | G | A | C | G | H | H | H | A | H | A | C | G |
| 2016A_P2-A7 | PHI2 | PHI1 | mother | 2016A_P2-A7 | | H | H | C | H | A | C | T | H | H | H | H | T | H | H | C | C | H | T | C | T | H | C | H | H | T | H | G | G | C | A | G | T | T | C | C | C | H | T | H | G | H | H | T | H | H | H | H | H | H | H | H | H | H | H | A | H | H | G | H | H | H | H | C | G | H | H | T | A | H | H | G | A | C | H | H | H | H | A | H | A | H | H |
| 2016A_P2-A7_9 | PHI2 | PHI1 | progeny | 2016A_P2-A7 | | T | G | H | H | A | C | T | C | C | H | T | T | T | H | C | C | C | T | C | T | H | C | H | H | T | G | G | G | C | A | G | T | T | C | C | C | C | T | A | G | C | A | T | H | H | H | H | H | H | G | A | C | C | H | A | H | H | G | H | C | G | H | C | G | H | T | A | A | T | A | G | A | C | A | C | H | H | A | H | A | H | H |
| 2016A_P2-B10 | PHI2 | PHI1 | mother | 2016A_P2-B10 | | H | H | C | H | A | C | T | H | H | H | H | T | H | H | C | C | H | T | C | T | H | C | H | H | T | H | H | H | H | H | H | T | T | C | H | H | H | T | H | G | H | H | T | H | H | H | H | H | H | H | H | H | H | G | A | H | H | G | H | H | H | H | C | G | H | H | H | A | H | H | G | A | C | H | H | H | H | A | H | A | H | H |
| 2016A_P2-B10_6 | PHI2 | PHI1 | progeny | 2016A_P2-B10 | | G | A | C | G | A | C | T | H | H | A | C | T | G | G | C | C | H | T | C | T | H | C | G | T | T | A | A | A | A | G | A | T | T | C | H | T | C | T | A | G | G | H | T | H | H | H | H | H | H | H | A | C | C | G | A | H | H | G | G | G | C | A | C | G | H | H | A | A | T | A | G | A | C | H | H | H | C | A | H | A | H | H |
| 2016A_P2-B4 | PHI2 | PHI1 | mother | 2016A_P2-B4 | | H | H | C | H | A | C | T | H | H | H | H | T | H | H | C | C | H | T | C | T | T | C | H | H | T | H | H | H | H | H | H | T | T | C | H | H | H | T | H | G | G | H | H | H | H | H | H | H | H | H | A | H | C | G | A | H | H | G | H | H | H | H | C | G | H | H | H | A | H | H | G | A | C | G | H | H | H | A | H | A | H | H |
| 2016A_P2-B4_1 | PHI2 | PHI1 | progeny | 2016A_P2-B4 | | G | A | C | G | A | C | T | T | T | A | C | T | H | H | C | C | H | T | C | T | T | C | H | C | T | G | G | G | C | A | G | T | T | C | H | H | C | T | A | G | G | A | C | H | H | A | T | G | C | G | A | C | C | G | A | H | H | G | H | H | H | H | C | G | H | H | H | A | T | A | G | A | C | G | C | C | T | A | C | A | A | H |
| 2016A_P2-B4_5 | PHI2 | PHI1 | progeny | 2016A_P2-B4 | | G | A | C | G | A | C | T | T | T | A | H | T | T | T | C | C | H | T | C | T | T | C | H | H | T | H | A | A | A | G | A | T | T | C | G | H | C | T | H | G | G | H | H | H | H | A | T | G | C | G | A | C | C | G | A | H | H | G | H | H | H | H | C | G | H | H | H | A | T | A | G | A | C | G | C | C | T | A | C | A | H | H |
| 2016A_P2-B4_6 | PHI2 | PHI1 | progeny | 2016A_P2-B4 | | G | A | C | G | A | C | T | H | T | A | H | T | H | G | C | C | H | T | C | T | H | C | H | T | T | A | A | A | A | G | A | T | T | C | H | H | C | T | A | G | H | A | H | G | H | H | T | H | C | G | A | H | C | G | A | G | H | G | H | G | H | A | H | G | H | H | T | A | T | A | G | A | C | G | C | C | T | A | H | A | H | H |
| 2016A_P2-B7 | PHI2 | PHI1 | mother | 2016A_P2-B7 | | H | H | C | H | A | C | T | H | H | H | H | T | H | H | C | C | H | T | C | T | H | C | H | H | T | H | G | G | C | A | G | T | T | C | C | C | H | T | H | G | H | H | T | H | H | H | H | H | H | H | H | H | H | H | A | H | H | G | H | H | H | H | C | G | H | H | H | A | H | H | G | A | C | H | H | H | H | A | H | A | H | H |
| 2016A_P2-B7_1 | PHI2 | PHI1 | progeny | 2016A_P2-B7 | | G | A | C | G | A | C | T | T | T | A | C | T | G | G | C | C | T | T | C | T | H | C | H | H | T | H | G | G | C | A | G | T | T | C | C | C | C | T | A | G | C | A | T | G | H | H | H | H | H | H | H | H | H | H | A | H | H | G | H | H | H | H | C | G | H | T | A | A | H | H | G | A | C | H | H | H | H | A | H | A | H | H |
| 2016A_P2-C11 | PHI2 | PHI1 | mother | 2016A_P2-C11 | | H | H | C | H | A | C | T | H | H | H | H | T | H | H | H | C | H | T | H | T | H | C | H | H | T | H | H | H | H | H | H | T | T | C | C | H | H | T | H | G | H | H | T | H | H | H | H | H | H | H | H | H | H | H | A | H | H | G | H | H | H | H | C | G | H | H | H | A | H | H | G | A | C | H | H | H | H | A | H | A | H | H |
| 2016A_P2-C11_1 | PHI2 | PHI1 | progeny | 2016A_P2-C11 | | T | A | H | G | H | C | T | T | H | A | H | T | G | H | H | C | H | H | H | T | H | C | H | H | T | H | G | H | H | G | H | T | T | C | C | H | H | H | T | H | H | H | T | H | C | H | H | H | H | H | H | H | H | H | A | H | H | G | H | G | G | A | C | G | G | T | H | H | H | A | G | A | C | H | H | A | H | H | H | H | A | G |
| 2016A_P2-C11_4 | PHI2 | PHI1 | progeny | 2016A_P2-C11 | | G | A | C | G | A | C | T | T | T | A | C | T | H | H | H | C | H | T | H | T | G | C | G | H | T | A | A | A | A | G | A | T | H | C | C | H | H | T | H | G | H | A | T | A | T | G | C | A | T | T | G | T | A | H | A | G | T | G | G | G | H | H | C | G | H | H | A | A | T | A | G | A | C | H | H | H | H | A | H | A | H | H |
| 2016A_P2-D1 | PHI2 | PHI1 | mother | 2016A_P2-D1 | | H | H | C | H | A | C | T | H | H | H | H | T | H | H | C | C | H | T | C | T | H | C | H | H | T | H | G | G | C | A | G | T | T | C | C | C | H | T | H | G | H | H | T | H | H | H | H | H | H | H | H | H | H | H | A | H | H | G | H | H | H | H | C | G | H | H | H | A | H | H | G | A | C | H | H | H | H | A | H | A | H | H |
| 2016A_P2-D1_10 | PHI2 | PHI1 | progeny | 2016A_P2-D1 | | T | G | C | H | A | C | T | C | H | H | H | T | H | H | C | C | H | T | C | T | H | C | H | H | T | H | G | G | C | A | G | T | T | C | C | C | C | T | A | G | C | H | T | H | H | H | H | H | T | T | G | T | A | T | A | C | H | G | H | H | C | A | C | H | A | T | A | A | T | A | G | A | C | A | C | C | C | A | G | A | C | G |
| 2016A_P2-D1_12 | PHI2 | PHI1 | progeny | 2016A_P2-D1 | | G | A | C | G | A | C | T | T | H | H | H | T | H | H | C | C | H | T | C | T | H | C | H | H | T | H | G | G | C | A | G | T | T | C | C | C | H | T | H | G | C | A | T | H | C | A | T | G | C | G | A | C | C | H | A | H | H | G | H | H | G | C | C | G | H | H | H | A | T | A | G | A | C | A | H | H | H | A | H | A | C | G |
| 2016A_P2-D1_3 | PHI2 | PHI1 | progeny | 2016A_P2-D1 | | H | H | C | H | A | C | T | H | H | H | C | T | G | G | C | C | H | T | C | T | H | C | H | H | T | H | G | G | C | A | G | T | H | C | C | C | H | T | H | G | H | H | T | G | C | A | T | G | C | G | A | C | C | H | A | H | H | G | G | G | H | H | C | G | A | T | H | A | H | H | G | A | C | H | H | A | C | A | G | A | C | H |
| 2016A_P2-D1_4 | PHI2 | PHI1 | progeny | 2016A_P2-D1 | | H | H | C | H | A | C | T | H | H | H | H | T | H | H | C | C | H | T | C | T | H | C | H | H | T | H | G | G | C | A | G | T | T | C | C | C | H | T | H | G | H | A | T | H | H | H | H | H | H | T | G | T | A | T | A | C | C | G | A | H | H | H | C | G | H | H | H | A | H | H | G | A | C | H | H | H | H | A | H | A | H | H |
| 2016A_P2-D1_7 | PHI2 | PHI1 | progeny | 2016A_P2-D1 | | H | H | C | H | A | C | T | H | H | H | H | T | H | H | C | C | H | T | C | T | H | C | H | H | T | H | G | G | C | A | G | T | T | C | C | C | H | T | H | G | H | H | T | H | H | H | H | H | H | H | H | H | H | H | A | H | H | G | H | H | H | H | C | G | H | H | H | A | H | G | G | A | C | G | T | A | C | A | C | A | A | A |
| 2016A_P2-D1_8 | PHI2 | PHI1 | progeny | 2016A_P2-D1 | | H | H | C | H | A | C | T | H | H | H | H | T | H | H | C | C | H | T | C | T | H | C | H | H | T | H | G | G | C | A | G | T | T | C | C | C | H | T | H | G | H | H | T | H | H | H | H | H | H | H | H | H | H | H | A | H | H | G | H | H | H | H | C | G | H | H | H | A | H | H | G | A | C | H | H | H | H | A | G | A | C | G |
| 2016A_P2-D10 | PHI2 | PHI1 | mother | 2016A_P2-D10 | | H | H | C | H | A | C | T | H | H | H | H | T | H | H | C | C | H | T | C | T | T | C | H | H | T | H | G | G | C | A | G | T | T | C | C | C | T | T | H | G | G | H | T | H | H | H | H | H | H | H | A | H | C | G | A | H | H | G | H | H | H | H | C | G | G | H | H | A | H | H | G | A | C | G | H | H | H | A | H | A | C | G |
| 2016A_P2-D10_3 | PHI2 | PHI1 | progeny | 2016A_P2-D10 | | H | H | C | H | A | C | T | H | H | H | H | T | H | H | C | C | H | T | C | T | T | C | G | T | T | A | G | G | C | A | G | T | T | C | C | C | T | T | H | G | G | H | T | G | C | H | H | H | H | H | A | H | C | G | A | H | H | G | H | H | H | H | C | G | G | H | A | A | H | H | G | A | C | G | H | H | H | A | H | A | C | G |
| 2016A_P2-D5 | PHI2 | PHI1 | mother | 2016A_P2-D5 | | H | H | C | H | A | C | T | H | H | H | H | T | H | H | C | C | H | H | H | T | H | C | H | H | T | H | G | G | C | A | G | T | T | C | C | C | H | T | H | G | H | H | T | H | H | H | H | H | H | H | H | H | H | H | A | H | H | G | H | H | H | H | C | G | H | H | H | A | H | H | G | A | C | H | H | H | H | A | H | A | H | H |
| 2016A_P2-D5_4 | PHI2 | PHI1 | progeny | 2016A_P2-D5 | | H | G | H | H | H | C | T | H | H | H | H | T | H | G | C | C | H | H | H | T | H | C | H | H | T | G | H | H | C | H | G | H | T | C | H | H | H | H | H | G | H | H | T | H | H | H | H | H | H | H | H | H | H | H | A | H | C | G | A | H | G | H | C | G | H | H | A | H | H | A | G | A | C | H | H | C | T | H | H | H | H | H |
| 2016A_P2-D7 | PHI2 | PHI1 | mother | 2016A_P2-D7 | | H | H | C | H | A | C | T | H | H | H | H | T | H | H | C | C | H | T | C | T | H | C | H | H | T | G | G | G | C | A | G | T | T | C | C | C | H | T | H | G | H | H | T | H | H | H | H | H | H | H | H | H | H | H | A | H | H | G | H | H | H | H | C | G | H | H | H | A | H | H | G | A | C | H | H | H | H | A | H | A | H | H |
| 2016A_P2-D7_3 | PHI2 | PHI1 | progeny | 2016A_P2-D7 | | G | A | C | G | A | C | T | T | T | A | H | T | H | H | C | C | H | T | C | T | H | C | H | H | T | G | G | G | C | A | G | T | T | C | C | C | H | T | H | G | H | H | T | H | H | H | C | A | T | T | G | T | A | T | A | C | C | G | H | H | H | A | C | G | H | H | H | A | H | H | G | A | C | H | H | H | H | A | H | A | A | A |
| 2016A_P2-F2 | PHI2 | PHI1 | mother | 2016A_P2-F2 | | H | H | C | H | A | C | T | H | H | H | H | T | H | H | H | C | C | T | H | T | H | C | H | H | T | H | H | H | H | H | H | T | T | C | C | C | H | T | H | G | H | H | T | H | H | H | H | H | H | H | H | H | H | H | A | H | H | G | H | H | H | H | C | G | H | H | H | A | H | H | G | A | C | H | H | H | H | A | H | A | H | H |
| 2016A_P2-F2_2 | PHI2 | PHI1 | progeny | 2016A_P2-F2 | | G | A | C | G | A | C | T | T | T | A | C | T | G | G | A | C | C | T | H | T | H | C | H | H | T | H | H | H | H | H | H | T | T | C | C | C | H | T | H | G | H | H | T | H | H | H | T | G | C | G | A | C | C | G | A | H | H | G | G | G | C | A | C | G | A | H | H | A | H | H | G | A | C | H | H | H | H | A | H | A | H | H |
| 2016A_P2-F7 | PHI2 | PHI1 | mother | 2016A_P2-F7 | | H | H | C | H | A | C | T | H | H | H | H | T | H | H | C | C | H | T | C | T | H | C | H | H | T | H | G | G | C | A | G | T | T | C | C | C | H | T | H | G | H | H | T | H | H | H | H | H | H | H | H | H | H | H | A | H | H | G | H | H | H | H | C | G | H | H | H | A | H | H | G | A | C | H | H | H | H | A | H | A | H | H |
| 2016A_P2-F7_1 | PHI2 | PHI1 | progeny | 2016A_P2-F7 | | H | H | C | H | A | C | T | H | H | H | H | T | H | H | C | C | H | T | C | T | H | C | H | H | T | H | G | G | C | A | G | T | T | C | C | C | H | T | H | G | H | H | T | H | H | H | H | H | H | H | H | H | H | H | A | C | C | G | A | C | H | H | C | G | H | H | H | A | H | H | G | A | C | H | H | H | H | A | H | A | H | H |
| 2016A_P2-G1 | PHI2 | PHI1 | mother | 2016A_P2-G1 | | H | H | C | H | A | C | T | H | H | H | H | T | H | H | H | C | H | T | H | T | H | C | H | H | T | H | H | H | H | H | H | T | T | C | H | H | H | T | H | G | H | H | T | H | H | H | H | H | H | H | H | H | H | H | A | H | H | G | H | H | H | H | C | G | H | H | H | A | H | H | G | A | C | H | H | H | H | A | H | A | H | H |
| 2016A_P2-G1_4 | PHI2 | PHI1 | progeny | 2016A_P2-G1 | | G | A | C | G | A | C | T | T | T | A | C | T | T | T | C | C | H | T | H | T | H | C | H | H | T | G | G | G | C | A | G | T | T | C | C | C | H | T | H | G | H | H | T | H | H | H | H | H | H | H | H | H | H | H | A | H | H | G | H | H | H | H | C | G | H | H | H | A | H | H | G | A | C | H | H | H | H | A | H | A | H | H |
| 2016A_P2-G1_6 | PHI2 | PHI1 | progeny | 2016A_P2-G1 | | G | G | H | G | H | C | T | T | H | A | H | T | H | G | C | C | T | H | C | T | T | C | C | C | T | G | H | H | C | H | G | H | T | C | H | H | T | H | H | G | H | H | T | H | H | H | H | H | H | H | H | H | A | H | A | G | C | G | A | H | G | H | C | H | H | H | H | A | C | H | G | A | C | H | H | C | H | H | H | H | H | H |
| 2016A_P2-G2 | PHI2 | PHI1 | mother | 2016A_P2-G2 | | H | H | C | H | A | C | T | H | H | H | H | T | H | H | H | H | C | T | H | T | G | C | H | H | T | H | H | H | H | H | H | T | T | C | C | C | H | T | H | G | H | A | T | H | H | H | H | H | H | H | H | H | H | H | A | H | H | G | H | H | H | H | C | G | H | H | T | A | H | H | G | A | C | H | H | H | H | A | H | A | H | H |
| 2016A_P2-G2_5 | PHI2 | PHI1 | progeny | 2016A_P2-G2 | | G | A | C | G | A | C | T | T | T | A | C | T | G | T | A | T | C | T | A | T | G | C | G | H | T | A | G | G | C | A | G | T | T | C | C | C | T | T | T | G | G | C | T | A | T | G | C | G | C | G | A | C | C | G | A | G | C | G | A | C | G | C | C | G | A | T | A | A | T | A | G | A | C | A | C | C | T | A | C | A | A | A |
| 2016A_P2-G2_6 | PHI2 | PHI1 | progeny | 2016A_P2-G2 | | T | G | C | H | A | C | T | C | C | G | T | T | T | T | A | T | C | T | A | T | T | C | C | T | T | A | A | A | A | G | A | T | H | C | C | C | C | T | A | G | C | A | T | G | C | A | T | G | C | G | G | T | A | T | A | C | C | G | G | G | C | A | C | G | G | C | T | A | C | G | G | A | C | G | T | A | C | A | G | A | A | A |
| 2016A_P2-G7 | PHI2 | PHI1 | mother | 2016A_P2-G7 | | H | H | C | H | A | C | T | H | H | H | H | T | H | H | C | C | H | T | C | T | H | C | H | H | T | H | G | G | C | A | G | T | T | C | C | C | H | T | H | G | H | H | T | H | H | H | H | H | H | H | H | H | H | H | A | H | H | G | H | H | H | H | C | G | H | H | H | A | H | H | G | A | C | H | H | H | H | A | H | A | H | H |
| 2016A_P2-G7_13 | PHI2 | PHI1 | progeny | 2016A_P2-G7 | | H | H | C | H | A | C | T | C | C | G | T | T | T | T | C | C | C | T | C | T | H | C | H | H | T | H | G | G | C | A | G | T | T | C | C | C | C | T | A | G | C | A | T | H | H | H | H | H | H | H | H | H | H | H | A | H | H | G | G | G | C | A | C | G | H | H | H | A | H | H | G | A | C | H | H | H | H | A | H | A | C | G |
| 2016A_P2-G7_15 | PHI2 | PHI1 | progeny | 2016A_P2-G7 | | T | G | C | H | A | C | T | C | C | G | C | T | G | G | C | C | C | T | C | T | T | C | H | H | T | H | G | G | C | A | G | T | T | C | C | C | H | T | H | G | H | C | T | H | H | H | H | H | H | T | G | T | A | T | A | C | C | G | A | H | H | H | C | G | A | T | A | A | H | H | G | A | C | H | H | H | H | A | H | A | C | G |
| 2016A_P2-G7_6 | PHI2 | PHI1 | progeny | 2016A_P2-G7 | | H | H | C | H | A | C | T | H | H | H | T | T | H | H | C | C | T | H | C | T | G | C | G | H | T | G | G | G | C | A | G | T | H | C | C | C | T | T | T | G | H | H | T | G | C | A | H | H | H | H | H | H | H | H | A | H | H | G | H | H | C | H | C | G | H | H | H | A | H | H | G | A | C | H | H | H | H | A | H | A | H | G |
| 2016A_P2-G7_8 | PHI2 | PHI1 | progeny | 2016A_P2-G7 | | T | G | C | H | A | C | T | C | C | H | H | T | H | H | C | C | T | T | C | T | H | C | H | H | T | H | G | G | C | A | G | T | T | C | C | C | H | T | H | G | H | H | T | H | H | H | H | H | H | H | H | H | H | H | A | H | H | G | A | H | H | H | C | G | A | H | H | A | H | H | G | A | C | H | H | H | H | A | H | A | H | H |
| 2016A_P2-G9 | PHI2 | PHI1 | mother | 2016A_P2-G9 | | H | H | C | H | A | C | T | H | H | H | H | T | H | H | C | C | H | T | C | T | H | C | C | H | T | H | H | H | H | H | H | T | T | C | C | C | T | T | H | G | G | H | H | H | H | H | H | H | H | H | A | H | C | G | A | G | H | G | H | H | H | H | C | G | G | H | H | A | H | H | G | A | C | H | H | H | H | A | H | A | H | H |
| 2016A_P2-G9_3 | PHI2 | PHI1 | progeny | 2016A_P2-G9 | | G | A | C | G | A | C | T | T | T | H | H | T | H | H | C | C | H | T | C | T | H | C | C | C | T | G | G | G | C | A | G | T | T | C | C | C | T | T | T | G | G | C | T | H | T | G | C | A | T | T | A | T | C | G | A | C | C | G | A | C | G | C | C | G | G | H | H | A | T | A | G | A | C | A | H | H | H | A | H | A | C | G |
| 2016A_P2-G9_5 | PHI2 | PHI1 | progeny | 2016A_P2-G9 | | G | H | C | H | A | C | T | H | H | H | H | T | H | H | C | C | H | T | C | T | H | C | C | H | T | A | A | A | A | G | A | T | T | C | C | C | T | T | H | G | G | H | H | H | T | G | C | A | T | T | A | T | C | G | A | H | H | G | H | H | H | H | C | G | G | C | T | A | H | H | G | A | C | H | H | H | H | A | H | A | H | H |
| 2016A_P2-G9_6 | PHI2 | PHI1 | progeny | 2016A_P2-G9 | | G | A | C | G | A | C | T | T | H | H | T | T | T | H | C | C | H | T | C | T | H | C | C | H | T | H | G | G | C | A | H | T | T | C | C | C | T | T | H | G | G | H | H | H | H | H | T | G | C | G | A | C | C | G | A | H | C | G | A | C | H | H | C | G | G | H | H | A | H | H | G | A | C | H | C | C | T | A | C | A | H | H |
| 2016A_P2-H6 | PHI2 | PHI1 | mother | 2016A_P2-H6 | | H | H | C | H | A | C | T | H | H | H | H | T | H | H | C | H | H | H | H | T | H | C | C | H | T | H | H | H | H | H | H | T | T | C | H | H | H | T | H | G | H | H | H | H | H | H | H | H | H | H | A | H | C | G | A | H | H | G | H | H | H | H | C | G | H | H | H | A | H | H | G | A | C | H | H | H | H | A | H | A | C | G |
| 2016A_P2-H6_1 | PHI2 | PHI1 | progeny | 2016A_P2-H6 | | H | H | C | H | A | C | T | H | H | H | C | T | G | G | C | C | T | H | H | T | H | C | C | C | T | G | G | G | C | A | G | T | T | C | C | H | H | T | H | G | H | H | C | H | H | A | T | G | C | G | A | C | C | G | A | H | H | G | H | C | G | C | C | G | H | H | H | A | T | A | G | A | C | A | H | H | H | A | C | A | C | G |
| 2016A_P2-H6_16 | PHI2 | PHI1 | progeny | 2016A_P2-H6 | | H | H | C | H | A | C | T | H | H | H | H | T | H | H | C | H | H | H | H | T | H | C | C | H | T | H | H | H | H | H | H | T | T | C | H | H | H | T | H | G | H | H | H | H | H | H | H | H | H | H | A | H | C | G | A | H | H | G | H | H | H | H | C | G | H | H | H | A | H | H | G | A | C | H | H | H | H | A | H | A | C | G |
| 2016A_P2-H6_17 | PHI2 | PHI1 | progeny | 2016A_P2-H6 | | G | A | C | G | A | C | T | T | T | A | H | T | H | H | C | H | H | H | H | T | G | C | C | H | T | H | H | H | H | H | H | T | T | C | H | H | T | T | H | G | H | C | T | H | H | H | H | H | H | H | A | H | C | G | A | H | H | G | H | H | H | H | C | G | A | T | H | A | H | H | G | A | C | H | H | H | H | A | H | A | C | G |
| 2016A_P2-H6_9 | PHI2 | PHI1 | progeny | 2016A_P2-H6 | | H | H | C | H | A | C | T | H | H | H | H | T | H | H | C | H | H | H | H | T | H | C | C | H | T | H | H | H | H | H | H | T | T | C | H | H | H | T | H | G | H | H | H | H | H | H | H | H | H | H | A | H | C | G | A | H | H | G | H | H | H | H | C | G | H | H | H | A | H | H | G | A | C | H | H | H | H | A | H | A | C | G |
| 2016A_P3-B5 | PHI10 | PHI1 | mother | 2016A_P3-B5 | | H | H | H | H | H | H | H | C | H | G | H | H | H | T | H | H | C | C | H | T | H | H | H | H | H | G | G | G | C | A | G | H | H | H | G | T | C | T | A | G | C | A | T | A | H | H | C | H | H | T | G | T | A | T | A | C | H | H | H | C | H | H | C | H | H | H | H | H | H | H | H | A | H | A | C | C | H | H | H | H | H | A |
| 2016A_P3-B5_1 | PHI10 | PHI1 | progeny | 2016A_P3-B5 | | T | G | C | H | A | C | T | C | C | G | T | T | T | T | A | C | C | C | A | T | T | C | G | T | T | G | G | G | C | A | G | T | T | C | G | T | C | T | A | G | C | A | T | A | T | G | C | G | C | T | G | T | A | T | A | C | T | C | G | C | C | C | C | G | A | T | A | A | T | A | G | A | C | A | C | C | T | A | C | A | A | A |
| 2016A_P3-B8 | PHI10 | PHI1 | mother | 2016A_P3-B8 | | H | H | H | H | H | H | H | C | H | G | H | H | H | T | H | H | C | C | A | T | H | H | H | H | H | H | H | H | H | H | H | H | C | H | H | H | C | T | A | G | C | A | T | A | H | H | C | H | H | T | H | T | H | T | A | C | H | H | H | C | H | H | C | H | H | H | H | H | H | H | H | A | H | A | C | C | H | A | H | H | H | A |
| 2016A_P3-B8_1 | PHI10 | PHI1 | progeny | 2016A_P3-B8 | | T | G | C | H | A | C | T | C | C | G | T | T | T | T | A | C | C | C | A | T | T | C | G | T | T | A | A | A | A | G | A | T | H | C | C | C | C | T | A | G | C | A | T | A | C | A | C | G | C | T | G | T | A | T | A | C | T | G | A | C | G | C | C | G | A | T | A | A | C | G | A | A | T | A | C | C | C | A | G | G | C | A |
| 2016A_P3-B8_2 | PHI10 | PHI1 | progeny | 2016A_P3-B8 | | G | A | T | G | G | G | C | C | T | G | C | C | G | T | A | C | C | C | A | T | G | C | G | T | T | A | A | A | A | G | A | T | H | C | C | C | C | T | A | G | C | A | T | A | T | G | C | A | T | T | A | T | C | T | A | C | C | G | A | C | G | C | C | A | G | C | T | C | T | A | G | A | C | A | C | C | T | A | C | A | C | A |
| 2016A_P3-B8_4 | PHI10 | PHI1 | progeny | 2016A_P3-B8 | | G | A | T | G | G | G | C | C | T | G | T | T | T | T | C | T | C | C | A | T | G | C | G | T | T | A | A | A | A | G | A | T | C | T | G | T | C | T | A | G | C | A | T | A | T | G | C | G | C | T | G | T | A | T | A | C | C | G | A | C | G | C | C | G | A | T | A | A | T | A | G | A | C | A | C | C | T | A | C | A | A | A |
| 2016A_P3-C12 | PHI10 | PHI1 | mother | 2016A_P3-C12 | | H | H | H | H | H | H | H | C | H | H | H | H | H | T | H | H | C | H | H | T | H | H | H | H | H | G | G | G | C | A | G | H | C | H | H | H | C | T | A | G | C | A | T | A | H | H | C | H | H | T | G | T | A | T | A | C | H | H | H | C | H | H | C | H | H | H | H | H | H | H | H | A | H | A | C | C | T | H | H | H | H | A |
| 2016A_P3-C12_2 | PHI10 | PHI1 | progeny | 2016A_P3-C12 | | T | G | C | H | A | C | T | C | C | H | T | T | T | T | C | C | C | C | A | T | G | C | G | T | T | G | G | G | C | A | G | T | H | C | C | T | C | T | A | G | C | A | T | A | T | G | C | A | T | T | G | T | A | T | A | C | C | C | G | C | C | A | C | G | A | T | A | A | T | A | G | A | C | A | C | C | T | A | C | A | A | A |
| 2016A_P3-D9 | PHI10 | PHI1 | mother | 2016A_P3-D9 | | H | H | H | H | H | H | H | C | H | H | H | H | H | T | H | C | C | H | A | T | H | H | H | H | H | G | G | G | C | A | G | H | C | H | G | T | C | T | A | G | C | A | T | A | H | H | C | H | H | T | H | T | H | T | A | C | H | H | H | H | H | H | C | H | H | H | H | H | H | H | H | A | H | A | C | C | H | A | H | H | H | A |
| 2016A_P3-D9_2 | PHI10 | PHI1 | progeny | 2016A_P3-D9 | | T | G | C | H | A | C | T | C | C | H | T | T | T | T | C | C | C | T | A | T | G | C | G | C | C | G | G | G | C | A | G | G | C | H | G | T | C | T | A | G | C | A | T | A | C | A | C | G | C | T | G | T | A | T | A | C | T | C | G | G | C | A | C | A | G | C | T | C | T | A | G | A | C | A | C | C | T | A | C | A | A | A |
| 2016A_P3-H12 | PHI4 | PHI1 | mother | 2016A_P3-H12 | | T | G | C | H | H | H | H | C | C | H | H | H | H | T | H | C | C | T | C | T | G | C | H | H | H | G | G | G | C | A | G | H | H | H | H | T | C | T | A | G | H | H | H | H | T | G | H | H | H | H | H | H | H | H | A | H | H | G | H | C | G | C | H | G | H | T | H | H | T | H | G | A | H | H | H | H | T | H | H | A | A | A |
| 2016A_P3-H12_1 | PHI4 | PHI1 | progeny | 2016A_P3-H12 | | H | G | C | H | G | H | H | C | C | H | H | H | H | T | H | C | C | T | C | T | G | C | H | H | H | G | G | G | C | A | G | G | H | H | H | H | C | T | A | G | H | C | H | H | T | G | H | H | T | H | H | H | H | H | A | H | H | G | G | C | G | C | H | G | H | T | T | H | T | H | G | A | H | H | H | H | T | H | H | A | A | A |
| 2016A_P3-H8 | PHI4 | PHI1 | mother | 2016A_P3-H8 | | T | G | C | H | H | H | H | C | C | H | H | H | H | T | H | C | H | T | C | T | G | C | H | H | H | G | G | G | C | A | G | H | H | H | H | T | C | T | A | G | H | H | H | H | T | G | H | H | H | H | H | H | H | H | A | H | H | G | H | C | G | C | H | G | H | T | H | H | T | H | G | A | H | H | H | H | T | H | H | A | A | A |
| 2016A_P3-H8_1 | PHI4 | PHI1 | progeny | 2016A_P3-H8 | | T | G | C | H | H | H | H | C | C | H | H | H | T | T | H | C | H | T | C | T | G | C | H | H | H | G | G | G | C | A | G | H | H | H | H | T | C | T | A | G | H | H | H | H | T | G | H | H | H | H | H | H | H | H | A | C | H | G | A | C | G | C | H | G | H | T | H | A | T | H | G | A | H | H | H | H | T | H | H | A | A | A |
| 2016A_P4-A1 | PHI4 | PHI1 | mother | 2016A_P4-A1 | | T | G | C | H | H | H | H | C | C | H | H | H | H | H | H | H | C | H | H | T | G | H | H | H | H | H | H | H | H | H | G | H | C | H | H | T | C | T | A | G | H | H | H | H | T | G | H | H | H | H | H | H | H | H | A | C | H | H | H | C | G | C | H | G | H | T | H | H | T | H | G | A | H | H | H | H | T | H | H | A | A | A |
| 2016A_P4-A1_1 | PHI4 | PHI1 | progeny | 2016A_P4-A1 | | T | G | C | H | H | H | H | C | C | H | H | H | T | H | H | H | C | H | H | T | G | H | H | H | H | H | H | H | H | H | G | H | C | H | H | T | C | T | A | G | H | H | H | H | T | G | H | H | H | H | H | H | H | H | A | H | H | H | A | C | G | C | H | G | H | T | T | H | T | H | G | A | H | H | H | H | T | H | H | A | A | A |
| 2016A_P4-A3 | PHI4 | PHI1 | mother | 2016A_P4-A3 | | T | G | C | H | H | H | H | C | C | H | H | H | H | H | H | C | H | T | C | T | G | H | H | H | H | H | H | H | H | H | H | H | H | H | H | H | C | T | A | G | H | H | H | H | T | G | H | H | H | H | H | H | H | H | A | H | H | H | H | C | G | C | H | G | H | T | H | H | T | H | G | A | H | H | H | H | T | H | H | H | H | A |
| 2016A_P4-A3_1 | PHI4 | PHI1 | progeny | 2016A_P4-A3 | | T | G | C | H | H | H | H | C | C | H | H | H | H | H | H | C | H | T | C | T | G | H | H | H | H | H | H | H | H | H | H | H | H | H | H | H | C | T | A | G | H | H | H | H | T | G | H | H | H | H | H | H | H | H | A | H | H | H | H | C | G | C | H | G | H | T | H | H | T | H | G | A | H | H | H | H | T | H | H | H | H | A |
| 2016A_P4-A3_5 | PHI4 | PHI1 | progeny | 2016A_P4-A3 | | T | G | C | H | A | C | T | C | C | H | T | T | T | H | H | C | H | T | C | T | G | H | H | H | H | H | H | H | H | H | H | H | H | H | H | C | C | T | A | H | H | H | H | H | T | G | H | H | H | H | H | H | H | H | A | H | H | H | H | C | G | C | H | G | H | T | H | H | T | H | G | A | H | H | H | H | T | H | H | H | C | A |
| 2016A_P4-B4 | PHI4 | PHI1 | mother | 2016A_P4-B4 | | T | G | C | H | H | H | H | C | C | H | H | H | H | H | H | C | H | T | C | T | G | H | H | H | H | G | G | G | C | A | G | H | C | H | H | H | C | T | A | G | H | H | H | H | T | G | H | H | H | H | H | H | H | H | A | C | H | H | H | C | G | C | H | G | H | T | H | H | T | H | G | A | H | H | H | H | T | H | H | A | A | A |
| 2016A_P4-B4_3 | PHI4 | PHI1 | progeny | 2016A_P4-B4 | | T | G | C | H | A | H | H | C | C | H | T | T | T | T | C | C | C | T | C | T | G | H | H | H | H | G | G | G | C | A | G | H | C | H | H | H | C | T | A | G | H | C | H | H | T | G | H | H | H | H | H | C | H | H | A | C | H | C | A | C | H | C | H | G | H | T | H | H | T | H | G | A | H | H | H | H | T | H | H | A | A | A |
| 2016A_P4-C11 | PHI4 | PHI1 | mother | 2016A_P4-C11 | | T | G | C | H | H | C | H | C | C | H | H | H | H | T | A | C | H | T | C | T | G | H | H | C | H | H | G | G | C | A | G | H | C | T | H | H | C | T | A | G | H | H | H | H | T | H | H | H | H | H | H | H | H | H | A | H | H | G | H | C | G | H | H | H | A | T | A | A | T | H | G | A | H | G | H | H | T | H | G | A | A | A |
| 2016A_P4-C11_3 | PHI4 | PHI1 | progeny | 2016A_P4-C11 | | T | G | C | H | H | H | H | C | C | H | H | H | H | T | H | C | H | T | C | T | G | H | H | H | H | G | G | G | C | A | G | T | T | C | C | C | C | T | A | G | C | A | H | H | T | G | H | H | H | H | H | H | H | H | A | H | H | G | H | C | G | C | H | G | H | T | H | H | T | H | G | A | H | H | H | H | T | H | H | A | A | A |
| 2016A_P4-C2 | PHI4 | PHI1 | mother | 2016A_P4-C2 | | H | H | C | H | A | C | T | C | C | H | H | T | H | H | H | C | C | T | C | T | G | H | H | H | H | G | G | G | C | A | G | H | H | H | G | T | C | T | A | G | H | H | H | H | T | G | H | H | H | H | H | H | H | H | A | C | H | H | H | C | G | C | H | G | H | T | H | H | T | H | G | A | H | H | H | H | T | H | H | H | A | A |
| 2016A_P4-C2_2 | PHI4 | PHI1 | progeny | 2016A_P4-C2 | | G | A | C | G | A | C | T | C | C | H | H | T | H | H | H | C | C | T | C | T | G | H | H | H | H | G | G | G | C | A | G | T | T | C | G | T | C | T | A | G | H | A | T | H | T | G | H | H | H | T | G | T | A | H | A | C | H | H | H | C | G | C | H | G | G | T | T | C | T | G | G | A | T | G | T | A | T | H | H | H | A | A |
| 2016A_P4-C2_3 | PHI4 | PHI1 | progeny | 2016A_P4-C2 | | H | A | C | H | A | C | T | C | C | H | T | T | T | G | A | H | C | T | C | H | H | C | H | T | T | G | G | H | C | A | G | T | T | C | G | T | C | T | A | H | C | A | C | H | H | H | T | H | C | G | A | H | H | G | A | C | T | G | G | C | H | C | G | G | T | T | H | C | H | G | G | A | T | G | T | H | T | H | G | H | A | H |
| 2016A_P4-C2_5 | PHI4 | PHI1 | progeny | 2016A_P4-C2 | | H | H | C | H | A | C | T | C | C | H | H | T | H | H | H | C | C | T | C | T | G | H | H | H | H | G | G | G | C | A | G | H | H | H | G | T | C | T | A | G | H | H | H | G | T | G | H | H | H | T | G | T | A | H | A | C | H | H | H | C | G | C | H | G | H | T | H | H | T | A | G | A | C | H | H | H | T | A | C | H | A | A |
| 2016A_P4-C2_6 | PHI4 | PHI1 | progeny | 2016A_P4-C2 | | G | A | C | G | A | C | T | C | C | A | C | T | G | G | A | C | C | T | C | T | G | C | G | T | T | G | G | G | C | A | G | T | T | C | G | T | C | T | A | G | H | H | H | G | T | G | T | G | C | G | A | C | C | G | A | C | T | C | G | C | G | C | G | G | G | T | T | C | T | G | G | A | T | G | T | A | T | C | G | H | A | A |
| 2016A_P4-C4 | PHI4 | PHI1 | mother | 2016A_P4-C4 | | T | G | C | H | H | H | H | C | C | H | H | H | H | T | H | C | C | T | C | T | G | H | H | H | H | G | G | G | C | A | G | H | H | H | H | H | C | T | A | G | H | H | H | H | T | G | H | H | H | H | H | H | H | H | A | C | H | G | H | C | G | C | H | G | H | T | H | H | T | H | G | A | H | H | H | H | T | H | H | A | A | A |
| 2016A_P4-C4_9 | PHI4 | PHI1 | progeny | 2016A_P4-C4 | | T | G | C | H | A | H | H | C | C | H | H | H | T | T | H | C | C | T | C | T | G | H | H | H | H | G | G | G | C | A | G | H | H | H | H | H | C | T | A | G | H | A | H | H | T | G | H | H | H | H | H | H | H | H | A | C | H | G | H | C | G | C | H | H | H | T | H | H | T | H | G | A | H | H | H | H | T | H | H | A | A | A |
| 2016A_P4-C9 | PHI4 | PHI1 | mother | 2016A_P4-C9 | | T | G | C | H | H | H | H | C | C | H | H | H | H | H | H | C | C | T | C | T | G | H | H | H | H | G | G | G | C | A | G | H | C | H | H | H | C | T | A | G | H | H | H | H | T | G | H | H | H | H | A | H | C | H | A | C | H | H | H | C | G | C | H | G | H | T | H | H | T | H | G | A | H | H | H | H | T | H | H | A | A | A |
| 2016A_P4-C9_1 | PHI4 | PHI1 | progeny | 2016A_P4-C9 | | T | G | C | H | H | H | H | C | C | H | H | H | H | H | H | C | C | T | C | T | G | H | H | H | H | G | G | G | C | A | G | H | C | H | H | H | C | T | A | G | H | H | H | H | T | G | H | H | H | H | A | H | C | H | A | C | H | H | H | C | G | C | H | G | A | T | A | A | T | A | G | A | C | A | C | C | T | A | H | A | A | A |
| 2016A_P4-C9_3 | PHI4 | PHI1 | progeny | 2016A_P4-C9 | | T | G | C | H | H | H | H | C | C | H | H | H | H | H | H | C | C | T | C | T | G | H | H | H | H | G | G | G | C | A | G | H | C | H | H | H | C | T | A | G | H | H | H | G | T | G | T | G | C | G | A | C | C | G | A | C | T | C | G | C | G | C | H | G | A | T | A | A | T | H | G | A | H | H | H | H | T | H | H | A | A | A |
| 2016A_P4-D12 | PHI4 | PHI1 | mother | 2016A_P4-D12 | | T | G | C | H | H | H | H | C | C | H | H | H | H | T | H | C | H | T | C | T | H | C | H | H | H | G | G | G | C | A | G | H | C | H | H | H | C | T | A | G | H | H | H | H | T | G | H | H | H | H | H | H | H | H | A | H | H | H | H | C | G | C | H | G | H | T | H | H | T | H | G | A | H | H | H | H | T | H | H | A | A | A |
| 2016A_P4-D12_5 | PHI4 | PHI1 | progeny | 2016A_P4-D12 | | T | G | C | H | H | H | H | C | C | H | H | H | H | T | H | C | H | T | C | T | H | C | H | H | H | G | G | G | C | A | G | H | C | H | H | H | C | T | A | G | H | H | H | H | T | G | H | H | H | H | H | H | H | H | A | H | H | H | H | C | G | C | H | G | H | T | H | H | T | H | G | A | H | H | H | H | T | H | H | A | A | A |
| 2016A_P4-D5 | PHI4 | PHI1 | mother | 2016A_P4-D5 | | H | H | C | H | A | C | T | C | C | H | H | T | H | H | H | H | C | T | C | T | G | H | H | H | H | G | H | H | H | H | G | H | C | H | H | T | C | T | A | G | H | H | H | H | T | G | H | H | H | H | H | H | H | H | A | C | H | H | H | H | H | H | H | G | H | T | H | H | T | H | G | A | H | H | H | H | T | H | H | A | A | A |
| 2016A_P4-D5_1 | PHI4 | PHI1 | progeny | 2016A_P4-D5 | | H | H | C | H | A | C | T | C | C | H | H | T | H | H | H | H | C | T | C | T | G | H | H | H | H | G | H | H | H | H | G | H | C | H | H | T | C | T | A | G | H | H | C | H | T | G | H | H | H | H | H | H | H | H | A | C | H | H | H | H | H | H | H | G | H | T | H | H | T | H | G | A | H | H | H | H | T | H | H | A | A | A |
| 2016A_P4-F5 | PHI4 | PHI1 | mother | 2016A_P4-F5 | | H | H | C | H | A | C | T | C | C | H | H | T | H | H | H | H | C | H | H | T | G | H | H | H | H | H | H | H | H | H | G | H | C | H | H | T | C | T | A | G | H | H | H | H | T | G | H | H | H | H | H | H | H | H | A | C | H | H | H | C | G | H | H | G | H | T | H | H | T | H | G | A | H | H | H | H | T | H | H | A | A | A |
| 2016A_P4-F5_1 | PHI4 | PHI1 | progeny | 2016A_P4-F5 | | H | H | C | H | A | C | T | C | C | A | C | T | G | H | H | H | C | H | H | T | G | H | H | H | H | H | H | H | H | H | G | H | C | H | H | T | C | T | A | G | H | H | H | H | T | G | H | H | H | H | H | H | H | H | A | C | H | H | H | C | G | H | H | G | H | T | H | H | T | H | G | A | H | H | H | H | T | H | H | A | A | A |
| 2016A_P4-F5_2 | PHI4 | PHI1 | progeny | 2016A_P4-F5 | | G | A | C | G | A | C | T | C | C | A | T | T | T | T | C | T | C | C | A | T | G | H | H | H | H | H | H | H | H | H | G | H | C | H | H | T | C | T | A | G | C | A | T | H | T | G | H | H | H | H | H | H | H | H | A | C | H | H | H | C | G | H | H | G | H | T | H | H | T | H | G | A | H | H | H | H | T | H | H | A | A | A |
| 2016A_P4-F5_5 | PHI4 | PHI1 | progeny | 2016A_P4-F5 | | H | H | C | H | H | C | H | H | C | H | H | T | H | G | A | C | C | T | C | H | H | H | H | T | H | H | G | A | H | H | G | G | C | H | C | H | C | H | A | H | H | A | C | H | H | H | H | H | T | H | H | H | H | H | A | H | H | H | H | C | H | A | G | H | H | T | H | A | H | H | H | H | H | G | H | H | T | H | C | H | A | H |
| 2016A_P4-F5_6 | PHI4 | PHI1 | progeny | 2016A_P4-F5 | | H | H | C | H | A | C | T | C | C | H | H | T | H | H | H | H | C | H | H | T | G | H | H | H | H | H | H | H | H | H | G | H | C | H | H | T | C | T | A | G | H | C | T | H | T | G | T | G | C | G | A | C | C | H | A | C | H | H | H | C | G | C | C | G | H | T | H | H | T | H | G | A | H | H | H | H | T | H | H | A | A | A |
| 2016A_P4-F7 | PHI7 | PHI1 | mother | 2016A_P4-F7 | | H | H | H | H | H | H | H | H | H | H | H | H | H | H | H | C | H | T | C | H | H | H | H | H | H | G | G | G | C | A | G | H | C | H | H | H | H | H | H | H | H | H | H | H | H | H | H | H | T | H | A | H | C | H | H | C | H | H | H | H | H | H | G | H | H | H | H | H | H | H | H | H | H | H | H | H | H | H | H | H | H | H |
| 2016A_P4-F7_1 | PHI7 | PHI1 | progeny | 2016A_P4-F7 | | G | A | T | G | G | G | C | T | H | H | H | H | H | H | H | C | C | T | C | C | H | H | H | H | H | G | G | G | C | A | G | T | H | C | G | T | C | T | A | H | H | H | H | H | H | A | T | G | T | G | A | C | C | G | T | C | H | H | H | H | H | H | H | G | G | C | T | C | C | G | A | G | T | G | T | A | C | C | G | G | H | H |
| 2016A_P4-G11 | PHI7 | PHI1 | mother | 2016A_P4-G11 | | H | H | H | H | H | H | H | H | H | H | H | H | H | H | H | T | H | H | H | H | H | H | H | H | H | H | G | G | C | A | G | H | H | H | H | H | H | H | H | H | H | H | T | H | H | H | H | H | T | H | H | H | H | H | H | C | H | H | H | H | H | H | G | H | H | H | H | H | H | H | H | H | H | H | H | H | H | H | H | H | H | H |
| 2016A_P4-G11_2 | PHI7 | PHI1 | progeny | 2016A_P4-G11 | | H | H | H | H | H | H | H | H | H | H | H | H | H | H | H | T | H | H | H | H | H | H | H | H | H | H | G | G | C | A | G | H | H | H | H | H | H | H | H | H | H | H | T | H | H | H | H | H | T | H | H | H | H | H | H | C | H | H | H | H | H | H | G | H | H | H | H | H | H | H | H | H | H | H | H | H | H | H | H | H | H | H |
| 2016A_P4-G11_4 | PHI7 | PHI1 | progeny | 2016A_P4-G11 | | H | H | H | H | H | H | H | H | H | H | H | H | H | H | H | T | H | H | H | H | H | H | H | H | H | H | G | G | C | A | G | H | H | H | H | H | H | H | H | H | H | H | T | H | H | H | H | H | T | H | H | H | H | H | H | C | H | H | H | H | H | H | G | H | H | H | H | H | H | H | H | H | H | H | H | H | H | H | H | H | H | H |

### Table S5. Results of the SNP array analasis (continue)

| plant_id | mom_gen | dad_gen | mother_progeny | family_ |
|---|---|---|---|---|
| 2016A_P4-H1 | PHI7 | PHI1 | mother | 2016A_P4-H1 |
| 2016A_P4-H1_2 | PHI7 | PHI1 | progeny | 2016A_P4-H1 |
| 2016A_P4-H1_3 | PHI7 | PHI1 | progeny | 2016A_P4-H1 |
| 2016A_P4-H1_4 | PHI7 | PHI1 | progeny | 2016A_P4-H1 |
| 2016A_P4-H10 | PHI7 | PHI1 | mother | 2016A_P4-H10 |
| 2016A_P4-H10_4 | PHI7 | PHI1 | progeny | 2016A_P4-H10 |
| 2016A_P4-H5 | PHI7 | PHI1 | mother | 2016A_P4-H5 |
| 2016A_P4-H5_1 | PHI7 | PHI1 | progeny | 2016A_P4-H5 |
| 2016A_P4-H5_2 | PHI7 | PHI1 | progeny | 2016A_P4-H5 |
| 2016A_P5-A1 | PHI7 | PHI1 | mother | 2016A_P5-A1 |
| 2016A_P5-A1_3 | PHI7 | PHI1 | progeny | 2016A_P5-A1 |
| 2016A_P5-A1_4 | PHI7 | PHI1 | progeny | 2016A_P5-A1 |
| 2016A_P5-B11 | PHI7 | PHI1 | mother | 2016A_P5-B11 |
| 2016A_P5-B11_1 | PHI7 | PHI1 | progeny | 2016A_P5-B11 |
| 2016A_P5-B11_7 | PHI7 | PHI1 | progeny | 2016A_P5-B11 |
| 2016A_P5-C8 | PHI7 | PHI1 | mother | 2016A_P5-C8 |
| 2016A_P5-C8_1 | PHI7 | PHI1 | progeny | 2016A_P5-C8 |
| 2016A_P5-C8_2 | PHI7 | PHI1 | progeny | 2016A_P5-C8 |
| 2016A_P5-F1 | PHI6 | PHI1 | mother | 2016A_P5-F1 |
| 2016A_P5-F1_1 | PHI6 | PHI1 | progeny | 2016A_P5-F1 |
| 2016A_P5-F10 | PHI6 | PHI1 | mother | 2016A_P5-F10 |
| 2016A_P5-F10_2 | PHI6 | PHI1 | progeny | 2016A_P5-F10 |
| 2016A_P5-F10_4 | PHI6 | PHI1 | progeny | 2016A_P5-F10 |
| 2016A_P5-F10_7 | PHI6 | PHI1 | progeny | 2016A_P5-F10 |
| 2016A_P5-F2 | PHI6 | PHI1 | mother | 2016A_P5-F2 |
| 2016A_P5-F2_3 | PHI6 | PHI1 | progeny | 2016A_P5-F2 |
| 2016A_P5-G10 | PHI6 | PHI1 | mother | 2016A_P5-G10 |
| 2016A_P5-G10_10 | PHI6 | PHI1 | progeny | 2016A_P5-G10 |
| 2016A_P5-G10_11 | PHI6 | PHI1 | progeny | 2016A_P5-G10 |
| 2016A_P5-G10_6 | PHI6 | PHI1 | progeny | 2016A_P5-G10 |
| 2016A_P5-G10_7 | PHI6 | PHI1 | progeny | 2016A_P5-G10 |
| 2016A_P5-G10_8 | PHI6 | PHI1 | progeny | 2016A_P5-G10 |
| 2016A_P5-G10_9 | PHI6 | PHI1 | progeny | 2016A_P5-G10 |
| 2016A_P5-G12 | PHI6 | PHI1 | mother | 2016A_P5-G12 |
| 2016A_P5-G12_1 | PHI6 | PHI1 | progeny | 2016A_P5-G12 |
| 2016A_P5-G8 | PHI6 | PHI1 | mother | 2016A_P5-G8 |
| 2016A_P5-G8_3 | PHI6 | PHI1 | progeny | 2016A_P5-G8 |
| 2016A_P5-G8_4 | PHI6 | PHI1 | progeny | 2016A_P5-G8 |
| 2016A_P5-G8_5 | PHI6 | PHI1 | progeny | 2016A_P5-G8 |
| 2016A_P5-G9 | PHI6 | PHI1 | mother | 2016A_P5-G9 |
| 2016A_P5-G9_1 | PHI6 | PHI1 | progeny | 2016A_P5-G9 |
| 2016A_P5-G9_5 | PHI6 | PHI1 | progeny | 2016A_P5-G9 |
| 2016A_P5-G9_6 | PHI6 | PHI1 | progeny | 2016A_P5-G9 |
| 2016A_P5-H10 | PHI6 | PHI1 | mother | 2016A_P5-H10 |
| 2016A_P5-H10_2 | PHI6 | PHI1 | progeny | 2016A_P5-H10 |
| 2016A_P5-H10_3 | PHI6 | PHI1 | progeny | 2016A_P5-H10 |
| 2016A_P5-H10_5 | PHI6 | PHI1 | progeny | 2016A_P5-H10 |
| 2016A_P5-H11 | PHI6 | PHI1 | mother | 2016A_P5-H11 |
| 2016A_P5-H11_2 | PHI6 | PHI1 | progeny | 2016A_P5-H11 |
| 2016A_P5-H11_3 | PHI6 | PHI1 | progeny | 2016A_P5-H11 |
| 2016A_P5-H11_5 | PHI6 | PHI1 | progeny | 2016A_P5-H11 |
| 2016A_P5-H11_6 | PHI6 | PHI1 | progeny | 2016A_P5-H11 |
| 2016A_P5-H12_11 | PHI6 | PHI1 | progeny | 2016A_P5-H12 |
| 2016A_P5-H12_14 | PHI6 | PHI1 | progeny | 2016A_P5-H12 |
| 2016A_P5-H12_16 | PHI6 | PHI1 | progeny | 2016A_P5-H12 |
| 2016A_P5-H12_17 | PHI6 | PHI1 | progeny | 2016A_P5-H12 |
| 2016A_P5-H12_3 | PHI6 | PHI1 | progeny | 2016A_P5-H12 |
| 2016A_P5-H12_9 | PHI6 | PHI1 | progeny | 2016A_P5-H12 |
| 2016A_P5-H3 | PHI6 | PHI1 | mother | 2016A_P5-H3 |
| 2016A_P5-H3_1 | PHI6 | PHI1 | progeny | 2016A_P5-H3 |
| 2016A_P5-H3_11 | PHI6 | PHI1 | progeny | 2016A_P5-H3 |
| 2016A_P5-H3_2 | PHI6 | PHI1 | progeny | 2016A_P5-H3 |
| 2016A_P5-H3_3 | PHI6 | PHI1 | progeny | 2016A_P5-H3 |
| 2016A_P5-H3_4 | PHI6 | PHI1 | progeny | 2016A_P5-H3 |
| 2016A_P5-H3_5 | PHI6 | PHI1 | progeny | 2016A_P5-H3 |
| 2016A_P5-H3_6 | PHI6 | PHI1 | progeny | 2016A_P5-H3 |
| 2016A_P5-H3_9 | PHI6 | PHI1 | progeny | 2016A_P5-H3 |
| 2016A_P5-H5 | PHI6 | PHI1 | mother | 2016A_P5-H5 |
| 2016A_P5-H5_1 | PHI6 | PHI1 | progeny | 2016A_P5-H5 |
| 2016A_P5-H5_10 | PHI6 | PHI1 | progeny | 2016A_P5-H5 |
| 2016A_P5-H5_12 | PHI6 | PHI1 | progeny | 2016A_P5-H5 |
| 2016A_P5-H5_3 | PHI6 | PHI1 | progeny | 2016A_P5-H5 |
| 2016A_P5-H5_4 | PHI6 | PHI1 | progeny | 2016A_P5-H5 |
| 2016A_P5-H5_5 | PHI6 | PHI1 | progeny | 2016A_P5-H5 |
| 2016A_P5-H5_6 | PHI6 | PHI1 | progeny | 2016A_P5-H5 |
| 2016A_P5-H5_7 | PHI6 | PHI1 | progeny | 2016A_P5-H5 |
| 2016A_P5-H5_8 | PHI6 | PHI1 | progeny | 2016A_P5-H5 |
| 2016A_P5-H6 | PHI6 | PHI1 | mother | 2016A_P5-H6 |
| 2016A_P5-H6_1 | PHI6 | PHI1 | progeny | 2016A_P5-H6 |
| 2016A_P5-H8 | PHI6 | PHI1 | mother | 2016A_P5-H8 |
| 2016A_P5-H8_1 | PHI6 | PHI1 | progeny | 2016A_P5-H8 |
| 2016A_P5-H8_10 | PHI6 | PHI1 | progeny | 2016A_P5-H8 |
| 2016A_P5-H8_16 | PHI6 | PHI1 | progeny | 2016A_P5-H8 |
| 2016A_P5-H8_4 | PHI6 | PHI1 | progeny | 2016A_P5-H8 |
| 2016A_P5H12 | PHI6 | PHI1 | mother | 2016A_P5H12 |
| 2016A_P6-A8 | PHI6 | PHI1 | mother | 2016A_P6-A8 |
| 2016A_P6-A8_1 | PHI6 | PHI1 | progeny | 2016A_P6-A8 |
| 2016A_P6-A8_11 | PHI6 | PHI1 | progeny | 2016A_P6-A8 |
| 2016A_P6-A8_12 | PHI6 | PHI1 | progeny | 2016A_P6-A8 |
| 2016A_P6-A8_16 | PHI6 | PHI1 | progeny | 2016A_P6-A8 |
| 2016A_P6-A8_8 | PHI6 | PHI1 | progeny | 2016A_P6-A8 |
| 2016A_P6-A9 | PHI6 | PHI1 | mother | 2016A_P6-A9 |
| 2016A_P6-A9_1 | PHI6 | PHI1 | progeny | 2016A_P6-A9 |
| 2016A_P6-A9_2 | PHI6 | PHI1 | progeny | 2016A_P6-A9 |
| 2016A_P6-A9_4 | PHI6 | PHI1 | progeny | 2016A_P6-A9 |
| 2016A_P6-A9_6 | PHI6 | PHI1 | progeny | 2016A_P6-A9 |
| 2016A_P6-A9_8 | PHI6 | PHI1 | progeny | 2016A_P6-A9 |
| 2016A_P6-B10 | PHI6 | PHI1 | mother | 2016A_P6-B10 |
| 2016A_P6-B10_11 | PHI6 | PHI1 | progeny | 2016A_P6-B10 |
| 2016A_P6-B10_12 | PHI6 | PHI1 | progeny | 2016A_P6-B10 |
| 2016A_P6-B10_13 | PHI6 | PHI1 | progeny | 2016A_P6-B10 |
| 2016A_P6-B10_15 | PHI6 | PHI1 | progeny | 2016A_P6-B10 |
| 2016A_P6-B10_4 | PHI6 | PHI1 | progeny | 2016A_P6-B10 |
| 2016A_P6-B10_8 | PHI6 | PHI1 | progeny | 2016A_P6-B10 |
| 2016A_P6-B12 | PHI6 | PHI1 | mother | 2016A_P6-B12 |
| 2016A_P6-B12_4 | PHI6 | PHI1 | progeny | 2016A_P6-B12 |
| 2016A_P6-B12_7 | PHI6 | PHI1 | progeny | 2016A_P6-B12 |
| 2016A_P6-B7 | PHI6 | PHI1 | mother | 2016A_P6-B7 |
| 2016A_P6-B7_1 | PHI6 | PHI1 | progeny | 2016A_P6-B7 |
| 2016A_P6-B7_2 | PHI6 | PHI1 | progeny | 2016A_P6-B7 |
| 2016A_P6-B8 | PHI6 | PHI1 | mother | 2016A_P6-B8 |
| 2016A_P6-B8_2 | PHI6 | PHI1 | progeny | 2016A_P6-B8 |
| 2016A_P6-B8_3 | PHI6 | PHI1 | progeny | 2016A_P6-B8 |
| 2016A_P6-C10 | PHI6 | PHI1 | mother | 2016A_P6-C10 |
| 2016A_P6-C10_1 | PHI6 | PHI1 | progeny | 2016A_P6-C10 |
| 2016A_P6-C10_2 | PHI6 | PHI1 | progeny | 2016A_P6-C10 |
| 2016A_P6-C10_3 | PHI6 | PHI1 | progeny | 2016A_P6-C10 |
| 2016A_P6-C12 | PHI6 | PHI1 | mother | 2016A_P6-C12 |
| 2016A_P6-C12_1 | PHI6 | PHI1 | progeny | 2016A_P6-C12 |
| 2016A_P6-C12_2 | PHI6 | PHI1 | progeny | 2016A_P6-C12 |
| 2016A_P6-C12_3 | PHI6 | PHI1 | progeny | 2016A_P6-C12 |
| 2016A_P6-C12_4 | PHI6 | PHI1 | progeny | 2016A_P6-C12 |
| 2016A_P6-D11 | PHI6 | PHI1 | mother | 2016A_P6-D11 |
| 2016A_P6-D11_1 | PHI6 | PHI1 | progeny | 2016A_P6-D11 |
| 2016A_P6-E5 | PHI8 | PHI1 | mother | 2016A_P6-E5 |
| 2016A_P6-E5_4 | PHI8 | PHI1 | progeny | 2016A_P6-E5 |
| 2016A_P6-F7 | PHI8 | PHI1 | mother | 2016A_P6-F7 |
| 2016A_P6-F7_12 | PHI8 | PHI1 | progeny | 2016A_P6-F7 |
| 2016A_P6-G9 | PHI8 | PHI1 | mother | 2016A_P6-G9 |
| 2016A_P6-G9_4 | PHI8 | PHI1 | progeny | 2016A_P6-G9 |
| 2016A_P6-H1 | PHI8 | PHI1 | mother | 2016A_P6-H1 |
| 2016A_P6-H1_3 | PHI8 | PHI1 | progeny | 2016A_P6-H1 |
| 2016A_P6-H3_3 | PHI8 | PHI1 | progeny | 2016A_P6-H3 |
| 2016A_P7-A8 | PHI8 | PHI1 | mother | 2016A_P7-A8 |

| Chr | 7 | 7 | 7 | 7 | 7 | 7 | 7 | 7 | 7 | 7 | 7 | 7 | 7 | 7 | 7 | 7 | 7 | 7 | 7 | 8 | 8 | 8 | 8 | 8 | 8 | 8 | 8 | 8 | 8 | 8 | 8 | 8 | 8 | 8 | 8 | 8 | 8 | 8 | 8 | 8 | 8 | 8 | 8 | 9 | 9 | 9 | 9 | 9 | 9 | 9 | 9 | 9 | 9 | 9 | 9 | 9 | 9 | 9 | 9 | 9 | 9 | 9 | 9 | 10 | 10 | 10 | 10 | 10 | 10 | 10 | 10 | 10 | 10 | 10 | 10 | 10 | 10 | 10 | 10 | 10 | 10 | 10 |
|---|---|---|---|---|---|---|---|---|---|---|---|---|---|---|---|---|---|---|---|---|---|---|---|---|---|---|---|---|---|---|---|---|---|---|---|---|---|---|---|---|---|---|---|---|---|---|---|---|---|---|---|---|---|---|---|---|---|---|---|---|---|---|---|---|---|---|---|---|---|---|---|---|---|---|---|---|---|---|---|---|---|---|
| Position, cM | 76 | 78 | 77 | 80 | 83 | 84 | 84 | 98 | 114 | 121 | 129 | 142 | 151 | 170 | 177 | 186 | 196 | 204 | 206 | 3 | 15 | 31 | 46 | 48 | 54 | 68 | 85 | 84 | 85 | 87 | 95 | 105 | 114 | 122 | 130 | 137 | 149 | 158 | 168 | 174 | 182 | 197 | 210 | 12 | 26 | 38 | 46 | 56 | 60 | 72 | 76 | 76 | 77 | 89 | 101 | 109 | 124 | 134 | 141 | 152 | 163 | 174 | 181 | 4.1 | 10 | 18 | 30 | 44 | 58 | 62 | 61 | 61 | 62 | 71 | 81 | 93 | 98 | 114 | 128 | 140 | 154 | 162 |
| 2016A_P4-H1 | H | H | H | H | H | H | H | H | H | H | H | H | H | H | H | C | H | T | C | H | H | H | H | H | H | G | G | G | C | A | G | H | C | H | C | C | H | H | H | H | H | H | H | H | H | H | H | H | T | H | H | H | H | H | H | C | H | H | H | H | H | H | H | H | H | H | H | H | H | H | H | H | H | H | H | H | H | H | H | H | H | H |
| 2016A_P4-H1_2 | G | A | T | G | G | G | C | H | H | H | H | H | H | H | H | C | H | T | C | H | H | H | H | H | H | G | G | G | C | A | G | G | C | T | C | C | C | T | A | G | C | A | T | H | H | H | H | G | T | G | A | C | C | G | T | C | H | H | H | H | H | H | H | H | H | H | H | H | T | A | G | A | C | A | C | C | T | H | H | H | C | G |
| 2016A_P4-H1_3 | H | H | H | H | H | H | H | H | H | H | H | H | H | H | H | C | H | T | C | T | G | C | G | T | T | G | G | G | C | A | G | T | H | C | C | C | H | H | H | H | H | H | H | G | C | A | T | H | T | H | H | H | H | H | H | C | H | H | H | H | H | H | H | H | H | H | H | H | H | A | H | H | C | A | C | C | T | A | C | A | A | A |
| 2016A_P4-H1_4 | H | H | H | H | H | H | H | H | H | H | H | H | H | H | H | C | H | T | C | H | H | H | H | H | H | G | G | G | C | A | G | H | C | H | C | C | H | H | H | H | H | H | H | A | T | G | C | A | T | T | H | H | H | H | H | C | H | H | H | H | H | H | H | H | H | H | H | H | H | H | H | H | H | H | H | H | H | H | H | H | H | H |
| 2016A_P4-H10 | H | H | H | H | H | H | H | H | H | H | H | H | H | H | H | C | H | T | C | T | H | H | H | H | H | G | G | G | C | A | G | H | C | H | H | H | H | H | H | H | H | H | H | H | H | H | H | H | T | H | A | H | C | H | H | C | H | H | H | H | H | H | H | G | H | H | H | H | H | H | H | H | H | H | H | H | H | H | H | H | H | H |
| 2016A_P4-H10_4 | H | H | H | H | H | H | H | H | H | H | H | H | H | H | H | C | H | T | C | T | H | H | H | H | H | G | G | G | C | A | G | H | C | H | H | H | H | H | A | G | C | A | T | H | H | H | H | H | T | H | A | H | C | H | H | C | H | H | H | H | H | H | H | G | H | H | H | H | H | H | H | H | H | H | H | H | H | H | H | H | H | H |
| 2016A_P4-H5 | H | H | H | H | H | H | H | H | H | H | H | H | H | H | H | C | H | T | C | H | H | H | H | H | H | G | G | G | C | A | G | H | H | H | C | H | H | H | H | H | H | H | T | H | H | H | H | H | T | H | H | H | H | H | H | C | H | H | H | H | H | H | H | H | H | H | H | H | H | H | H | H | H | H | H | H | H | H | H | H | H | H |
| 2016A_P4-H5_1 – 2016A_P7-A8 | [illegible] | | | | | | | | | | | | | | | | | | | | | | | | | | | | | | | | | | | | | | | | | | | | | | | | | | | | | | | | | | | | | | | | | | | | | | | | | | | | | | | | | |

**Table S5. Results of the SNP array analasis (continue)**

| | | | | | Chr | 7 | 7 | 7 | 7 | 7 | 7 | 7 | 7 | 7 | 7 | 7 | 7 | 7 | 7 | 7 | 7 | 7 | 7 | 7 |
|---|---|---|---|---|---|---|---|---|---|---|---|---|---|---|---|---|---|---|---|---|---|---|---|---|
| | | | | | Position, cM | 76 | 78 | 77 | 80 | 83 | 84 | 84 | 98 | 114 | 121 | 129 | 142 | 151 | 170 | 177 | 186 | 196 | 204 | 206 |
| plant_id | mom_gen | dad_gen | mother_progeny | family_ | | | | | | | | | | | | | | | | | | | | |
| 2016A_P7-A8_1 | PHI8 | PHI1 | progeny | 2016A_P7-A8 | | G | A | T | G | G | G | C | T | T | A | C | C | H | H | C | C | C | T | C |
| 2016A_P7-C11 | PHI5 | PHI1 | mother | 2016A_P7-C11 | | H | H | H | H | H | H | H | H | H | H | H | H | H | H | H | C | C | T | C |
| 2016A_P7-C11_1 | PHI5 | PHI1 | progeny | 2016A_P7-C11 | | G | A | T | G | G | G | C | T | T | A | C | C | G | G | A | C | C | T | C |
| 2016A_P7-D10 | PHI5 | PHI1 | mother | 2016A_P7-D10 | | H | H | H | H | H | H | H | H | H | H | H | H | H | H | H | H | C | H | C |
| 2016A_P7-D10_1 | PHI5 | PHI1 | progeny | 2016A_P7-D10 | | G | A | T | G | G | G | C | T | T | A | C | T | T | T | H | C | C | T | C |
| 2016A_P7-D2 | PHI5 | PHI1 | mother | 2016A_P7-D2 | | T | H | H | H | A | H | H | H | H | H | H | T | H | T | A | H | H | T | C |
| 2016A_P7-D2_1 | PHI5 | PHI1 | progeny | 2016A_P7-D2 | | G | A | T | G | G | G | C | T | T | A | C | C | G | G | C | C | C | T | C |
| 2016A_P7-D8 | PHI5 | PHI1 | mother | 2016A_P7-D8 | | H | H | H | H | H | H | H | H | H | H | H | H | H | H | H | C | C | T | C |
| 2016A_P7-D8_2 | PHI5 | PHI1 | progeny | 2016A_P7-D8 | | G | A | T | G | G | G | C | T | T | A | H | C | G | T | H | C | C | T | C |
| 2016A_P7-G3 | PHI5 | PHI1 | mother | 2016A_P7-G3 | | H | H | H | H | H | H | H | H | H | H | H | H | H | H | H | C | C | T | C |
| 2016A_P7-G3_1 | PHI5 | PHI1 | progeny | 2016A_P7-G3 | | H | H | H | H | G | H | T | H | C | H | H | H | G | H | H | C | C | T | C |
| 2016A_P7-G3_3 | PHI5 | PHI1 | progeny | 2016A_P7-G3 | | T | G | C | A | A | C | T | C | C | G | T | T | T | T | C | C | C | T | C |
| 2016A_P7-G3_4 | PHI5 | PHI1 | progeny | 2016A_P7-G3 | | G | A | T | G | G | G | C | T | T | A | C | C | G | G | H | C | C | T | C |
| 2016A_P7-G7 | PHI5 | PHI1 | mother | 2016A_P7-G7 | | H | H | H | H | H | H | H | H | H | H | H | H | H | H | H | C | H | T | C |
| 2016A_P7-G7_1 | PHI5 | PHI1 | progeny | 2016A_P7-G7 | | H | H | H | H | H | H | H | H | H | H | H | H | H | H | H | C | H | T | C |
| 2016A_P7-H9 | PHI5 | PHI1 | mother | 2016A_P7-H9 | | H | H | H | H | H | H | H | H | H | H | H | H | H | H | H | C | H | T | C |
| 2016A_P7-H9_1 | PHI5 | PHI1 | progeny | 2016A_P7-H9 | | H | H | H | H | H | H | H | H | H | H | T | H | H | H | H | H | C | H | H |
| 2016A_P7-H9_2 | PHI5 | PHI1 | progeny | 2016A_P7-H9 | | H | H | H | H | H | H | H | H | H | H | H | H | H | H | H | H | C | H | H |
| 2016A_P8-A1 | PHI5 | PHI1 | mother | 2016A_P8-A1 | | H | H | H | H | H | H | H | H | H | H | H | H | H | H | H | H | H | H | H |
| 2016A_P8-A1_1 | PHI5 | PHI1 | progeny | 2016A_P8-A1 | | T | G | C | H | A | C | T | C | C | G | T | T | T | T | C | T | C | C | A |
| 2016A_P8-A5 | PHI5 | PHI1 | mother | 2016A_P8-A5 | | H | H | H | H | H | H | H | H | H | H | H | H | H | H | H | C | C | T | C |
| 2016A_P8-A5_1 | PHI5 | PHI1 | progeny | 2016A_P8-A5 | | H | H | H | H | H | H | H | H | H | H | H | H | H | H | H | C | H | T | C |
| 2016A_P8-A6 | PHI5 | PHI1 | mother | 2016A_P8-A6 | | H | H | H | H | H | H | H | H | H | H | H | H | H | H | H | C | C | T | C |
| 2016A_P8-A6_1 | PHI5 | PHI1 | progeny | 2016A_P8-A6 | | H | H | H | H | H | H | H | H | H | H | T | T | T | H | H | C | C | T | C |
| 2016A_P8-D5 | PHI9 | PHI1 | mother | 2016A_P8-D5 | | H | H | H | H | H | H | H | H | C | H | H | T | T | H | H | H | H | H | H |
| 2016A_P8-D5_3 | PHI9 | PHI1 | progeny | 2016A_P8-D5 | | H | H | H | H | H | H | H | H | C | H | H | T | T | H | H | H | H | H | H |
| 2016A_P8-D5_5 | PHI9 | PHI1 | progeny | 2016A_P8-D5 | | T | G | C | H | A | C | T | C | C | G | T | T | T | H | H | H | H | H | H |
| 2016A_P8-E1 | PHI9 | PHI1 | mother | 2016A_P8-E1 | | H | H | H | H | H | H | H | H | C | H | H | T | T | H | H | H | H | H | H |
| 2016A_P8-E1_2 | PHI9 | PHI1 | progeny | 2016A_P8-E1 | | H | H | H | H | H | H | H | H | C | H | H | T | T | H | H | C | H | T | C |
| 2016A_P9-A10 | PHI9 | PHI1 | mother | 2016A_P9-A10 | | H | H | H | H | H | H | H | H | C | H | H | T | T | H | C | C | H | T | C |
| 2016A_P9-A10_1 | PHI9 | PHI1 | progeny | 2016A_P9-A10 | | T | G | C | H | A | C | T | C | C | G | T | T | T | T | C | C | C | T | C |
| 2016A_P9-A10_3 | PHI9 | PHI1 | progeny | 2016A_P9-A10 | | T | G | C | H | A | C | T | C | C | G | T | T | T | T | C | C | C | T | C |
| 2016A_P9-A7 | PHI9 | PHI1 | mother | 2016A_P9-A7 | | H | H | H | H | H | H | H | H | C | H | H | T | T | H | H | H | H | H | H |
| 2016A_P9-A7_1 | PHI9 | PHI1 | progeny | 2016A_P9-A7 | | H | H | H | H | H | H | H | H | C | H | H | T | T | H | A | H | H | H | H |
| 2016A_P9-A7_2 | PHI9 | PHI1 | progeny | 2016A_P9-A7 | | H | H | H | H | H | H | H | H | C | H | H | T | T | H | H | H | H | H | H |
| 2016A_P9-B1 | PHI9 | PHI1 | mother | 2016A_P9-B1 | | H | H | H | H | H | H | H | H | C | H | H | T | T | H | C | C | H | H | H |
| 2016A_P9-B1_2 | PHI9 | PHI1 | progeny | 2016A_P9-B1 | | G | A | T | G | G | G | C | T | C | A | C | T | T | G | C | C | C | C | A |
| 2016A_P9-B7 | PHI9 | PHI1 | mother | 2016A_P9-B7 | | H | H | H | H | H | H | H | H | C | H | H | T | T | H | C | C | H | T | C |
| 2016A_P9-B7_10 | PHI9 | PHI1 | progeny | 2016A_P9-B7 | | G | A | T | G | G | G | C | T | C | A | C | T | T | T | C | C | C | T | C |
| 2016A_P9-B7_5 | PHI9 | PHI1 | progeny | 2016A_P9-B7 | | T | G | C | H | A | C | T | T | C | A | T | T | T | T | C | C | C | T | C |
| 2016A_P9-B7_9 | PHI9 | PHI1 | progeny | 2016A_P9-B7 | | G | A | T | G | G | G | C | T | C | A | C | T | T | G | C | C | T | T | C |

| | Chr | 8 | 8 | 8 | 8 | 8 | 8 | 8 | 8 | 8 | 8 | 8 | 8 | 8 | 8 | 8 | 8 | 8 | 8 | 8 | 8 | 8 | 8 | 8 | 8 |
|---|---|---|---|---|---|---|---|---|---|---|---|---|---|---|---|---|---|---|---|---|---|---|---|---|---|
| | Position, cM | 3 | 15 | 31 | 46 | 48 | 54 | 68 | 85 | 84 | 85 | 87 | 95 | 105 | 114 | 122 | 130 | 137 | 149 | 158 | 168 | 174 | 182 | 197 | 210 |
| plant_id | | | | | | | | | | | | | | | | | | | | | | | | | |
| 2016A_P7-A8_1 | | H | H | H | H | H | H | G | G | G | C | A | G | H | C | H | H | H | H | H | H | H | H | H | T |
| 2016A_P7-C11 | | H | H | H | H | H | H | H | H | H | H | H | H | H | H | H | C | C | H | H | H | H | H | H | H |
| 2016A_P7-C11_1 | | H | H | T | C | C | C | G | G | G | C | A | G | G | H | H | C | C | H | H | H | H | H | H | H |
| 2016A_P7-D10 | | T | H | C | H | H | H | H | H | H | H | H | G | H | C | H | C | H | H | H | H | H | H | H | H |
| 2016A_P7-D10_1 | | C | T | T | C | C | C | G | G | G | C | A | G | G | C | T | C | C | T | C | T | T | G | C | C |
| 2016A_P7-D2 | | H | H | H | H | H | H | G | G | G | C | A | G | T | H | H | C | H | H | H | H | G | H | C | H |
| 2016A_P7-D2_1 | | T | G | C | G | T | T | G | G | G | C | A | G | T | T | C | C | T | C | T | A | G | C | A | T |
| 2016A_P7-D8 | | H | H | H | H | H | H | G | G | G | C | A | G | H | C | H | C | H | H | H | H | H | H | H | H |
| 2016A_P7-D8_2 | | H | H | H | H | H | H | G | G | G | C | A | G | H | C | H | C | H | H | H | H | H | H | H | H |
| 2016A_P7-G3 | | H | H | H | H | H | H | G | G | G | C | A | G | H | C | H | C | H | H | H | H | H | H | H | H |
| 2016A_P7-G3_1 | | H | H | H | H | H | H | G | G | G | C | A | G | H | C | H | C | C | H | H | H | H | H | H | H |
| 2016A_P7-G3_3 | | H | T | T | C | C | C | G | G | G | C | A | G | G | C | T | C | C | T | C | T | T | G | C | C |
| 2016A_P7-G3_4 | | H | H | C | G | T | T | G | G | G | C | A | G | T | H | C | C | H | T | C | T | T | G | C | C |
| 2016A_P7-G7 | | H | H | H | H | H | H | G | G | G | C | A | G | H | H | H | C | C | H | H | H | H | H | H | H |
| 2016A_P7-G7_1 | | H | H | H | H | H | H | G | G | G | C | A | G | H | H | H | C | C | H | H | H | H | H | A | T |
| 2016A_P7-H9 | | H | H | H | H | H | H | G | G | G | C | A | G | H | C | H | C | H | H | H | H | H | H | H | H |
| 2016A_P7-H9_1 | | T | H | C | H | H | H | G | G | G | C | A | G | H | C | H | C | C | H | H | H | H | H | H | H |
| 2016A_P7-H9_2 | | T | H | C | H | H | H | G | G | G | C | A | G | H | C | H | C | C | H | H | H | H | H | H | H |
| 2016A_P8-A1 | | H | H | H | H | H | H | H | H | H | H | H | H | H | H | H | H | H | H | H | H | H | H | H | H |
| 2016A_P8-A1_1 | | T | G | C | G | H | H | H | H | H | H | H | H | H | H | H | H | H | H | H | H | H | H | H | H |
| 2016A_P8-A5 | | H | H | H | H | H | H | G | G | G | C | A | G | H | C | H | C | H | H | H | H | H | H | H | H |
| 2016A_P8-A5_1 | | H | H | H | H | H | H | G | H | H | C | H | G | H | H | H | H | H | H | H | H | H | H | H | H |
| 2016A_P8-A6 | | H | H | H | H | H | H | G | G | G | C | A | G | H | H | H | C | H | H | H | H | H | H | H | H |
| 2016A_P8-A6_1 | | H | H | H | H | C | C | G | G | G | C | A | G | G | C | T | C | C | T | C | T | H | H | H | H |
| 2016A_P8-D5 | | H | H | C | H | H | T | G | G | G | C | A | G | T | T | C | H | T | H | T | A | H | H | H | H |
| 2016A_P8-D5_3 | | H | H | C | H | H | T | G | G | G | C | A | G | T | T | C | H | T | H | T | A | H | H | H | H |
| 2016A_P8-D5_5 | | C | T | C | C | C | T | G | G | G | C | A | G | T | T | C | H | T | H | T | A | H | H | H | H |
| 2016A_P8-E1 | | H | H | H | H | H | H | G | H | H | H | H | H | T | T | C | C | H | H | T | A | H | H | C | H |
| 2016A_P8-E1_2 | | H | H | H | H | H | H | G | G | G | C | A | G | T | T | H | H | H | H | T | A | H | H | H | T |
| 2016A_P9-A10 | | H | H | H | H | H | H | H | G | G | C | A | G | T | T | H | H | T | H | T | A | H | H | H | T |
| 2016A_P9-A10_1 | | C | T | T | C | C | C | G | G | G | C | A | G | T | H | T | C | T | T | T | A | T | G | C | T |
| 2016A_P9-A10_3 | | C | T | T | C | C | T | A | G | G | C | A | G | T | T | C | G | T | C | T | A | T | G | C | T |
| 2016A_P9-A7 | | H | H | H | H | H | H | H | H | H | H | H | H | T | T | H | C | H | H | T | A | H | H | H | H |
| 2016A_P9-A7_1 | | H | G | H | H | H | H | H | H | H | H | H | H | T | T | H | C | H | H | T | A | H | H | H | H |
| 2016A_P9-A7_2 | | H | H | C | H | H | H | H | H | H | H | H | H | T | T | H | C | H | H | T | A | H | H | H | H |
| 2016A_P9-B1 | | H | H | H | H | H | H | H | H | H | H | H | H | T | T | H | C | T | H | T | A | H | H | H | H |
| 2016A_P9-B1_2 | | T | G | C | G | T | T | A | A | A | A | G | A | T | T | C | C | T | C | T | A | G | C | C | C |
| 2016A_P9-B7 | | H | H | C | H | H | T | G | G | G | C | A | G | T | T | H | C | H | H | T | A | H | H | H | H |
| 2016A_P9-B7_10 | | C | T | C | C | C | T | G | G | G | C | A | G | T | T | T | C | T | T | T | A | T | G | C | C |
| 2016A_P9-B7_5 | | T | G | C | G | C | T | G | G | G | C | A | G | T | T | T | C | T | T | T | A | G | C | A | T |
| 2016A_P9-B7_9 | | C | T | C | G | T | T | G | G | G | C | A | G | T | T | C | C | C | C | T | A | G | C | A | T |

| | Chr | 9 | 9 | 9 | 9 | 9 | 9 | 9 | 9 | 9 | 9 | 9 | 9 | 9 | 9 | 9 | 9 | 9 | 9 | 9 | 9 |
|---|---|---|---|---|---|---|---|---|---|---|---|---|---|---|---|---|---|---|---|---|---|
| | Position, cM | 12 | 26 | 38 | 46 | 56 | 60 | 72 | 76 | 76 | 77 | 89 | 101 | 109 | 124 | 134 | 141 | 152 | 163 | 174 | 181 |
| plant_id | | | | | | | | | | | | | | | | | | | | | |
| 2016A_P7-A8_1 | | H | H | H | H | H | H | H | A | H | C | H | A | H | H | G | A | H | H | H | H |
| 2016A_P7-C11 | | H | H | H | H | H | H | H | A | H | C | H | A | C | H | G | H | H | H | C | H |
| 2016A_P7-C11_1 | | H | T | G | C | A | T | T | A | T | C | T | A | C | C | G | A | H | H | C | H |
| 2016A_P7-D10 | | H | H | H | H | H | H | H | H | H | H | H | H | C | H | G | H | H | H | H | C |
| 2016A_P7-D10_1 | | G | T | G | C | A | T | T | G | T | A | T | A | C | C | G | A | H | G | C | G |
| 2016A_P7-D2 | | H | H | H | H | H | T | H | H | T | H | H | A | H | H | G | H | H | H | C | H |
| 2016A_P7-D2_1 | | A | T | G | C | A | T | T | G | T | A | T | A | C | C | G | G | G | C | C | C |
| 2016A_P7-D8 | | H | H | H | H | H | H | H | H | H | H | H | A | H | H | G | H | H | H | C | H |
| 2016A_P7-D8_2 | | G | C | A | T | G | C | G | A | C | C | G | A | H | H | G | H | H | H | C | H |
| 2016A_P7-G3 | | H | H | H | H | H | H | H | H | H | H | H | A | H | H | G | H | H | H | C | H |
| 2016A_P7-G3_1 | | H | H | H | H | H | H | H | H | H | H | H | A | G | H | G | H | H | H | C | H |
| 2016A_P7-G3_3 | | H | T | G | C | A | T | T | G | T | A | T | A | H | H | G | H | H | H | C | H |
| 2016A_P7-G3_4 | | H | H | H | C | A | T | T | G | T | H | H | A | G | T | G | G | G | C | C | G |
| 2016A_P7-G7 | | H | H | H | H | H | H | H | H | H | H | H | A | H | H | G | H | H | H | C | H |
| 2016A_P7-G7_1 | | H | H | H | C | A | T | T | G | T | A | T | A | C | H | G | H | H | H | C | H |
| 2016A_P7-H9 | | H | H | H | H | H | H | H | H | H | H | H | H | C | H | H | H | H | H | C | H |
| 2016A_P7-H9_1 | | H | H | H | H | H | H | H | A | H | C | H | H | C | H | H | H | H | H | H | C |
| 2016A_P7-H9_2 | | H | H | H | H | H | H | H | A | H | C | H | H | C | H | H | H | H | H | H | C |
| 2016A_P8-A1 | | H | H | H | H | H | H | H | H | H | H | H | H | C | H | H | H | H | H | C | H |
| 2016A_P8-A1_1 | | H | T | G | C | A | T | T | G | T | A | T | A | C | C | G | A | H | G | C | H |
| 2016A_P8-A5 | | H | H | H | H | H | H | H | H | H | H | H | A | H | H | G | H | H | H | C | H |
| 2016A_P8-A5_1 | | H | H | H | H | H | H | H | H | H | H | H | A | H | H | G | H | H | H | H | H |
| 2016A_P8-A6 | | H | H | H | H | H | H | H | H | H | H | H | A | H | H | G | H | H | H | C | H |
| 2016A_P8-A6_1 | | A | T | G | C | A | T | T | G | T | A | T | A | C | C | G | A | H | H | C | H |
| 2016A_P8-D5 | | H | H | G | C | H | H | H | H | H | H | H | H | H | H | H | H | H | H | H | H |
| 2016A_P8-D5_3 | | H | H | G | C | H | H | H | H | H | H | H | H | H | H | H | H | H | H | H | H |
| 2016A_P8-D5_5 | | A | T | G | C | A | T | T | G | T | H | H | H | H | H | H | H | H | H | H | H |
| 2016A_P8-E1 | | H | H | G | C | H | H | H | H | H | H | H | H | G | H | C | H | H | H | H | H |
| 2016A_P8-E1_2 | | H | H | G | C | H | H | H | H | H | H | H | H | H | H | H | H | H | H | H | H |
| 2016A_P9-A10 | | H | H | G | C | H | H | H | H | H | H | H | H | H | H | H | H | H | H | H | H |
| 2016A_P9-A10_1 | | A | T | G | C | A | T | G | A | C | C | G | T | G | T | C | A | C | G | C | C |
| 2016A_P9-A10_3 | | G | C | G | C | G | C | G | A | C | C | G | T | C | C | G | A | C | G | C | C |
| 2016A_P9-A7 | | H | H | G | C | H | H | H | H | H | H | H | H | H | H | H | H | H | H | H | H |
| 2016A_P9-A7_1 | | H | H | G | C | H | H | H | H | H | H | H | A | H | H | H | H | H | H | H | H |
| 2016A_P9-A7_2 | | H | H | G | C | H | H | H | H | H | H | H | H | H | H | H | H | H | H | H | H |
| 2016A_P9-B1 | | H | H | G | C | H | H | H | H | H | H | H | H | H | H | H | H | H | H | H | H |
| 2016A_P9-B1_2 | | A | T | G | C | A | T | T | G | T | A | T | A | C | C | G | A | C | G | C | C |
| 2016A_P9-B7 | | H | H | G | C | H | H | H | H | H | H | H | H | H | H | H | H | H | H | H | H |
| 2016A_P9-B7_10 | | A | T | G | C | G | C | G | A | C | C | G | T | G | T | C | G | G | C | A | G |
| 2016A_P9-B7_5 | | G | C | G | C | A | T | T | G | T | A | T | A | C | C | C | G | G | C | A | G |
| 2016A_P9-B7_9 | | A | T | G | C | A | T | T | G | T | A | T | A | C | C | C | G | G | C | A | G |

| | Chr | 10 | 10 | 10 | 10 | 10 | 10 | 10 | 10 | 10 | 10 | 10 | 10 | 10 | 10 | 10 | 10 | 10 | 10 | 10 |
|---|---|---|---|---|---|---|---|---|---|---|---|---|---|---|---|---|---|---|---|---|
| | Position, cM | 4.1 | 10 | 18 | 30 | 44 | 58 | 62 | 61 | 61 | 62 | 71 | 81 | 93 | 98 | 114 | 128 | 140 | 154 | 162 |
| plant_id | | | | | | | | | | | | | | | | | | | | |
| 2016A_P7-A8_1 | | H | H | C | T | C | C | G | A | G | T | G | T | A | C | C | H | A | H | H |
| 2016A_P7-C11 | | H | H | H | A | H | H | H | H | H | H | H | H | H | H | H | H | G | A | A |
| 2016A_P7-C11_1 | | G | H | T | A | A | T | A | G | A | C | A | C | C | T | A | C | A | A | A |
| 2016A_P7-D10 | | H | H | H | A | H | H | H | H | H | H | H | H | H | H | H | H | H | A | A |
| 2016A_P7-D10_1 | | A | G | C | A | C | C | H | H | H | H | G | T | A | C | C | G | G | H | A |
| 2016A_P7-D2 | | G | H | H | A | C | H | H | H | H | H | H | H | H | H | H | H | H | A | A |
| 2016A_P7-D2_1 | | G | A | T | A | A | T | A | G | A | C | G | T | A | C | C | G | H | A | A |
| 2016A_P7-D8 | | H | H | H | A | H | H | H | H | H | H | H | H | H | H | H | H | H | A | A |
| 2016A_P7-D8_2 | | H | H | H | A | H | T | A | G | A | C | A | C | C | H | H | H | H | A | A |
| 2016A_P7-G3 | | H | H | H | H | H | H | H | H | H | H | H | H | H | H | H | H | H | A | A |
| 2016A_P7-G3_1 | | H | H | H | H | H | H | H | H | H | H | H | H | H | H | H | H | A | A | A |
| 2016A_P7-G3_3 | | H | H | H | A | A | T | A | G | A | C | A | C | C | H | H | H | H | A | A |
| 2016A_P7-G3_4 | | H | H | H | A | C | C | G | A | G | T | H | H | H | H | H | H | H | A | A |
| 2016A_P7-G7 | | H | H | H | A | H | H | H | H | H | H | H | H | H | H | H | H | H | A | A |
| 2016A_P7-G7_1 | | H | H | H | A | H | T | A | G | A | C | A | C | C | T | A | C | A | A | A |
| 2016A_P7-H9 | | H | H | H | A | H | H | H | H | H | H | H | H | H | H | H | H | H | A | A |
| 2016A_P7-H9_1 | | H | H | C | A | H | H | H | H | H | H | H | H | H | H | H | H | H | A | H |
| 2016A_P7-H9_2 | | H | H | H | A | H | H | H | H | H | H | H | H | H | H | H | H | H | A | A |
| 2016A_P8-A1 | | H | H | H | A | H | H | H | H | H | H | H | H | H | H | H | H | H | A | A |
| 2016A_P8-A1_1 | | H | H | H | A | H | H | H | H | H | H | H | H | H | H | H | H | H | A | A |
| 2016A_P8-A5 | | H | H | H | A | H | H | H | H | H | H | H | H | H | H | H | H | H | A | A |
| 2016A_P8-A5_1 | | H | G | H | A | C | C | H | H | H | H | H | T | H | H | C | G | H | H | H |
| 2016A_P8-A6 | | H | H | H | A | H | H | H | H | H | H | H | H | H | H | H | H | H | A | A |
| 2016A_P8-A6_1 | | H | H | H | A | H | C | G | A | G | T | G | T | A | C | C | G | G | A | A |
| 2016A_P8-D5 | | H | H | H | H | H | T | H | G | A | H | H | H | H | H | H | H | H | C | H |
| 2016A_P8-D5_3 | | H | H | H | H | H | T | H | G | A | H | H | H | H | H | H | H | H | C | H |
| 2016A_P8-D5_5 | | H | H | H | H | A | T | A | G | A | C | A | H | H | H | H | H | H | C | H |
| 2016A_P8-E1 | | H | H | H | H | H | T | H | G | A | H | H | H | H | H | H | H | A | C | H |
| 2016A_P8-E1_2 | | H | H | H | H | H | T | H | G | A | H | H | H | H | H | H | H | H | C | H |
| 2016A_P9-A10 | | H | H | H | H | H | T | H | G | A | H | H | H | H | H | H | H | A | H | H |
| 2016A_P9-A10_1 | | G | A | T | A | A | T | A | G | A | C | A | C | C | T | C | G | A | C | G |
| 2016A_P9-A10_3 | | A | G | C | T | C | T | G | G | A | T | G | T | A | C | C | C | A | C | G |
| 2016A_P9-A7 | | H | H | H | H | H | T | H | G | A | H | H | H | H | H | H | H | A | H | A |
| 2016A_P9-A7_1 | | H | H | H | T | H | T | H | G | A | H | H | H | H | H | H | C | A | A | A |
| 2016A_P9-A7_2 | | H | H | H | H | H | T | H | G | A | H | H | H | H | H | H | H | A | H | A |
| 2016A_P9-B1 | | H | H | H | H | H | T | H | G | A | H | H | H | H | H | H | H | A | H | A |
| 2016A_P9-B1_2 | | A | G | C | T | C | T | G | G | A | T | G | T | A | C | C | C | A | A | A |
| 2016A_P9-B7 | | H | H | H | H | H | T | H | G | A | H | H | H | H | H | H | H | A | H | H |
| 2016A_P9-B7_10 | | G | A | T | A | A | T | G | G | A | T | G | T | A | C | C | G | A | C | G |
| 2016A_P9-B7_5 | | G | A | T | A | A | T | A | G | A | C | A | C | C | T | A | C | A | C | G |
| 2016A_P9-B7_9 | | A | G | C | T | C | T | G | G | A | T | G | T | A | C | A | C | A | A | A |

plant_id – unique identifier of the plant
mom_gen – genetic backgroiund of mother in the hybred
dad_gen – genetic background of father in the hybred
family_ – identify mother and all the progeny plants from the same mother
mother_progeny – identify each plant as mother of the family or progeny
Chr – number of the chromosome, where marker is located
Distance – position of the marker on the corresponding chromosome in cM
H – heterozygous marker
A, T, G and C – marker homozygous for the corresponding nucleotide
EQV, LS, -9 , etc. – missing values

**Table S6. Results of the SNP array analasis**

| | | | | | Chr | 1 | 1 | 1 |
|---|---|---|---|---|---|---|---|---|
| | | | | | Position, CM | 82 | 105 | 156 |
| plant_id | mom_gen | dad_gen | mother_pr | family_ | | | | |
| 2016A_P1-A3 | PHI1 | PHI3 | mother | 2016A_P1-A3 | | C | H | H |
| 2016A_P1-A3_1 | PHI1 | PHI3 | progeny | 2016A_P1-A3 | | C | H | H |
| 2016A_P1-D5 | PHI1 | PHI3 | mother | 2016A_P1-D5 | | C | H | H |
| 2016A_P1-D5_1 | PHI1 | PHI3 | progeny | 2016A_P1-D5 | | C | H | H |
| 2016A_P1-D8 | PHI1 | PHI3 | mother | 2016A_P1-D8 | | H | H | H |
| 2016A_P1-D8_3 | PHI1 | PHI3 | progeny | 2016A_P1-D8 | | C | C | C |
| 2016A_P1-E3 | PHI1 | PHI3 | mother | 2016A_P1-E3 | | C | H | H |
| 2016A_P1-E3_3 | PHI1 | PHI3 | progeny | 2016A_P1-E3 | | C | C | C |
| 2016A_P1-E3_5 | PHI1 | PHI3 | progeny | 2016A_P1-E3 | | C | H | H |
| 2016A_P1-E9 | PHI1 | PHI3 | mother | 2016A_P1-E9 | | H | H | H |
| 2016A_P1-E9_1 | PHI1 | PHI3 | progeny | 2016A_P1-E9 | | C | C | C |
| 2016A_P1-F7 | PHI1 | PHI3 | mother | 2016A_P1-F7 | | H | H | H |
| 2016A_P1-F7_4 | PHI1 | PHI3 | progeny | 2016A_P1-F7 | | C | C | C |
| 2016A_P1-F8 | PHI2 | PHI1 | mother | 2016A_P1-F8 | | H | H | C |
| 2016A_P1-F8_4 | PHI2 | PHI1 | progeny | 2016A_P1-F8 | | H | H | C |
| 2016A_P1-G12 | PHI2 | PHI1 | mother | 2016A_P1-G12 | | H | H | C |
| 2016A_P1-G12_1 | PHI2 | PHI1 | progeny | 2016A_P1-G12 | | H | H | C |
| 2016A_P1-G12_2 | PHI2 | PHI1 | progeny | 2016A_P1-G12 | | T | H | C |
| 2016A_P1-G12_6 | PHI2 | PHI1 | progeny | 2016A_P1-G12 | | T | T | C |
| 2016A_P1-H11 | PHI2 | PHI1 | mother | 2016A_P1-H11 | | H | H | C |
| 2016A_P1-H11_3 | PHI2 | PHI1 | progeny | 2016A_P1-H11 | | T | H | C |
| 2016A_P1-H11_4 | PHI2 | PHI1 | progeny | 2016A_P1-H11 | | T | H | C |
| 2016A_P1-H8 | PHI2 | PHI1 | mother | 2016A_P1-H8 | | H | H | C |
| 2016A_P1-H8_1 | PHI2 | PHI1 | progeny | 2016A_P1-H8 | | C | C | C |
| 2016A_P1-H8_16 | PHI2 | PHI1 | progeny | 2016A_P1-H8 | | H | C | C |
| 2016A_P1-H8_4 | PHI2 | PHI1 | progeny | 2016A_P1-H8 | | H | H | C |
| 2016A_P1-H8_5 | PHI2 | PHI1 | progeny | 2016A_P1-H8 | | H | H | C |
| 2016A_P1-H8_7 | PHI2 | PHI1 | progeny | 2016A_P1-H8 | | H | H | C |
| 2016A_P2-A11 | PHI2 | PHI1 | mother | 2016A_P2-A11 | | H | H | C |
| 2016A_P2-A11_4 | PHI2 | PHI1 | progeny | 2016A_P2-A11 | | H | H | C |
| 2016A_P2-A11_5 | PHI2 | PHI1 | progeny | 2016A_P2-A11 | | H | H | C |
| 2016A_P2-A12 | PHI2 | PHI1 | mother | 2016A_P2-A12 | | H | H | C |
| 2016A_P2-A12_12 | PHI2 | PHI1 | progeny | 2016A_P2-A12 | | C | C | C |
| 2016A_P2-A12_4 | PHI2 | PHI1 | progeny | 2016A_P2-A12 | | T | C | C |
| 2016A_P2-A12_7 | PHI2 | PHI1 | progeny | 2016A_P2-A12 | | H | C | C |
| 2016A_P2-A12_8 | PHI2 | PHI1 | progeny | 2016A_P2-A12 | | H | H | H |
| 2016A_P2-A12_9 | PHI2 | PHI1 | progeny | 2016A_P2-A12 | | H | H | C |
| 2016A_P2-A7 | PHI2 | PHI1 | mother | 2016A_P2-A7 | | H | H | C |
| 2016A_P2-A7_9 | PHI2 | PHI1 | progeny | 2016A_P2-A7 | | H | C | C |
| 2016A_P2-B10 | PHI2 | PHI1 | mother | 2016A_P2-B10 | | H | H | C |
| 2016A_P2-B10_6 | PHI2 | PHI1 | progeny | 2016A_P2-B10 | | C | C | C |
| 2016A_P2-B4 | PHI2 | PHI1 | mother | 2016A_P2-B4 | | H | H | C |
| 2016A_P2-B4_1 | PHI2 | PHI1 | progeny | 2016A_P2-B4 | | C | C | C |
| 2016A_P2-B4_5 | PHI2 | PHI1 | progeny | 2016A_P2-B4 | | H | H | C |
| 2016A_P2-B4_6 | PHI2 | PHI1 | progeny | 2016A_P2-B4 | | C | C | C |
| 2016A_P2-B7 | PHI2 | PHI1 | mother | 2016A_P2-B7 | | H | H | C |
| 2016A_P2-B7_1 | PHI2 | PHI1 | progeny | 2016A_P2-B7 | | H | H | C |
| 2016A_P2-C11 | PHI2 | PHI1 | mother | 2016A_P2-C11 | | H | H | C |
| 2016A_P2-C11_1 | PHI2 | PHI1 | progeny | 2016A_P2-C11 | | H | H | C |
| 2016A_P2-C11_4 | PHI2 | PHI1 | progeny | 2016A_P2-C11 | | H | H | C |
| 2016A_P2-D1 | PHI2 | PHI1 | mother | 2016A_P2-D1 | | H | H | C |
| 2016A_P2-D1_10 | PHI2 | PHI1 | progeny | 2016A_P2-D1 | | H | H | C |
| 2016A_P2-D1_12 | PHI2 | PHI1 | progeny | 2016A_P2-D1 | | H | C | C |
| 2016A_P2-D1_3 | PHI2 | PHI1 | progeny | 2016A_P2-D1 | | H | H | C |
| 2016A_P2-D1_4 | PHI2 | PHI1 | progeny | 2016A_P2-D1 | | H | H | C |
| 2016A_P2-D1_7 | PHI2 | PHI1 | progeny | 2016A_P2-D1 | | H | H | C |
| 2016A_P2-D1_8 | PHI2 | PHI1 | progeny | 2016A_P2-D1 | | H | H | C |
| 2016A_P2-D10 | PHI2 | PHI1 | mother | 2016A_P2-D10 | | H | H | C |
| 2016A_P2-D5 | PHI2 | PHI1 | mother | 2016A_P2-D5 | | H | H | C |
| 2016A_P2-D5_4 | PHI2 | PHI1 | progeny | 2016A_P2-D5 | | H | H | C |
| 2016A_P2-D7 | PHI2 | PHI1 | mother | 2016A_P2-D7 | | H | H | C |
| 2016A_P2-D7_3 | PHI2 | PHI1 | progeny | 2016A_P2-D7 | | H | T | C |
| 2016A_P2-F2 | PHI2 | PHI1 | mother | 2016A_P2-F2 | | H | H | C |
| 2016A_P2-F2_2 | PHI2 | PHI1 | progeny | 2016A_P2-F2 | | H | H | C |
| 2016A_P2-F7 | PHI2 | PHI1 | mother | 2016A_P2-F7 | | H | H | C |
| 2016A_P2-F7_1 | PHI2 | PHI1 | progeny | 2016A_P2-F7 | | H | H | C |
| 2016A_P2-G1 | PHI2 | PHI1 | mother | 2016A_P2-G1 | | H | H | C |
| 2016A_P2-G1_4 | PHI2 | PHI1 | progeny | 2016A_P2-G1 | | T | T | C |
| 2016A_P2-G1_6 | PHI2 | PHI1 | progeny | 2016A_P2-G1 | | H | C | C |
| 2016A_P2-G2 | PHI2 | PHI1 | mother | 2016A_P2-G2 | | H | H | C |
| 2016A_P2-G2_5 | PHI2 | PHI1 | progeny | 2016A_P2-G2 | | C | C | C |
| 2016A_P2-G2_6 | PHI2 | PHI1 | progeny | 2016A_P2-G2 | | C | C | C |
| 2016A_P2-G7 | PHI2 | PHI1 | mother | 2016A_P2-G7 | | H | H | C |
| 2016A_P2-G7_13 | PHI2 | PHI1 | progeny | 2016A_P2-G7 | | C | H | C |
| 2016A_P2-G7_15 | PHI2 | PHI1 | progeny | 2016A_P2-G7 | | H | H | C |
| 2016A_P2-G7_6 | PHI2 | PHI1 | progeny | 2016A_P2-G7 | | H | H | C |
| 2016A_P2-G7_8 | PHI2 | PHI1 | progeny | 2016A_P2-G7 | | H | H | C |
| 2016A_P2-G9 | PHI2 | PHI1 | mother | 2016A_P2-G9 | | H | H | C |
| 2016A_P2-G9_3 | PHI2 | PHI1 | progeny | 2016A_P2-G9 | | H | T | C |
| 2016A_P2-G9_5 | PHI2 | PHI1 | progeny | 2016A_P2-G9 | | C | H | C |
| 2016A_P2-G9_6 | PHI2 | PHI1 | progeny | 2016A_P2-G9 | | H | H | C |
| 2016A_P2-H6 | PHI2 | PHI1 | mother | 2016A_P2-H6 | | H | H | C |
| 2016A_P2-H6_1 | PHI2 | PHI1 | progeny | 2016A_P2-H6 | | H | T | C |
| 2016A_P2-H6_16 | PHI2 | PHI1 | progeny | 2016A_P2-H6 | | H | H | C |
| 2016A_P2-H6_17 | PHI2 | PHI1 | progeny | 2016A_P2-H6 | | T | H | C |
| 2016A_P2-H6_9 | PHI2 | PHI1 | progeny | 2016A_P2-H6 | | H | H | C |
| 2016A_P3-B5 | PHI10 | PHI1 | mother | 2016A_P3-B5 | | H | C | C |
| 2016A_P3-B5_1 | PHI10 | PHI1 | progeny | 2016A_P3-B5 | | C | C | C |
| 2016A_P3-B8 | PHI10 | PHI1 | mother | 2016A_P3-B8 | | H | C | C |
| 2016A_P3-B8_1 | PHI10 | PHI1 | progeny | 2016A_P3-B8 | | T | C | C |
| 2016A_P3-B8_2 | PHI10 | PHI1 | progeny | 2016A_P3-B8 | | T | C | C |
| 2016A_P3-B8_4 | PHI10 | PHI1 | progeny | 2016A_P3-B8 | | C | C | C |
| 2016A_P3-C12 | PHI10 | PHI1 | mother | 2016A_P3-C12 | | H | C | C |
| 2016A_P3-C12_2 | PHI10 | PHI1 | progeny | 2016A_P3-C12 | | T | H | C |
| 2016A_P3-D9 | PHI10 | PHI1 | mother | 2016A_P3-D9 | | H | C | C |
| 2016A_P3-D9_2 | PHI10 | PHI1 | progeny | 2016A_P3-D9 | | C | C | C |
| 2016A_P3-H12 | PHI4 | PHI1 | mother | 2016A_P3-H12 | | C | C | C |
| 2016A_P3-H12_1 | PHI4 | PHI1 | progeny | 2016A_P3-H12 | | C | C | C |
| 2016A_P3-H8 | PHI4 | PHI1 | mother | 2016A_P3-H8 | | C | C | C |
| 2016A_P3-H8_1 | PHI4 | PHI1 | progeny | 2016A_P3-H8 | | C | C | C |
| 2016A_P4-A1 | PHI4 | PHI1 | mother | 2016A_P4-A1 | | C | C | C |
| 2016A_P4-A1_1 | PHI4 | PHI1 | progeny | 2016A_P4-A1 | | C | C | C |
| 2016A_P4-A3 | PHI4 | PHI1 | mother | 2016A_P4-A3 | | C | C | C |
| 2016A_P4-A3_1 | PHI4 | PHI1 | progeny | 2016A_P4-A3 | | C | C | C |
| 2016A_P4-A3_5 | PHI4 | PHI1 | progeny | 2016A_P4-A3 | | C | C | C |
| 2016A_P4-B4 | PHI4 | PHI1 | mother | 2016A_P4-B4 | | C | C | C |
| 2016A_P4-B4_3 | PHI4 | PHI1 | progeny | 2016A_P4-B4 | | C | C | C |
| 2016A_P4-C11 | PHI4 | PHI1 | mother | 2016A_P4-C11 | | C | C | C |
| 2016A_P4-C11_3 | PHI4 | PHI1 | progeny | 2016A_P4-C11 | | C | C | C |
| 2016A_P4-C2 | PHI4 | PHI1 | mother | 2016A_P4-C2 | | C | C | C |
| 2016A_P4-C2_2 | PHI4 | PHI1 | progeny | 2016A_P4-C2 | | C | C | C |
| 2016A_P4-C2_3 | PHI4 | PHI1 | progeny | 2016A_P4-C2 | | C | C | C |
| 2016A_P4-C2_5 | PHI4 | PHI1 | progeny | 2016A_P4-C2 | | C | C | C |
| 2016A_P4-C2_6 | PHI4 | PHI1 | progeny | 2016A_P4-C2 | | C | C | C |
| 2016A_P4-C4 | PHI4 | PHI1 | mother | 2016A_P4-C4 | | C | C | C |
| 2016A_P4-C4_9 | PHI4 | PHI1 | progeny | 2016A_P4-C4 | | C | C | C |
| 2016A_P4-C9 | PHI4 | PHI1 | mother | 2016A_P4-C9 | | C | C | C |
| 2016A_P4-C9_1 | PHI4 | PHI1 | progeny | 2016A_P4-C9 | | C | C | C |
| 2016A_P4-C9_3 | PHI4 | PHI1 | progeny | 2016A_P4-C9 | | C | C | C |
| 2016A_P4-D12 | PHI4 | PHI1 | mother | 2016A_P4-D12 | | C | C | C |
| 2016A_P4-D12_5 | PHI4 | PHI1 | progeny | 2016A_P4-D12 | | C | C | C |
| 2016A_P4-D5 | PHI4 | PHI1 | mother | 2016A_P4-D5 | | C | C | C |
| 2016A_P4-D5_1 | PHI4 | PHI1 | progeny | 2016A_P4-D5 | | C | C | C |
| 2016A_P4-F5 | PHI4 | PHI1 | mother | 2016A_P4-F5 | | C | C | C |
| 2016A_P4-F5_1 | PHI4 | PHI1 | progeny | 2016A_P4-F5 | | C | C | C |
| 2016A_P4-F5_2 | PHI4 | PHI1 | progeny | 2016A_P4-F5 | | C | C | C |
| 2016A_P4-F5_5 | PHI4 | PHI1 | progeny | 2016A_P4-F5 | | C | C | C |
| 2016A_P4-F5_6 | PHI4 | PHI1 | progeny | 2016A_P4-F5 | | C | C | C |
| 2016A_P4-F7 | PHI7 | PHI1 | mother | 2016A_P4-F7 | | C | H | H |
| 2016A_P4-F7_1 | PHI7 | PHI1 | progeny | 2016A_P4-F7 | | C | C | C |
| 2016A_P4-G11 | PHI7 | PHI1 | mother | 2016A_P4-G11 | | C | H | H |
| 2016A_P4-G11_2 | PHI7 | PHI1 | progeny | 2016A_P4-G11 | | C | H | H |
| 2016A_P4-G11_4 | PHI7 | PHI1 | progeny | 2016A_P4-G11 | | C | H | H |
| 2016A_P4-H1 | PHI7 | PHI1 | mother | 2016A_P4-H1 | | C | H | H |
| 2016A_P4-H1_2 | PHI7 | PHI1 | progeny | 2016A_P4-H1 | | C | H | C |
| 2016A_P4-H1_3 | PHI7 | PHI1 | progeny | 2016A_P4-H1 | | C | H | H |
| 2016A_P4-H1_4 | PHI7 | PHI1 | progeny | 2016A_P4-H1 | | C | H | H |
| 2016A_P4-H10 | PHI7 | PHI1 | mother | 2016A_P4-H10 | | C | H | H |
| 2016A_P4-H10_4 | PHI7 | PHI1 | progeny | 2016A_P4-H10 | | C | H | H |
| 2016A_P4-H5 | PHI7 | PHI1 | mother | 2016A_P4-H5 | | C | H | H |
| 2016A_P4-H5_1 | PHI7 | PHI1 | progeny | 2016A_P4-H5 | | C | H | H |
| 2016A_P4-H5_2 | PHI7 | PHI1 | progeny | 2016A_P4-H5 | | C | T | H |
| 2016A_P5-A1 | PHI7 | PHI1 | mother | 2016A_P5-A1 | | C | H | H |
| 2016A_P5-A1_3 | PHI7 | PHI1 | progeny | 2016A_P5-A1 | | C | C | C |
| 2016A_P5-A1_4 | PHI7 | PHI1 | progeny | 2016A_P5-A1 | | C | C | C |
| 2016A_P5-B11 | PHI7 | PHI1 | mother | 2016A_P5-B11 | | C | H | H |
| 2016A_P5-B11_1 | PHI7 | PHI1 | progeny | 2016A_P5-B11 | | C | H | H |
| 2016A_P5-B11_7 | PHI7 | PHI1 | progeny | 2016A_P5-B11 | | C | H | H |
| 2016A_P5-C8 | PHI7 | PHI1 | mother | 2016A_P5-C8 | | C | H | H |
| 2016A_P5-C8_1 | PHI7 | PHI1 | progeny | 2016A_P5-C8 | | C | H | H |
| 2016A_P5-C8_2 | PHI7 | PHI1 | progeny | 2016A_P5-C8 | | C | H | G |
| 2016A_P5-F1 | PHI6 | PHI1 | mother | 2016A_P5-F1 | | H | H | H |
| 2016A_P5-F1_1 | PHI6 | PHI1 | progeny | 2016A_P5-F1 | | H | T | G |
| 2016A_P5-F10 | PHI6 | PHI1 | mother | 2016A_P5-F10 | | H | H | H |
| 2016A_P5-F10_2 | PHI6 | PHI1 | progeny | 2016A_P5-F10 | | C | C | H |
| 2016A_P5-F10_4 | PHI6 | PHI1 | progeny | 2016A_P5-F10 | | C | H | H |
| 2016A_P5-F10_7 | PHI6 | PHI1 | progeny | 2016A_P5-F10 | | H | H | H |
| 2016A_P5-F2 | PHI6 | PHI1 | mother | 2016A_P5-F2 | | H | H | H |
| 2016A_P5-F2_3 | PHI6 | PHI1 | progeny | 2016A_P5-F2 | | T | H | H |
| 2016A_P5-G10 | PHI6 | PHI1 | mother | 2016A_P5-G10 | | H | H | H |
| 2016A_P5-G10_10 | PHI6 | PHI1 | progeny | 2016A_P5-G10 | | H | H | H |
| 2016A_P5-G10_11 | PHI6 | PHI1 | progeny | 2016A_P5-G10 | | H | H | H |
| 2016A_P5-G10_6 | PHI6 | PHI1 | progeny | 2016A_P5-G10 | | H | H | H |
| 2016A_P5-G10_7 | PHI6 | PHI1 | progeny | 2016A_P5-G10 | | H | H | H |
| 2016A_P5-G10_8 | PHI6 | PHI1 | progeny | 2016A_P5-G10 | | H | H | G |
| 2016A_P5-G10_9 | PHI6 | PHI1 | progeny | 2016A_P5-G10 | | H | H | H |
| 2016A_P5-G12 | PHI6 | PHI1 | mother | 2016A_P5-G12 | | H | H | H |
| 2016A_P5-G12_1 | PHI6 | PHI1 | progeny | 2016A_P5-G12 | | C | C | C |
| 2016A_P5-G8 | PHI6 | PHI1 | mother | 2016A_P5-G8 | | H | H | H |

| Chr | 1 | 1 | 1 | 1 | 1 | 1 | 1 | 1 | 1 | 1 | 1 | 1 | 1 | 2 | 2 | 2 | 2 | 2 | 2 | 2 | 2 | 2 | 2 | 3 | 3 | 3 | 3 | 3 | 3 | 3 | 3 | 3 | 3 | 3 | 4 | 4 | 4 | 4 | 4 | 5 | 5 | 5 | 6 | 6 | 6 | 7 | 7 | 7 | 7 | 7 | 7 | 7 | 7 | 7 | 7 | 8 | 8 | 8 | 8 | 8 | 8 | 8 | 8 | 8 | 8 | 8 | 8 | 8 | 8 | 8 | 8 | 8 | 8 | 9 | 9 | 9 | 9 | 9 | 9 | 9 | 9 | 9 | 9 | 9 | 9 | 9 | 9 | 9 | 10 | 10 | 10 | 10 | 10 |
|---|---|---|---|---|---|---|---|---|---|---|---|---|---|---|---|---|---|---|---|---|---|---|---|---|---|---|---|---|---|---|---|---|---|---|---|---|---|---|---|---|---|---|---|---|---|---|---|---|---|---|---|---|---|---|---|---|---|---|---|---|---|---|---|---|---|---|---|---|---|---|---|---|---|---|---|---|---|---|---|---|---|---|---|---|---|---|---|---|---|---|---|---|---|
| Position, CM | 159 | 162 | 170 | 185 | 225 | 233 | 257 | 261 | 273 | 291 | 304 | 310 | 314 | 8.2 | 57 | 97 | 154 | 202 | 210 | 220 | 229 | 247 | 253 | 17 | 20 | 50 | 98 | 114 | 148 | 152 | 154 | 182 | 195 | 245 | 29 | 99 | 158 | 201 | 240 | 24 | 95 | 130 | 6.2 | 26 | 163 | 8 | 37 | 87 | 146 | 173 | 182 | 195 | 200 | 205 | 211 | 67 | 83 | 84 | 85 | 85 | 88 | 91 | 95 | 97 | 105 | 108 | 109 | 112 | 113 | 119 | 125 | 152 | 192 | 12 | 14 | 18 | 35 | 39 | 45 | 77 | 100 | 147 | 152 | 164 | 176 | 179 | 181 | 181 | 13 | 31 | 60 | 112 | 154 |
| 2016A_P1-A3 | T | H | C | H | C | G | H | T | A | C | A | T | T | T | G | H | H | T | H | H | H | A | H | T | T | H | A | T | H | H | A | H | A | A | H | H | H | T | H | H | H | G | H | H | H | H | G | H | H | T | C | H | H | H | H | A | H | A | H | A | T | G | G | H | A | H | H | H | G | A | H | H | H | H | H | H | H | H | A | H | G | H | H | A | G | G | C | T | H | H | H | A | H |
| 2016A_P1-A3_1 | T | H | C | H | C | G | H | T | A | C | A | T | T | T | G | H | H | T | H | H | H | A | A | T | T | G | A | T | A | A | A | C | A | A | H | H | H | T | H | H | H | G | H | H | H | H | G | H | H | T | C | H | H | H | H | A | H | A | H | A | T | G | G | H | A | H | H | H | G | A | H | H | H | H | H | H | H | H | A | H | G | H | H | A | G | G | C | T | H | H | H | A | H |
| 2016A_P1-D5 | T | H | C | H | C | G | H | T | A | C | A | T | T | T | G | H | H | T | H | H | H | A | H | T | T | H | A | T | H | H | A | H | A | A | H | H | H | T | H | H | H | H | H | H | H | H | G | H | H | T | C | H | H | H | H | A | H | A | H | A | T | G | G | H | A | H | H | H | G | A | H | H | H | H | H | H | H | H | A | H | G | H | H | A | G | G | C | T | H | H | H | A | H |
| 2016A_P1-D5_1 | T | H | C | H | C | G | H | T | A | C | A | T | T | T | G | A | C | T | C | C | T | A | G | T | T | H | A | T | H | H | A | H | A | A | H | H | H | T | H | H | H | H | C | G | G | H | G | H | H | T | C | H | H | H | H | A | H | A | H | A | T | G | G | H | A | H | H | H | G | A | H | H | H | H | H | H | H | H | A | H | G | H | H | A | G | G | C | T | H | H | H | A | H |
| 2016A_P1-D8 | T | H | C | H | C | G | H | T | A | C | A | T | T | T | G | H | H | T | H | H | H | A | H | T | T | H | A | H | H | H | A | H | A | H | H | H | H | T | H | H | H | H | H | H | H | H | G | H | H | T | C | H | A | H | H | H | H | H | H | H | H | H | H | H | A | H | H | H | G | A | H | H | H | H | H | H | H | H | A | H | G | H | H | A | H | H | H | H | H | H | H | A | H |
| 2016A_P1-D8_3 | T | T | C | T | C | G | T | T | A | C | A | T | T | T | G | A | C | T | H | H | H | A | H | T | T | H | A | H | H | H | A | H | A | H | H | H | H | T | H | A | C | G | H | G | H | C | G | T | H | T | C | H | A | H | H | C | C | C | G | G | G | A | T | A | A | C | C | A | G | A | G | T | C | T | T | G | H | A | A | A | G | T | T | A | A | A | T | C | H | H | G | A | C |
| 2016A_P1-E3 | T | H | C | H | C | G | H | T | A | C | A | T | T | T | G | H | H | T | H | H | H | A | H | T | T | H | A | H | H | H | A | H | A | H | H | H | H | T | H | H | H | H | H | H | H | H | G | H | H | T | C | H | H | H | H | A | H | A | H | A | T | G | G | H | H | H | T | H | G | A | H | H | H | H | H | H | H | H | H | H | G | H | H | A | H | H | H | T | H | H | H | A | H |
| 2016A_P1-E3_3 | T | T | C | T | C | G | H | T | A | C | A | T | T | T | G | H | T | T | T | H | H | A | H | T | T | G | A | T | H | H | A | H | A | H | H | G | H | T | H | A | C | G | A | A | H | H | G | T | H | T | C | H | H | H | H | A | H | A | H | A | T | G | G | H | H | H | T | H | G | A | H | H | H | H | H | H | H | H | H | A | G | H | H | A | H | H | H | T | H | G | A | H | G |
| 2016A_P1-E3_5 | T | H | C | H | C | G | C | T | A | C | A | H | T | H | H | H | H | T | H | H | H | A | H | T | T | H | A | H | H | H | H | H | H | H | H | H | H | H | H | H | H | H | H | H | H | H | H | H | H | T | C | H | H | C | H | H | H | H | H | A | H | H | H | H | H | H | H | H | G | A | H | H | C | H | H | H | H | H | H | H | G | H | H | A | H | H | H | T | H | H | H | A | H |
| 2016A_P1-E9 | T | H | C | H | C | G | H | T | A | C | A | T | T | T | G | H | H | T | H | H | H | A | H | T | T | H | A | T | H | A | H | H | A | A | H | H | H | T | H | H | H | H | H | H | H | H | G | H | H | T | C | H | H | H | H | A | H | A | H | A | T | G | G | H | A | H | H | H | G | A | H | H | H | H | H | H | H | H | A | H | G | H | H | A | G | G | C | T | H | H | H | A | H |
| 2016A_P1-E9_1 | T | T | C | T | C | G | H | T | A | C | A | T | T | T | G | G | T | T | C | C | T | A | G | T | T | C | A | [illegible] | [illegible] | [illegible] | [illegible] | [illegible] | [illegible] | [illegible] | [illegible] | [illegible] | [illegible] | [illegible] | [illegible] | [illegible] | [illegible] | [illegible] | [illegible] | [illegible] | [illegible] | [illegible] | [illegible] | [illegible] | [illegible] | [illegible] | [illegible] | [illegible] | [illegible] | [illegible] | [illegible] | [illegible] | [illegible] | [illegible] | [illegible] | [illegible] | [illegible] | [illegible] | [illegible] | [illegible] | [illegible] | [illegible] | [illegible] | [illegible] | [illegible] | [illegible] | [illegible] | [illegible] | [illegible] | [illegible] | [illegible] | [illegible] | [illegible] | [illegible] | [illegible] | [illegible] | [illegible] | [illegible] | [illegible] | [illegible] | [illegible] | [illegible] | [illegible] | [illegible] | [illegible] | [illegible] | [illegible] | [illegible] | [illegible] |
| 2016A_P1-F7 | T | H | C | H | C | G | H | T | A | C | A | T | T | T | G | H | H | T | H | H | H | A | H | T | T | H | A | [illegible] | [illegible] | [illegible] | [illegible] | [illegible] | [illegible] | [illegible] | [illegible] | [illegible] | [illegible] | [illegible] | [illegible] | [illegible] | [illegible] | [illegible] | [illegible] | [illegible] | [illegible] | [illegible] | [illegible] | [illegible] | [illegible] | [illegible] | [illegible] | [illegible] | [illegible] | [illegible] | [illegible] | [illegible] | [illegible] | [illegible] | [illegible] | [illegible] | [illegible] | [illegible] | [illegible] | [illegible] | [illegible] | [illegible] | [illegible] | [illegible] | [illegible] | [illegible] | [illegible] | [illegible] | [illegible] | [illegible] | [illegible] | [illegible] | [illegible] | [illegible] | [illegible] | [illegible] | [illegible] | [illegible] | [illegible] | [illegible] | [illegible] | [illegible] | [illegible] | [illegible] | [illegible] | [illegible] | [illegible] | [illegible] | [illegible] |
| 2016A_P1-F7_4 | T | T | C | T | C | G | T | T | A | C | A | T | T | T | G | H | H | T | H | H | H | A | H | T | T | C | A | [illegible] | [illegible] | [illegible] | [illegible] | [illegible] | [illegible] | [illegible] | [illegible] | [illegible] | [illegible] | [illegible] | [illegible] | [illegible] | [illegible] | [illegible] | [illegible] | [illegible] | [illegible] | [illegible] | [illegible] | [illegible] | [illegible] | [illegible] | [illegible] | [illegible] | [illegible] | [illegible] | [illegible] | [illegible] | [illegible] | [illegible] | [illegible] | [illegible] | [illegible] | [illegible] | [illegible] | [illegible] | [illegible] | [illegible] | [illegible] | [illegible] | [illegible] | [illegible] | [illegible] | [illegible] | [illegible] | [illegible] | [illegible] | [illegible] | [illegible] | [illegible] | [illegible] | [illegible] | [illegible] | [illegible] | [illegible] | [illegible] | [illegible] | [illegible] | [illegible] | [illegible] | [illegible] | [illegible] | [illegible] | [illegible] | [illegible] |

### Table S6. Results of the SNP array analasis (continue)

| | | | | | | | | | | | | | | | | | | | | | | | | | | | | | | | | | | | | | | | | | | | | | | | | | | | | | | | | | | | | | | | | | | | | | | | | | | | | | | | | | | | | | | | | | | | | | | | | | | | | | |
|---|---|---|---|---|---|---|---|---|---|---|---|---|---|---|---|---|---|---|---|---|---|---|---|---|---|---|---|---|---|---|---|---|---|---|---|---|---|---|---|---|---|---|---|---|---|---|---|---|---|---|---|---|---|---|---|---|---|---|---|---|---|---|---|---|---|---|---|---|---|---|---|---|---|---|---|---|---|---|---|---|---|---|---|---|---|---|---|---|---|---|---|---|---|---|---|---|---|---|---|---|---|
| | | | | | Chr | 1 | 1 | 1 | 1 | 1 | 1 | 1 | 1 | 1 | 1 | 1 | 1 | 1 | 1 | 1 | 1 | 2 | 2 | 2 | 2 | 2 | 2 | 2 | 2 | 2 | 2 | 3 | 3 | 3 | 3 | 3 | 3 | 3 | 3 | 3 | 3 | 3 | 4 | 4 | 4 | 4 | 4 | 5 | 5 | 5 | 6 | 6 | 6 | 7 | 7 | 7 | 7 | 7 | 7 | 7 | 7 | 7 | 7 | 8 | 8 | 8 | 8 | 8 | 8 | 8 | 8 | 8 | 8 | 8 | 8 | 8 | 8 | 8 | 8 | 8 | 8 | 9 | 9 | 9 | 9 | 9 | 9 | 9 | 9 | 9 | 9 | 9 | 9 | 9 | 9 | 9 | 10 | 10 | 10 | 10 | 10 |
| | | | | | Position, CM | 82 | 105 | 156 | 159 | 162 | 170 | 185 | 225 | 233 | 257 | 261 | 273 | 291 | 304 | 310 | 314 | 8.2 | 57 | 97 | 154 | 202 | 210 | 220 | 229 | 247 | 253 | 17 | 20 | 50 | 98 | 114 | 148 | 152 | 154 | 182 | 195 | 245 | 29 | 99 | 158 | 201 | 240 | 24 | 95 | 130 | 6.2 | 26 | 163 | 8 | 37 | 87 | 146 | 173 | 182 | 195 | 200 | 205 | 211 | 67 | 83 | 84 | 85 | 85 | 88 | 91 | 95 | 97 | 105 | 108 | 109 | 112 | 113 | 119 | 125 | 152 | 192 | 12 | 14 | 18 | 35 | 39 | 45 | 77 | 100 | 147 | 152 | 164 | 176 | 179 | 181 | 181 | 13 | 31 | 60 | 112 | 154 |
| plant_id | mom_gen | dad_gen | mother_pr | family_ | | | | | | | | | | | | | | | | | | | | | | | | | | | | | | | | | | | | | | | | | | | | | | | | | | | | | | | | | | | | | | | | | | | | | | | | | | | | | | | | | | | | | | | | | | | | | | | | | |
| 2016A_P5-G8_3 | PHI6 | PHI1 | progeny | 2016A_P5-G8 | | H | C | C | T | T | C | T | C | G | H | H | A | H | A | T | H | H | G | A | C | T | H | C | H | A | G | G | T | G | A | C | H | A | H | H | A | A | H | G | H | H | T | H | C | G | H | H | G | T | C | C | H | T | C | G | G | T | G | H | C | C | G | G | G | G | G | A | G | C | T | G | G | G | G | H | H | H | H | H | H | A | A | G | G | H | H | H | H | H | H | H | G | H | A | H | H |
| 2016A_P5-G8_4 | PHI6 | PHI1 | progeny | 2016A_P5-G8 | | H | H | H | T | H | C | H | C | G | H | H | A | H | A | T | H | H | G | A | H | T | H | C | H | A | G | H | T | C | A | T | H | A | H | H | A | A | H | H | A | H | T | H | C | H | H | A | G | T | G | H | A | T | C | H | H | T | G | H | H | H | H | H | H | G | G | H | H | H | T | G | G | H | H | H | H | H | H | H | H | A | A | G | H | C | C | A | G | G | C | T | H | H | H | A | G |
| 2016A_P5-G8_5 | PHI6 | PHI1 | progeny | 2016A_P5-G8 | | T | T | H | T | H | C | H | C | G | H | H | A | H | A | T | H | H | G | A | H | T | H | C | H | A | G | H | T | G | A | C | G | A | A | C | A | A | G | G | A | H | T | H | C | G | C | G | G | T | H | T | H | T | C | H | H | T | G | A | T | A | A | A | T | G | G | G | A | T | T | G | G | A | C | A | G | H | H | H | H | A | A | G | A | H | H | H | H | H | H | H | H | H | G | A | C |
| 2016A_P5-G9 | PHI6 | PHI1 | mother | 2016A_P5-G9 | | H | H | H | T | H | C | H | C | G | H | H | A | H | A | T | H | H | G | H | H | T | H | C | H | A | G | H | T | H | A | H | G | H | A | H | H | A | H | H | H | H | H | H | H | H | H | H | G | H | H | H | H | T | C | H | A | H | G | A | H | A | H | A | T | G | G | H | H | H | H | H | G | A | H | H | H | H | H | H | H | A | A | H | H | H | C | H | G | G | C | T | H | H | H | H | H |
| 2016A_P5-G9_1 | PHI6 | PHI1 | progeny | 2016A_P5-G9 | | H | T | G | T | C | C | C | C | G | H | H | A | H | A | T | H | H | G | G | T | T | H | C | H | A | G | T | T | C | A | T | G | G | A | A | H | A | H | A | H | H | H | H | C | C | C | G | G | H | H | C | H | T | C | H | A | H | G | A | T | A | A | A | T | G | G | G | A | T | T | G | G | A | C | H | H | T | T | G | H | A | A | A | G | H | C | H | G | G | C | T | H | H | A | H | H |
| 2016A_P5-G9_5 | PHI6 | PHI1 | progeny | 2016A_P5-G9 | | H | H | G | T | C | C | C | C | G | H | H | A | H | A | T | H | H | G | H | H | T | C | C | T | A | G | H | T | H | A | C | G | A | A | C | G | A | H | G | A | C | T | G | A | C | H | G | G | H | H | C | A | T | C | H | A | C | G | A | T | A | A | A | T | G | G | G | A | T | T | G | G | A | C | A | H | H | H | H | H | A | A | G | A | H | C | H | G | G | C | T | H | C | G | H | H |
| 2016A_P5-G9_6 | PHI6 | PHI1 | progeny | 2016A_P5-G9 | | T | T | G | T | C | C | C | C | G | C | C | A | T | A | T | H | H | G | A | C | T | C | C | H | A | G | H | T | H | A | H | G | G | A | A | A | A | H | G | A | C | T | H | A | C | H | G | G | H | G | T | G | T | C | H | A | H | G | A | T | A | A | A | T | G | G | G | A | T | T | G | G | A | C | H | C | H | H | H | H | A | A | A | G | H | C | H | G | G | C | T | C | C | G | H | H |
| 2016A_P5-H10 | PHI6 | PHI1 | mother | 2016A_P5-H10 | | H | H | H | T | H | C | H | C | G | H | H | A | H | A | T | H | H | G | H | H | T | H | C | H | A | G | H | H | H | H | T | G | H | A | H | A | A | H | H | G | H | H | H | H | G | H | H | G | H | H | H | H | H | C | H | H | T | G | A | H | A | H | A | T | G | G | H | H | H | T | G | G | H | H | H | H | H | H | H | H | A | A | H | H | H | H | H | H | H | H | H | H | H | H | H | H |
| 2016A_P5-H10_2 | PHI6 | PHI1 | progeny | 2016A_P5-H10 | | H | H | H | T | H | C | H | C | G | H | H | A | H | A | T | H | H | G | H | H | T | H | C | H | A | G | H | H | H | H | T | G | H | A | H | A | A | H | H | G | H | H | H | H | G | H | H | G | H | H | H | H | H | C | H | H | T | G | A | H | A | H | A | T | G | G | H | H | H | T | G | G | H | H | H | H | H | H | H | H | A | A | H | H | H | H | H | H | H | H | H | H | H | H | H | H |
| 2016A_P5-H10_3 | PHI6 | PHI1 | progeny | 2016A_P5-H10 | | T | T | G | T | C | C | C | C | G | H | H | A | H | A | T | H | H | G | A | C | T | C | C | T | A | G | H | H | H | H | T | G | H | A | H | A | A | G | G | G | C | H | A | C | G | H | A | G | H | G | T | H | H | C | T | A | T | G | A | T | A | A | A | T | G | G | G | A | T | T | G | G | A | C | H | H | H | H | H | H | A | A | G | A | H | H | H | H | H | H | H | H | C | G | A | H |
| 2016A_P5-H10_5 | PHI6 | PHI1 | progeny | 2016A_P5-H10 | | H | H | H | T | H | C | H | C | G | H | H | A | H | A | T | H | H | G | H | H | T | H | C | H | A | G | H | H | H | H | T | G | H | A | H | A | A | H | H | G | H | H | H | H | G | H | H | G | H | H | H | H | H | C | H | H | T | G | A | H | A | H | A | T | G | G | H | H | H | T | G | G | H | H | H | H | H | H | H | H | A | A | H | H | H | H | H | H | H | H | H | H | H | H | H | H |
| 2016A_P5-H11 | PHI6 | PHI1 | mother | 2016A_P5-H11 | | H | H | H | T | H | C | H | C | G | H | H | A | H | A | T | H | H | G | H | H | T | H | C | H | A | H | H | H | H | H | H | H | H | A | H | H | A | H | H | H | H | T | H | H | H | H | H | G | T | H | H | H | T | C | H | H | T | G | A | H | A | H | A | T | G | G | H | H | H | T | G | G | H | H | H | H | H | H | H | H | A | A | H | H | H | H | H | H | H | H | H | H | H | H | H | H |
| 2016A_P5-H11_2 | PHI6 | PHI1 | progeny | 2016A_P5-H11 | | H | T | G | T | C | C | C | C | G | H | H | A | H | A | T | H | H | G | A | C | T | H | C | H | A | H | G | A | G | H | H | G | A | A | C | H | A | H | G | H | H | T | H | C | G | H | G | G | T | H | T | G | T | C | T | A | T | G | A | C | A | G | A | T | G | G | H | G | C | T | G | G | G | G | T | C | H | H | H | H | A | A | G | A | H | H | H | H | H | H | H | H | C | G | A | H |
| 2016A_P5-H11_3 | PHI6 | PHI1 | progeny | 2016A_P5-H11 | | H | T | G | T | C | C | C | C | G | H | H | A | H | A | T | H | H | G | H | H | T | H | C | H | A | H | H | H | C | A | C | A | A | G | H | H | A | H | A | G | H | T | H | A | G | C | H | G | T | H | C | H | T | C | H | H | T | G | A | T | A | A | A | T | G | G | G | A | T | T | G | G | A | C | H | H | T | T | G | H | A | A | A | G | H | H | H | H | H | H | H | H | H | H | A | C |
| 2016A_P5-H11_5 | PHI6 | PHI1 | progeny | 2016A_P5-H11 | | H | H | H | T | H | C | H | C | G | H | H | A | H | A | T | H | H | G | H | H | T | H | C | C | A | A | H | H | H | H | H | H | A | H | H | H | A | H | H | H | H | T | H | H | G | H | H | G | T | H | H | H | T | C | H | H | T | G | A | H | A | H | A | T | G | G | H | H | H | T | G | G | H | H | C | H | H | H | H | H | A | A | A | G | H | H | H | H | H | H | H | H | H | H | H | H |
| 2016A_P5-H11_6 | PHI6 | PHI1 | progeny | 2016A_P5-H11 | | H | C | C | T | H | C | T | C | H | H | H | H | H | A | H | H | C | H | G | H | H | H | C | H | A | H | H | H | H | C | H | G | A | H | C | G | A | H | G | H | C | T | H | H | G | C | H | G | T | H | C | H | T | C | H | H | H | G | H | H | H | G | A | H | H | H | H | G | C | H | G | G | H | G | A | H | H | H | H | H | H | A | A | H | H | C | A | H | H | H | H | H | H | A | H | H |
| 2016A_P5-H12_11 | PHI6 | PHI1 | progeny | 2016A_P5-H12 | | H | T | G | T | C | C | C | C | G | H | T | A | H | A | T | H | C | G | G | H | T | H | C | H | A | G | T | T | H | A | C | H | A | H | H | H | A | C | A | H | H | T | H | H | H | H | H | G | T | H | H | H | T | C | G | A | C | G | A | H | A | H | A | T | G | G | H | A | H | H | H | G | A | H | H | H | H | H | H | H | A | A | H | H | H | C | H | G | G | C | T | H | H | H | A | H |
| 2016A_P5-H12_14 | PHI6 | PHI1 | progeny | 2016A_P5-H12 | | H | H | G | T | C | C | H | C | G | H | H | A | H | A | T | H | T | G | H | H | T | H | C | H | A | G | H | T | H | A | C | G | A | A | H | H | A | H | H | H | H | T | G | A | C | H | G | G | T | C | C | H | T | C | T | A | T | G | A | T | A | A | A | T | G | G | G | A | T | T | G | G | A | C | T | C | H | H | H | H | A | A | A | H | H | C | H | G | G | C | T | C | C | G | A | H |
| 2016A_P5-H12_16 | PHI6 | PHI1 | progeny | 2016A_P5-H12 | | H | H | C | [illegible] |
| 2016A_P5-H12_17 | PHI6 | PHI1 | progeny | 2016A_P5-H12 | | T | T | G | [illegible] |
| 2016A_P5-H12_3 | PHI6 | PHI1 | progeny | 2016A_P5-H12 | | H | C | C | [illegible] |
| 2016A_P5-H12_9 | PHI6 | PHI1 | progeny | 2016A_P5-H12 | | H | H | G | [illegible] |
| 2016A_P5-H3 | PHI6 | PHI1 | mother | 2016A_P5-H3 | | H | H | H | [illegible] |
| 2016A_P5-H3_1 | PHI6 | PHI1 | progeny | 2016A_P5-H3 | | T | T | G | [illegible] |
| 2016A_P5-H3_11 | PHI6 | PHI1 | progeny | 2016A_P5-H3 | | H | H | H | [illegible] |
| 2016A_P5-H3_2 | PHI6 | PHI1 | progeny | 2016A_P5-H3 | | H | C | C | [illegible] |
| 2016A_P5-H3_3 | PHI6 | PHI1 | progeny | 2016A_P5-H3 | | H | H | G | [illegible] |
| 2016A_P5-H3_4 | PHI6 | PHI1 | progeny | 2016A_P5-H3 | | H | C | H | [illegible] |
| 2016A_P5-H3_5 | PHI6 | PHI1 | progeny | 2016A_P5-H3 | | H | H | H | [illegible] |
| 2016A_P5-H3_6 | PHI6 | PHI1 | progeny | 2016A_P5-H3 | | H | C | C | [illegible] |
| 2016A_P5-H3_9 | PHI6 | PHI1 | progeny | 2016A_P5-H3 | | H | H | C | [illegible] |
| 2016A_P5-H5 | PHI6 | PHI1 | mother | 2016A_P5-H5 | | H | H | H | [illegible] |
| 2016A_P5-H5_1 | PHI6 | PHI1 | progeny | 2016A_P5-H5 | | H | H | G | [illegible] |
| 2016A_P5-H5_10 | PHI6 | PHI1 | progeny | 2016A_P5-H5 | | H | H | H | [illegible] |
| 2016A_P5-H5_12 | PHI6 | PHI1 | progeny | 2016A_P5-H5 | | T | T | G | [illegible] |
| 2016A_P5-H5_3 | PHI6 | PHI1 | progeny | 2016A_P5-H5 | | C | H | H | [illegible] |
| 2016A_P5-H5_4 | PHI6 | PHI1 | progeny | 2016A_P5-H5 | | C | C | C | [illegible] |
| 2016A_P5-H5_5 | PHI6 | PHI1 | progeny | 2016A_P5-H5 | | C | C | C | [illegible] |
| 2016A_P5-H5_6 | PHI6 | PHI1 | progeny | 2016A_P5-H5 | | H | T | G | [illegible] |
| 2016A_P5-H5_7 | PHI6 | PHI1 | progeny | 2016A_P5-H5 | | H | H | H | [illegible] |
| 2016A_P5-H5_8 | PHI6 | PHI1 | progeny | 2016A_P5-H5 | | H | H | H | [illegible] |
| 2016A_P5-H6 | PHI6 | PHI1 | mother | 2016A_P5-H6 | | H | H | H | [illegible] |
| 2016A_P5-H6_1 | PHI6 | PHI1 | progeny | 2016A_P5-H6 | | C | C | C | [illegible] |
| 2016A_P5-H8 | PHI6 | PHI1 | mother | 2016A_P5-H8 | | H | H | H | [illegible] |
| 2016A_P5-H8_1 | PHI6 | PHI1 | progeny | 2016A_P5-H8 | | T | H | H | [illegible] |
| 2016A_P5-H8_10 | PHI6 | PHI1 | progeny | 2016A_P5-H8 | | H | H | H | [illegible] |
| 2016A_P5-H8_16 | PHI6 | PHI1 | progeny | 2016A_P5-H8 | | H | H | H | [illegible] |
| 2016A_P5-H8_4 | PHI6 | PHI1 | progeny | 2016A_P5-H8 | | H | H | H | [illegible] |
| 2016A_P5H12 | PHI6 | PHI1 | mother | 2016A_P5H12 | | H | H | H | [illegible] |
| 2016A_P6-A8 | PHI6 | PHI1 | mother | 2016A_P6-A8 | | H | H | H | [illegible] |
| 2016A_P6-A8_1 | PHI6 | PHI1 | progeny | 2016A_P6-A8 | | H | C | C | [illegible] |
| 2016A_P6-A8_11 | PHI6 | PHI1 | progeny | 2016A_P6-A8 | | H | H | H | [illegible] |
| 2016A_P6-A8_12 | PHI6 | PHI1 | progeny | 2016A_P6-A8 | | H | C | C | [illegible] |
| 2016A_P6-A8_16 | PHI6 | PHI1 | progeny | 2016A_P6-A8 | | H | H | C | [illegible] |
| 2016A_P6-A8_8 | PHI6 | PHI1 | progeny | 2016A_P6-A8 | | H | H | H | [illegible] |
| 2016A_P6-A9 | PHI6 | PHI1 | mother | 2016A_P6-A9 | | H | H | H | [illegible] |
| 2016A_P6-A9_1 | PHI6 | PHI1 | progeny | 2016A_P6-A9 | | H | H | H | [illegible] |
| 2016A_P6-A9_2 | PHI6 | PHI1 | progeny | 2016A_P6-A9 | | T | H | G | [illegible] |
| 2016A_P6-A9_4 | PHI6 | PHI1 | progeny | 2016A_P6-A9 | | H | H | H | [illegible] |
| 2016A_P6-A9_6 | PHI6 | PHI1 | progeny | 2016A_P6-A9 | | C | C | C | [illegible] |
| 2016A_P6-A9_8 | PHI6 | PHI1 | progeny | 2016A_P6-A9 | | T | H | C | [illegible] |
| 2016A_P6-B10 | PHI6 | PHI1 | mother | 2016A_P6-B10 | | H | H | H | [illegible] |
| 2016A_P6-B10_11 | PHI6 | PHI1 | progeny | 2016A_P6-B10 | | H | C | H | [illegible] |
| 2016A_P6-B10_12 | PHI6 | PHI1 | progeny | 2016A_P6-B10 | | H | C | C | [illegible] |
| 2016A_P6-B10_13 | PHI6 | PHI1 | progeny | 2016A_P6-B10 | | C | C | C | [illegible] |
| 2016A_P6-B10_15 | PHI6 | PHI1 | progeny | 2016A_P6-B10 | | H | H | H | [illegible] |
| 2016A_P6-B10_4 | PHI6 | PHI1 | progeny | 2016A_P6-B10 | | H | H | H | [illegible] |
| 2016A_P6-B10_8 | PHI6 | PHI1 | progeny | 2016A_P6-B10 | | H | H | H | [illegible] |
| 2016A_P6-B12 | PHI6 | PHI1 | mother | 2016A_P6-B12 | | H | H | H | [illegible] |
| 2016A_P6-B12_4 | PHI6 | PHI1 | progeny | 2016A_P6-B12 | | H | H | H | [illegible] |
| 2016A_P6-B12_7 | PHI6 | PHI1 | progeny | 2016A_P6-B12 | | H | C | C | [illegible] |
| 2016A_P6-B7 | PHI6 | PHI1 | mother | 2016A_P6-B7 | | H | H | H | [illegible] |
| 2016A_P6-B7_1 | PHI6 | PHI1 | progeny | 2016A_P6-B7 | | H | C | C | [illegible] |
| 2016A_P6-B7_2 | PHI6 | PHI1 | progeny | 2016A_P6-B7 | | T | T | H | [illegible] |
| 2016A_P6-B8 | PHI6 | PHI1 | mother | 2016A_P6-B8 | | H | H | H | [illegible] |
| 2016A_P6-B8_2 | PHI6 | PHI1 | progeny | 2016A_P6-B8 | | H | H | H | [illegible] |
| 2016A_P6-B8_3 | PHI6 | PHI1 | progeny | 2016A_P6-B8 | | T | H | H | [illegible] |
| 2016A_P6-C10 | PHI6 | PHI1 | mother | 2016A_P6-C10 | | H | H | H | [illegible] |
| 2016A_P6-C10_1 | PHI6 | PHI1 | progeny | 2016A_P6-C10 | | H | C | H | [illegible] |
| 2016A_P6-C10_2 | PHI6 | PHI1 | progeny | 2016A_P6-C10 | | C | C | C | [illegible] |
| 2016A_P6-C10_3 | PHI6 | PHI1 | progeny | 2016A_P6-C10 | | H | H | C | [illegible] |
| 2016A_P6-C12 | PHI6 | PHI1 | mother | 2016A_P6-C12 | | H | H | H | [illegible] |
| 2016A_P6-C12_1 | PHI6 | PHI1 | progeny | 2016A_P6-C12 | | H | H | H | [illegible] |
| 2016A_P6-C12_2 | PHI6 | PHI1 | progeny | 2016A_P6-C12 | | H | H | H | [illegible] |
| 2016A_P6-C12_3 | PHI6 | PHI1 | progeny | 2016A_P6-C12 | | H | H | H | [illegible] |
| 2016A_P6-C12_4 | PHI6 | PHI1 | progeny | 2016A_P6-C12 | | H | H | H | [illegible] |
| 2016A_P6-D11 | PHI6 | PHI1 | mother | 2016A_P6-D11 | | H | H | H | [illegible] |
| 2016A_P6-D11_1 | PHI6 | PHI1 | progeny | 2016A_P6-D11 | | H | H | C | [illegible] |
| 2016A_P6-E5 | PHI8 | PHI1 | mother | 2016A_P6-E5 | | H | H | H | [illegible] |
| 2016A_P6-E5_4 | PHI8 | PHI1 | progeny | 2016A_P6-E5 | | H | H | H | [illegible] |
| 2016A_P6-F7 | PHI8 | PHI1 | mother | 2016A_P6-F7 | | H | H | H | [illegible] |
| 2016A_P6-F7_12 | PHI8 | PHI1 | progeny | 2016A_P6-F7 | | C | H | H | [illegible] |
| 2016A_P6-G9 | PHI8 | PHI1 | mother | 2016A_P6-G9 | | H | H | H | [illegible] |
| 2016A_P6-G9_4 | PHI8 | PHI1 | progeny | 2016A_P6-G9 | | H | H | H | [illegible] |
| 2016A_P6-H1 | PHI8 | PHI1 | mother | 2016A_P6-H1 | | H | H | H | [illegible] |
| 2016A_P6-H1_3 | PHI8 | PHI1 | progeny | 2016A_P6-H1 | | H | H | H | [illegible] |
| 2016A_P6-H3_3 | PHI8 | PHI1 | progeny | 2016A_P6-H3 | | T | H | H | [illegible] |
| 2016A_P7-A8 | PHI8 | PHI1 | mother | 2016A_P7-A8 | | H | H | H | [illegible] |
| 2016A_P7-A8_1 | PHI8 | PHI1 | progeny | 2016A_P7-A8 | | H | H | H | [illegible] |
| 2016A_P7-C11 | PHI5 | PHI1 | mother | 2016A_P7-C11 | | H | H | H | [illegible] |
| 2016A_P7-C11_1 | PHI5 | PHI1 | progeny | 2016A_P7-C11 | | T | T | G | [illegible] |
| 2016A_P7-D10 | PHI5 | PHI1 | mother | 2016A_P7-D10 | | H | H | H | [illegible] |
| 2016A_P7-D10_1 | PHI5 | PHI1 | progeny | 2016A_P7-D10 | | C | T | G | [illegible] |
| 2016A_P7-D2 | PHI5 | PHI1 | mother | 2016A_P7-D2 | | H | H | H | [illegible] |
| 2016A_P7-D2_1 | PHI5 | PHI1 | progeny | 2016A_P7-D2 | | C | C | C | [illegible] |
| 2016A_P7-D8 | PHI5 | PHI1 | mother | 2016A_P7-D8 | | H | H | H | [illegible] |
| 2016A_P7-D8_2 | PHI5 | PHI1 | progeny | 2016A_P7-D8 | | H | C | C | [illegible] |
| 2016A_P7-G3 | PHI5 | PHI1 | mother | 2016A_P7-G3 | | H | H | H | [illegible] |
| 2016A_P7-G3_1 | PHI5 | PHI1 | progeny | 2016A_P7-G3 | | H | H | H | [illegible] |
| 2016A_P7-G3_3 | PHI5 | PHI1 | progeny | 2016A_P7-G3 | | H | H | G | [illegible] |
| 2016A_P7-G3_4 | PHI5 | PHI1 | progeny | 2016A_P7-G3 | | H | H | G | [illegible] |
| 2016A_P7-G7 | PHI5 | PHI1 | mother | 2016A_P7-G7 | | H | H | H | [illegible] |
| 2016A_P7-G7_1 | PHI5 | PHI1 | progeny | 2016A_P7-G7 | | H | H | H | [illegible] |
| 2016A_P7-H9 | PHI5 | PHI1 | mother | 2016A_P7-H9 | | H | H | H | [illegible] |
| 2016A_P7-H9_1 | PHI5 | PHI1 | progeny | 2016A_P7-H9 | | H | H | H | [illegible] |
| 2016A_P7-H9_2 | PHI5 | PHI1 | progeny | 2016A_P7-H9 | | H | H | H | [illegible] |
| 2016A_P8-A1 | PHI5 | PHI1 | mother | 2016A_P8-A1 | | H | H | H | [illegible] |
| 2016A_P8-A1_1 | PHI5 | PHI1 | progeny | 2016A_P8-A1 | | H | H | H | [illegible] |
| 2016A_P8-A5 | PHI5 | PHI1 | mother | 2016A_P8-A5 | | H | H | H | [illegible] |
| 2016A_P8-A5_1 | PHI5 | PHI1 | progeny | 2016A_P8-A5 | | H | H | H | [illegible] |
| 2016A_P8-A6 | PHI5 | PHI1 | mother | 2016A_P8-A6 | | H | H | H | [illegible] |
| 2016A_P8-A6_1 | PHI5 | PHI1 | progeny | 2016A_P8-A6 | | C | C | C | [illegible] |
| 2016A_P8-D5 | PHI9 | PHI1 | mother | 2016A_P8-D5 | | C | C | C | [illegible] |
| 2016A_P8-D5_3 | PHI9 | PHI1 | progeny | 2016A_P8-D5 | | C | C | C | [illegible] |
| 2016A_P8-D5_5 | PHI9 | PHI1 | progeny | 2016A_P8-D5 | | C | C | C | [illegible] |
| 2016A_P8-E1 | PHI9 | PHI1 | mother | 2016A_P8-E1 | | C | C | C | [illegible] |
| 2016A_P8-E1_2 | PHI9 | PHI1 | progeny | 2016A_P8-E1 | | C | C | C | [illegible] |
| 2016A_P9-A10 | PHI9 | PHI1 | mother | 2016A_P9-A10 | | C | C | C | [illegible] |
| 2016A_P9-A10_1 | PHI9 | PHI1 | progeny | 2016A_P9-A10 | | C | C | C | [illegible] |
| 2016A_P9-A10_3 | PHI9 | PHI1 | progeny | 2016A_P9-A10 | | C | C | C | [illegible] |
| 2016A_P9-A7 | PHI9 | PHI1 | mother | 2016A_P9-A7 | | C | C | C | [illegible] |
| 2016A_P9-A7_1 | PHI9 | PHI1 | progeny | 2016A_P9-A7 | | C | C | C | [illegible] |
| 2016A_P9-A7_2 | PHI9 | PHI1 | progeny | 2016A_P9-A7 | | C | C | C | [illegible] |
| 2016A_P9-B1 | PHI9 | PHI1 | mother | 2016A_P9-B1 | | H | C | C | [illegible] |
| 2016A_P9-B1_2 | PHI9 | PHI1 | progeny | 2016A_P9-B1 | | T | C | C | [illegible] |
| 2016A_P9-B7 | PHI9 | PHI1 | mother | 2016A_P9-B7 | | C | C | C | [illegible] |
| 2016A_P9-B7_10 | PHI9 | PHI1 | progeny | 2016A_P9-B7 | | C | C | C | [illegible] |
| 2016A_P9-B7_5 | PHI9 | PHI1 | progeny | 2016A_P9-B7 | | C | C | C | [illegible] |
| 2016A_P9-B7_9 | PHI9 | PHI1 | progeny | 2016A_P9-B7 | | C | C | C | [illegible] |

plant_id – unique identifier of the plant
mom_gen – genetic backgroiund of mother in the hybred
dad_gen – genetic background of father in the hybred
family_ – identify mother and all the progeny plants from the same mother
mother_progeny – identify each plant as mother of the family or progeny
Chr – number of the chromosome, where marker is located
Distance – position of the marker on the corresponding chromosome in cM
H – heterozygous marker
A, T, G and C – marker homozygous for the corresponding nucleotide
EQV, LS, -9 , etc. – missing values

**Table S7. Results of the SNP array analasis**

| plant_id | Chr | 1 | 1 | 1 | 1 | 1 | 1 | 1 | 1 | 1 | 1 | 1 | 1 | 1 | 1 | 1 | 1 | 1 | 1 | 1 | 1 | 1 | 1 | 1 | 2 | 2 | 2 | 2 | 2 | 2 | 2 | 2 | 2 | 2 | 2 | 2 | 2 | 2 | 2 | 2 | 2 | 2 | 2 | 2 | 2 | 2 | 3 | 3 | 3 | 3 | 3 | 3 | 3 | 3 | 3 | 3 | 3 | 3 | 3 | 3 | 3 | 3 | 3 | 3 | 3 | 3 | 3 | 4 | 4 | 4 | 4 | 4 | 4 | 4 | 4 | 4 | 4 | 4 | 4 | 4 | 4 | 4 | 4 | 4 | 4 | 4 | 4 | 4 | 4 | 4 | 5 | 5 | 5 | 5 | 5 | 5 | 5 | 5 | 5 | 5 | 5 | 5 | 5 | 5 | 5 | 5 | 5 | 5 | 5 | 5 | 6 | 6 | 6 | 6 | 6 | 6 | 6 | 6 |
|---|---|---|---|---|---|---|---|---|---|---|---|---|---|---|---|---|---|---|---|---|---|---|---|---|---|---|---|---|---|---|---|---|---|---|---|---|---|---|---|---|---|---|---|---|---|---|---|---|---|---|---|---|---|---|---|---|---|---|---|---|---|---|---|---|---|---|---|---|---|---|---|---|---|---|---|---|---|---|---|---|---|---|---|---|---|---|---|---|---|---|---|---|---|---|---|---|---|---|---|---|---|---|---|---|---|---|---|---|---|---|---|---|---|---|---|---|---|---|
| | Position, cM | 2.6 | 4.5 | 16 | 39 | 59 | 80 | 84 | 108 | 125 | 137 | 137 | 137 | 154 | 164 | 185 | 192 | 203 | 220 | 246 | 273 | 281 | 300 | 316 | 7.7 | 25 | 35 | 54 | 55 | 78 | 94 | 113 | 129 | 129 | 129 | 130 | 130 | 160 | 178 | 183 | 198 | 213 | 220 | 231 | 235 | 253 | 5.9 | 10 | 28 | 50 | 64 | 77 | 79 | 78 | 79 | 88 | 92 | 98 | 109 | 135 | 160 | 165 | 195 | 215 | 227 | 235 | 246 | 7.4 | 18 | 29 | 38 | 57 | 63 | 67 | 89 | 110 | 111 | 110 | 115 | 129 | 133 | 157 | 166 | 173 | 181 | 193 | 201 | 207 | 210 | 246 | 5.3 | 13 | 20 | 30 | 53 | 78 | 89 | 94 | 101 | 102 | 102 | 102 | 108 | 119 | 138 | 145 | 165 | 195 | 217 | 227 | 5.4 | 14 | 23 | 24 | 23 | 29 | 43 | 54 |
| 2016H-P4-H4 | | C | H | C | A | H | H | G | C | G | H | H | H | C | A | H | T | A | G | H | H | H | A | A | A | A | T | G | H | C | T | T | G | H | H | H | H | T | A | G | A | C | T | G | H | T | H | G | H | H | A | H | H | H | H | H | H | H | T | H | T | H | H | A | A | H | T | H | H | H | H | H | H | H | H | H | H | A | H | H | G | H | T | H | H | H | H | H | H | H | G | A | H | H | H | H | H | H | H | H | H | H | H | G | T | H | H | H | H | H | H | H | T | A | T | H | H | H |
| 2016H-P5-D5 | | C | H | C | A | H | H | G | C | G | H | H | H | H | H | H | T | A | G | H | H | H | A | A | A | A | T | G | H | C | T | T | G | H | H | H | H | T | A | G | A | C | T | G | H | T | H | G | H | H | A | H | H | H | H | H | H | H | T | H | T | H | A | H | A | H | T | H | H | H | H | H | H | H | H | H | H | A | H | H | G | A | H | H | H | H | H | H | H | H | G | A | H | H | H | H | H | H | H | H | H | H | H | G | T | H | G | H | H | H | H | H | T | A | T | H | H | H |
| 2016H-P4-F3 | | C | H | C | A | H | H | G | C | G | H | H | H | H | H | H | T | A | G | G | H | H | A | A | A | A | T | G | H | C | T | T | G | H | H | H | H | T | A | G | A | C | T | G | H | T | H | G | H | H | A | H | H | H | H | H | H | H | T | A | T | H | H | A | A | H | T | H | H | H | H | H | H | H | H | H | A | H | C | H | H | H | T | H | H | H | H | H | H | H | G | A | H | H | H | H | H | H | H | H | H | H | H | G | T | H | H | H | H | H | H | H | T | A | T | H | H | H |
| 2016H-P5-C9 | | C | H | C | A | H | H | G | C | G | H | H | H | C | A | H | T | A | G | H | H | H | A | A | A | A | T | G | H | C | T | T | G | H | H | H | H | T | A | G | A | C | T | G | H | T | H | G | H | H | A | H | H | H | H | H | H | H | T | A | T | H | A | H | H | H | H | H | H | H | H | H | H | H | H | H | H | A | H | H | H | A | H | H | H | H | H | H | H | H | G | A | H | H | H | H | H | H | H | H | H | H | T | G | T | H | H | H | H | H | H | H | T | A | T | H | H | H |
| 2016H-P5-B5 | | C | H | C | A | H | H | G | C | G | H | H | H | C | A | H | T | A | G | H | H | H | A | A | A | A | T | G | H | C | T | T | G | H | H | H | H | T | A | G | A | C | T | G | H | T | H | G | H | H | A | H | H | H | H | H | H | H | T | H | T | H | A | H | H | H | H | H | H | H | H | H | H | H | H | H | A | H | C | H | G | A | H | H | H | H | H | H | H | H | G | A | H | H | H | H | H | H | H | H | H | H | T | G | T | H | G | H | C | H | H | H | T | A | T | H | H | H |
| 2016H-P4-H1 | | C | H | C | A | H | H | G | C | G | H | H | H | C | A | H | T | A | G | H | H | H | A | A | A | A | T | G | H | C | T | T | G | H | H | H | H | T | A | G | A | C | T | G | H | T | H | G | H | H | A | H | H | H | H | H | H | H | T | H | T | H | A | H | H | H | H | H | H | H | H | H | H | H | H | H | H | A | H | H | H | H | T | H | H | H | H | H | H | H | G | A | H | H | H | H | H | H | H | H | H | H | T | G | T | H | G | H | H | H | H | H | T | A | T | H | H | H |
| 2016H-P3-E5 | | C | G | C | A | H | A | G | H | H | A | A | C | C | A | C | H | H | H | T | A | H | A | H | A | A | T | G | T | H | H | H | G | A | C | C | C | H | H | G | A | C | H | H | H | H | H | G | H | G | H | H | H | H | H | H | H | H | T | H | T | H | H | A | A | H | T | H | C | C | C | T | H | T | H | H | H | H | H | H | G | A | H | T | G | H | H | H | H | H | H | H | H | A | H | H | H | H | H | H | H | H | H | G | H | H | G | A | T | T | T | A | T | H | H | H | H | G |
| 2016H-P3-E11 | | C | G | C | A | H | A | G | H | H | A | A | C | C | A | C | H | H | H | T | A | H | A | H | A | A | T | G | T | H | H | H | G | A | C | C | C | H | H | G | A | C | H | H | H | H | H | G | H | G | H | H | H | H | H | H | H | H | T | H | T | H | H | A | A | H | T | H | C | C | C | T | H | T | H | H | T | A | G | H | G | A | H | T | G | H | H | H | H | H | H | H | H | A | H | H | H | H | H | H | H | H | H | G | T | H | G | A | T | T | T | A | T | A | T | T | H | G |

| plant_id | mom_gen | dad_gen | mother_progeny | family_ |
|---|---|---|---|---|
| 2016H-P4-H4 | PHI5 | PHI3 | mother | 2016H-P4-H4 |
| 2016H-P5-D5 | PHI5 | PHI3 | mother | 2016H-P5-D5 |
| 2016H-P4-F3 | PHI5 | PHI3 | mother | 2016H-P4-F3 |
| 2016H-P5-C9 | PHI5 | PHI3 | mother | 2016H-P5-C9 |
| 2016H-P5-B5 | PHI5 | PHI3 | mother | 2016H-P5-B5 |
| 2016H-P4-H1 | PHI5 | PHI3 | mother | 2016H-P4-H1 |
| 2016H-P3-E5 | PHI6 | PHI3 | mother | 2016H-P3-E5 |
| 2016H-P3-E11 | PHI6 | PHI3 | mother | 2016H-P3-E11 |
| 2016H-P3-H10 | PHI6 | PHI3 | mother | 2016H-P3-H10 |
| 2016H-P3-E3 | PHI6 | PHI3 | mother | 2016H-P3-E3 |
| 2016H-P4-A9 | PHI6 | PHI3 | mother | 2016H-P4-A9 |
| 2016H-P3-H7 | PHI6 | PHI3 | mother | 2016H-P3-H7 |
| 2016H-P3-E4 | PHI6 | PHI3 | mother | 2016H-P3-E4 |
| 2016H-P3-H9 | PHI6 | PHI3 | mother | 2016H-P3-H9 |
| 2016H-P4-A6 | PHI6 | PHI3 | mother | 2016H-P4-A6 |
| 2016H-P3-G11 | PHI6 | PHI3 | mother | 2016H-P3-G11 |
| 2016H-P3-G12 | PHI6 | PHI3 | mother | 2016H-P3-G12 |
| 2016H-P3-F1 | PHI6 | PHI3 | mother | 2016H-P3-F1 |
| 2016H-P3-F11 | PHI6 | PHI3 | mother | 2016H-P3-F11 |
| 2016H-P3-F3 | PHI6 | PHI3 | mother | 2016H-P3-F3 |
| 2016H-P3-H8 | PHI6 | PHI3 | mother | 2016H-P3-H8 |
| 2016H-P3-G6 | PHI6 | PHI3 | mother | 2016H-P3-G6 |
| 2016H-P3-F6 | PHI6 | PHI3 | mother | 2016H-P3-F6 |
| 2016H-P3-G2 | PHI6 | PHI3 | mother | 2016H-P3-G2 |
| 2016H-P3-F8 | PHI6 | PHI3 | mother | 2016H-P3-F8 |
| 2016H-P3-G7 | PHI6 | PHI3 | mother | 2016H-P3-G7 |
| 2016H-P3-C4 | PHI7 | PHI3 | mother | 2016H-P3-C4 |
| 2016H-P3-A9 | PHI7 | PHI3 | mother | 2016H-P3-A9 |
| 2016H-P3-C2 | PHI7 | PHI3 | mother | 2016H-P3-C2 |
| 2016H-P2-H11 | PHI7 | PHI3 | mother | 2016H-P2-H11 |
| 2016H-P3-B2 | PHI7 | PHI3 | mother | 2016H-P3-B2 |
| 2016H-P3-C9 | PHI7 | PHI3 | mother | 2016H-P3-C9 |
| 2016H-P3-A8 | PHI7 | PHI3 | mother | 2016H-P3-A8 |
| 2016H-P3-A11 | PHI7 | PHI3 | mother | 2016H-P3-A11 |
| 2016H-P3-B9 | PHI7 | PHI3 | mother | 2016H-P3-B9 |
| 2016H-P3-A12 | PHI7 | PHI3 | mother | 2016H-P3-A12 |
| 2016H-P2-C9 | PHI4 | PHI3 | mother | 2016H-P2-C9 |
| 2016H-P2-B7 | PHI4 | PHI3 | mother | 2016H-P2-B7 |
| 2016H-P2-F3 | PHI4 | PHI3 | mother | 2016H-P2-F3 |
| 2016H-P2-G6 | PHI4 | PHI3 | mother | 2016H-P2-G6 |
| 2016H-P2-E7 | PHI4 | PHI3 | mother | 2016H-P2-E7 |
| 2016H-P2-A12 | PHI4 | PHI3 | mother | 2016H-P2-A12 |
| 2016H-P2-C1 | PHI4 | PHI3 | mother | 2016H-P2-C1 |
| 2016H-P2-E9 | PHI4 | PHI3 | mother | 2016H-P2-E9 |
| 2016H-P2-E8 | PHI4 | PHI3 | mother | 2016H-P2-E8 |
| 2016H-P2-F11 | PHI4 | PHI3 | mother | 2016H-P2-F11 |
| 2016H-P2-D6 | PHI4 | PHI3 | mother | 2016H-P2-D6 |
| 2016H-P2-D9 | PHI4 | PHI3 | mother | 2016H-P2-D9 |
| 2016H-P2-E1 | PHI4 | PHI3 | mother | 2016H-P2-E1 |
| 2016H-P2-D3 | PHI4 | PHI3 | mother | 2016H-P2-D3 |
| 2016H-P2-D2 | PHI4 | PHI3 | mother | 2016H-P2-D2 |
| 2016H-P2-G8 | PHI4 | PHI3 | mother | 2016H-P2-G8 |
| 2016H-P2-C7 | PHI4 | PHI3 | mother | 2016H-P2-C7 |
| 2016H-P2-C6 | PHI4 | PHI3 | mother | 2016H-P2-C6 |
| 2016H-P1-G1 | PHI10 | PHI3 | mother | 2016H-P1-G1 |
| 2016H-P1-F4 | PHI10 | PHI3 | mother | 2016H-P1-F4 |
| 2016H-P1-F5 | PHI10 | PHI3 | mother | 2016H-P1-F5 |
| 2016H-P1-F11 | PHI10 | PHI3 | mother | 2016H-P1-F11 |
| 2016H-P1-G9 | PHI10 | PHI3 | mother | 2016H-P1-G9 |
| 2016H-P1-G2 | PHI10 | PHI3 | mother | 2016H-P1-G2 |
| 2016H-P1-G5 | PHI10 | PHI3 | mother | 2016H-P1-G5 |
| 2016H-P1-F9 | PHI10 | PHI3 | mother | 2016H-P1-F9 |
| 2016H-P1-F3 | PHI10 | PHI3 | mother | 2016H-P1-F3 |
| 2016H-P1-F10 | PHI10 | PHI3 | mother | 2016H-P1-F10 |
| 2016H-P5-G3 | PHI9 | PHI3 | mother | 2016H-P5-G3 |
| 2016H-P5-E5 | PHI9 | PHI3 | mother | 2016H-P5-E5 |
| 2016H-P5-G4 | PHI9 | PHI3 | mother | 2016H-P5-G4 |
| 2016H-P5-F10 | PHI9 | PHI3 | mother | 2016H-P5-F10 |
| 2016H-P5-F2 | PHI9 | PHI3 | mother | 2016H-P5-F2 |
| 2016H-P5-E7 | PHI9 | PHI3 | mother | 2016H-P5-E7 |
| 2016H-P1-C9 | PHI2 | PHI3 | mother | 2016H-P1-C9 |
| 2016H-P1-C2 | PHI2 | PHI3 | mother | 2016H-P1-C2 |
| 2016H-P1-C6 | PHI2 | PHI3 | mother | 2016H-P1-C6 |
| 2016H-P1-A4 | PHI2 | PHI3 | mother | 2016H-P1-A4 |
| 2016H-P1-A9 | PHI2 | PHI3 | mother | 2016H-P1-A9 |
| 2016H-P1-A11 | PHI2 | PHI3 | mother | 2016H-P1-A11 |
| 2016H-P4-H4_2 | PHI5 | PHI3 | progeny | 2016H-P4-H4 |
| 2016H-P5-D5_1 | PHI5 | PHI3 | progeny | 2016H-P5-D5 |
| 2016H-P5-D5_2 | PHI5 | PHI3 | progeny | 2016H-P5-D5 |
| 2016H-P5-D5_3 | PHI5 | PHI3 | progeny | 2016H-P5-D5 |
| 2016H-P4-F3_1 | PHI5 | PHI3 | progeny | 2016H-P4-F3 |
| 2016H-P5-C9_1 | PHI5 | PHI3 | progeny | 2016H-P5-C9 |
| 2016H-P5-C9_2 | PHI5 | PHI3 | progeny | 2016H-P5-C9 |
| 2016H-P5-B6_1 | PHI5 | PHI3 | progeny | 2016H-P5-B6 |
| 2016H-P5-B6_2 | PHI5 | PHI3 | progeny | 2016H-P5-B6 |
| 2016H-P5-B5_1 | PHI5 | PHI3 | progeny | 2016H-P5-B5 |
| 2016H-P5-B5_2 | PHI5 | PHI3 | progeny | 2016H-P5-B5 |
| 2016H-P4-H1_1 | PHI5 | PHI3 | progeny | 2016H-P4-H1 |
| 2016H-P3-E5_1 | PHI6 | PHI3 | progeny | 2016H-P3-E5 |
| 2016H-P3-E11_1 | PHI6 | PHI3 | progeny | 2016H-P3-E11 |
| 2016H-P3-H10_1 | PHI6 | PHI3 | progeny | 2016H-P3-H10 |
| 2016H-P3-E3_2 | PHI6 | PHI3 | progeny | 2016H-P3-E3 |
| 2016H-P4-A9_1 | PHI6 | PHI3 | progeny | 2016H-P4-A9 |
| 2016H-P3-H7_1 | PHI6 | PHI3 | progeny | 2016H-P3-H7 |
| 2016H-P3-E4_3 | PHI6 | PHI3 | progeny | 2016H-P3-E4 |
| 2016H-P3-E4_6 | PHI6 | PHI3 | progeny | 2016H-P3-E4 |
| 2016H-P3-H9_1 | PHI6 | PHI3 | progeny | 2016H-P3-H9 |
| 2016H-P3-H9_2 | PHI6 | PHI3 | progeny | 2016H-P3-H9 |
| 2016H-P3-H9_3 | PHI6 | PHI3 | progeny | 2016H-P3-H9 |
| 2016H-P3-H9_4 | PHI6 | PHI3 | progeny | 2016H-P3-H9 |
| 2016H-P3-H9_5 | PHI6 | PHI3 | progeny | 2016H-P3-H9 |
| 2016H-P4-A6_2 | PHI6 | PHI3 | progeny | 2016H-P4-A6 |
| 2016H-P4-A6_3 | PHI6 | PHI3 | progeny | 2016H-P4-A6 |
| 2016H-P3-G11_1 | PHI6 | PHI3 | progeny | 2016H-P3-G11 |
| 2016H-P3-G11_2 | PHI6 | PHI3 | progeny | 2016H-P3-G11 |
| 2016H-P3-G11_3 | PHI6 | PHI3 | progeny | 2016H-P3-G11 |
| 2016H-P3-G11_4 | PHI6 | PHI3 | progeny | 2016H-P3-G11 |
| 2016H-P3-G11_5 | PHI6 | PHI3 | progeny | 2016H-P3-G11 |
| 2016H-P3-G11_6 | PHI6 | PHI3 | progeny | 2016H-P3-G11 |
| 2016H-P3-G11_7 | PHI6 | PHI3 | progeny | 2016H-P3-G11 |
| 2016H-P3-G11_8 | PHI6 | PHI3 | progeny | 2016H-P3-G11 |
| 2016H-P3-G11_9 | PHI6 | PHI3 | progeny | 2016H-P3-G11 |
| 2016H-P3-G12_2 | PHI6 | PHI3 | progeny | 2016H-P3-G12 |
| 2016H-P3-G12_3 | PHI6 | PHI3 | progeny | 2016H-P3-G12 |
| 2016H-P3-G12_4 | PHI6 | PHI3 | progeny | 2016H-P3-G12 |
| 2016H-P3-G12_5 | PHI6 | PHI3 | progeny | 2016H-P3-G12 |
| 2016H-P3-G12_6 | PHI6 | PHI3 | progeny | 2016H-P3-G12 |
| 2016H-P3-G12_8 | PHI6 | PHI3 | progeny | 2016H-P3-G12 |
| 2016H-P3-G12_9 | PHI6 | PHI3 | progeny | 2016H-P3-G12 |
| 2016H-P3-G12_10 | PHI6 | PHI3 | progeny | 2016H-P3-G12 |
| 2016H-P3-G12_11 | PHI6 | PHI3 | progeny | 2016H-P3-G12 |
| 2016H-P3-G12_12 | PHI6 | PHI3 | progeny | 2016H-P3-G12 |
| 2016H-P3-G12_13 | PHI6 | PHI3 | progeny | 2016H-P3-G12 |
| 2016H-P3-F1_1 | PHI6 | PHI3 | progeny | 2016H-P3-F1 |
| 2016H-P3-F1_2 | PHI6 | PHI3 | progeny | 2016H-P3-F1 |
| 2016H-P3-F1_3 | PHI6 | PHI3 | progeny | 2016H-P3-F1 |
| 2016H-P3-F1_4 | PHI6 | PHI3 | progeny | 2016H-P3-F1 |
| 2016H-P3-F1_5 | PHI6 | PHI3 | progeny | 2016H-P3-F1 |
| 2016H-P3-F1_6 | PHI6 | PHI3 | progeny | 2016H-P3-F1 |
| 2016H-P3-F1_7 | PHI6 | PHI3 | progeny | 2016H-P3-F1 |
| 2016H-P3-F1_8 | PHI6 | PHI3 | progeny | 2016H-P3-F1 |
| 2016H-P3-F1_9 | PHI6 | PHI3 | progeny | 2016H-P3-F1 |
| 2016H-P3-F1_10 | PHI6 | PHI3 | progeny | 2016H-P3-F1 |
| 2016H-P3-F1_11 | PHI6 | PHI3 | progeny | 2016H-P3-F1 |
| 2016H-P3-F1_12 | PHI6 | PHI3 | progeny | 2016H-P3-F1 |
| 2016H-P3-F1_14 | PHI6 | PHI3 | progeny | 2016H-P3-F1 |
| 2016H-P3-F1_15 | PHI6 | PHI3 | progeny | 2016H-P3-F1 |
| 2016H-P3-F1_16 | PHI6 | PHI3 | progeny | 2016H-P3-F1 |
| 2016H-P3-F1_17 | PHI6 | PHI3 | progeny | 2016H-P3-F1 |
| 2016H-P3-F11_3 | PHI6 | PHI3 | progeny | 2016H-P3-F11 |
| 2016H-P3-F11_4 | PHI6 | PHI3 | progeny | 2016H-P3-F11 |
| 2016H-P3-F11_5 | PHI6 | PHI3 | progeny | 2016H-P3-F11 |
| 2016H-P3-F11_7 | PHI6 | PHI3 | progeny | 2016H-P3-F11 |
| 2016H-P3-F11_8 | PHI6 | PHI3 | progeny | 2016H-P3-F11 |
| 2016H-P3-F11_11 | PHI6 | PHI3 | progeny | 2016H-P3-F11 |
| 2016H-P3-F11_14 | PHI6 | PHI3 | progeny | 2016H-P3-F11 |

### Table S7. Results of the SNP array analasis (continue)

| Chr | 1 | 1 | 1 | 1 | 1 | 1 | 1 | 1 | 1 | 1 | 1 | 1 | 1 | 1 | 1 | 1 | 1 | 1 | 1 | 1 | 1 | 1 | 1 | 2 | 2 | 2 | 2 | 2 | 2 | 2 | 2 | 2 | 2 | 2 | 2 | 2 | 2 | 2 | 2 | 2 | 2 | 2 | 2 | 2 | 2 | 3 | 3 | 3 | 3 | 3 | 3 | 3 | 3 | 3 | 3 | 3 | 3 | 3 | 3 | 3 | 3 | 3 | 3 | 3 | 3 | 3 | 4 | 4 | 4 | 4 | 4 | 4 | 4 | 4 | 4 | 4 | 4 | 4 | 4 | 4 | 4 | 4 | 4 | 4 | 4 | 4 | 4 | 4 | 4 | 5 | 5 | 5 | 5 | 5 | 5 | 5 | 5 | 5 | 5 | 5 | 5 | 5 | 5 | 5 | 5 | 5 | 5 | 5 | 5 | 6 | 6 | 6 | 6 | 6 | 6 | 6 | 6 |
|---|---|---|---|---|---|---|---|---|---|---|---|---|---|---|---|---|---|---|---|---|---|---|---|---|---|---|---|---|---|---|---|---|---|---|---|---|---|---|---|---|---|---|---|---|---|---|---|---|---|---|---|---|---|---|---|---|---|---|---|---|---|---|---|---|---|---|---|---|---|---|---|---|---|---|---|---|---|---|---|---|---|---|---|---|---|---|---|---|---|---|---|---|---|---|---|---|---|---|---|---|---|---|---|---|---|---|---|---|---|---|---|---|---|---|---|---|---|
| Position, cM | 2.6 | 4.5 | 16 | 39 | 59 | 80 | 84 | 108 | 125 | 137 | 137 | 137 | 154 | 164 | 185 | 192 | 203 | 220 | 246 | 273 | 281 | 300 | 316 | 7.7 | 25 | 35 | 54 | 55 | 78 | 94 | 113 | 129 | 129 | 129 | 130 | 130 | 160 | 178 | 183 | 198 | 213 | 220 | 231 | 235 | 253 | 5.9 | 10 | 28 | 50 | 64 | 77 | 79 | 78 | 79 | 88 | 92 | 98 | 109 | 135 | 160 | 165 | 195 | 215 | 227 | 235 | 246 | 7.4 | 18 | 29 | 38 | 57 | 63 | 67 | 89 | 110 | 111 | 110 | 115 | 129 | 133 | 157 | 166 | 173 | 181 | 193 | 201 | 207 | 210 | 246 | 5.3 | 13 | 20 | 30 | 53 | 78 | 89 | 94 | 101 | 102 | 102 | 102 | 108 | 119 | 138 | 145 | 165 | 195 | 217 | 227 | 5.4 | 14 | 23 | 24 | 23 | 29 | 43 | 54 |

| plant_id | mom_gen | dad_gen | mother_progeny | family_ | | genotype calls |
|---|---|---|---|---|---|---|
| 2016H-P3-F3_3 | PHI6 | PHI3 | progeny | 2016H-P3-F3 | | [illegible] |
| 2016H-P3-H8_5 | PHI6 | PHI3 | progeny | 2016H-P3-H8 | | [illegible] |
| 2016H-P3-G6_5 | PHI6 | PHI3 | progeny | 2016H-P3-G6 | | [illegible] |
| 2016H-P3-F6_4 | PHI6 | PHI3 | progeny | 2016H-P3-F6 | | [illegible] |
| 2016H-P3-F6_5 | PHI6 | PHI3 | progeny | 2016H-P3-F6 | | [illegible] |
| 2016H-P3-F6_6 | PHI6 | PHI3 | progeny | 2016H-P3-F6 | | [illegible] |
| 2016H-P3-G2_1 | PHI6 | PHI3 | progeny | 2016H-P3-G2 | | [illegible] |
| 2016H-P3-G2_2 | PHI6 | PHI3 | progeny | 2016H-P3-G2 | | [illegible] |
| 2016H-P3-F8_1 | PHI6 | PHI3 | progeny | 2016H-P3-F8 | | [illegible] |
| 2016H-P3-F8_2 | PHI6 | PHI3 | progeny | 2016H-P3-F8 | | [illegible] |
| 2016H-P3-G7_6 | PHI6 | PHI3 | progeny | 2016H-P3-G7 | | [illegible] |
| 2016H-P3-A12_1 | PHI7 | PHI3 | progeny | 2016H-P3-A12 | | [illegible] |
| 2016H-P3-A12_2 | PHI7 | PHI3 | progeny | 2016H-P3-A12 | | [illegible] |
| 2016H-P3-C4_1 | PHI7 | PHI3 | progeny | 2016H-P3-C4 | | [illegible] |
| 2016H-P3-A9_1 | PHI7 | PHI3 | progeny | 2016H-P3-A9 | | [illegible] |
| 2016H-P3-C2_1 | PHI7 | PHI3 | progeny | 2016H-P3-C2 | | [illegible] |
| 2016H-P2-H11_2 | PHI7 | PHI3 | progeny | 2016H-P2-H11 | | [illegible] |
| 2016H-P3-B2_1 | PHI7 | PHI3 | progeny | 2016H-P3-B2 | | [illegible] |
| 2016H-P3-B2_2 | PHI7 | PHI3 | progeny | 2016H-P3-B2 | | [illegible] |
| 2016H-P3-A12_3 | PHI7 | PHI3 | progeny | 2016H-P3-A12 | | [illegible] |
| 2016H-P3-A12_12 | PHI7 | PHI3 | progeny | 2016H-P3-A12 | | [illegible] |
| 2016H-P3-C9_2 | PHI7 | PHI3 | progeny | 2016H-P3-C9 | | [illegible] |
| 2016H-P3-A8_2 | PHI7 | PHI3 | progeny | 2016H-P3-A8 | | [illegible] |
| 2016H-P3-A8_3 | PHI7 | PHI3 | progeny | 2016H-P3-A8 | | [illegible] |
| 2016H-P3-A11_4 | PHI7 | PHI3 | progeny | 2016H-P3-A11 | | [illegible] |
| 2016H-P3-A11_5 | PHI7 | PHI3 | progeny | 2016H-P3-A11 | | [illegible] |
| 2016H-P3-A11_6 | PHI7 | PHI3 | progeny | 2016H-P3-A11 | | [illegible] |
| 2016H-P3-B9_2 | PHI7 | PHI3 | progeny | 2016H-P3-B9 | | [illegible] |
| 2016H-P2-C6_3 | PHI4 | PHI3 | progeny | 2016H-P2-C6 | | [illegible] |
| 2016H-P2-C9_1 | PHI4 | PHI3 | progeny | 2016H-P2-C9 | | [illegible] |
| 2016H-P2-C9_4 | PHI4 | PHI3 | progeny | 2016H-P2-C9 | | [illegible] |
| 2016H-P2-B7_1 | PHI4 | PHI3 | progeny | 2016H-P2-B7 | | [illegible] |
| 2016H-P2-B7_2 | PHI4 | PHI3 | progeny | 2016H-P2-B7 | | [illegible] |
| 2016H-P2-B7_3 | PHI4 | PHI3 | progeny | 2016H-P2-B7 | | [illegible] |
| 2016H-P2-B7_4 | PHI4 | PHI3 | progeny | 2016H-P2-B7 | | [illegible] |
| 2016H-P2-F3_2 | PHI4 | PHI3 | progeny | 2016H-P2-F3 | | [illegible] |
| 2016H-P2-F3_3 | PHI4 | PHI3 | progeny | 2016H-P2-F3 | | [illegible] |
| 2016H-P2-G6_1 | PHI4 | PHI3 | progeny | 2016H-P2-G6 | | [illegible] |
| 2016H-P2-G6_2 | PHI4 | PHI3 | progeny | 2016H-P2-G6 | | [illegible] |
| 2016H-P2-G6_3 | PHI4 | PHI3 | progeny | 2016H-P2-G6 | | [illegible] |
| 2016H-P2-G6_4 | PHI4 | PHI3 | progeny | 2016H-P2-G6 | | [illegible] |
| 2016H-P2-E7_1 | PHI4 | PHI3 | progeny | 2016H-P2-E7 | | [illegible] |
| 2016H-P2-E7_2 | PHI4 | PHI3 | progeny | 2016H-P2-E7 | | [illegible] |
| 2016H-P2-A12_1 | PHI4 | PHI3 | progeny | 2016H-P2-A12 | | [illegible] |
| 2016H-P2-A12_2 | PHI4 | PHI3 | progeny | 2016H-P2-A12 | | [illegible] |
| 2016H-P2-C1_4 | PHI4 | PHI3 | progeny | 2016H-P2-C1 | | [illegible] |
| 2016H-P2-C1_5 | PHI4 | PHI3 | progeny | 2016H-P2-C1 | | [illegible] |
| 2016H-P2-C1_6 | PHI4 | PHI3 | progeny | 2016H-P2-C1 | | [illegible] |
| 2016H-P2-E9_1 | PHI4 | PHI3 | progeny | 2016H-P2-E9 | | [illegible] |
| 2016H-P2-E9_3 | PHI4 | PHI3 | progeny | 2016H-P2-E9 | | [illegible] |
| 2016H-P2-E8_6 | PHI4 | PHI3 | progeny | 2016H-P2-E8 | | [illegible] |
| 2016H-P2-F11_1 | PHI4 | PHI3 | progeny | 2016H-P2-F11 | | [illegible] |
| 2016H-P2-F11_2 | PHI4 | PHI3 | progeny | 2016H-P2-F11 | | [illegible] |
| 2016H-P2-D6_4 | PHI4 | PHI3 | progeny | 2016H-P2-D6 | | [illegible] |
| 2016H-P2-D6_5 | PHI4 | PHI3 | progeny | 2016H-P2-D6 | | [illegible] |
| 2016H-P2-D6_6 | PHI4 | PHI3 | progeny | 2016H-P2-D6 | | [illegible] |
| 2016H-P2-D9_5 | PHI4 | PHI3 | progeny | 2016H-P2-D9 | | [illegible] |
| 2016H-P2-E1_1 | PHI4 | PHI3 | progeny | 2016H-P2-E1 | | [illegible] |
| 2016H-P2-E1_2 | PHI4 | PHI3 | progeny | 2016H-P2-E1 | | [illegible] |
| 2016H-P2-E1_3 | PHI4 | PHI3 | progeny | 2016H-P2-E1 | | [illegible] |
| 2016H-P2-D3_1 | PHI4 | PHI3 | progeny | 2016H-P2-D3 | | [illegible] |
| 2016H-P2-D3_3 | PHI4 | PHI3 | progeny | 2016H-P2-D3 | | [illegible] |
| 2016H-P2-D3_4 | PHI4 | PHI3 | progeny | 2016H-P2-D3 | | [illegible] |
| 2016H-P2-D3_5 | PHI4 | PHI3 | progeny | 2016H-P2-D3 | | [illegible] |
| 2016H-P2-D3_6 | PHI4 | PHI3 | progeny | 2016H-P2-D3 | | [illegible] |
| 2016H-P2-D2_1 | PHI4 | PHI3 | progeny | 2016H-P2-D2 | | [illegible] |
| 2016H-P2-D2_2 | PHI4 | PHI3 | progeny | 2016H-P2-D2 | | [illegible] |
| 2016H-P2-G8_2 | PHI4 | PHI3 | progeny | 2016H-P2-G8 | | [illegible] |
| 2016H-P2-G8_3 | PHI4 | PHI3 | progeny | 2016H-P2-G8 | | [illegible] |
| 2016H-P2-C7_1 | PHI4 | PHI3 | progeny | 2016H-P2-C7 | | [illegible] |
| 2016H-P2-C6_1 | PHI4 | PHI3 | progeny | 2016H-P2-C6 | | [illegible] |
| 2016H-P2-C6_2 | PHI4 | PHI3 | progeny | 2016H-P2-C6 | | [illegible] |
| 2016H-P1-G1_3 | PHI10 | PHI3 | progeny | 2016H-P1-G1 | | [illegible] |
| 2016H-P1-G1_4 | PHI10 | PHI3 | progeny | 2016H-P1-G1 | | [illegible] |
| 2016H-P1-G1_5 | PHI10 | PHI3 | progeny | 2016H-P1-G1 | | [illegible] |
| 2016H-P1-G1_6 | PHI10 | PHI3 | progeny | 2016H-P1-G1 | | [illegible] |
| 2016H-P1-F4_2 | PHI10 | PHI3 | progeny | 2016H-P1-F4 | | [illegible] |
| 2016H-P1-F4_3 | PHI10 | PHI3 | progeny | 2016H-P1-F4 | | [illegible] |
| 2016H-P1-F4_4 | PHI10 | PHI3 | progeny | 2016H-P1-F4 | | [illegible] |
| 2016H-P1-F5_1 | PHI10 | PHI3 | progeny | 2016H-P1-F5 | | [illegible] |
| 2016H-P1-F11_6 | PHI10 | PHI3 | progeny | 2016H-P1-F11 | | [illegible] |
| 2016H-P1-G9_11 | PHI10 | PHI3 | progeny | 2016H-P1-G9 | | [illegible] |
| 2016H-P1-G2_1 | PHI10 | PHI3 | progeny | 2016H-P1-G2 | | [illegible] |
| 2016H-P1-G2_2 | PHI10 | PHI3 | progeny | 2016H-P1-G2 | | [illegible] |
| 2016H-P1-G5_2 | PHI10 | PHI3 | progeny | 2016H-P1-G5 | | [illegible] |
| 2016H-P1-G5_4 | PHI10 | PHI3 | progeny | 2016H-P1-G5 | | [illegible] |
| 2016H-P1-F9_1 | PHI10 | PHI3 | progeny | 2016H-P1-F9 | | [illegible] |
| 2016H-P1-F9_2 | PHI10 | PHI3 | progeny | 2016H-P1-F9 | | [illegible] |
| 2016H-P1-F3_4 | PHI10 | PHI3 | progeny | 2016H-P1-F3 | | [illegible] |
| 2016H-P1-F3_9 | PHI10 | PHI3 | progeny | 2016H-P1-F3 | | [illegible] |
| 2016H-P1-F10_2 | PHI10 | PHI3 | progeny | 2016H-P1-F10 | | [illegible] |
| 2016H-P1-F10_6 | PHI10 | PHI3 | progeny | 2016H-P1-F10 | | [illegible] |
| 2016H-P1-F10_7 | PHI10 | PHI3 | progeny | 2016H-P1-F10 | | [illegible] |
| 2016H-P1-F10_9 | PHI10 | PHI3 | progeny | 2016H-P1-F10 | | [illegible] |
| 2016H-P5-G3_4 | PHI9 | PHI3 | progeny | 2016H-P5-G3 | | [illegible] |
| 2016H-P5-E5_1 | PHI9 | PHI3 | progeny | 2016H-P5-E5 | | [illegible] |
| 2016H-P5-E5_2 | PHI9 | PHI3 | progeny | 2016H-P5-E5 | | [illegible] |
| 2016H-P5-E5_3 | PHI9 | PHI3 | progeny | 2016H-P5-E5 | | [illegible] |
| 2016H-P5-E5_7 | PHI9 | PHI3 | progeny | 2016H-P5-E5 | | [illegible] |
| 2016H-P5-E5_9 | PHI9 | PHI3 | progeny | 2016H-P5-E5 | | [illegible] |
| 2016H-P5-E5_10 | PHI9 | PHI3 | progeny | 2016H-P5-E5 | | [illegible] |
| 2016H-P5-E5_11 | PHI9 | PHI3 | progeny | 2016H-P5-E5 | | [illegible] |
| 2016H-P5-G4_1 | PHI9 | PHI3 | progeny | 2016H-P5-G4 | | [illegible] |
| 2016H-P5-F10_1 | PHI9 | PHI3 | progeny | 2016H-P5-F10 | | [illegible] |
| 2016H-P5-F2_1 | PHI9 | PHI3 | progeny | 2016H-P5-F2 | | [illegible] |
| 2016H-P5-E7_3 | PHI9 | PHI3 | progeny | 2016H-P5-E7 | | [illegible] |
| 2016H-P1-C9_1 | PHI2 | PHI3 | progeny | 2016H-P1-C9 | | [illegible] |
| 2016H-P1-C9_2 | PHI2 | PHI3 | progeny | 2016H-P1-C9 | | [illegible] |
| 2016H-P1-C9_3 | PHI2 | PHI3 | progeny | 2016H-P1-C9 | | [illegible] |
| 2016H-P1-C2_3 | PHI2 | PHI3 | progeny | 2016H-P1-C2 | | [illegible] |
| 2016H-P1-C6_1 | PHI2 | PHI3 | progeny | 2016H-P1-C6 | | [illegible] |
| 2016H-P1-A4_1 | PHI2 | PHI3 | progeny | 2016H-P1-A4 | | [illegible] |
| 2016H-P1-A4_2 | PHI2 | PHI3 | progeny | 2016H-P1-A4 | | [illegible] |
| 2016H-P1-A4_4 | PHI2 | PHI3 | progeny | 2016H-P1-A4 | | [illegible] |
| 2016H-P1-A4_5 | PHI2 | PHI3 | progeny | 2016H-P1-A4 | | [illegible] |
| 2016H-P1-A4_9 | PHI2 | PHI3 | progeny | 2016H-P1-A4 | | [illegible] |
| 2016H-P1-A4_10 | PHI2 | PHI3 | progeny | 2016H-P1-A4 | | [illegible] |
| 2016H-P1-A9_1 | PHI2 | PHI3 | progeny | 2016H-P1-A9 | | [illegible] |
| 2016H-P1-A9_3 | PHI2 | PHI3 | progeny | 2016H-P1-A9 | | [illegible] |
| 2016H-P1-A11_1 | PHI2 | PHI3 | progeny | 2016H-P1-A11 | | [illegible] |
| 2016H-P1-A11_3 | PHI2 | PHI3 | progeny | 2016H-P1-A11 | | [illegible] |
| 2016H-P1-A11_4 | PHI2 | PHI3 | progeny | 2016H-P1-A11 | | [illegible] |
| 2016H-P1-A11_5 | PHI2 | PHI3 | progeny | 2016H-P1-A11 | | [illegible] |

plant_id – unique identifier of the plant
mom_gen – genetic backgroiund of mother in the hybred
dad_gen – genetic background of father in the hybred
family_ – identify mother and all the progeny plants from the same mother
mother_progeny – identify each plant as mother of the family or progeny
Chr – number of the chromosome, where marker is located
Distance – position of the marker on the corresponding chromosome in cM
H – heterozygous marker
A, T, G and C – marker homozygous for the corresponding nucleotide
EQV, LS, -9 , etc. – missing values

**Table S7. Results of the SNP array analasis (continue)**

Chr: 6 | 6 | 6 | 6 | 6 | 6 | 6 | 6 | 6 | 6 | 6 | 7 | 7 | 7 | 7 | 7 | 7 | 7 | 7 | 7 | 7 | 7 | 7 | 7 | 7 | 7 | 7 | 7 | 7 | 7 | 7 | 8 | 8 | 8 | 8 | 8 | 8 | 8 | 8 | 8 | 8 | 8 | 8 | 8 | 8 | 8 | 8 | 9 | 9 | 9 | 9 | 9 | 9 | 9 | 9 | 9 | 9 | 9 | 9 | 9 | 9 | 9 | 9 | 10 | 10 | 10 | 10 | 10 | 10 | 10 | 10 | 10 | 10 | 10 | 10

Position, cM: 60 | 76 | 82 | 96 | 105 | 124 | 134 | 147 | 155 | 163 | 181 | 4.2 | 5.6 | 23 | 46 | 52 | 73 | 75 | 75 | 77 | 76 | 91 | 114 | 127 | 131 | 141 | 167 | 175 | 180 | 200 | 211 | 1.8 | 11 | 30 | 51 | 67 | 83 | 85 | 108 | 131 | 132 | 149 | 168 | 174 | 186 | 194 | 210 | 0.3 | 12 | 41 | 60 | 70 | 76 | 76 | 77 | 83 | 100 | 113 | 130 | 142 | 163 | 164 | 181 | 9.5 | 25 | 36 | 55 | 61 | 61 | 78 | 93 | 125 | 127 | 140 | 157

| plant_id | mom_gen | dad_gen | mother_progeny | family_ | |
|---|---|---|---|---|---|
| 2016H-P4-H4 | PHI5 | PHI3 | mother | 2016H-P4-H4 | [illegible] |
| 2016H-P5-D5 | PHI5 | PHI3 | mother | 2016H-P5-D5 | [illegible] |
| 2016H-P4-F3 | PHI5 | PHI3 | mother | 2016H-P4-F3 | [illegible] |
| 2016H-P5-C9 | PHI5 | PHI3 | mother | 2016H-P5-C9 | [illegible] |
| 2016H-P5-B5 | PHI5 | PHI3 | mother | 2016H-P5-B5 | [illegible] |
| 2016H-P4-H1 | PHI5 | PHI3 | mother | 2016H-P4-H1 | [illegible] |
| 2016H-P3-E5 | PHI6 | PHI3 | mother | 2016H-P3-E5 | [illegible] |
| 2016H-P3-E11 | PHI6 | PHI3 | mother | 2016H-P3-E11 | [illegible] |
| 2016H-P3-H10 | PHI6 | PHI3 | mother | 2016H-P3-H10 | [illegible] |
| 2016H-P3-E3 | PHI6 | PHI3 | mother | 2016H-P3-E3 | [illegible] |
| 2016H-P4-A9 | PHI6 | PHI3 | mother | 2016H-P4-A9 | [illegible] |
| 2016H-P3-H7 | PHI6 | PHI3 | mother | 2016H-P3-H7 | [illegible] |
| 2016H-P3-E4 | PHI6 | PHI3 | mother | 2016H-P3-E4 | [illegible] |
| 2016H-P3-H9 | PHI6 | PHI3 | mother | 2016H-P3-H9 | [illegible] |
| 2016H-P4-A6 | PHI6 | PHI3 | mother | 2016H-P4-A6 | [illegible] |
| 2016H-P3-G11 | PHI6 | PHI3 | mother | 2016H-P3-G11 | [illegible] |
| 2016H-P3-G12 | PHI6 | PHI3 | mother | 2016H-P3-G12 | [illegible] |
| 2016H-P3-F1 | PHI6 | PHI3 | mother | 2016H-P3-F1 | [illegible] |
| 2016H-P3-F11 | PHI6 | PHI3 | mother | 2016H-P3-F11 | [illegible] |
| 2016H-P3-F3 | PHI6 | PHI3 | mother | 2016H-P3-F3 | [illegible] |
| 2016H-P3-H8 | PHI6 | PHI3 | mother | 2016H-P3-H8 | [illegible] |
| 2016H-P3-G6 | PHI6 | PHI3 | mother | 2016H-P3-G6 | [illegible] |
| 2016H-P3-F6 | PHI6 | PHI3 | mother | 2016H-P3-F6 | [illegible] |
| 2016H-P3-G2 | PHI6 | PHI3 | mother | 2016H-P3-G2 | [illegible] |
| 2016H-P3-F8 | PHI6 | PHI3 | mother | 2016H-P3-F8 | [illegible] |
| 2016H-P3-G7 | PHI6 | PHI3 | mother | 2016H-P3-G7 | [illegible] |
| 2016H-P3-C4 | PHI7 | PHI3 | mother | 2016H-P3-C4 | [illegible] |
| 2016H-P3-A9 | PHI7 | PHI3 | mother | 2016H-P3-A9 | [illegible] |
| 2016H-P3-C2 | PHI7 | PHI3 | mother | 2016H-P3-C2 | [illegible] |
| 2016H-P2-H11 | PHI7 | PHI3 | mother | 2016H-P2-H11 | [illegible] |
| 2016H-P3-B2 | PHI7 | PHI3 | mother | 2016H-P3-B2 | [illegible] |
| 2016H-P3-C9 | PHI7 | PHI3 | mother | 2016H-P3-C9 | [illegible] |
| 2016H-P3-A8 | PHI7 | PHI3 | mother | 2016H-P3-A8 | [illegible] |
| 2016H-P3-A11 | PHI7 | PHI3 | mother | 2016H-P3-A11 | [illegible] |
| 2016H-P3-B9 | PHI7 | PHI3 | mother | 2016H-P3-B9 | [illegible] |
| 2016H-P3-A12 | PHI7 | PHI3 | mother | 2016H-P3-A12 | [illegible] |
| 2016H-P2-C9 | PHI4 | PHI3 | mother | 2016H-P2-C9 | [illegible] |
| 2016H-P2-B7 | PHI4 | PHI3 | mother | 2016H-P2-B7 | [illegible] |
| 2016H-P2-F3 | PHI4 | PHI3 | mother | 2016H-P2-F3 | [illegible] |
| 2016H-P2-G6 | PHI4 | PHI3 | mother | 2016H-P2-G6 | [illegible] |
| 2016H-P2-E7 | PHI4 | PHI3 | mother | 2016H-P2-E7 | [illegible] |
| 2016H-P2-A12 | PHI4 | PHI3 | mother | 2016H-P2-A12 | [illegible] |
| 2016H-P2-C1 | PHI4 | PHI3 | mother | 2016H-P2-C1 | [illegible] |
| 2016H-P2-E9 | PHI4 | PHI3 | mother | 2016H-P2-E9 | [illegible] |
| 2016H-P2-E8 | PHI4 | PHI3 | mother | 2016H-P2-E8 | [illegible] |
| 2016H-P2-F11 | PHI4 | PHI3 | mother | 2016H-P2-F11 | [illegible] |
| 2016H-P2-D6 | PHI4 | PHI3 | mother | 2016H-P2-D6 | [illegible] |
| 2016H-P2-D9 | PHI4 | PHI3 | mother | 2016H-P2-D9 | [illegible] |
| 2016H-P2-E1 | PHI4 | PHI3 | mother | 2016H-P2-E1 | [illegible] |
| 2016H-P2-D3 | PHI4 | PHI3 | mother | 2016H-P2-D3 | [illegible] |
| 2016H-P2-D2 | PHI4 | PHI3 | mother | 2016H-P2-D2 | [illegible] |
| 2016H-P2-G8 | PHI4 | PHI3 | mother | 2016H-P2-G8 | [illegible] |
| 2016H-P2-C7 | PHI4 | PHI3 | mother | 2016H-P2-C7 | [illegible] |
| 2016H-P2-C6 | PHI4 | PHI3 | mother | 2016H-P2-C6 | [illegible] |
| 2016H-P1-G1 | PHI10 | PHI3 | mother | 2016H-P1-G1 | [illegible] |
| 2016H-P1-F4 | PHI10 | PHI3 | mother | 2016H-P1-F4 | [illegible] |
| 2016H-P1-F5 | PHI10 | PHI3 | mother | 2016H-P1-F5 | [illegible] |
| 2016H-P1-F11 | PHI10 | PHI3 | mother | 2016H-P1-F11 | [illegible] |
| 2016H-P1-G9 | PHI10 | PHI3 | mother | 2016H-P1-G9 | [illegible] |
| 2016H-P1-G2 | PHI10 | PHI3 | mother | 2016H-P1-G2 | [illegible] |
| 2016H-P1-G5 | PHI10 | PHI3 | mother | 2016H-P1-G5 | [illegible] |
| 2016H-P1-F9 | PHI10 | PHI3 | mother | 2016H-P1-F9 | [illegible] |
| 2016H-P1-F3 | PHI10 | PHI3 | mother | 2016H-P1-F3 | [illegible] |
| 2016H-P1-F10 | PHI10 | PHI3 | mother | 2016H-P1-F10 | [illegible] |
| 2016H-P5-G3 | PHI9 | PHI3 | mother | 2016H-P5-G3 | [illegible] |
| 2016H-P5-E5 | PHI9 | PHI3 | mother | 2016H-P5-E5 | [illegible] |
| 2016H-P5-G4 | PHI9 | PHI3 | mother | 2016H-P5-G4 | [illegible] |
| 2016H-P5-F10 | PHI9 | PHI3 | mother | 2016H-P5-F10 | [illegible] |
| 2016H-P5-F2 | PHI9 | PHI3 | mother | 2016H-P5-F2 | [illegible] |
| 2016H-P5-E7 | PHI9 | PHI3 | mother | 2016H-P5-E7 | [illegible] |
| 2016H-P1-C9 | PHI2 | PHI3 | mother | 2016H-P1-C9 | [illegible] |
| 2016H-P1-C2 | PHI2 | PHI3 | mother | 2016H-P1-C2 | [illegible] |
| 2016H-P1-C6 | PHI2 | PHI3 | mother | 2016H-P1-C6 | [illegible] |
| 2016H-P1-A4 | PHI2 | PHI3 | mother | 2016H-P1-A4 | [illegible] |
| 2016H-P1-A9 | PHI2 | PHI3 | mother | 2016H-P1-A9 | [illegible] |
| 2016H-P1-A11 | PHI2 | PHI3 | mother | 2016H-P1-A11 | [illegible] |
| 2016H-P4-H4_2 | PHI5 | PHI3 | progeny | 2016H-P4-H4 | [illegible] |
| 2016H-P5-D5_1 | PHI5 | PHI3 | progeny | 2016H-P5-D5 | [illegible] |
| 2016H-P5-D5_2 | PHI5 | PHI3 | progeny | 2016H-P5-D5 | [illegible] |
| 2016H-P5-D5_3 | PHI5 | PHI3 | progeny | 2016H-P5-D5 | [illegible] |
| 2016H-P4-F3_1 | PHI5 | PHI3 | progeny | 2016H-P4-F3 | [illegible] |
| 2016H-P5-C9_1 | PHI5 | PHI3 | progeny | 2016H-P5-C9 | [illegible] |
| 2016H-P5-C9_2 | PHI5 | PHI3 | progeny | 2016H-P5-C9 | [illegible] |
| 2016H-P5-B6_1 | PHI5 | PHI3 | progeny | 2016H-P5-B6 | [illegible] |
| 2016H-P5-B6_2 | PHI5 | PHI3 | progeny | 2016H-P5-B6 | [illegible] |
| 2016H-P5-B5_1 | PHI5 | PHI3 | progeny | 2016H-P5-B5 | [illegible] |
| 2016H-P5-B5_2 | PHI5 | PHI3 | progeny | 2016H-P5-B5 | [illegible] |
| 2016H-P4-H1_1 | PHI5 | PHI3 | progeny | 2016H-P4-H1 | [illegible] |
| 2016H-P3-E5_1 | PHI6 | PHI3 | progeny | 2016H-P3-E5 | [illegible] |
| 2016H-P3-E11_1 | PHI6 | PHI3 | progeny | 2016H-P3-E11 | [illegible] |
| 2016H-P3-H10_1 | PHI6 | PHI3 | progeny | 2016H-P3-H10 | [illegible] |
| 2016H-P3-E3_2 | PHI6 | PHI3 | progeny | 2016H-P3-E3 | [illegible] |
| 2016H-P4-A9_1 | PHI6 | PHI3 | progeny | 2016H-P4-A9 | [illegible] |
| 2016H-P3-H7_1 | PHI6 | PHI3 | progeny | 2016H-P3-H7 | [illegible] |
| 2016H-P3-E4_3 | PHI6 | PHI3 | progeny | 2016H-P3-E4 | [illegible] |
| 2016H-P3-E4_6 | PHI6 | PHI3 | progeny | 2016H-P3-E4 | [illegible] |
| 2016H-P3-H9_1 | PHI6 | PHI3 | progeny | 2016H-P3-H9 | [illegible] |
| 2016H-P3-H9_2 | PHI6 | PHI3 | progeny | 2016H-P3-H9 | [illegible] |
| 2016H-P3-H9_3 | PHI6 | PHI3 | progeny | 2016H-P3-H9 | [illegible] |
| 2016H-P3-H9_4 | PHI6 | PHI3 | progeny | 2016H-P3-H9 | [illegible] |
| 2016H-P3-H9_5 | PHI6 | PHI3 | progeny | 2016H-P3-H9 | [illegible] |
| 2016H-P4-A6_2 | PHI6 | PHI3 | progeny | 2016H-P4-A6 | [illegible] |
| 2016H-P4-A6_3 | PHI6 | PHI3 | progeny | 2016H-P4-A6 | [illegible] |
| 2016H-P3-G11_1 | PHI6 | PHI3 | progeny | 2016H-P3-G11 | [illegible] |
| 2016H-P3-G11_2 | PHI6 | PHI3 | progeny | 2016H-P3-G11 | [illegible] |
| 2016H-P3-G11_3 | PHI6 | PHI3 | progeny | 2016H-P3-G11 | [illegible] |
| 2016H-P3-G11_4 | PHI6 | PHI3 | progeny | 2016H-P3-G11 | [illegible] |
| 2016H-P3-G11_5 | PHI6 | PHI3 | progeny | 2016H-P3-G11 | [illegible] |
| 2016H-P3-G11_6 | PHI6 | PHI3 | progeny | 2016H-P3-G11 | [illegible] |
| 2016H-P3-G11_7 | PHI6 | PHI3 | progeny | 2016H-P3-G11 | [illegible] |
| 2016H-P3-G11_8 | PHI6 | PHI3 | progeny | 2016H-P3-G11 | [illegible] |
| 2016H-P3-G11_9 | PHI6 | PHI3 | progeny | 2016H-P3-G11 | [illegible] |
| 2016H-P3-G12_2 | PHI6 | PHI3 | progeny | 2016H-P3-G12 | [illegible] |
| 2016H-P3-G12_3 | PHI6 | PHI3 | progeny | 2016H-P3-G12 | [illegible] |
| 2016H-P3-G12_4 | PHI6 | PHI3 | progeny | 2016H-P3-G12 | [illegible] |
| 2016H-P3-G12_5 | PHI6 | PHI3 | progeny | 2016H-P3-G12 | [illegible] |
| 2016H-P3-G12_6 | PHI6 | PHI3 | progeny | 2016H-P3-G12 | [illegible] |
| 2016H-P3-G12_8 | PHI6 | PHI3 | progeny | 2016H-P3-G12 | [illegible] |
| 2016H-P3-G12_9 | PHI6 | PHI3 | progeny | 2016H-P3-G12 | [illegible] |
| 2016H-P3-G12_10 | PHI6 | PHI3 | progeny | 2016H-P3-G12 | [illegible] |
| 2016H-P3-G12_11 | PHI6 | PHI3 | progeny | 2016H-P3-G12 | [illegible] |
| 2016H-P3-G12_12 | PHI6 | PHI3 | progeny | 2016H-P3-G12 | [illegible] |
| 2016H-P3-G12_13 | PHI6 | PHI3 | progeny | 2016H-P3-G12 | [illegible] |
| 2016H-P3-F1_1 | PHI6 | PHI3 | progeny | 2016H-P3-F1 | [illegible] |
| 2016H-P3-F1_2 | PHI6 | PHI3 | progeny | 2016H-P3-F1 | [illegible] |
| 2016H-P3-F1_3 | PHI6 | PHI3 | progeny | 2016H-P3-F1 | [illegible] |
| 2016H-P3-F1_4 | PHI6 | PHI3 | progeny | 2016H-P3-F1 | [illegible] |
| 2016H-P3-F1_5 | PHI6 | PHI3 | progeny | 2016H-P3-F1 | [illegible] |
| 2016H-P3-F1_6 | PHI6 | PHI3 | progeny | 2016H-P3-F1 | [illegible] |
| 2016H-P3-F1_7 | PHI6 | PHI3 | progeny | 2016H-P3-F1 | [illegible] |
| 2016H-P3-F1_8 | PHI6 | PHI3 | progeny | 2016H-P3-F1 | [illegible] |
| 2016H-P3-F1_9 | PHI6 | PHI3 | progeny | 2016H-P3-F1 | [illegible] |
| 2016H-P3-F1_10 | PHI6 | PHI3 | progeny | 2016H-P3-F1 | [illegible] |
| 2016H-P3-F1_11 | PHI6 | PHI3 | progeny | 2016H-P3-F1 | [illegible] |
| 2016H-P3-F1_12 | PHI6 | PHI3 | progeny | 2016H-P3-F1 | [illegible] |
| 2016H-P3-F1_14 | PHI6 | PHI3 | progeny | 2016H-P3-F1 | [illegible] |
| 2016H-P3-F1_15 | PHI6 | PHI3 | progeny | 2016H-P3-F1 | [illegible] |
| 2016H-P3-F1_16 | PHI6 | PHI3 | progeny | 2016H-P3-F1 | [illegible] |
| 2016H-P3-F1_17 | PHI6 | PHI3 | progeny | 2016H-P3-F1 | [illegible] |
| 2016H-P3-F11_3 | PHI6 | PHI3 | progeny | 2016H-P3-F11 | [illegible] |
| 2016H-P3-F11_4 | PHI6 | PHI3 | progeny | 2016H-P3-F11 | [illegible] |
| 2016H-P3-F11_5 | PHI6 | PHI3 | progeny | 2016H-P3-F11 | [illegible] |
| 2016H-P3-F11_7 | PHI6 | PHI3 | progeny | 2016H-P3-F11 | [illegible] |
| 2016H-P3-F11_8 | PHI6 | PHI3 | progeny | 2016H-P3-F11 | [illegible] |
| 2016H-P3-F11_11 | PHI6 | PHI3 | progeny | 2016H-P3-F11 | [illegible] |
| 2016H-P3-F11_14 | PHI6 | PHI3 | progeny | 2016H-P3-F11 | [illegible] |

### Table S7. Results of the SNP array analasis (continue)

Marker columns (Chr / Position, cM):

| Chr | 6 | 6 | 6 | 6 | 6 | 6 | 6 | 6 | 6 | 6 | 6 | 7 | 7 | 7 | 7 | 7 | 7 | 7 | 7 | 7 | 7 | 7 | 7 | 7 | 7 | 7 | 7 | 7 | 7 | 7 | 7 | 8 | 8 | 8 | 8 | 8 | 8 | 8 | 8 | 8 | 8 | 8 | 8 | 8 | 8 | 8 | 8 | 9 | 9 | 9 | 9 | 9 | 9 | 9 | 9 | 9 | 9 | 9 | 9 | 9 | 9 | 9 | 9 | 10 | 10 | 10 | 10 | 10 | 10 | 10 | 10 | 10 | 10 | 10 | 10 |
|---|---|---|---|---|---|---|---|---|---|---|---|---|---|---|---|---|---|---|---|---|---|---|---|---|---|---|---|---|---|---|---|---|---|---|---|---|---|---|---|---|---|---|---|---|---|---|---|---|---|---|---|---|---|---|---|---|---|---|---|---|---|---|---|---|---|---|---|---|---|---|---|---|---|---|---|
| Position, cM | 60 | 76 | 82 | 96 | 105 | 124 | 134 | 147 | 155 | 163 | 181 | 4.2 | 5.6 | 23 | 46 | 52 | 73 | 75 | 75 | 77 | 76 | 91 | 114 | 127 | 131 | 141 | 167 | 175 | 180 | 200 | 211 | 1.8 | 11 | 30 | 51 | 67 | 83 | 85 | 108 | 131 | 132 | 149 | 168 | 174 | 186 | 194 | 210 | 0.3 | 12 | 41 | 60 | 70 | 76 | 76 | 77 | 83 | 100 | 113 | 130 | 142 | 163 | 164 | 181 | 9.5 | 25 | 36 | 55 | 61 | 61 | 78 | 93 | 125 | 127 | 140 | 157 |

| plant_id | mom_gen | dad_gen | mother_progeny | family_ | genotypes |
|---|---|---|---|---|---|
| 2016H-P3-F3_3 | PHI6 | PHI3 | progeny | 2016H-P3-F3 | [illegible] |
| 2016H-P3-H8_5 | PHI6 | PHI3 | progeny | 2016H-P3-H8 | [illegible] |
| 2016H-P3-G6_5 | PHI6 | PHI3 | progeny | 2016H-P3-G6 | [illegible] |
| 2016H-P3-F6_4 | PHI6 | PHI3 | progeny | 2016H-P3-F6 | [illegible] |
| 2016H-P3-F6_5 | PHI6 | PHI3 | progeny | 2016H-P3-F6 | [illegible] |
| 2016H-P3-F6_6 | PHI6 | PHI3 | progeny | 2016H-P3-F6 | [illegible] |
| 2016H-P3-G2_1 | PHI6 | PHI3 | progeny | 2016H-P3-G2 | [illegible] |
| 2016H-P3-G2_2 | PHI6 | PHI3 | progeny | 2016H-P3-G2 | [illegible] |
| 2016H-P3-F8_1 | PHI6 | PHI3 | progeny | 2016H-P3-F8 | [illegible] |
| 2016H-P3-F8_2 | PHI6 | PHI3 | progeny | 2016H-P3-F8 | [illegible] |
| 2016H-P3-G7_6 | PHI6 | PHI3 | progeny | 2016H-P3-G7 | [illegible] |
| 2016H-P3-A12_1 | PHI7 | PHI3 | progeny | 2016H-P3-A12 | [illegible] |
| 2016H-P3-A12_2 | PHI7 | PHI3 | progeny | 2016H-P3-A12 | [illegible] |
| 2016H-P3-C4_1 | PHI7 | PHI3 | progeny | 2016H-P3-C4 | [illegible] |
| 2016H-P3-A9_1 | PHI7 | PHI3 | progeny | 2016H-P3-A9 | [illegible] |
| 2016H-P3-C2_1 | PHI7 | PHI3 | progeny | 2016H-P3-C2 | [illegible] |
| 2016H-P2-H11_2 | PHI7 | PHI3 | progeny | 2016H-P2-H11 | [illegible] |
| 2016H-P3-B2_1 | PHI7 | PHI3 | progeny | 2016H-P3-B2 | [illegible] |
| 2016H-P3-B2_2 | PHI7 | PHI3 | progeny | 2016H-P3-B2 | [illegible] |
| 2016H-P3-A12_3 | PHI7 | PHI3 | progeny | 2016H-P3-A12 | [illegible] |
| 2016H-P3-A12_12 | PHI7 | PHI3 | progeny | 2016H-P3-A12 | [illegible] |
| 2016H-P3-C9_2 | PHI7 | PHI3 | progeny | 2016H-P3-C9 | [illegible] |
| 2016H-P3-A8_2 | PHI7 | PHI3 | progeny | 2016H-P3-A8 | [illegible] |
| 2016H-P3-A8_3 | PHI7 | PHI3 | progeny | 2016H-P3-A8 | [illegible] |
| 2016H-P3-A11_4 | PHI7 | PHI3 | progeny | 2016H-P3-A11 | [illegible] |
| 2016H-P3-A11_5 | PHI7 | PHI3 | progeny | 2016H-P3-A11 | [illegible] |
| 2016H-P3-A11_6 | PHI7 | PHI3 | progeny | 2016H-P3-A11 | [illegible] |
| 2016H-P3-B9_2 | PHI7 | PHI3 | progeny | 2016H-P3-B9 | [illegible] |
| 2016H-P2-C6_3 | PHI4 | PHI3 | progeny | 2016H-P2-C6 | [illegible] |
| 2016H-P2-C9_1 | PHI4 | PHI3 | progeny | 2016H-P2-C9 | [illegible] |
| 2016H-P2-C9_4 | PHI4 | PHI3 | progeny | 2016H-P2-C9 | [illegible] |
| 2016H-P2-B7_1 | PHI4 | PHI3 | progeny | 2016H-P2-B7 | [illegible] |
| 2016H-P2-B7_2 | PHI4 | PHI3 | progeny | 2016H-P2-B7 | [illegible] |
| 2016H-P2-B7_3 | PHI4 | PHI3 | progeny | 2016H-P2-B7 | [illegible] |
| 2016H-P2-B7_4 | PHI4 | PHI3 | progeny | 2016H-P2-B7 | [illegible] |
| 2016H-P2-F3_2 | PHI4 | PHI3 | progeny | 2016H-P2-F3 | [illegible] |
| 2016H-P2-F3_3 | PHI4 | PHI3 | progeny | 2016H-P2-F3 | [illegible] |
| 2016H-P2-G6_1 | PHI4 | PHI3 | progeny | 2016H-P2-G6 | [illegible] |
| 2016H-P2-G6_2 | PHI4 | PHI3 | progeny | 2016H-P2-G6 | [illegible] |
| 2016H-P2-G6_3 | PHI4 | PHI3 | progeny | 2016H-P2-G6 | [illegible] |
| 2016H-P2-G6_4 | PHI4 | PHI3 | progeny | 2016H-P2-G6 | [illegible] |
| 2016H-P2-E7_1 | PHI4 | PHI3 | progeny | 2016H-P2-E7 | [illegible] |
| 2016H-P2-E7_2 | PHI4 | PHI3 | progeny | 2016H-P2-E7 | [illegible] |
| 2016H-P2-A12_1 | PHI4 | PHI3 | progeny | 2016H-P2-A12 | [illegible] |
| 2016H-P2-A12_2 | PHI4 | PHI3 | progeny | 2016H-P2-A12 | [illegible] |
| 2016H-P2-C1_4 | PHI4 | PHI3 | progeny | 2016H-P2-C1 | [illegible] |
| 2016H-P2-C1_5 | PHI4 | PHI3 | progeny | 2016H-P2-C1 | [illegible] |
| 2016H-P2-C1_6 | PHI4 | PHI3 | progeny | 2016H-P2-C1 | [illegible] |
| 2016H-P2-E9_1 | PHI4 | PHI3 | progeny | 2016H-P2-E9 | [illegible] |
| 2016H-P2-E9_3 | PHI4 | PHI3 | progeny | 2016H-P2-E9 | [illegible] |
| 2016H-P2-E8_6 | PHI4 | PHI3 | progeny | 2016H-P2-E8 | [illegible] |
| 2016H-P2-F11_1 | PHI4 | PHI3 | progeny | 2016H-P2-F11 | [illegible] |
| 2016H-P2-F11_2 | PHI4 | PHI3 | progeny | 2016H-P2-F11 | [illegible] |
| 2016H-P2-D6_4 | PHI4 | PHI3 | progeny | 2016H-P2-D6 | [illegible] |
| 2016H-P2-D6_5 | PHI4 | PHI3 | progeny | 2016H-P2-D6 | [illegible] |
| 2016H-P2-D6_6 | PHI4 | PHI3 | progeny | 2016H-P2-D6 | [illegible] |
| 2016H-P2-D9_5 | PHI4 | PHI3 | progeny | 2016H-P2-D9 | [illegible] |
| 2016H-P2-E1_1 | PHI4 | PHI3 | progeny | 2016H-P2-E1 | [illegible] |
| 2016H-P2-E1_2 | PHI4 | PHI3 | progeny | 2016H-P2-E1 | [illegible] |
| 2016H-P2-E1_3 | PHI4 | PHI3 | progeny | 2016H-P2-E1 | [illegible] |
| 2016H-P2-D3_1 | PHI4 | PHI3 | progeny | 2016H-P2-D3 | [illegible] |
| 2016H-P2-D3_3 | PHI4 | PHI3 | progeny | 2016H-P2-D3 | [illegible] |
| 2016H-P2-D3_4 | PHI4 | PHI3 | progeny | 2016H-P2-D3 | [illegible] |
| 2016H-P2-D3_5 | PHI4 | PHI3 | progeny | 2016H-P2-D3 | [illegible] |
| 2016H-P2-D3_6 | PHI4 | PHI3 | progeny | 2016H-P2-D3 | [illegible] |
| 2016H-P2-D2_1 | PHI4 | PHI3 | progeny | 2016H-P2-D2 | [illegible] |
| 2016H-P2-D2_2 | PHI4 | PHI3 | progeny | 2016H-P2-D2 | [illegible] |
| 2016H-P2-G8_2 | PHI4 | PHI3 | progeny | 2016H-P2-G8 | [illegible] |
| 2016H-P2-G8_3 | PHI4 | PHI3 | progeny | 2016H-P2-G8 | [illegible] |
| 2016H-P2-C7_1 | PHI4 | PHI3 | progeny | 2016H-P2-C7 | [illegible] |
| 2016H-P2-C6_1 | PHI4 | PHI3 | progeny | 2016H-P2-C6 | [illegible] |
| 2016H-P2-C6_2 | PHI4 | PHI3 | progeny | 2016H-P2-C6 | [illegible] |
| 2016H-P1-G1_3 | PHI10 | PHI3 | progeny | 2016H-P1-G1 | [illegible] |
| 2016H-P1-G1_4 | PHI10 | PHI3 | progeny | 2016H-P1-G1 | [illegible] |
| 2016H-P1-G1_5 | PHI10 | PHI3 | progeny | 2016H-P1-G1 | [illegible] |
| 2016H-P1-G1_6 | PHI10 | PHI3 | progeny | 2016H-P1-G1 | [illegible] |
| 2016H-P1-F4_2 | PHI10 | PHI3 | progeny | 2016H-P1-F4 | [illegible] |
| 2016H-P1-F4_3 | PHI10 | PHI3 | progeny | 2016H-P1-F4 | [illegible] |
| 2016H-P1-F4_4 | PHI10 | PHI3 | progeny | 2016H-P1-F4 | [illegible] |
| 2016H-P1-F5_1 | PHI10 | PHI3 | progeny | 2016H-P1-F5 | [illegible] |
| 2016H-P1-F11_6 | PHI10 | PHI3 | progeny | 2016H-P1-F11 | [illegible] |
| 2016H-P1-G9_11 | PHI10 | PHI3 | progeny | 2016H-P1-G9 | [illegible] |
| 2016H-P1-G2_1 | PHI10 | PHI3 | progeny | 2016H-P1-G2 | [illegible] |
| 2016H-P1-G2_2 | PHI10 | PHI3 | progeny | 2016H-P1-G2 | [illegible] |
| 2016H-P1-G5_2 | PHI10 | PHI3 | progeny | 2016H-P1-G5 | [illegible] |
| 2016H-P1-G5_4 | PHI10 | PHI3 | progeny | 2016H-P1-G5 | [illegible] |
| 2016H-P1-F9_1 | PHI10 | PHI3 | progeny | 2016H-P1-F9 | [illegible] |
| 2016H-P1-F9_2 | PHI10 | PHI3 | progeny | 2016H-P1-F9 | [illegible] |
| 2016H-P1-F3_4 | PHI10 | PHI3 | progeny | 2016H-P1-F3 | [illegible] |
| 2016H-P1-F3_9 | PHI10 | PHI3 | progeny | 2016H-P1-F3 | [illegible] |
| 2016H-P1-F10_2 | PHI10 | PHI3 | progeny | 2016H-P1-F10 | [illegible] |
| 2016H-P1-F10_6 | PHI10 | PHI3 | progeny | 2016H-P1-F10 | [illegible] |
| 2016H-P1-F10_7 | PHI10 | PHI3 | progeny | 2016H-P1-F10 | [illegible] |
| 2016H-P1-F10_9 | PHI10 | PHI3 | progeny | 2016H-P1-F10 | [illegible] |
| 2016H-P5-G3_4 | PHI9 | PHI3 | progeny | 2016H-P5-G3 | [illegible] |
| 2016H-P5-E5_1 | PHI9 | PHI3 | progeny | 2016H-P5-E5 | [illegible] |
| 2016H-P5-E5_2 | PHI9 | PHI3 | progeny | 2016H-P5-E5 | [illegible] |
| 2016H-P5-E5_3 | PHI9 | PHI3 | progeny | 2016H-P5-E5 | [illegible] |
| 2016H-P5-E5_7 | PHI9 | PHI3 | progeny | 2016H-P5-E5 | [illegible] |
| 2016H-P5-E5_9 | PHI9 | PHI3 | progeny | 2016H-P5-E5 | [illegible] |
| 2016H-P5-E5_10 | PHI9 | PHI3 | progeny | 2016H-P5-E5 | [illegible] |
| 2016H-P5-E5_11 | PHI9 | PHI3 | progeny | 2016H-P5-E5 | [illegible] |
| 2016H-P5-G4_1 | PHI9 | PHI3 | progeny | 2016H-P5-G4 | [illegible] |
| 2016H-P5-F10_1 | PHI9 | PHI3 | progeny | 2016H-P5-F10 | [illegible] |
| 2016H-P5-F2_1 | PHI9 | PHI3 | progeny | 2016H-P5-F2 | [illegible] |
| 2016H-P5-E7_3 | PHI9 | PHI3 | progeny | 2016H-P5-E7 | [illegible] |
| 2016H-P1-C9_1 | PHI2 | PHI3 | progeny | 2016H-P1-C9 | [illegible] |
| 2016H-P1-C9_2 | PHI2 | PHI3 | progeny | 2016H-P1-C9 | [illegible] |
| 2016H-P1-C9_3 | PHI2 | PHI3 | progeny | 2016H-P1-C9 | [illegible] |
| 2016H-P1-C2_3 | PHI2 | PHI3 | progeny | 2016H-P1-C2 | [illegible] |
| 2016H-P1-C6_1 | PHI2 | PHI3 | progeny | 2016H-P1-C6 | [illegible] |
| 2016H-P1-A4_1 | PHI2 | PHI3 | progeny | 2016H-P1-A4 | [illegible] |
| 2016H-P1-A4_2 | PHI2 | PHI3 | progeny | 2016H-P1-A4 | [illegible] |
| 2016H-P1-A4_4 | PHI2 | PHI3 | progeny | 2016H-P1-A4 | [illegible] |
| 2016H-P1-A4_5 | PHI2 | PHI3 | progeny | 2016H-P1-A4 | [illegible] |
| 2016H-P1-A4_9 | PHI2 | PHI3 | progeny | 2016H-P1-A4 | [illegible] |
| 2016H-P1-A4_10 | PHI2 | PHI3 | progeny | 2016H-P1-A4 | [illegible] |
| 2016H-P1-A9_1 | PHI2 | PHI3 | progeny | 2016H-P1-A9 | [illegible] |
| 2016H-P1-A9_3 | PHI2 | PHI3 | progeny | 2016H-P1-A9 | [illegible] |
| 2016H-P1-A11_1 | PHI2 | PHI3 | progeny | 2016H-P1-A11 | [illegible] |
| 2016H-P1-A11_3 | PHI2 | PHI3 | progeny | 2016H-P1-A11 | [illegible] |
| 2016H-P1-A11_4 | PHI2 | PHI3 | progeny | 2016H-P1-A11 | [illegible] |
| 2016H-P1-A11_5 | PHI2 | PHI3 | progeny | 2016H-P1-A11 | [illegible] |

plant_id – unique identifier of the plant
mom_gen – genetic backgroiund of mother in the hybred
dad_gen – genetic background of father in the hybred
family_ – identify mother and all the progeny plants from the same r
mother_progeny – identify each plant as mother of the family or progeny
Chr – number of the chromosome, where marker is located
Distance – position of the marker on the corresponding chromosome
H – heterozygous marker
A, T, G and C – marker homozygous for the corresponding nucleotide
EQV, LS, -9 , etc. – missing values

**Table S8. Results of the SNP array analasis**

| | | | | | Chr | 1 | 1 | 1 | 1 | [illegible] |
|---|---|---|---|---|---|---|---|---|---|---|
| | | | | | Position, cM | 13 | 23 | 48 | 50 | [illegible] |
| plant_id | mom_gen | dad_gen | mother_progeny | family_ | | | | | | |
| 2016H_P4-H4 | PHI5 | PHI3 | mother | P4-H4 | | H | H | H | A | [illegible] |
| 2016H_P5-D5 | PHI5 | PHI3 | mother | P5-D5 | | H | H | H | H | [illegible] |
| 2016H_P4-F3 | PHI5 | PHI3 | mother | P4-F3 | | H | H | H | H | [illegible] |
| 2016H_P5-C9 | PHI5 | PHI3 | mother | P5-C9 | | H | H | H | A | [illegible] |
| 2016H_P5-B5 | PHI5 | PHI3 | mother | P5-B5 | | H | H | H | H | [illegible] |
| 2016H_P4-H1 | PHI5 | PHI3 | mother | P4-H1 | | H | H | H | A | [illegible] |
| 2016H_P3-E5 | PHI6 | PHI3 | mother | P3-E5 | | H | H | H | H | [illegible] |
| 2016H_P3-E11 | PHI6 | PHI3 | mother | P3-E11 | | H | H | H | A | [illegible] |
| 2016H_P3-H10 | PHI6 | PHI3 | mother | P3-H10 | | H | H | H | A | [illegible] |
| 2016H_P3-E3 | PHI6 | PHI3 | mother | P3-E3 | | H | H | H | H | [illegible] |
| 2016H_P4-A9 | PHI6 | PHI3 | mother | P4-A9 | | H | H | H | A | [illegible] |
| 2016H_P3-H7 | PHI6 | PHI3 | mother | P3-H7 | | H | H | H | H | [illegible] |
| 2016H_P3-E4 | PHI6 | PHI3 | mother | P3-E4 | | H | H | H | A | [illegible] |
| 2016H_P3-H9 | PHI6 | PHI3 | mother | P3-H9 | | H | H | H | A | [illegible] |
| 2016H_P4-A6 | PHI6 | PHI3 | mother | P4-A6 | | H | H | H | A | [illegible] |
| 2016H_P3-G11 | PHI6 | PHI3 | mother | P3-G11 | | H | H | H | A | [illegible] |
| 2016H_P3-G12 | PHI6 | PHI3 | mother | P3-G12 | | H | H | H | A | [illegible] |
| 2016H_P3-F1 | PHI6 | PHI3 | mother | P3-F1 | | H | H | H | H | [illegible] |
| 2016H_P3-F11 | PHI6 | PHI3 | mother | P3-F11 | | H | H | H | A | [illegible] |
| 2016H_P3-F3 | PHI6 | PHI3 | mother | P3-F3 | | H | H | H | H | [illegible] |
| 2016H_P3-H8 | PHI6 | PHI3 | mother | P3-H8 | | H | H | H | H | [illegible] |
| 2016H_P3-G6 | PHI6 | PHI3 | mother | P3-G6 | | H | H | H | H | [illegible] |
| 2016H_P3-F6 | PHI6 | PHI3 | mother | P3-F6 | | H | H | H | H | [illegible] |
| 2016H_P3-G2 | PHI6 | PHI3 | mother | P3-G2 | | H | H | H | H | [illegible] |
| 2016H_P3-F8 | PHI6 | PHI3 | mother | P3-F8 | | H | H | H | H | [illegible] |
| 2016H_P3-G7 | PHI6 | PHI3 | mother | P3-G7 | | H | H | H | A | [illegible] |
| 2016H_P3-C4 | PHI7 | PHI3 | mother | P3-C4 | | H | H | A | H | [illegible] |
| 2016H_P3-A9 | PHI7 | PHI3 | mother | P3-A9 | | H | H | A | H | [illegible] |
| 2016H_P3-C2 | PHI7 | PHI3 | mother | P3-C2 | | H | H | A | A | [illegible] |
| 2016H_P2-H11 | PHI7 | PHI3 | mother | P2-H11 | | H | H | A | A | [illegible] |
| 2016H_P3-B2 | PHI7 | PHI3 | mother | P3-B2 | | H | H | A | H | [illegible] |
| 2016H_P3-C9 | PHI7 | PHI3 | mother | P3-C9 | | H | H | A | H | [illegible] |
| 2016H_P3-A8 | PHI7 | PHI3 | mother | P3-A8 | | H | H | A | A | [illegible] |
| 2016H_P3-A11 | PHI7 | PHI3 | mother | P3-A11 | | H | H | A | H | [illegible] |
| 2016H_P3-B9 | PHI7 | PHI3 | mother | P3-B9 | | H | H | A | H | [illegible] |
| 2016H_P3-A12 | PHI7 | PHI3 | mother | P3-A12 | | H | H | A | H | [illegible] |
| 2016H_P2-C9 | PHI4 | PHI3 | mother | P2-C9 | | H | H | H | A | [illegible] |
| 2016H_P2-B7 | PHI4 | PHI3 | mother | P2-B7 | | H | H | H | A | [illegible] |
| 2016H_P2-F3 | PHI4 | PHI3 | mother | P2-F3 | | G | H | H | A | [illegible] |
| 2016H_P2-G6 | PHI4 | PHI3 | mother | P2-G6 | | H | H | H | A | [illegible] |
| 2016H_P2-E7 | PHI4 | PHI3 | mother | P2-E7 | | G | G | H | H | [illegible] |
| 2016H_P2-A12 | PHI4 | PHI3 | mother | P2-A12 | | G | G | H | A | [illegible] |
| 2016H_P2-C1 | PHI4 | PHI3 | mother | P2-C1 | | H | H | H | A | [illegible] |
| 2016H_P2-E9 | PHI4 | PHI3 | mother | P2-E9 | | G | G | H | H | [illegible] |
| 2016H_P2-E8 | PHI4 | PHI3 | mother | P2-E8 | | G | G | H | H | [illegible] |
| 2016H_P2-F11 | PHI4 | PHI3 | mother | P2-F11 | | H | H | H | A | [illegible] |
| 2016H_P2-D6 | PHI4 | PHI3 | mother | P2-D6 | | H | H | H | A | [illegible] |
| 2016H_P2-D9 | PHI4 | PHI3 | mother | P2-D9 | | G | G | H | H | [illegible] |
| 2016H_P2-E1 | PHI4 | PHI3 | mother | P2-E1 | | G | H | H | H | [illegible] |
| 2016H_P2-D3 | PHI4 | PHI3 | mother | P2-D3 | | H | H | H | H | [illegible] |
| 2016H_P2-D2 | PHI4 | PHI3 | mother | P2-D2 | | G | G | H | A | [illegible] |
| 2016H_P2-G8 | PHI4 | PHI3 | mother | P2-G8 | | G | G | H | A | [illegible] |
| 2016H_P2-C7 | PHI4 | PHI3 | mother | P2-C7 | | G | G | H | H | [illegible] |
| 2016H_P2-C6 | PHI4 | PHI3 | mother | P2-C6 | | H | H | H | A | [illegible] |
| 2016H_P1-G1 | PHI10 | PHI3 | mother | P1-G1 | | H | G | H | A | [illegible] |
| 2016H_P1-F4 | PHI10 | PHI3 | mother | P1-F4 | | H | H | H | H | [illegible] |
| 2016H_P1-F5 | PHI10 | PHI3 | mother | P1-F5 | | H | G | H | H | [illegible] |
| 2016H_P1-F11 | PHI10 | PHI3 | mother | P1-F11 | | H | G | H | H | [illegible] |
| 2016H_P1-G9 | PHI10 | PHI3 | mother | P1-G9 | | H | G | H | H | [illegible] |
| 2016H_P1-G2 | PHI10 | PHI3 | mother | P1-G2 | | H | G | H | A | [illegible] |
| 2016H_P1-G5 | PHI10 | PHI3 | mother | P1-G5 | | H | G | H | A | [illegible] |
| 2016H_P1-F9 | PHI10 | PHI3 | mother | P1-F9 | | H | G | H | A | [illegible] |
| 2016H_P1-F3 | PHI10 | PHI3 | mother | P1-F4 | | H | G | H | H | [illegible] |
| 2016H_P1-F10 | PHI10 | PHI3 | mother | P1-F10 | | H | G | H | A | [illegible] |
| 2016H_P5-G3 | PHI9 | PHI3 | mother | P5-G3 | | G | G | A | H | [illegible] |
| 2016H_P5-E5 | PHI9 | PHI3 | mother | P5-E5 | | G | G | H | H | [illegible] |
| 2016H_P5-G4 | PHI9 | PHI3 | mother | P5-G4 | | G | G | A | H | [illegible] |
| 2016H_P5-F10 | PHI9 | PHI3 | mother | P5-F10 | | G | G | H | A | [illegible] |
| 2016H_P5-F2 | PHI9 | PHI3 | mother | P5-F2 | | G | G | H | H | [illegible] |
| 2016H_P5-E7 | PHI9 | PHI3 | mother | P5-E7 | | G | G | H | H | [illegible] |
| 2016H_P1-C9 | PHI2 | PHI3 | mother | P1-C9 | | H | H | H | H | [illegible] |
| 2016H_P1-C2 | PHI2 | PHI3 | mother | P1-C2 | | H | H | H | A | [illegible] |
| 2016H_P1-C6 | PHI2 | PHI3 | mother | P1-C6 | | H | H | H | H | [illegible] |
| 2016H_P1-A4 | PHI2 | PHI3 | mother | P1-A4 | | H | H | H | A | [illegible] |
| 2016H_P1-A9 | PHI2 | PHI3 | mother | P1-A9 | | H | H | H | H | [illegible] |
| 2016H_P1-A11 | PHI2 | PHI3 | mother | P1-A11 | | H | H | H | A | [illegible] |
| 2016H_P4-H4_2 | PHI5 | PHI3 | progeny | P4-H4 | | H | H | H | A | [illegible] |
| 2016H_P5-D5_1 | PHI5 | PHI3 | progeny | P5-D5 | | H | H | H | H | [illegible] |
| 2016H_P5-D5_2 | PHI5 | PHI3 | progeny | P5-D5 | | H | H | H | H | [illegible] |
| 2016H_P5-D5_3 | PHI5 | PHI3 | progeny | P5-D5 | | H | H | H | H | [illegible] |
| 2016H_P4-F3_1 | PHI5 | PHI3 | progeny | P4-F3 | | H | H | H | H | [illegible] |
| 2016H_P5-C9_1 | PHI5 | PHI3 | progeny | P5-C9 | | H | H | H | A | [illegible] |
| 2016H_P5-C9_2 | PHI5 | PHI3 | progeny | P5-C9 | | C | A | H | A | [illegible] |
| 2016H_P5-B6_1 | PHI5 | PHI3 | progeny | P5-B6 | | C | A | T | A | [illegible] |
| 2016H_P5-B6_2 | PHI5 | PHI3 | progeny | P5-B6 | | H | H | H | A | [illegible] |
| 2016H_P5-B5_1 | PHI5 | PHI3 | progeny | P5-B5 | | H | H | H | H | [illegible] |
| 2016H_P5-B5_2 | PHI5 | PHI3 | progeny | P5-B5 | | C | A | T | A | [illegible] |
| 2016H_P4-H1_1 | PHI5 | PHI3 | progeny | P4-H1 | | H | H | A | A | [illegible] |
| 2016H_P3-E5_1 | PHI6 | PHI3 | progeny | P3-E5 | | H | H | T | A | [illegible] |
| 2016H_P3-E11_1 | PHI6 | PHI3 | progeny | P3-E11 | | G | G | A | A | [illegible] |
| 2016H_P3-H10_1 | PHI6 | PHI3 | progeny | P3-H10 | | H | H | H | A | [illegible] |
| 2016H_P3-E3_2 | PHI6 | PHI3 | progeny | P3-E3 | | H | H | H | A | [illegible] |
| 2016H_P4-A9_1 | PHI6 | PHI3 | progeny | P4-A9 | | H | H | H | A | [illegible] |
| 2016H_P3-H7_1 | PHI6 | PHI3 | progeny | P3-H7 | | H | H | H | A | [illegible] |
| 2016H_P3-E4_3 | PHI6 | PHI3 | progeny | P3-E4 | | H | A | T | A | [illegible] |
| 2016H_P3-E4_6 | PHI6 | PHI3 | progeny | P3-E4 | | C | A | H | H | [illegible] |
| 2016H_P3-H9_1 | PHI6 | PHI3 | progeny | P3-H9 | | G | A | T | A | [illegible] |
| 2016H_P3-H9_2 | PHI6 | PHI3 | progeny | P3-H9 | | C | A | H | H | [illegible] |
| 2016H_P3-H9_3 | PHI6 | PHI3 | progeny | P3-H9 | | G | G | A | A | [illegible] |
| 2016H_P3-H9_4 | PHI6 | PHI3 | progeny | P3-H9 | | C | A | A | A | [illegible] |
| 2016H_P3-H9_5 | PHI6 | PHI3 | progeny | P3-H9 | | C | A | A | A | [illegible] |
| 2016H_P4-A6_2 | PHI6 | PHI3 | progeny | P4-A6 | | H | H | H | A | [illegible] |
| 2016H_P4-A6_3 | PHI6 | PHI3 | progeny | P4-A6 | | H | H | T | A | [illegible] |
| 2016H_P3-G11_1 | PHI6 | PHI3 | progeny | P3-G11 | | G | G | A | A | [illegible] |
| 2016H_P3-G11_2 | PHI6 | PHI3 | progeny | P3-G11 | | C | A | T | A | [illegible] |
| 2016H_P3-G11_3 | PHI6 | PHI3 | progeny | P3-G11 | | G | G | A | A | [illegible] |
| 2016H_P3-G11_4 | PHI6 | PHI3 | progeny | P3-G11 | | C | A | T | A | [illegible] |
| 2016H_P3-G11_5 | PHI6 | PHI3 | progeny | P3-G11 | | G | G | A | A | [illegible] |
| 2016H_P3-G11_6 | PHI6 | PHI3 | progeny | P3-G11 | | G | G | A | A | [illegible] |
| 2016H_P3-G11_7 | PHI6 | PHI3 | progeny | P3-G11 | | G | G | T | A | [illegible] |
| 2016H_P3-G11_8 | PHI6 | PHI3 | progeny | P3-G11 | | C | A | T | A | [illegible] |
| 2016H_P3-G11_9 | PHI6 | PHI3 | progeny | P3-G11 | | G | G | A | A | [illegible] |
| 2016H_P3-G12_2 | PHI6 | PHI3 | progeny | P3-G12 | | G | G | A | G | [illegible] |
| 2016H_P3-G12_3 | PHI6 | PHI3 | progeny | P3-G12 | | G | G | A | G | [illegible] |
| 2016H_P3-G12_4 | PHI6 | PHI3 | progeny | P3-G12 | | C | A | A | G | [illegible] |
| 2016H_P3-G12_5 | PHI6 | PHI3 | progeny | P3-G12 | | G | G | A | G | [illegible] |
| 2016H_P3-G12_6 | PHI6 | PHI3 | progeny | P3-G12 | | C | A | T | A | [illegible] |
| 2016H_P3-G12_8 | PHI6 | PHI3 | progeny | P3-G12 | | G | G | H | G | [illegible] |
| 2016H_P3-G12_9 | PHI6 | PHI3 | progeny | P3-G12 | | G | G | A | G | [illegible] |
| 2016H_P3-G12_10 | PHI6 | PHI3 | progeny | P3-G12 | | C | A | T | A | [illegible] |
| 2016H_P3-G12_11 | PHI6 | PHI3 | progeny | P3-G12 | | G | G | A | G | [illegible] |
| 2016H_P3-G12_12 | PHI6 | PHI3 | progeny | P3-G12 | | C | A | T | A | [illegible] |
| 2016H_P3-G12_13 | PHI6 | PHI3 | progeny | P3-G12 | | G | G | T | A | [illegible] |
| 2016H_P3-F1_1 | PHI6 | PHI3 | progeny | P3-F1 | | G | G | A | G | [illegible] |
| 2016H_P3-F1_2 | PHI6 | PHI3 | progeny | P3-F1 | | C | A | T | A | [illegible] |
| 2016H_P3-F1_3 | PHI6 | PHI3 | progeny | P3-F1 | | G | G | A | G | [illegible] |
| 2016H_P3-F1_4 | PHI6 | PHI3 | progeny | P3-F1 | | C | A | T | A | [illegible] |
| 2016H_P3-F1_5 | PHI6 | PHI3 | progeny | P3-F1 | | G | G | A | G | [illegible] |
| 2016H_P3-F1_6 | PHI6 | PHI3 | progeny | P3-F1 | | C | A | T | A | [illegible] |
| 2016H_P3-F1_7 | PHI6 | PHI3 | progeny | P3-F1 | | C | A | T | A | [illegible] |
| 2016H_P3-F1_8 | PHI6 | PHI3 | progeny | P3-F1 | | C | A | A | G | [illegible] |
| 2016H_P3-F1_9 | PHI6 | PHI3 | progeny | P3-F1 | | G | G | A | G | [illegible] |
| 2016H_P3-F1_10 | PHI6 | PHI3 | progeny | P3-F1 | | C | A | T | A | [illegible] |
| 2016H_P3-F1_11 | PHI6 | PHI3 | progeny | P3-F1 | | C | A | A | G | [illegible] |
| 2016H_P3-F1_12 | PHI6 | PHI3 | progeny | P3-F1 | | C | A | T | A | [illegible] |
| 2016H_P3-F1_14 | PHI6 | PHI3 | progeny | P3-F1 | | C | A | T | A | [illegible] |
| 2016H_P3-F1_15 | PHI6 | PHI3 | progeny | P3-F1 | | C | A | T | A | [illegible] |
| 2016H_P3-F1_16 | PHI6 | PHI3 | progeny | P3-F1 | | C | A | T | A | [illegible] |
| 2016H_P3-F1_17 | PHI6 | PHI3 | progeny | P3-F1 | | C | A | T | A | [illegible] |
| 2016H_P3-F11_3 | PHI6 | PHI3 | progeny | P3-F11 | | G | G | T | A | [illegible] |
| 2016H_P3-F11_4 | PHI6 | PHI3 | progeny | P3-F11 | | C | A | T | A | [illegible] |
| 2016H_P3-F11_5 | PHI6 | PHI3 | progeny | P3-F11 | | G | G | A | G | [illegible] |
| 2016H_P3-F11_7 | PHI6 | PHI3 | progeny | P3-F11 | | G | G | T | A | [illegible] |
| 2016H_P3-F11_8 | PHI6 | PHI3 | progeny | P3-F11 | | G | G | A | G | [illegible] |
| 2016H_P3-F11_11 | PHI6 | PHI3 | progeny | P3-F11 | | G | G | T | A | [illegible] |
| 2016H_P3-F11_14 | PHI6 | PHI3 | progeny | P3-F11 | | G | G | A | G | [illegible] |

**Table S8. Results of the SNP array analasis (continue)**

| Chr | 1 | 1 | 1 | 1 | 1 | 1 | 1 | 1 | 1 | 1 | 1 | 1 | 1 | 1 | 1 | 1 | 1 | 1 | 1 | 1 | 1 | 1 | 1 | 1 | 1 | 1 | 1 | 1 | 2 | 2 | 2 | 2 | 2 | 2 | 2 | 2 | 2 | 2 | 2 | 2 | 2 | 2 | 2 | 2 | 2 | 2 | 2 | 2 | 2 | 2 | 2 | 2 | 2 | 2 | 2 | 2 | 2 | 3 | 3 | 3 | 3 | 3 | 3 | 3 | 3 | 3 | 3 | 3 | 3 | 3 | 3 | 3 | 3 | 3 | 4 | 4 | 4 | 4 | 4 | 4 | 4 | 4 | 4 | 4 | 4 | 4 | 4 | 4 | 4 | 4 | 4 | 4 | 4 | 5 | 5 | 5 | 5 | 5 | 5 | 5 | 5 | 5 | 5 | 5 | 5 | 5 | 5 | 5 | 5 | 5 | 5 | 5 | 5 | 5 | 5 | 5 | 6 | 6 | 6 | 6 | 6 | 6 |
|---|---|---|---|---|---|---|---|---|---|---|---|---|---|---|---|---|---|---|---|---|---|---|---|---|---|---|---|---|---|---|---|---|---|---|---|---|---|---|---|---|---|---|---|---|---|---|---|---|---|---|---|---|---|---|---|---|---|---|---|---|---|---|---|---|---|---|---|---|---|---|---|---|---|---|---|---|---|---|---|---|---|---|---|---|---|---|---|---|---|---|---|---|---|---|---|---|---|---|---|---|---|---|---|---|---|---|---|---|---|---|---|---|---|---|---|---|---|---|---|---|---|
| Position, cM | 13 | 23 | 48 | 50 | 55 | 65 | 77 | 80 | 81 | 85 | 97 | 113 | 138 | 138 | 144 | 159 | 173 | 188 | 197 | 208 | 216 | 217 | 235 | 240 | 280 | 291 | 305 | 316 | 2.2 | 8.2 | 12 | 37 | 43 | 49 | 57 | 72 | 75 | 95 | 97 | 105 | 114 | 120 | 125 | 127 | 135 | 138 | 142 | 154 | 162 | 167 | 196 | 208 | 216 | 228 | 233 | 245 | 254 | 47 | 75 | 85 | 96 | 112 | 126 | 133 | 157 | 163 | 180 | 189 | 191 | 208 | 211 | 216 | 228 | 243 | 4.3 | 22 | 29 | 53 | 63 | 67 | 75 | 80 | 84 | 88 | 97 | 103 | 112 | 128 | 146 | 225 | 237 | 242 | 246 | 4.9 | 12 | 24 | 28 | 67 | 71 | 80 | 83 | 94 | 108 | 117 | 126 | 134 | 148 | 163 | 167 | 180 | 187 | 188 | 191 | 209 | 215 | 224 | 5.6 | 8.9 | 13 | 26 | 46 | 85 |

| plant_id | mom_gen | dad_gen | mother_progeny | family_ |
|---|---|---|---|---|
| 2016H_P3-F3_3 | PHI6 | PHI3 | progeny | P3-F3 |
| 2016H_P3-H8_5 | PHI6 | PHI3 | progeny | P3-H8 |
| 2016H_P3-G6_5 | PHI6 | PHI3 | progeny | P3-G6 |
| 2016H_P3-F6_4 | PHI6 | PHI3 | progeny | P3-F6 |
| 2016H_P3-F6_5 | PHI6 | PHI3 | progeny | P3-F6 |
| 2016H_P3-F6_6 | PHI6 | PHI3 | progeny | P3-F6 |
| 2016H_P3-G2_1 | PHI6 | PHI3 | progeny | P3-G2 |
| 2016H_P3-G2_2 | PHI6 | PHI3 | progeny | P3-G2 |
| 2016H_P3-F8_1 | PHI6 | PHI3 | progeny | P3-F8 |
| 2016H_P3-F8_2 | PHI6 | PHI3 | progeny | P3-F8 |
| 2016H_P3-G7_6 | PHI6 | PHI3 | progeny | P3-G7 |
| 2016H_P3-A12_1 | PHI7 | PHI3 | progeny | P3-A12 |
| 2016H_P3-A12_2 | PHI7 | PHI3 | progeny | P3-A12 |
| 2016H_P3-C4_1 | PHI7 | PHI3 | progeny | P3-C4 |
| 2016H_P3-A9_1 | PHI7 | PHI3 | progeny | P3-A9 |
| 2016H_P3-C2_1 | PHI7 | PHI3 | progeny | P3-C2 |
| 2016H_P2-H11_2 | PHI7 | PHI3 | progeny | P2-H11 |
| 2016H_P3-B2_1 | PHI7 | PHI3 | progeny | P3-B2 |
| 2016H_P3-B2_2 | PHI7 | PHI3 | progeny | P3-B2 |
| 2016H_P3-A12_3 | PHI7 | PHI3 | progeny | P3-A12 |
| 2016H_P3-A12_12 | PHI7 | PHI3 | progeny | P3-A12 |
| 2016H_P3-C9_2 | PHI7 | PHI3 | progeny | P3-C9 |
| 2016H_P3-A8_2 | PHI7 | PHI3 | progeny | P3-A8 |
| 2016H_P3-A8_3 | PHI7 | PHI3 | progeny | P3-A8 |
| 2016H_P3-A11_4 | PHI7 | PHI3 | progeny | P3-A11 |
| 2016H_P3-A11_5 | PHI7 | PHI3 | progeny | P3-A11 |
| 2016H_P3-A11_6 | PHI7 | PHI3 | progeny | P3-A11 |
| 2016H_P3-B9_2 | PHI7 | PHI3 | progeny | P3-B9 |
| 2016H_P2-C6_3 | PHI4 | PHI3 | progeny | P2-C6 |
| 2016H_P2-C9_1 | PHI4 | PHI3 | progeny | P2-C9 |
| 2016H_P2-C9_4 | PHI4 | PHI3 | progeny | P2-C9 |
| 2016H_P2-B7_1 | PHI4 | PHI3 | progeny | P2-B7 |
| 2016H_P2-B7_2 | PHI4 | PHI3 | progeny | P2-B7 |
| 2016H_P2-B7_3 | PHI4 | PHI3 | progeny | P2-B7 |
| 2016H_P2-B7_4 | PHI4 | PHI3 | progeny | P2-B7 |
| 2016H_P2-F3_2 | PHI4 | PHI3 | progeny | P2-F3 |
| 2016H_P2-F3_3 | PHI4 | PHI3 | progeny | P2-F3 |
| 2016H_P2-G6_1 | PHI4 | PHI3 | progeny | P2-G6 |
| 2016H_P2-G6_2 | PHI4 | PHI3 | progeny | P2-G6 |
| 2016H_P2-G6_3 | PHI4 | PHI3 | progeny | P2-G6 |
| 2016H_P2-G6_4 | PHI4 | PHI3 | progeny | P2-G6 |
| 2016H_P2-E7_1 | PHI4 | PHI3 | progeny | P2-E7 |
| 2016H_P2-E7_2 | PHI4 | PHI3 | progeny | P2-E7 |
| 2016H_P2-A12_1 | PHI4 | PHI3 | progeny | P2-A12 |
| 2016H_P2-A12_2 | PHI4 | PHI3 | progeny | P2-A12 |
| 2016H_P2-C1_4 | PHI4 | PHI3 | progeny | P2-C1 |
| 2016H_P2-C1_5 | PHI4 | PHI3 | progeny | P2-C1 |
| 2016H_P2-C1_6 | PHI4 | PHI3 | progeny | P2-C1 |
| 2016H_P2-E9_1 | PHI4 | PHI3 | progeny | P2-E9 |
| 2016H_P2-E9_3 | PHI4 | PHI3 | progeny | P2-E9 |
| 2016H_P2-E8_6 | PHI4 | PHI3 | progeny | P2-E8 |
| 2016H_P2-F11_1 | PHI4 | PHI3 | progeny | P2-F11 |
| 2016H_P2-F11_2 | PHI4 | PHI3 | progeny | P2-F11 |
| 2016H_P2-D6_4 | PHI4 | PHI3 | progeny | P2-D6 |
| 2016H_P2-D6_5 | PHI4 | PHI3 | progeny | P2-D6 |
| 2016H_P2-D6_6 | PHI4 | PHI3 | progeny | P2-D6 |
| 2016H_P2-D9_5 | PHI4 | PHI3 | progeny | P2-D9 |
| 2016H_P2-E1_1 | PHI4 | PHI3 | progeny | P2-E1 |
| 2016H_P2-E1_2 | PHI4 | PHI3 | progeny | P2-E1 |
| 2016H_P2-E1_3 | PHI4 | PHI3 | progeny | P2-E1 |
| 2016H_P2-D3_1 | PHI4 | PHI3 | progeny | P2-D3 |
| 2016H_P2-D3_3 | PHI4 | PHI3 | progeny | P2-D3 |
| 2016H_P2-D3_4 | PHI4 | PHI3 | progeny | P2-D3 |
| 2016H_P2-D3_5 | PHI4 | PHI3 | progeny | P2-D3 |
| 2016H_P2-D3_6 | PHI4 | PHI3 | progeny | P2-D3 |
| 2016H_P2-D2_1 | PHI4 | PHI3 | progeny | P2-D2 |
| 2016H_P2-D2_2 | PHI4 | PHI3 | progeny | P2-D2 |
| 2016H_P2-G8_2 | PHI4 | PHI3 | progeny | P2-G8 |
| 2016H_P2-G8_3 | PHI4 | PHI3 | progeny | P2-G8 |
| 2016H_P2-C7_1 | PHI4 | PHI3 | progeny | P2-C7 |
| 2016H_P2-C6_1 | PHI4 | PHI3 | progeny | P2-C6 |
| 2016H_P2-C6_2 | PHI4 | PHI3 | progeny | P2-C6 |
| 2016H_P1-G1_3 | PHI10 | PHI3 | progeny | P1-G1 |
| 2016H_P1-G1_4 | PHI10 | PHI3 | progeny | P1-G1 |
| 2016H_P1-G1_5 | PHI10 | PHI3 | progeny | P1-G1 |
| 2016H_P1-G1_6 | PHI10 | PHI3 | progeny | P1-G1 |
| 2016H_P1-F4_2 | PHI10 | PHI3 | progeny | P1-F4 |
| 2016H_P1-F4_3 | PHI10 | PHI3 | progeny | P1-F4 |
| 2016H_P1-F4_4 | PHI10 | PHI3 | progeny | P1-F4 |
| 2016H_P1-F5_1 | PHI10 | PHI3 | progeny | P1-F5 |
| 2016H_P1-F11_6 | PHI10 | PHI3 | progeny | P1-F11 |
| 2016H_P1-G9_11 | PHI10 | PHI3 | progeny | P1-G9 |
| 2016H_P1-G2_1 | PHI10 | PHI3 | progeny | P1-G2 |
| 2016H_P1-G2_2 | PHI10 | PHI3 | progeny | P1-G2 |
| 2016H_P1-G5_2 | PHI10 | PHI3 | progeny | P1-G5 |
| 2016H_P1-G5_4 | PHI10 | PHI3 | progeny | P1-G5 |
| 2016H_P1-F9_1 | PHI10 | PHI3 | progeny | P1-F9 |
| 2016H_P1-F9_2 | PHI10 | PHI3 | progeny | P1-F9 |
| 2016H_P1-F3_4 | PHI10 | PHI3 | progeny | P1-F3 |
| 2016H_P1-F3_9 | PHI10 | PHI3 | progeny | P1-F3 |
| 2016H_P1-F10_2 | PHI10 | PHI3 | progeny | P1-F10 |
| 2016H_P1-F10_6 | PHI10 | PHI3 | progeny | P1-F10 |
| 2016H_P1-F10_7 | PHI10 | PHI3 | progeny | P1-F10 |
| 2016H_P1-F10_9 | PHI10 | PHI3 | progeny | P1-F10 |
| 2016H_P5-G3_4 | PHI9 | PHI3 | progeny | P5-G3 |
| 2016H_P5-E5_1 | PHI9 | PHI3 | progeny | P5-E5 |
| 2016H_P5-E5_2 | PHI9 | PHI3 | progeny | P5-E5 |
| 2016H_P5-E5_3 | PHI9 | PHI3 | progeny | P5-E5 |
| 2016H_P5-E5_7 | PHI9 | PHI3 | progeny | P5-E5 |
| 2016H_P5-E5_9 | PHI9 | PHI3 | progeny | P5-E5 |
| 2016H_P5-E5_10 | PHI9 | PHI3 | progeny | P5-E5 |
| 2016H_P5-E5_11 | PHI9 | PHI3 | progeny | P5-E5 |
| 2016H_P5-G4_1 | PHI9 | PHI3 | progeny | P5-G4 |
| 2016H_P5-F10_1 | PHI9 | PHI3 | progeny | P5-F10 |
| 2016H_P5-F2_1 | PHI9 | PHI3 | progeny | P5-F2 |
| 2016H_P5-E7_3 | PHI9 | PHI3 | progeny | P5-E7 |
| 2016H_P1-C9_1 | PHI2 | PHI3 | progeny | P1-C9 |
| 2016H_P1-C9_2 | PHI2 | PHI3 | progeny | P1-C9 |
| 2016H_P1-C9_3 | PHI2 | PHI3 | progeny | P1-C9 |
| 2016H_P1-C2_3 | PHI2 | PHI3 | progeny | P1-C2 |
| 2016H_P1-C6_1 | PHI2 | PHI3 | progeny | P1-C6 |
| 2016H_P1-A4_1 | PHI2 | PHI3 | progeny | P1-A4 |
| 2016H_P1-A4_2 | PHI2 | PHI3 | progeny | P1-A4 |
| 2016H_P1-A4_4 | PHI2 | PHI3 | progeny | P1-A4 |
| 2016H_P1-A4_5 | PHI2 | PHI3 | progeny | P1-A4 |
| 2016H_P1-A4_9 | PHI2 | PHI3 | progeny | P1-A4 |
| 2016H_P1-A4_10 | PHI2 | PHI3 | progeny | P1-A4 |
| 2016H_P1-A9_1 | PHI2 | PHI3 | progeny | P1-A9 |
| 2016H_P1-A9_3 | PHI2 | PHI3 | progeny | P1-A9 |
| 2016H_P1-A11_1 | PHI2 | PHI3 | progeny | P1-A11 |
| 2016H_P1-A11_3 | PHI2 | PHI3 | progeny | P1-A11 |
| 2016H_P1-A11_4 | PHI2 | PHI3 | progeny | P1-A11 |
| 2016H_P1-A11_5 | PHI2 | PHI3 | progeny | P1-A11 |

plant_id – unique identifier of the plant
mom_gen – genetic backgroiund of mother in the hybred
dad_gen – genetic background of father in the hybred
family_ – identify mother and all the progeny plants from the same mother
mother_progeny – identify each plant as mother of the family or progeny
Chr – number of the chromosome, where marker is located
Distance – position of the marker on the corresponding chromosome in cM
H – heterozygous marker
A, T, G and C – marker homozygous for the corresponding nucleotide
EQV, LS, -9 , etc. – missing values

**Table S8. Results of the SNP array analasis (continue)**

| | | | | | Chr | 6 | 6 | 6 | 6 | 6 | 6 | 6 | 7 | 7 | 7 | 7 | 7 | 7 | 7 | 7 | 7 | 7 | 7 | 7 | 7 | 7 | 7 | 7 | 7 | 7 | 7 | 7 | 7 | 7 | 7 | 7 | 7 | 8 | 8 | 8 | 8 | 8 | 8 | 8 | 9 | 9 | 9 | 9 | 9 | 9 | 9 | 9 | 9 | 9 | 10 | 10 | 10 | 10 | 10 | 10 | 10 | 10 | 10 | 10 | 10 | 10 | 10 | 10 | 10 | 10 | 10 | 10 | 10 | 10 | 10 |
|---|---|---|---|---|---|---|---|---|---|---|---|---|---|---|---|---|---|---|---|---|---|---|---|---|---|---|---|---|---|---|---|---|---|---|---|---|---|---|---|---|---|---|---|---|---|---|---|---|---|---|---|---|---|---|---|---|---|---|---|---|---|---|---|---|---|---|---|---|---|---|---|---|---|---|---|
| | | | | | Position, cM | 89 | 130 | 142 | 151 | 155 | 180 | 180 | 8 | 17 | 32 | 37 | 52 | 64 | 75 | 74 | 85 | 92 | 109 | 111 | 130 | 135 | 141 | 150 | 162 | 177 | 182 | 186 | 192 | 196 | 201 | 205 | 205 | 40 | 50 | 113 | 164 | 177 | 187 | 195 | 35 | 39 | 45 | 60 | 127 | 143 | 159 | 166 | 178 | 180 | 11 | 13 | 18 | 27 | 31 | 40 | 48 | 54 | 70 | 76 | 87 | 91 | 103 | 108 | 112 | 121 | 130 | 143 | 154 | 162 | 161 |
| plant_id | mom_gen | dad_gen | mother_progeny | family_ | | | | | | | | | | | | | | | | | | | | | | | | | | | | | | | | | | | | | | | | | | | | | | | | | | | | | | | | | | | | | | | | | | | | | | | |
| 2016H_P4-H4 | PHI5 | PHI3 | mother | P4-H4 | | G | H | T | C | G | G | A | T | C | H | H | H | T | A | H | H | H | C | H | C | C | T | G | G | H | C | H | H | H | C | C | H | G | H | C | H | H | H | H | C | G | G | C | H | H | T | H | H | C | T | H | H | G | H | H | C | H | H | H | C | H | H | H | H | T | A | H | H | H | G |
| 2016H_P5-D5 | PHI5 | PHI3 | mother | P5-D5 | | G | H | T | C | H | G | A | T | C | H | H | H | T | A | H | H | H | C | H | C | C | T | G | G | H | C | H | A | G | C | C | H | G | H | C | H | H | H | H | C | G | H | C | H | H | T | H | H | C | T | H | H | G | H | H | C | H | H | H | C | H | H | H | H | T | A | H | H | H | H |
| 2016H_P4-F3 | PHI5 | PHI3 | mother | P4-F3 | | G | H | H | C | H | G | A | T | C | H | H | H | T | A | H | H | H | C | H | C | C | T | G | G | H | C | H | A | G | C | C | H | G | H | C | H | H | H | H | C | G | H | C | H | H | T | H | H | C | T | H | H | G | H | H | C | H | H | H | C | H | H | H | H | T | A | H | H | H | G |
| 2016H_P5-C9 | PHI5 | PHI3 | mother | P5-C9 | | G | H | T | C | G | G | A | T | C | H | H | H | T | A | H | H | H | C | H | C | C | T | G | G | H | C | C | A | G | C | H | C | G | H | C | H | H | H | H | C | G | H | C | H | H | H | H | H | T | H | H | H | G | H | H | C | H | H | H | C | H | H | H | H | T | A | H | H | H | H |
| 2016H_P5-B5 | PHI5 | PHI3 | mother | P5-B5 | | G | H | H | C | H | G | A | T | C | H | H | H | T | A | H | H | H | C | H | C | C | T | G | G | H | C | C | A | G | C | H | C | G | H | C | H | H | H | H | C | G | H | C | H | H | T | H | H | C | T | H | H | G | H | H | C | H | H | H | C | H | H | H | H | T | A | H | H | H | H |
| 2016H_P4-H1 | PHI5 | PHI3 | mother | P4-H1 | | G | C | T | C | G | G | A | T | C | H | H | H | T | A | H | H | H | C | H | C | C | T | G | G | H | C | C | H | H | C | C | H | G | H | C | H | H | H | H | C | G | H | C | H | H | H | H | H | A | T | H | H | G | H | H | C | H | H | H | C | H | H | H | H | T | A | H | H | H | G |
| 2016H_P3-E5 | PHI6 | PHI3 | mother | P3-E5 | | H | H | H | H | G | G | A | T | C | H | H | H | H | H | C | A | H | H | T | C | H | T | G | G | H | C | C | A | G | H | C | C | G | H | C | A | G | T | A | H | H | A | C | C | G | T | H | T | H | H | G | H | H | G | C | C | H | G | A | C | H | A | H | A | T | H | C | G | H | H |
| 2016H_P3-E11 | PHI6 | PHI3 | mother | P3-E11 | | H | H | H | H | G | G | A | T | C | H | H | H | H | H | C | A | H | H | T | C | H | T | G | G | C | C | C | H | H | H | C | C | G | H | C | A | G | T | A | H | H | A | C | C | G | T | H | H | H | H | G | H | H | G | C | C | H | G | A | C | H | H | C | C | T | H | C | G | H | H |
| 2016H_P3-H10 | PHI6 | PHI3 | mother | P3-H10 | | H | H | H | H | G | G | A | T | C | H | H | H | H | H | C | A | H | H | T | C | H | T | G | G | C | C | C | A | G | C | H | C | G | H | C | A | G | T | A | H | H | H | C | C | G | T | H | H | H | H | G | H | H | G | C | C | H | H | A | C | H | A | H | H | T | H | C | G | H | G |
| 2016H_P3-E3 | PHI6 | PHI3 | mother | P3-E3 | | H | H | H | H | G | G | A | T | C | H | H | H | H | H | C | A | H | H | T | C | H | T | G | G | H | C | H | A | G | H | C | C | T | T | C | A | G | T | A | H | H | H | C | C | G | T | H | H | H | H | G | H | H | G | C | C | H | G | A | C | H | A | H | H | T | C | C | G | H | G |
| 2016H_P4-A9 | PHI6 | PHI3 | mother | P4-A9 | | H | H | H | H | G | G | A | T | C | H | H | H | H | H | C | A | H | H | T | C | H | T | G | G | H | C | C | H | G | H | C | C | G | H | C | A | G | T | A | H | H | H | C | C | G | T | H | H | H | H | G | H | H | G | C | C | H | G | A | C | H | A | H | A | T | C | C | G | H | H |
| 2016H_P3-H7 | PHI6 | PHI3 | mother | P3-H7 | | H | H | H | H | G | G | A | T | C | H | H | H | H | H | C | A | H | H | T | C | H | T | G | G | C | C | H | A | G | H | C | C | G | H | C | A | G | T | A | H | H | A | C | C | G | T | H | T | H | H | G | H | H | G | C | C | H | G | A | C | T | H | C | H | T | H | C | G | H | G |
| 2016H_P3-E4 | PHI6 | PHI3 | mother | P3-E4 | | G | C | H | C | H | H | H | T | C | H | H | H | H | H | C | A | H | H | T | C | H | T | G | G | H | C | C | H | H | C | H | C | G | H | C | A | G | T | A | H | H | A | C | C | G | T | H | T | C | H | G | H | H | G | C | C | H | G | A | C | T | H | C | C | T | C | C | G | H | H |
| 2016H_P3-H9 | PHI6 | PHI3 | mother | P3-H9 | | H | H | H | H | G | G | A | T | C | H | H | H | H | H | C | A | H | H | T | C | H | T | G | G | C | C | H | H | H | C | C | C | G | T | C | A | G | T | A | H | H | A | C | C | G | T | H | H | H | C | H | G | H | G | C | C | H | G | A | C | T | H | C | H | T | H | C | G | H | G |
| 2016H_P4-A6 | PHI6 | PHI3 | mother | P4-A6 | | H | H | H | H | G | G | A | T | C | H | H | H | H | H | C | A | H | H | T | C | H | T | G | G | C | C | C | A | G | H | C | C | G | H | C | A | G | T | A | H | H | A | C | C | G | T | H | H | T | C | H | G | H | G | C | C | H | G | A | C | H | A | H | H | T | C | C | G | H | G |
| 2016H_P3-G11 | PHI6 | PHI3 | mother | P3-G11 | | H | H | H | H | G | G | A | T | C | H | H | H | H | H | C | A | H | H | T | C | H | T | G | G | C | C | H | H | H | C | C | C | G | H | C | A | G | T | A | H | H | A | C | C | G | T | H | T | A | H | G | H | H | G | C | C | H | G | A | C | H | A | H | A | T | H | C | G | H | H |
| 2016H_P3-G12 | PHI6 | PHI3 | mother | P3-G12 | | H | H | H | H | G | G | A | T | C | H | H | H | H | H | C | A | H | H | T | C | H | T | G | G | C | C | H | A | G | H | C | C | G | H | C | A | G | T | A | H | H | A | C | C | G | T | H | T | H | H | G | H | G | G | C | C | H | G | A | C | H | A | H | H | T | H | C | G | H | G |
| 2016H_P3-F1 | PHI6 | PHI3 | mother | P3-F1 | | H | H | H | H | G | G | A | T | C | H | H | H | H | H | C | A | H | H | T | C | H | T | G | G | C | C | C | A | G | H | C | C | G | H | C | A | G | T | A | H | H | H | C | C | G | T | H | T | A | H | G | H | H | G | C | C | H | G | A | C | H | A | H | A | T | H | C | G | H | G |
| 2016H_P3-F11 | PHI6 | PHI3 | mother | P3-F11 | | H | H | H | H | G | G | A | T | C | H | H | H | H | H | C | A | H | H | T | C | H | T | G | G | C | C | C | A | G | H | C | C | G | H | C | A | G | T | A | H | H | H | C | C | G | T | H | T | C | H | G | H | H | G | C | C | H | G | A | C | H | A | H | H | T | H | C | G | H | G |
| 2016H_P3-F3 | PHI6 | PHI3 | mother | P3-F3 | | H | H | H | H | G | G | A | T | C | H | H | H | H | H | C | A | H | H | T | C | H | T | G | G | C | C | H | A | G | H | C | C | G | H | C | A | G | T | A | H | H | H | C | C | G | T | H | H | H | H | G | H | H | G | C | C | H | H | A | C | H | A | H | H | T | C | C | G | H | G |
| 2016H_P3-H8 | PHI6 | PHI3 | mother | P3-H8 | | H | H | H | H | G | G | A | T | C | H | H | H | H | H | C | A | H | H | T | C | H | T | G | G | C | C | H | A | G | H | C | C | G | H | C | A | G | T | A | H | H | A | C | C | G | T | H | T | H | H | G | H | H | G | C | C | H | G | A | C | T | H | C | H | T | H | C | G | H | G |
| 2016H_P3-G6 | PHI6 | PHI3 | mother | P3-G6 | | H | H | H | C | H | H | H | T | C | H | H | H | H | H | C | A | H | H | T | C | H | T | G | G | C | C | H | H | H | C | C | C | G | H | C | A | G | T | A | H | H | A | C | C | G | T | H | T | C | H | G | H | G | G | C | C | H | G | A | C | T | H | H | A | T | C | C | G | H | G |
| 2016H_P3-F6 | PHI6 | PHI3 | mother | P3-F6 | | H | H | H | H | G | G | A | T | C | H | H | H | H | H | C | A | H | H | T | C | H | T | G | G | C | C | C | A | G | H | C | C | G | H | C | A | G | T | A | H | H | H | C | C | G | T | H | T | H | H | G | H | H | G | C | C | H | G | A | C | H | A | C | C | T | H | C | G | H | H |
| 2016H_P3-G2 | PHI6 | PHI3 | mother | P3-G2 | | H | H | H | H | G | G | A | T | C | H | H | H | H | H | C | A | H | H | T | C | H | T | G | G | C | C | H | A | G | H | C | C | G | H | C | A | G | T | A | H | H | H | C | C | G | T | H | H | A | H | G | H | H | G | C | C | H | G | A | C | H | A | H | A | T | H | C | G | H | G |
| 2016H_P3-F8 | PHI6 | PHI3 | mother | P3-F8 | | H | H | H | C | H | H | H | T | C | H | H | H | H | H | C | A | H | H | T | C | H | T | G | G | C | C | H | H | H | C | C | C | G | H | C | A | G | T | A | H | H | A | C | C | G | T | H | H | A | H | G | H | H | G | C | C | H | G | A | C | T | A | H | A | T | C | C | G | H | H |
| 2016H_P3-G7 | PHI6 | PHI3 | mother | P3-G7 | | H | H | H | H | G | G | A | T | C | H | H | H | H | H | C | A | H | H | T | C | H | T | G | G | C | C | H | A | G | H | C | C | G | H | C | A | G | T | A | H | H | A | C | C | G | T | H | H | H | H | G | H | H | G | C | C | H | G | A | C | H | A | H | H | T | H | C | G | H | H |
| 2016H_P3-C4 | PHI7 | PHI3 | mother | P3-C4 | | G | H | T | C | H | H | H | T | C | H | H | H | H | H | C | A | H | C | H | H | C | T | G | G | H | C | C | A | G | H | C | C | G | T | C | A | G | T | A | C | G | H | H | H | H | G | H | H | H | T | G | G | G | H | C | C | H | G | A | C | H | A | H | H | T | A | C | G | G | G |
| 2016H_P3-A9 | PHI7 | PHI3 | mother | P3-A9 | | G | H | T | C | H | H | H | T | C | H | H | H | H | H | C | A | H | C | H | H | C | T | G | G | H | C | C | A | G | H | C | C | G | T | C | A | G | T | A | C | G | H | H | H | H | G | H | A | T | G | G | G | H | C | C | C | H | H | A | C | H | A | H | H | T | H | C | G | G | G |
| 2016H_P3-C2 | PHI7 | PHI3 | mother | P3-C2 | | G | H | T | C | H | H | H | T | C | H | H | H | H | H | C | A | H | C | H | H | C | T | G | G | H | C | C | A | G | H | C | C | G | T | C | A | G | T | A | C | G | G | H | H | H | H | G | A | T | G | G | G | H | C | C | C | H | H | A | C | H | A | H | H | T | H | C | G | G | G |
| 2016H_P2-H11 | PHI7 | PHI3 | mother | P2-H11 | | G | H | T | C | H | H | H | T | C | H | H | H | H | H | C | A | H | C | H | H | C | T | G | G | H | C | C | A | G | H | C | C | G | T | C | A | G | T | A | C | G | H | H | H | H | H | G | H | H | T | G | G | G | H | C | C | H | H | A | C | H | A | H | H | T | A | C | G | G | G |
| 2016H_P3-B2 | PHI7 | PHI3 | mother | P3-B2 | | G | H | T | C | H | H | H | T | C | H | H | H | H | H | C | A | H | C | H | H | C | T | G | G | H | C | H | A | G | H | C | C | G | T | C | A | G | T | A | C | G | H | H | H | H | G | H | H | H | T | G | G | G | H | C | C | H | H | A | C | H | A | H | H | T | A | C | G | G | G |
| 2016H_P3-C9 | PHI7 | PHI3 | mother | P3-C9 | | G | H | T | C | H | H | H | T | C | H | H | H | H | H | C | A | H | C | H | H | C | T | G | G | H | C | C | A | G | H | C | C | G | T | C | A | G | T | A | C | G | H | H | H | H | G | H | H | H | T | G | G | G | H | C | C | H | H | A | C | H | A | H | H | T | A | C | G | G | G |
| 2016H_P3-A8 | PHI7 | PHI3 | mother | P3-A8 | | G | H | T | C | H | H | H | T | C | H | H | H | H | H | C | A | H | C | H | H | C | T | G | G | H | C | C | A | G | H | C | C | G | T | C | A | G | T | A | C | G | H | H | H | H | G | H | H | H | T | G | G | G | H | C | C | H | H | A | C | H | A | H | H | T | A | C | G | G | G |
| 2016H_P3-A11 | PHI7 | PHI3 | mother | P3-A11 | | G | H | T | C | H | H | H | T | C | H | H | H | H | H | C | A | H | C | H | H | C | T | G | G | C | H | H | A | G | H | C | C | G | T | C | A | G | T | A | C | G | H | H | H | H | G | H | H | H | T | G | G | G | H | C | C | H | H | A | C | H | A | H | H | T | C | C | G | G | G |
| 2016H_P3-B9 | PHI7 | PHI3 | mother | P3-B9 | | G | H | T | C | H | H | H | T | C | H | H | H | H | H | C | A | H | C | H | H | C | T | G | G | C | H | H | A | G | H | C | C | G | T | C | H | G | T | A | C | G | G | H | H | H | H | G | C | A | T | G | G | G | H | C | C | H | G | A | C | H | A | H | H | T | H | C | G | G | G |
| 2016H_P3-A12 | PHI7 | PHI3 | mother | P3-A12 | | G | H | T | C | H | H | H | T | C | H | H | H | H | H | C | A | H | C | H | H | C | T | G | G | C | H | H | A | G | H | C | C | G | T | C | A | G | T | A | C | G | G | H | H | H | H | G | H | H | T | G | G | G | H | C | C | H | H | A | C | H | A | H | H | T | A | C | G | G | G |
| 2016H_P2-C9 | PHI4 | PHI3 | mother | P2-C9 | | G | H | T | C | H | H | H | T | H | H | H | H | H | H | H | A | H | H | T | H | H | H | G | G | H | C | C | H | H | C | C | C | G | T | H | H | G | T | A | C | H | G | C | H | H | H | T | H | T | G | G | H | G | C | C | H | H | C | A | C | H | A | H | A | T | A | H | G | G | H |
| 2016H_P2-B7 | PHI4 | PHI3 | mother | P2-B7 | | G | H | T | C | H | H | H | T | H | H | H | H | H | H | H | A | H | H | T | H | H | H | G | G | H | C | C | H | H | C | C | C | G | T | H | H | G | T | A | C | H | H | C | H | H | H | T | H | T | G | G | H | G | C | C | H | H | C | A | C | G | A | H | A | T | H | H | G | G | H |
| 2016H_P2-F3 | PHI4 | PHI3 | mother | P2-F3 | | G | H | T | C | H | H | H | T | H | H | H | H | H | T | H | A | H | H | T | H | H | H | G | G | H | C | C | A | G | C | H | C | G | T | H | H | G | T | A | C | H | H | C | H | H | H | T | H | T | G | G | H | G | C | C | H | H | C | A | C | G | A | H | A | T | A | H | G | G | H |
| 2016H_P2-G6 | PHI4 | PHI3 | mother | P2-G6 | | G | H | T | C | H | H | H | T | H | H | H | H | H | H | H | A | H | H | T | H | H | H | G | G | H | C | C | H | H | C | C | C | G | T | H | H | G | T | A | C | H | G | C | H | H | H | T | H | T | G | G | H | G | C | C | H | H | C | A | C | G | A | H | A | T | H | H | G | G | H |
| 2016H_P2-E7 | PHI4 | PHI3 | mother | P2-E7 | | G | H | T | C | H | H | H | T | H | H | H | H | H | H | H | A | H | H | T | C | H | H | H | G | H | C | C | H | H | C | C | C | G | T | H | H | G | T | A | C | H | G | C | H | H | H | H | H | H | G | G | H | G | C | C | H | H | C | A | C | G | A | H | A | T | A | H | G | G | H |
| 2016H_P2-A12 | PHI4 | PHI3 | mother | P2-A12 | | G | C | H | C | H | H | H | T | H | H | H | H | H | T | H | A | H | H | T | H | H | H | G | G | H | C | C | A | G | H | C | C | G | T | H | H | G | T | A | C | H | H | C | H | H | T | H | T | C | G | G | H | G | C | C | H | H | C | A | C | G | A | H | A | T | A | H | G | G | H |
| 2016H_P2-C1 | PHI4 | PHI3 | mother | P2-C1 | | G | H | T | C | H | H | H | T | H | H | H | H | H | H | H | A | H | H | T | H | H | H | G | G | H | C | C | H | H | C | C | C | G | T | H | H | G | T | A | C | H | H | C | H | H | T | H | H | H | G | G | H | G | C | C | H | H | C | A | C | G | A | H | A | T | H | H | G | G | H |
| 2016H_P2-E9 | PHI4 | PHI3 | mother | P2-E9 | | G | H | T | C | H | H | H | T | H | H | H | H | H | H | H | A | H | H | T | H | H | H | G | G | H | C | C | H | H | C | C | C | G | T | H | H | G | T | A | C | H | H | C | H | H | T | H | H | C | G | G | H | G | C | C | H | H | C | A | C | G | A | H | A | T | H | H | G | G | H |
| 2016H_P2-E8 | PHI4 | PHI3 | mother | P2-E8 | | G | C | H | C | H | H | H | T | H | H | H | H | H | H | H | A | H | H | T | H | H | H | G | G | H | C | C | A | G | C | C | C | G | T | H | H | G | T | A | C | H | G | C | H | H | T | H | H | C | G | G | H | G | C | C | H | H | C | A | C | G | A | H | A | T | A | H | G | G | H |
| 2016H_P2-F11 | PHI4 | PHI3 | mother | P2-F11 | | G | H | T | C | H | H | H | T | H | H | H | H | H | H | H | A | H | H | T | H | H | H | G | G | H | C | C | H | H | C | C | C | G | T | H | H | G | T | A | C | H | H | C | H | H | T | H | T | C | G | G | H | G | C | C | H | H | C | A | C | G | A | H | A | T | H | H | G | G | H |
| 2016H_P2-D6 | PHI4 | PHI3 | mother | P2-D6 | | G | C | H | C | H | H | H | T | H | H | H | H | H | H | H | A | H | H | T | H | H | H | G | G | H | C | C | H | H | C | C | C | G | T | H | H | G | T | A | C | H | H | C | H | H | T | H | H | C | G | G | H | G | C | C | H | H | C | A | C | H | A | H | A | T | H | H | G | G | H |
| 2016H_P2-D9 | PHI4 | PHI3 | mother | P2-D9 | | G | H | T | C | H | H | H | T | H | H | H | H | H | H | H | A | H | H | T | H | H | H | G | G | H | C | C | H | H | C | C | C | G | T | H | H | G | T | A | C | H | H | G | C | H | H | H | T | C | G | G | H | G | C | C | H | H | C | A | C | G | A | H | A | T | H | H | G | G | H |
| 2016H_P2-E1 | PHI4 | PHI3 | mother | P2-E1 | | G | H | T | C | H | H | H | T | H | H | H | H | H | H | H | A | H | H | T | C | H | H | H | G | H | C | C | H | H | C | C | C | G | T | H | H | G | T | A | C | H | H | C | H | H | T | H | H | C | G | G | H | G | C | C | H | H | C | A | C | G | A | H | A | T | A | H | G | G | H |
| 2016H_P2-D3 | PHI4 | PHI3 | mother | P2-D3 | | G | H | H | C | H | H | H | T | H | H | H | H | H | H | H | A | H | H | T | H | H | H | G | G | H | C | C | H | H | C | C | C | G | T | H | H | G | T | A | C | H | H | C | H | H | T | H | H | H | G | G | H | G | C | C | H | H | C | A | C | G | A | H | A | T | H | H | G | G | H |
| 2016H_P2-D2 | PHI4 | PHI3 | mother | P2-D2 | | G | H | T | C | H | H | H | T | H | H | H | H | H | H | H | A | H | H | T | C | H | H | H | G | H | C | C | H | H | C | C | C | G | T | H | H | G | T | A | C | H | H | C | H | H | T | H | H | C | G | G | H | G | C | C | H | H | C | A | C | G | A | H | A | T | H | H | G | G | H |
| 2016H_P2-G8 | PHI4 | PHI3 | mother | P2-G8 | | G | C | H | C | H | H | H | T | H | H | H | H | H | T | H | A | H | H | T | H | H | H | G | G | H | C | C | A | G | C | H | C | G | T | H | H | G | T | A | C | H | H | C | H | H | T | H | H | H | G | G | H | G | C | C | H | H | C | A | C | G | A | H | A | T | A | H | G | G | H |
| 2016H_P2-C7 | PHI4 | PHI3 | mother | P2-C7 | | G | H | T | C | H | H | H | T | H | H | H | H | H | H | H | A | H | H | T | C | H | H | H | G | H | C | C | H | H | C | C | C | G | T | H | H | G | T | A | C | H | G | C | H | H | T | H | T | C | G | G | H | G | C | C | H | H | C | A | C | G | A | H | A | T | A | H | G | G | H |
| 2016H_P2-C6 | PHI4 | PHI3 | mother | P2-C6 | | G | C | H | C | H | H | H | T | H | H | H | H | H | T | H | A | H | H | T | H | H | H | G | G | H | C | C | H | H | C | C | C | G | T | H | H | G | T | A | C | H | H | C | H | H | T | H | T | C | G | G | H | G | C | C | H | H | C | A | C | G | A | H | A | T | A | H | G | G | H |
| 2016H_P1-G1 | PHI10 | PHI3 | mother | P1-G1 | | G | H | T | H | H | H | H | T | H | H | H | H | G | H | H | A | G | H | T | H | H | H | G | G | H | C | C | A | H | C | H | C | G | T | C | H | G | H | A | C | H | H | C | H | H | H | H | C | A | G | G | H | G | H | H | H | H | H | H | H | H | A | H | H | T | C | C | H | G | H |
| 2016H_P1-F4 | PHI10 | PHI3 | mother | P1-F4 | | H | H | H | H | H | H | H | H | H | H | H | H | H | H | H | H | H | H | H | H | H | H | H | H | H | H | H | H | H | H | H | H | H | H | H | H | H | H | H | H | H | H | H | H | H | H | H | H | H | H | H | H | H | H | H | H | H | H | H | H | H | H | H | H | T | H | H | H | H | H |
| 2016H_P1-F5 | PHI10 | PHI3 | mother | P1-F5 | | G | H | T | H | H | H | H | T | H | A | H | G | H | H | H | A | G | H | T | H | H | H | G | G | H | C | C | A | H | C | H | C | G | T | C | H | G | H | A | C | H | A | C | H | H | H | H | C | A | G | G | H | G | H | H | H | H | H | C | H | H | A | H | A | T | C | C | H | G | H |
| 2016H_P1-F11 | PHI10 | PHI3 | mother | P1-F11 | | G | H | T | H | H | H | H | T | H | A | H | G | H | H | H | A | G | H | T | C | H | H | G | G | H | C | C | A | H | C | H | C | G | T | C | H | G | H | A | C | H | H | C | H | H | H | H | C | A | G | G | H | G | H | H | H | H | H | H | H | H | A | H | A | T | C | C | H | G | H |
| 2016H_P1-G9 | PHI10 | PHI3 | mother | P1-G9 | | G | H | T | H | H | H | H | T | H | A | H | H | T | H | H | A | H | H | T | H | H | H | G | G | H | C | C | A | H | C | H | C | G | T | C | H | G | H | A | C | H | A | C | H | H | H | H | C | A | G | G | H | G | H | H | H | H | H | H | H | H | A | H | A | T | H | C | H | G | H |
| 2016H_P1-G2 | PHI10 | PHI3 | mother | P1-G2 | | G | H | T | H | H | H | H | T | H | H | H | H | H | H | H | A | G | H | T | C | H | H | G | G | H | C | C | A | H | C | H | C | G | T | C | H | G | H | A | C | H | H | C | H | H | H | H | C | A | G | G | H | G | H | H | H | H | H | H | H | H | A | H | H | T | C | C | H | G | H |
| 2016H_P1-G5 | PHI10 | PHI3 | mother | P1-G5 | | G | H | T | H | H | H | H | T | H | A | H | H | T | H | H | A | H | H | T | H | H | H | G | G | H | C | C | A | H | C | H | C | G | T | C | H | G | H | A | C | H | H | C | H | H | H | H | C | A | G | G | H | G | H | H | H | H | H | H | H | H | A | H | H | T | H | C | H | G | H |
| 2016H_P1-F9 | PHI10 | PHI3 | mother | P1-F9 | | G | H | T | H | H | H | H | T | H | H | H | G | H | H | H | A | G | H | T | C | H | H | G | G | H | C | C | A | H | C | H | C | G | T | C | H | G | H | A | C | H | H | C | H | G | H | H | H | H | G | G | H | G | H | H | H | H | H | H | H | H | A | H | A | T | C | C | H | G | H |
| 2016H_P1-F3 | PHI10 | PHI3 | mother | P1-F4 | | G | H | T | H | H | H | H | T | H | H | H | G | H | H | H | A | G | H | T | C | H | H | G | G | H | C | C | A | H | C | H | C | G | T | C | H | G | H | A | C | H | A | C | H | G | H | H | H | H | G | G | H | G | H | H | C | H | H | H | H | H | A | H | H | T | C | C | H | G | H |
| 2016H_P1-F10 | PHI10 | PHI3 | mother | P1-F10 | | G | H | T | H | H | H | H | T | H | H | H | G | H | H | H | A | G | H | T | C | H | H | G | G | H | C | C | A | H | C | H | C | G | T | C | H | G | H | A | C | H | A | C | H | G | T | H | T | H | G | G | H | G | H | H | H | H | H | H | H | H | A | H | H | T | H | C | H | G | H |
| 2016H_P5-G3 | PHI9 | PHI3 | mother | P5-G3 | | G | H | T | C | H | H | H | H | H | A | T | G | H | A | H | H | H | H | H | T | C | H | H | H | G | C | H | G | A | C | C | C | T | T | H | A | G | T | A | C | H | A | C | H | G | T | H | C | A | H | G | G | H | G | C | C | A | H | H | C | G | G | H | H | C | C | C | G | H | G |
| 2016H_P5-E5 | PHI9 | PHI3 | mother | P5-E5 | | G | H | T | C | H | H | H | H | H | A | T | G | H | A | H | H | H | H | H | T | C | H | H | H | G | C | C | G | A | C | C | C | T | T | H | A | G | T | A | C | H | A | C | H | G | T | H | C | A | H | G | G | H | G | C | C | A | H | H | C | G | H | A | A | H | C | C | G | H | G |
| 2016H_P5-G4 | PHI9 | PHI3 | mother | P5-G4 | | G | H | T | C | H | H | H | H | H | A | T | G | H | A | H | H | H | H | H | T | C | H | H | H | G | C | H | H | C | C | C | C | T | T | H | H | G | T | A | C | H | A | C | H | G | T | H | C | A | H | G | G | H | G | C | C | A | H | H | C | G | G | A | A | H | C | C | G | H | G |
| 2016H_P5-F10 | PHI9 | PHI3 | mother | P5-F10 | | G | H | T | C | H | H | H | H | H | A | T | G | H | A | H | H | H | H | H | T | C | H | H | H | G | H | C | G | A | C | C | C | T | T | H | A | G | T | A | C | H | A | C | H | G | T | H | H | H | H | G | G | H | G | C | C | A | H | H | C | G | H | A | A | H | C | C | G | H | G |
| 2016H_P5-F2 | PHI9 | PHI3 | mother | P5-F2 | | G | H | H | C | H | H | H | H | H | A | T | G | H | A | H | H | H | H | H | T | C | H | H | H | G | C | C | G | A | C | C | C | T | T | H | A | G | T | A | C | H | H | C | H | G | T | H | H | H | H | G | G | H | G | C | C | A | H | H | C | H | G | H | H | H | C | C | G | H | G |
| 2016H_P5-E7 | PHI9 | PHI3 | mother | P5-E7 | | G | H | H | C | H | H | H | H | H | A | T | G | H | A | H | H | H | H | H | T | C | H | H | H | G | H | C | G | A | C | C | C | T | T | H | A | G | T | A | C | H | H | C | H | G | T | H | H | H | H | G | G | H | G | C | C | A | H | H | C | H | H | A | A | H | C | C | G | H | G |
| 2016H_P1-C9 | PHI2 | PHI3 | mother | P1-C9 | | G | C | H | C | H | H | H | H | H | A | H | H | T | H | H | A | H | H | T | C | H | H | H | H | H | C | C | H | H | C | H | C | T | T | H | A | G | H | A | H | H | H | C | H | H | H | H | H | H | G | G | H | G | H | C | A | H | A | C | T | H | C | H | T | H | H | H | H | H | H |
| 2016H_P1-C2 | PHI2 | PHI3 | mother | P1-C2 | | G | C | H | C | H | H | H | H | H | A | H | H | T | H | H | A | H | H | T | C | H | H | H | H | C | C | C | H | H | C | H | C | T | T | H | A | G | H | A | H | H | H | C | H | H | H | C | A | H | G | G | H | G | H | H | A | H | A | C | T | H | C | H | T | H | H | H | H | H | H |
| 2016H_P1-C6 | PHI2 | PHI3 | mother | P1-C6 | | G | C | H | C | H | H | H | H | H | A | H | H | T | H | H | A | H | H | T | C | H | H | H | H | C | C | C | A | G | H | C | C | T | T | H | A | G | H | A | H | H | A | C | H | H | H | H | C | A | G | G | H | G | H | C | A | H | A | C | T | H | C | H | T | H | H | H | H | H | H |
| 2016H_P1-A4 | PHI2 | PHI3 | mother | P1-A4 | | G | C | H | C | H | H | H | H | H | A | H | H | T | H | H | A | H | H | T | C | H | H | H | H | C | C | C | H | H | C | H | C | T | T | H | A | G | H | A | H | H | A | C | H | H | H | H | C | A | G | G | H | G | H | C | A | H | H | C | H | H | A | H | A | T | C | H | H | H | G |
| 2016H_P1-A9 | PHI2 | PHI3 | mother | P1-A9 | | G | C | H | C | H | H | H | H | H | A | H | H | T | H | H | A | H | H | T | C | H | H | H | H | C | C | C | A | G | H | C | C | T | T | H | A | G | H | A | H | H | A | C | H | H | H | H | C | A | H | G | G | H | G | H | C | A | H | A | C | H | A | H | A | T | H | H | H | H | G |
| 2016H_P1-A11 | PHI2 | PHI3 | mother | P1-A11 | | G | C | H | C | H | H | H | H | H | A | H | H | T | H | H | A | H | H | T | C | H | H | H | H | C | C | C | H | H | C | H | C | T | T | H | A | G | H | A | H | H | A | C | H | H | H | H | C | A | H | G | G | H | G | H | C | A | H | A | C | T | A | C | H | T | H | H | H | H | G |
| 2016H_P4-H4_2 | PHI5 | PHI3 | progeny | P4-H4 | | G | H | T | H | H | H | H | H | H | C | H | C | H | H | H | H | H | H | H | H | C | H | T | G | G | H | C | H | H | G | H | H | T | T | H | A | G | T | A | H | H | G | H | C | H | H | T | H | H | H | G | G | G | G | H | C | H | H | H | H | H | H | H | H | H | T | H | H | H | G |
| 2016H_P5-D5_1 | PHI5 | PHI3 | progeny | P5-D5 | | G | C | H | H | H | H | H | H | H | C | H | H | H | H | H | H | H | C | H | H | C | H | T | G | G | H | C | C | H | G | H | H | H | H | H | H | G | H | H | G | H | C | H | A | T | T | C | A | H | H | H | H | G | H | H | C | H | H | H | H | H | G | A | A | A | T | H | H | H | T |
| 2016H_P5-D5_2 | PHI5 | PHI3 | progeny | P5-D5 | | G | C | H | H | H | H | H | T | C | H | H | H | H | H | A | H | H | G | H | H | C | H | T | G | G | H | C | H | H | H | H | C | H | H | H | H | H | H | H | G | H | C | H | H | T | T | H | H | H | H | H | H | G | H | H | C | H | H | H | H | H | H | H | H | H | T | H | H | H | H |
| 2016H_P5-D5_3 | PHI5 | PHI3 | progeny | P5-D5 | | G | C | H | H | H | H | H | T | C | H | H | H | H | H | A | H | H | G | H | T | C | H | T | G | G | C | C | H | H | A | C | C | H | H | H | H | A | H | G | A | C | H | H | T | G | H | H | H | H | G | G | G | G | G | H | C | A | H | H | H | H | A | A | A | T | H | H | H | H | H |
| 2016H_P4-F3_1 | PHI5 | PHI3 | progeny | P4-F3 | | G | H | H | C | H | G | A | T | C | H | H | H | H | T | A | H | H | H | C | H | C | C | T | G | G | H | C | H | A | G | C | C | H | G | H | C | H | H | H | H | H | C | G | H | C | H | H | T | H | H | H | G | H | H | C | H | H | H | H | H | H | H | H | T | A | H | H | H | H | H |
| 2016H_P5-C9_1 | PHI5 | PHI3 | progeny | P5-C9 | | G | H | T | C | G | G | A | T | C | G | C | A | T | A | T | G | C | C | H | C | C | T | G | G | H | C | C | A | G | H | C | C | G | H | C | G | A | A | C | C | G | H | C | H | H | H | H | T | C | A | T | H | H | G | A | T | A | G | C | G | C | T | G | C | C | T | A | H | H | H |
| 2016H_P5-C9_2 | PHI5 | PHI3 | progeny | P5-C9 | | G | C | H | C | H | H | H | T | C | H | H | H | H | H | H | H | H | C | H | H | C | H | T | G | G | H | C | H | G | H | C | C | H | H | H | H | H | H | H | H | H | G | H | C | H | H | H | H | H | H | H | H | G | H | H | C | H | H | H | H | H | G | A | A | A | T | H | H | H | T |
| 2016H_P5-B6_1 | PHI5 | PHI3 | progeny | P5-B6 | | G | C | T | H | H | H | H | T | C | G | C | A | T | H | H | T | H | C | H | H | C | H | T | G | G | H | C | H | A | H | C | C | H | H | H | H | H | H | H | H | H | G | H | C | H | T | H | H | H | H | H | H | G | H | H | C | H | H | H | H | H | G | A | A | A | T | H | H | H | H |
| 2016H_P5-B6_2 | PHI5 | PHI3 | progeny | P5-B6 | | G | H | T | C | G | H | H | T | C | G | C | A | H | H | T | G | H | H | H | C | H | T | G | G | H | C | H | H | H | C | C | T | T | H | A | G | T | A | H | G | A | C | C | H | T | H | H | H | H | H | H | H | H | H | H | C | G | H | H | H | H | C | C | T | H | G | A | C | G |
| 2016H_P5-B5_1 | PHI5 | PHI3 | progeny | P5-B5 | | G | H | H | C | H | H | H | H | C | H | H | H | H | A | H | H | H | H | T | C | H | T | G | G | H | C | H | H | H | H | C | C | T | T | H | A | G | T | A | H | G | H | C | H | H | T | H | H | H | H | H | H | H | H | H | G | H | T | A | H | C | H | H | H | H | H | H | T | H | H |
| 2016H_P5-B5_2 | PHI5 | PHI3 | progeny | P5-B5 | | G | H | H | C | H | G | A | T | C | H | H | H | T | A | H | H | H | C | H | C | C | T | G | G | H | C | C | A | G | H | C | C | G | G | C | G | A | A | C | C | G | A | C | C | G | T | H | H | C | T | C | A | G | A | T | A | G | C | G | C | T | G | C | C | T | A | G | A | C | G |
| 2016H_P4-H1_1 | PHI5 | PHI3 | progeny | P4-H1 | | G | C | T | H | G | H | H | H | H | H | C | A | H | H | T | G | H | H | H | C | H | T | G | G | H | C | H | H | H | C | C | C | H | H | H | H | H | H | H | G | H | C | H | H | H | H | H | H | H | H | H | H | H | H | H | H | H | H | H | H | H | H | H | H | H | T | H | H | G |
| 2016H_P3-E5_1 | PHI6 | PHI3 | progeny | P3-E5 | | G | C | H | C | H | H | H | T | C | A | T | G | T | A | C | A | G | C | T | C | C | T | G | G | H | C | H | H | H | C | C | C | G | H | H | G | T | A | H | H | A | C | C | G | T | H | T | C | H | H | G | H | G | G | C | C | H | G | A | C | T | H | C | C | T | C | C | G | H | H |
| 2016H_P3-E11_1 | PHI6 | PHI3 | progeny | P3-E11 | | H | H | H | H | G | G | A | T | C | H | H | H | G | C | C | A | C | G | T | C | H | T | G | G | C | C | C | A | G | H | C | C | C | G | G | C | A | G | T | A | H | H | A | C | C | G | T | H | H | T | G | G | G | G | C | C | A | G | A | C | T | G | C | H | T | A | C | G | H | H |
| 2016H_P3-H10_1 | PHI6 | PHI3 | progeny | P3-H10 | | H | H | H | H | G | G | A | T | C | A | T | G | T | A | C | A | G | C | T | C | H | T | G | G | C | C | C | A | G | H | C | C | G | G | G | C | A | G | T | A | C | G | G | C | C | G | T | H | H | H | H | H | G | G | C | C | A | G | G | A | C | H | A | A | A | T | C | G | G | G |
| 2016H_P3-E3_2 | PHI6 | PHI3 | progeny | P3-E3 | | C | T | C | T | G | G | A | T | C | A | T | G | T | A | C | A | G | C | T | C | C | T | G | G | H | C | C | A | G | C | C | C | G | G | G | C | A | G | T | A | H | H | A | C | C | G | T | H | T | A | H | G | G | G | C | C | A | G | G | A | C | T | A | H | A | T | H | C | G | H |
| 2016H_P4-A9_1 | PHI6 | PHI3 | progeny | P4-A9 | | G | H | H | H | G | G | A | T | C | H | H | G | T | A | C | A | G | C | T | C | C | T | G | G | H | C | C | H | G | T | C | C | G | G | G | C | A | G | T | A | H | H | A | C | C | G | T | T | T | H | H | G | H | G | G | C | A | G | G | A | C | T | A | C | H | T | C | G | H | H |
| 2016H_P3-H7_1 | PHI6 | PHI3 | progeny | P3-H7 | | H | H | H | H | G | G | A | T | C | H | H | H | H | C | C | A | C | G | T | C | H | T | G | G | C | C | C | A | G | T | C | C | G | H | C | A | G | T | A | H | H | H | C | C | G | T | T | T | A | G | G | A | G | G | C | C | A | G | G | A | C | T | A | C | H | T | H | C | G | H |
| 2016H_P3-E4_3 | PHI6 | PHI3 | progeny | P3-E4 | | G | H | H | H | G | G | A | T | C | H | H | G | T | A | C | A | G | H | T | C | H | T | G | G | H | C | H | A | G | H | C | C | T | T | C | A | G | T | A | H | H | H | C | C | G | T | G | H | H | H | G | H | G | G | C | A | G | G | A | C | T | A | C | C | T | C | C | G | H | G |
| 2016H_P3-E4_6 | PHI6 | PHI3 | progeny | P3-E4 | | G | C | H | H | G | G | A | T | C | H | H | H | H | H | C | A | H | H | T | C | H | T | G | G | H | C | H | A | G | H | C | C | T | T | C | A | G | T | A | H | H | H | C | C | G | T | H | H | H | T | G | G | G | G | C | A | G | G | A | C | G | A | A | A | T | C | C | G | H | G |
| 2016H_P3-H9_1 | PHI6 | PHI3 | progeny | P3-H9 | | G | C | T | C | G | G | A | T | C | G | C | A | G | C | C | A | C | G | T | C | T | T | G | G | C | C | T | A | G | H | C | C | H | T | C | A | G | T | A | C | G | A | C | C | G | T | T | T | C | C | G | G | G | G | C | A | G | G | A | C | T | A | C | C | T | C | C | G | G | T |
| 2016H_P3-H9_2 | PHI6 | PHI3 | progeny | P3-H9 | | G | C | H | H | G | G | A | T | C | H | H | H | H | H | C | A | H | H | T | C | H | T | G | G | H | C | H | A | G | H | C | C | T | T | C | A | G | T | A | H | H | H | C | C | G | T | H | H | H | T | G | G | G | G | C | A | G | G | A | C | H | A | A | A | T | C | C | G | H | G |
| 2016H_P3-H9_3 | PHI6 | PHI3 | progeny | P3-H9 | | G | C | T | C | G | G | A | T | C | G | C | A | T | A | C | A | G | C | T | C | C | T | G | G | C | C | G | A | C | H | C | C | H | T | C | A | G | T | A | C | G | A | C | C | G | T | T | T | H | G | G | A | G | G | C | H | G | G | A | C | T | A | C | C | T | C | C | G | H | G |
| 2016H_P3-H9_4 | PHI6 | PHI3 | progeny | P3-H9 | | C | C | T | C | G | G | A | T | C | G | T | G | T | A | C | A | G | C | T | C | C | T | G | G | C | C | C | A | G | C | C | C | H | T | C | A | G | T | A | C | G | A | C | C | G | T | G | T | C | T | G | G | G | G | C | A | G | G | A | C | T | G | C | H | T | C | C | G | H | G |
| 2016H_P3-H9_5 | PHI6 | PHI3 | progeny | P3-H9 | | G | C | T | C | G | G | A | T | C | A | T | G | G | C | C | A | C | G | T | C | T | T | G | G | C | C | T | A | G | T | C | C | G | T | C | A | G | T | A | C | G | A | C | C | G | T | T | T | C | T | G | G | G | G | C | A | G | G | A | C | T | G | C | H | T | A | C | G | H | G |
| 2016H_P4-A6_2 | PHI6 | PHI3 | progeny | P4-A6 | | H | H | H | H | G | G | A | T | C | H | H | H | H | H | C | A | H | H | T | C | H | T | G | G | C | C | C | A | G | H | C | C | G | H | C | A | G | T | A | T | H | A | C | C | G | T | H | T | C | H | G | H | G | G | C | H | G | G | A | C | H | A | H | H | T | C | C | G | H | G |
| 2016H_P4-A6_3 | PHI6 | PHI3 | progeny | P4-A6 | | G | H | H | C | G | G | A | T | C | G | C | A | G | C | C | A | C | G | T | C | T | T | G | G | C | C | C | A | G | H | C | C | G | H | C | A | G | T | A | H | H | A | C | C | G | T | H | T | C | H | G | H | G | G | C | A | G | G | A | C | T | A | H | H | T | C | C | G | H | G |
| 2016H_P3-G11_1 | PHI6 | PHI3 | progeny | P3-G11 | | C | T | C | T | G | G | A | T | C | G | C | A | G | C | C | A | C | G | T | C | T | T | G | G | C | C | T | A | G | T | C | C | H | T | C | A | G | T | A | T | A | A | C | C | G | T | G | T | C | T | G | G | G | G | C | A | G | G | A | C | G | A | A | A | T | C | C | G | G | C |
| 2016H_P3-G11_2 | PHI6 | PHI3 | progeny | P3-G11 | | G | T | C | T | G | G | A | T | C | A | T | G | T | A | C | A | G | C | T | C | C | T | G | G | C | C | T | A | G | T | C | C | G | G | C | A | G | T | A | T | A | A | C | C | G | T | T | T | A | G | G | A | G | G | C | A | G | G | A | C | T | A | C | C | T | A | C | G | G | G |
| 2016H_P3-G11_3 | PHI6 | PHI3 | progeny | P3-G11 | | G | C | T | C | G | G | A | T | C | G | C | A | G | C | C | A | C | G | T | C | T | T | G | G | C | C | T | A | G | T | C | C | H | T | C | A | G | T | A | T | A | A | C | C | G | T | T | T | A | G | G | A | G | G | C | A | G | G | A | C | T | A | C | C | T | C | C | G | G | G |
| 2016H_P3-G11_4 | PHI6 | PHI3 | progeny | P3-G11 | | C | T | C | T | G | G | A | T | C | A | T | G | G | C | C | A | C | G | T | C | T | T | G | G | C | C | T | A | G | T | C | C | G | G | C | A | G | T | A | T | A | A | C | C | G | T | G | T | C | T | G | G | G | G | C | A | G | G | A | C | T | A | A | A | T | A | C | G | G | G |
| 2016H_P3-G11_5 | PHI6 | PHI3 | progeny | P3-G11 | | G | C | T | C | G | G | A | T | C | G | C | A | G | C | C | A | C | G | T | C | T | T | G | G | C | C | T | A | G | T | C | C | H | T | C | A | G | T | A | C | G | A | C | C | G | T | G | T | A | T | G | G | G | G | C | A | G | G | A | C | G | A | A | A | T | A | C | G | G | G |
| 2016H_P3-G11_6 | PHI6 | PHI3 | progeny | P3-G11 | | G | C | T | C | G | G | A | T | C | G | C | G | T | A | C | A | G | C | T | C | C | T | G | G | C | C | T | A | G | T | C | C | H | T | C | A | G | T | A | C | G | A | C | C | G | T | T | T | A | T | G | G | G | G | C | A | G | G | A | C | G | A | A | A | T | A | C | G | G | G |
| 2016H_P3-G11_7 | PHI6 | PHI3 | progeny | P3-G11 | | G | C | T | C | G | G | A | T | C | G | C | A | G | C | C | A | C | C | T | C | C | T | G | G | C | C | T | A | G | T | C | C | H | T | C | A | G | T | A | C | G | A | C | C | G | T | T | T | A | G | G | A | G | G | C | A | G | G | A | C | T | A | C | C | T | C | C | G | C | G |
| 2016H_P3-G11_8 | PHI6 | PHI3 | progeny | P3-G11 | | G | C | T | C | G | G | A | T | C | G | C | A | G | C | C | A | C | G | T | C | T | T | G | G | C | C | T | A | G | T | C | C | G | G | C | A | G | T | A | C | G | A | C | C | G | T | G | T | C | T | G | G | G | G | C | A | G | G | A | C | G | A | A | A | T | C | C | G | C | G |
| 2016H_P3-G11_9 | PHI6 | PHI3 | progeny | P3-G11 | | C | C | T | C | G | G | A | T | C | G | C | A | G | C | C | A | C | G | T | C | T | T | G | G | C | C | T | A | G | T | C | C | G | G | C | A | G | T | A | C | G | A | C | C | G | T | T | T | A | G | G | A | G | G | C | A | G | G | A | C | T | A | C | C | T | C | C | G | G | G |
| 2016H_P3-G12_2 | PHI6 | PHI3 | progeny | P3-G12 | | C | C | T | C | G | G | A | T | C | A | T | G | T | A | C | A | G | C | T | C | C | T | G | G | C | C | G | A | C | H | C | C | G | G | C | A | G | T | A | C | G | G | C | C | G | T | G | C | C | G | G | A | G | G | C | A | G | G | A | C | G | A | A | A | T | C | C | G | G | G |
| 2016H_P3-G12_3 | PHI6 | PHI3 | progeny | P3-G12 | | G | C | T | C | G | G | A | T | C | G | C | A | G | C | C | A | C | G | T | C | T | T | G | G | C | C | G | A | C | C | C | C | T | T | C | A | G | T | A | T | A | A | C | C | G | T | T | G | C | G | G | A | G | G | C | A | G | G | A | C | T | A | C | H | T | C | C | G | C | G |
| 2016H_P3-G12_4 | PHI6 | PHI3 | progeny | P3-G12 | | G | C | T | C | G | G | A | T | C | A | T | G | T | A | C | A | G | C | T | C | C | T | G | G | C | C | G | A | C | C | C | C | T | T | C | A | G | T | A | T | A | A | C | C | G | T | T | T | C | G | G | A | G | G | C | A | G | G | A | C | T | A | C | H | T | C | C | G | G | G |
| 2016H_P3-G12_5 | PHI6 | PHI3 | progeny | P3-G12 | | G | C | T | C | G | G | A | T | C | A | T | G | T | A | C | A | G | C | T | C | C | T | G | G | C | C | T | A | G | T | C | C | H | T | C | A | G | T | A | C | G | G | C | C | G | T | G | C | C | G | G | A | G | G | C | A | G | G | A | C | T | A | C | H | T | C | C | G | G | G |
| 2016H_P3-G12_6 | PHI6 | PHI3 | progeny | P3-G12 | | C | T | C | T | G | A | C | T | C | G | C | A | G | C | C | A | C | G | T | C | T | T | G | G | C | C | G | A | C | C | C | C | H | T | C | A | G | T | A | C | G | A | C | C | G | T | T | T | C | T | G | G | G | G | C | A | G | G | A | C | T | A | C | H | T | C | C | G | G | G |
| 2016H_P3-G12_8 | PHI6 | PHI3 | progeny | P3-G12 | | G | T | C | T | G | A | C | T | C | G | C | A | G | C | C | A | C | G | T | C | T | T | G | G | C | C | G | A | C | C | C | C | T | T | C | A | G | T | A | C | G | G | C | C | G | T | T | G | C | T | G | G | G | G | C | A | G | G | A | C | H | A | A | A | T | C | C | G | C | G |
| 2016H_P3-G12_9 | PHI6 | PHI3 | progeny | P3-G12 | | G | C | T | C | G | G | A | T | C | A | T | G | T | A | C | A | G | C | T | C | C | T | G | G | C | C | T | A | G | T | C | C | T | T | C | A | G | T | A | C | G | G | C | C | G | T | T | T | C | G | G | A | G | G | C | A | G | G | A | C | T | A | C | C | T | C | C | G | G | G |
| 2016H_P3-G12_10 | PHI6 | PHI3 | progeny | P3-G12 | | C | C | T | C | G | G | A | T | C | A | T | G | T | A | C | A | G | C | T | C | C | T | G | G | C | C | T | A | G | T | C | C | T | T | C | A | G | T | A | C | G | G | C | C | G | T | G | C | C | G | G | A | G | G | C | A | G | G | A | C | T | A | C | H | T | C | C | G | C | G |
| 2016H_P3-G12_11 | PHI6 | PHI3 | progeny | P3-G12 | | C | T | C | T | G | A | C | T | C | A | T | G | T | A | C | A | G | C | T | C | C | T | G | G | C | C | T | G | A | C | C | C | H | T | C | A | G | T | A | C | G | G | C | C | G | T | G | C | C | T | G | G | G | G | C | A | G | G | A | C | T | A | C | H | T | C | C | G | C | G |
| 2016H_P3-G12_12 | PHI6 | PHI3 | progeny | P3-G12 | | G | C | T | C | G | A | C | T | C | A | T | G | T | A | C | A | G | C | T | C | C | T | G | G | C | C | T | A | G | T | C | C | H | T | C | A | G | T | A | C | G | G | C | C | G | T | G | C | C | G | G | A | G | G | C | A | G | G | A | C | T | A | C | H | T | C | C | G | G | G |
| 2016H_P3-G12_13 | PHI6 | PHI3 | progeny | P3-G12 | | C | C | T | C | G | G | A | T | C | G | C | A | G | C | C | A | C | G | T | C | T | T | G | G | C | C | G | A | C | C | C | C | H | T | C | A | G | T | A | C | G | A | C | C | G | T | G | C | C | G | G | A | G | G | C | A | G | G | A | C | G | A | C | C | T | C | C | G | C | G |
| 2016H_P3-F1_1 | PHI6 | PHI3 | progeny | P3-F1 | | C | T | C | C | G | G | A | T | C | G | C | G | T | A | C | A | G | C | T | C | C | T | G | G | C | C | T | A | G | T | C | C | T | T | C | A | G | T | A | C | G | G | C | C | G | T | G | C | C | G | G | G | G | C | C | A | G | G | A | C | T | G | C | H | T | C | C | G | G | G |
| 2016H_P3-F1_2 | PHI6 | PHI3 | progeny | P3-F1 | | C | T | C | C | G | A | C | T | C | A | T | G | T | C | C | A | C | G | T | C | T | T | G | G | C | C | C | A | A | C | C | C | T | T | C | A | G | T | A | C | G | G | C | C | G | T | G | C | C | G | G | A | G | G | C | A | G | G | A | C | T | G | C | H | T | C | C | G | G | G |
| 2016H_P3-F1_3 | PHI6 | PHI3 | progeny | P3-F1 | | G | C | T | C | G | G | A | T | C | A | C | A | G | C | C | A | C | G | T | C | T | T | G | G | C | C | T | A | G | T | C | C | T | T | C | A | G | T | A | C | G | G | C | C | G | T | G | C | C | G | G | A | G | G | C | A | G | G | A | C | T | A | C | C | T | C | C | G | G | G |
| 2016H_P3-F1_4 | PHI6 | PHI3 | progeny | P3-F1 | | G | C | T | C | G | A | C | T | C | G | C | A | T | A | C | A | G | C | T | C | T | T | G | G | C | C | T | A | G | T | C | C | T | T | C | A | G | T | A | T | A | A | C | C | G | T | T | T | C | G | G | A | G | G | C | A | G | G | A | C | T | A | C | C | T | C | C | G | G | G |
| 2016H_P3-F1_5 | PHI6 | PHI3 | progeny | P3-F1 | | C | T | C | T | G | A | C | T | C | G | C | G | T | A | C | A | G | C | T | C | C | T | G | G | C | C | C | A | A | T | C | C | H | T | C | A | G | T | A | C | G | G | C | C | G | T | G | C | C | G | G | A | G | G | C | H | G | G | A | C | T | A | C | C | T | C | C | G | G | G |
| 2016H_P3-F1_6 | PHI6 | PHI3 | progeny | P3-F1 | | C | T | C | T | G | G | A | T | C | G | C | A | G | C | C | A | C | C | T | C | C | T | G | G | C | C | C | A | A | C | C | C | T | T | C | A | G | T | A | T | A | G | C | C | G | T | T | T | C | T | G | G | G | G | C | C | A | G | A | C | T | G | C | H | T | C | C | G | C | G |
| 2016H_P3-F1_7 | PHI6 | PHI3 | progeny | P3-F1 | | C | T | C | T | G | A | C | T | C | A | T | G | T | A | C | A | G | C | T | C | C | T | G | G | C | C | T | A | G | T | C | C | T | T | H | G | T | A | T | A | A | C | C | G | T | T | T | C | T | G | G | G | G | G | C | A | G | G | A | C | T | A | C | C | T | C | C | G | G | G |
| 2016H_P3-F1_8 | PHI6 | PHI3 | progeny | P3-F1 | | G | C | T | C | G | G | A | T | C | G | C | A | G | C | C | A | C | C | T | C | C | T | G | G | C | C | C | A | A | T | C | C | H | T | C | A | G | T | A | C | G | G | C | C | G | T | T | T | C | G | G | G | G | C | C | A | G | G | A | C | T | A | C | H | T | C | C | G | G | G |
| 2016H_P3-F1_9 | PHI6 | PHI3 | progeny | P3-F1 | | C | C | T | C | G | G | A | T | C | G | C | A | G | C | C | A | C | G | T | C | T | T | G | G | C | C | C | A | A | C | C | C | H | T | C | A | G | T | A | T | A | A | C | C | G | T | G | C | C | G | G | A | G | G | C | A | G | G | A | C | T | A | C | H | T | C | C | G | G | G |
| 2016H_P3-F1_10 | PHI6 | PHI3 | progeny | P3-F1 | | C | T | T | C | G | A | C | T | C | G | C | A | G | C | C | A | C | G | T | C | T | T | G | G | C | C | T | A | G | T | C | C | H | T | C | A | G | T | A | C | G | G | C | C | G | T | G | C | C | G | G | G | G | C | C | A | G | G | A | C | T | G | C | H | T | C | C | G | G | G |
| 2016H_P3-F1_11 | PHI6 | PHI3 | progeny | P3-F1 | | C | T | C | T | G | G | A | T | C | A | T | G | T | A | C | A | G | C | T | C | C | T | G | G | C | C | C | A | A | C | C | C | T | T | C | A | G | T | A | C | G | G | C | C | G | T | G | C | C | G | G | A | G | G | C | A | G | G | A | C | T | A | C | C | T | C | C | G | C | G |
| 2016H_P3-F1_12 | PHI6 | PHI3 | progeny | P3-F1 | | G | C | C | T | G | A | C | T | C | G | C | A | T | A | C | A | G | C | T | C | C | T | G | G | C | C | C | A | G | T | C | C | H | T | C | A | G | T | A | T | A | A | C | C | G | T | G | C | C | G | G | A | G | G | C | A | G | G | A | C | T | A | C | C | T | C | C | G | G | G |
| 2016H_P3-F1_14 | PHI6 | PHI3 | progeny | P3-F1 | | G | C | T | C | G | G | A | T | C | G | C | A | G | C | C | A | C | G | T | C | T | T | G | G | C | C | C | A | G | T | C | C | T | T | C | A | G | T | A | T | A | A | C | C | G | T | T | T | C | G | G | G | G | C | C | A | G | G | A | C | T | A | C | C | T | C | C | G | C | G |
| 2016H_P3-F1_15 | PHI6 | PHI3 | progeny | P3-F1 | | G | C | C | T | G | A | C | T | C | A | T | G | T | A | C | A | G | C | T | C | T | T | G | G | C | C | T | A | G | T | C | C | H | T | C | A | G | T | A | T | A | A | C | C | G | T | G | T | C | G | G | A | G | G | C | A | G | G | A | C | T | A | C | C | T | C | C | G | C | G |
| 2016H_P3-F1_16 | PHI6 | PHI3 | progeny | P3-F1 | | C | T | C | T | G | A | C | T | C | A | T | G | G | C | C | A | C | C | T | C | C | T | G | G | C | C | C | A | A | C | C | C | T | T | C | A | G | T | A | C | G | G | C | C | G | T | G | C | C | G | G | A | G | G | C | A | G | G | A | C | T | A | C | C | T | C | C | G | C | G |
| 2016H_P3-F1_17 | PHI6 | PHI3 | progeny | P3-F1 | | C | T | T | C | G | G | A | T | C | A | T | G | T | A | C | A | G | C | T | C | T | T | G | G | C | C | T | A | G | T | C | C | H | T | C | A | G | T | A | C | G | A | C | C | G | T | G | C | C | G | G | G | G | G | C | H | G | G | A | C | T | G | C | H | T | C | C | G | G | G |
| 2016H_P3-F11_3 | PHI6 | PHI3 | progeny | P3-F11 | | G | C | C | C | C | A | C | T | C | A | C | A | G | C | C | A | C | G | T | C | T | T | G | G | C | C | T | A | G | C | T | C | G | G | C | A | G | T | A | T | A | A | C | C | G | T | T | T | C | G | G | A | G | G | C | A | G | G | A | C | G | A | A | A | T | A | C | G | C | G |
| 2016H_P3-F11_4 | PHI6 | PHI3 | progeny | P3-F11 | | G | C | C | C | C | A | C | T | C | G | C | A | G | C | C | A | C | C | T | C | C | T | G | G | C | C | C | A | G | C | T | C | G | G | H | A | G | T | A | T | A | A | C | C | G | T | G | T | C | G | G | A | G | G | C | A | G | G | A | C | T | A | C | C | T | C | C | G | G | T |
| 2016H_P3-F11_5 | PHI6 | PHI3 | progeny | P3-F11 | | G | C | C | C | C | A | C | T | C | A | T | A | G | C | C | A | C | G | T | C | T | T | G | G | C | C | C | A | G | C | C | C | G | G | C | A | G | T | A | T | A | A | C | C | G | T | G | T | C | G | G | G | G | G | C | C | A | G | A | C | H | A | A | A | T | A | C | G | G | T |
| 2016H_P3-F11_7 | PHI6 | PHI3 | progeny | P3-F11 | | G | C | T | C | G | G | A | T | C | G | C | A | T | A | C | A | G | C | T | C | T | T | G | G | C | C | C | A | G | C | C | C | G | G | C | A | G | T | A | C | G | A | C | C | G | T | T | T | C | T | G | G | G | G | C | C | A | G | A | C | H | A | A | A | T | C | C | G | C | G |
| 2016H_P3-F11_8 | PHI6 | PHI3 | progeny | P3-F11 | | G | C | C | C | C | A | C | T | C | A | T | A | G | C | C | A | C | G | T | C | C | T | G | G | C | C | T | A | G | C | T | C | G | G | C | A | G | T | A | T | A | A | C | C | G | T | G | T | C | G | G | A | G | G | C | A | G | G | A | C | T | A | C | C | T | C | C | G | C | G |
| 2016H_P3-F11_11 | PHI6 | PHI3 | progeny | P3-F11 | | G | C | C | C | G | G | A | T | C | G | C | G | T | A | C | A | G | C | T | C | C | T | G | G | C | C | C | A | G | C | T | C | H | T | C | A | G | T | A | C | G | A | C | C | G | T | G | T | C | G | G | A | G | G | C | A | G | G | A | C | T | A | C | C | T | C | C | G | C | G |
| 2016H_P3-F11_14 | PHI6 | PHI3 | progeny | P3-F11 | | G | C | C | C | C | A | C | T | C | A | T | G | T | A | C | A | G | C | T | C | C | T | G | G | C | C | C | A | G | C | C | C | G | G | C | A | G | T | A | T | A | A | C | C | G | T | T | T | C | G | G | G | G | G | C | C | G | G | A | C | T | A | C | C | T | C | C | G | C | G |

**Table S8. Results of the SNP array analasis (continue)**

| | | | | | Chr | 6 | 6 | 6 | 6 | 6 | 6 | 6 | 7 | 7 | 7 | 7 | 7 | 7 | 7 | 7 | 7 | 7 | 7 | 7 | 7 | 7 | 7 | 7 | 7 | 7 | 7 | 7 | 7 | 7 | 7 | 7 | 7 | 8 | 8 | 8 | 8 | 8 | 8 | 8 | 9 | 9 | 9 | 9 | 9 | 9 | 9 | 9 | 9 | 9 | 10 | 10 | 10 | 10 | 10 | 10 | 10 | 10 | 10 | 10 | 10 | 10 | 10 | 10 | 10 | 10 | 10 | 10 | 10 | 10 | 10 |
|---|---|---|---|---|---|---|---|---|---|---|---|---|---|---|---|---|---|---|---|---|---|---|---|---|---|---|---|---|---|---|---|---|---|---|---|---|---|---|---|---|---|---|---|---|---|---|---|---|---|---|---|---|---|---|---|---|---|---|---|---|---|---|---|---|---|---|---|---|---|---|---|---|---|---|---|
| | | | | | Position, cM | 89 | 130 | 142 | 151 | 155 | 180 | 180 | 8 | 17 | 32 | 37 | 52 | 64 | 75 | 74 | 85 | 92 | 109 | 111 | 130 | 135 | 141 | 150 | 162 | 177 | 182 | 186 | 192 | 196 | 201 | 205 | 205 | 40 | 50 | 113 | 164 | 177 | 187 | 195 | 35 | 39 | 45 | 60 | 127 | 143 | 159 | 166 | 178 | 180 | 11 | 13 | 18 | 27 | 31 | 40 | 48 | 54 | 70 | 76 | 87 | 91 | 103 | 108 | 112 | 121 | 130 | 143 | 154 | 162 | 161 |
| plant_id | mom_gen | dad_gen | mother_progeny | family_ | | | | | | | | | | | | | | | | | | | | | | | | | | | | | | | | | | | | | | | | | | | | | | | | | | | | | | | | | | | | | | | | | | | | | | | |
| 2016H_P3-F3_3 | PHI6 | PHI3 | progeny | P3-F3 | | G | C | T | C | G | G | A | T | C | A | T | G | T | A | C | A | G | C | T | C | C | T | G | G | C | C | C | G | A | C | C | C | H | T | C | A | G | T | A | C | G | A | C | C | G | T | T | C | A | T | G | G | G | G | C | A | G | G | A | C | T | A | C | A | T | C | C | G | G | T |
| 2016H_P3-H8_5 | PHI6 | PHI3 | progeny | P3-H8 | | C | T | C | T | G | G | A | T | C | G | C | A | G | C | C | A | C | G | T | C | T | T | G | G | C | C | C | A | G | C | C | C | G | G | C | A | G | T | A | T | A | A | C | C | G | T | T | T | A | T | G | G | G | G | C | C | A | G | A | C | T | G | C | H | T | A | C | G | C | G |
| 2016H_P3-G6_5 | PHI6 | PHI3 | progeny | P3-G6 | | C | T | C | C | C | G | A | T | C | G | C | A | G | C | C | A | C | C | T | C | C | T | G | G | C | C | C | G | G | T | C | C | G | G | C | A | G | T | A | C | G | A | C | C | G | T | T | T | C | G | G | A | G | G | C | A | G | G | A | C | T | G | A | A | T | C | C | G | G | G |
| 2016H_P3-F6_4 | PHI6 | PHI3 | progeny | P3-F6 | | C | C | T | C | G | G | A | T | C | A | T | G | T | A | C | A | G | C | T | C | C | T | G | G | C | C | C | A | G | C | C | C | G | G | C | A | G | T | A | T | A | A | C | C | G | T | G | T | C | T | G | G | G | G | C | C | A | G | A | C | G | A | A | A | T | C | C | G | C | G |
| 2016H_P3-F6_5 | PHI6 | PHI3 | progeny | P3-F6 | | C | T | C | T | G | G | A | T | C | G | C | A | G | C | C | A | C | G | T | C | T | T | G | G | C | C | C | A | G | C | C | C | G | G | C | A | G | T | A | T | A | A | C | C | G | T | G | T | C | T | G | G | G | G | C | C | A | G | A | C | T | A | C | C | T | C | C | G | C | G |
| 2016H_P3-F6_6 | PHI6 | PHI3 | progeny | P3-F6 | | C | T | C | T | G | G | A | T | C | A | T | G | T | A | C | A | G | C | T | C | C | T | G | G | C | C | T | A | G | T | C | C | G | G | C | A | G | T | A | C | G | A | C | C | G | T | T | T | C | T | G | G | G | G | C | A | G | G | A | C | T | A | C | C | T | C | C | G | C | G |
| 2016H_P3-G2_1 | PHI6 | PHI3 | progeny | P3-G2 | | G | C | T | C | G | G | A | T | C | A | T | G | T | C | C | A | C | G | T | C | T | T | G | G | C | C | C | A | G | C | C | C | H | T | C | A | G | T | A | C | G | G | C | C | G | T | T | T | A | T | G | G | G | G | C | C | A | G | A | C | H | A | A | A | T | A | C | G | C | G |
| 2016H_P3-G2_2 | PHI6 | PHI3 | progeny | P3-G2 | | G | C | T | C | G | G | A | T | C | G | C | A | G | C | C | A | C | G | T | C | T | T | G | G | C | C | T | A | G | T | C | C | G | G | C | A | G | T | A | C | G | G | C | C | G | T | T | T | A | T | G | G | G | G | C | C | A | G | A | C | G | A | A | A | T | A | C | G | G | T |
| 2016H_P3-F8_1 | PHI6 | PHI3 | progeny | P3-F8 | | G | C | T | T | G | G | A | T | C | G | C | A | G | C | C | A | C | G | T | C | T | T | G | G | C | C | C | A | G | C | C | C | H | T | C | A | G | T | A | T | A | A | C | C | G | T | G | C | A | T | G | G | G | G | C | C | A | C | A | C | G | A | A | A | T | C | C | G | C | G |
| 2016H_P3-F8_2 | PHI6 | PHI3 | progeny | P3-F8 | | G | C | T | T | G | G | A | T | C | G | C | A | G | C | C | A | C | G | T | C | T | T | G | G | C | C | T | A | G | T | C | C | H | T | C | A | G | T | A | T | A | A | C | C | G | T | G | C | A | T | G | G | G | G | C | C | A | C | A | C | G | A | A | A | T | C | C | G | C | G |
| 2016H_P3-G7_6 | PHI6 | PHI3 | progeny | P3-G7 | | C | T | C | T | G | G | A | T | C | A | C | A | G | C | C | A | C | C | T | C | C | T | G | G | C | C | C | A | G | C | T | C | H | T | C | A | G | T | A | C | G | A | C | C | G | T | G | C | A | G | G | A | G | G | C | A | G | G | A | C | T | A | C | C | T | C | C | G | C | G |
| 2016H_P3-A12_1 | PHI7 | PHI3 | progeny | P3-A12 | | G | T | T | C | C | A | C | T | C | G | C | G | T | A | C | A | G | C | T | C | C | T | G | G | C | C | C | A | G | C | C | C | G | T | C | H | G | T | A | C | G | G | C | C | A | C | G | C | A | T | G | G | G | A | C | A | G | G | A | C | T | A | C | C | T | A | C | G | G | G |
| 2016H_P3-A12_2 | PHI7 | PHI3 | progeny | P3-A12 | | G | C | T | C | C | A | C | T | C | G | C | A | G | C | C | A | C | C | C | T | C | T | G | G | C | A | T | A | G | T | C | C | H | T | C | A | G | T | A | C | G | G | C | C | G | T | G | C | A | T | G | G | G | A | C | A | G | G | A | C | T | A | A | A | T | A | C | G | G | G |
| 2016H_P3-C4_1 | PHI7 | PHI3 | progeny | P3-C4 | | G | H | T | C | H | A | C | T | C | G | C | A | G | C | C | A | C | C | C | T | C | T | G | G | H | C | H | A | G | H | C | C | H | T | C | A | G | T | A | C | G | H | C | C | G | T | G | T | C | T | G | G | G | H | C | A | G | G | A | C | T | A | H | H | T | A | C | G | G | G |
| 2016H_P3-A9_1 | PHI7 | PHI3 | progeny | P3-A9 | | G | H | T | C | H | H | H | T | C | H | H | H | H | H | C | A | H | C | H | H | C | T | G | G | C | C | C | A | G | H | C | C | G | T | C | A | G | T | A | C | G | G | T | T | A | C | G | C | A | T | G | G | G | G | C | C | A | C | A | C | G | A | A | A | T | C | C | G | G | G |
| 2016H_P3-C2_1 | PHI7 | PHI3 | progeny | P3-C2 | | G | H | T | C | H | H | H | T | C | H | H | H | H | H | C | A | H | C | H | H | C | T | G | G | H | C | C | A | G | H | C | C | G | T | C | A | G | T | A | C | G | G | H | H | H | H | G | C | A | T | G | G | G | H | C | C | H | H | A | C | H | A | H | H | T | H | C | G | G | G |
| 2016H_P2-H11_2 | PHI7 | PHI3 | progeny | P2-H11 | | G | T | T | C | C | A | C | T | C | A | H | G | T | A | C | A | G | C | T | C | C | T | G | G | C | C | C | A | G | C | C | C | G | T | C | A | G | T | A | C | G | G | T | T | H | H | G | H | C | T | G | G | G | A | C | C | A | C | A | C | G | A | A | A | T | A | C | G | G | G |
| 2016H_P3-B2_1 | PHI7 | PHI3 | progeny | P3-B2 | | G | C | T | C | G | G | A | T | C | H | H | H | H | H | H | A | H | C | H | H | C | T | G | G | H | C | H | A | G | H | C | C | G | T | C | A | G | T | A | C | G | H | H | H | H | H | G | H | H | T | G | G | G | H | C | C | H | G | A | C | T | A | C | C | T | A | C | H | G | G |
| 2016H_P3-B2_2 | PHI7 | PHI3 | progeny | P3-B2 | | G | C | T | C | G | G | A | T | C | G | C | A | G | C | C | A | C | C | C | H | C | T | G | G | H | C | C | A | G | C | C | C | H | T | C | H | G | T | A | C | G | A | C | H | H | H | G | H | H | T | G | G | G | H | C | C | H | G | A | C | T | A | C | C | T | A | C | G | G | G |
| 2016H_P3-A12_3 | PHI7 | PHI3 | progeny | P3-A12 | | G | T | T | C | C | A | C | T | C | A | T | G | T | A | C | H | G | C | T | C | C | T | G | G | C | C | T | A | G | T | C | C | G | T | C | A | G | T | A | C | G | G | T | C | G | T | G | C | A | T | G | G | G | A | C | A | G | G | A | C | T | A | C | C | T | A | C | G | G | G |
| 2016H_P3-A12_12 | PHI7 | PHI3 | progeny | P3-A12 | | G | T | T | C | C | G | A | T | C | H | H | H | H | A | C | A | G | C | H | C | C | T | G | G | C | C | C | A | G | C | C | C | H | T | C | A | G | T | A | C | G | G | C | H | H | H | G | H | H | T | G | G | G | G | C | C | A | C | A | C | H | A | C | C | T | A | C | G | G | G |
| 2016H_P3-C9_2 | PHI7 | PHI3 | progeny | P3-C9 | | G | H | T | C | G | G | A | T | C | A | T | G | T | A | C | A | G | C | T | C | C | T | G | G | C | C | C | A | G | C | C | C | G | T | C | A | G | T | A | C | G | H | H | C | G | T | G | H | C | T | G | G | G | G | C | C | A | C | A | C | H | A | H | H | T | A | C | G | G | G |
| 2016H_P3-A8_2 | PHI7 | PHI3 | progeny | P3-A8 | | G | H | T | C | H | H | H | T | C | H | H | H | H | H | C | A | H | C | H | H | C | T | G | G | H | C | C | A | G | H | C | C | G | T | C | A | G | T | A | C | G | H | H | T | H | H | G | H | H | T | G | G | G | G | C | C | A | C | A | C | G | A | H | H | T | H | C | G | G | G |
| 2016H_P3-A8_3 | PHI7 | PHI3 | progeny | P3-A8 | | G | H | T | C | H | A | C | T | C | A | T | G | T | A | C | A | G | C | H | H | C | T | G | G | H | C | C | A | G | H | C | C | G | T | C | A | G | T | A | C | G | A | C | H | H | H | G | H | H | T | G | G | G | H | C | C | H | C | A | C | G | A | H | H | T | H | C | G | G | G |
| 2016H_P3-A11_4 | PHI7 | PHI3 | progeny | P3-A11 | | G | C | T | C | C | G | A | T | C | A | H | G | T | C | C | A | H | C | T | C | C | T | G | G | C | C | C | A | G | C | C | C | G | T | C | A | G | T | A | C | G | G | C | C | G | T | G | C | A | T | G | G | G | A | C | A | G | G | A | C | T | A | C | C | T | A | C | G | G | G |
| 2016H_P3-A11_5 | PHI7 | PHI3 | progeny | P3-A11 | | G | C | T | C | G | G | A | T | C | A | T | G | T | A | C | A | G | C | T | C | C | T | G | G | C | C | C | A | G | C | C | C | H | T | C | A | G | T | A | C | G | A | C | C | G | T | G | C | A | T | G | G | G | G | C | C | A | C | A | C | G | A | A | A | T | C | C | G | G | G |
| 2016H_P3-A11_6 | PHI7 | PHI3 | progeny | P3-A11 | | G | C | T | C | C | A | C | T | C | G | C | A | G | C | C | A | C | C | C | T | C | T | G | G | C | A | T | A | G | T | C | C | H | T | C | A | G | T | A | C | G | G | T | C | G | T | G | T | C | T | G | G | G | G | C | C | A | C | A | C | T | A | C | C | T | A | C | G | G | G |
| 2016H_P3-B9_2 | PHI7 | PHI3 | progeny | P3-B9 | | G | C | T | C | G | G | A | T | C | A | T | G | T | A | C | A | C | C | C | T | C | T | G | G | C | A | T | A | G | C | C | C | G | T | C | A | G | T | A | C | G | G | T | T | A | T | G | C | A | T | G | G | G | A | C | A | G | G | A | C | T | A | A | A | T | C | C | H | G | G |
| 2016H_P2-C6_3 | PHI4 | PHI3 | progeny | P2-C6 | | G | C | H | C | H | H | H | T | H | H | H | H | T | H | H | A | H | H | T | H | H | H | G | G | H | C | C | A | G | C | H | C | G | T | H | H | G | T | A | C | H | H | C | H | H | T | H | T | C | H | G | G | H | G | C | C | H | H | A | C | H | A | H | A | T | A | H | G | G | H |
| 2016H_P2-C9_1 | PHI4 | PHI3 | progeny | P2-C9 | | G | H | T | C | H | H | H | T | H | H | H | A | G | C | T | A | C | G | T | H | H | H | G | G | H | C | C | H | H | C | C | C | G | T | H | H | G | T | A | C | A | G | C | T | H | T | H | T | H | H | G | G | H | G | C | C | H | C | A | C | G | A | H | A | T | A | H | G | G | H |
| 2016H_P2-C9_4 | PHI4 | PHI3 | progeny | P2-C9 | | G | H | T | C | H | H | H | T | H | H | H | H | H | H | H | A | H | H | T | H | H | H | G | G | H | C | C | H | H | C | C | C | H | T | C | A | G | T | A | C | H | G | C | H | H | T | H | T | H | H | G | G | H | G | C | C | H | C | A | C | G | A | H | A | T | A | H | G | G | H |
| 2016H_P2-B7_1 | PHI4 | PHI3 | progeny | P2-B7 | | G | H | T | C | H | H | H | T | H | H | H | H | H | H | H | A | H | H | T | H | H | H | H | G | H | C | C | H | H | C | C | C | G | T | H | H | G | T | A | C | H | H | C | H | H | T | H | T | C | H | G | G | H | G | C | C | H | H | A | C | G | A | H | A | T | H | H | G | G | H |
| 2016H_P2-B7_2 | PHI4 | PHI3 | progeny | P2-B7 | | G | H | T | C | C | H | H | T | H | H | H | H | T | A | C | A | G | C | T | C | C | T | G | G | C | C | C | A | G | C | C | C | G | T | T | H | G | T | A | C | G | A | C | C | H | T | H | T | C | G | G | G | A | G | C | C | G | C | A | C | G | A | H | A | T | H | H | G | G | T |
| 2016H_P2-B7_3 | PHI4 | PHI3 | progeny | P2-B7 | | G | H | T | C | H | H | H | T | H | H | H | H | H | H | H | A | H | H | T | H | H | H | H | G | H | C | C | H | H | C | C | C | G | T | H | H | G | T | A | C | H | H | C | H | H | T | H | T | C | H | G | G | H | G | C | C | H | H | A | C | G | A | H | A | T | H | H | G | G | H |
| 2016H_P2-B7_4 | PHI4 | PHI3 | progeny | P2-B7 | | G | H | T | C | H | H | H | T | H | H | H | H | H | H | H | A | H | H | T | H | H | H | H | G | H | C | C | H | H | C | C | C | G | T | H | H | G | T | A | C | G | A | C | H | H | T | H | T | C | H | G | G | H | G | C | C | H | H | A | C | G | A | H | A | T | H | H | G | G | H |
| 2016H_P2-F3_2 | PHI4 | PHI3 | progeny | P2-F3 | | G | H | T | C | H | H | H | T | H | H | C | A | G | C | T | A | C | G | T | T | T | G | G | G | A | C | C | A | G | C | H | C | G | T | C | A | G | T | A | C | H | H | C | H | H | T | H | T | C | T | G | G | G | G | C | C | A | C | A | C | H | A | H | A | T | A | C | G | G | G |
| 2016H_P2-F3_3 | PHI4 | PHI3 | progeny | P2-F3 | | G | H | T | C | H | H | H | T | H | H | H | H | T | H | H | A | H | H | T | H | H | H | G | G | H | C | C | A | G | C | H | C | G | T | H | H | G | T | A | C | H | H | C | C | G | T | G | T | C | G | G | G | A | G | C | C | G | C | A | C | G | A | H | A | T | A | H | G | G | H |
| 2016H_P2-G6_1 | PHI4 | PHI3 | progeny | P2-G6 | | G | H | T | C | H | H | H | T | H | H | H | H | H | H | H | A | H | H | T | H | H | H | G | G | H | C | C | H | H | C | C | C | G | T | H | H | G | T | A | C | H | G | C | H | H | T | H | T | C | H | G | G | H | G | C | C | H | C | A | C | G | A | H | A | T | H | H | G | G | H |
| 2016H_P2-G6_2 | PHI4 | PHI3 | progeny | P2-G6 | | G | H | T | C | H | H | H | T | H | H | H | H | H | H | H | A | H | H | T | H | H | H | G | G | H | C | C | H | H | C | C | C | G | T | H | H | G | T | A | C | H | G | C | H | H | T | H | T | C | H | G | G | H | G | C | C | H | C | A | C | H | A | H | A | T | H | H | G | G | H |
| 2016H_P2-G6_3 | PHI4 | PHI3 | progeny | P2-G6 | | G | H | T | C | H | H | H | T | H | H | H | H | H | H | H | A | H | H | T | H | H | H | G | G | H | C | C | H | A | C | C | C | G | T | H | H | G | T | A | C | H | G | C | H | H | T | H | T | C | H | G | G | H | G | C | C | H | C | A | C | G | A | H | A | T | H | H | G | G | H |
| 2016H_P2-G6_4 | PHI4 | PHI3 | progeny | P2-G6 | | G | H | T | C | H | H | H | T | H | H | H | H | H | H | H | A | C | G | T | T | T | G | G | G | A | C | C | G | A | C | C | C | G | T | H | H | G | T | A | C | G | G | C | H | H | T | H | T | C | H | G | G | H | G | C | C | H | C | A | C | H | A | H | A | T | H | H | G | G | H |
| 2016H_P2-E7_1 | PHI4 | PHI3 | progeny | P2-E7 | | G | H | T | C | H | H | H | T | H | H | H | H | H | H | H | A | H | H | T | C | C | T | G | G | C | C | C | H | H | C | C | C | G | T | H | H | G | T | A | C | A | G | C | H | H | T | H | H | H | G | G | G | A | G | C | C | G | C | A | C | G | A | H | A | T | A | C | G | G | G |
| 2016H_P2-E7_2 | PHI4 | PHI3 | progeny | P2-E7 | | G | H | T | C | H | H | H | T | C | A | T | G | T | A | C | A | H | H | T | C | H | H | H | G | H | C | C | A | G | C | C | C | H | T | C | H | G | T | A | C | G | G | C | H | H | T | H | C | H | H | G | G | H | G | C | C | H | C | A | C | G | A | H | A | T | A | H | G | G | H |
| 2016H_P2-A12_1 | PHI4 | PHI3 | progeny | P2-A12 | | G | C | T | C | H | H | H | T | H | H | H | H | H | H | H | A | H | H | T | H | H | H | G | G | H | C | C | A | G | C | H | C | G | T | H | H | G | T | A | C | H | H | C | H | H | T | H | T | C | H | G | G | H | G | C | C | H | C | A | C | G | A | H | A | T | A | H | G | G | H |
| 2016H_P2-A12_2 | PHI4 | PHI3 | progeny | P2-A12 | | G | C | H | C | H | A | C | T | C | A | T | G | T | A | C | A | G | C | T | C | C | T | G | G | A | C | C | A | G | C | T | C | G | T | H | H | G | T | A | C | H | H | C | H | H | T | H | T | C | H | G | G | H | G | C | C | H | C | A | C | G | A | H | A | T | A | H | G | G | H |
| 2016H_P2-C1_4 | PHI4 | PHI3 | progeny | P2-C1 | | G | H | T | C | H | H | H | T | H | H | H | H | H | H | H | A | H | H | T | H | H | H | G | G | H | C | C | A | H | C | C | C | G | T | H | H | G | T | A | C | A | G | C | H | H | T | H | T | C | H | G | G | H | G | C | C | H | H | A | C | G | A | H | A | T | A | H | G | G | H |
| 2016H_P2-C1_5 | PHI4 | PHI3 | progeny | P2-C1 | | G | H | T | C | H | H | H | T | H | H | H | H | H | H | H | A | H | H | T | H | H | H | G | G | H | C | C | A | H | C | C | C | G | T | T | H | G | T | A | C | G | A | C | C | H | T | H | T | C | T | G | G | G | G | C | C | A | G | A | C | G | A | A | A | T | A | H | G | G | H |
| 2016H_P2-C1_6 | PHI4 | PHI3 | progeny | P2-C1 | | G | H | T | C | H | H | H | T | H | H | H | H | H | H | H | A | H | H | T | H | H | H | G | G | H | C | C | A | H | C | C | C | G | T | H | H | G | T | A | C | H | H | C | H | H | T | H | T | C | H | G | G | H | G | C | C | H | H | A | C | G | A | H | A | T | A | H | G | G | T |
| 2016H_P2-E9_1 | PHI4 | PHI3 | progeny | P2-E9 | | G | H | T | C | H | H | H | T | H | H | H | H | H | H | H | A | H | H | T | H | H | H | G | G | H | C | C | H | H | C | C | C | G | T | T | H | G | T | A | C | H | A | C | C | G | T | H | H | C | H | G | G | H | G | C | C | H | C | A | C | G | A | A | A | T | A | C | G | G | G |
| 2016H_P2-E9_3 | PHI4 | PHI3 | progeny | P2-E9 | | G | H | T | C | H | H | H | T | H | A | T | G | T | A | C | A | G | C | T | C | C | T | G | G | C | C | C | A | G | C | C | C | G | T | H | H | G | T | A | C | H | H | C | H | H | T | H | H | C | H | G | G | H | G | C | C | H | C | A | C | G | A | C | A | T | A | G | G | G | T |
| 2016H_P2-E8_6 | PHI4 | PHI3 | progeny | P2-E8 | | G | C | C | C | C | A | C | T | H | H | H | H | H | H | H | A | H | H | T | H | H | H | G | G | H | C | C | A | G | C | C | C | G | T | H | H | G | T | A | C | G | G | C | C | G | T | G | C | A | H | G | G | H | G | C | C | H | C | A | C | G | A | H | A | T | A | H | G | G | H |
| 2016H_P2-F11_1 | PHI4 | PHI3 | progeny | P2-F11 | | G | H | T | C | H | H | H | T | H | H | H | H | H | H | H | A | H | H | T | H | H | H | G | G | H | C | C | H | H | C | C | C | G | T | H | H | G | T | A | C | H | H | C | H | H | T | H | T | C | H | G | G | H | G | C | C | H | H | A | C | G | A | H | A | T | H | H | G | G | H |
| 2016H_P2-F11_2 | PHI4 | PHI3 | progeny | P2-F11 | | G | C | T | C | G | G | A | T | H | H | H | H | H | H | H | A | H | H | T | H | H | H | G | G | H | C | C | A | G | C | C | C | G | T | H | H | G | T | A | C | H | H | C | H | H | T | H | T | C | H | G | G | H | G | C | C | H | H | A | C | G | A | H | A | T | H | H | G | G | H |
| 2016H_P2-D6_4 | PHI4 | PHI3 | progeny | P2-D6 | | G | C | C | C | C | A | C | T | C | A | T | G | G | C | T | A | C | G | T | T | T | G | G | G | A | C | C | G | A | C | C | C | G | T | T | A | G | T | A | C | G | A | C | C | A | T | T | T | C | G | G | G | A | G | C | C | G | C | A | C | H | A | C | A | T | A | G | G | G | T |
| 2016H_P2-D6_5 | PHI4 | PHI3 | progeny | P2-D6 | | G | C | T | C | C | A | C | T | C | A | T | G | T | A | C | A | G | C | T | C | C | T | G | G | C | C | C | A | G | C | C | C | H | T | T | G | G | T | A | C | A | G | C | C | G | T | T | T | C | T | G | G | G | G | C | C | G | C | A | C | G | A | C | A | T | A | G | G | G | T |
| 2016H_P2-D6_6 | PHI4 | PHI3 | progeny | P2-D6 | | G | C | T | C | G | G | A | T | G | G | C | A | G | C | T | A | C | G | T | T | T | G | G | G | A | C | C | G | A | C | C | C | G | T | T | G | G | T | A | C | A | G | C | C | G | T | G | C | C | T | G | G | G | G | C | C | G | C | A | C | G | A | C | A | T | C | C | G | G | T |
| 2016H_P2-D9_5 | PHI4 | PHI3 | progeny | P2-D9 | | G | T | T | C | C | A | C | T | G | G | C | A | G | C | T | A | C | G | T | T | T | G | G | G | A | C | C | G | A | C | C | C | H | T | C | A | G | T | A | C | A | G | C | C | G | T | T | T | C | G | G | G | A | G | C | C | G | C | A | C | G | A | C | A | T | A | G | G | G | T |
| 2016H_P2-E1_1 | PHI4 | PHI3 | progeny | P2-E1 | | G | C | T | [illegible] | [illegible] | [illegible] | [illegible] | [illegible] | [illegible] | [illegible] | [illegible] | [illegible] | [illegible] | [illegible] | [illegible] | [illegible] | [illegible] | [illegible] | [illegible] | [illegible] | [illegible] | [illegible] | [illegible] | [illegible] | [illegible] | [illegible] | [illegible] | [illegible] | [illegible] | [illegible] | [illegible] | [illegible] | [illegible] | [illegible] | [illegible] | [illegible] | [illegible] | [illegible] | [illegible] | [illegible] | [illegible] | [illegible] | [illegible] | [illegible] | [illegible] | [illegible] | [illegible] | [illegible] | [illegible] | [illegible] | [illegible] | [illegible] | [illegible] | [illegible] | [illegible] | [illegible] | [illegible] | [illegible] | [illegible] | [illegible] | [illegible] | [illegible] | [illegible] | [illegible] | [illegible] | [illegible] |
| 2016H_P2-E1_2 | PHI4 | PHI3 | progeny | P2-E1 | | G | C | T | [illegible] | [illegible] | [illegible] | [illegible] | [illegible] | [illegible] | [illegible] | [illegible] | [illegible] | [illegible] | [illegible] | [illegible] | [illegible] | [illegible] | [illegible] | [illegible] | [illegible] | [illegible] | [illegible] | [illegible] | [illegible] | [illegible] | [illegible] | [illegible] | [illegible] | [illegible] | [illegible] | [illegible] | [illegible] | [illegible] | [illegible] | [illegible] | [illegible] | [illegible] | [illegible] | [illegible] | [illegible] | [illegible] | [illegible] | [illegible] | [illegible] | [illegible] | [illegible] | [illegible] | [illegible] | [illegible] | [illegible] | [illegible] | [illegible] | [illegible] | [illegible] | [illegible] | [illegible] | [illegible] | [illegible] | [illegible] | [illegible] | [illegible] | [illegible] | [illegible] | [illegible] | [illegible] | [illegible] |
| 2016H_P2-E1_3 | PHI4 | PHI3 | progeny | P2-E1 | | G | C | T | [illegible] | [illegible] | [illegible] | [illegible] | [illegible] | [illegible] | [illegible] | [illegible] | [illegible] | [illegible] | [illegible] | [illegible] | [illegible] | [illegible] | [illegible] | [illegible] | [illegible] | [illegible] | [illegible] | [illegible] | [illegible] | [illegible] | [illegible] | [illegible] | [illegible] | [illegible] | [illegible] | [illegible] | [illegible] | [illegible] | [illegible] | [illegible] | [illegible] | [illegible] | [illegible] | [illegible] | [illegible] | [illegible] | [illegible] | [illegible] | [illegible] | [illegible] | [illegible] | [illegible] | [illegible] | [illegible] | [illegible] | [illegible] | [illegible] | [illegible] | [illegible] | [illegible] | [illegible] | [illegible] | [illegible] | [illegible] | [illegible] | [illegible] | [illegible] | [illegible] | [illegible] | [illegible] | [illegible] |
| 2016H_P2-D3_1 | PHI4 | PHI3 | progeny | P2-D3 | | G | C | T | [illegible] | [illegible] | [illegible] | [illegible] | [illegible] | [illegible] | [illegible] | [illegible] | [illegible] | [illegible] | [illegible] | [illegible] | [illegible] | [illegible] | [illegible] | [illegible] | [illegible] | [illegible] | [illegible] | [illegible] | [illegible] | [illegible] | [illegible] | [illegible] | [illegible] | [illegible] | [illegible] | [illegible] | [illegible] | [illegible] | [illegible] | [illegible] | [illegible] | [illegible] | [illegible] | [illegible] | [illegible] | [illegible] | [illegible] | [illegible] | [illegible] | [illegible] | [illegible] | [illegible] | [illegible] | [illegible] | [illegible] | [illegible] | [illegible] | [illegible] | [illegible] | [illegible] | [illegible] | [illegible] | [illegible] | [illegible] | [illegible] | [illegible] | [illegible] | [illegible] | [illegible] | [illegible] | [illegible] |
| 2016H_P2-D3_3 | PHI4 | PHI3 | progeny | P2-D3 | | G | C | T | [illegible] | [illegible] | [illegible] | [illegible] | [illegible] | [illegible] | [illegible] | [illegible] | [illegible] | [illegible] | [illegible] | [illegible] | [illegible] | [illegible] | [illegible] | [illegible] | [illegible] | [illegible] | [illegible] | [illegible] | [illegible] | [illegible] | [illegible] | [illegible] | [illegible] | [illegible] | [illegible] | [illegible] | [illegible] | [illegible] | [illegible] | [illegible] | [illegible] | [illegible] | [illegible] | [illegible] | [illegible] | [illegible] | [illegible] | [illegible] | [illegible] | [illegible] | [illegible] | [illegible] | [illegible] | [illegible] | [illegible] | [illegible] | [illegible] | [illegible] | [illegible] | [illegible] | [illegible] | [illegible] | [illegible] | [illegible] | [illegible] | [illegible] | [illegible] | [illegible] | [illegible] | [illegible] | [illegible] |
| 2016H_P2-D3_4 | PHI4 | PHI3 | progeny | P2-D3 | | G | T | T | [illegible] | [illegible] | [illegible] | [illegible] | [illegible] | [illegible] | [illegible] | [illegible] | [illegible] | [illegible] | [illegible] | [illegible] | [illegible] | [illegible] | [illegible] | [illegible] | [illegible] | [illegible] | [illegible] | [illegible] | [illegible] | [illegible] | [illegible] | [illegible] | [illegible] | [illegible] | [illegible] | [illegible] | [illegible] | [illegible] | [illegible] | [illegible] | [illegible] | [illegible] | [illegible] | [illegible] | [illegible] | [illegible] | [illegible] | [illegible] | [illegible] | [illegible] | [illegible] | [illegible] | [illegible] | [illegible] | [illegible] | [illegible] | [illegible] | [illegible] | [illegible] | [illegible] | [illegible] | [illegible] | [illegible] | [illegible] | [illegible] | [illegible] | [illegible] | [illegible] | [illegible] | [illegible] | [illegible] |
| 2016H_P2-D3_5 | PHI4 | PHI3 | progeny | P2-D3 | | G | C | T | [illegible] | [illegible] | [illegible] | [illegible] | [illegible] | [illegible] | [illegible] | [illegible] | [illegible] | [illegible] | [illegible] | [illegible] | [illegible] | [illegible] | [illegible] | [illegible] | [illegible] | [illegible] | [illegible] | [illegible] | [illegible] | [illegible] | [illegible] | [illegible] | [illegible] | [illegible] | [illegible] | [illegible] | [illegible] | [illegible] | [illegible] | [illegible] | [illegible] | [illegible] | [illegible] | [illegible] | [illegible] | [illegible] | [illegible] | [illegible] | [illegible] | [illegible] | [illegible] | [illegible] | [illegible] | [illegible] | [illegible] | [illegible] | [illegible] | [illegible] | [illegible] | [illegible] | [illegible] | [illegible] | [illegible] | [illegible] | [illegible] | [illegible] | [illegible] | [illegible] | [illegible] | [illegible] | [illegible] |
| 2016H_P2-D3_6 | PHI4 | PHI3 | progeny | P2-D3 | | G | T | C | [illegible] | [illegible] | [illegible] | [illegible] | [illegible] | [illegible] | [illegible] | [illegible] | [illegible] | [illegible] | [illegible] | [illegible] | [illegible] | [illegible] | [illegible] | [illegible] | [illegible] | [illegible] | [illegible] | [illegible] | [illegible] | [illegible] | [illegible] | [illegible] | [illegible] | [illegible] | [illegible] | [illegible] | [illegible] | [illegible] | [illegible] | [illegible] | [illegible] | [illegible] | [illegible] | [illegible] | [illegible] | [illegible] | [illegible] | [illegible] | [illegible] | [illegible] | [illegible] | [illegible] | [illegible] | [illegible] | [illegible] | [illegible] | [illegible] | [illegible] | [illegible] | [illegible] | [illegible] | [illegible] | [illegible] | [illegible] | [illegible] | [illegible] | [illegible] | [illegible] | [illegible] | [illegible] | [illegible] |
| 2016H_P2-D2_1 | PHI4 | PHI3 | progeny | P2-D2 | | G | H | T | [illegible] | [illegible] | [illegible] | [illegible] | [illegible] | [illegible] | [illegible] | [illegible] | [illegible] | [illegible] | [illegible] | [illegible] | [illegible] | [illegible] | [illegible] | [illegible] | [illegible] | [illegible] | [illegible] | [illegible] | [illegible] | [illegible] | [illegible] | [illegible] | [illegible] | [illegible] | [illegible] | [illegible] | [illegible] | [illegible] | [illegible] | [illegible] | [illegible] | [illegible] | [illegible] | [illegible] | [illegible] | [illegible] | [illegible] | [illegible] | [illegible] | [illegible] | [illegible] | [illegible] | [illegible] | [illegible] | [illegible] | [illegible] | [illegible] | [illegible] | [illegible] | [illegible] | [illegible] | [illegible] | [illegible] | [illegible] | [illegible] | [illegible] | [illegible] | [illegible] | [illegible] | [illegible] | [illegible] |
| 2016H_P2-D2_2 | PHI4 | PHI3 | progeny | P2-D2 | | G | H | T | [illegible] | [illegible] | [illegible] | [illegible] | [illegible] | [illegible] | [illegible] | [illegible] | [illegible] | [illegible] | [illegible] | [illegible] | [illegible] | [illegible] | [illegible] | [illegible] | [illegible] | [illegible] | [illegible] | [illegible] | [illegible] | [illegible] | [illegible] | [illegible] | [illegible] | [illegible] | [illegible] | [illegible] | [illegible] | [illegible] | [illegible] | [illegible] | [illegible] | [illegible] | [illegible] | [illegible] | [illegible] | [illegible] | [illegible] | [illegible] | [illegible] | [illegible] | [illegible] | [illegible] | [illegible] | [illegible] | [illegible] | [illegible] | [illegible] | [illegible] | [illegible] | [illegible] | [illegible] | [illegible] | [illegible] | [illegible] | [illegible] | [illegible] | [illegible] | [illegible] | [illegible] | [illegible] | [illegible] |
| 2016H_P2-G8_2 | PHI4 | PHI3 | progeny | P2-G8 | | G | C | H | [illegible] | [illegible] | [illegible] | [illegible] | [illegible] | [illegible] | [illegible] | [illegible] | [illegible] | [illegible] | [illegible] | [illegible] | [illegible] | [illegible] | [illegible] | [illegible] | [illegible] | [illegible] | [illegible] | [illegible] | [illegible] | [illegible] | [illegible] | [illegible] | [illegible] | [illegible] | [illegible] | [illegible] | [illegible] | [illegible] | [illegible] | [illegible] | [illegible] | [illegible] | [illegible] | [illegible] | [illegible] | [illegible] | [illegible] | [illegible] | [illegible] | [illegible] | [illegible] | [illegible] | [illegible] | [illegible] | [illegible] | [illegible] | [illegible] | [illegible] | [illegible] | [illegible] | [illegible] | [illegible] | [illegible] | [illegible] | [illegible] | [illegible] | [illegible] | [illegible] | [illegible] | [illegible] | [illegible] |
| 2016H_P2-G8_3 | PHI4 | PHI3 | progeny | P2-G8 | | G | C | H | [illegible] | [illegible] | [illegible] | [illegible] | [illegible] | [illegible] | [illegible] | [illegible] | [illegible] | [illegible] | [illegible] | [illegible] | [illegible] | [illegible] | [illegible] | [illegible] | [illegible] | [illegible] | [illegible] | [illegible] | [illegible] | [illegible] | [illegible] | [illegible] | [illegible] | [illegible] | [illegible] | [illegible] | [illegible] | [illegible] | [illegible] | [illegible] | [illegible] | [illegible] | [illegible] | [illegible] | [illegible] | [illegible] | [illegible] | [illegible] | [illegible] | [illegible] | [illegible] | [illegible] | [illegible] | [illegible] | [illegible] | [illegible] | [illegible] | [illegible] | [illegible] | [illegible] | [illegible] | [illegible] | [illegible] | [illegible] | [illegible] | [illegible] | [illegible] | [illegible] | [illegible] | [illegible] | [illegible] |
| 2016H_P2-C7_1 | PHI4 | PHI3 | progeny | P2-C7 | | G | H | T | [illegible] | [illegible] | [illegible] | [illegible] | [illegible] | [illegible] | [illegible] | [illegible] | [illegible] | [illegible] | [illegible] | [illegible] | [illegible] | [illegible] | [illegible] | [illegible] | [illegible] | [illegible] | [illegible] | [illegible] | [illegible] | [illegible] | [illegible] | [illegible] | [illegible] | [illegible] | [illegible] | [illegible] | [illegible] | [illegible] | [illegible] | [illegible] | [illegible] | [illegible] | [illegible] | [illegible] | [illegible] | [illegible] | [illegible] | [illegible] | [illegible] | [illegible] | [illegible] | [illegible] | [illegible] | [illegible] | [illegible] | [illegible] | [illegible] | [illegible] | [illegible] | [illegible] | [illegible] | [illegible] | [illegible] | [illegible] | [illegible] | [illegible] | [illegible] | [illegible] | [illegible] | [illegible] | [illegible] |
| 2016H_P2-C6_1 | PHI4 | PHI3 | progeny | P2-C6 | | G | C | H | [illegible] | [illegible] | [illegible] | [illegible] | [illegible] | [illegible] | [illegible] | [illegible] | [illegible] | [illegible] | [illegible] | [illegible] | [illegible] | [illegible] | [illegible] | [illegible] | [illegible] | [illegible] | [illegible] | [illegible] | [illegible] | [illegible] | [illegible] | [illegible] | [illegible] | [illegible] | [illegible] | [illegible] | [illegible] | [illegible] | [illegible] | [illegible] | [illegible] | [illegible] | [illegible] | [illegible] | [illegible] | [illegible] | [illegible] | [illegible] | [illegible] | [illegible] | [illegible] | [illegible] | [illegible] | [illegible] | [illegible] | [illegible] | [illegible] | [illegible] | [illegible] | [illegible] | [illegible] | [illegible] | [illegible] | [illegible] | [illegible] | [illegible] | [illegible] | [illegible] | [illegible] | [illegible] | [illegible] |
| 2016H_P2-C6_2 | PHI4 | PHI3 | progeny | P2-C6 | | G | C | T | [illegible] | [illegible] | [illegible] | [illegible] | [illegible] | [illegible] | [illegible] | [illegible] | [illegible] | [illegible] | [illegible] | [illegible] | [illegible] | [illegible] | [illegible] | [illegible] | [illegible] | [illegible] | [illegible] | [illegible] | [illegible] | [illegible] | [illegible] | [illegible] | [illegible] | [illegible] | [illegible] | [illegible] | [illegible] | [illegible] | [illegible] | [illegible] | [illegible] | [illegible] | [illegible] | [illegible] | [illegible] | [illegible] | [illegible] | [illegible] | [illegible] | [illegible] | [illegible] | [illegible] | [illegible] | [illegible] | [illegible] | [illegible] | [illegible] | [illegible] | [illegible] | [illegible] | [illegible] | [illegible] | [illegible] | [illegible] | [illegible] | [illegible] | [illegible] | [illegible] | [illegible] | [illegible] | [illegible] |
| 2016H_P1-G1_3 | PHI10 | PHI3 | progeny | P1-G1 | | G | T | T | [illegible] | [illegible] | [illegible] | [illegible] | [illegible] | [illegible] | [illegible] | [illegible] | [illegible] | [illegible] | [illegible] | [illegible] | [illegible] | [illegible] | [illegible] | [illegible] | [illegible] | [illegible] | [illegible] | [illegible] | [illegible] | [illegible] | [illegible] | [illegible] | [illegible] | [illegible] | [illegible] | [illegible] | [illegible] | [illegible] | [illegible] | [illegible] | [illegible] | [illegible] | [illegible] | [illegible] | [illegible] | [illegible] | [illegible] | [illegible] | [illegible] | [illegible] | [illegible] | [illegible] | [illegible] | [illegible] | [illegible] | [illegible] | [illegible] | [illegible] | [illegible] | [illegible] | [illegible] | [illegible] | [illegible] | [illegible] | [illegible] | [illegible] | [illegible] | [illegible] | [illegible] | [illegible] | [illegible] |
| 2016H_P1-G1_4 | PHI10 | PHI3 | progeny | P1-G1 | | G | C | T | [illegible] | [illegible] | [illegible] | [illegible] | [illegible] | [illegible] | [illegible] | [illegible] | [illegible] | [illegible] | [illegible] | [illegible] | [illegible] | [illegible] | [illegible] | [illegible] | [illegible] | [illegible] | [illegible] | [illegible] | [illegible] | [illegible] | [illegible] | [illegible] | [illegible] | [illegible] | [illegible] | [illegible] | [illegible] | [illegible] | [illegible] | [illegible] | [illegible] | [illegible] | [illegible] | [illegible] | [illegible] | [illegible] | [illegible] | [illegible] | [illegible] | [illegible] | [illegible] | [illegible] | [illegible] | [illegible] | [illegible] | [illegible] | [illegible] | [illegible] | [illegible] | [illegible] | [illegible] | [illegible] | [illegible] | [illegible] | [illegible] | [illegible] | [illegible] | [illegible] | [illegible] | [illegible] | [illegible] |
| 2016H_P1-G1_5 | PHI10 | PHI3 | progeny | P1-G1 | | G | C | T | [illegible] | [illegible] | [illegible] | [illegible] | [illegible] | [illegible] | [illegible] | [illegible] | [illegible] | [illegible] | [illegible] | [illegible] | [illegible] | [illegible] | [illegible] | [illegible] | [illegible] | [illegible] | [illegible] | [illegible] | [illegible] | [illegible] | [illegible] | [illegible] | [illegible] | [illegible] | [illegible] | [illegible] | [illegible] | [illegible] | [illegible] | [illegible] | [illegible] | [illegible] | [illegible] | [illegible] | [illegible] | [illegible] | [illegible] | [illegible] | [illegible] | [illegible] | [illegible] | [illegible] | [illegible] | [illegible] | [illegible] | [illegible] | [illegible] | [illegible] | [illegible] | [illegible] | [illegible] | [illegible] | [illegible] | [illegible] | [illegible] | [illegible] | [illegible] | [illegible] | [illegible] | [illegible] | [illegible] |
| 2016H_P1-G1_6 | PHI10 | PHI3 | progeny | P1-G1 | | G | T | T | [illegible] | [illegible] | [illegible] | [illegible] | [illegible] | [illegible] | [illegible] | [illegible] | [illegible] | [illegible] | [illegible] | [illegible] | [illegible] | [illegible] | [illegible] | [illegible] | [illegible] | [illegible] | [illegible] | [illegible] | [illegible] | [illegible] | [illegible] | [illegible] | [illegible] | [illegible] | [illegible] | [illegible] | [illegible] | [illegible] | [illegible] | [illegible] | [illegible] | [illegible] | [illegible] | [illegible] | [illegible] | [illegible] | [illegible] | [illegible] | [illegible] | [illegible] | [illegible] | [illegible] | [illegible] | [illegible] | [illegible] | [illegible] | [illegible] | [illegible] | [illegible] | [illegible] | [illegible] | [illegible] | [illegible] | [illegible] | [illegible] | [illegible] | [illegible] | [illegible] | [illegible] | [illegible] | [illegible] |
| 2016H_P1-F4_2 | PHI10 | PHI3 | progeny | P1-F4 | | G | T | T | [illegible] | [illegible] | [illegible] | [illegible] | [illegible] | [illegible] | [illegible] | [illegible] | [illegible] | [illegible] | [illegible] | [illegible] | [illegible] | [illegible] | [illegible] | [illegible] | [illegible] | [illegible] | [illegible] | [illegible] | [illegible] | [illegible] | [illegible] | [illegible] | [illegible] | [illegible] | [illegible] | [illegible] | [illegible] | [illegible] | [illegible] | [illegible] | [illegible] | [illegible] | [illegible] | [illegible] | [illegible] | [illegible] | [illegible] | [illegible] | [illegible] | [illegible] | [illegible] | [illegible] | [illegible] | [illegible] | [illegible] | [illegible] | [illegible] | [illegible] | [illegible] | [illegible] | [illegible] | [illegible] | [illegible] | [illegible] | [illegible] | [illegible] | [illegible] | [illegible] | [illegible] | [illegible] | [illegible] |
| 2016H_P1-F4_3 | PHI10 | PHI3 | progeny | P1-F4 | | G | T | T | [illegible] | [illegible] | [illegible] | [illegible] | [illegible] | [illegible] | [illegible] | [illegible] | [illegible] | [illegible] | [illegible] | [illegible] | [illegible] | [illegible] | [illegible] | [illegible] | [illegible] | [illegible] | [illegible] | [illegible] | [illegible] | [illegible] | [illegible] | [illegible] | [illegible] | [illegible] | [illegible] | [illegible] | [illegible] | [illegible] | [illegible] | [illegible] | [illegible] | [illegible] | [illegible] | [illegible] | [illegible] | [illegible] | [illegible] | [illegible] | [illegible] | [illegible] | [illegible] | [illegible] | [illegible] | [illegible] | [illegible] | [illegible] | [illegible] | [illegible] | [illegible] | [illegible] | [illegible] | [illegible] | [illegible] | [illegible] | [illegible] | [illegible] | [illegible] | [illegible] | [illegible] | [illegible] | [illegible] |
| 2016H_P1-F4_4 | PHI10 | PHI3 | progeny | P1-F4 | | G | T | T | [illegible] | [illegible] | [illegible] | [illegible] | [illegible] | [illegible] | [illegible] | [illegible] | [illegible] | [illegible] | [illegible] | [illegible] | [illegible] | [illegible] | [illegible] | [illegible] | [illegible] | [illegible] | [illegible] | [illegible] | [illegible] | [illegible] | [illegible] | [illegible] | [illegible] | [illegible] | [illegible] | [illegible] | [illegible] | [illegible] | [illegible] | [illegible] | [illegible] | [illegible] | [illegible] | [illegible] | [illegible] | [illegible] | [illegible] | [illegible] | [illegible] | [illegible] | [illegible] | [illegible] | [illegible] | [illegible] | [illegible] | [illegible] | [illegible] | [illegible] | [illegible] | [illegible] | [illegible] | [illegible] | [illegible] | [illegible] | [illegible] | [illegible] | [illegible] | [illegible] | [illegible] | [illegible] | [illegible] |
| 2016H_P1-F5_1 | PHI10 | PHI3 | progeny | P1-F5 | | G | C | T | [illegible] | [illegible] | [illegible] | [illegible] | [illegible] | [illegible] | [illegible] | [illegible] | [illegible] | [illegible] | [illegible] | [illegible] | [illegible] | [illegible] | [illegible] | [illegible] | [illegible] | [illegible] | [illegible] | [illegible] | [illegible] | [illegible] | [illegible] | [illegible] | [illegible] | [illegible] | [illegible] | [illegible] | [illegible] | [illegible] | [illegible] | [illegible] | [illegible] | [illegible] | [illegible] | [illegible] | [illegible] | [illegible] | [illegible] | [illegible] | [illegible] | [illegible] | [illegible] | [illegible] | [illegible] | [illegible] | [illegible] | [illegible] | [illegible] | [illegible] | [illegible] | [illegible] | [illegible] | [illegible] | [illegible] | [illegible] | [illegible] | [illegible] | [illegible] | [illegible] | [illegible] | [illegible] | [illegible] |
| 2016H_P1-F11_6 | PHI10 | PHI3 | progeny | P1-F11 | | G | C | T | [illegible] | [illegible] | [illegible] | [illegible] | [illegible] | [illegible] | [illegible] | [illegible] | [illegible] | [illegible] | [illegible] | [illegible] | [illegible] | [illegible] | [illegible] | [illegible] | [illegible] | [illegible] | [illegible] | [illegible] | [illegible] | [illegible] | [illegible] | [illegible] | [illegible] | [illegible] | [illegible] | [illegible] | [illegible] | [illegible] | [illegible] | [illegible] | [illegible] | [illegible] | [illegible] | [illegible] | [illegible] | [illegible] | [illegible] | [illegible] | [illegible] | [illegible] | [illegible] | [illegible] | [illegible] | [illegible] | [illegible] | [illegible] | [illegible] | [illegible] | [illegible] | [illegible] | [illegible] | [illegible] | [illegible] | [illegible] | [illegible] | [illegible] | [illegible] | [illegible] | [illegible] | [illegible] | [illegible] |
| 2016H_P1-G9_11 | PHI10 | PHI3 | progeny | P1-G9 | | G | C | T | [illegible] | [illegible] | [illegible] | [illegible] | [illegible] | [illegible] | [illegible] | [illegible] | [illegible] | [illegible] | [illegible] | [illegible] | [illegible] | [illegible] | [illegible] | [illegible] | [illegible] | [illegible] | [illegible] | [illegible] | [illegible] | [illegible] | [illegible] | [illegible] | [illegible] | [illegible] | [illegible] | [illegible] | [illegible] | [illegible] | [illegible] | [illegible] | [illegible] | [illegible] | [illegible] | [illegible] | [illegible] | [illegible] | [illegible] | [illegible] | [illegible] | [illegible] | [illegible] | [illegible] | [illegible] | [illegible] | [illegible] | [illegible] | [illegible] | [illegible] | [illegible] | [illegible] | [illegible] | [illegible] | [illegible] | [illegible] | [illegible] | [illegible] | [illegible] | [illegible] | [illegible] | [illegible] | [illegible] |
| 2016H_P1-G2_1 | PHI10 | PHI3 | progeny | P1-G2 | | G | T | T | [illegible] | [illegible] | [illegible] | [illegible] | [illegible] | [illegible] | [illegible] | [illegible] | [illegible] | [illegible] | [illegible] | [illegible] | [illegible] | [illegible] | [illegible] | [illegible] | [illegible] | [illegible] | [illegible] | [illegible] | [illegible] | [illegible] | [illegible] | [illegible] | [illegible] | [illegible] | [illegible] | [illegible] | [illegible] | [illegible] | [illegible] | [illegible] | [illegible] | [illegible] | [illegible] | [illegible] | [illegible] | [illegible] | [illegible] | [illegible] | [illegible] | [illegible] | [illegible] | [illegible] | [illegible] | [illegible] | [illegible] | [illegible] | [illegible] | [illegible] | [illegible] | [illegible] | [illegible] | [illegible] | [illegible] | [illegible] | [illegible] | [illegible] | [illegible] | [illegible] | [illegible] | [illegible] | [illegible] |
| 2016H_P1-G2_2 | PHI10 | PHI3 | progeny | P1-G2 | | G | C | T | [illegible] | [illegible] | [illegible] | [illegible] | [illegible] | [illegible] | [illegible] | [illegible] | [illegible] | [illegible] | [illegible] | [illegible] | [illegible] | [illegible] | [illegible] | [illegible] | [illegible] | [illegible] | [illegible] | [illegible] | [illegible] | [illegible] | [illegible] | [illegible] | [illegible] | [illegible] | [illegible] | [illegible] | [illegible] | [illegible] | [illegible] | [illegible] | [illegible] | [illegible] | [illegible] | [illegible] | [illegible] | [illegible] | [illegible] | [illegible] | [illegible] | [illegible] | [illegible] | [illegible] | [illegible] | [illegible] | [illegible] | [illegible] | [illegible] | [illegible] | [illegible] | [illegible] | [illegible] | [illegible] | [illegible] | [illegible] | [illegible] | [illegible] | [illegible] | [illegible] | [illegible] | [illegible] | [illegible] |
| 2016H_P1-G5_2 | PHI10 | PHI3 | progeny | P1-G5 | | G | C | T | [illegible] | [illegible] | [illegible] | [illegible] | [illegible] | [illegible] | [illegible] | [illegible] | [illegible] | [illegible] | [illegible] | [illegible] | [illegible] | [illegible] | [illegible] | [illegible] | [illegible] | [illegible] | [illegible] | [illegible] | [illegible] | [illegible] | [illegible] | [illegible] | [illegible] | [illegible] | [illegible] | [illegible] | [illegible] | [illegible] | [illegible] | [illegible] | [illegible] | [illegible] | [illegible] | [illegible] | [illegible] | [illegible] | [illegible] | [illegible] | [illegible] | [illegible] | [illegible] | [illegible] | [illegible] | [illegible] | [illegible] | [illegible] | [illegible] | [illegible] | [illegible] | [illegible] | [illegible] | [illegible] | [illegible] | [illegible] | [illegible] | [illegible] | [illegible] | [illegible] | [illegible] | [illegible] | [illegible] |
| 2016H_P1-G5_4 | PHI10 | PHI3 | progeny | P1-G5 | | G | C | T | [illegible] | [illegible] | [illegible] | [illegible] | [illegible] | [illegible] | [illegible] | [illegible] | [illegible] | [illegible] | [illegible] | [illegible] | [illegible] | [illegible] | [illegible] | [illegible] | [illegible] | [illegible] | [illegible] | [illegible] | [illegible] | [illegible] | [illegible] | [illegible] | [illegible] | [illegible] | [illegible] | [illegible] | [illegible] | [illegible] | [illegible] | [illegible] | [illegible] | [illegible] | [illegible] | [illegible] | [illegible] | [illegible] | [illegible] | [illegible] | [illegible] | [illegible] | [illegible] | [illegible] | [illegible] | [illegible] | [illegible] | [illegible] | [illegible] | [illegible] | [illegible] | [illegible] | [illegible] | [illegible] | [illegible] | [illegible] | [illegible] | [illegible] | [illegible] | [illegible] | [illegible] | [illegible] | [illegible] |
| 2016H_P1-F9_1 | PHI10 | PHI3 | progeny | P1-F9 | | G | C | T | [illegible] | [illegible] | [illegible] | [illegible] | [illegible] | [illegible] | [illegible] | [illegible] | [illegible] | [illegible] | [illegible] | [illegible] | [illegible] | [illegible] | [illegible] | [illegible] | [illegible] | [illegible] | [illegible] | [illegible] | [illegible] | [illegible] | [illegible] | [illegible] | [illegible] | [illegible] | [illegible] | [illegible] | [illegible] | [illegible] | [illegible] | [illegible] | [illegible] | [illegible] | [illegible] | [illegible] | [illegible] | [illegible] | [illegible] | [illegible] | [illegible] | [illegible] | [illegible] | [illegible] | [illegible] | [illegible] | [illegible] | [illegible] | [illegible] | [illegible] | [illegible] | [illegible] | [illegible] | [illegible] | [illegible] | [illegible] | [illegible] | [illegible] | [illegible] | [illegible] | [illegible] | [illegible] | [illegible] |
| 2016H_P1-F9_2 | PHI10 | PHI3 | progeny | P1-F9 | | G | T | T | [illegible] | [illegible] | [illegible] | [illegible] | [illegible] | [illegible] | [illegible] | [illegible] | [illegible] | [illegible] | [illegible] | [illegible] | [illegible] | [illegible] | [illegible] | [illegible] | [illegible] | [illegible] | [illegible] | [illegible] | [illegible] | [illegible] | [illegible] | [illegible] | [illegible] | [illegible] | [illegible] | [illegible] | [illegible] | [illegible] | [illegible] | [illegible] | [illegible] | [illegible] | [illegible] | [illegible] | [illegible] | [illegible] | [illegible] | [illegible] | [illegible] | [illegible] | [illegible] | [illegible] | [illegible] | [illegible] | [illegible] | [illegible] | [illegible] | [illegible] | [illegible] | [illegible] | [illegible] | [illegible] | [illegible] | [illegible] | [illegible] | [illegible] | [illegible] | [illegible] | [illegible] | [illegible] | [illegible] |
| 2016H_P1-F3_4 | PHI10 | PHI3 | progeny | P1-F3 | | G | C | T | [illegible] | [illegible] | [illegible] | [illegible] | [illegible] | [illegible] | [illegible] | [illegible] | [illegible] | [illegible] | [illegible] | [illegible] | [illegible] | [illegible] | [illegible] | [illegible] | [illegible] | [illegible] | [illegible] | [illegible] | [illegible] | [illegible] | [illegible] | [illegible] | [illegible] | [illegible] | [illegible] | [illegible] | [illegible] | [illegible] | [illegible] | [illegible] | [illegible] | [illegible] | [illegible] | [illegible] | [illegible] | [illegible] | [illegible] | [illegible] | [illegible] | [illegible] | [illegible] | [illegible] | [illegible] | [illegible] | [illegible] | [illegible] | [illegible] | [illegible] | [illegible] | [illegible] | [illegible] | [illegible] | [illegible] | [illegible] | [illegible] | [illegible] | [illegible] | [illegible] | [illegible] | [illegible] | [illegible] |
| 2016H_P1-F3_9 | PHI10 | PHI3 | progeny | P1-F3 | | G | T | T | [illegible] | [illegible] | [illegible] | [illegible] | [illegible] | [illegible] | [illegible] | [illegible] | [illegible] | [illegible] | [illegible] | [illegible] | [illegible] | [illegible] | [illegible] | [illegible] | [illegible] | [illegible] | [illegible] | [illegible] | [illegible] | [illegible] | [illegible] | [illegible] | [illegible] | [illegible] | [illegible] | [illegible] | [illegible] | [illegible] | [illegible] | [illegible] | [illegible] | [illegible] | [illegible] | [illegible] | [illegible] | [illegible] | [illegible] | [illegible] | [illegible] | [illegible] | [illegible] | [illegible] | [illegible] | [illegible] | [illegible] | [illegible] | [illegible] | [illegible] | [illegible] | [illegible] | [illegible] | [illegible] | [illegible] | [illegible] | [illegible] | [illegible] | [illegible] | [illegible] | [illegible] | [illegible] | [illegible] |
| 2016H_P1-F10_2 | PHI10 | PHI3 | progeny | P1-F10 | | G | C | T | [illegible] | [illegible] | [illegible] | [illegible] | [illegible] | [illegible] | [illegible] | [illegible] | [illegible] | [illegible] | [illegible] | [illegible] | [illegible] | [illegible] | [illegible] | [illegible] | [illegible] | [illegible] | [illegible] | [illegible] | [illegible] | [illegible] | [illegible] | [illegible] | [illegible] | [illegible] | [illegible] | [illegible] | [illegible] | [illegible] | [illegible] | [illegible] | [illegible] | [illegible] | [illegible] | [illegible] | [illegible] | [illegible] | [illegible] | [illegible] | [illegible] | [illegible] | [illegible] | [illegible] | [illegible] | [illegible] | [illegible] | [illegible] | [illegible] | [illegible] | [illegible] | [illegible] | [illegible] | [illegible] | [illegible] | [illegible] | [illegible] | [illegible] | [illegible] | [illegible] | [illegible] | [illegible] | [illegible] |
| 2016H_P1-F10_6 | PHI10 | PHI3 | progeny | P1-F10 | | G | T | T | [illegible] | [illegible] | [illegible] | [illegible] | [illegible] | [illegible] | [illegible] | [illegible] | [illegible] | [illegible] | [illegible] | [illegible] | [illegible] | [illegible] | [illegible] | [illegible] | [illegible] | [illegible] | [illegible] | [illegible] | [illegible] | [illegible] | [illegible] | [illegible] | [illegible] | [illegible] | [illegible] | [illegible] | [illegible] | [illegible] | [illegible] | [illegible] | [illegible] | [illegible] | [illegible] | [illegible] | [illegible] | [illegible] | [illegible] | [illegible] | [illegible] | [illegible] | [illegible] | [illegible] | [illegible] | [illegible] | [illegible] | [illegible] | [illegible] | [illegible] | [illegible] | [illegible] | [illegible] | [illegible] | [illegible] | [illegible] | [illegible] | [illegible] | [illegible] | [illegible] | [illegible] | [illegible] | [illegible] |
| 2016H_P1-F10_7 | PHI10 | PHI3 | progeny | P1-F10 | | G | T | T | [illegible] | [illegible] | [illegible] | [illegible] | [illegible] | [illegible] | [illegible] | [illegible] | [illegible] | [illegible] | [illegible] | [illegible] | [illegible] | [illegible] | [illegible] | [illegible] | [illegible] | [illegible] | [illegible] | [illegible] | [illegible] | [illegible] | [illegible] | [illegible] | [illegible] | [illegible] | [illegible] | [illegible] | [illegible] | [illegible] | [illegible] | [illegible] | [illegible] | [illegible] | [illegible] | [illegible] | [illegible] | [illegible] | [illegible] | [illegible] | [illegible] | [illegible] | [illegible] | [illegible] | [illegible] | [illegible] | [illegible] | [illegible] | [illegible] | [illegible] | [illegible] | [illegible] | [illegible] | [illegible] | [illegible] | [illegible] | [illegible] | [illegible] | [illegible] | [illegible] | [illegible] | [illegible] | [illegible] |
| 2016H_P1-F10_9 | PHI10 | PHI3 | progeny | P1-F10 | | G | C | T | [illegible] | [illegible] | [illegible] | [illegible] | [illegible] | [illegible] | [illegible] | [illegible] | [illegible] | [illegible] | [illegible] | [illegible] | [illegible] | [illegible] | [illegible] | [illegible] | [illegible] | [illegible] | [illegible] | [illegible] | [illegible] | [illegible] | [illegible] | [illegible] | [illegible] | [illegible] | [illegible] | [illegible] | [illegible] | [illegible] | [illegible] | [illegible] | [illegible] | [illegible] | [illegible] | [illegible] | [illegible] | [illegible] | [illegible] | [illegible] | [illegible] | [illegible] | [illegible] | [illegible] | [illegible] | [illegible] | [illegible] | [illegible] | [illegible] | [illegible] | [illegible] | [illegible] | [illegible] | [illegible] | [illegible] | [illegible] | [illegible] | [illegible] | [illegible] | [illegible] | [illegible] | [illegible] | [illegible] |
| 2016H_P5-G3_4 | PHI9 | PHI3 | progeny | P5-G3 | | G | C | T | [illegible] | [illegible] | [illegible] | [illegible] | [illegible] | [illegible] | [illegible] | [illegible] | [illegible] | [illegible] | [illegible] | [illegible] | [illegible] | [illegible] | [illegible] | [illegible] | [illegible] | [illegible] | [illegible] | [illegible] | [illegible] | [illegible] | [illegible] | [illegible] | [illegible] | [illegible] | [illegible] | [illegible] | [illegible] | [illegible] | [illegible] | [illegible] | [illegible] | [illegible] | [illegible] | [illegible] | [illegible] | [illegible] | [illegible] | [illegible] | [illegible] | [illegible] | [illegible] | [illegible] | [illegible] | [illegible] | [illegible] | [illegible] | [illegible] | [illegible] | [illegible] | [illegible] | [illegible] | [illegible] | [illegible] | [illegible] | [illegible] | [illegible] | [illegible] | [illegible] | [illegible] | [illegible] | [illegible] |
| 2016H_P5-E5_1 | PHI9 | PHI3 | progeny | P5-E5 | | G | T | T | [illegible] | [illegible] | [illegible] | [illegible] | [illegible] | [illegible] | [illegible] | [illegible] | [illegible] | [illegible] | [illegible] | [illegible] | [illegible] | [illegible] | [illegible] | [illegible] | [illegible] | [illegible] | [illegible] | [illegible] | [illegible] | [illegible] | [illegible] | [illegible] | [illegible] | [illegible] | [illegible] | [illegible] | [illegible] | [illegible] | [illegible] | [illegible] | [illegible] | [illegible] | [illegible] | [illegible] | [illegible] | [illegible] | [illegible] | [illegible] | [illegible] | [illegible] | [illegible] | [illegible] | [illegible] | [illegible] | [illegible] | [illegible] | [illegible] | [illegible] | [illegible] | [illegible] | [illegible] | [illegible] | [illegible] | [illegible] | [illegible] | [illegible] | [illegible] | [illegible] | [illegible] | [illegible] | [illegible] |
| 2016H_P5-E5_2 | PHI9 | PHI3 | progeny | P5-E5 | | G | T | T | [illegible] | [illegible] | [illegible] | [illegible] | [illegible] | [illegible] | [illegible] | [illegible] | [illegible] | [illegible] | [illegible] | [illegible] | [illegible] | [illegible] | [illegible] | [illegible] | [illegible] | [illegible] | [illegible] | [illegible] | [illegible] | [illegible] | [illegible] | [illegible] | [illegible] | [illegible] | [illegible] | [illegible] | [illegible] | [illegible] | [illegible] | [illegible] | [illegible] | [illegible] | [illegible] | [illegible] | [illegible] | [illegible] | [illegible] | [illegible] | [illegible] | [illegible] | [illegible] | [illegible] | [illegible] | [illegible] | [illegible] | [illegible] | [illegible] | [illegible] | [illegible] | [illegible] | [illegible] | [illegible] | [illegible] | [illegible] | [illegible] | [illegible] | [illegible] | [illegible] | [illegible] | [illegible] | [illegible] |
| 2016H_P5-E5_3 | PHI9 | PHI3 | progeny | P5-E5 | | G | T | T | [illegible] | [illegible] | [illegible] | [illegible] | [illegible] | [illegible] | [illegible] | [illegible] | [illegible] | [illegible] | [illegible] | [illegible] | [illegible] | [illegible] | [illegible] | [illegible] | [illegible] | [illegible] | [illegible] | [illegible] | [illegible] | [illegible] | [illegible] | [illegible] | [illegible] | [illegible] | [illegible] | [illegible] | [illegible] | [illegible] | [illegible] | [illegible] | [illegible] | [illegible] | [illegible] | [illegible] | [illegible] | [illegible] | [illegible] | [illegible] | [illegible] | [illegible] | [illegible] | [illegible] | [illegible] | [illegible] | [illegible] | [illegible] | [illegible] | [illegible] | [illegible] | [illegible] | [illegible] | [illegible] | [illegible] | [illegible] | [illegible] | [illegible] | [illegible] | [illegible] | [illegible] | [illegible] | [illegible] |
| 2016H_P5-E5_7 | PHI9 | PHI3 | progeny | P5-E5 | | G | T | T | [illegible] | [illegible] | [illegible] | [illegible] | [illegible] | [illegible] | [illegible] | [illegible] | [illegible] | [illegible] | [illegible] | [illegible] | [illegible] | [illegible] | [illegible] | [illegible] | [illegible] | [illegible] | [illegible] | [illegible] | [illegible] | [illegible] | [illegible] | [illegible] | [illegible] | [illegible] | [illegible] | [illegible] | [illegible] | [illegible] | [illegible] | [illegible] | [illegible] | [illegible] | [illegible] | [illegible] | [illegible] | [illegible] | [illegible] | [illegible] | [illegible] | [illegible] | [illegible] | [illegible] | [illegible] | [illegible] | [illegible] | [illegible] | [illegible] | [illegible] | [illegible] | [illegible] | [illegible] | [illegible] | [illegible] | [illegible] | [illegible] | [illegible] | [illegible] | [illegible] | [illegible] | [illegible] | [illegible] |
| 2016H_P5-E5_9 | PHI9 | PHI3 | progeny | P5-E5 | | G | C | T | [illegible] | [illegible] | [illegible] | [illegible] | [illegible] | [illegible] | [illegible] | [illegible] | [illegible] | [illegible] | [illegible] | [illegible] | [illegible] | [illegible] | [illegible] | [illegible] | [illegible] | [illegible] | [illegible] | [illegible] | [illegible] | [illegible] | [illegible] | [illegible] | [illegible] | [illegible] | [illegible] | [illegible] | [illegible] | [illegible] | [illegible] | [illegible] | [illegible] | [illegible] | [illegible] | [illegible] | [illegible] | [illegible] | [illegible] | [illegible] | [illegible] | [illegible] | [illegible] | [illegible] | [illegible] | [illegible] | [illegible] | [illegible] | [illegible] | [illegible] | [illegible] | [illegible] | [illegible] | [illegible] | [illegible] | [illegible] | [illegible] | [illegible] | [illegible] | [illegible] | [illegible] | [illegible] | [illegible] |
| 2016H_P5-E5_10 | PHI9 | PHI3 | progeny | P5-E5 | | G | T | T | [illegible] | [illegible] | [illegible] | [illegible] | [illegible] | [illegible] | [illegible] | [illegible] | [illegible] | [illegible] | [illegible] | [illegible] | [illegible] | [illegible] | [illegible] | [illegible] | [illegible] | [illegible] | [illegible] | [illegible] | [illegible] | [illegible] | [illegible] | [illegible] | [illegible] | [illegible] | [illegible] | [illegible] | [illegible] | [illegible] | [illegible] | [illegible] | [illegible] | [illegible] | [illegible] | [illegible] | [illegible] | [illegible] | [illegible] | [illegible] | [illegible] | [illegible] | [illegible] | [illegible] | [illegible] | [illegible] | [illegible] | [illegible] | [illegible] | [illegible] | [illegible] | [illegible] | [illegible] | [illegible] | [illegible] | [illegible] | [illegible] | [illegible] | [illegible] | [illegible] | [illegible] | [illegible] | [illegible] |
| 2016H_P5-E5_11 | PHI9 | PHI3 | progeny | P5-E5 | | G | T | H | [illegible] | [illegible] | [illegible] | [illegible] | [illegible] | [illegible] | [illegible] | [illegible] | [illegible] | [illegible] | [illegible] | [illegible] | [illegible] | [illegible] | [illegible] | [illegible] | [illegible] | [illegible] | [illegible] | [illegible] | [illegible] | [illegible] | [illegible] | [illegible] | [illegible] | [illegible] | [illegible] | [illegible] | [illegible] | [illegible] | [illegible] | [illegible] | [illegible] | [illegible] | [illegible] | [illegible] | [illegible] | [illegible] | [illegible] | [illegible] | [illegible] | [illegible] | [illegible] | [illegible] | [illegible] | [illegible] | [illegible] | [illegible] | [illegible] | [illegible] | [illegible] | [illegible] | [illegible] | [illegible] | [illegible] | [illegible] | [illegible] | [illegible] | [illegible] | [illegible] | [illegible] | [illegible] | [illegible] |
| 2016H_P5-G4_1 | PHI9 | PHI3 | progeny | P5-G4 | | G | C | T | [illegible] | [illegible] | [illegible] | [illegible] | [illegible] | [illegible] | [illegible] | [illegible] | [illegible] | [illegible] | [illegible] | [illegible] | [illegible] | [illegible] | [illegible] | [illegible] | [illegible] | [illegible] | [illegible] | [illegible] | [illegible] | [illegible] | [illegible] | [illegible] | [illegible] | [illegible] | [illegible] | [illegible] | [illegible] | [illegible] | [illegible] | [illegible] | [illegible] | [illegible] | [illegible] | [illegible] | [illegible] | [illegible] | [illegible] | [illegible] | [illegible] | [illegible] | [illegible] | [illegible] | [illegible] | [illegible] | [illegible] | [illegible] | [illegible] | [illegible] | [illegible] | [illegible] | [illegible] | [illegible] | [illegible] | [illegible] | [illegible] | [illegible] | [illegible] | [illegible] | [illegible] | [illegible] | [illegible] |
| 2016H_P5-F10_1 | PHI9 | PHI3 | progeny | P5-F10 | | G | H | T | [illegible] | [illegible] | [illegible] | [illegible] | [illegible] | [illegible] | [illegible] | [illegible] | [illegible] | [illegible] | [illegible] | [illegible] | [illegible] | [illegible] | [illegible] | [illegible] | [illegible] | [illegible] | [illegible] | [illegible] | [illegible] | [illegible] | [illegible] | [illegible] | [illegible] | [illegible] | [illegible] | [illegible] | [illegible] | [illegible] | [illegible] | [illegible] | [illegible] | [illegible] | [illegible] | [illegible] | [illegible] | [illegible] | [illegible] | [illegible] | [illegible] | [illegible] | [illegible] | [illegible] | [illegible] | [illegible] | [illegible] | [illegible] | [illegible] | [illegible] | [illegible] | [illegible] | [illegible] | [illegible] | [illegible] | [illegible] | [illegible] | [illegible] | [illegible] | [illegible] | [illegible] | [illegible] | [illegible] |
| 2016H_P5-F2_1 | PHI9 | PHI3 | progeny | P5-F2 | | G | H | T | [illegible] | [illegible] | [illegible] | [illegible] | [illegible] | [illegible] | [illegible] | [illegible] | [illegible] | [illegible] | [illegible] | [illegible] | [illegible] | [illegible] | [illegible] | [illegible] | [illegible] | [illegible] | [illegible] | [illegible] | [illegible] | [illegible] | [illegible] | [illegible] | [illegible] | [illegible] | [illegible] | [illegible] | [illegible] | [illegible] | [illegible] | [illegible] | [illegible] | [illegible] | [illegible] | [illegible] | [illegible] | [illegible] | [illegible] | [illegible] | [illegible] | [illegible] | [illegible] | [illegible] | [illegible] | [illegible] | [illegible] | [illegible] | [illegible] | [illegible] | [illegible] | [illegible] | [illegible] | [illegible] | [illegible] | [illegible] | [illegible] | [illegible] | [illegible] | [illegible] | [illegible] | [illegible] | [illegible] |
| 2016H_P5-E7_3 | PHI9 | PHI3 | progeny | P5-E7 | | G | T | T | [illegible] | [illegible] | [illegible] | [illegible] | [illegible] | [illegible] | [illegible] | [illegible] | [illegible] | [illegible] | [illegible] | [illegible] | [illegible] | [illegible] | [illegible] | [illegible] | [illegible] | [illegible] | [illegible] | [illegible] | [illegible] | [illegible] | [illegible] | [illegible] | [illegible] | [illegible] | [illegible] | [illegible] | [illegible] | [illegible] | [illegible] | [illegible] | [illegible] | [illegible] | [illegible] | [illegible] | [illegible] | [illegible] | [illegible] | [illegible] | [illegible] | [illegible] | [illegible] | [illegible] | [illegible] | [illegible] | [illegible] | [illegible] | [illegible] | [illegible] | [illegible] | [illegible] | [illegible] | [illegible] | [illegible] | [illegible] | [illegible] | [illegible] | [illegible] | [illegible] | [illegible] | [illegible] | [illegible] |
| 2016H_P1-C9_1 | PHI2 | PHI3 | progeny | P1-C9 | | G | C | H | [illegible] | [illegible] | [illegible] | [illegible] | [illegible] | [illegible] | [illegible] | [illegible] | [illegible] | [illegible] | [illegible] | [illegible] | [illegible] | [illegible] | [illegible] | [illegible] | [illegible] | [illegible] | [illegible] | [illegible] | [illegible] | [illegible] | [illegible] | [illegible] | [illegible] | [illegible] | [illegible] | [illegible] | [illegible] | [illegible] | [illegible] | [illegible] | [illegible] | [illegible] | [illegible] | [illegible] | [illegible] | [illegible] | [illegible] | [illegible] | [illegible] | [illegible] | [illegible] | [illegible] | [illegible] | [illegible] | [illegible] | [illegible] | [illegible] | [illegible] | [illegible] | [illegible] | [illegible] | [illegible] | [illegible] | [illegible] | [illegible] | [illegible] | [illegible] | [illegible] | [illegible] | [illegible] | [illegible] |
| 2016H_P1-C9_2 | PHI2 | PHI3 | progeny | P1-C9 | | G | C | T | [illegible] | [illegible] | [illegible] | [illegible] | [illegible] | [illegible] | [illegible] | [illegible] | [illegible] | [illegible] | [illegible] | [illegible] | [illegible] | [illegible] | [illegible] | [illegible] | [illegible] | [illegible] | [illegible] | [illegible] | [illegible] | [illegible] | [illegible] | [illegible] | [illegible] | [illegible] | [illegible] | [illegible] | [illegible] | [illegible] | [illegible] | [illegible] | [illegible] | [illegible] | [illegible] | [illegible] | [illegible] | [illegible] | [illegible] | [illegible] | [illegible] | [illegible] | [illegible] | [illegible] | [illegible] | [illegible] | [illegible] | [illegible] | [illegible] | [illegible] | [illegible] | [illegible] | [illegible] | [illegible] | [illegible] | [illegible] | [illegible] | [illegible] | [illegible] | [illegible] | [illegible] | [illegible] | [illegible] |
| 2016H_P1-C9_3 | PHI2 | PHI3 | progeny | P1-C9 | | G | C | H | [illegible] | [illegible] | [illegible] | [illegible] | [illegible] | [illegible] | [illegible] | [illegible] | [illegible] | [illegible] | [illegible] | [illegible] | [illegible] | [illegible] | [illegible] | [illegible] | [illegible] | [illegible] | [illegible] | [illegible] | [illegible] | [illegible] | [illegible] | [illegible] | [illegible] | [illegible] | [illegible] | [illegible] | [illegible] | [illegible] | [illegible] | [illegible] | [illegible] | [illegible] | [illegible] | [illegible] | [illegible] | [illegible] | [illegible] | [illegible] | [illegible] | [illegible] | [illegible] | [illegible] | [illegible] | [illegible] | [illegible] | [illegible] | [illegible] | [illegible] | [illegible] | [illegible] | [illegible] | [illegible] | [illegible] | [illegible] | [illegible] | [illegible] | [illegible] | [illegible] | [illegible] | [illegible] | [illegible] |
| 2016H_P1-C2_3 | PHI2 | PHI3 | progeny | P1-C2 | | G | C | H | [illegible] | [illegible] | [illegible] | [illegible] | [illegible] | [illegible] | [illegible] | [illegible] | [illegible] | [illegible] | [illegible] | [illegible] | [illegible] | [illegible] | [illegible] | [illegible] | [illegible] | [illegible] | [illegible] | [illegible] | [illegible] | [illegible] | [illegible] | [illegible] | [illegible] | [illegible] | [illegible] | [illegible] | [illegible] | [illegible] | [illegible] | [illegible] | [illegible] | [illegible] | [illegible] | [illegible] | [illegible] | [illegible] | [illegible] | [illegible] | [illegible] | [illegible] | [illegible] | [illegible] | [illegible] | [illegible] | [illegible] | [illegible] | [illegible] | [illegible] | [illegible] | [illegible] | [illegible] | [illegible] | [illegible] | [illegible] | [illegible] | [illegible] | [illegible] | [illegible] | [illegible] | [illegible] | [illegible] |
| 2016H_P1-C6_1 | PHI2 | PHI3 | progeny | P1-C6 | | G | C | H | [illegible] | [illegible] | [illegible] | [illegible] | [illegible] | [illegible] | [illegible] | [illegible] | [illegible] | [illegible] | [illegible] | [illegible] | [illegible] | [illegible] | [illegible] | [illegible] | [illegible] | [illegible] | [illegible] | [illegible] | [illegible] | [illegible] | [illegible] | [illegible] | [illegible] | [illegible] | [illegible] | [illegible] | [illegible] | [illegible] | [illegible] | [illegible] | [illegible] | [illegible] | [illegible] | [illegible] | [illegible] | [illegible] | [illegible] | [illegible] | [illegible] | [illegible] | [illegible] | [illegible] | [illegible] | [illegible] | [illegible] | [illegible] | [illegible] | [illegible] | [illegible] | [illegible] | [illegible] | [illegible] | [illegible] | [illegible] | [illegible] | [illegible] | [illegible] | [illegible] | [illegible] | [illegible] | [illegible] |
| 2016H_P1-A4_1 | PHI2 | PHI3 | progeny | P1-A4 | | G | C | H | [illegible] | [illegible] | [illegible] | [illegible] | [illegible] | [illegible] | [illegible] | [illegible] | [illegible] | [illegible] | [illegible] | [illegible] | [illegible] | [illegible] | [illegible] | [illegible] | [illegible] | [illegible] | [illegible] | [illegible] | [illegible] | [illegible] | [illegible] | [illegible] | [illegible] | [illegible] | [illegible] | [illegible] | [illegible] | [illegible] | [illegible] | [illegible] | [illegible] | [illegible] | [illegible] | [illegible] | [illegible] | [illegible] | [illegible] | [illegible] | [illegible] | [illegible] | [illegible] | [illegible] | [illegible] | [illegible] | [illegible] | [illegible] | [illegible] | [illegible] | [illegible] | [illegible] | [illegible] | [illegible] | [illegible] | [illegible] | [illegible] | [illegible] | [illegible] | [illegible] | [illegible] | [illegible] | [illegible] |
| 2016H_P1-A4_2 | PHI2 | PHI3 | progeny | P1-A4 | | G | C | H | [illegible] | [illegible] | [illegible] | [illegible] | [illegible] | [illegible] | [illegible] | [illegible] | [illegible] | [illegible] | [illegible] | [illegible] | [illegible] | [illegible] | [illegible] | [illegible] | [illegible] | [illegible] | [illegible] | [illegible] | [illegible] | [illegible] | [illegible] | [illegible] | [illegible] | [illegible] | [illegible] | [illegible] | [illegible] | [illegible] | [illegible] | [illegible] | [illegible] | [illegible] | [illegible] | [illegible] | [illegible] | [illegible] | [illegible] | [illegible] | [illegible] | [illegible] | [illegible] | [illegible] | [illegible] | [illegible] | [illegible] | [illegible] | [illegible] | [illegible] | [illegible] | [illegible] | [illegible] | [illegible] | [illegible] | [illegible] | [illegible] | [illegible] | [illegible] | [illegible] | [illegible] | [illegible] | [illegible] |
| 2016H_P1-A4_4 | PHI2 | PHI3 | progeny | P1-A4 | | G | C | H | [illegible] | [illegible] | [illegible] | [illegible] | [illegible] | [illegible] | [illegible] | [illegible] | [illegible] | [illegible] | [illegible] | [illegible] | [illegible] | [illegible] | [illegible] | [illegible] | [illegible] | [illegible] | [illegible] | [illegible] | [illegible] | [illegible] | [illegible] | [illegible] | [illegible] | [illegible] | [illegible] | [illegible] | [illegible] | [illegible] | [illegible] | [illegible] | [illegible] | [illegible] | [illegible] | [illegible] | [illegible] | [illegible] | [illegible] | [illegible] | [illegible] | [illegible] | [illegible] | [illegible] | [illegible] | [illegible] | [illegible] | [illegible] | [illegible] | [illegible] | [illegible] | [illegible] | [illegible] | [illegible] | [illegible] | [illegible] | [illegible] | [illegible] | [illegible] | [illegible] | [illegible] | [illegible] | [illegible] |
| 2016H_P1-A4_5 | PHI2 | PHI3 | progeny | P1-A4 | | G | C | H | [illegible] | [illegible] | [illegible] | [illegible] | [illegible] | [illegible] | [illegible] | [illegible] | [illegible] | [illegible] | [illegible] | [illegible] | [illegible] | [illegible] | [illegible] | [illegible] | [illegible] | [illegible] | [illegible] | [illegible] | [illegible] | [illegible] | [illegible] | [illegible] | [illegible] | [illegible] | [illegible] | [illegible] | [illegible] | [illegible] | [illegible] | [illegible] | [illegible] | [illegible] | [illegible] | [illegible] | [illegible] | [illegible] | [illegible] | [illegible] | [illegible] | [illegible] | [illegible] | [illegible] | [illegible] | [illegible] | [illegible] | [illegible] | [illegible] | [illegible] | [illegible] | [illegible] | [illegible] | [illegible] | [illegible] | [illegible] | [illegible] | [illegible] | [illegible] | [illegible] | [illegible] | [illegible] | [illegible] |
| 2016H_P1-A4_9 | PHI2 | PHI3 | progeny | P1-A4 | | G | C | H | [illegible] | [illegible] | [illegible] | [illegible] | [illegible] | [illegible] | [illegible] | [illegible] | [illegible] | [illegible] | [illegible] | [illegible] | [illegible] | [illegible] | [illegible] | [illegible] | [illegible] | [illegible] | [illegible] | [illegible] | [illegible] | [illegible] | [illegible] | [illegible] | [illegible] | [illegible] | [illegible] | [illegible] | [illegible] | [illegible] | [illegible] | [illegible] | [illegible] | [illegible] | [illegible] | [illegible] | [illegible] | [illegible] | [illegible] | [illegible] | [illegible] | [illegible] | [illegible] | [illegible] | [illegible] | [illegible] | [illegible] | [illegible] | [illegible] | [illegible] | [illegible] | [illegible] | [illegible] | [illegible] | [illegible] | [illegible] | [illegible] | [illegible] | [illegible] | [illegible] | [illegible] | [illegible] | [illegible] |
| 2016H_P1-A4_10 | PHI2 | PHI3 | progeny | P1-A4 | | G | C | H | [illegible] | [illegible] | [illegible] | [illegible] | [illegible] | [illegible] | [illegible] | [illegible] | [illegible] | [illegible] | [illegible] | [illegible] | [illegible] | [illegible] | [illegible] | [illegible] | [illegible] | [illegible] | [illegible] | [illegible] | [illegible] | [illegible] | [illegible] | [illegible] | [illegible] | [illegible] | [illegible] | [illegible] | [illegible] | [illegible] | [illegible] | [illegible] | [illegible] | [illegible] | [illegible] | [illegible] | [illegible] | [illegible] | [illegible] | [illegible] | [illegible] | [illegible] | [illegible] | [illegible] | [illegible] | [illegible] | [illegible] | [illegible] | [illegible] | [illegible] | [illegible] | [illegible] | [illegible] | [illegible] | [illegible] | [illegible] | [illegible] | [illegible] | [illegible] | [illegible] | [illegible] | [illegible] | [illegible] |
| 2016H_P1-A9_1 | PHI2 | PHI3 | progeny | P1-A9 | | G | C | H | [illegible] | [illegible] | [illegible] | [illegible] | [illegible] | [illegible] | [illegible] | [illegible] | [illegible] | [illegible] | [illegible] | [illegible] | [illegible] | [illegible] | [illegible] | [illegible] | [illegible] | [illegible] | [illegible] | [illegible] | [illegible] | [illegible] | [illegible] | [illegible] | [illegible] | [illegible] | [illegible] | [illegible] | [illegible] | [illegible] | [illegible] | [illegible] | [illegible] | [illegible] | [illegible] | [illegible] | [illegible] | [illegible] | [illegible] | [illegible] | [illegible] | [illegible] | [illegible] | [illegible] | [illegible] | [illegible] | [illegible] | [illegible] | [illegible] | [illegible] | [illegible] | [illegible] | [illegible] | [illegible] | [illegible] | [illegible] | [illegible] | [illegible] | [illegible] | [illegible] | [illegible] | [illegible] | [illegible] |
| 2016H_P1-A9_3 | PHI2 | PHI3 | progeny | P1-A9 | | G | C | C | [illegible] | [illegible] | [illegible] | [illegible] | [illegible] | [illegible] | [illegible] | [illegible] | [illegible] | [illegible] | [illegible] | [illegible] | [illegible] | [illegible] | [illegible] | [illegible] | [illegible] | [illegible] | [illegible] | [illegible] | [illegible] | [illegible] | [illegible] | [illegible] | [illegible] | [illegible] | [illegible] | [illegible] | [illegible] | [illegible] | [illegible] | [illegible] | [illegible] | [illegible] | [illegible] | [illegible] | [illegible] | [illegible] | [illegible] | [illegible] | [illegible] | [illegible] | [illegible] | [illegible] | [illegible] | [illegible] | [illegible] | [illegible] | [illegible] | [illegible] | [illegible] | [illegible] | [illegible] | [illegible] | [illegible] | [illegible] | [illegible] | [illegible] | [illegible] | [illegible] | [illegible] | [illegible] | [illegible] |
| 2016H_P1-A11_1 | PHI2 | PHI3 | progeny | P1-A11 | | G | C | H | [illegible] | [illegible] | [illegible] | [illegible] | [illegible] | [illegible] | [illegible] | [illegible] | [illegible] | [illegible] | [illegible] | [illegible] | [illegible] | [illegible] | [illegible] | [illegible] | [illegible] | [illegible] | [illegible] | [illegible] | [illegible] | [illegible] | [illegible] | [illegible] | [illegible] | [illegible] | [illegible] | [illegible] | [illegible] | [illegible] | [illegible] | [illegible] | [illegible] | [illegible] | [illegible] | [illegible] | [illegible] | [illegible] | [illegible] | [illegible] | [illegible] | [illegible] | [illegible] | [illegible] | [illegible] | [illegible] | [illegible] | [illegible] | [illegible] | [illegible] | [illegible] | [illegible] | [illegible] | [illegible] | [illegible] | [illegible] | [illegible] | [illegible] | [illegible] | [illegible] | [illegible] | [illegible] | [illegible] |
| 2016H_P1-A11_3 | PHI2 | PHI3 | progeny | P1-A11 | | G | C | H | [illegible] | [illegible] | [illegible] | [illegible] | [illegible] | [illegible] | [illegible] | [illegible] | [illegible] | [illegible] | [illegible] | [illegible] | [illegible] | [illegible] | [illegible] | [illegible] | [illegible] | [illegible] | [illegible] | [illegible] | [illegible] | [illegible] | [illegible] | [illegible] | [illegible] | [illegible] | [illegible] | [illegible] | [illegible] | [illegible] | [illegible] | [illegible] | [illegible] | [illegible] | [illegible] | [illegible] | [illegible] | [illegible] | [illegible] | [illegible] | [illegible] | [illegible] | [illegible] | [illegible] | [illegible] | [illegible] | [illegible] | [illegible] | [illegible] | [illegible] | [illegible] | [illegible] | [illegible] | [illegible] | [illegible] | [illegible] | [illegible] | [illegible] | [illegible] | [illegible] | [illegible] | [illegible] | [illegible] |
| 2016H_P1-A11_4 | PHI2 | PHI3 | progeny | P1-A11 | | G | C | H | [illegible] | [illegible] | [illegible] | [illegible] | [illegible] | [illegible] | [illegible] | [illegible] | [illegible] | [illegible] | [illegible] | [illegible] | [illegible] | [illegible] | [illegible] | [illegible] | [illegible] | [illegible] | [illegible] | [illegible] | [illegible] | [illegible] | [illegible] | [illegible] | [illegible] | [illegible] | [illegible] | [illegible] | [illegible] | [illegible] | [illegible] | [illegible] | [illegible] | [illegible] | [illegible] | [illegible] | [illegible] | [illegible] | [illegible] | [illegible] | [illegible] | [illegible] | [illegible] | [illegible] | [illegible] | [illegible] | [illegible] | [illegible] | [illegible] | [illegible] | [illegible] | [illegible] | [illegible] | [illegible] | [illegible] | [illegible] | [illegible] | [illegible] | [illegible] | [illegible] | [illegible] | [illegible] | [illegible] |
| 2016H_P1-A11_5 | PHI2 | PHI3 | progeny | P1-A11 | | G | C | H | [illegible] | [illegible] | [illegible] | [illegible] | [illegible] | [illegible] | [illegible] | [illegible] | [illegible] | [illegible] | [illegible] | [illegible] | [illegible] | [illegible] | [illegible] | [illegible] | [illegible] | [illegible] | [illegible] | [illegible] | [illegible] | [illegible] | [illegible] | [illegible] | [illegible] | [illegible] | [illegible] | [illegible] | [illegible] | [illegible] | [illegible] | [illegible] | [illegible] | [illegible] | [illegible] | [illegible] | [illegible] | [illegible] | [illegible] | [illegible] | [illegible] | [illegible] | [illegible] | [illegible] | [illegible] | [illegible] | [illegible] | [illegible] | [illegible] | [illegible] | [illegible] | [illegible] | [illegible] | [illegible] | [illegible] | [illegible] | [illegible] | [illegible] | [illegible] | [illegible] | [illegible] | [illegible] | [illegible] |

plant_id – unique identifier of the plant
mom_gen – genetic backgroiund of mother in the hybred
dad_gen – genetic background of father in the hybred
family_ – identify mother and all the progeny plants from the s
mother_progeny – identify each plant as mother of the family or proger
Chr – number of the chromosome, where marker is locate
Distance – position of the marker on the corresponding chromo
H – heterozygous marker
A, T, G and C – marker homozygous for the corresponding nucleoti
EQV, LS, -9 , etc. – missing values

**Table S9: Maintenance of maternal heterozygosity in putative clonal progeny**

| # | Plant ID | Genetic background of hybred | Maintenance of maternal heterozygosity , % |
|---|---|---|---|
| 1 | 2013_P1-8-4 | PHI1 X PHI2 | 99 |
| 2 | 2013_P1-25-1 | PHI1 X PHI2 | 95 |
| 3 | 2013_P1-1-1 | PHI1 X PHI2 | 93 |
| 4 | 2013_P1-19-3 | PHI1 X PHI2 | 90 |
| 5 | 2013_P1-8-1 | PHI1 X PHI2 | 87 |
| 6 | 2013_P1-8-5 | PHI1 X PHI2 | 87 |
| 7 | 2013_P1-6-1 | PHI1 X PHI2 | 85 |
| 8 | 2013_P1-19-2 | PHI1 X PHI2 | 82 |
| 9 | 2013_P1-22-1 | PHI1 X PHI2 | 79 |
| 10 | 2013_P1-25-4 | PHI1 X PHI2 | 79 |
| 11 | 2013_P1-31-1 | PHI1 X PHI2 | 77 |
| 12 | 2013_P1-8-3 | PHI1 X PHI2 | 77 |
| 13 | 2013_P1-19-4 | PHI1 X PHI2 | 76 |
| 14 | 2013_P1-9-3 | PHI1 X PHI2 | 74 |
| 15 | 2013_P1-10-1 | PHI1 X PHI2 | 73 |
| 16 | 2013_P1-25-6 | PHI1 X PHI2 | 72 |
| 17 | 2013_P1-25-3 | PHI1 X PHI2 | 70 |
| 18 | 2013_P1-25-5 | PHI1 X PHI2 | 70 |
| 19 | 2013_P1-7-1 | PHI1 X PHI2 | 66 |
| 20 | 2013_P1-8-2 | PHI1 X PHI2 | 61 |
| 21 | 2013_P1-17-1 | PHI1 X PHI2 | 58 |
| 22 | 2013_P1-25-2 | PHI1 X PHI2 | 54 |
| 23 | 2013_P1-17-2 | PHI1 X PHI2 | 54 |
| 24 | 2013_P1-9-1 | PHI1 X PHI2 | 52 |
| 25 | 2013_P1-16-2 | PHI1 X PHI2 | 52 |
| 26 | 2013_P1-31-2 | PHI1 X PHI2 | 51 |
| 27 | 2013_P1-27-1 | PHI1 X PHI2 | 51 |
| 28 | 2013_P1-9-2 | PHI1 X PHI2 | 50 |
| 29 | 2013_P1-19-5 | PHI1 X PHI2 | 50 |
| 30 | 2013_P1-21-3 | PHI1 X PHI2 | 50 |
| 31 | 2013_P1-21-1 | PHI1 X PHI2 | 50 |
| 32 | 2013_P1-20-1 | PHI1 X PHI2 | 50 |
| 33 | 2013_P1-21-2 | PHI1 X PHI2 | 48 |
| 34 | 2013_P1-26-1 | PHI1 X PHI2 | 46 |
| 35 | 2013_P1-28-1 | PHI1 X PHI2 | 45 |
| 36 | 2013_P1-19-1 | PHI1 X PHI2 | 43 |
| 37 | 2013_P1-21-4 | PHI1 X PHI2 | 42 |
| 38 | 2013_P1-16-1 | PHI1 X PHI2 | 42 |
| 39 | 2013_P1-20-2 | PHI1 X PHI2 | 41 |
| 40 | 2013_P1-22-2 | PHI1 X PHI2 | 36 |
| 41 | 2015_P3-B5-1 | PHI1 X PHI3 | 100 |
| 42 | 2015_P3-F3-1 | PHI1 X PHI3 | 100 |
| 43 | 2015_P3-F2-1 | PHI1 X PHI3 | 100 |
| 44 | 2015_P3-F4-1 | PHI1 X PHI3 | 100 |
| 45 | 2015_P3-F4-3 | PHI1 X PHI3 | 98 |
| 46 | 2015_P3-C11-3 | PHI1 X PHI3 | 98 |
| 47 | 2015_P3-A7-1 | PHI1 X PHI3 | 96 |
| 48 | 2015_P3-A8-2 | PHI1 X PHI3 | 94 |
| 49 | 2015_P3-A6-1 | PHI1 X PHI3 | 92 |
| 50 | 2015_P3-B2-1 | PHI1 X PHI3 | 90 |
| 51 | 2015_P3-E4-1 | PHI1 X PHI3 | 89 |
| 52 | 2015_P3-F3-2 | PHI1 X PHI3 | 89 |
| 53 | 2015_P3-C10-2 | PHI1 X PHI3 | 85 |
| 54 | 2016A_P1-A3_1 | PHI1 X PHI3 | 84 |
| 55 | 2015_P3-D5-1 | PHI1 X PHI3 | 83 |
| 56 | 2015_P3-A10-1 | PHI1 X PHI3 | 83 |
| 57 | 2016A_P1-D5_1 | PHI1 X PHI3 | 81 |
| 58 | 2015_P3-E10-1 | PHI1 X PHI3 | 80 |
| 59 | 2015_P3-A8-1 | PHI1 X PHI3 | 74 |
| 60 | 2015_P3-C8-1 | PHI1 X PHI3 | 73 |
| 61 | 2015_P3-A1-1 | PHI1 X PHI3 | 68 |
| 62 | 2015_P3-C2-1 | PHI1 X PHI3 | 66 |
| 63 | 2015_P3-F4-4 | PHI1 X PHI3 | 63 |
| 64 | 2016A_P1-E3_5 | PHI1 X PHI3 | 63 |
| 65 | 2015_P3-E8-2 | PHI1 X PHI3 | 58 |

**Table S9: Maintenance of maternal heterozygosity in putative clonal progeny**

| # | Plant ID | Genetic background of hybred | Maintenance of maternal heterozygosity , % |
|---|---|---|---|
| 66 | 2015_P3-C1-1 | PHI1 X PHI3 | 58 |
| 67 | 2015_P3-D3-1 | PHI1 X PHI3 | 56 |
| 68 | 2016A_P1-F7_4 | PHI1 X PHI3 | 56 |
| 69 | 2015_P3-D6-1 | PHI1 X PHI3 | 54 |
| 70 | 2015_P3-D7-2 | PHI1 X PHI3 | 54 |
| 71 | 2016A_P1-E3_3 | PHI1 X PHI3 | 54 |
| 72 | 2015_P3-A2-2 | PHI1 X PHI3 | 54 |
| 73 | 2015_P3-E8-3 | PHI1 X PHI3 | 52 |
| 74 | 2015_P3-D10-1 | PHI1 X PHI3 | 51 |
| 75 | 2015_P3-B11-1 | PHI1 X PHI3 | 51 |
| 76 | 2015_P3-E9-1 | PHI1 X PHI3 | 51 |
| 77 | 2015_P3-A2-3 | PHI1 X PHI3 | 51 |
| 78 | 2015_P3-E9-3 | PHI1 X PHI3 | 50 |
| 79 | 2015_P3-E8-1 | PHI1 X PHI3 | 50 |
| 80 | 2015_P3-A3-1 | PHI1 X PHI3 | 50 |
| 81 | 2015_P3-B3-1 | PHI1 X PHI3 | 49 |
| 82 | 2015_P3-C7-1 | PHI1 X PHI3 | 48 |
| 83 | 2015_P3-A2-1 | PHI1 X PHI3 | 48 |
| 84 | 2015_P3-F4-2 | PHI1 X PHI3 | 47 |
| 85 | 2015_P3-E5-1 | PHI1 X PHI3 | 47 |
| 86 | 2015_P3-B4-1 | PHI1 X PHI3 | 46 |
| 87 | 2015_P3-D4-1 | PHI1 X PHI3 | 45 |
| 88 | 2015_P3-E7-1 | PHI1 X PHI3 | 41 |
| 89 | 2015_P3-E11-1 | PHI1 X PHI3 | 41 |
| 90 | 2015_P3-E5-2 | PHI1 X PHI3 | 40 |
| 91 | 2015_P3-B2-2 | PHI1 X PHI3 | 40 |
| 92 | 2016A_P1-E9_1 | PHI1 X PHI3 | 40 |
| 93 | 2015_P3-D4-3 | PHI1 X PHI3 | 39 |
| 94 | 2015_P3-A2-4 | PHI1 X PHI3 | 39 |
| 95 | 2015_P3-C8-3 | PHI1 X PHI3 | 38 |
| 96 | 2015_P3-A8-3 | PHI1 X PHI3 | 38 |
| 97 | 2015_P3-E9-4 | PHI1 X PHI3 | 38 |
| 98 | 2015_P3-E12-1 | PHI1 X PHI3 | 38 |
| 99 | 2015_P3-D7-1 | PHI1 X PHI3 | 36 |
| 100 | 2015_P3-B7-1 | PHI1 X PHI3 | 36 |
| 101 | 2015_P3-F1-1 | PHI1 X PHI3 | 35 |
| 102 | 2016A_P1-D8_3 | PHI1 X PHI3 | 34 |
| 103 | 2015_P3-E4-2 | PHI1 X PHI3 | 34 |
| 104 | 2015_P3-D4-2 | PHI1 X PHI3 | 32 |
| 105 | 2015_P3-D3-2 | PHI1 X PHI3 | 30 |
| 106 | 2015_P3-E9-2 | PHI1 X PHI3 | 29 |
| 107 | 2015_P3-C8-2 | PHI1 X PHI3 | 25 |
| 108 | 2015_P3-E5-3 | PHI1 X PHI3 | 25 |
| 109 | 2015_P3-A11-3 | PHI1 X PHI3 | 22 |
| 110 | 2015_P3-A11-2 | PHI1 X PHI3 | 20 |
| 111 | 2015_P3-A11-1 | PHI1 X PHI3 | 17 |
| 115 | 2015_P4-A9-3 | PHI2 X PHI1 | 100 |
| 116 | 2015_P3-H3-1 | PHI2 X PHI1 | 100 |
| 117 | 2015_P3-G1-4 | PHI2 X PHI1 | 100 |
| 118 | 2016A_P2-H6_16 | PHI2 X PHI1 | 100 |
| 119 | 2015_P3-H5-4 | PHI2 X PHI1 | 100 |
| 120 | 2016A_P1-H8_4 | PHI2 X PHI1 | 100 |
| 121 | 2016A_P1-H8_7 | PHI2 X PHI1 | 100 |
| 122 | 2016A_P2-H6_9 | PHI2 X PHI1 | 99 |
| 123 | 2015_P4-A5-1 | PHI2 X PHI1 | 99 |
| 124 | 2015_P4-A3-1 | PHI2 X PHI1 | 97 |
| 125 | 2015_P3-F9-5 | PHI2 X PHI1 | 97 |
| 126 | 2015_P3-G9-6 | PHI2 X PHI1 | 96 |
| 127 | 2016A_P2-D1_7 | PHI2 X PHI1 | 96 |
| 128 | 2015_P4-B6-2 | PHI2 X PHI1 | 95 |
| 129 | 2016A_P2-D10_3 | PHI2 X PHI1 | 94 |
| 130 | 2015_P3-H10-4 | PHI2 X PHI1 | 94 |
| 131 | 2015_P3-G9-11 | PHI2 X PHI1 | 94 |
| 132 | 2015_P4-A1-1 | PHI2 X PHI1 | 93 |
| 133 | 2015_P3-F7-5 | PHI2 X PHI1 | 93 |

**Table S9: Maintenance of maternal heterozygosity in putative clonal progeny**

| # | Plant ID | Genetic background of hybred | Maintenance of maternal heterozygosity , % |
|---|---|---|---|
| 134 | 2015_P3-H7-1 | PHI2 X PHI1 | 93 |
| 135 | 2015_P3-F9-2 | PHI2 X PHI1 | 92 |
| 136 | 2015_P3-F7-1 | PHI2 X PHI1 | 90 |
| 137 | 2016A_P1-H8_5 | PHI2 X PHI1 | 90 |
| 138 | 2015_P4-A2-1 | PHI2 X PHI1 | 89 |
| 139 | 2015_P3-F9-4 | PHI2 X PHI1 | 89 |
| 140 | 2015_P3-G1-3 | PHI2 X PHI1 | 89 |
| 141 | 2016A_P2-D5_4 | PHI2 X PHI1 | 87 |
| 142 | 2015_P3-H11-1 | PHI2 X PHI1 | 87 |
| 143 | 2015_P3-G4-2 | PHI2 X PHI1 | 87 |
| 144 | 2015_P3-H6-1 | PHI2 X PHI1 | 87 |
| 145 | 2015_P3-G8-1 | PHI2 X PHI1 | 87 |
| 146 | 2015_P3-G1-1 | PHI2 X PHI1 | 87 |
| 147 | 2015_P3-G3-1 | PHI2 X PHI1 | 86 |
| 148 | 2016A_P2-F7_1 | PHI2 X PHI1 | 86 |
| 149 | 2016A_P2-B7_1 | PHI2 X PHI1 | 86 |
| 150 | 2015_P3-G9-10 | PHI2 X PHI1 | 85 |
| 151 | 2016A_P2-G7_6 | PHI2 X PHI1 | 84 |
| 152 | 2015_P3-F6-2 | PHI2 X PHI1 | 84 |
| 153 | 2016A_P2-G7_8 | PHI2 X PHI1 | 84 |
| 154 | 2015_P3-F6-3 | PHI2 X PHI1 | 83 |
| 155 | 2016A_P1-G12_2 | PHI2 X PHI1 | 83 |
| 156 | 2015_P3-G9-7 | PHI2 X PHI1 | 83 |
| 157 | 2015_P3-F12-3 | PHI2 X PHI1 | 82 |
| 158 | 2016A_P2-A11_5 | PHI2 X PHI1 | 82 |
| 159 | 2015_P3-F9-3 | PHI2 X PHI1 | 82 |
| 160 | 2015_P4-A5-2 | PHI2 X PHI1 | 82 |
| 161 | 2015_P3-G8-2 | PHI2 X PHI1 | 82 |
| 162 | 2015_P3-H10-6 | PHI2 X PHI1 | 82 |
| 163 | 2015_P4-A1-2 | PHI2 X PHI1 | 81 |
| 164 | 2015_P4-A4-1 | PHI2 X PHI1 | 81 |
| 165 | 2015_P3-H5-1 | PHI2 X PHI1 | 80 |
| 166 | 2016A_P2-A12_8 | PHI2 X PHI1 | 80 |
| 167 | 2016A_P1-H8_1 | PHI2 X PHI1 | 79 |
| 168 | 2016A_P2-G7_13 | PHI2 X PHI1 | 79 |
| 169 | 2015_P3-F8-1 | PHI2 X PHI1 | 78 |
| 170 | 2015_P3-G9-2 | PHI2 X PHI1 | 78 |
| 171 | 2015_P4-A9-4 | PHI2 X PHI1 | 78 |
| 172 | 2015_P3-G4-5 | PHI2 X PHI1 | 77 |
| 173 | 2015_P3-H5-2 | PHI2 X PHI1 | 77 |
| 174 | 2015_P4-B3-2 | PHI2 X PHI1 | 77 |
| 175 | 2015_P3-F12-2 | PHI2 X PHI1 | 76 |
| 176 | 2015_P3-F11-1 | PHI2 X PHI1 | 76 |
| 177 | 2015_P3-H5-3 | PHI2 X PHI1 | 76 |
| 178 | 2015_P4-A7-1 | PHI2 X PHI1 | 76 |
| 179 | 2015_P4-A9-2 | PHI2 X PHI1 | 76 |
| 180 | 2015_P3-G6-1 | PHI2 X PHI1 | 76 |
| 181 | 2016A_P2-D1_8 | PHI2 X PHI1 | 75 |
| 182 | 2015_P4-A9-1 | PHI2 X PHI1 | 75 |
| 183 | 2016A_P2-A12_12 | PHI2 X PHI1 | 75 |
| 184 | 2016A_P2-G9_5 | PHI2 X PHI1 | 75 |
| 185 | 2015_P3-G10-9 | PHI2 X PHI1 | 74 |
| 186 | 2015_P3-H10-1 | PHI2 X PHI1 | 74 |
| 187 | 2015_P3-G4-4 | PHI2 X PHI1 | 74 |
| 188 | 2015_P3-G9-8 | PHI2 X PHI1 | 74 |
| 189 | 2015_P3-H10-5 | PHI2 X PHI1 | 73 |
| 190 | 2016A_P2-A11_4 | PHI2 X PHI1 | 73 |
| 191 | 2016A_P2-C11_1 | PHI2 X PHI1 | 73 |
| 192 | 2016A_P2-D1_4 | PHI2 X PHI1 | 73 |
| 193 | 2016A_P1-G12_6 | PHI2 X PHI1 | 72 |
| 194 | 2016A_P1-G12_1 | PHI2 X PHI1 | 72 |
| 195 | 2015_P3-G9-5 | PHI2 X PHI1 | 72 |
| 196 | 2015_P3-G10-8 | PHI2 X PHI1 | 72 |
| 197 | 2015_P3-G9-3 | PHI2 X PHI1 | 71 |
| 198 | 2015_P4-A4-4 | PHI2 X PHI1 | 71 |

**Table S9: Maintenance of maternal heterozygosity in putative clonal progeny**

| # | Plant ID | Genetic background of hybred | Maintenance of maternal heterozygosity , % |
|---|---|---|---|
| 199 | 2016A_P2-F2_2 | PHI2 X PHI1 | 70 |
| 200 | 2015_P4-B6-1 | PHI2 X PHI1 | 70 |
| 201 | 2015_P3-H8-1 | PHI2 X PHI1 | 69 |
| 202 | 2016A_P2-D1_3 | PHI2 X PHI1 | 68 |
| 203 | 2016A_P2-H6_17 | PHI2 X PHI1 | 68 |
| 204 | 2016A_P2-G9_6 | PHI2 X PHI1 | 68 |
| 205 | 2016A_P2-G1_6 | PHI2 X PHI1 | 67 |
| 206 | 2016A_P1-H11_3 | PHI2 X PHI1 | 67 |
| 207 | 2015_P3-H3-2 | PHI2 X PHI1 | 66 |
| 208 | 2015_P3-F9-1 | PHI2 X PHI1 | 65 |
| 209 | 2015_P3-G12-3 | PHI2 X PHI1 | 65 |
| 210 | 2015_P3-G4-3 | PHI2 X PHI1 | 65 |
| 211 | 2015_P4-B3-3 | PHI2 X PHI1 | 65 |
| 212 | 2015_P3-F6-1 | PHI2 X PHI1 | 64 |
| 213 | 2015_P4-A6-3 | PHI2 X PHI1 | 62 |
| 214 | 2015_P3-G10-2 | PHI2 X PHI1 | 62 |
| 215 | 2015_P4-A6-4 | PHI2 X PHI1 | 61 |
| 216 | 2015_P3-F12-1 | PHI2 X PHI1 | 61 |
| 217 | 2016A_P2-A12_7 | PHI2 X PHI1 | 61 |
| 218 | 2015_P3-G12-1 | PHI2 X PHI1 | 60 |
| 219 | 2015_P4-B3-1 | PHI2 X PHI1 | 60 |
| 220 | 2016A_P2-G7_15 | PHI2 X PHI1 | 60 |
| 221 | 2015_P4-A4-3 | PHI2 X PHI1 | 60 |
| 222 | 2015_P3-F5-1 | PHI2 X PHI1 | 58 |
| 223 | 2015_P3-G12-2 | PHI2 X PHI1 | 58 |
| 224 | 2016A_P2-A7_9 | PHI2 X PHI1 | 57 |
| 225 | 2016A_P2-A12_4 | PHI2 X PHI1 | 57 |
| 226 | 2016A_P1-H11_4 | PHI2 X PHI1 | 57 |
| 227 | 2015_P3-G10-1 | PHI2 X PHI1 | 56 |
| 228 | 2015_P4-B4-2 | PHI2 X PHI1 | 56 |
| 229 | 2016A_P2-A12_9 | PHI2 X PHI1 | 55 |
| 230 | 2015_P3-H9-1 | PHI2 X PHI1 | 55 |
| 231 | 2015_P3-H10-7 | PHI2 X PHI1 | 54 |
| 232 | 2015_P3-G10-3 | PHI2 X PHI1 | 54 |
| 233 | 2016A_P1-H8_16 | PHI2 X PHI1 | 53 |
| 234 | 2015_P3-G10-10 | PHI2 X PHI1 | 53 |
| 235 | 2015_P3-G10-4 | PHI2 X PHI1 | 53 |
| 236 | 2015_P3-H9-3 | PHI2 X PHI1 | 53 |
| 237 | 2016A_P2-D7_3 | PHI2 X PHI1 | 53 |
| 238 | 2015_P3-G2-1 | PHI2 X PHI1 | 53 |
| 239 | 2015_P3-H9-2 | PHI2 X PHI1 | 53 |
| 240 | 2016A_P2-C11_4 | PHI2 X PHI1 | 52 |
| 241 | 2015_P4-A4-2 | PHI2 X PHI1 | 52 |
| 242 | 2015_P3-G10-7 | PHI2 X PHI1 | 52 |
| 243 | 2015_P3-G9-9 | PHI2 X PHI1 | 51 |
| 244 | 2015_P3-G12-4 | PHI2 X PHI1 | 51 |
| 245 | 2016A_P1-F8_4 | PHI2 X PHI1 | 51 |
| 246 | 2015_P3-G9-12 | PHI2 X PHI1 | 50 |
| 247 | 2015_P4-A6-1 | PHI2 X PHI1 | 50 |
| 248 | 2015_P4-A5-3 | PHI2 X PHI1 | 50 |
| 249 | 2016A_P2-H6_1 | PHI2 X PHI1 | 50 |
| 250 | 2015_P3-H9-6 | PHI2 X PHI1 | 50 |
| 251 | 2015_P3-G9-1 | PHI2 X PHI1 | 49 |
| 252 | 2016A_P2-B4_5 | PHI2 X PHI1 | 48 |
| 253 | 2015_P4-B4-1 | PHI2 X PHI1 | 48 |
| 254 | 2015_P3-F12-4 | PHI2 X PHI1 | 48 |
| 255 | 2015_P3-G10-6 | PHI2 X PHI1 | 48 |
| 256 | 2015_P3-H10-3 | PHI2 X PHI1 | 48 |
| 257 | 2015_P3-G4-1 | PHI2 X PHI1 | 47 |
| 258 | 2016A_P2-B10_6 | PHI2 X PHI1 | 47 |
| 259 | 2016A_P2-D1_12 | PHI2 X PHI1 | 46 |
| 260 | 2015_P4-A6-2 | PHI2 X PHI1 | 46 |
| 261 | 2015_P3-G1-2 | PHI2 X PHI1 | 46 |
| 262 | 2016A_P2-D1_10 | PHI2 X PHI1 | 45 |
| 263 | 2015_P3-H9-4 | PHI2 X PHI1 | 44 |

**Table S9: Maintenance of maternal heterozygosity in putative clonal progeny**

| # | Plant ID | Genetic background of hybred | Maintenance of maternal heterozygosity , % |
|---|---|---|---|
| 264 | 2015_P3-F7-2 | PHI2 X PHI1 | 43 |
| 265 | 2015_P3-F7-4 | PHI2 X PHI1 | 43 |
| 266 | 2015_P3-H10-2 | PHI2 X PHI1 | 42 |
| 267 | 2015_P3-F7-3 | PHI2 X PHI1 | 42 |
| 268 | 2015_P3-H9-5 | PHI2 X PHI1 | 42 |
| 269 | 2015_P3-G6-2 | PHI2 X PHI1 | 41 |
| 270 | 2016A_P2-G9_3 | PHI2 X PHI1 | 40 |
| 271 | 2016A_P2-G1_4 | PHI2 X PHI1 | 39 |
| 272 | 2016A_P2-B4_1 | PHI2 X PHI1 | 37 |
| 273 | 2015_P3-G9-4 | PHI2 X PHI1 | 36 |
| 274 | 2016A_P2-B4_6 | PHI2 X PHI1 | 34 |
| 275 | 2015_P3-G10-5 | PHI2 X PHI1 | 32 |
| 276 | 2015_P4-B9-1 | PHI2 X PHI3 | 97 |
| 277 | 2015_P4-B9-2 | PHI2 X PHI3 | 95 |
| 278 | 2015_P4-E5-2 | PHI2 X PHI3 | 91 |
| 279 | 2015_P4-B7-2 | PHI2 X PHI3 | 90 |
| 280 | 2016H_P1-A9_1 | PHI2 X PHI3 | 87 |
| 281 | 2015_P4-C1-2 | PHI2 X PHI3 | 86 |
| 282 | 2015_P4-D4-1 | PHI2 X PHI3 | 85 |
| 283 | 2015_P4-C1-1 | PHI2 X PHI3 | 85 |
| 284 | 2015_P4-C10-1 | PHI2 X PHI3 | 85 |
| 285 | 2016H_P1-A11_1 | PHI2 X PHI3 | 81 |
| 286 | 2016H_P1-A4_2 | PHI2 X PHI3 | 76 |
| 287 | 2015_P4-B11-3 | PHI2 X PHI3 | 76 |
| 288 | 2015_P4-C8-1 | PHI2 X PHI3 | 73 |
| 289 | 2016H_P1-A4_1 | PHI2 X PHI3 | 72 |
| 290 | 2015_P4-D12-2 | PHI2 X PHI3 | 70 |
| 291 | 2015_P4-E1-1 | PHI2 X PHI3 | 69 |
| 292 | 2016H_P1-C6_1 | PHI2 X PHI3 | 68 |
| 293 | 2015_P4-C11-2 | PHI2 X PHI3 | 67 |
| 294 | 2015_P4-D7-1 | PHI2 X PHI3 | 63 |
| 295 | 2015_P4-C11-1 | PHI2 X PHI3 | 63 |
| 296 | 2016H_P1-A4_5 | PHI2 X PHI3 | 58 |
| 297 | 2015_P4-C9-2 | PHI2 X PHI3 | 58 |
| 298 | 2016H_P1-A11_3 | PHI2 X PHI3 | 57 |
| 299 | 2015_P4-B11-1 | PHI2 X PHI3 | 55 |
| 300 | 2015_P4-B7-1 | PHI2 X PHI3 | 54 |
| 301 | 2015_P4-E6-2 | PHI2 X PHI3 | 52 |
| 302 | 2015_P4-E1-2 | PHI2 X PHI3 | 51 |
| 303 | 2016H_P1-A11_5 | PHI2 X PHI3 | 51 |
| 304 | 2016H_P1-C9_2 | PHI2 X PHI3 | 50 |
| 305 | 2016H_P1-A4_10 | PHI2 X PHI3 | 50 |
| 306 | 2015_P4-C8-2 | PHI2 X PHI3 | 47 |
| 307 | 2016H_P1-A4_4 | PHI2 X PHI3 | 46 |
| 308 | 2016H_P1-C9_3 | PHI2 X PHI3 | 46 |
| 309 | 2015_P4-E5-1 | PHI2 X PHI3 | 45 |
| 310 | 2015_P4-C2-1 | PHI2 X PHI3 | 45 |
| 311 | 2016H_P1-A4_9 | PHI2 X PHI3 | 45 |
| 312 | 2016H_P1-A11_4 | PHI2 X PHI3 | 44 |
| 313 | 2016H_P1-C9_1 | PHI2 X PHI3 | 44 |
| 314 | 2016H_P1-C2_3 | PHI2 X PHI3 | 41 |
| 315 | 2015_P4-C9-1 | PHI2 X PHI3 | 40 |
| 316 | 2016H_P1-A9_3 | PHI2 X PHI3 | 37 |
| 317 | 2015_P4-D12-1 | PHI2 X PHI3 | 36 |
| 318 | 2015_P4-D3-1 | PHI2 X PHI3 | 32 |
| 319 | 2015_P4-E5-3 | PHI2 X PHI3 | 31 |
| 320 | 2015_P4-E6-1 | PHI2 X PHI3 | 18 |
| 328 | 2016A_P4-D12_5 | PHI4 X PHI1 | 99 |
| 329 | 2016A_P4-A1_1 | PHI4 X PHI1 | 97 |
| 330 | 2016A_P3-H8_1 | PHI4 X PHI1 | 97 |
| 331 | 2016A_P4-A3_1 | PHI4 X PHI1 | 96 |
| 332 | 2016A_P4-C4_9 | PHI4 X PHI1 | 93 |
| 333 | 2016A_P4-C9_3 | PHI4 X PHI1 | 90 |
| 334 | 2016A_P4-F5_2 | PHI4 X PHI1 | 90 |
| 335 | 2016A_P4-B4_3 | PHI4 X PHI1 | 88 |

**Table S9: Maintenance of maternal heterozygosity in putative clonal progeny**

| # | Plant ID | Genetic background of hybred | Maintenance of maternal heterozygosity , % |
|---|---|---|---|
| 336 | 2016A_P4-F5_6 | PHI4 X PHI1 | 88 |
| 337 | 2016A_P4-A3_5 | PHI4 X PHI1 | 87 |
| 338 | 2016A_P3-H12_1 | PHI4 X PHI1 | 87 |
| 339 | 2016A_P4-D5_1 | PHI4 X PHI1 | 83 |
| 340 | 2016A_P4-F5_1 | PHI4 X PHI1 | 82 |
| 341 | 2016A_P4-C9_1 | PHI4 X PHI1 | 77 |
| 342 | 2016A_P4-C11_3 | PHI4 X PHI1 | 77 |
| 343 | 2016A_P4-C2_5 | PHI4 X PHI1 | 66 |
| 344 | 2016A_P4-F5_5 | PHI4 X PHI1 | 65 |
| 345 | 2016A_P4-C2_3 | PHI4 X PHI1 | 45 |
| 346 | 2016A_P4-C2_2 | PHI4 X PHI1 | 33 |
| 347 | 2016A_P4-C2_6 | PHI4 X PHI1 | 18 |
| 348 | 2016H_P2-B7_1 | PHI4 X PHI3 | 100 |
| 349 | 2016H_P2-G6_2 | PHI4 X PHI3 | 100 |
| 350 | 2016H_P2-C6_3 | PHI4 X PHI3 | 99 |
| 351 | 2016H_P2-B7_4 | PHI4 X PHI3 | 98 |
| 352 | 2016H_P2-G6_3 | PHI4 X PHI3 | 97 |
| 353 | 2016H_P2-D2_2 | PHI4 X PHI3 | 95 |
| 354 | 2016H_P2-C6_1 | PHI4 X PHI3 | 94 |
| 355 | 2016H_P2-B7_3 | PHI4 X PHI3 | 94 |
| 356 | 2016H_P2-C1_4 | PHI4 X PHI3 | 94 |
| 357 | 2016H_P2-G8_2 | PHI4 X PHI3 | 94 |
| 358 | 2016H_P2-G6_4 | PHI4 X PHI3 | 91 |
| 359 | 2016H_P2-G6_1 | PHI4 X PHI3 | 91 |
| 360 | 2016H_P2-C9_1 | PHI4 X PHI3 | 90 |
| 361 | 2016H_P2-C9_4 | PHI4 X PHI3 | 90 |
| 362 | 2016H_P2-C6_2 | PHI4 X PHI3 | 86 |
| 363 | 2016H_P2-A12_2 | PHI4 X PHI3 | 86 |
| 364 | 2016H_P2-C7_1 | PHI4 X PHI3 | 86 |
| 365 | 2016H_P2-E7_2 | PHI4 X PHI3 | 85 |
| 366 | 2016H_P2-F11_1 | PHI4 X PHI3 | 84 |
| 367 | 2016H_P2-C1_6 | PHI4 X PHI3 | 84 |
| 368 | 2016H_P2-D2_1 | PHI4 X PHI3 | 81 |
| 369 | 2016H_P2-F3_3 | PHI4 X PHI3 | 69 |
| 370 | 2016H_P2-A12_1 | PHI4 X PHI3 | 68 |
| 371 | 2016H_P2-G8_3 | PHI4 X PHI3 | 67 |
| 372 | 2016H_P2-E8_6 | PHI4 X PHI3 | 65 |
| 373 | 2016H_P2-E7_1 | PHI4 X PHI3 | 62 |
| 374 | 2016H_P2-C1_5 | PHI4 X PHI3 | 62 |
| 375 | 2016H_P2-E9_3 | PHI4 X PHI3 | 57 |
| 376 | 2016H_P2-E9_1 | PHI4 X PHI3 | 49 |
| 377 | 2016H_P2-F11_2 | PHI4 X PHI3 | 49 |
| 378 | 2016H_P2-F3_2 | PHI4 X PHI3 | 37 |
| 379 | 2016H_P2-B7_2 | PHI4 X PHI3 | 31 |
| 402 | 2016A_P5-G10_10 | PHI6 X PHI1 | 100 |
| 403 | 2016A_P5-G10_11 | PHI6 X PHI1 | 100 |
| 404 | 2016A_P5-H5_7 | PHI6 X PHI1 | 97 |
| 405 | 2016A_P5-H10_2 | PHI6 X PHI1 | 96 |
| 406 | 2016A_P5-G10_9 | PHI6 X PHI1 | 96 |
| 407 | 2016A_P5-G10_7 | PHI6 X PHI1 | 96 |
| 408 | 2016A_P5-F10_7 | PHI6 X PHI1 | 94 |
| 409 | 2016A_P6-C12_2 | PHI6 X PHI1 | 93 |
| 410 | 2016A_P5-G10_6 | PHI6 X PHI1 | 92 |
| 411 | 2016A_P6-B8_2 | PHI6 X PHI1 | 92 |
| 412 | 2016A_P5-H10_5 | PHI6 X PHI1 | 91 |
| 413 | 2016A_P6-C12_1 | PHI6 X PHI1 | 88 |
| 414 | 2016A_P6-A8_11 | PHI6 X PHI1 | 87 |
| 415 | 2016A_P5-H11_5 | PHI6 X PHI1 | 87 |
| 416 | 2016A_P5-H8_10 | PHI6 X PHI1 | 87 |
| 417 | 2016A_P6-B12_4 | PHI6 X PHI1 | 85 |
| 418 | 2016A_P5-H5_10 | PHI6 X PHI1 | 85 |
| 419 | 2016A_P5-H3_9 | PHI6 X PHI1 | 85 |
| 420 | 2016A_P5-H8_4 | PHI6 X PHI1 | 84 |
| 421 | 2016A_P5-H5_8 | PHI6 X PHI1 | 82 |
| 422 | 2016A_P6-C10_1 | PHI6 X PHI1 | 81 |

**Table S9: Maintenance of maternal heterozygosity in putative clonal progeny**

| # | Plant ID | Genetic background of hybred | Maintenance of maternal heterozygosity , % |
|---|---|---|---|
| 423 | 2016A_P6-B10_15 | PHI6 X PHI1 | 81 |
| 424 | 2016A_P5-F10_2 | PHI6 X PHI1 | 80 |
| 425 | 2016A_P5-G10_8 | PHI6 X PHI1 | 79 |
| 426 | 2016A_P5-F10_4 | PHI6 X PHI1 | 79 |
| 427 | 2016A_P6-C12_4 | PHI6 X PHI1 | 78 |
| 428 | 2016A_P6-B8_3 | PHI6 X PHI1 | 76 |
| 429 | 2016A_P5-H6_1 | PHI6 X PHI1 | 76 |
| 430 | 2016A_P6-B10_4 | PHI6 X PHI1 | 75 |
| 431 | 2016A_P5-H8_1 | PHI6 X PHI1 | 72 |
| 432 | 2016A_P5-H11_6 | PHI6 X PHI1 | 71 |
| 433 | 2016A_P6-A9_4 | PHI6 X PHI1 | 70 |
| 434 | 2016A_P6-A8_1 | PHI6 X PHI1 | 70 |
| 435 | 2016A_P6-B7_2 | PHI6 X PHI1 | 69 |
| 436 | 2016A_P5-F1_1 | PHI6 X PHI1 | 69 |
| 437 | 2016A_P6-B10_11 | PHI6 X PHI1 | 63 |
| 438 | 2016A_P5-H3_11 | PHI6 X PHI1 | 62 |
| 439 | 2016A_P5-F2_3 | PHI6 X PHI1 | 62 |
| 440 | 2016A_P5-H3_6 | PHI6 X PHI1 | 61 |
| 441 | 2016A_P5-G8_4 | PHI6 X PHI1 | 60 |
| 442 | 2016A_P5-H3_3 | PHI6 X PHI1 | 58 |
| 443 | 2016A_P5-H11_3 | PHI6 X PHI1 | 57 |
| 444 | 2016A_P6-B7_1 | PHI6 X PHI1 | 57 |
| 445 | 2016A_P5-H5_1 | PHI6 X PHI1 | 57 |
| 446 | 2016A_P5-H3_5 | PHI6 X PHI1 | 57 |
| 447 | 2016A_P5-H5_6 | PHI6 X PHI1 | 54 |
| 448 | 2016A_P6-A8_8 | PHI6 X PHI1 | 54 |
| 449 | 2016A_P5-H3_1 | PHI6 X PHI1 | 53 |
| 450 | 2016A_P6-A9_2 | PHI6 X PHI1 | 53 |
| 451 | 2016A_P5-G9_1 | PHI6 X PHI1 | 52 |
| 452 | 2016A_P5-H11_2 | PHI6 X PHI1 | 50 |
| 453 | 2016A_P6-A9_1 | PHI6 X PHI1 | 50 |
| 454 | 2016A_P5-H5_4 | PHI6 X PHI1 | 49 |
| 455 | 2016A_P6-A9_6 | PHI6 X PHI1 | 49 |
| 456 | 2016A_P5-H3_4 | PHI6 X PHI1 | 48 |
| 457 | 2016A_P6-C12_3 | PHI6 X PHI1 | 48 |
| 458 | 2016A_P6-B12_7 | PHI6 X PHI1 | 48 |
| 459 | 2016A_P6-B10_8 | PHI6 X PHI1 | 47 |
| 460 | 2016A_P5-H5_5 | PHI6 X PHI1 | 47 |
| 461 | 2016A_P5-G8_3 | PHI6 X PHI1 | 47 |
| 462 | 2016A_P5-H8_16 | PHI6 X PHI1 | 45 |
| 463 | 2016A_P6-C10_2 | PHI6 X PHI1 | 45 |
| 464 | 2016A_P5-H5_12 | PHI6 X PHI1 | 44 |
| 465 | 2016A_P6-A9_8 | PHI6 X PHI1 | 43 |
| 466 | 2016A_P6-A8_16 | PHI6 X PHI1 | 43 |
| 467 | 2016A_P5-G8_5 | PHI6 X PHI1 | 42 |
| 468 | 2016A_P5-H10_3 | PHI6 X PHI1 | 40 |
| 469 | 2016A_P5-G9_5 | PHI6 X PHI1 | 40 |
| 470 | 2016A_P6-B10_12 | PHI6 X PHI1 | 40 |
| 471 | 2016A_P6-C10_3 | PHI6 X PHI1 | 39 |
| 472 | 2016A_P6-A8_12 | PHI6 X PHI1 | 38 |
| 473 | 2016A_P5-G12_1 | PHI6 X PHI1 | 38 |
| 474 | 2016A_P5-H3_2 | PHI6 X PHI1 | 37 |
| 475 | 2016A_P6-D11_1 | PHI6 X PHI1 | 35 |
| 476 | 2016A_P5-G9_6 | PHI6 X PHI1 | 35 |
| 477 | 2016A_P5-H5_3 | PHI6 X PHI1 | 33 |
| 478 | 2016A_P6-B10_13 | PHI6 X PHI1 | 33 |

### Table S10. Positions and values of all informative SNPs on the chromosomes of individuals with maternal heterozygosity maintenance over 97%

| Plant ID | Female_IL | Male_IL |
|---|---|---|
| 2013_P1-8-4 | PHI1 | PHI2 |
| 2015_P3-C11-3 | PHI1 | PHI3 |
| 2015_P3-F2-1 | PHI1 | PHI3 |
| 2015_P3-F3-1 | PHI1 | PHI3 |
| 2015_P3-F4-3 | PHI1 | PHI3 |
| 2015_P3-F4-1 | PHI1 | PHI3 |
| 2015_P3-B5-1 | PHI1 | PHI3 |
| 2015_P3-F9-5 | PHI2 | PHI1 |
| 2015_P3-G1-4 | PHI2 | PHI1 |
| 2015_P3-H3-1 | PHI2 | PHI1 |
| 2015_P3-H5-4 | PHI2 | PHI1 |
| 2015_P4-A3-1 | PHI2 | PHI1 |
| 2015_P4-A5-1 | PHI2 | PHI1 |
| 2015_P4-A9-3 | PHI2 | PHI1 |
| 2016A_P2-H6_16 | PHI2 | PHI1 |
| 2016A_P2-H6_9 | PHI2 | PHI1 |
| 2015_P4-B9-1 | PHI2 | PHI3 |
| 2016A_P3-H8_1 | PHI4 | PHI1 |
| 2016A_P4-A1_1 | PHI4 | PHI1 |
| 2016A_P4-D12_5 | PHI4 | PHI1 |
| 2016H-P2-B7_1 | PHI4 | PHI3 |
| 2016H-P2-B7_4 | PHI4 | PHI3 |
| 2016H-P2-C6_3 | PHI4 | PHI3 |
| 2016H-P2-G6_2 | PHI4 | PHI3 |
| 2016H-P2-G6_3 | PHI4 | PHI3 |
| 2016A_P5-G10_10 | PHI6 | PHI1 |
| 2016A_P5-G10_11 | PHI6 | PHI1 |
| 2016A_P5-H5_7 | PHI6 | PHI1 |

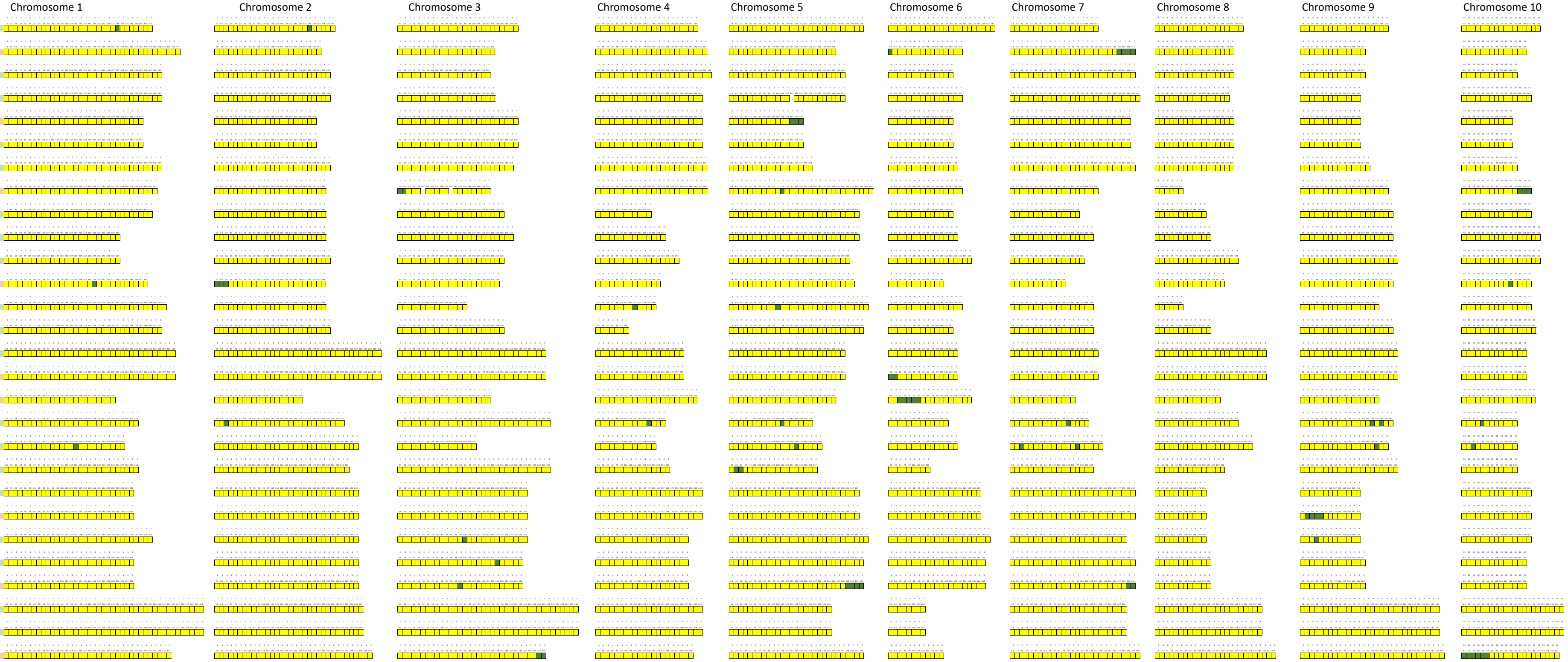

- clonal to mothers
- lost heterozygosity

- heterozygous SNP
- homozygous SNP

### Table S11. List of primers

| Gene | Primer name | PCR program | Fragment size | Sequence | Remark |
|---|---|---|---|---|---|
| *Nrf4* | NRM4-Zyg-GSP-F | 95˚C – 5'; (94˚C – 15"; 61,5 ˚C – 15"; 72˚C – 30" ) X 40; 72˚C – 5' | 459 bp | TTCAATGTCATGGGCCGGATTCG | Genotyping wild-type allele of *Nrf4* gene for *nrf4-1 & nrf4-2* |
| | NRM4-Zyg-GSP-R | | | AGTGCAATTCCGTAGCGCGATGA | |
| *Nrf4* | NRM4-GSP-new | 95˚C – 5'; (94˚C – 40"; 63 ˚C – 50"; 72˚C – 60" ) X 40; 72˚C – 5' | 328 bp | AACTCGCCGATTGCACACCTTAC | Genotyping mutant *nrf4-1* allele |
| | Cust Mu | | | AATCCCGTCCGCTCTTCGTCTAT | |
| *Nrf4* | DO153493 | 95˚C – 5'; (94˚C – 15"; 65˚C – 15"; 72˚C – 30" ) X 35; 72˚C – 5' | 1333 bp | TAGACTTAGACTGAAACTGCGGGCCTCTT | Genotyping wild-type allele of *Nrf4* gene for *nrf4-A09; nrf4-C07* |
| | DO153496 | | | GATCCAGGGCACCACCATTGCCT | |
| *Nrf4* | DO153496 | 95˚C – 5'; (94˚C – 15"; 65 ˚C – 15"; 72˚C – 30" ) X 35; 72˚C – 5' | A09 – 570 bp<br>C07 – bp | GATCCAGGGCACCACCATTGCCT | Genotyping mutant *nrf4-A09* and *nrf4-C07* alleles |
| | MuTIR | | | AGAGAAGCCAACGCCAWCGCCTCYATTTCGTC | |
| *Mtl* | MZA-18157-F | 95˚C – 5'; (94˚C – 15"; 60 ˚C – 15"; 72˚C – 20" ) X 35; 72˚C – 5' | 600-700 bp | CTTTGATCCTCTGTATTGAAG | SSR marker closely linked to *Mtl* gene |
| | MZA-18157-R | | | AGGCTGTATCTGCATATGATG | |
| *Nrf4* | pco657953f | 95˚C – 5'; (94˚C – 20"; 63˚C – 15"; 72˚C – 30") X 35; 72˚C – 5' | gDNA - 867bp;<br>cDNA - 654bp | AAACTTTTCGAAAATCCCCCACAGAATCCC | RT-PCR of *Nrf4* mRNA |
| | pco657953r | | | AAAACCACTGCATGATCACCTCCGC | |
| *ZmAct1* | Actin1-F | 95˚C – 5'; (94˚C – 15"; 60˚C – 15"; 72˚C – 20") X 35; 72˚C – 5' | gDNA - 253bp;<br>cDNA - 168bp | AATGGCACTGGAATGGTCAAG | RT-PCR of *ZmAct1 mRNA* |
| | Actin2-R | | | CAGTGTCAGGATGCCTCTCTT | |
